# Vulnerability to climate change for narrowly ranged species: the case of Ecuadorian endemic *Magnolia mercedesiarum*

**DOI:** 10.1101/482000

**Authors:** V. Shalisko, J. A. Vázquez-García, A. R. Villalobos-Arámbula, M. A. Muñiz-Castro

## Abstract

Species vulnerability to climate change has been inferred using species distribution models from an example of the recently discovered *Magnolia mercedesiarum* (sect. *Talauma*, Magnoliaceae), a narrowly ranged species endemic to moist tropical forests in the eastern Ecuadorian Andes. The environmental conditions within the current species distribution area has been compared with conditions projected to 2050 and 2070, using data from the HadGEM2-ES model in two CO2 emission scenarios: RCP4.5 and RCP8.5. The ecological niche modelling allowed determination of parameters of climatic environmental conditions that control current species distribution to produce a hypothesis on probable changes in spatial pattern of suitable habitats in future scenarios. Within the current species distribution area of *M. mercedesiarum*, significant reduction of habitat suitability was projected for both emission scenarios, combined with a lack of nearby areas with adequate environmental conditions. Several disjunct sites of high habitat suitability were found to emerge in the Colombian Andes, but they seem unreachable by this tree species in the scope of a few decades, due to intrinsic dispersal limitations. The reduction of habitat suitability and improbability of distribution area shift to adjacent geographic locations could mean a high species vulnerability to climate change. The species could be at risk of extinction if it does not possess hidden phenotypical plasticity and potential for fast adaptation to climate change.

## Introduction

*Magnolia mercedesiarum* D. A. Neill, A. Vázquez & F. Arroyo (subsect. *Talauma*, Magnoliaceae) is a broadleaf evergreen tree (Fig. 1), naturally occurring in the moist tropical mountainous forests in the eastern slopes of the Andes in Ecuador. As stated in Vázquez-GarcÍa et al. (2018), this species is currently known from only four localities in the Napo and SucumbÍos Ecuadorian provinces and has an extremely narrow distribution range; thus, it fulfills the International Union for Conservation of Nature (IUCN 2012) Red List criteria B1 ab (i, ii, iii) for an endangered (EN) species.

**Figure 1.**
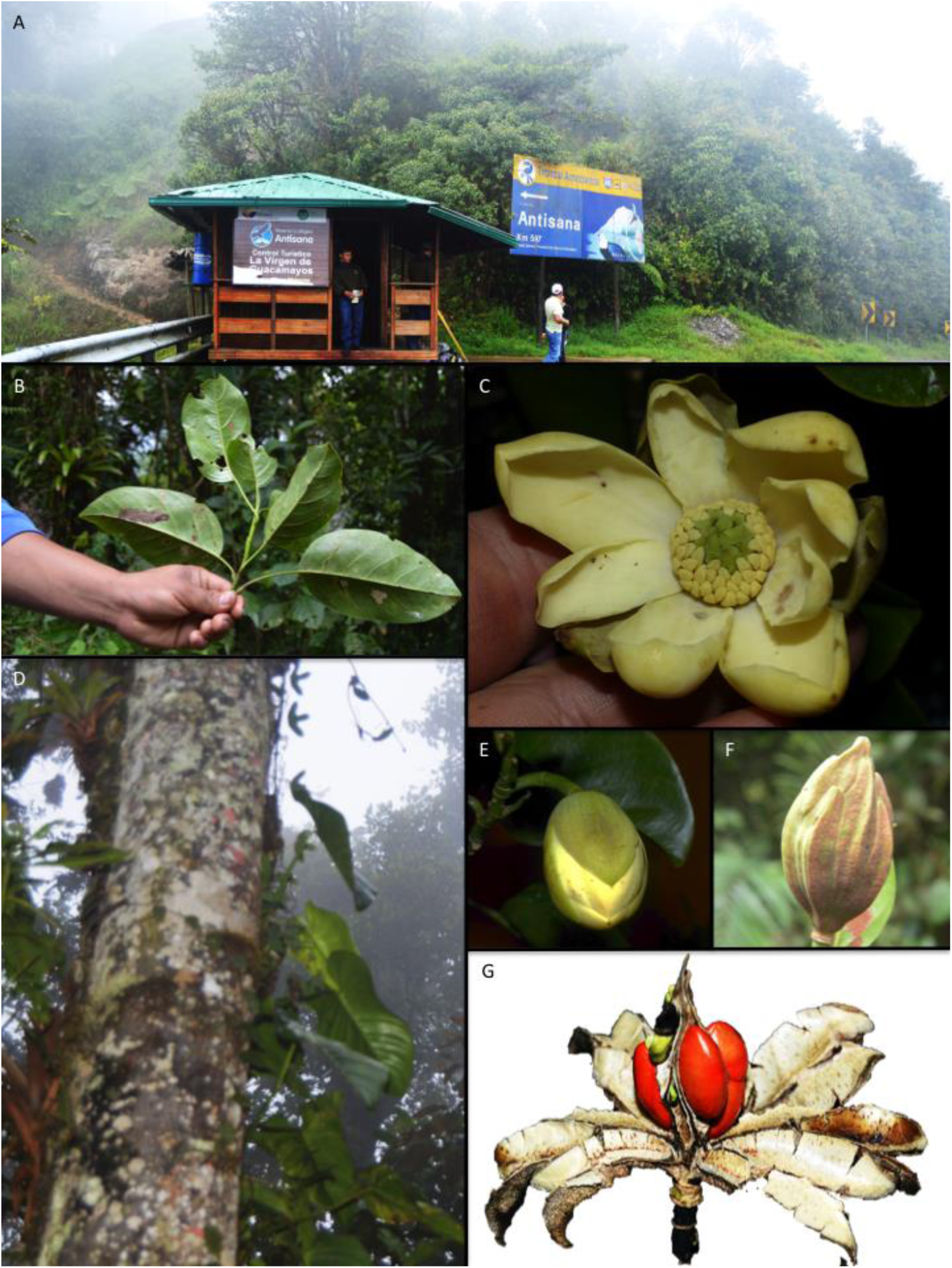
Magnolia mercedesiarum. A) The Antisana Ecological Reserve, one of the areas where this species is protected. B) Leaves, C) Open flower in female phase, D) Tree trunk, E) Flower bud showing sepals, F) Mature fruit, G) Fruit during seed dehiscing.

The aim of this study is to analyze the vulnerability of *M. mercedesiarum* to climate change predicted for the next decades in selected scenarios influenced by anthropogenic greenhouse gas emissions.

The species vulnerability is understood in geographic terms as being inversely proportional to the extent of reachable suitable habitat and is directly related to probable diminution of distribution area. The tree population size in given constant biotic conditions is expected to be directly proportional to the size of the area with appropriate abiotic habitat conditions or, in other words, to its projected fundamental niche. Both population size and area of occupancy of species are included as criteria for evaluation of species vulnerability and conservation status (IUCN, 2012). The analysis of species vulnerability is addressed through: a) comparing climate conditions in *M. mercedesiarum* current distribution area with conditions predicted in climate change scenarios to visualize changes in temperature and precipitation as envelope climatic conditions; b) developing the species distribution model for *M. mercedesiarum* on current environmental conditions; c) producing projections of probable pattern of suitable habitat in climate change scenarios across northeastern South America, and forecasting the change in species distribution in areas adjacent to current presence locations; d) evaluation of the species potential to reach the disjunct areas with potentially suitable habitat, by considering dispersal constraints shared within Neotropical *Magnolia*.

Hutchinson’s (1957) contribution to the theory of ecological niche made possible development of ecological niche models in environmental factors space, which is crucial for correlative species distribution modelling. Correlative models seek to relate species observations with potential environmental predictors to produce a hypothesis on species ecological niche in the form of statistical response surfaces and further prediction of its potential or real distribution (Franklin & Miller, 2009; Guillera-Arriota, 2017). Modern methods of predictive species distribution analysis strongly rely on Hutchinson’s niche–biotope duality, by including implicit reprojection of observations and predictions between environmental and geographic spaces (Colwell & Rangel, 2009). The ecological niche model can be inferred in the environmental space by statistical or machine learning methods, that typically include some grade of extrapolation in the environmental factors space (Mesgaran et al., 2014). Niche models are used to make projections to geographical space, not only to reconstruct the current species distribution, but to make assumptions on probable past or future distributions in case the environmental predictors are available for past of future conditions (Guisan & Thuiller, 2005; Elith & Leathwick, 2009). The particular question of usefulness of envelope SDMs based on climatic predictors, known as bioclimatic envelope models, was addressed in Araújo and Peterson (2012), who emphasized the need to explicitly state the assumptions about species ecological niches and put the results of SDM into a clear conceptual framework to avoid confusion during their interpretation and understand prediction limitations.

The hypothetical potentially abiotically-suitable area and area of true species distribution were recognized by Hutchinson (1978), correspondingly, as fundamental and realized ecological niches. As stated by Soberón and Nakamura (2009), these are different entities that correspond to actual and potential distribution areas. According to these authors, the fundamental niche is the fraction of the environmental space of scenopoetic variables for which the intrinsic population growth rate of a species would be positive. The geographic space where such scenopoetic conditions actually occur in, in a given region and time, can be projected to environmental space as potential species niche, being a smaller subset of the fundamental niche. This fraction of geographic space has suitable abiotic habitat conditions for species distribution. The realized niche is the subset of the potential niche that corresponds to the fraction of geographical space that the species actually use in specific time, being able to reach it by dispersion and after considering the effect of biotic interactions. Pulliam (2000) considers that areas that correspond to a realized niche could be geographically larger than those for the fundamental one, due to constant immigration of individuals from the source areas with suitable habitat conditions into closely located sink areas, where environmental conditions are less suitable, and incompatible with indefinite species persistence in the absence of immigration. The process for bioclimatic envelope modelling includes production of the fundamental niche model in the environmental space, and further rendering of a potential niche in actual geographic space for a given time instant, defining, in this way, areas with habitat abiotic conditions potentially suitable for species presence. The actual possibility of species occurrence in this area cannot be obtained in this manner because it additionally depends on dispersal, disturbance, resource and biotic factors that can be modelled by means other than bioclimatic envelope modelling (Guisan & Thuiller, 2005).

It should be clarified that the reprojections between models constructed in a hypervolume of environmental factors and geographic space are particularly useful to deal with a Grinnellian niche that depends on scenopoetic variables at a coarse scale (Soberón, 2007). Scenopoetic variables could be defined as environmental conditions favorable for survival and reproduction of individuals linked to specific geographic locations, which cannot be easily consumed or altered by individuals, such as topographic, climatic or edaphic parameters. The spatial resolution is important for the reason that during the analysis at coarse resolution, the effect of biotic interactions, competition for resources, and other local bionomic variables is effectively averaged within the grid cells, so the scenopoetic variables could retain considerable predictive value in niche modelling. The climatic regulators are likely to control gradual distribution patterns over a large geographic extent, while a patchy species distribution observed in fine resolution is related to patchy availability of resources and patterns of other bionomic variables driven by local topography and habitat fragmentation (Guisan & Thuiller, 2005). The reduction of bionomic variable importance in coarse resolution is known as the Eltonian noise hypothesis (Soberón & Nakamura, 2009), that allows development of ecological niche models and related SDM without considering biotic interactions at the local scale. Here, we follow such a hypothesis by explicitly excluding resource factors and biotic interactions at the selected spatial scale of analysis.

The application of Hutchinson’s duality principles to predict species distributions requires assuming the hypothesis of ecological niche conservatism in application to Grinnellian niches both in spatial and temporal scales (Soberón 2007). The ecological niche conservatism as discussed in Peterson (2003) and Wiens et al. (2010) looks reasonable in the scope of a few generations or over short periods of time when evolutionary effects in ecological niches could be minor. The review of options available for plant species’ niche evolution is given in Donoghue and Edwards (2014), who link the probability of species to adapt to new climatic conditions with its geographic opportunity to disperse, ecological interactions with another species, and intrinsic proclivity of species to evolve along the climate axis. The examples of species shift to novel biomes known from several plant linages demonstrates the potential possibility of drastic niche evolution in plant species, including trees, but available data demonstrate that such shifts could occur in the scope of geologic time scales. Although the capacity of species to adapt to new conditions that is related to evolutionary changes in ecological niches can be important in long time periods, it lies out of the scope of simple SDM without explicit simulation of evolutionary processes. Here, we follow the hypothesis that the fundamental niche of species is conserved in temporal frames of a few generations and the inherited physiological tolerances of species to environmental factors are conserved in the entire spatial and temporal scope of analysis (Araújo & Peterson, 2012). This is a required assumption for SDM that explicitly uses niche–biotope duality.

Another assumption referred to as the equilibrium postulate, is necessary for development of reliable ecological niche models, predictive modelling, and forecasting (Araújo & Peterson, 2012). The equilibrium postulate can be formulated as a requirement of absence of hidden phenotypic plasticity that is not expressed in observed species adaptation to current environmental conditions; in other words, species are assumed to fully reveal their ecological potential and stay in equilibrium with their environment (Colwell & Rangel, 2009). The species are expected to inhabit the entire spatial footprint of suitable habitat conditions (Araújo & Peterson, 2012). The last factor is frequently seen as problematic since dispersal and biotic factors could constrain the possibility of species establishment in some areas. The distribution of species occurring in the environment that experience fast changes could be strongly controlled by dispersal constraints, and such species my not reach equilibrium with their environment in the current moment (Guisan & Thuiller, 2005). In a situation when the species distribution does not reach the state of equilibrium with its environment, the inferred SDM may underrepresent the environmental conditions suitable for species existence. Partially, the equilibrium postulate is resolved by the Eltonian noise hypothesis that places the biotic interactions to the local scale, beyond the spatial resolution of SDM. The leading role of dispersal constraints in a definition of possibility for species to occupy the available habitat is discussed in Ozinga et al. (2005). It could be assumed that dispersal constraints represent the main factor preventing plant species from achieving full equilibrium with the current abiotic environment, on the coarse and medium scale of analysis.

The issue with underrepresentation of the environmental conditions in SDM produced from incomplete data or for species not fully in equilibrium with its environment can be resolved through extrapolation, which consists of model extension into novel regions of covariate space that are not represented in known presence data (Elith & Leathwick, 2009). The extrapolation is an immanent part of any statistical or machine learning modelling method, however the grade of extrapolation required to produce a predictive model is higher when not enough occurrence data are available, or the occurrence data are biased. The extrapolation into novel covariate space cannot be avoided, particularly in the case of model transferability in time and space; instead, the grade of extrapolation can be measured by several methods to address the related uncertainty (Elith et al., 2010; Mesgaran et al., 2014).

## Methods

### Changes in environmental parameters in species’ current distribution area

The probable current distribution area of *Magnolia mercedesiarum* was accepted from Vázquez-GarcÍa and co-workers’ (2018) estimation, based on SDM with conservative equal sensitivity and specificity (ESS) threshold (Appendix S1 and Supporting Information Data S1, S2). Considering that the current study does not propose to discover the actual species distribution, which is already known, but rather aims to provide information on probable distribution change in climate change scenarios, we take Vázquez-GarcÍa and colleages’ data as known true species distribution and use this to produce the SDM suitable to develop predictions of future distribution. Instead of configuring the model in a base of a few known presence localities for *M. mercedesiarum*, which could be biased due to an incomplete and spatially uneven set of field observations, we use the simulated presence points within a previously estimated distribution area. We prefer simulated presence data to known species observation localities because the latter could be spatially biased, spatially autocorrelated, and incomplete because the number of known separate localities of species presence is as few as four. The geographic sampling bias can have severe effects in SDM performance and results and can be difficult to correct (Fourcade et al., 2014). To sample the environmental variability in probable current distribution area, we had drawn 500 uniformly distributed random points within the presence grid cells. For the SDM process, this set was split into ten equal size folds for use in cross-validation replications, required for assessing model generality (Merrow et al., 2013). By introducing random sampling within the current distribution area, we simulate the uniform sampling effort, and therefore satisfy an assumption of uniform initial probability of grid cell selection for SDM training (Phillips et al., 2009).

The background point selection for SDM has been performed following the recommendations of Barbet-Massin et al. (2012) and Merrow et al. (2013), defining half of the randomly distributed background points within the probable dispersal radius of 500 km, and the rest of background points uniformly distributed across the entire extent of the terrestrial environment of the modeling area in northwestern South America. The overall number of 900 background points in each replication provide a reasonable representation of the environmental variation in the modelling area. The background points in presence only methods may include both true and false absences, thus representing biased absence data, with contribution of false absences that depend on species prevalence (Engler et al., 2004). Consequently, the use of background data for model evaluation cannot produce reliable results when the true species distribution is known. For model evaluation in this study, the background points outside of the known current distribution were counted as true absences, but all background points inside of the species presence area were excluded. The test presence points were defined by merging of two 50-point folds, different from the training points fold, to ensure test independence and a presence prevalence greater than 0.1; this process was repeated for each cross-validation replication.

The environmental information for current conditions was extracted from the WorldClim ver. 2 dataset (Fick & Hijmans, 2017), and includes mean monthly rainfall, minimal and maximum monthly temperatures at 30″ resolution grids (∼1 km at latitude of analysis). These data derive from meteorological observations during 1970 –2000 linked to a digital elevation model and represent average conditions that existed in sites where currently growing adult *Magnolia* tree individuals had successfully established. The temperature data were statistically downscaled to 7.5″ (∼250 m) resolution by two variable polynomic regressions from elevation and latitude; elevation was taken as median grid cell elevation from the GMTED2010 digital elevation model dataset (Danielson & Gesch, 2011). The rainfall data were resampled to 7.5″ resolution using a cubic convolution algorithm, controlling the absence of negative values. The set of 19 bioclimatic predictor variables (Nix, 1986; O’Donnell & Ignizio, 2012) was produced from rainfall and temperature data with the R ‘dismo’ package (Hijmans et al., 2017) as continuous floating-point layers. The selection of climatic predictors as a primary source of environmental data for model development is due to their importance in defining the coarse-grained features of distributions (Soberón & Nakamura, 2009; Araújo & Peterson, 2012) and availability for both current and projected future conditions.

The future condition projections in climate change scenarios were selected from products of the Hadley Global Environment Model 2 – Earth System (Martin et al., 2011) compatible with the Coupled Model Intercomparison Project Phase 5 [CMIP5] (Taylor et al., 2011). According to Collins et al. (2013), the simulations in CMIP5 framework follow four representative concentration pathway [RCP] scenarios, which correspond to target CO2 concentrations. For the analysis of *Magnolia mercedesiarum* vulnerability, we selected two scenarios, the moderate RCP4.5 and extreme RCP8.5. The monthly rainfall and temperature data for both scenarios are available for the years 2050 and 2070 in the WorldClim database at 30″ resolution grids (WorldClim, 2017). The downscaling procedure to 7.5″ grid and generation of 19 bioclimatic predictors was similar to those for current conditions.

The environmental change in current distribution area has been visualized by graphical comparison of sampled monthly minimum and maximum temperatures, rainfall, and 19 bioclimatic predictors, including current (1970–2000) data and future conditions in two scenarios projected for the years 2050 and 2070 (code in Supporting Information Data S3). The median, interquartile range, and outlier-excluded range for each variable has been represented as boxplot figures. The predictor variables are expected to be non-normally distributed, hence the significance of difference in its medians was tested in pairs between dates and scenarios by a Mann-Whitney U test with a significance level of 0.05. The null hypothesis that both samples belong to the same distribution with the same median is rejected in this test if U statistics are lower than the critical value for the selected sample size and significance level. The alternative hypothesis of significant differences between sample medians is accepted instead.

### Species distribution modelling

Breiman (2001) identified two main paradigms in statistical modelling: the first based on selecting the appropriate data model prior to model fitting, and the second that avoids starting with a defined data model but determining the relation between predictors and response variables during the algorithmic model selection and fitting process. The second approach, termed machine learning, has gained popularity in ecological modelling during the last two decades (Elith et al., 2008; Franklin & Miller, 2009). Among the machine learning methods available for SDM appears MaxEnt (Phillips et al., 2006), random forests (Breiman, 2001a), boosted regression trees (Elith et al., 2008), multivariate additive regression splines (Friedman, 1991), artificial neural networks and support vector machines, the latter two methods being discussed in Hastie et al. (2009). The performance of machine learning methods is generally better than that of techniques based on non-penalized regressions, envelopes, or multivariate distances (Elith et al., 2006). However, there is no single machine learning algorithm that performs better in all SDM cases. Here, we use the popular MaxEnt algorithm because it frequently outperforms other presence-only machine learning SDM methods, particularly when the sample size is small and is derived from observations obtained opportunistically (Elith et al., 2006; Ortega-Huerta & Peterson, 2008). In Mateo, Croat, FelicÍsimo, and Muñoz’ (2010) a study of *Anthurium* species distribution in Ecuador, MaxEnt was the presence-only technique that demonstrated performance similar to the two best presence-absence methods (generalized linear models and multivariate additive regression splines). Giovanelli et al. (2010) demonstrated that MaxEnt and support vector machines methods produce consistent predictions for distribution of spatially restricted species using calibration areas of different size with higher accuracy compared to several other SDM methods. The analysis of Aguirre-Gutierrez et al. (2013) reveals lower propitiousness of the MaxEnt algorithm to produce overfitted models, comparing with other widely used machine learning methods such as random forests, boosted regression trees, and artificial neural networks. In their analysis, the MaxEnt algorithm (along with two other methods) provided an advantage in terms of performance and consistency of predictions, which was particularly evident in the case of narrow and moderately wide distributions represented by sample sizes from a few up to 1700 observations, at fine and medium geographic scales with grid cells from one to several km². The model overfitting is a sensible issue, because it can severely reduce the transferability of SDM in space and time (Randin et al., 2006). Among the causes of MaxEnt’s good performance with small training sets is its integrated regularization procedure that balances model fit and generality, resulting in a gradual increase of mathematical complexity when more data are available, reducing in this way the risk of overfit (Phillips & DudÍk, 2008; Hastie et al., 2009; Elith et al., 2011). However, some concerns about the use of MaxEnt (e. g. from Halvorsen et al., 2016) are related to its susceptibility to spatial autocorrelation in a response variable, which is not fully understood, and its tendency to produce overfitted models when modelling is performed with default feature selection and regularization settings and is derived from a high number of presence points. Although MaxEnt’s primary purpose is to treat presence-only data, its use for presence-absence can be justified when presence-background and presence-absence models are compared. The models generated with specialized presence-absence methods could be very different from those which rely on presence-only methods, even when they a based on the same presence data (Mateo et al., 2010), and compatibility can become an issue. Accordingly, MaxEnt remains one of the best available machine learning methods that can be effectively tuned to work both with presence-background and presence-absence data (Philips & DudÍk, 2008), considering that its results remain different from those of statistical presence-absence methods because MaxEnt does not estimate the species presence probability, but the relative habitat suitability (Guillera-Arriota et al., 2014).

The SDM has been performed using MaxEnt 3.4.1 (Phillips et al., 2006, 2017) in the R ‘dismo’ environment (code in Supporting Information Data S4). The cross-validation workflow included producing ten correlative model replications based in the background and presence data with a prevalence of 0.1 in each case, following the recommendations of van Proosdij et al. (2016). The full set of 19 bioclimatic predictors for current conditions (coded as “bio1” to “bio19”) and mean elevation (“alt”) were included in the regularization and model selection process. We found no a priori reason to exclude any environmental predictor from modeling but decided to use only linear and quadratic features for production of derived predictors. The hinge, product, and threshold features were excluded to prevent model overfitting and improve its interpretability. The inclusion of product and hinge features may lead to model overfitting observed by Harlovsen et al. (2016) under default feature selection settings, when each environmental variable is transformed to six feature types, prior to model selection, and the selection algorithm tends to produce a very complex model in terms of number of parameters. Elith et al. (2011) note that hinge features, which could be available with at least 15 presence points, can lead to linear and threshold features becoming redundant. Finally, product and hinge features may produce response curves that are not consistent with theoretically expected unimodal or truncated unimodal forms (Oksanen & Minchin, 2002). We use the lax regularization parameter of 0.193 both for linear (βL) and quadratic (βQ) features. We had not performed separate stepwise selection of predictors because the MaxEnt algorithm uses an effective mechanism of feature shrinkage referred as the LASSO penalty (Tibshirani, 1996) and produces sparse solutions by setting variable coefficients to 0 in case of poor variable contribution, and thus, performs the model selection by itself (Merrow et al., 2013). This feature selection is effective, more stable than stepwise regression for correlation of predictors, and unlikely to be improved by pre-selecting variables (Elith et al., 2011). The proposal of Halvorsen et al. (2016) to use forward variable selection, instead of the LASSO penalty mechanism of MaxEnt, is derived from the study of a special case, when six feature types with default regularization settings were used on correlated predictor variables and requires further testing to be accepted as universally applicable.

The most common measure of the overall discriminatory capacity of SDMs is the area under the curve (AUC) of the receiver operating characteristic (Fielding & Bell 1997; McPherson et al., 2004). The AUC is threshold independent and near independent from prevalence of presences, although a prevalence below 0.1 can result in inflation of its values (McPherson et al., 2004). According to Manel et al. (2001), the AUC values above 0.9 can be considered high in terms of the model performance in discrimination between species presences and absences. The theoretical maximum value of AUC is 1, which can be obtained in a test with known presence and absence data that are independent from training data and referred to here as real AUC (Jiménes-Valverde, 2012; Halvorsen et al., 2016). This metric differs from the train AUC that can be estimated from presence and background dataset used in model training. The maximum attainable train AUC value is expected to have a value lower than 1 because it depends on the fraction of the geographical area of interest covered by true species distribution, and decreases with the number of presence records, given the fixed number of background points (Bean et al., 2012; van Proosdij et al., 2016).

This process occurs because part of the background points may actually correspond to presences, and the fact MaxEnt appends presence points to the background dataset during AUC calculation (Phillips et al., 2006; Elith et al., 2011). In this study, we treat the probable current distribution of *M. mercedesiarum* taken from Vázquez-GarcÍa et al. (2018) as the known realized distribution of species, therefore, in model evaluation, we deal with true presences and absences suitable for real AUC calculation. In this way, we determine the real AUC for model prediction in current conditions without violation of the underlying theory that requires use of true absences in model evaluation (Lobo et al., 2007; Jiménez-Valverde, 2012). We determine the real AUC value independently in each model replication, accessed mean model AUC and its standard deviation, along with the receiver operation characteristic plot. Additionally, we rank the real model AUC in a null-model test with presence and absence data (Olden et al., 2002; Raes & ter Steege, 2007). The significance of the inferred model is accessed by comparison with a set of 99 random models derived from permutated training points, to check the null hypothesis that the model is no better than those which can be obtained by chance. To reject the null hypothesis with a significance level of 0.05, the real model AUC rank is expected to be higher than 95.

Additionally, we evaluated the correlation between known data and model predictions, as the measure of significance of their relationship. The correlation between monotonous function of species presence probability and known presence-absence data could be assessed through Spearman’s rank correlation coefficient, which is in our case was preferred to Pearson’s product-moment biserial correlation because it avoids the assumption of linear relationships between two variables (Phillips et al., 2009). Spearman’s correlation coefficient does not consider real probability values, but their ranks, hence it is not sensible for precise calibration of presence probability, and thus can be used with non-calibrated logistic output of MaxEnt.

The threshold is required for transformation of continuous SDM output to presence-absence results. Such a threshold in the case of the logistic model output cannot be fixed to 0.5 because the presence probability distribution typically is skewed (Bean et al., 2012). We use the conservative criteria of threshold selection described by Liu et al. (2005) as the sensitivity and specificity equality approach. The ESS threshold is determined as a point in receiver operating characteristic curve where the absolute difference between sensitivity and specificity is minimal, weighting equally the commission and omission errors. This threshold criteria have been shown to be robust and insensitive against the modeling technique (Jiménez-Valverde, 2012). The ESS threshold was determined in each model replication from the same point dataset that is used for the receiver operation characteristic estimation. Sensitivity and specificity were reported to demonstrate the application of the ESS and assess the relative importance of model omission and commission errors as recommended by Lobo et al. (2007). True skill statistics (TSS) is a reliable threshold-dependent method of model evaluation, which integrates the effects of both sensitivity and specificity, and is negatively correlated with prevalence (Allouche et al., 2006). The TSS value was estimated for each replication and the mean TSS with its standard deviation is reported as another general model evaluation metric.

The mean logistic output from model prediction in ten cross-validation replications was interpreted as monotonously increasing function of the relative habitat suitability for species presence in grid cells (Philips & DudÍk, 2008; Phillips et al., 2009). The produced logistic output is monotonously related with another two available MaxEnt output types and yields the same results in ranked comparison of spatial grid cells, although it deviates from the true species presence probability because the last requires the adjustment of the model parameter *tau* responsible for precise calibration of the probability curve (Elith et al., 2011). The logistic output was transformed to the estimation of realized distribution of the species in current conditions by application of the ESS threshold, independently in each model replication.

### Forecasting of the species distribution in future scenarios

In order to produce predictions of species potential distribution, the set of ten models inferred in cross-validation replications for current conditions was reconfigured with projected environmental predictors. The substitution of predictor variables results in model transfer to a novel environment and is possible when the new dataset uses the same grid resolution, data type, and ranges of values as data in the initial configuration of the correlative model. The usefulness of such an approach has been demonstrated in several studies (e. g. Pearson & Dawson, 2003; Bertzky et al., 2005; Huntley et al., 2008; Randin et al., 2009; Franklin et al., 2013; West et al., 2015). A set of logistic output grids produced in forecasting was treated in the same way as for current conditions, applying ESS threshold independently in each model replication.

The species potential distribution dynamics in climate change scenarios has been evaluated by two methods. Within the current species distribution area, the ESS threshold was applied independently in ten model replications both for current and future scenarios to determine the number of grid cells and area suitable for species presence in each case. The area classified as suitable for species presence within a 10 km distance to known current distribution was included in the evaluation to address the model uncertainty. The habitat loss estimation consisted of subtracting the count of suitable cells in future scenarios from the count of suitable cells in current conditions predicted by the model, normalized by a count of cells in current conditions. In another approach, we estimated the overall size of suitable habitat in western South America by applying the ESS threshold in model replications for future conditions and taking the predominant value from presence-absence grids. The grid cell was considered to have a potentially suitable habitat in the given scenario if the majority of model replication had a logistic output value above the corresponding threshold.

To address the degree of model extrapolation into novel areas of environmental factors space, we follow the framework described in Mesgaran et al. (2014). The extrapolation is defined by these authors as a condition when the model produces predictions outside of the training range of individual covariates (type 1 novelty) or lies within ranges but constitutes novel combinations between covariates (type 2 novelty). The novelty detection in multivariate space is performed by generation of indices derived from scale-invariant Mahalanobis distance (Rousseeuw & van Zomeren, 1990). The ExDet ver. 1.1 tool (Mesgaran et al., 2014) allows detection of types 1 and 2 novelties by comparison of environmental space in a known species distribution area and in model projections, as well as to assess the contribution of environmental predictors in terms of novelty. The detection of extrapolation was performed within the areas identified by SDM as potentially suitable for species presence by majority criteria both for current and future conditions. The analysis of extrapolation in this study includes environmental factors with a contribution higher than 2% in the model covariate structure.

The species modelled presence probability in the current distribution area has been compared with the predicted for future scenarios using same sampling as in the receiver operation characteristic estimation. The species presence probability in current and future conditions data were united from ten replications to be visualized as boxplots. Similarly, to climatic predictors, the significance of difference between species presence probabilities was tested by mean of Mann-Whitney U test with a significance level of 0.05.

### Evaluation of species distribution ranges

The objective of evaluation of the dispersal potential of *Magnolia mercedesiarum* was addressed by comparison of current distribution ranges of representatives from section *Talauma* of *Magnolia* from northwestern South America. We assume that the dispersal characteristics of members of section *Talauma* are shared by the majority of its species, including *M. mercedesiarum*, and can be indirectly estimated by evaluating the size of current individual species’ ranges. The comparison of *M. mercedesiarum* distribution range with the rest of *Magnolia* could help to identify the probable evolutionary regime of dispersal for this species. The extents of *Magnolia* species range from Colombia (GarcÍa, 2007) and Ecuador (Vázquez-GarcÍa et al., 2016) were estimated by measuring the maximum Euclidean distance between observation points within each species, which is possible if taxa are known from more than one locality. The additional observations for given species were taken from the GBIF (2018).

## Results

### Changes in environmental parameters in species’ current distribution area

The current distribution of *Magnolia mercedesiarum* of 2701 km^2^ taken from Vázquez-GarcÍa et al. (2018), as shown in Fig. 2, was found to lie in the elevation range of 1075– 2576 m above the sea level; 90% of presence points are restricted to 1237–2264 m, with a median at 1736 m according to the GMTED2010 dataset. In a comparison of downscaled climatic conditions in 1970–2000 (WorldClim 2) and similarly downscaled future conditions in projection of the HadGEM2-ES model under RCP4.5 and RCP8.5 emission scenarios for all monthly temperature values, we detected an increase in the median (Fig. 3A, 3B, 4A, 4B). The monthly maximum temperature increase rate was lowest in the RCP4.5 scenario comparing current conditions with 2050 projections, ranging from 1.09°C in September to 1.59°C in April, while the monthly maximum temperature increase in the same scenario in 2050 is in the range from 3.05°C in July to 3.97°C in September. The RCP8.5 scenario corresponds to a faster temperature change, in this case the minimal monthly temperature increases for 2050 from 1.61°C in September to 2.10°C in April, while the monthly maximum temperature increase is as high as 3.69°C in July and 4.64°C in October. In both scenarios, the projection for 2070 demonstrates a further increase of temperature. The Mann-Whitney U test allowed to reject the null hypothesis in all temperature comparison cases, including comparison of current conditions with projections for 2050 and 2070, and comparison between 2050 and 2070 data, indicating the statistical significance of the temperature increase (Fig. 5, 6).

**Figure 2.**
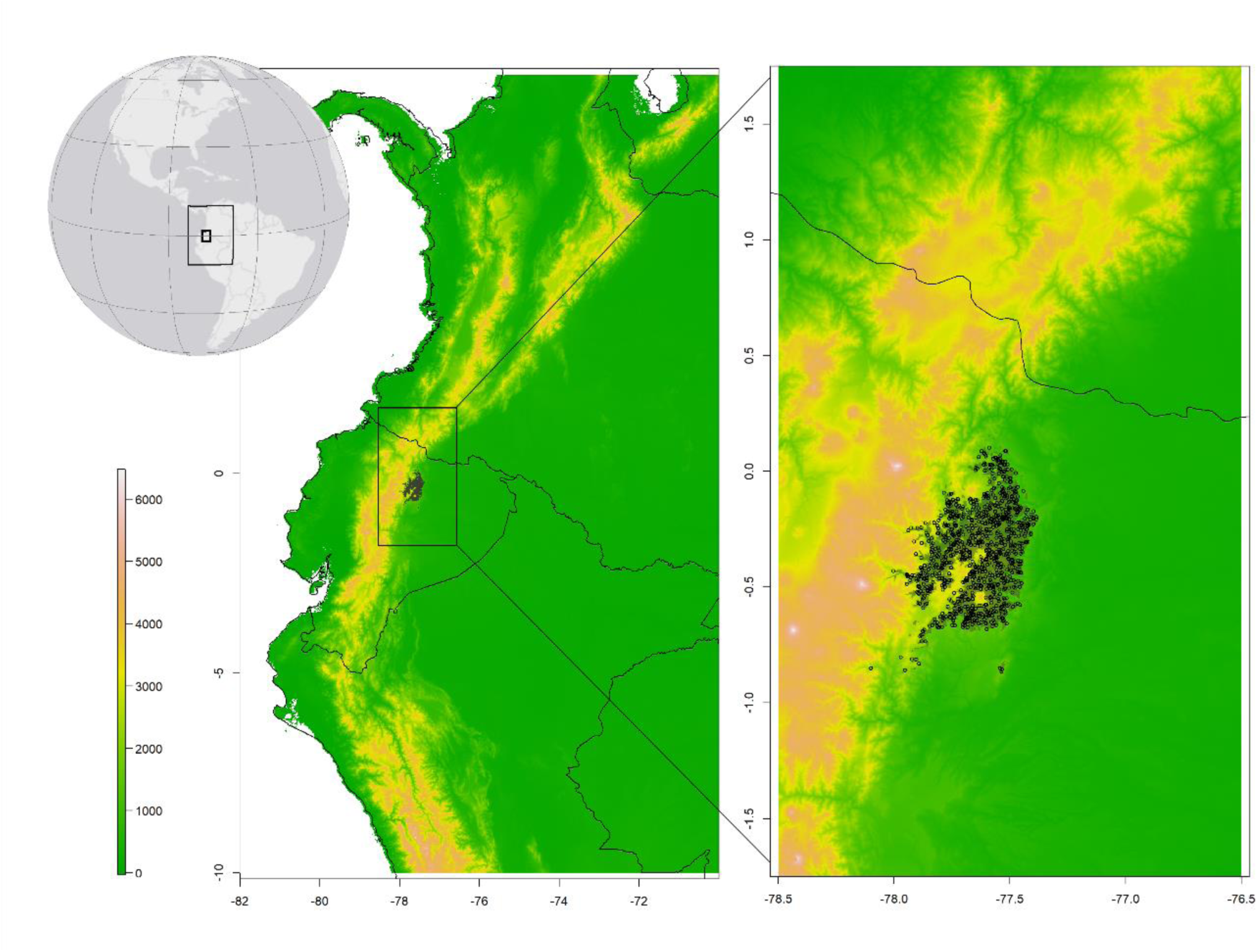
Current distribution of *Magnolia mercedesiarum* (shaded area) with GMTED2010 elevation data (scale in m above sea level). Points represent sampling sites within the current distribution area used for analysis. Rectangular areas on locator map indicate position of zoomed map fragments.

**Figure 3A.**
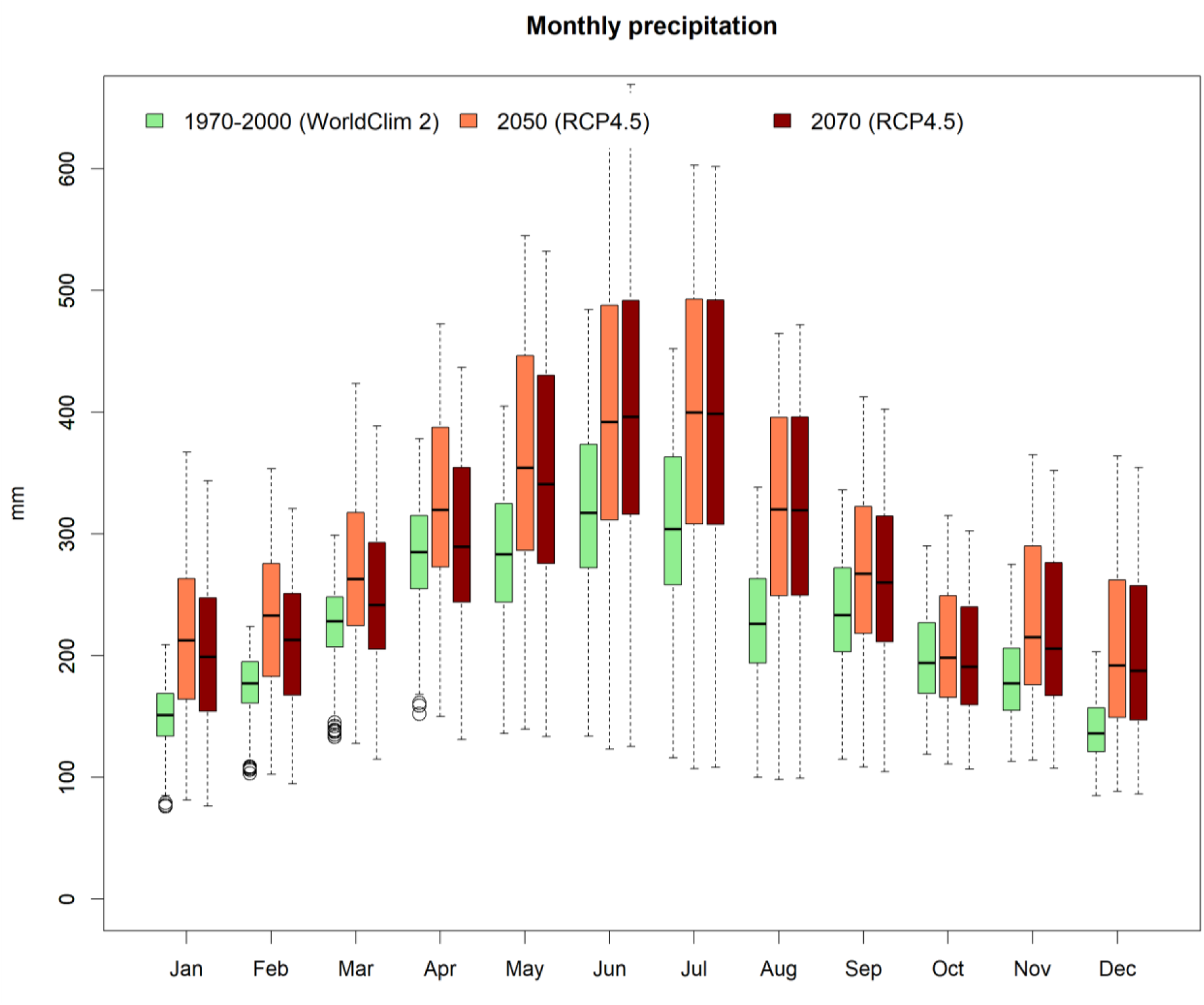
Monthly precipitation within current distribution of *M. mercedesiarum* in current conditions and projection of HadGEM2-ES model under RCP4.5 scenario for years 2050 and 2070.

**Figure 3B.**
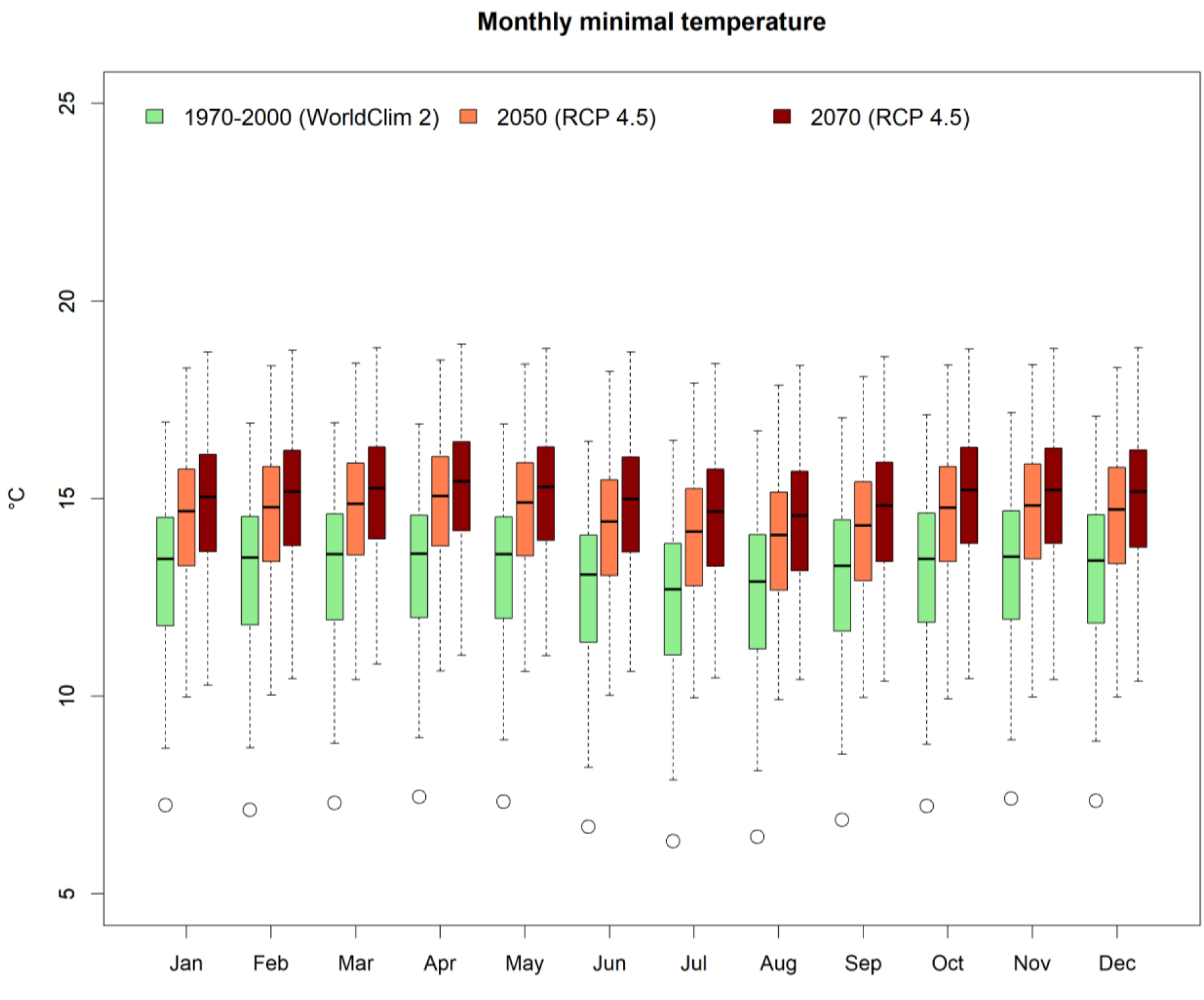
(continuation) Monthly minimum temperature within current distribution of *M. mercedesiarum* in current conditions and projection of HadGEM2-ES model under RCP4.5 scenario for years 2050 and 2070.

**Figure 3C.**
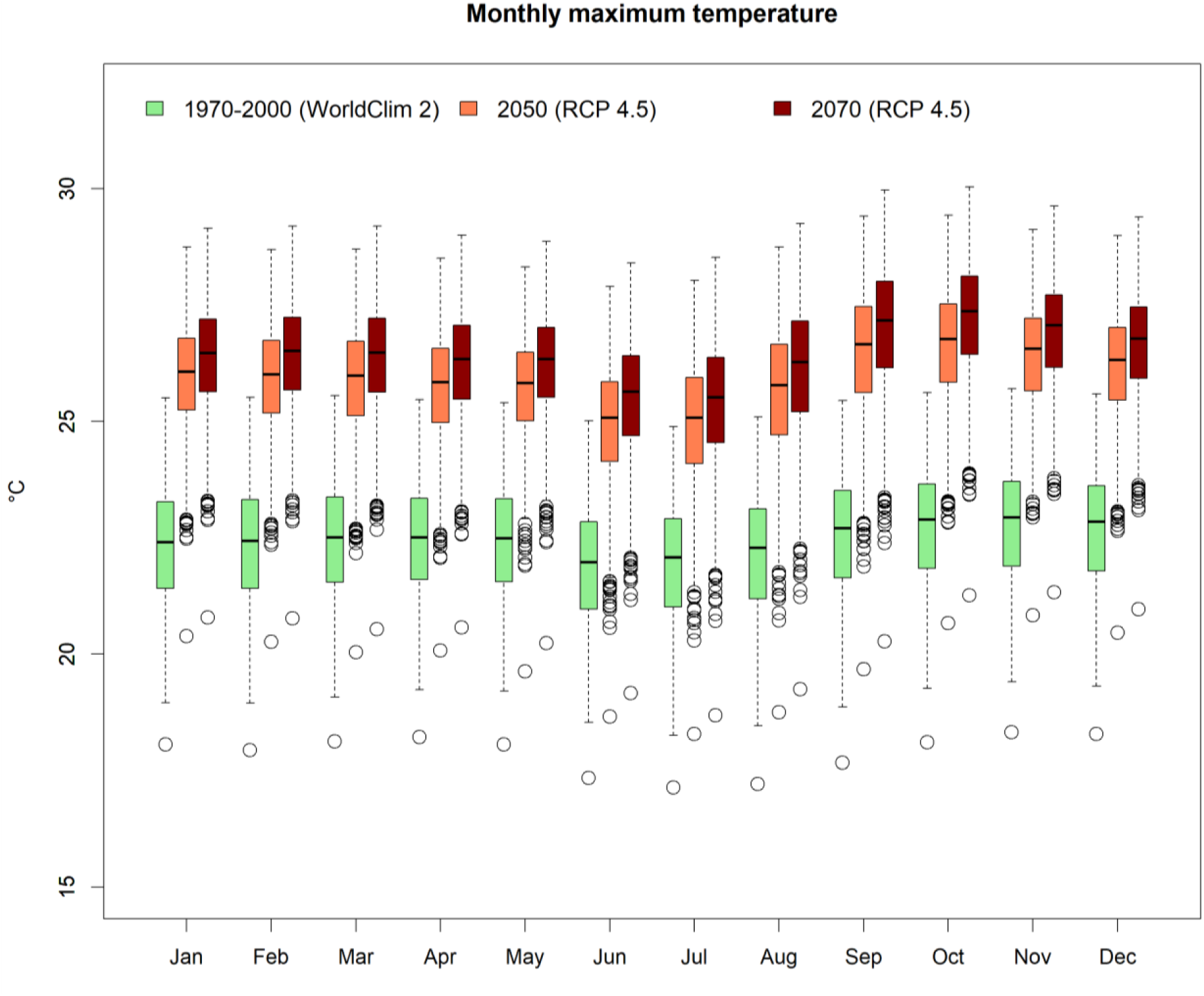
(continuation) Monthly maximum temperature within current distribution of *M. mercedesiarum* in current conditions and projection of HadGEM2-ES model under RCP4.5 scenario for years 2050 and 2070.

**Figure 4A.**
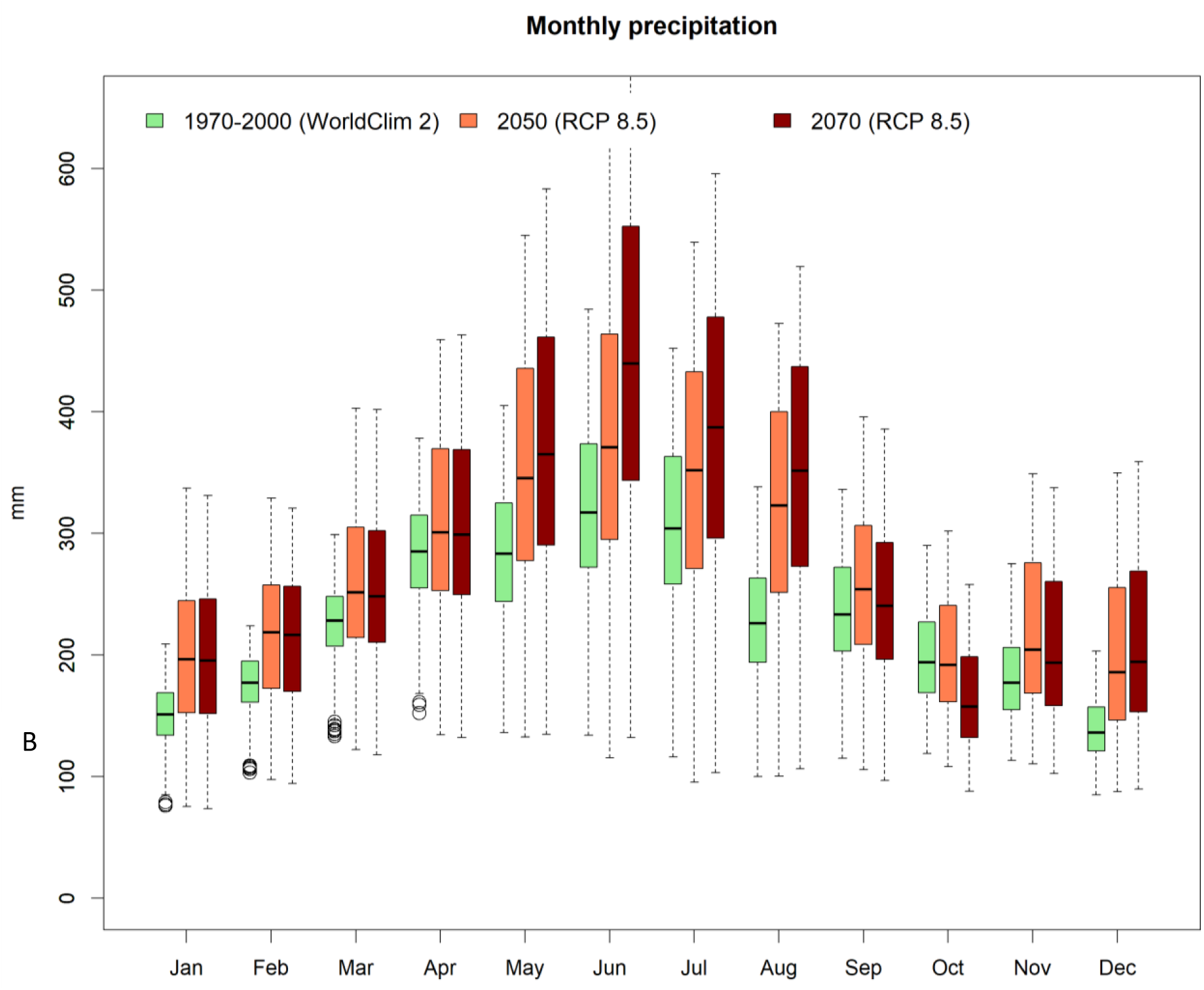
Monthly precipitation within the current distribution of *M. mercedesiarum* in current conditions and projection of HadGEM2-ES model under RCP8.5 scenario for years 2050 and 2070.

**Figure 4B.**
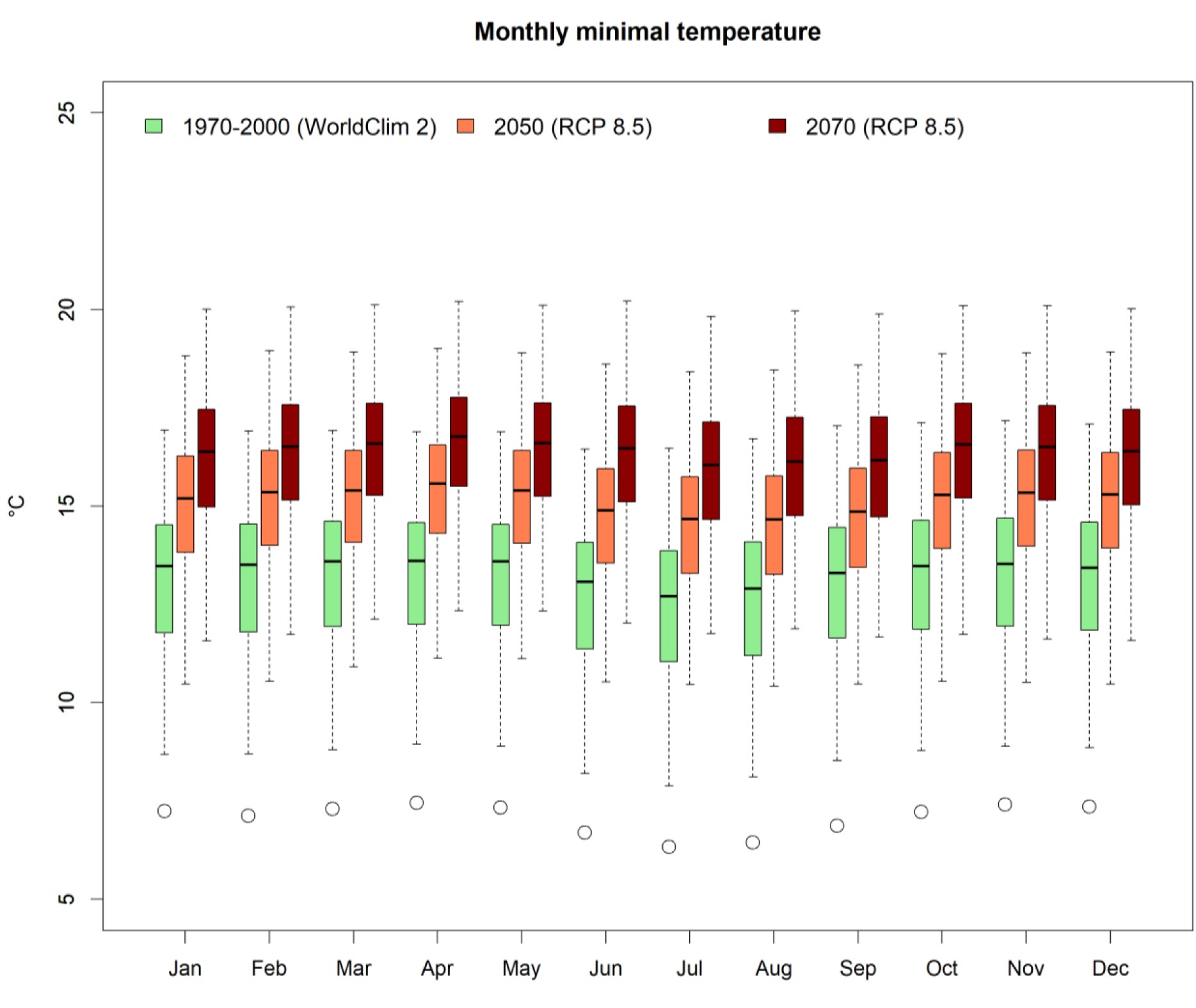
Monthly minimum temperature within the current distribution of *M. mercedesiarum* in current conditions and projection of HadGEM2-ES model under RCP8.5 scenario for years 2050 and 2070.

**Figure 4C.**
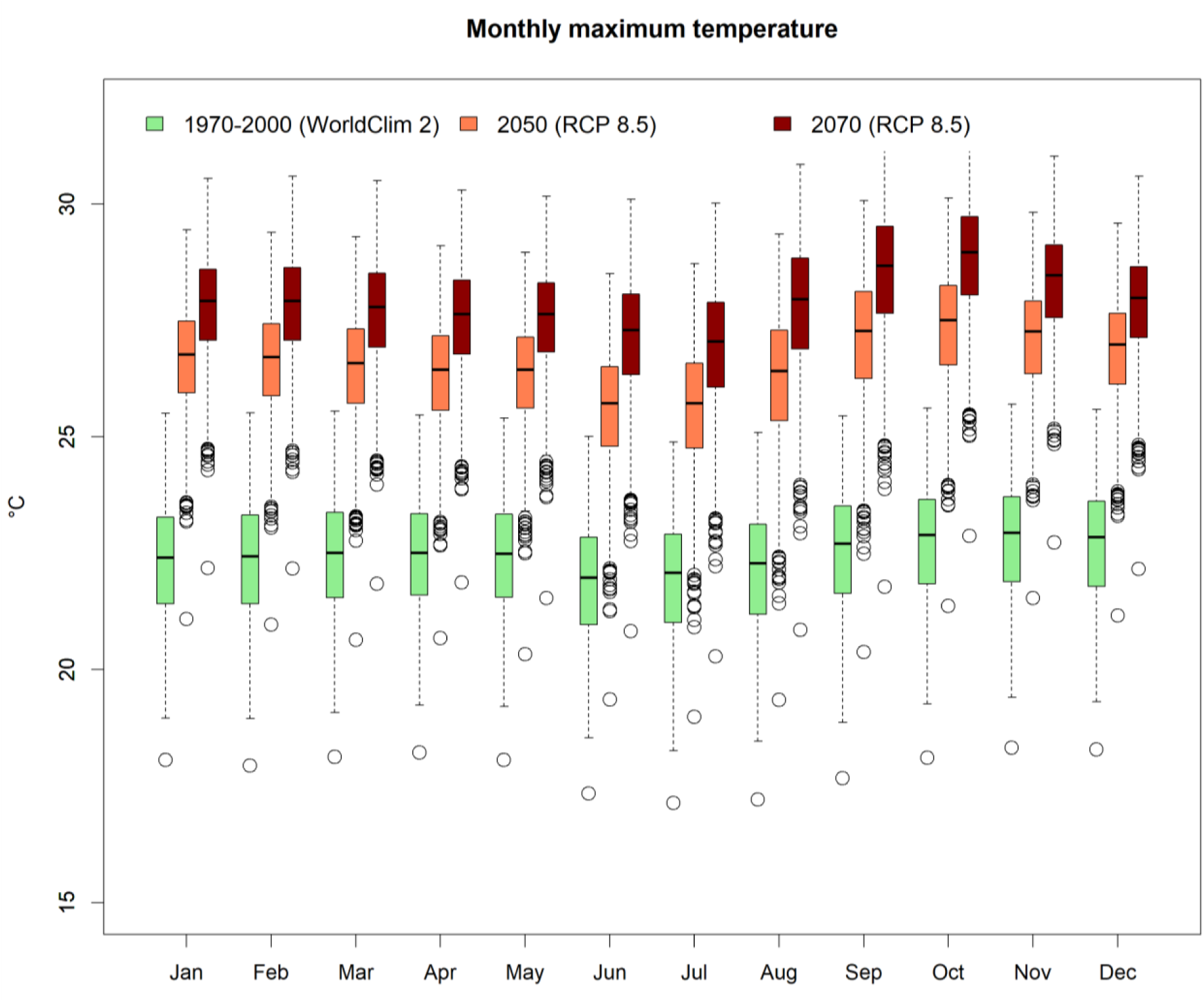
Monthly maximum temperature within the current distribution of *M. mercedesiarum* in current conditions and projection of HadGEM2-ES model under RCP8.5 scenario for years 2050 and 2070.

**Figure 5.**
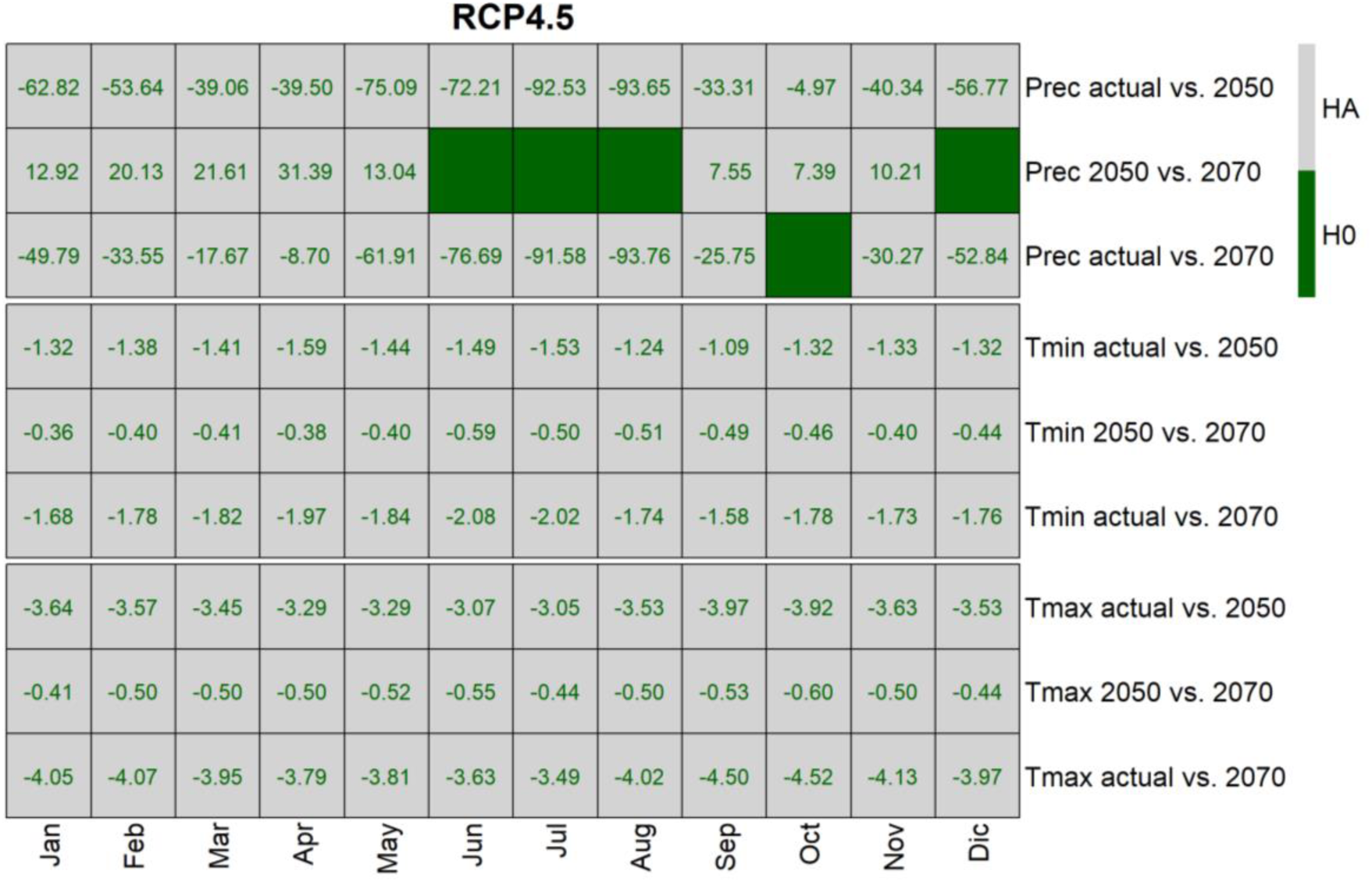
Results of Mann-Whitney U test for significance of difference in monthly precipitation, minimum and maximum temperature within the current distribution of *M. mercedesiarum* in current conditions and projection of HadGEM2-ES model under RCP4.5 scenario for years 2050 and 2070. The color-filled cells represent acceptance of null hypothesis (H0), blank cells correspond to acceptance of alternative hypothesis (HA) with numeric values reflecting difference of medians between two samples.

**Figure 6.**
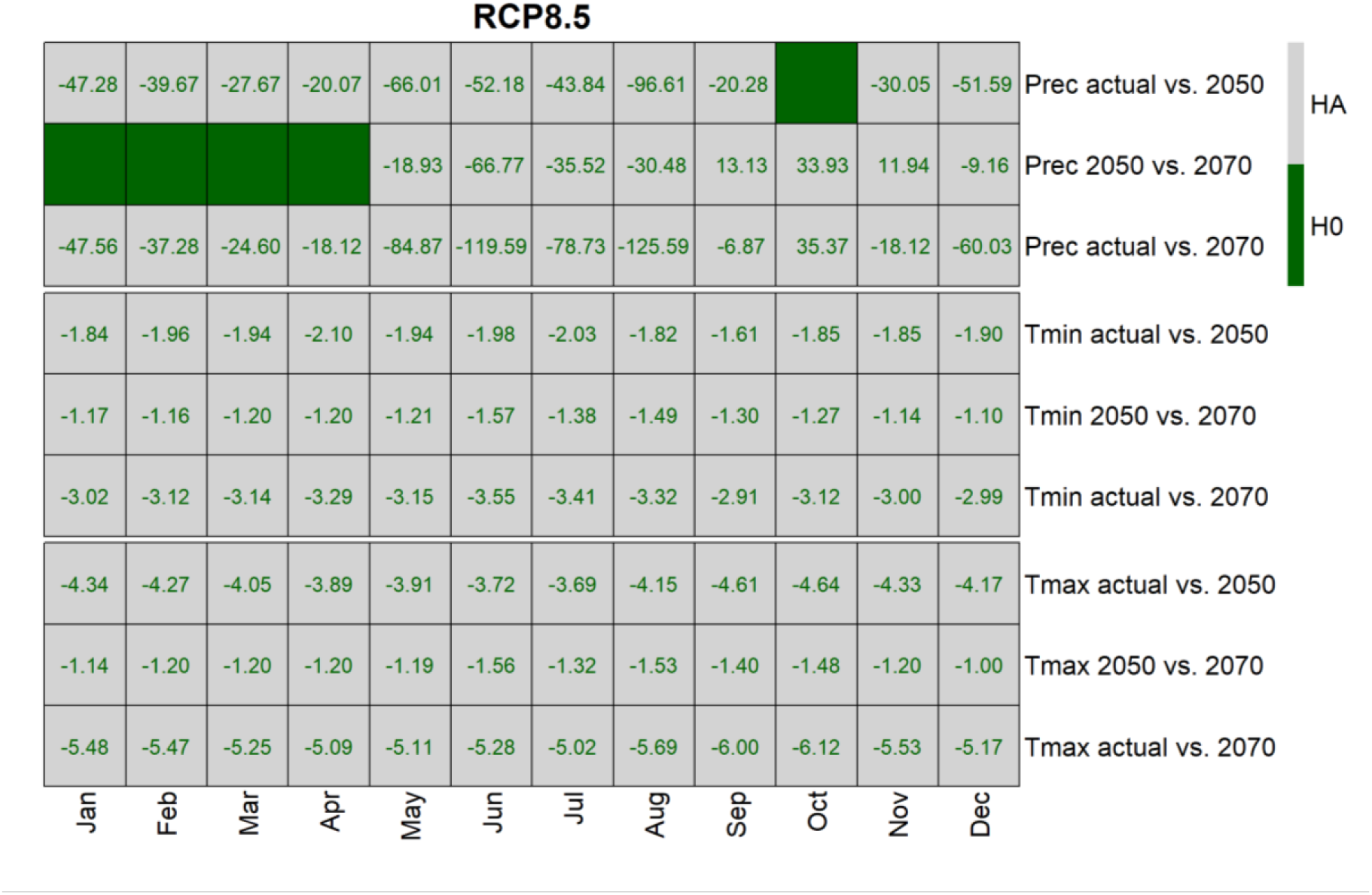
Results of Mann-Whitney U test for significance of difference in monthly precipitation, minimal and maximum temperature within the current distribution of *M. mercedesiarum* in current conditions and projection of HadGEM2-ES model under RCP8.5 scenario for years 2050 and 2070. The color-filled cells represent acceptance of null hypothesis (H0), blank cells correspond to acceptance of alternative hypothesis (HA) with numeric values reflecting difference of medians between two samples.

The rainfall change in species current distribution area is not particularly straightforward. The rainfall projections for years 2050 and 2070 generally have augmented spatial variability in comparison to current WorldClim 2 data and increase in absolute values of monthly rainfall from current conditions to 2050 (Fig. 3C, 4C, 5, 6). In the RCP4.5 scenario, the increase of the median of monthly rainfall between actual conditions and projection to 2050 was significant for all months, varying from 4.97 mm in October and to the highest value of 93.65 mm in August. When comparing actual rainfall with 2070 projections, a significant increase was found in all months except October; the increase is from 8.7 mm in April to 93.76 mm in August. The monthly rainfall increase in the RCP8.5 scenario is generally higher, with the exception of October, when we did not observe a significant change between actual and projected 2050 values. The lowest rainfall increase between actual and 2050 conditions was observed in April (20.05 mm) and the highest in August (96.61 mm), and when looking for changes from actual conditions to 2070 projections, it varies from 6.87 mm in September to 125.59 mm in August, with diminution of rainfall median in October (35.37 mm less). The annual rainfall (predictor BIO12) is significantly higher in future projections comparing with WorldClim 2 data, with a median of 2728 mm in 1970–2000, predicted 3379 mm (RCP4.5) or 3201 mm (RCP8.5) in 2050, and 3255 mm (RCP4.5) or 3293 mm (RCP8.5) in 2070.

The comparison of 19 bioclimatic predictors within the current distribution of *M. mercedesiarum* had shown significant differences between current conditions and projections for future scenarios for all variables (Appendix S2: Fig. S1, S2).

### Species distribution modelling in current conditions

The species relative habitat suitability produced in ten SDM cross-validation replications for current conditions appears to be strongly correlated with source distribution of *M. mercedesiarum*, represented by presence and absence points, as indicated by the mean Spearman’s rank correlation coefficient of 0.447. The correlation was highly significant because p-values were much lower than 0.05 in all replications. In threshold independent model evaluation, the mean real AUC value was 0.948 between ten replications, which can be interpreted as high overall ability of discrimination between species presence and absence. The null-model AUC rank test with the model real AUC located at the highest position for all replications demonstrated that the species is not independent from the used environmental predictors, and the inferred model is significantly better than by chance models with given presence prevalence (Fig. 7A).

**Figure 7.**
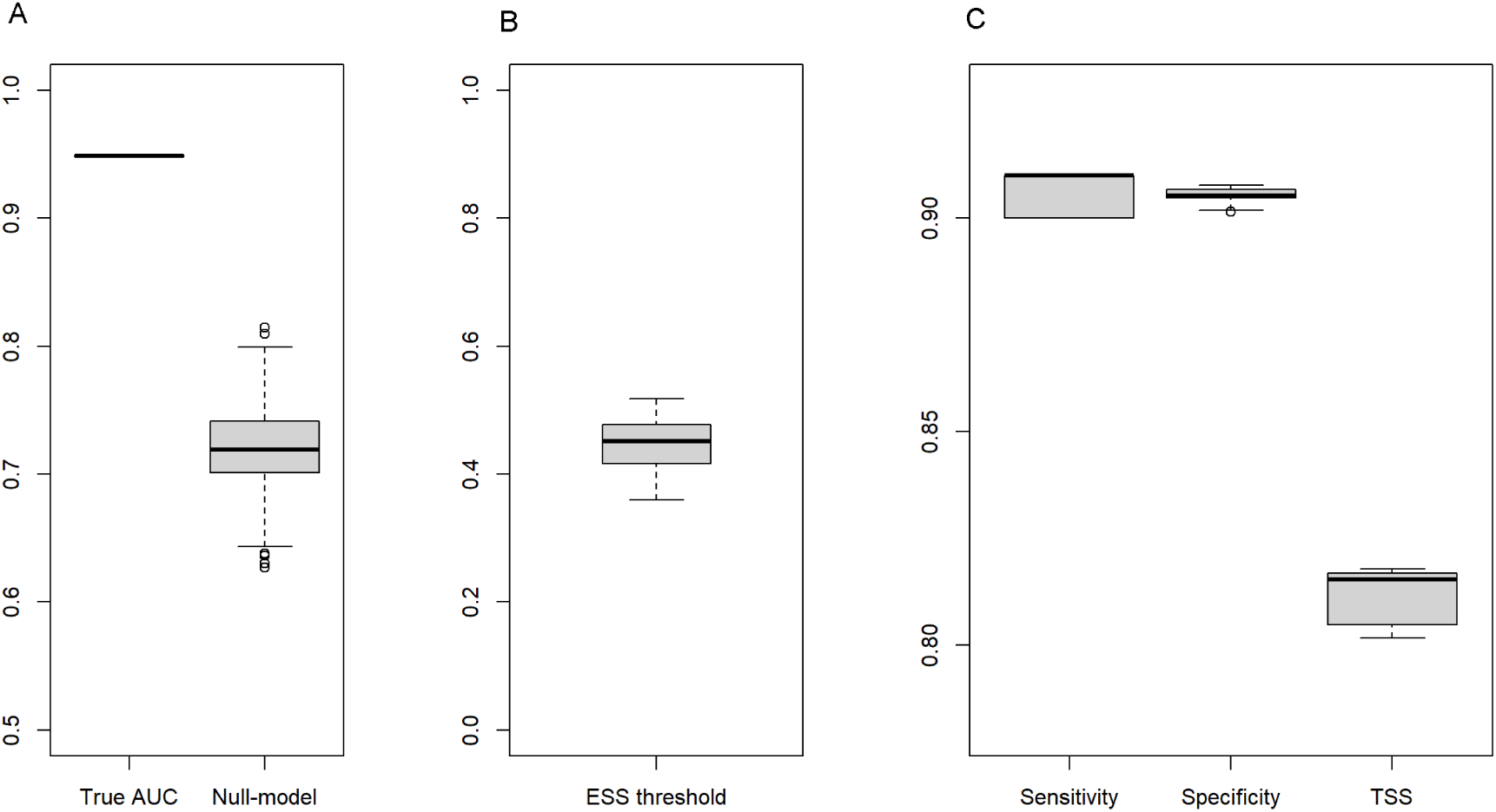
SDM evaluation metrics. A) Observed true AUC vs. set of null models AUC. B) ESS threshold. C) Sensitivity, specificity, and TSS.

The number of significant predictors selected by the LASSO process in independent model replications varied from 14 to 17. The analysis of generality of predictor contributions allowed to identify nine variables (“bio3”, “bio5”, “bio6”, “bio10”, “bio12”, “bio15”, “bio18”, “bio19” and “alt”) with non-zero contributions in all model replications; the same predictors demonstrated mean importance in permutations higher than 2% (Fig. 8). Six predictors from this list (“bio3”, “bio6”, “bio15”, “bio18”, “bio19” and “alt”) were found to contribute more than 2% each, and the sum of mean contributions for these six variables was estimated as 94.04%.

**Figure 8.**
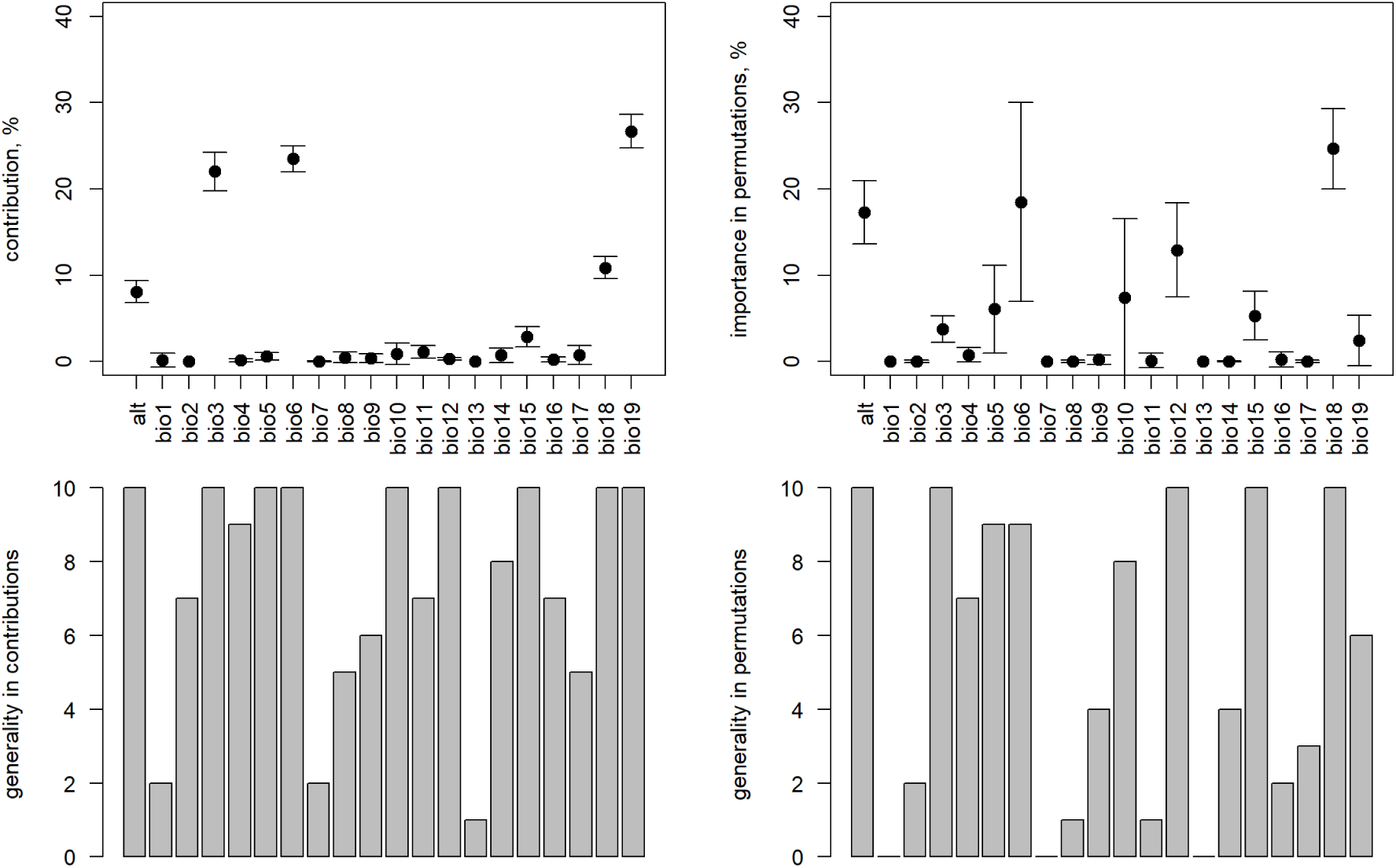
Predictor contribution, importance in permutations and generality in replications of SDM inferred from true presence and absence points.

The ESS threshold individually adjusted from the receiver operating characteristic curve in each model replication for current conditions varied from 0.360 to 0.518, with a mean value of 0.446 (Fig. 7B). The mean sensitivity and specificity values used in threshold estimation were correspondingly 0.906 and 0.905. The overall model performance under the selected threshold measured as TSS varied from 0.802 to 0.818, with a mean value of 0.811 (Fig. 7C). The threshold dependent evaluation demonstrated a high model reliability for current conditions because 89.6% of grid cells had a presence probability above the ESS threshold within the known species distribution area and 90.5% of grid cells were correctly classified within the buffer area of 500 km from current distribution.

The extent of area with high relative habitat suitability in model predictions for current conditions is larger than the known source distribution and includes a number of disjunct sites located in Colombia. Within the buffer zone of 500 km from the current distribution, the size of area with relative habitat suitability above the ESS threshold in the majority of replications is 5843 km^2^. The predicted high suitability area may include areas with environmental conditions similar to those of known species presence sites and areas where the model produces extrapolation to novel environments. The degree of extrapolation and detection of areas of novelty was analyzed using six covariates with contribution higher than 2%. The combined type 1 and 2 novelty indices for current conditions (Table 1, Appendix S2: Fig. S3) shows no significant extrapolation in high suitability areas immediately adjacent to known current species distribution, but some grade of novelty for disjunct areas with high habitat suitability. Only a few grid cells with type 2 novelty were found in the model extrapolation within the buffer of 500 km from known species presence locations. The high habitat suitability surfaces with evidence of extrapolation were estimated as 2101 km^2^ in the entire analysis extent and as 1150 km^2^ in the 500 km buffer zone. The contribution of type 1 novelty is of 98.2%, while type 2 novelty is observed in 1.8% of extrapolated grid cells.

**Table 1.** Areas with relative habitat suitability higher than ESS threshold by extrapolation type for six of the most important covariates. The inclusion of grid cells was defined by majority rule in ten cross-validation replications. No data are included for RCP8.5 in 2070 because no areas with relative suitability higher than ESS threshold were predicted. A buffer of 500 km was defined by Euclidean distance from known species observations.

|  | Area of habitat suitability above ESS threshold (km<sup>2</sup>) | No extrapolation (km<sup>2</sup>) | Type 1 novelty (km<sup>2</sup>) | Type 2 novelty (km<sup>2</sup>) |
| --- | --- | --- | --- | --- |
| 1970-2000 full extent | 7117 | 5016 | 2063 | 38 |
| 1970-2000 (500 km buffer) | 5843 | 4692 | 1147 | 4 |
| 2050 (rcp4.5, full extent) | 923 | 263 | 651 | 9 |
| 2050 (rcp4.5, 500 km buffer) | 566 | 263 | 303 | 0 |
| 2050 (rcp8.5, full extent) | 145 | 0 | 136 | 9 |
| 2050 (rcp8.5, 500 km buffer) | 70 | 0 | 70 | 0 |
| 2070 (rcp4.5, full extent) | 292 | 0 | 286 | 6 |
| 2070 (rcp4.5, 500 km buffer) | 103 | 0 | 103 | 0 |

### Forecasting of the species distribution in future scenarios

The modelled species relative habitat suitability in the current distribution area of *M. mercedesiarum*, assessed through logistic MaxEnt output, demonstrates drastic diminution in grid cells with high suitability in both future scenarios. According to the developed model, in current conditions the median logistic output value was 0.723 within the known species distribution area, which corresponds to high relative habitat suitability in most of the current distribution (Fig. 9). Under the RCP4.5 scenario in 2050, the median of grid cell logistic values reduced to 0.184, and in the RCP8.5 scenario for the same future moment, this median value was only 0.099. The habitat suitability in 2070 was lower than in 2050, with a median of logistic output as 0.086 and 0.027 under RCP4.5 and RCP8.5, respectively.

**Figure 9.**
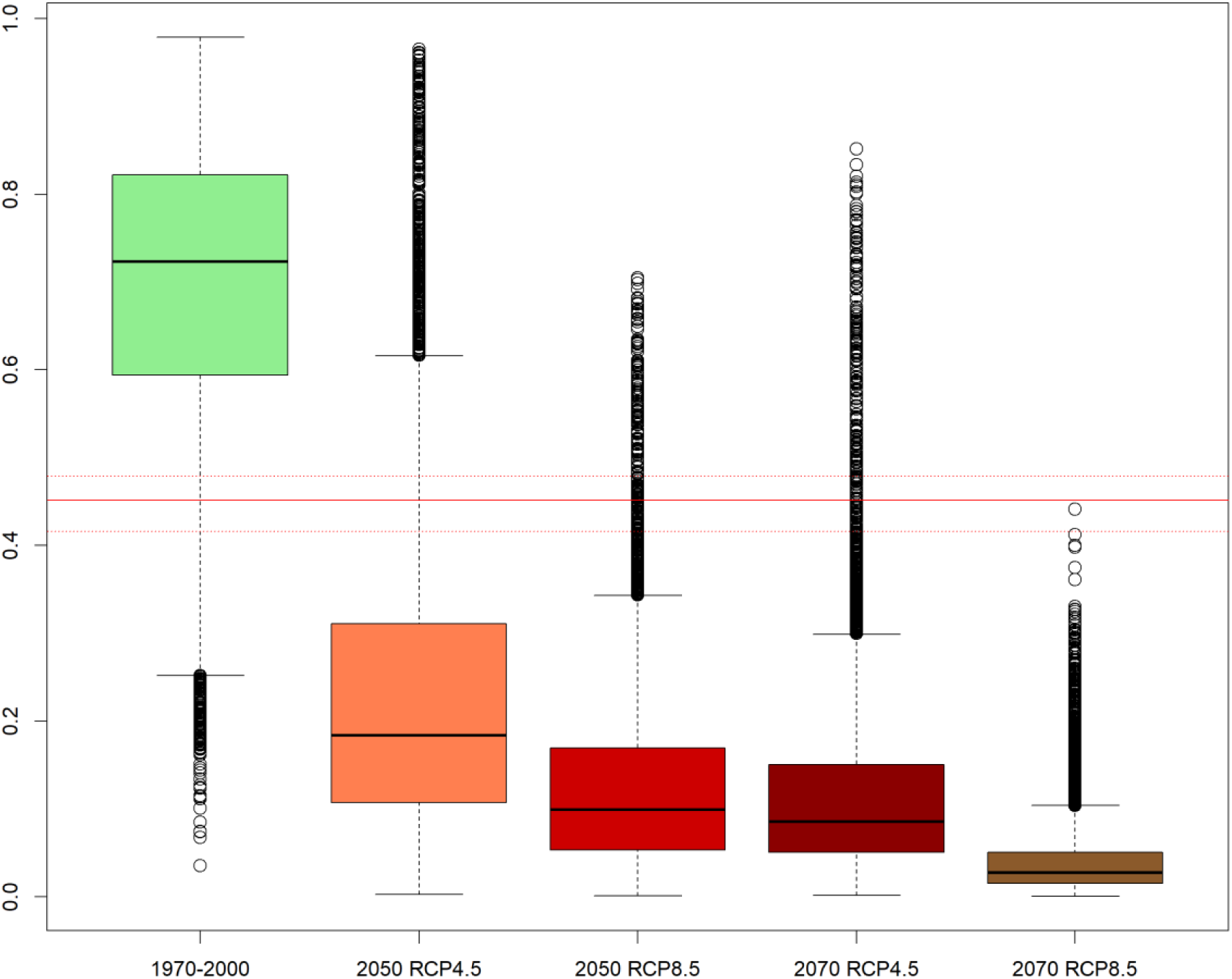
Logistic model output as proxy for species presence probability within the current distribution area of *M. mercedesiarum* in current conditions and in projection of HadGEM2-ES model under RCP4.5 and RCP8.5 scenarios. The horizontal line as ESS threshold median (continuous line) and 95% confidence interval (dotted lines).

The application of a threshold to identify areas with probable species presence showed that the minor fraction of grid cells within the known current distribution area retains values of relative habitat suitability above the ESS threshold (Fig. 10, Supporting Information Data S5). Under the RCP4.5 conditions in 2050, the persistence of species was estimated with a mean of 13.3% of total grid cells known to have species presence today. Considering the model uncertainty in classification of pixels for the current conditions within the surrounding area, the expected mean habitat loss under RCP4.5 scenario in 2050 was estimated as 89.6%, varying from 60.6% to 100.0% in independent model replications. The RCP8.5 scenario led to almost complete disappearance of suitable habitat during the same period because only a mean of 2.7% grid cells retained logistic output values above the ESS threshold. One model replication in this case was considerably different from the majority, with 26.4% of grid cells classified as being suitable for species presence. The mean habitat loss under this scenario in 2050 was estimated as 98.1%. For 2070, the mean predicted persistence of suitable grid cells was estimated as 2.6% and 0.1% of the known current distribution under RCP4.5 and RCP8.5 scenarios, respectively.

**Figure 10.**
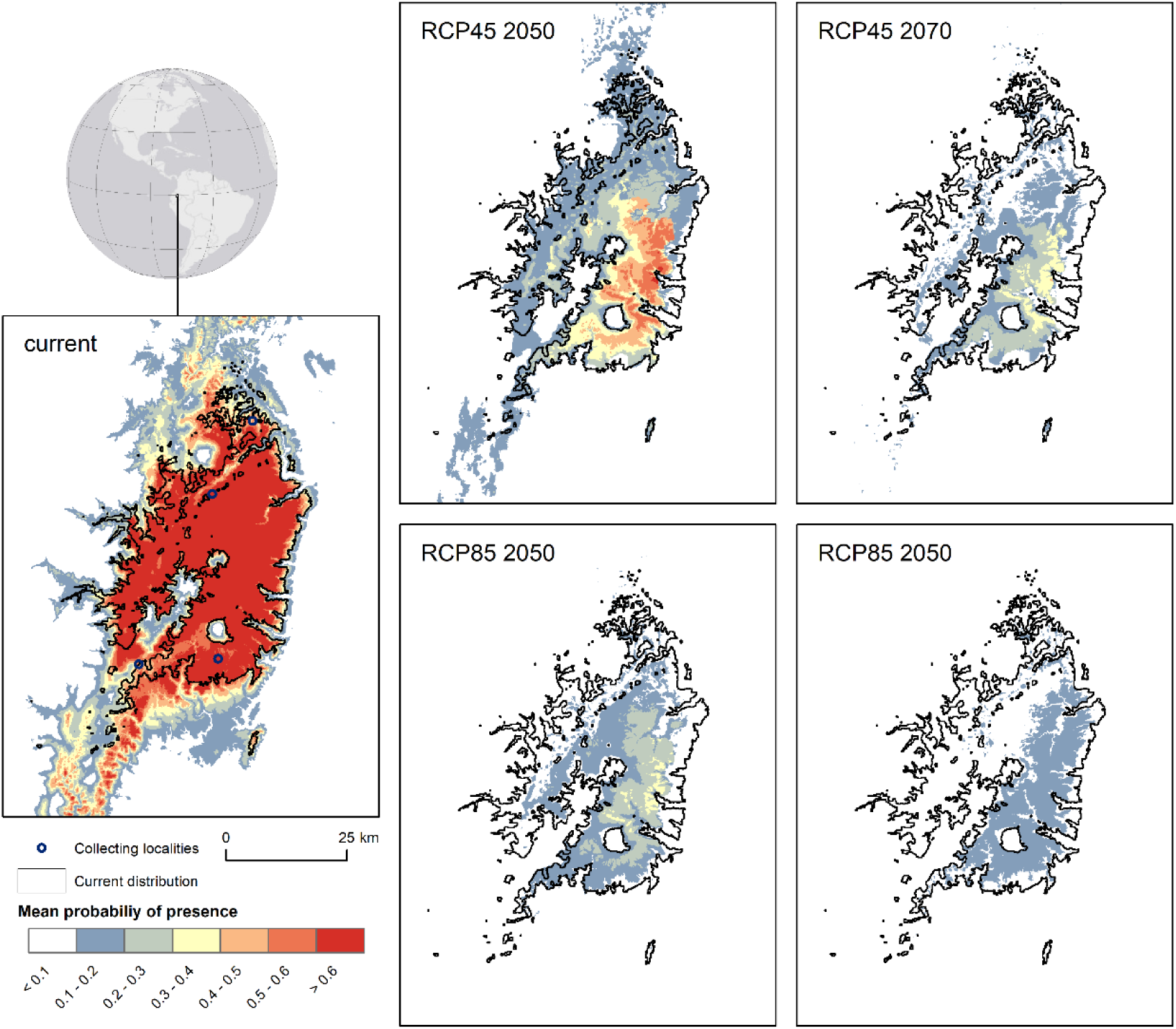
Maps of presence probability in area of known *M. mercedesiarum* distribution modelled for current conditions and for 2050 and 2070 under HadGEM2-ES RCP4.5 and RCP8.5 scenarios. Area closed by continuous line represents the current species distribution.

The overall suitable area for species presence in northwestern South America, estimated by a method of predominant value from presence grids in replications, includes grid cells in both, inside and outside of the current species distribution. While the source current distribution extent is known to have 2701 km^²^ and the model prediction of suitable area in current conditions with extrapolation exceeds 7117 km^²^ in future projections it is drastically reduced (Table 1). The estimated total suitable area under RCP4.5 in 2050 is 906 km^²^, but only 263 km^²^ of this area is produced without model extrapolation. In 2070, under the same scenario all highly suitable areas are a result of extrapolation and are predicted to have an area of 292 km^²^. Under RCP8.5 scenario for 2050, the highly suitable area is estimated to be 145 km^²^, the entire extent extrapolated, and no suitable area was found under this scenario for 2070.

A large fraction of positively classified cells in future scenarios was found to have positions separate from the current known distribution. Modelling allowed to detect several areas persistent between scenarios at the same geographical locations with a significant increase of relative habitat suitability compared to current conditions. Two main disjunct geographic localities of high habitat suitability were detected to the north of the current distribution area, both located in Colombia, on the eastern slopes of the Andean cordillera (Fig. 11, Appendix S2: Fig. S4, S5, Supporting Information Data S5). One of these localities with a center approximately at 2.5° N latitude and 77° W longitude is separated from the current distribution area by a distance of approximately 230 km by straight line. At this locality, the size of area with presence probability above the ESS threshold at least in single model replication was estimated in different scenarios as 319 km^2^ (RCP4.5 2050), 109 km^2^ (RCP4.5 2070), 74 km^2^ (RCP8.5 2050). Another persistent area with high presence probability is located northeast from the current distribution separated by a distance of approximately 590 km by straight line, with its center at 4.5° N latitude and 73.5° W longitude. In this location, the estimated size of area with suitable habitat was estimated as 337 km^2^ (RCP4.5 2050), 175 km^2^ (RCP4.5 2070) and 41 km^2^(RCP8.5 2050).

**Figure 11.**
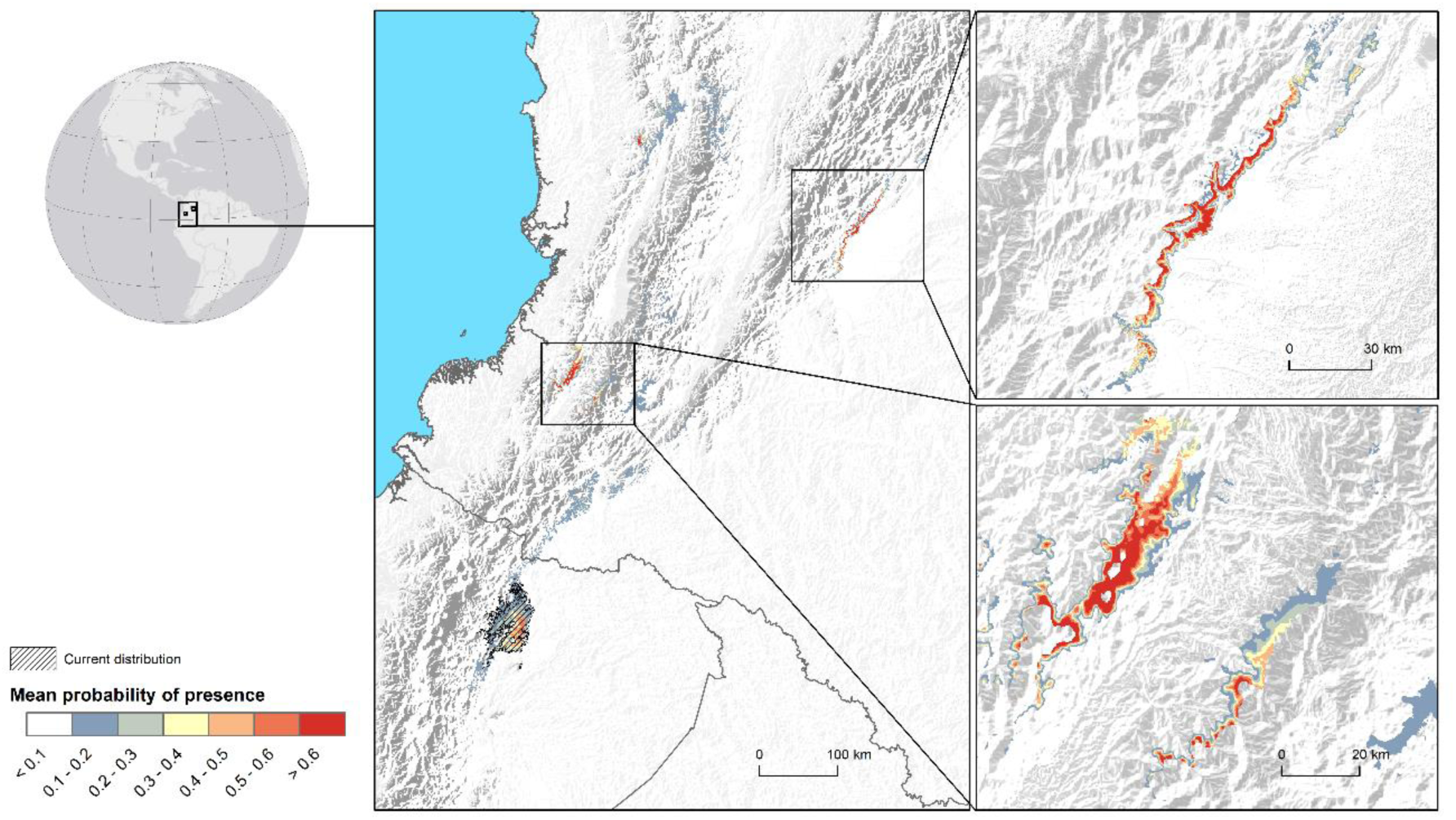
Potentially suitable habitat of *M. mercedesiarum* in disjunct locations outside of species current distribution, based on HadGEM2-ES model data for RCP4.5 scenario in 2050. Basemap shaded relief was derived from GMTED2010 elevation data. Rectangular areas and line pointing to locator map indicates position of zoomed map fragments.

### Evaluation of species distribution ranges

The vast majority of 55 known *Magnolia* species from Colombia and Ecuador listed in Table 2 have a narrow distribution range and can be classified as regional endemics. *Magnolia mercedesiarum* belongs to this majority with a maximum distance between observation localities reaching 85 km. After excluding species known from a single locality, the median of maximum species extent is estimated as 116 km; the histogram is evidently skewed to the left, with 23 species located in the first class for less than 100 km wide distributions (Fig. 12). Only 11 species have a distribution extent larger than 300 km, including a few shared with neighboring countries. However, few species presented moderately vast distributions, reaching the extent of 1549 km (*Magnolia rimachii*), 1562 km (*M. sambuensis*) and 1351 km (*M. chimantensis*).

**Table 2.**
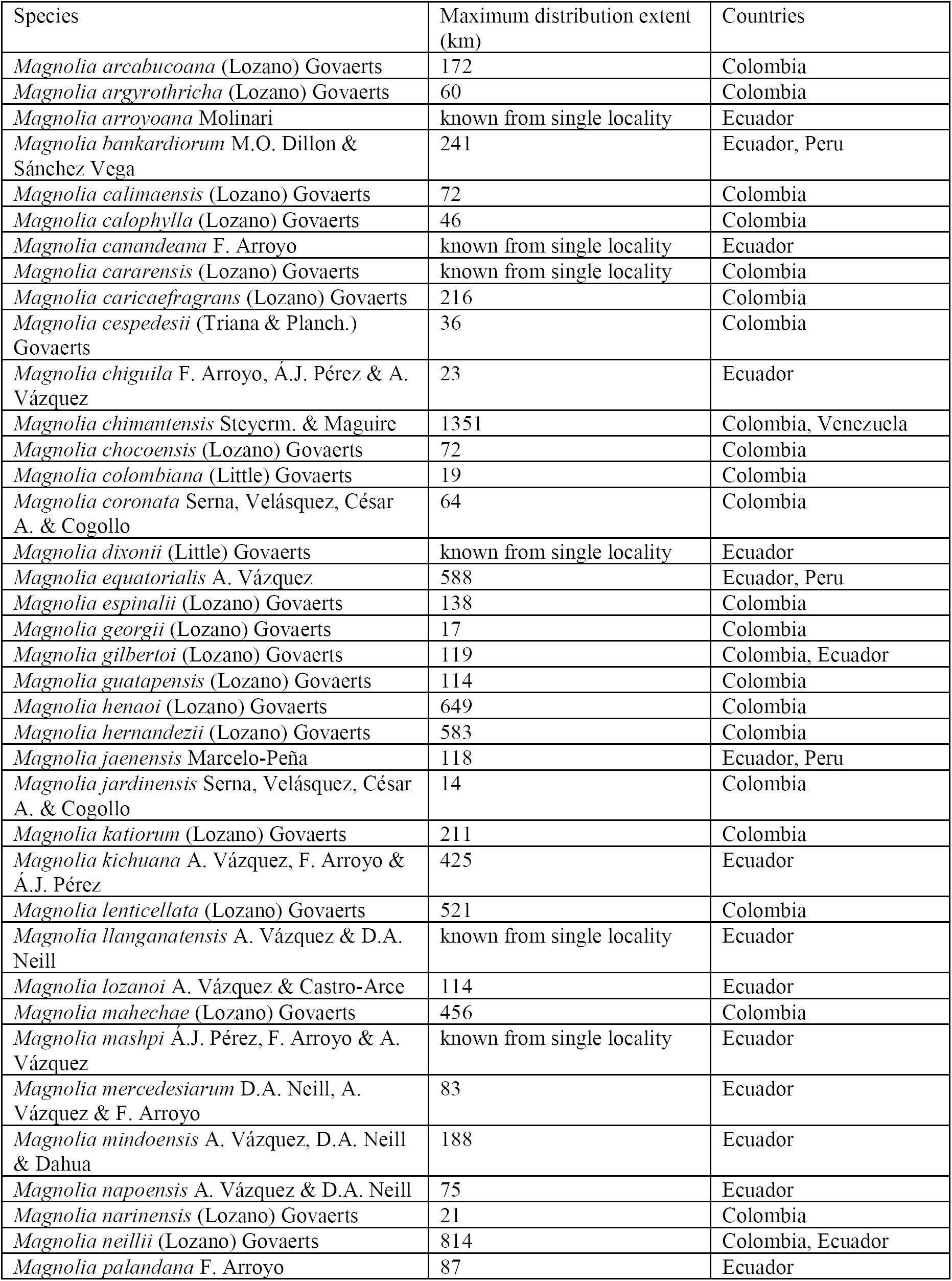
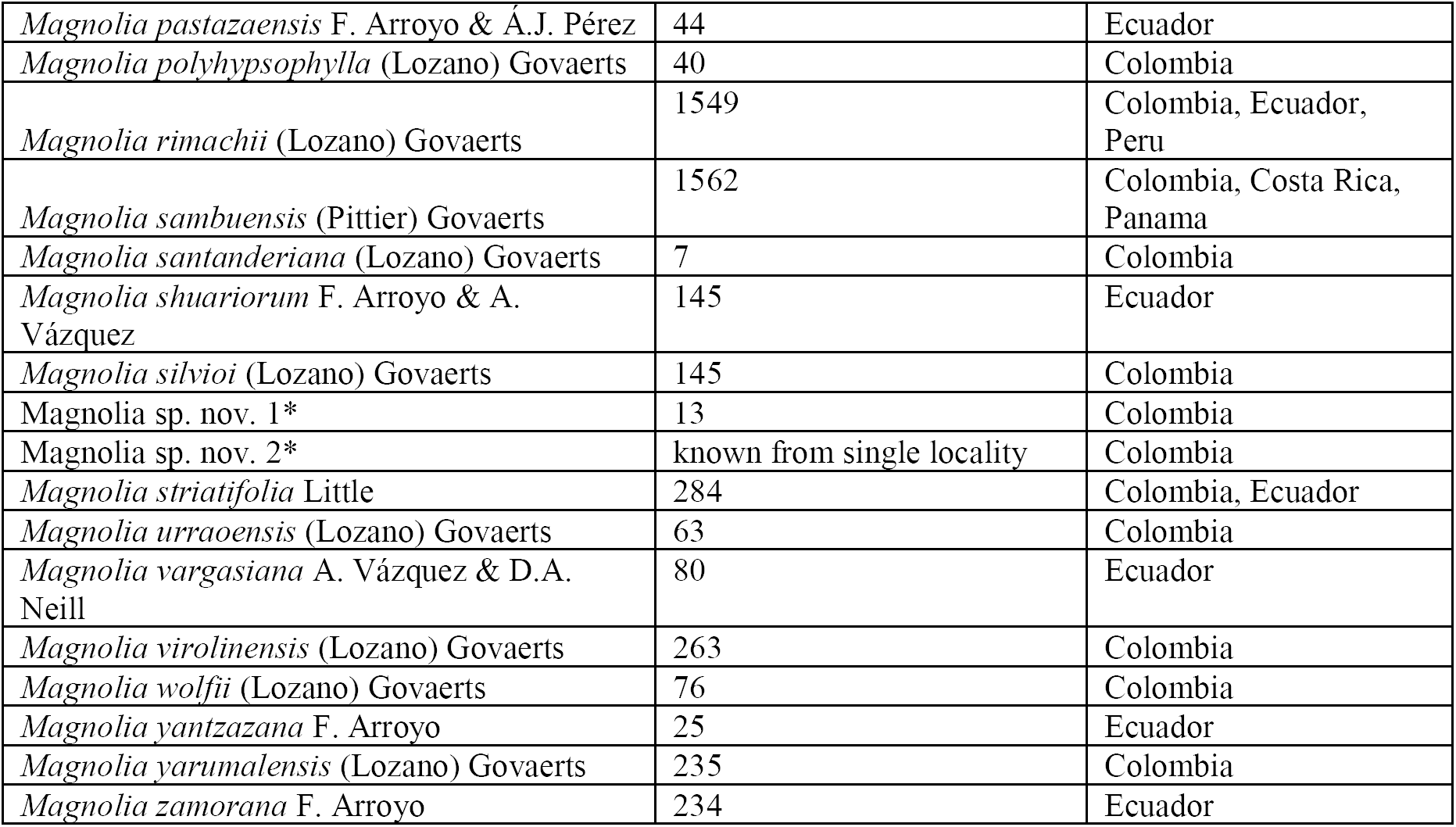
Distribution extent for *Magnolia* species from Colombia and Ecuador measured as maximum Euclidean distance between observation points. The asterisk (*) for two undescribed species as cited in GarcÍa (2007).

**Figure 12.**
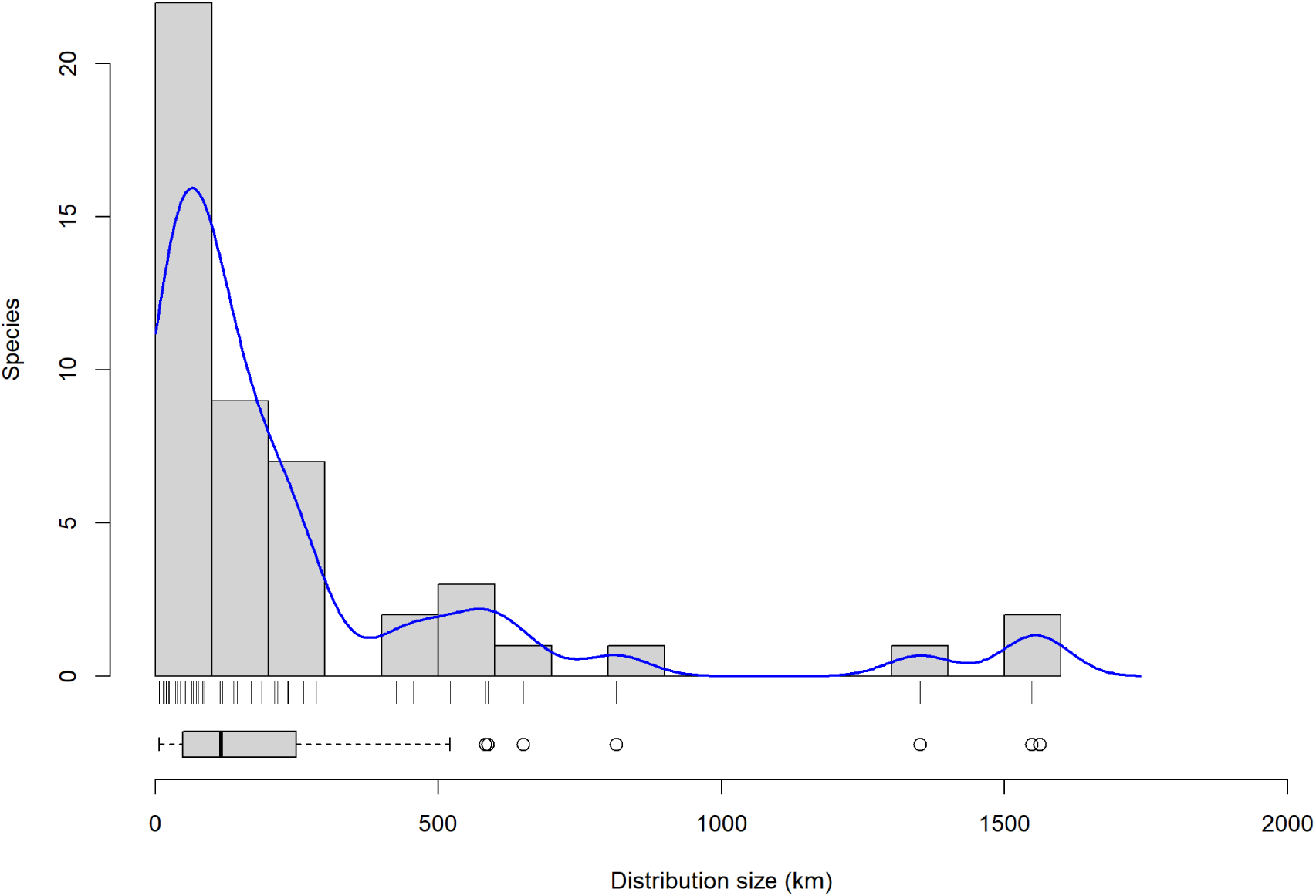
Histogram of distribution extent for *Magnolia* species from Colombia and Ecuador measured as maximum Euclidean distance between observation points for each species. Blue line represents density curve transformed to values of expected species counts.

## Discussion

The model of habitat conditions potentially suitable for a species was assessed in this study through the modelling of response to downscaled WorldClim 2 climatic data and represents the partial model of ecological niche for the species under consideration. The edaphic, geologic, and potentially many other environmental factors were excluded due to a lack of data with acceptable resolution and quality, and a lack of knowledge of the dynamics of such factors at the scope of decades. The effect of biotic interactions is assumed to be homogeneous at a coarse scale of 7.5″ grid cells within the entire continental area at the extent of analysis. The biotic interaction at a fine scale that can be easily recognized as heterogenous is the competition between trees, particularly young individuals and saplings competing for light, but the relative frequency of gaps in the canopy turns out to be essentially similar when comparing grid cells of more than six hectares each, with the condition that they belong to the same vegetation. This assumption is consistent with Soberón’s (2010) finding that the effect of biotic interactions, defined as competition between species and simulated at fine spatial scale, is effectively averaged at coarser scales (grid cells 8 or 16 times larger), where scenopoetic variables become only significant predictors. Soberón stresses that biotic factors could still have an effect at coarse resolution because this depends on relative spatial structure of different groups of factors in constitution of species’ ecological niche and correlation between them. We admit that the spatial variability in biotic interactions, such as canopy gap frequency, and other phenomena related to the biotic environment can be manifested at coarse scales, as a function of vegetation type. However, we do not include the vegetation type layer as a predictor in the analysis, for the same reasons as to why the edaphic and other non-climatic environmental layers were excluded.

Fortunately, the vegetation types are known to have strong correlation with climatic conditions (Donoghue & Edwards, 2014), hence this variability can be indirectly captured by pattern in rainfall and temperature data.

The bioclimatic envelope modeling fits well in the hierarchical modelling framework (Pearson & Dawson, 2003), providing information on regional-scale species distribution limitations, and when spatial resolution of predictors is fine enough, it provides the ability to assess the effect of climate variability correlated with topography at a local scale. Guisan and Thuiller (2005) consider that in the case of plant species as a kind of sessile organism, fine predictor resolution could significantly improve niche models because such species could strongly depend on predictor variability at a local scale. The selection of spatial resolution and scale is of critical importance for the SDM process not only due to its implications for the Eltonian noise hypothesis, but also to identify the groups of environmental factors that could have an effect and what patterns could be detected. The entire extent of SDM in the current study correspond to the regional scale, where the climatic and topographic variability is expected to have a leading role in the definition of species distributions, while the spatial resolution of 7.5″ that we use as the finest units of analysis correspond to transition between landscape and local spatial scales. Following Willis and Whittaker’s (2002) conceptualization, at the local spatial scale the effect of topographic variables is expected to mix with land-use, edaphic, and biotic factors, as well as the fact that climatic inference cannot be separated from the topographic component. Franklin et al. (2013) note that the climate variability on a scale of hundreds of meters is primary controlled by topography and can exhibit variability that is high enough to constitute refugia for local populations, as well as an increase in habitat connectivity for dispersal and migration. As demonstrated in Franklin and co-workers’ study, the precision in representation for ecologically significant microhabitats does increase with a change from coarse to fine scale, and somewhere between the 4 km and 800 m resolution there is a threshold beyond which further spatial generalization of climatic predictors severely affects predicted suitable habitat area. The generalization errors produced at coarse resolutions are especially high for narrow-range species. The finest resolution modelled by Franklin et al. was 90 m, but the difference between 90 m and 270 m resolutions in location of sites with suitable habitat was minor, at least for tree species considered in the study. The authors speculate that for some species with habitat somewhere in rugged terrain, the resolution of 10 m to 30 m may be needed to accurately capture microhabitats’ variability. However, the resolution required to model microhabitats for tree species may be coarser because the size of individual trees contributes to averaging of the effect from climatic fluctuations at a very fine scale. The tradeoff between the spatial resolution and the model’s ability to represent fine microhabitat structure is required because the finer resolution predictors significantly increase the computational requirements and processing time. In the case of *Magnolia mercedesiaruum*, the adult trees can be more than 20 m tall, similar to the majority of tree species in Franklin and colleages’ (2013) study. We assume that the resolution of approximately 250 m selected in this study is fine enough to capture topographic controlled climate variation in mountainous regions for the tree species under consideration. The further increase in predictors’ resolution may require addition of edaphic and land-use related variables at a fine scale because the effect of such variables on the local environment is expected to be high.

Although the environmental niche model produced in this study is partial, it demonstrates high correlation in geographic prediction with distribution estimation from Vázquez-GarcÍa et al. (2018), produced with the same predictors, treated here as true species distribution. Independently from the fidelity of the source species distribution estimation, the research question addressed in this article is how such narrow distribution could change in HadGEM2-ES climate projections for two CO2 emission scenarios.

To be able to answer this question, we followed the equilibrium assumption for relations between species and its environment in 1970–2000 climatic conditions. The 1970–2000 climatic conditions as represented in the WorldClim 2 dataset were treated as representative for this period of relative climatic stability. In particular, we assumed that the current distribution of *M. mercedesiarum* has formed during the period of relative climatic stability in the scope of centuries and species populations had persisted at the same geographical location at least for several generations; thus, it had the possibility to occupy the entire available potential habitat within its dispersal limitations and source-sink dynamics. The indirect argument in favor of stable status of populations is the manifested reproductive function. The lack of reproduction function may be an indication for populations that they may not belong to stable “core” areas with habitat condition characteristics for the species’ fundamental niche, but their belonging to less stable sink areas that depend on individuals’ immigration from sources, at the current moment or in the immediate past. All observations of *M. mercedesiarum* cited in Vázquez-GarcÍa et al. (2018) were made on trees bearing flowers or fruits, although no saplings were noted beneath the observed trees. Guisan and Thuiller (2005) affirm that populations without evidence of sexual reproduction are preferable to exclude from the training data in SDM because they can be assumed to lack the positive growth rate for population size and may be located beyond the specific climatic thresholds. In the case of *M. mercedesiarum*, the manifested reproductive function in all populations used as training data in Vázquez-GarcÍa and colleagues’ means that the estimated distribution represents the area with presumed positive population growth rate, and thus locations with source habitat conditions in the source-sink process (Pulliam, 2000).

The projection of the same niche model to future scenarios is possible by additionally assuming the immutability of potential ecological niches for the species in a given time scope of 50 or 70 years. We consider that such an assumption is highly probable because the distance in generation between currently growing trees and individuals in the future vegetation does not exceed one or two generations. Under the hypothesis of niche conservatism, the species tend to maintain an inherited response to environmental conditions. The assumption of conservatism works well in combination with an assumption of equilibrium in the model calibration time scope. If any of this key assumption is violated, the forecasting of future climatic conditions becomes problematic. One problem derived from violation of the equilibrium assumption is the possibility that the variability of environmental response in the population can be much wider than the manifested in the currently observed area of distribution. Such conditions may occur when a species that had recently experienced a partial loss or change of the distribution but had not reached an adaptive equilibrium with the new environment, maintaining hidden ecological plasticity. In this last case, the adaptation potential will allow fast adaptation to new conditions available within the dispersal range, even if such conditions do not occur within the species’ current distribution, but is compatible with an inherited fundamental niche. However, there are no data in favor for the presence of hidden ecological plasticity of *M. mercedesiarum*. At the same time, the projected diminution of suitable habitat conditions in 2050 and 2070 may open the possibility for the described situation, when species would manifest narrower ecological plasticity in their realized distribution than in their inherited fundamental niche.

The projected dramatic diminution of habitat suitability in the current area of species distribution, with possible loss of at least 89% of the area above the species presence threshold does not mean an immediate disappearance of the species within most parts of its current location. It is expected that in 50 years, many of the currently adult trees will still persist in the same place as today, tolerating the projected climatic change because adult trees could have broader tolerance to climatic environmental conditions than tree saplings. Talluto et al. (2017) data on the lags between environmental changes and distribution shifts for species with slow demography and limited dispersal confirm the presence of “extinction debts” in many tree species in North America. We assume that a similar “extinction debt” may occur in tree populations under climate change elsewhere. The delayed extinction corresponds to a transition of individual trees from source to sink zones in source-sink dynamics, and its exclusion from the geographical projection of fundamental niches. In the absence of closely located source areas, the extinction of such species in areas that had lost habitat conditions suitable for positive population dynamics may happen in the scope of the life of individual trees. In the case of tropical *Magnolia* species, such a delay may include several decades. The ecological niche projections from our point of view are more related to identification of sites where the species will be able to persist through reproduction, maintaining the capacity of tree saplings to establish, survive, and grow to adult trees in the competitive environment of a wet tropical mountain forest.

The capacity of species migration into novel localities depends on their intrinsic characteristics, such as the inherited dispersal syndrome. The dispersal syndrome is an evolutionary character that evolves within a given linage and has a profound impact on species dispersal capacity, population dynamics, and speciation (Gibbs et al., 2010). There is a tradeoff between speciation rate and dispersal capacity, which can result in narrow range distribution patterns for species with an evolutionary regime of short-range dispersal. Heinz’ et al. (2008) modeling demonstrates that the short-range dispersal has a positive evolutionary feedback with incipient speciation, both in sexually and asexually reproducing populations. According to their data, the evolutionary regime with short-range dispersal and speciation particularly often evolves in conditions of steep environmental gradients. Additionally, the model of Heinz and colleagues demonstrates the presence of an abrupt transition between evolutionary regimes of short-range dispersal with speciation and long-range dispersal without speciation. In the current study, the actual distribution for each species of *Magnolia* is assumed to be a result of dispersal from a single ancestral population, molded by environmental factors, dispersal limitations, and biotic interactions. The environmental diversification within the landscape with steep environmental gradients should result in fast allopatric speciation within the lineages with a predominantly short-range dispersal regime. The observed pattern of predominantly narrow-ranged species agrees with the short-range dispersal with speciation evolutionary regime from Heinz et al. (2008). As shown in Table 2 and Fig. 12, the majority of *Magnolia* section *Talauma* species, from the northern Andes, have extremely narrow distribution ranges, typical for species that experience isolation within their native habitat in consequence of low dispersal capacity and low ecological plasticity.

Most species of *Magnolia*, with their oily seeds closed within a red-colored fleshy sarcotesta, display a bird dispersal syndrome (Callaway, 1994). In *M. mexicana* (sect. *Talauma*) from Veracruz, México, a great diversity of bird species disperses the seeds. Gutiérrez-Zúñiga and Jimeno-Sevilla (2017) found that 64% of 33 bird species visit the *Magnolia* trees, however the species has a very narrow distribution range of almost 150 km. Dispersal of seeds is not always related to distance but also may depend on heterogeneity of vegetation structure (Debussche et al., 1982) or bird behavior. For instance, required snags or perching sites by birds may indeed limit seed dispersal of *Magnolia* (McClanahan & Wolfe, 1987). However, few species of *Magnolia* display clonal growth of rapid increase (Silvertown & Charlesworth, 2001). In *M. tomentosa*, from Ise Bay, Japan, many seeds are found on the forest floor near mother trees and its clonal growth is more effective for genetic structuring of populations over short distances than the short-ranged seed dispersal (Setsuko et al., 2004). These narrow ranges of section *Talauma* of *Magnolia* have resulted in remarkable patterns of allopatric speciation in the Caribbean (Howard, 1948), Mexico and central America (Vázquez-GarcÍa, 1994), Colombia (Lozano-Contreras, 1994), and Ecuador (Vázquez-GarcÍa et al., 2016), with few species shared between countries. Several species with intermediately wide distribution range, such as *M. equatorialis*, *M. neillii*, and *M. rimachii* known from the upper Amazonian region of Ecuador could hypothetically present relation of their seed dispersal mechanism with hydrological features because they are found within a single hydrographic system.

*Magnolia mercedesiarum* can be expected to inherit dispersion limitations typical for section *Talauma*, which prevents the majority of neotropical *Magnolia* species from fast colonizing of distant localities with a suitable habitat. So far, the pattern of known distribution for *M. mercedesiarum* indirectly indicates the low dispersal capacity of this species. These dispersal limitations could prevent the colonization of disjunct localities with a suitable habitat that appear in results of modeling in 2050 and 2070. There is no known mechanism that could allow natural transfer of *M. mercedesiarum* seeds to locations separated from the current distribution area by a distance of almost 370 km or more, with the presence of several orographic barriers between locations. No direct hydrological connection exists between the known species locality and distant areas of suitable habitat. The stochastic dispersion models could be useful to simulate the bird-mediated seed dispersion process in a precise form. The initial hypothesis for such modelling could postulate the low probability of seed dispersal into areas with suitable habitat separated by hundreds of kilometers in a time scope of 30 to 50 years.

## Conclusions

The potential reduction of area with suitable habitat, in rates from 89% to almost 100% in all climate change scenarios for 2050, can lead to further extinction of species in areas of known current distribution. The scope of this extinction will not have immediate visible effects on adult trees because the loss of suitable habitat conditions could primarily affect the reproduction and establishment of saplings. The species survival in 2070 under scenario RCP4.5 is constrained to its persistence as a healthy population only in small refugia with a size 2.6% of its current distribution area. The dispersal to separate distant high habitat suitability zones, projected in 2050 northwards from the current distribution is remotely probable, given the species’ seed dispersal syndrome.

This study contributes to conservation in tropical forests by identifying the vulnerability of tree species from the understorey of tropical mountainous forest in climate change projections. Contrary to common assumptions of distribution shift within altitude gradients in terrains with complex topography (e.g. Rumpf et al., 2018), we found another pattern of distribution change, mostly by reducing the size of the suitable area within the current distribution range and emerging new disjunct zones with high suitability but unreachable by natural dispersion. These findings can be used in planning of biodiversity protection for mountain ecosystems in the Neotropics.

Our approach for SDM here and in Vázquez-GarcÍa et al. (2018) contributes to improvement of methods for climate change response prediction for narrowly ranged species known from few observations. The use of climatic information at a high resolution (∼250 m) spatial scale allows the detection of spatial variability and refugia that could not be discovered at coarser spatial scales. At the same time, the selected spatial scale, still is controlled primarily by topographically driven abiotic factors, makes possible ecological niche modelling derived from bioclimatic variables downscaled from the output of global circulation models.

## Supporting information

## Acknowledgements

The research was partially funded by CONACYT scholarship No. 256271/275105 for Viacheslav Shalisko.

## Appendix S1

### Description of presence data and spatial autocorrelation test

The estimation of the current distribution area in Vázquez-GarcÍa et al. derives from a presence-only MaxEnt driven model, which was performed with 20 cross-validation iterations and was configured to account for spatial uncertainty and environmental variability in the location with four known populations of *M. mercedesiarum*. The collecting bias in this SDM for narrowly distributed species was addressed by using target background sampling and independent randomization during cross validation steps (Vázquez-GarcÍa, Neill, Shalisko, Arroyo, & Merino-Santi, 2018a). From the perspective of model accuracy and completeness, the minimum required number of presence records depends on species prevalence in the landscape but cannot be lower than 14 for narrowly ranged species in realistic environments (van Proosdij, Sosef, Wieringa, & Raes, 2016). Dealing with only four presence records, Vázquez-GarcÍa et al. performed the randomized sampling of environmental conditions around presence points with spatial resolution of 250 m, which allowed taking advantage of the expected spatial autocorrelation in species presences for improvement of environmental variability representation in the training dataset. Spatial autocorrelation derives from intrinsic properties of biological processes at the landscape level, that cause spatial aggregation of populations (Dormann et al., 2007).

The spatially autocorrelated presences could be a valuable source of information on species habitat variability for SDM based on Hutchinson’s duality because the model actually is not produced in the geographic space, but in niche hypervolume, so the highly spatially aggregated data that do not provide significant spatial information could provide information of predictors’ variability. However, there are two problems associated with autocorrelated data in statistical modeling: 1) such data provide less information on variability than completely independent observations, 2) the assumption of residuals’ independence could be violated, which can result in biased model accuracy parameter estimates and increase of type I error rates (Dormann et al., 2007). In this study, we considered that spatial autocorrelation of presence data cannot be avoided but should be evaluated. To determine multivariate spatial autocorrelation in the training dataset, we used the generalization of Wartenberg’s test on duality diagrams described in Smouse and Peakall (1999). For each iteration in cross-validation process, the full set of predictors at random training points within species distribution area were evaluated for multivariate spatial autocorrelation in a Monte-Carlo randomization process, to produce a multivariate spatial autocorrelation index and test it against the null-hypothesis of absence of spatial autocorrelation. The implementation of multivariate spatial autocorrelation was taken from the R ‘multispati.randtest’ function from the ‘ade4’ package (Dray, Dufour, & Thioulouse, 2018).

The spatial autocorrelation index for training data demonstrated positive spatial autocorrelation in all cross-validation iterations (null-hypothesis refuted since the p-level was less than the significance level of 0.05). The values of autocorrelation index and distribution of eigenvalues can be found in Supporting Information Data S2.

### Literature Cited in Appendix S1

Dormann, C. F., J. M. McPherson, M. B. Araújo, R. Bivand, J. Bolliger, G. Carl, A. Davies, R.G. Hirzel, W. Jetz, W. D. Kissling, I. Kühn, R. Ohlemüller, P. R. Peres-Neto, B. Reineking, B. Schröder, F. M. Schurr, and R. Wilson. 2007. Methods to account for spatial autocorrelation in the analysis of species distributional data: a review. Ecography 30: 609–628.

Dray, S., A.-B. Dufour, and J. Thioulouse. 2018. Analysis of ecological data: exploratory and Euclidean methods in environmental sciences: R package version 1.7-11 ‘ade4’. http://CRAN.R-project.org/package=ade4 Smouse, P. E., and R. Peakall. 1999. Spatial autocorrelation analysis of individual multiallele and multilocus genetic structure. Heredity 82: 561–573.

Vázquez-GarcÍa, J. A., D. A. Neill, V. Shalisko, F. Arroyo, and R. E. Merino-Santi. 2018a. Data from: *Magnolia mercedesiarum* (subsect. *Talauma*, Magnoliaceae): a new Andean species from northern Ecuador, with insights into its potential distribution. Dryad Digital Repository. https://doi.org/10.5061/dryad.s5f28

## Appendix S2

**Figure S1.**
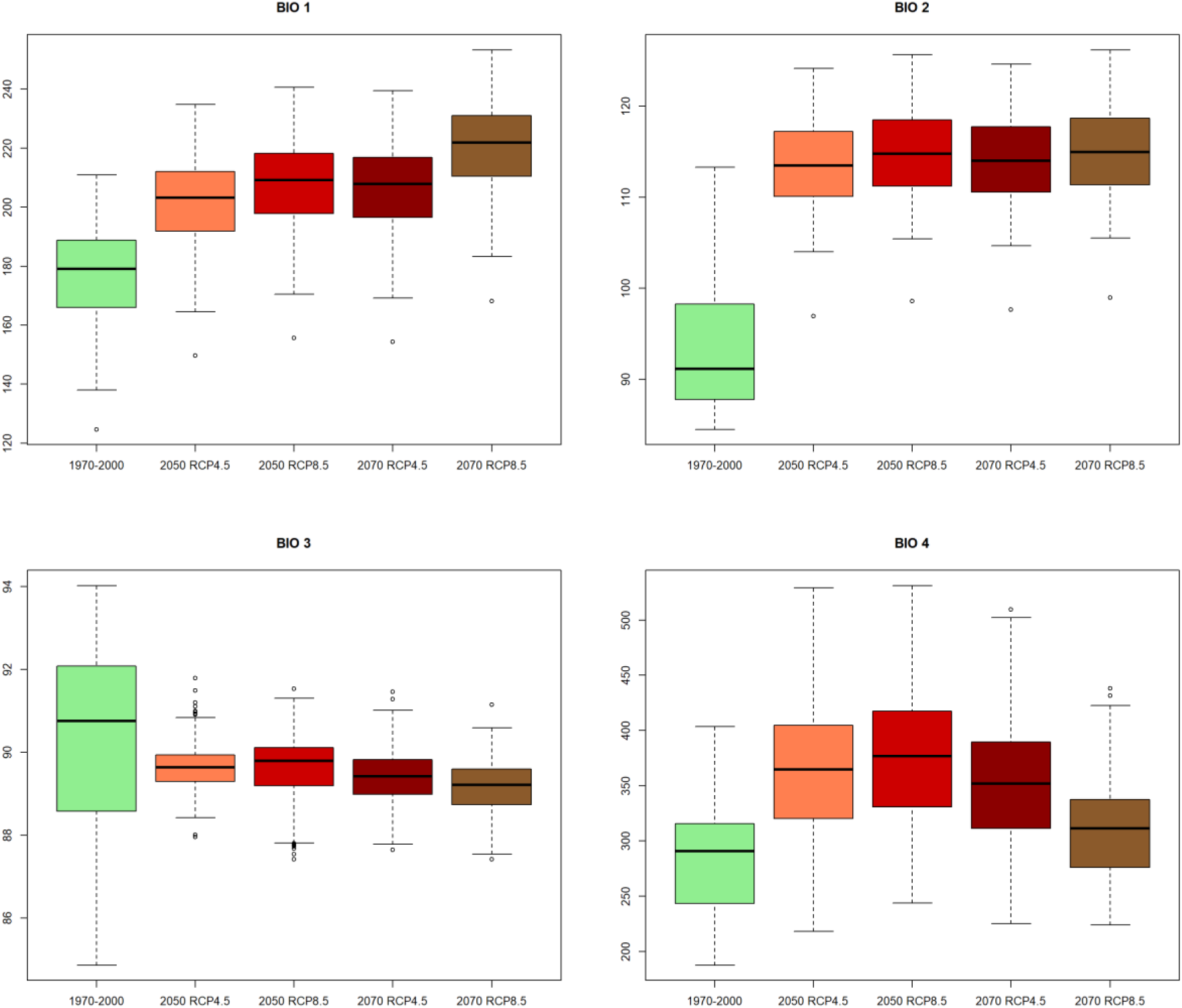
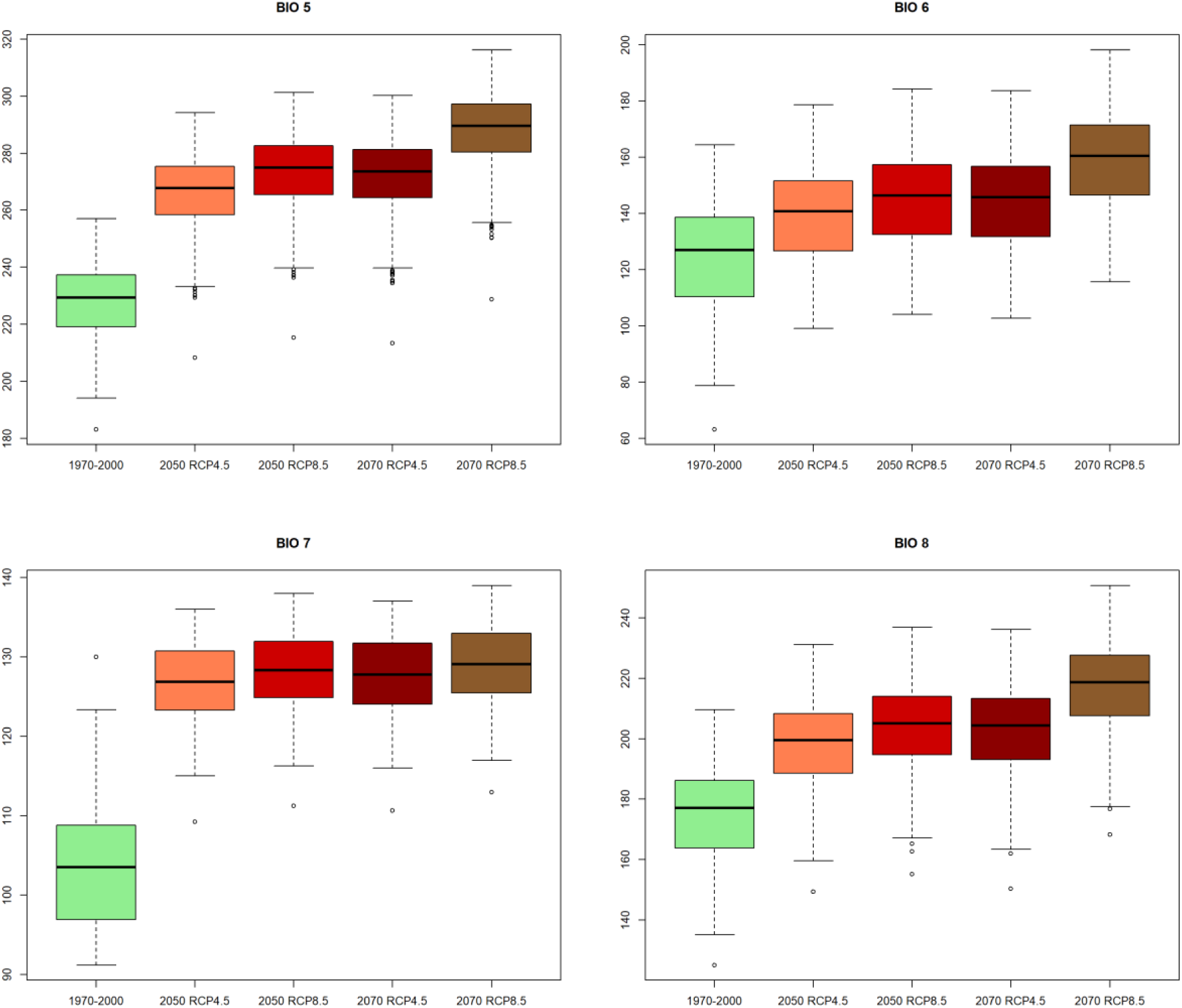
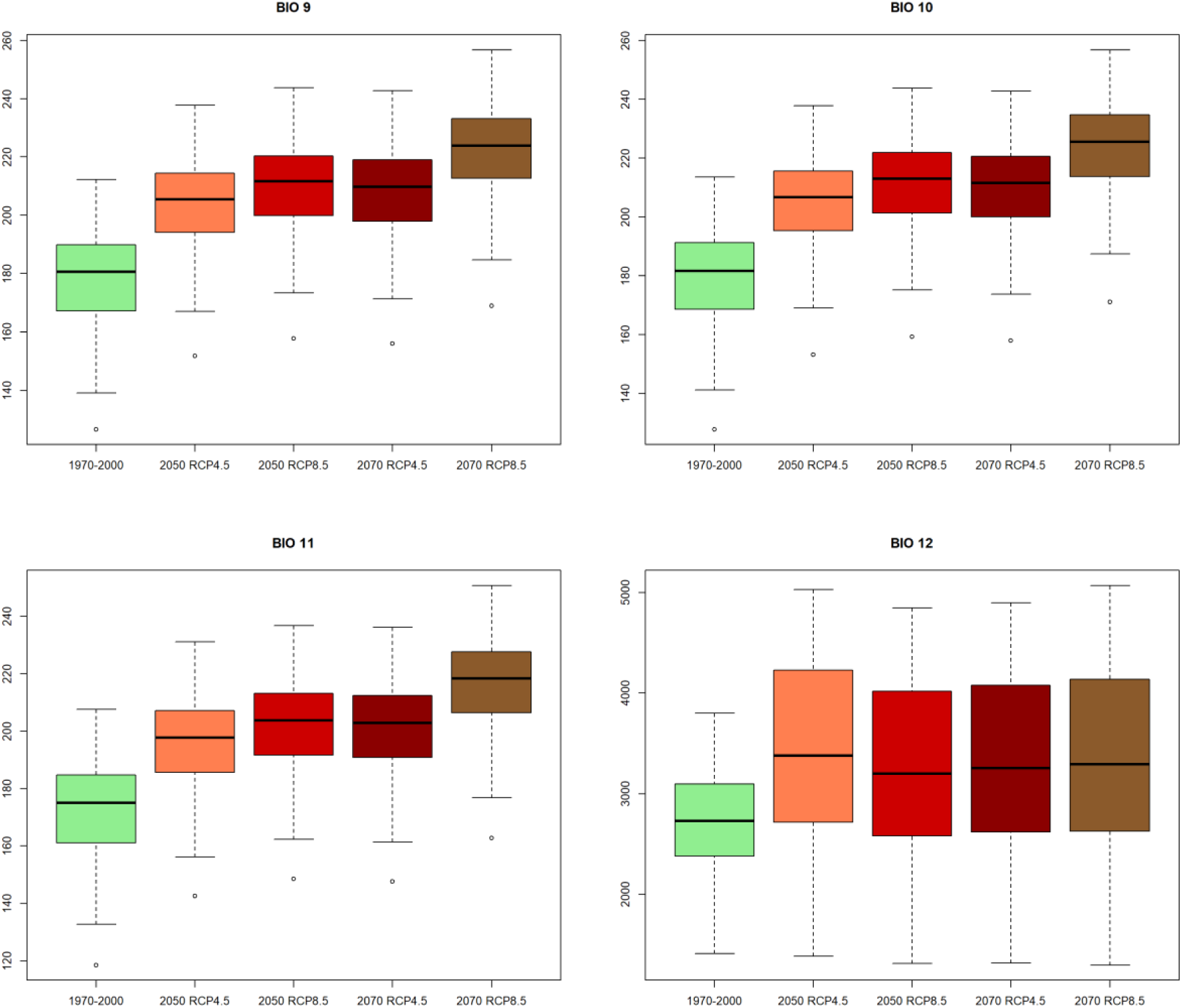
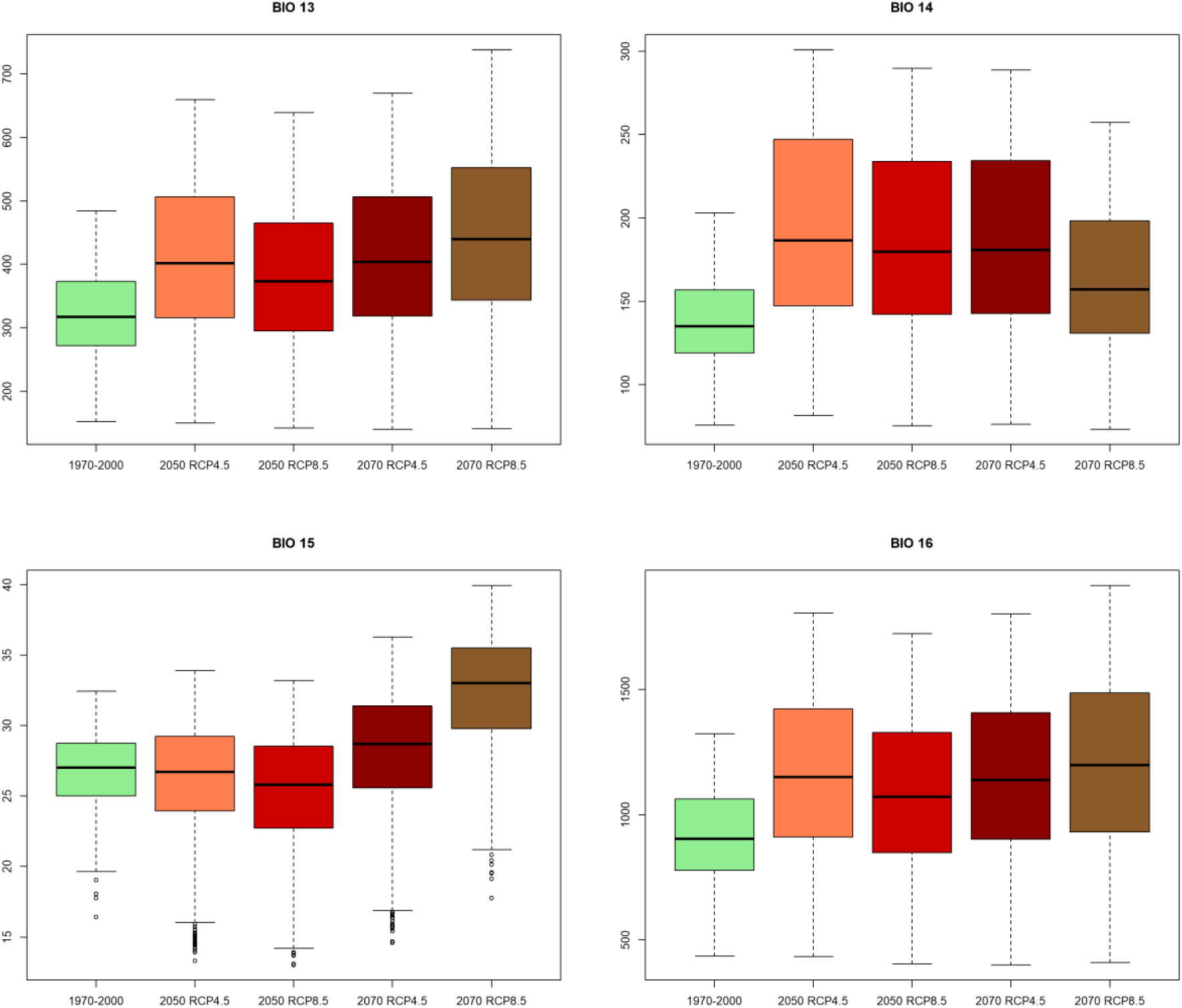
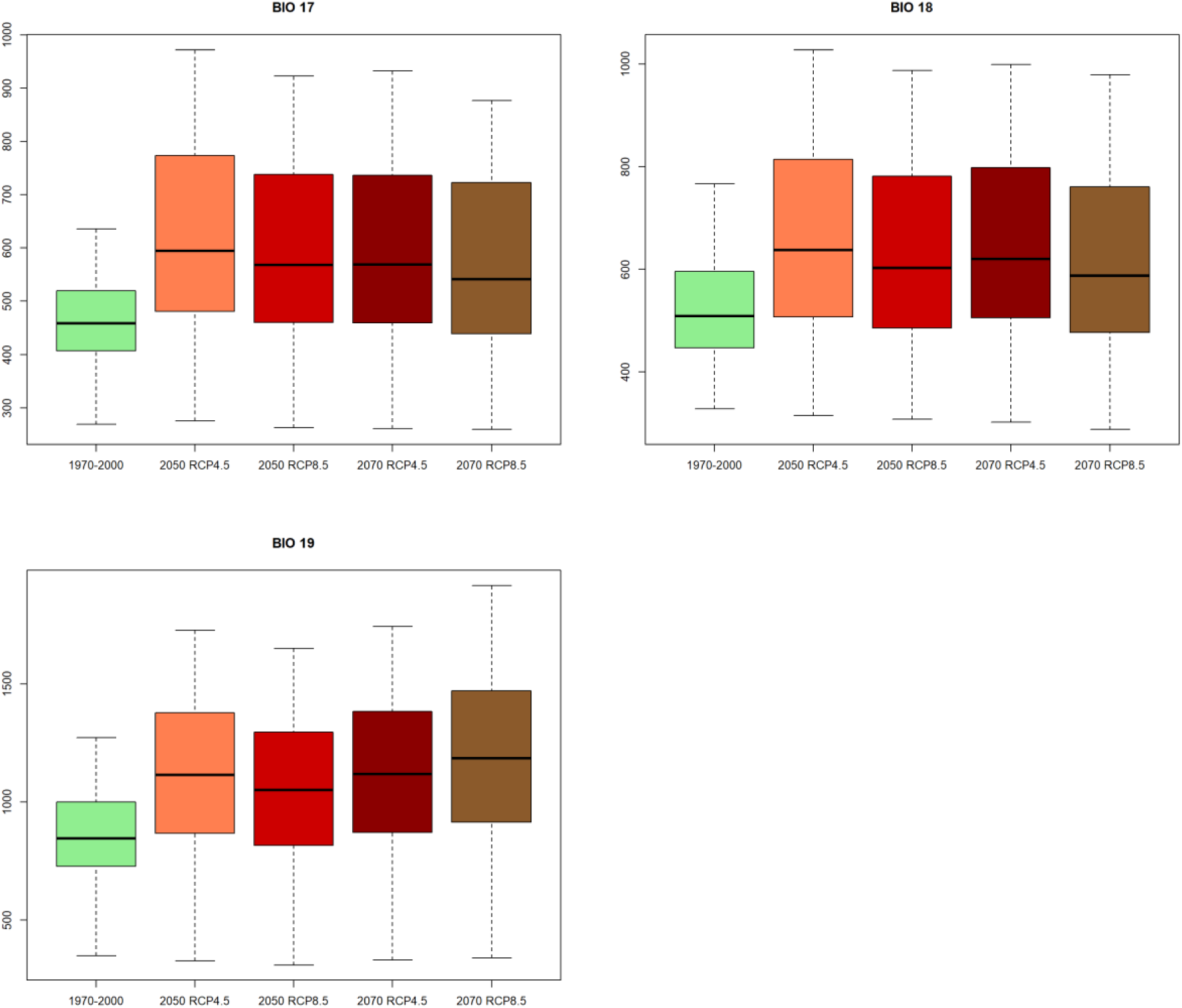
The comparison of 19 bioclimatic predictors within current distribution of M. mercedesiarum in current conditions and projection of HadGEM2-ES model under RCP4.5 and RCP8.5 scenarios for years 2050 and 2070.

**Figure S2.**
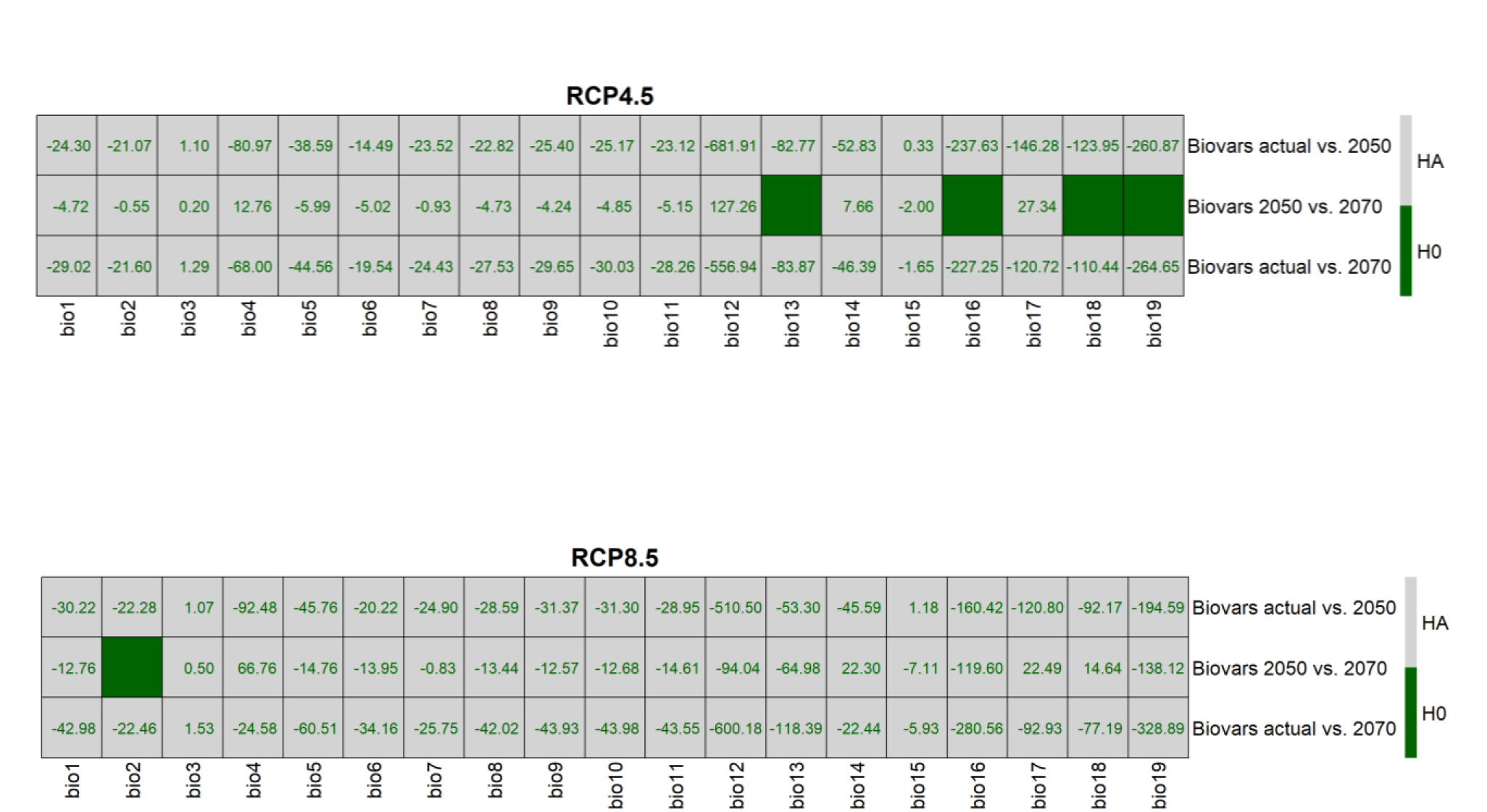
Results of Mann-Whitney U test for significance of difference in 19 bioclimatic predictors within the current distribution of *M. mercedesiarum* in current conditions and projection of HadGEM2-ES model under RCP4.5 and RCP8.5 scenarios for years 2050 and 2070. The color-filled cells represent acceptance of null hypothesis (H0), blank cells correspond to acceptance of alternative hypothesis (HA) with numeric values reflecting difference of medians between two samples.

**Figure S3.**
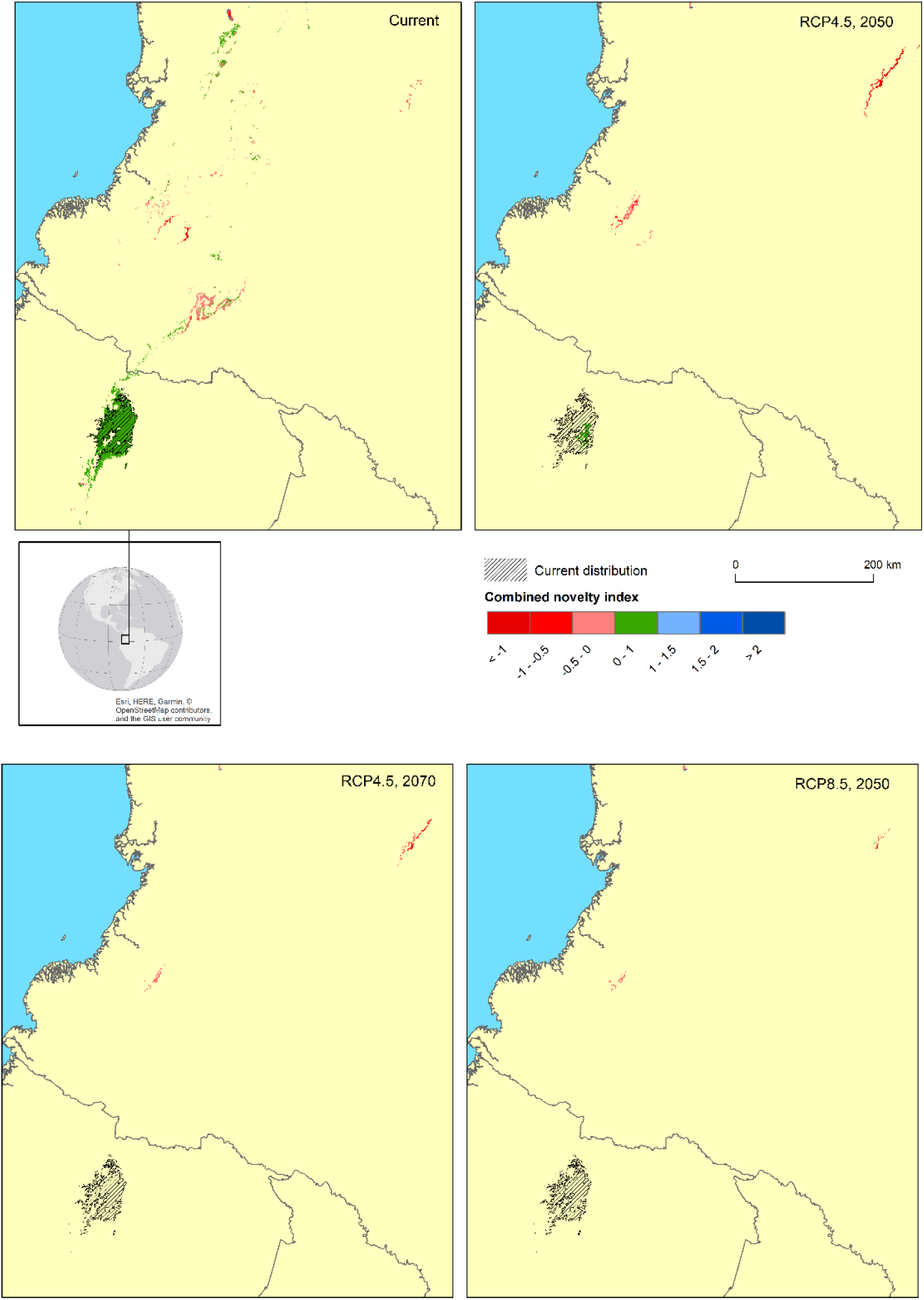
Combined novelty 1 and 2 index for areas with predicted relative habitat suitability higher than ESS threshold, majority in ten cross-validation replications. Green areas represent absence of extrapolation, red – type 1 novelty, blue – type 2 novelty. Results for prediction in current conditions (1970–2000 climatic data), RCP4.5 in 2050 and 2070, RCP8.5 in 2050. No areas with relative suitability higher than ESS threshold were predicted for RCP8.5 in 2070.

**Figure S4.**
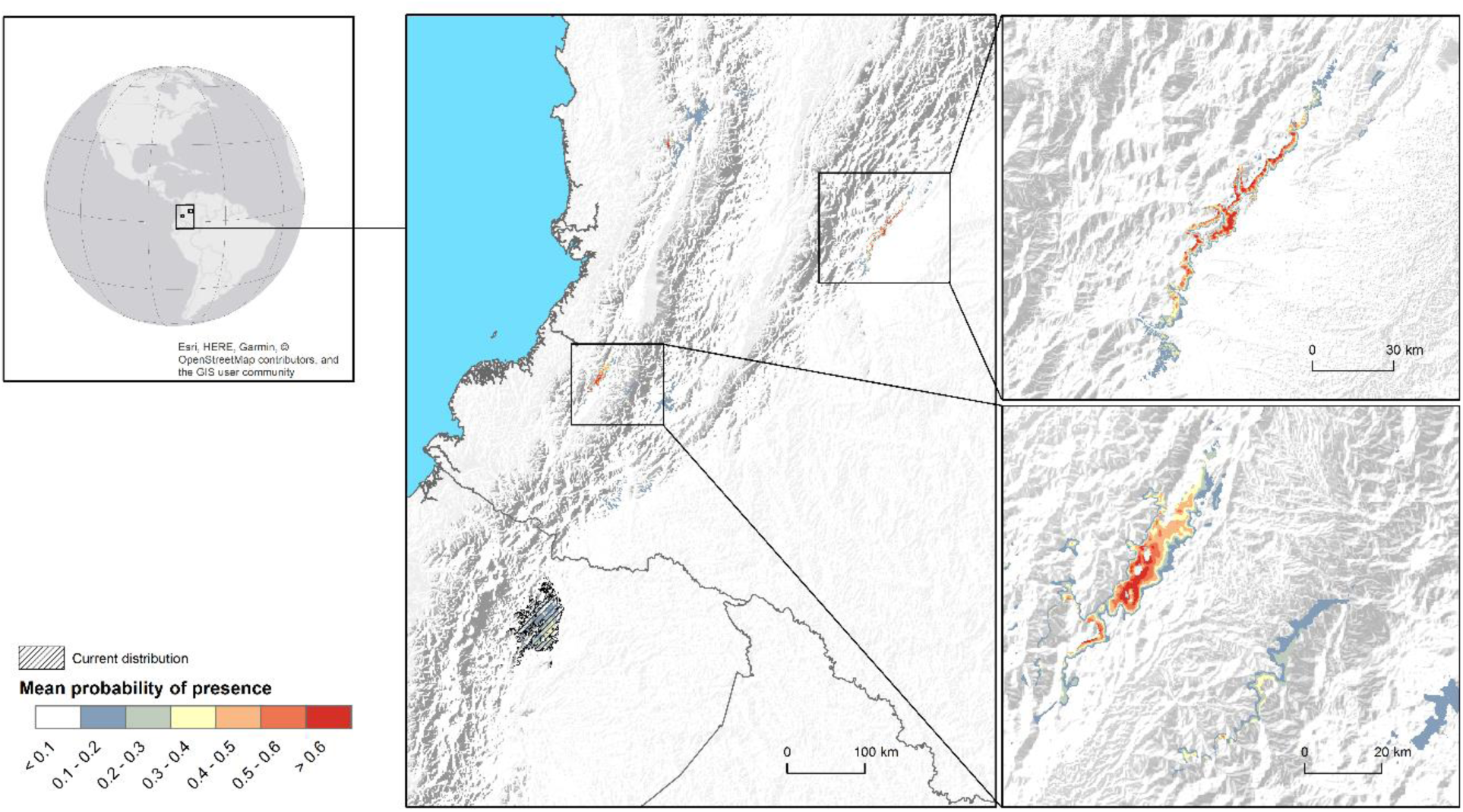
Potentially suitable habitat of *M. mercedesiarum* in disjunct locations outside of species current distribution, based on HadGEM2-ES model data for RCP4.5 scenario in 2070. Basemap same as in Figure 11.

**Figure S5.**
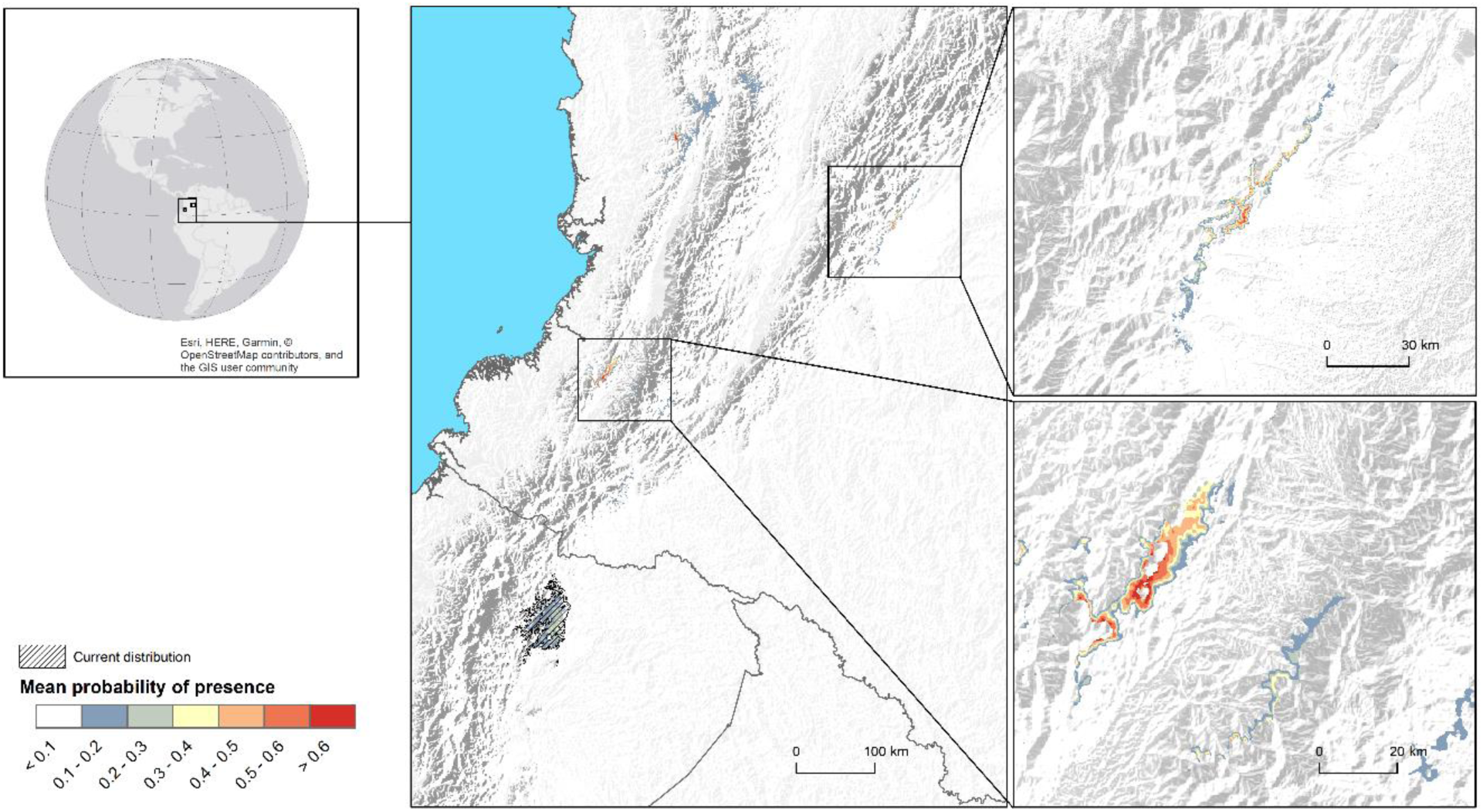
Potentially suitable habitat of *M. mercedesiarum* in disjunct locations outside of species current distribution, based on HadGEM2-ES model data for RCP8.5 scenario in 2050. Basemap same as in Figure 11.

