## Supplementary material for "Vulnerability to climate change for narrowly ranged species: the case of Ecuadorian endemic *Magnolia mercedesiarum*"

### Index of Supporting Information

- Appendix S1. Description of presence data and spatial autocorrelation test. Included in manuscript.
- Appendix S2. Additional figures. Included in manuscript.
- Data S1: *Magnolia mercedesiarum* current distribution
    - Polygon after application of ESS threshold, taken from Vazquez-García et al. (2018), file M_mercedesiarum_ETSS.shp
    - Raster mask of species presence with same resolution as predictor variables, file M_mercedesiarum_logistic_raster_clip.tif
- Data S2: Computer code for estimation of spatial autocorrelation index in predictor variables within the presence mask, and HTML report of script execution with results
    - Source code as R Markdown, file spatial_autocorrelation_test.Rmd
    - Result of code execution, file spatial_autocorrelation_test.html
- Data S3: Computer code for comparison of current and future climatic conditions with paired Wilcoxon rank sum test, and predicted habitat suitability within the presence mask, and HTML report of script execution with results, including code for preprocessing of monthly temperature rasters
    - Source code as R Markdown for statistical downscaling of monthly temperature rasters, file Downscaling_tmin_01_climond.Rmd
    - Source code as R Markdown for code for comparison of current and future climatic conditions, habitat suitability and Wilxon rank sum test, file M_mercedesiarum_climate_change_with_U.Rmd
    - Result of code execution, file M_mercedesiarum_climate_change_with_U.html
- Data S4: Computer code for production of cross-validated model in 10 iterations, prediction of habitat suitability in current and future conditions, model evaluation, and HTML report of script execution with results
    - Source code for model as self-documented R Markdown, set of files: integrator.Rmd, renderer.Rmd, evaluation_current.Rmd, predictor_current.Rmd, predictor_future_he45_50.Rmd,

    predictor_future_he45_70.Rmd, predictor_future_he85_50.Rmd, predictor_future_he85_70.Rmd
  - Results of code execution as report files in HTML format, set of files: integrator.html, step_29dup_1.html as well as nine similar files for ten cross-validation iterations, evaluation_current.html, predictor_current.html, predictor_future_he45_50.html, predictor_future_he45_70.html, predictor_future_he85_50.html, predictor_future_he85_70.html
- Data S5: Vector polygon files for estimated habitat suitability in current and future conditions, considering majority criteria and ESS threshold individually for each iteration,
  - Habitat suitability on current conditions, ESRI SHP file polygon_climond2_mayority_select.shp
  - Habitat suitability on current conditions within 500 km buffer from known presence, ESRI SHP file polygon_climond2_mayority_select_buffer500km.shp
  - Habitat suitability on future conditions RCP45 at 2050, ESRI SHP file polygon_he45_50_mayority_select.shp
  - Habitat suitability on future conditions RCP45 at 2070, ESRI SHP file polygon_he45_70_mayority_select.shp
  - Habitat suitability on future conditions RCP85 at 2050, ESRI SHP file polygon_he85_50_mayority_select.shp
  - Habitat suitability on future conditions RCP85 at 2070, ESRI SHP file polygon_he85_70_mayority_select.shp
