## Supplementary figures and images for "Vulnerability to climate change for narrowly ranged species: the case of Ecuadorian endemic *Magnolia mercedesiarum*"

### elevation2-1.png

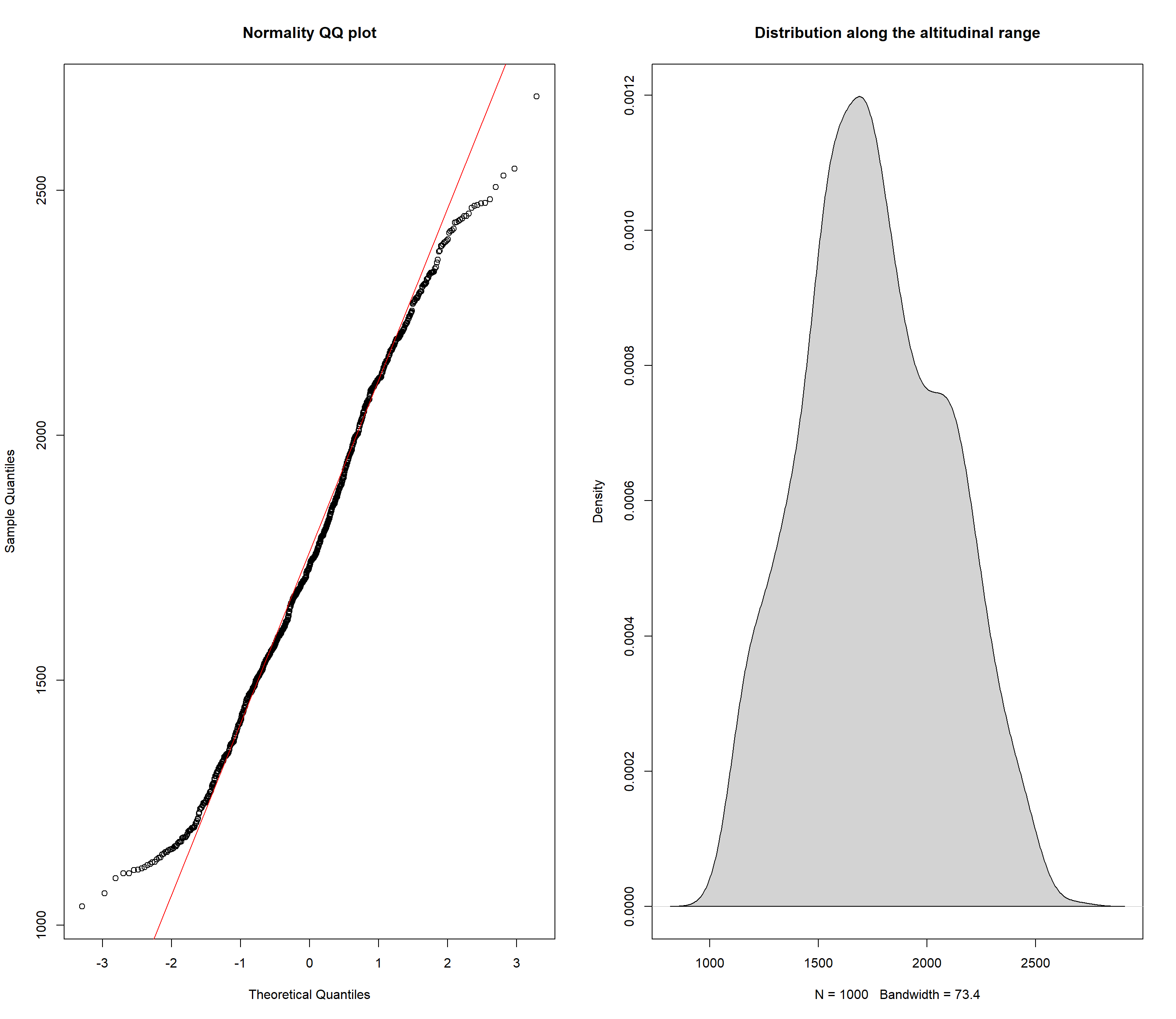

### elevation-1.png

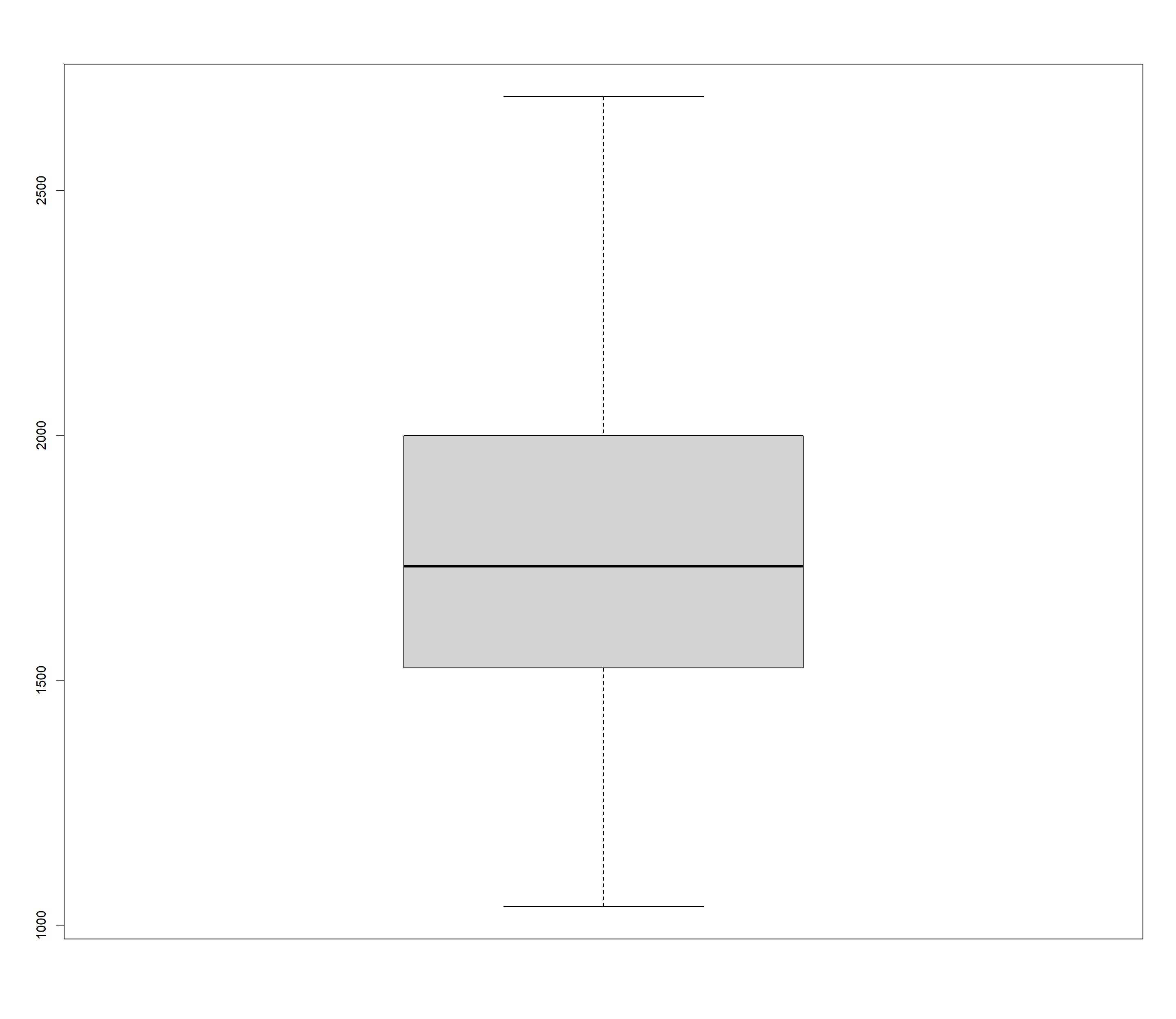

### evaluate-model-1.png

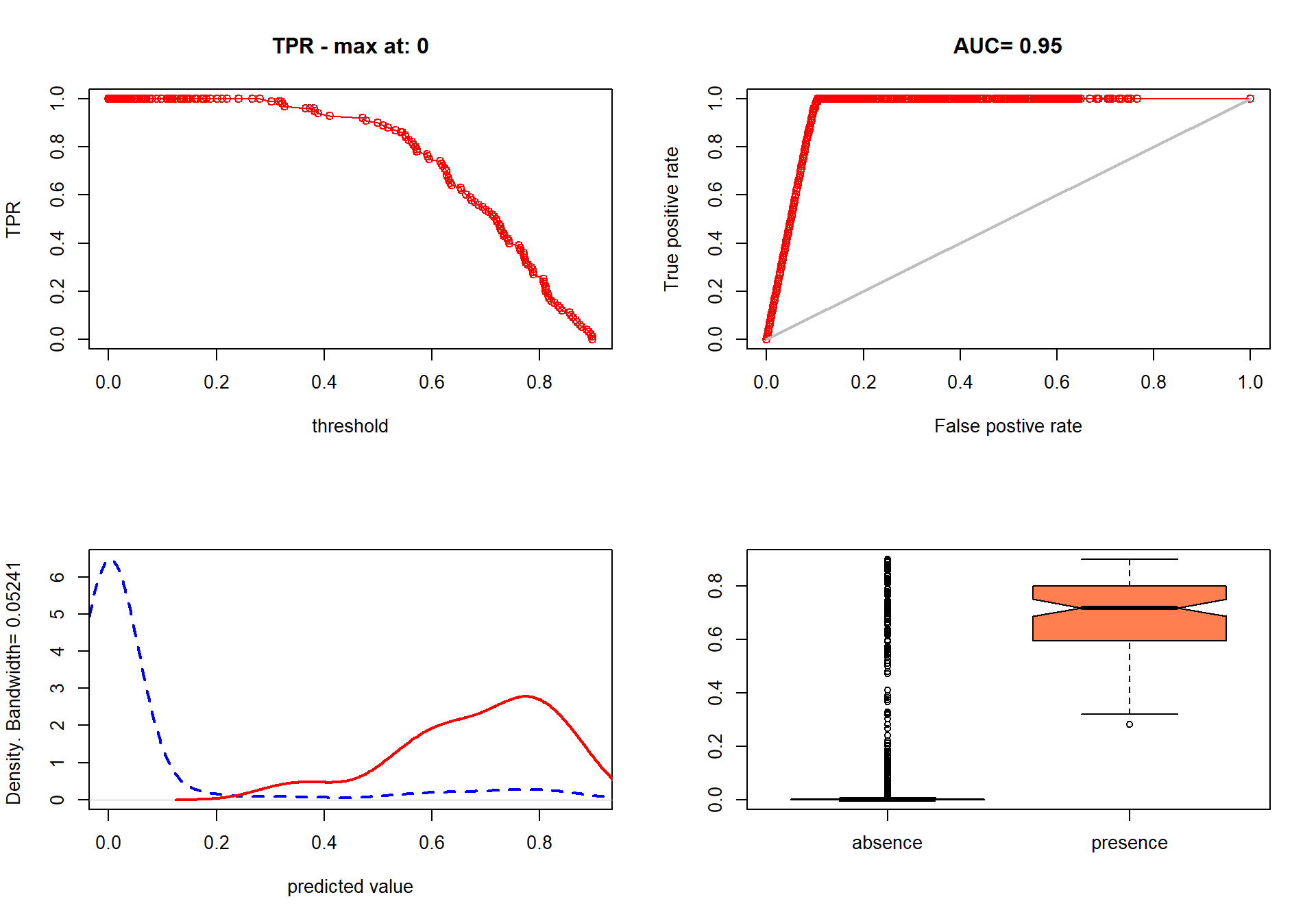

### evaluate-model-2.png

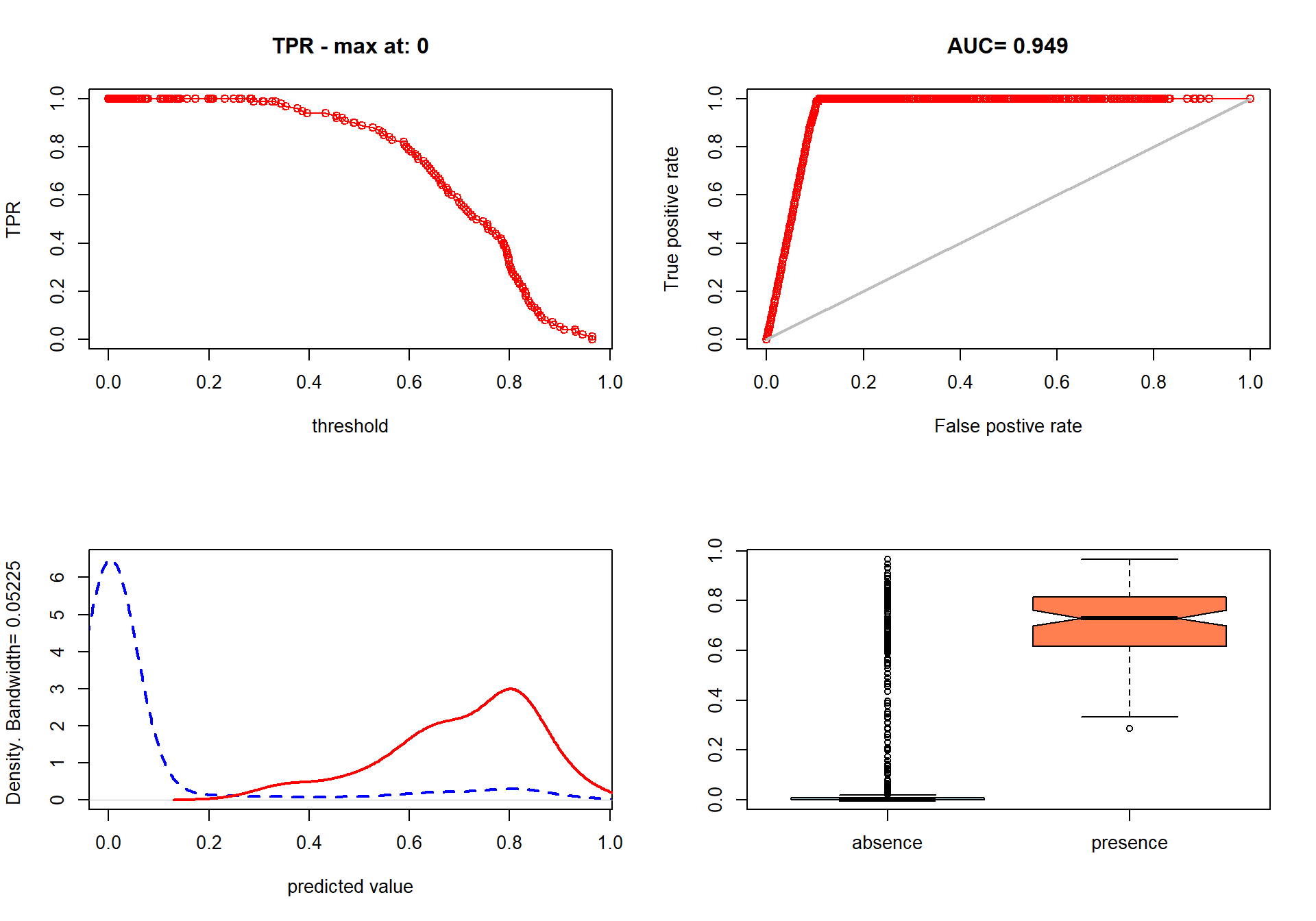

### evaluate-model-3.png

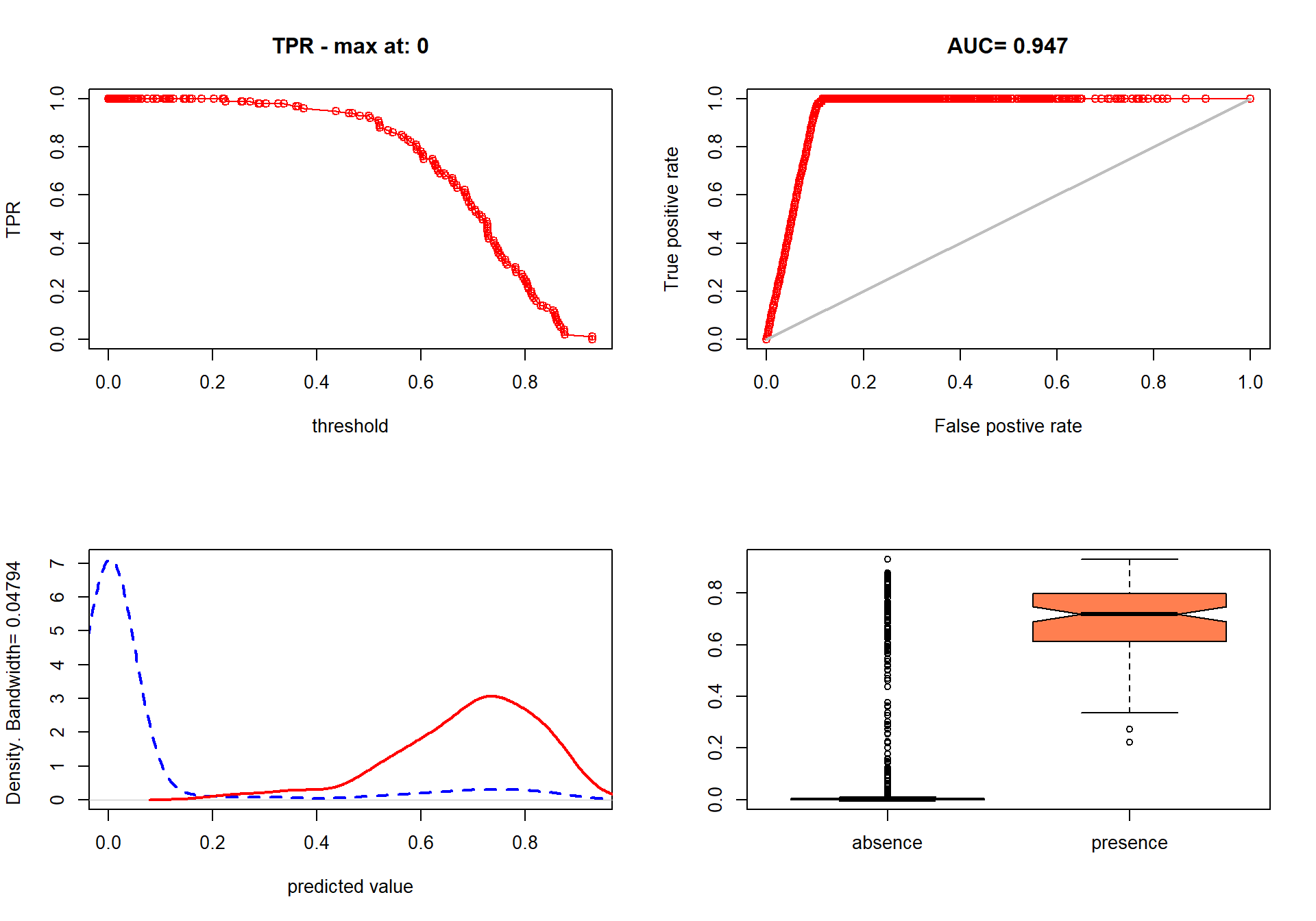

### evaluate-model-4.png

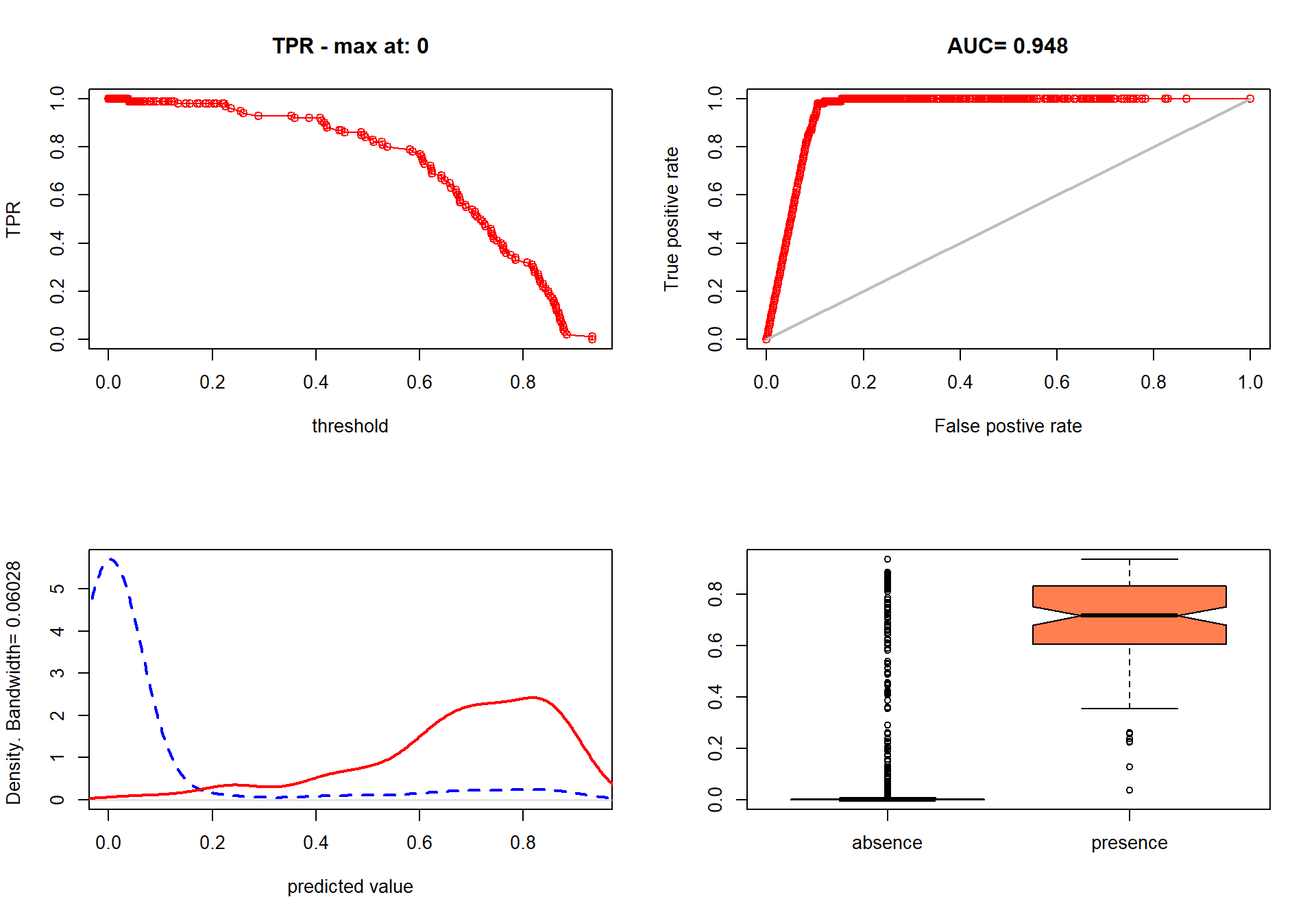

### evaluate-model-5.png

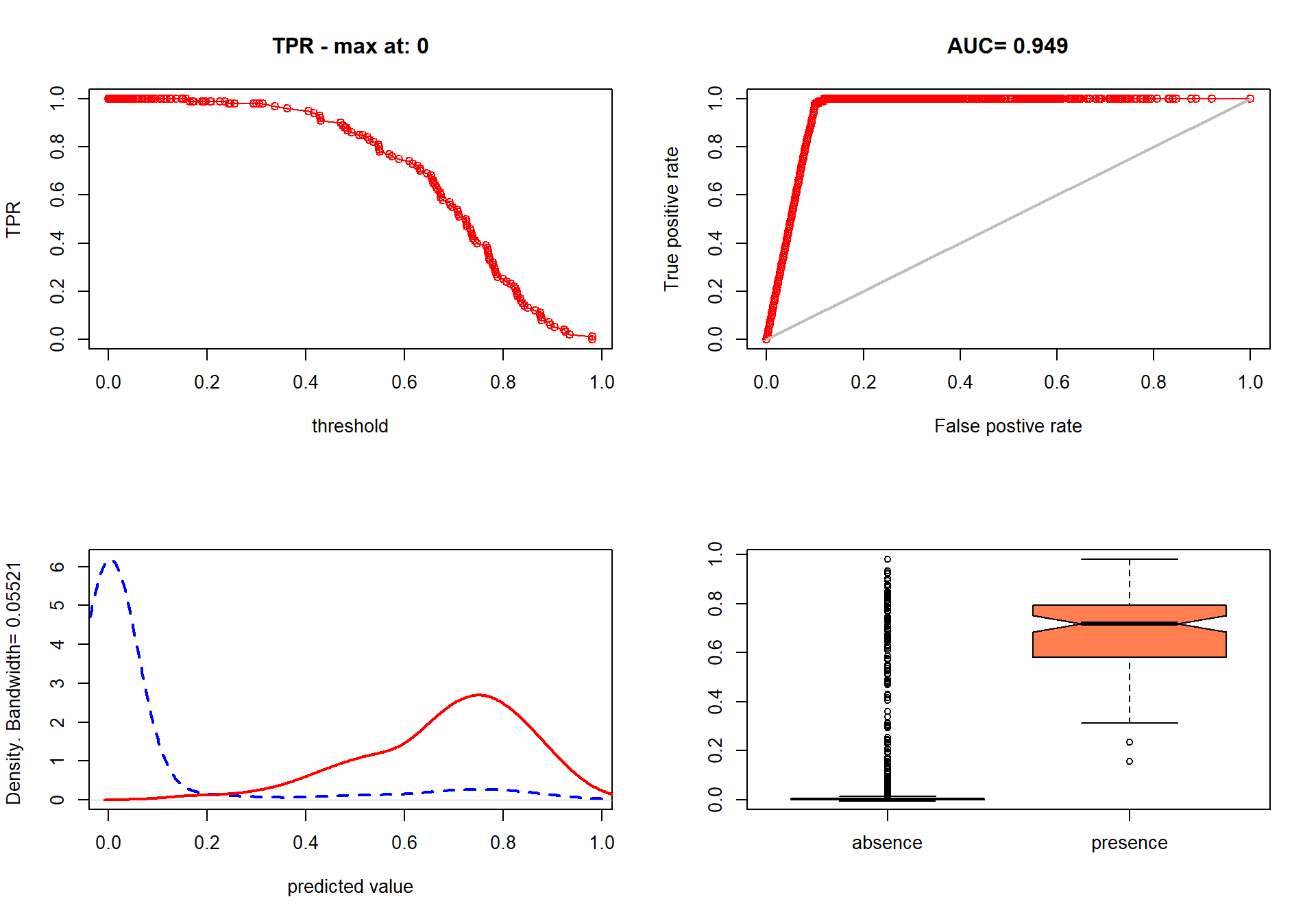

### evaluate-model-6.png

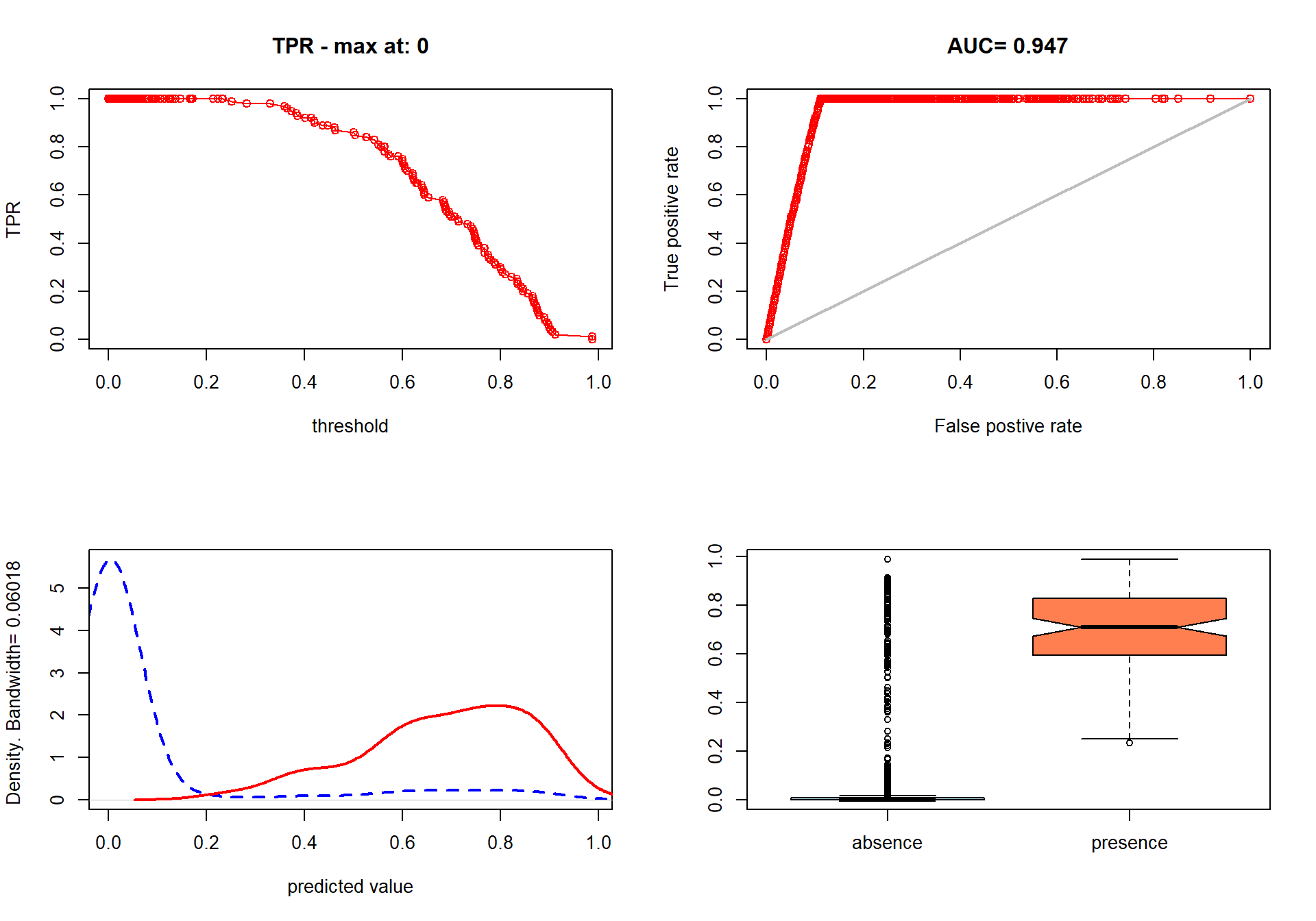

### evaluate-model-7.png

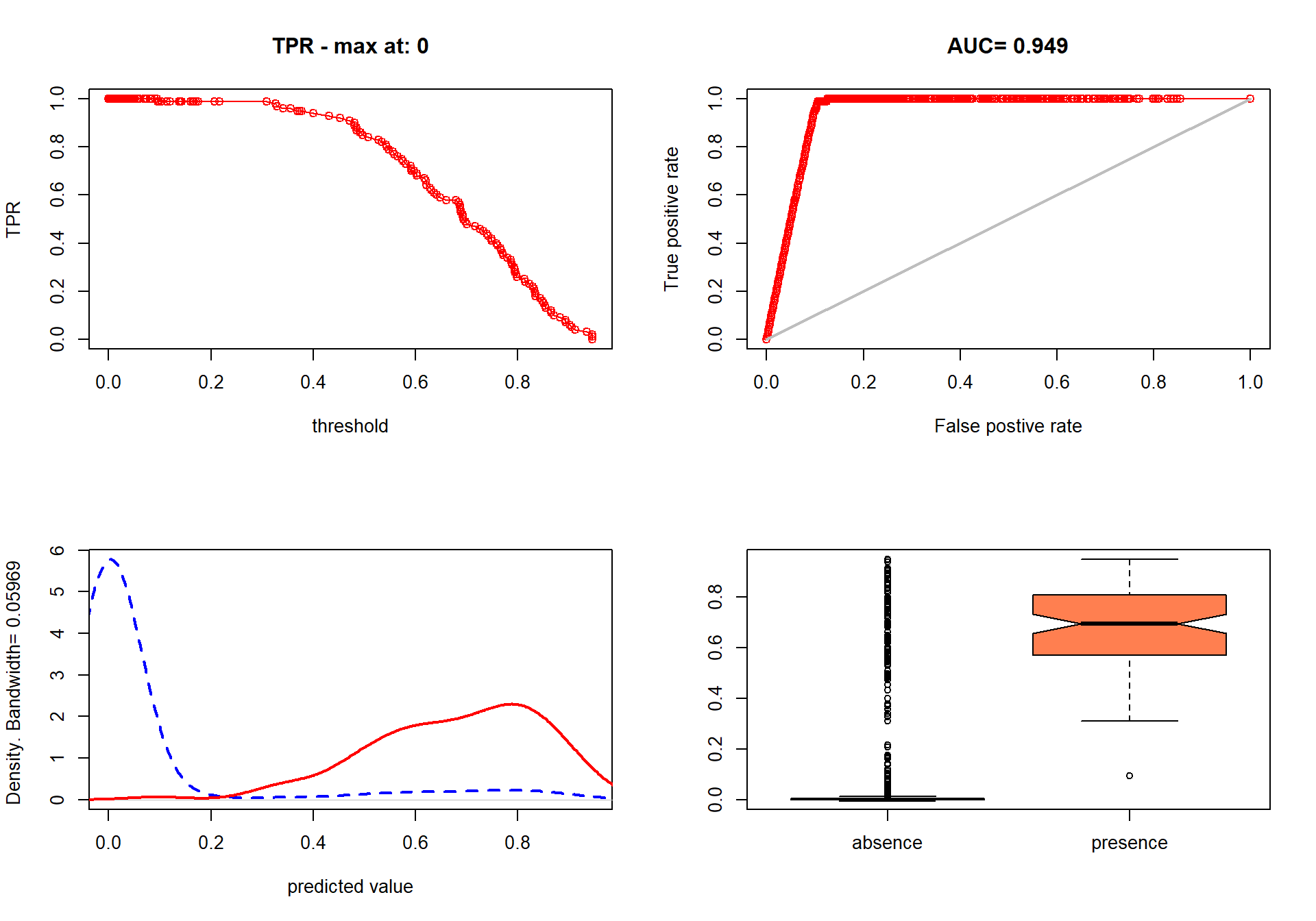

### evaluate-model-8.png

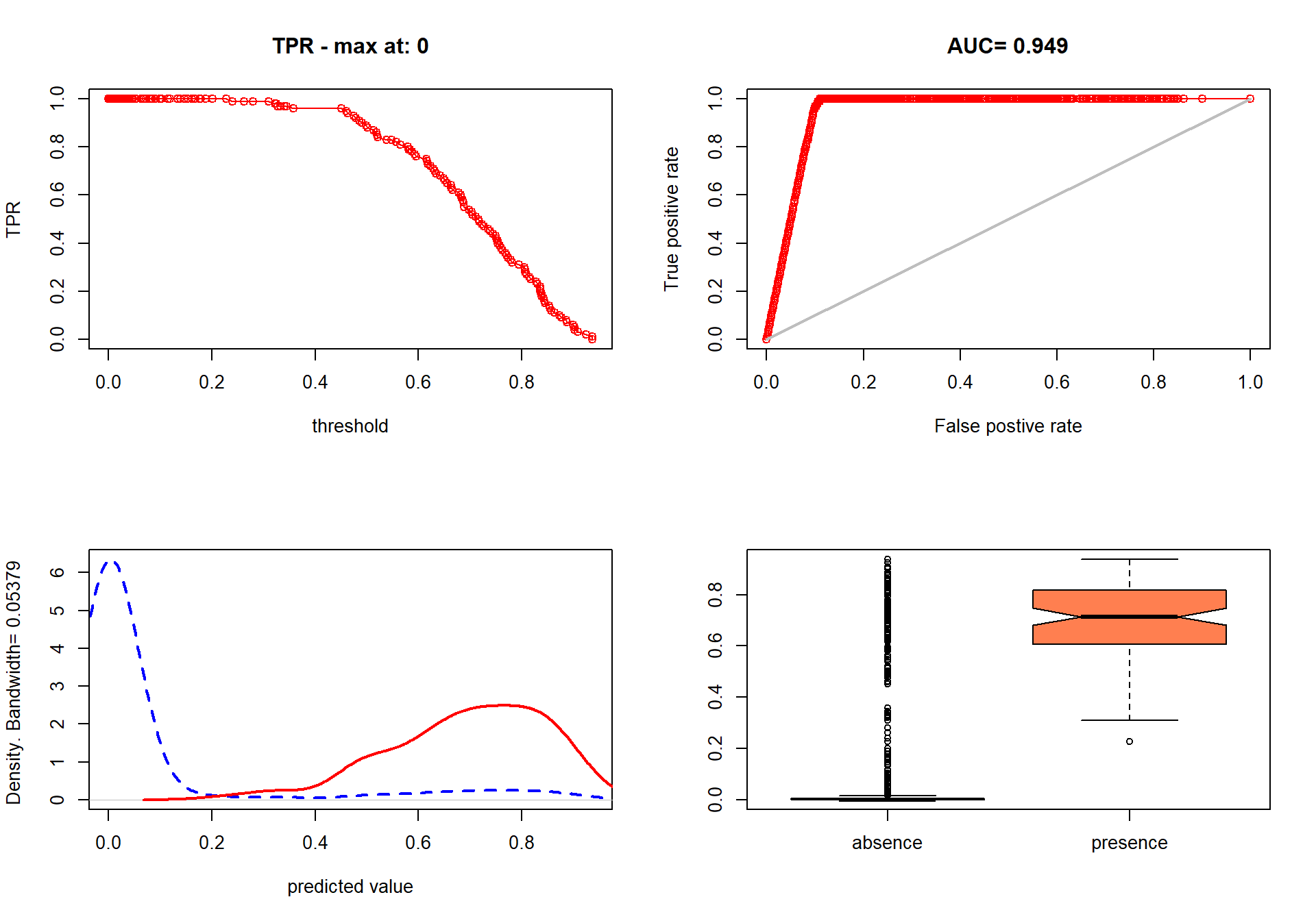

### evaluate-model-9.png

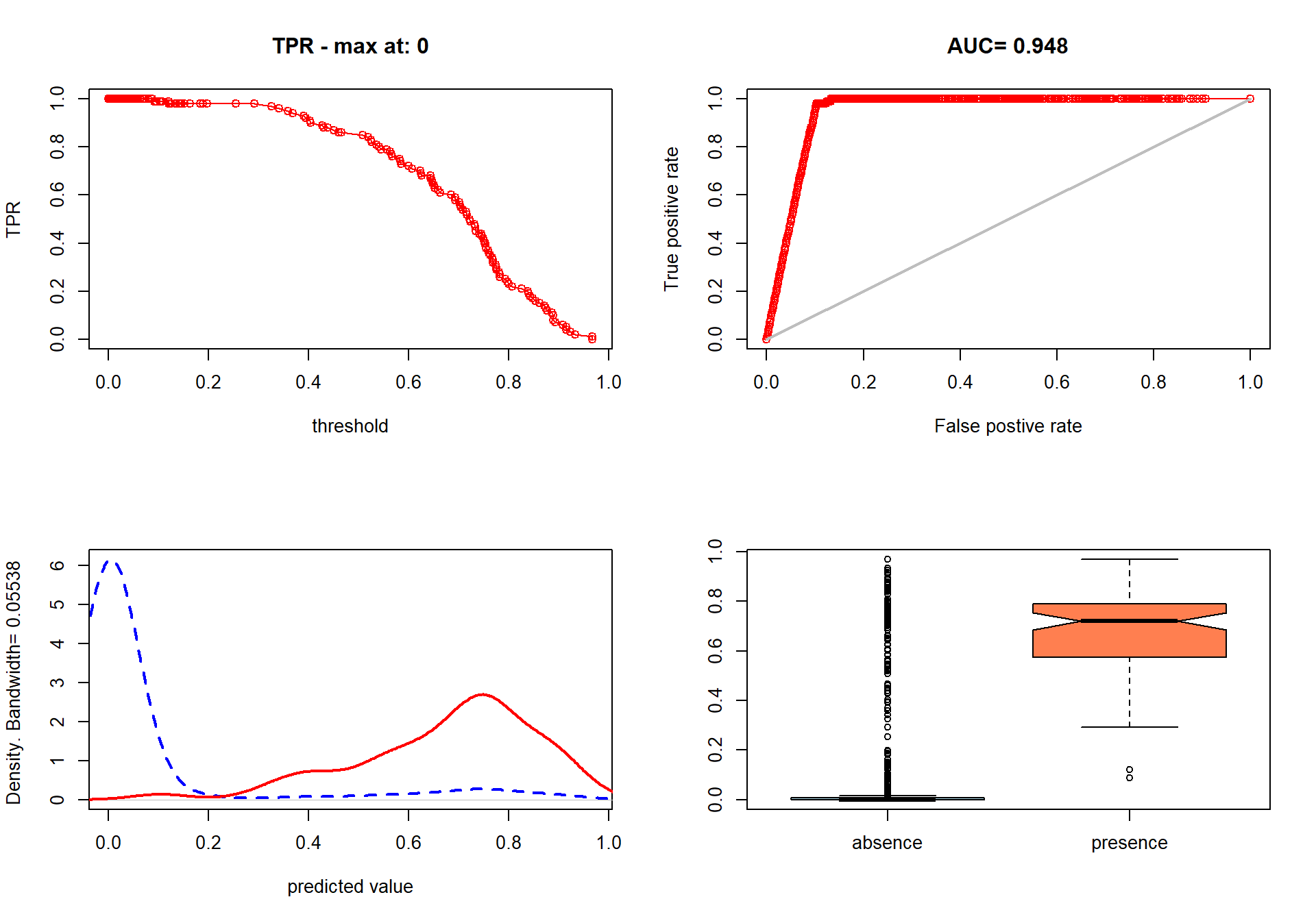

### evaluate-model-10.png

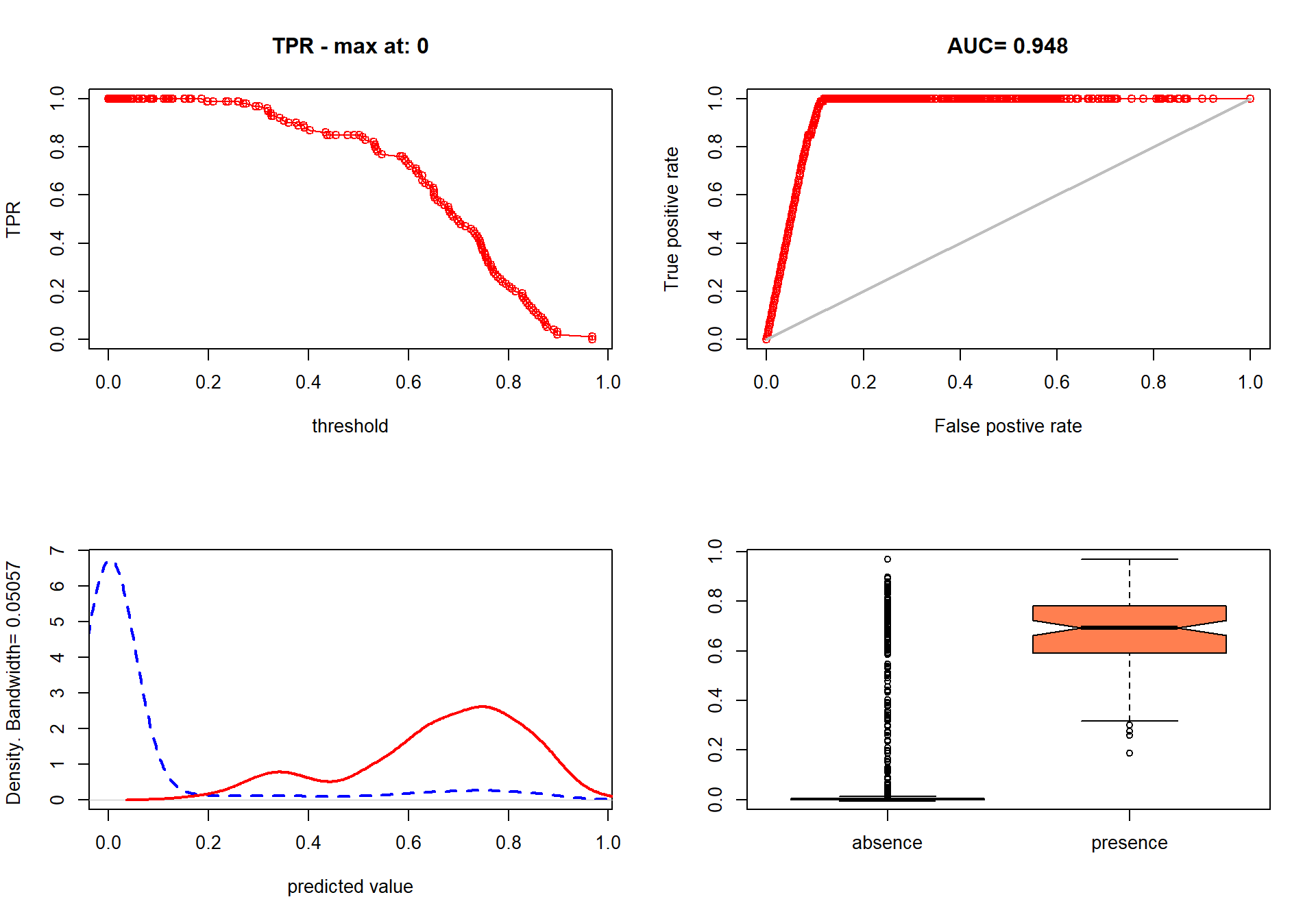

### load-tgs-raster-1.png

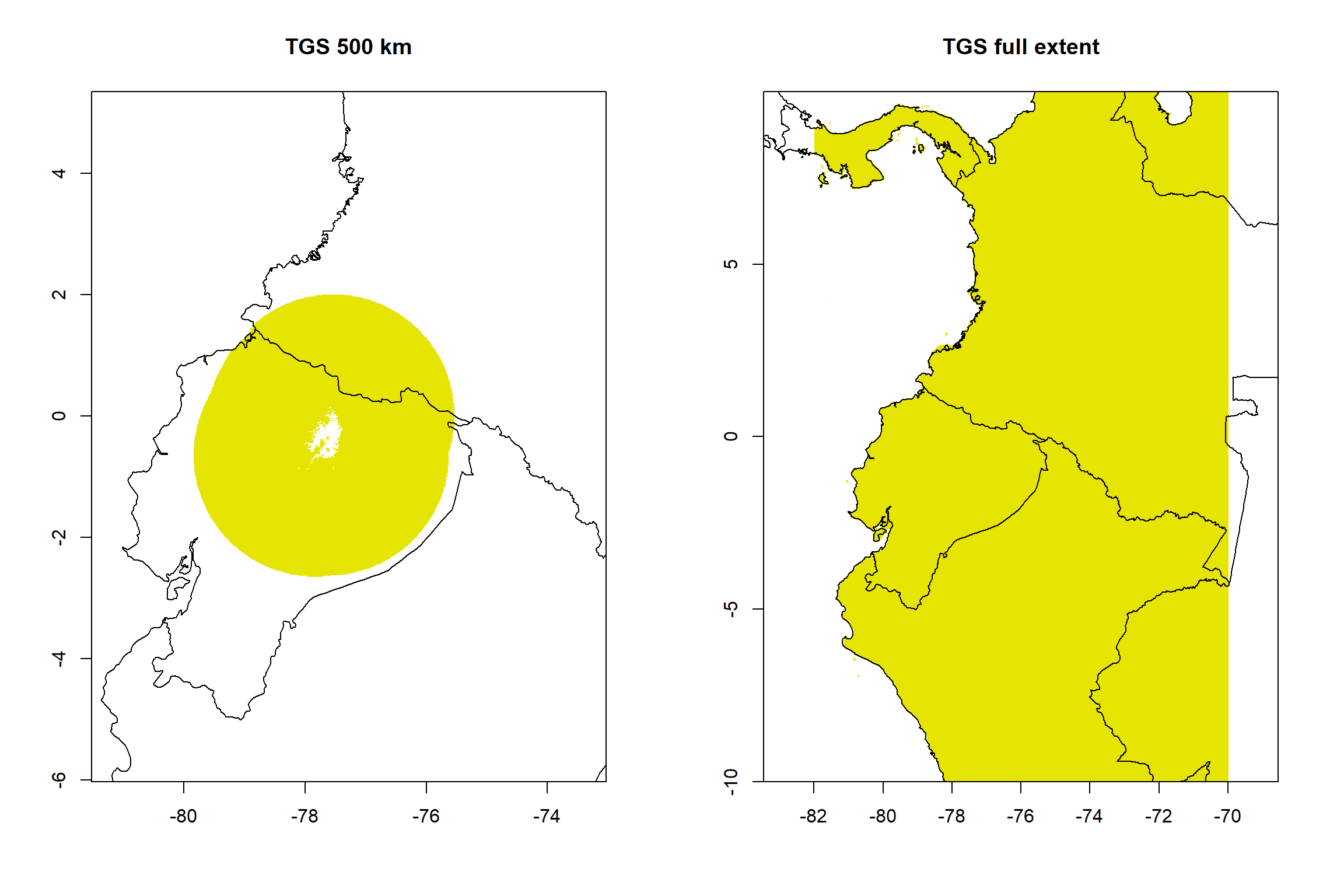

### M_mercedesiarum_logistic_raster_clip.tif

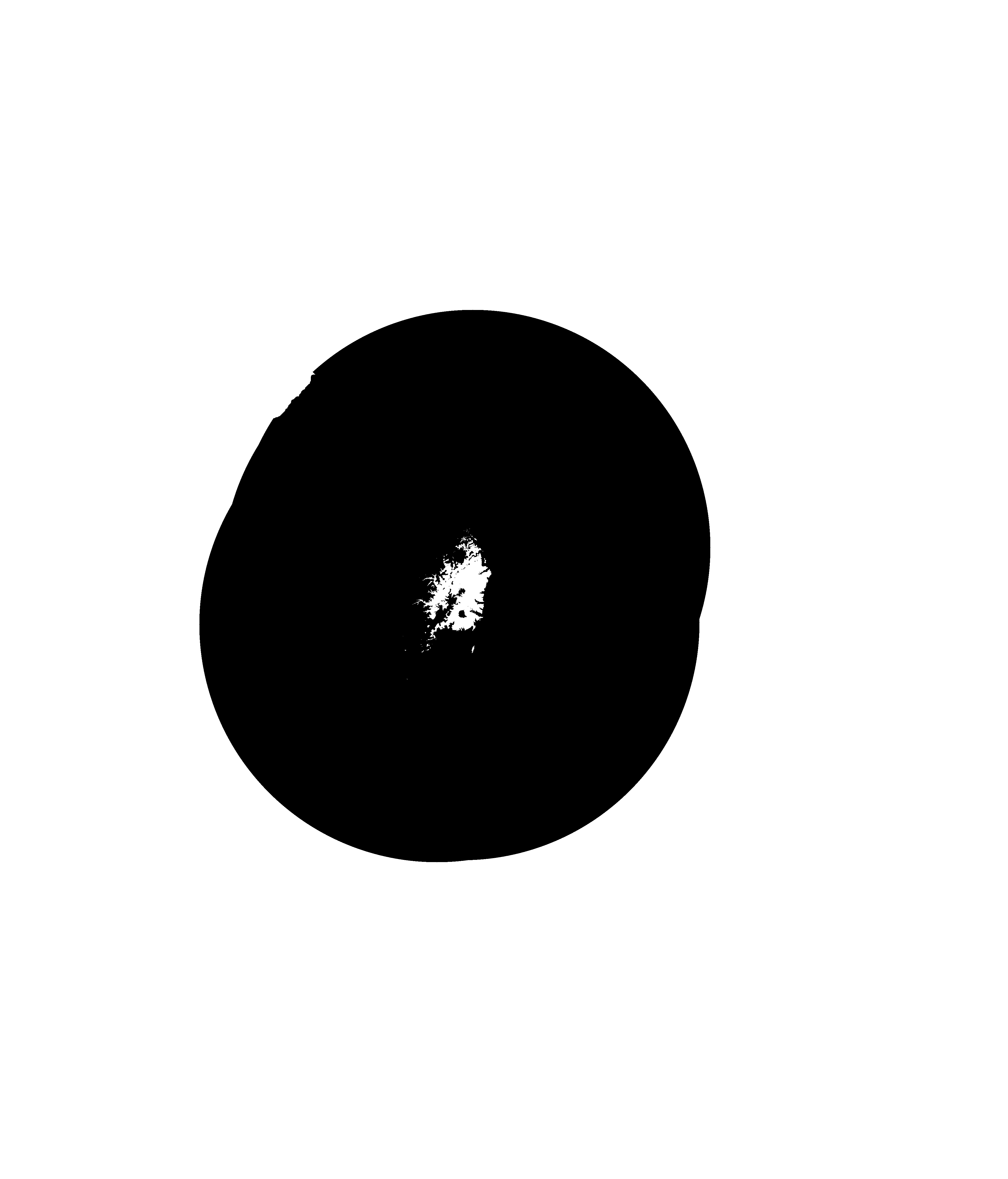

### points-visualization-1.png

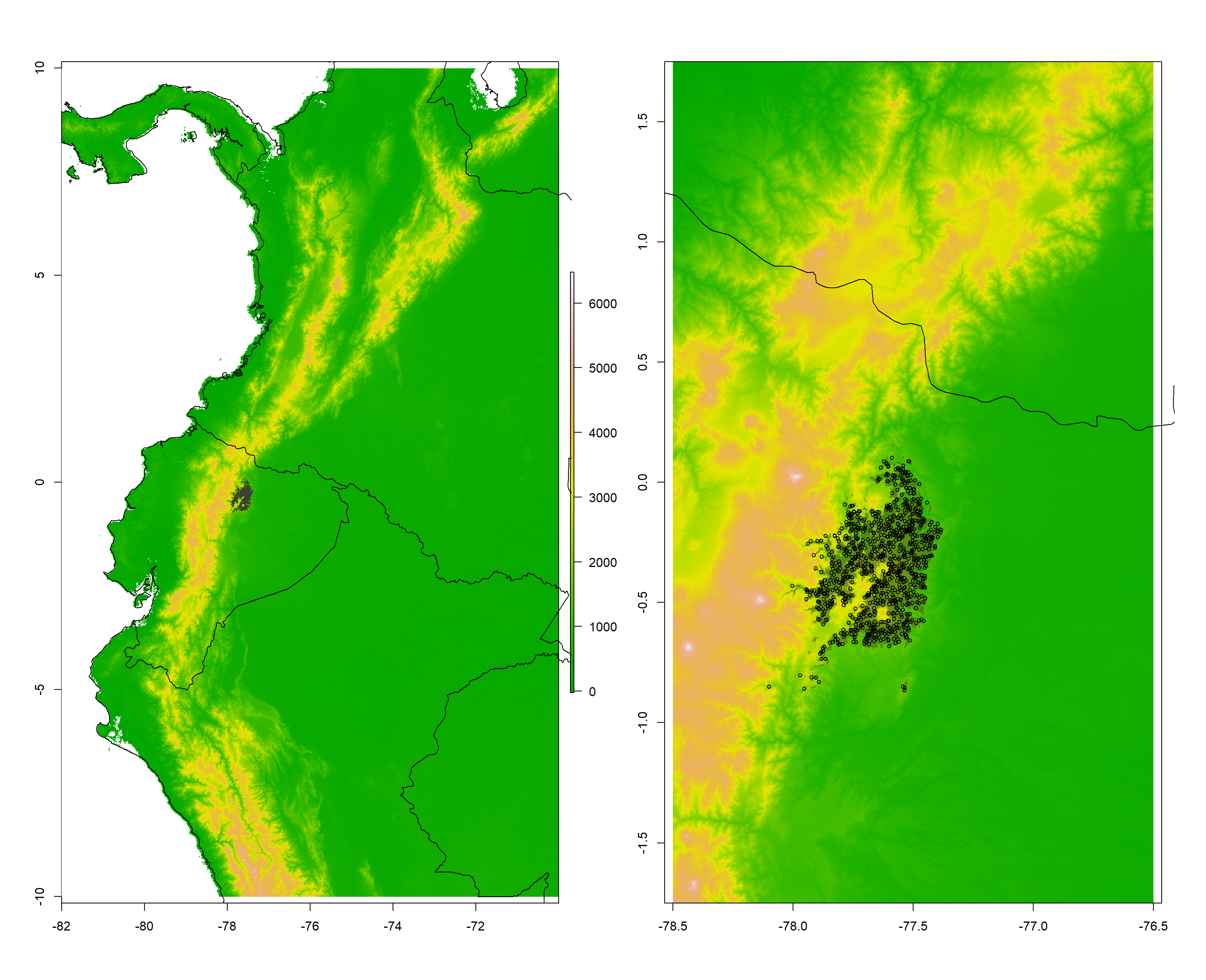

### review-model-1.png

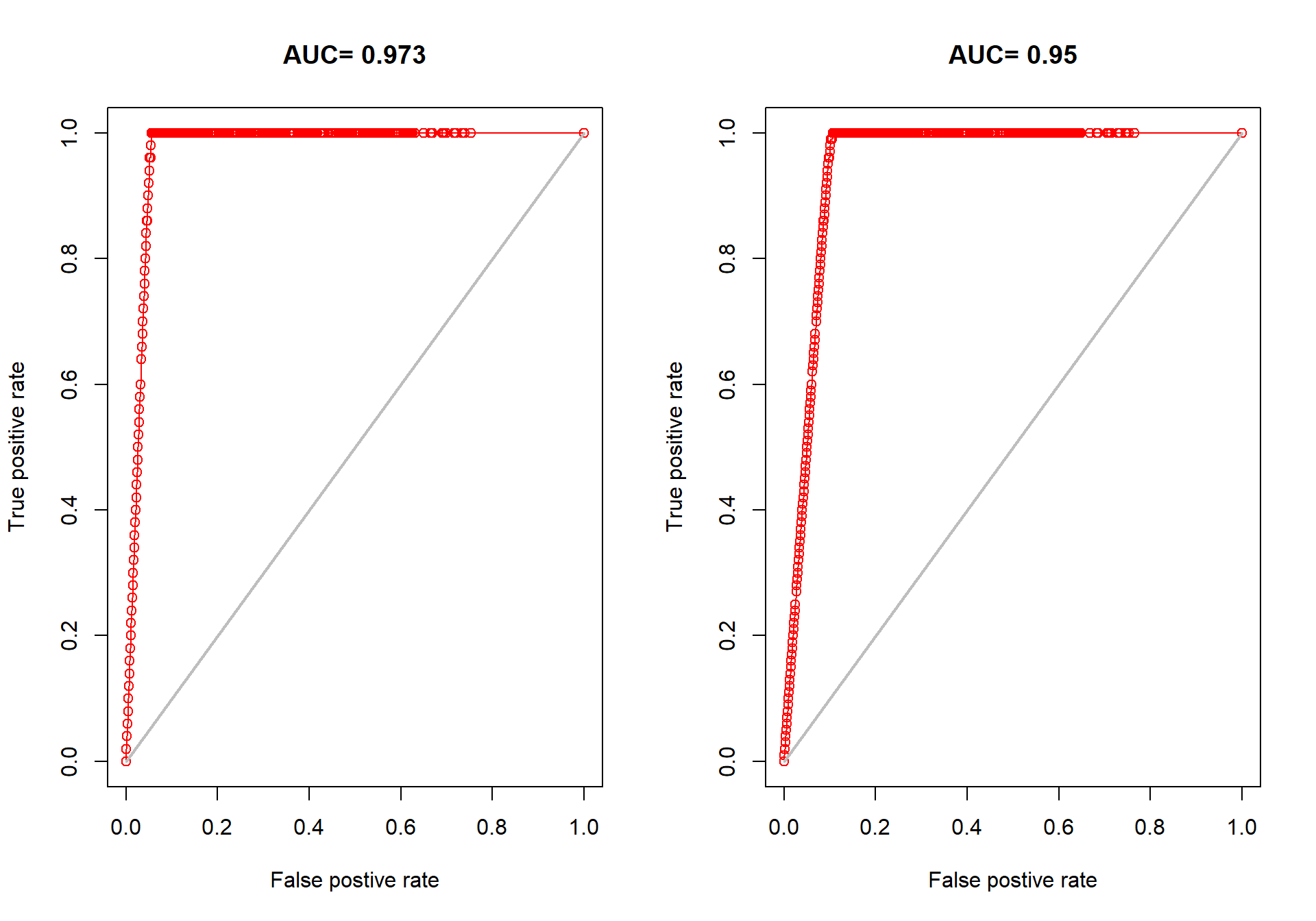

### review-model-2.png

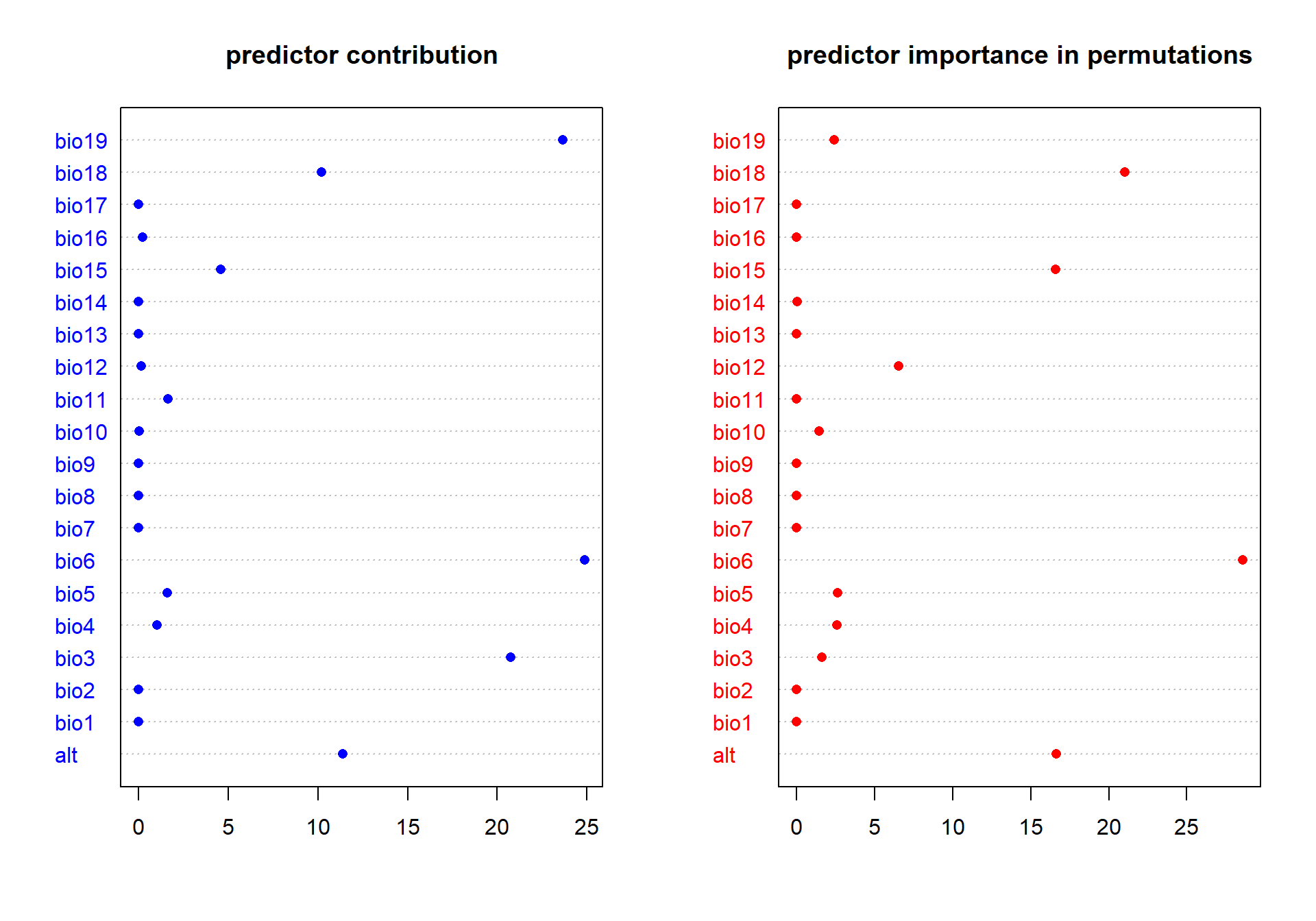

### review-model-3.png

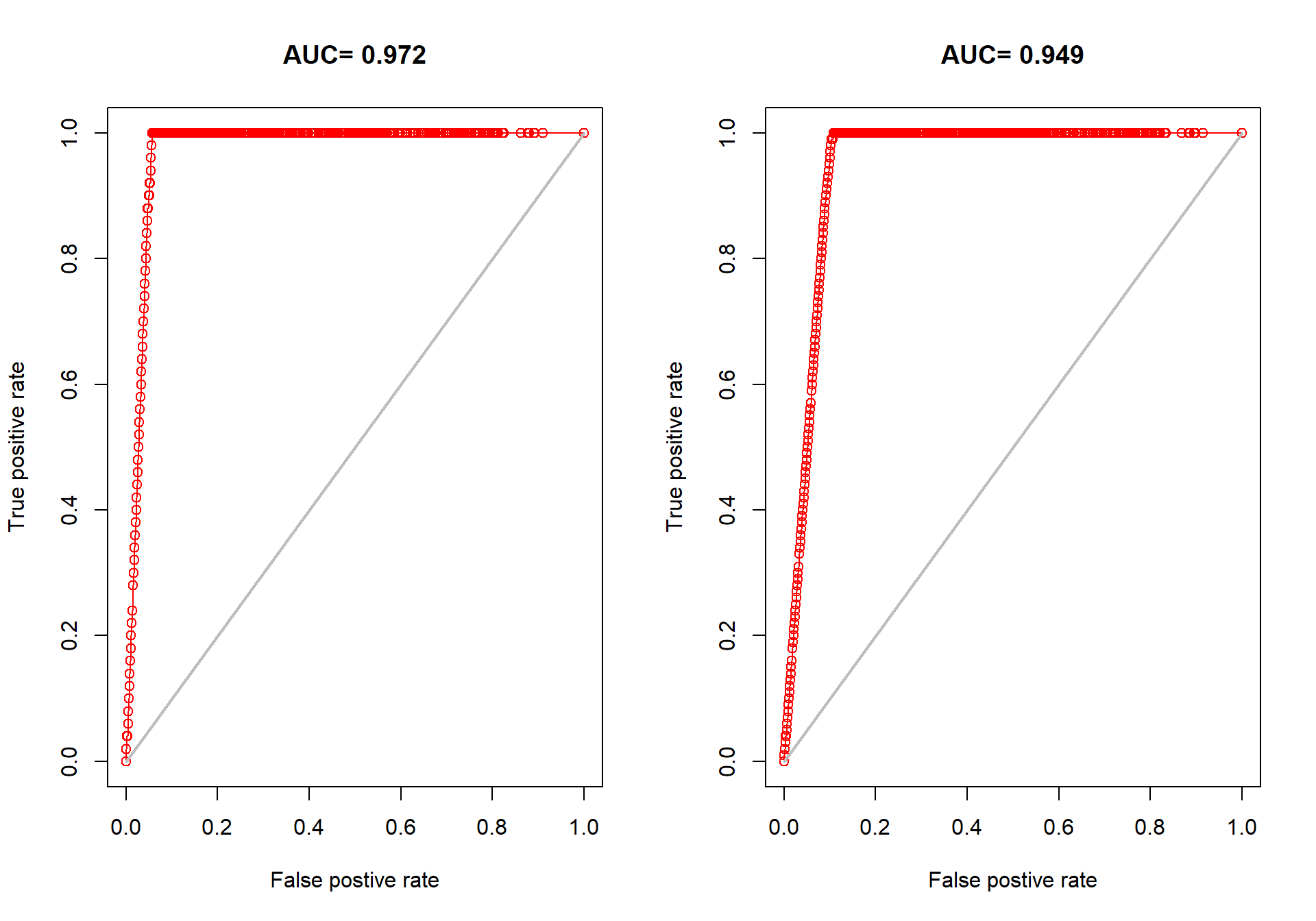

### review-model-4.png

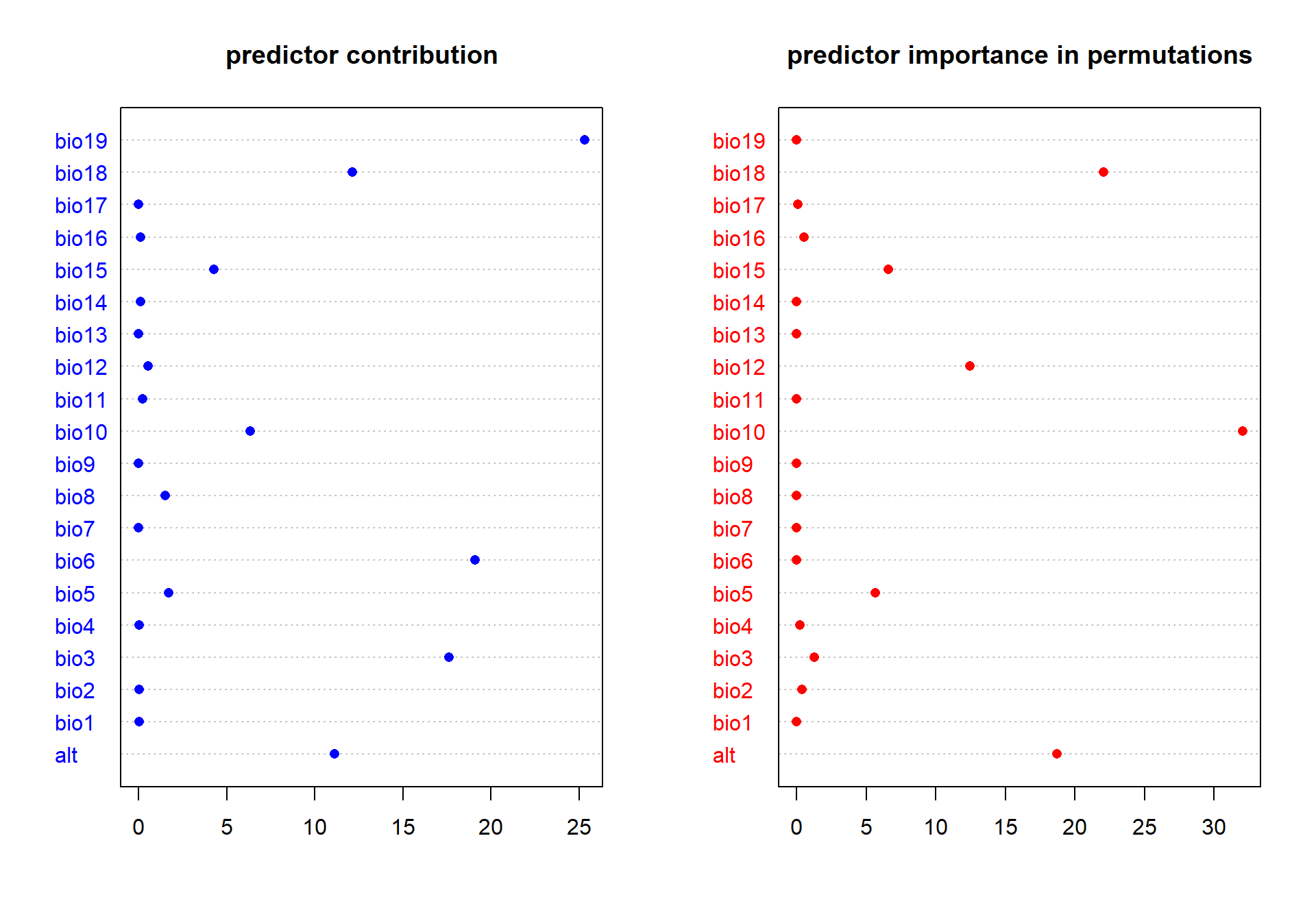

### review-model-5.png

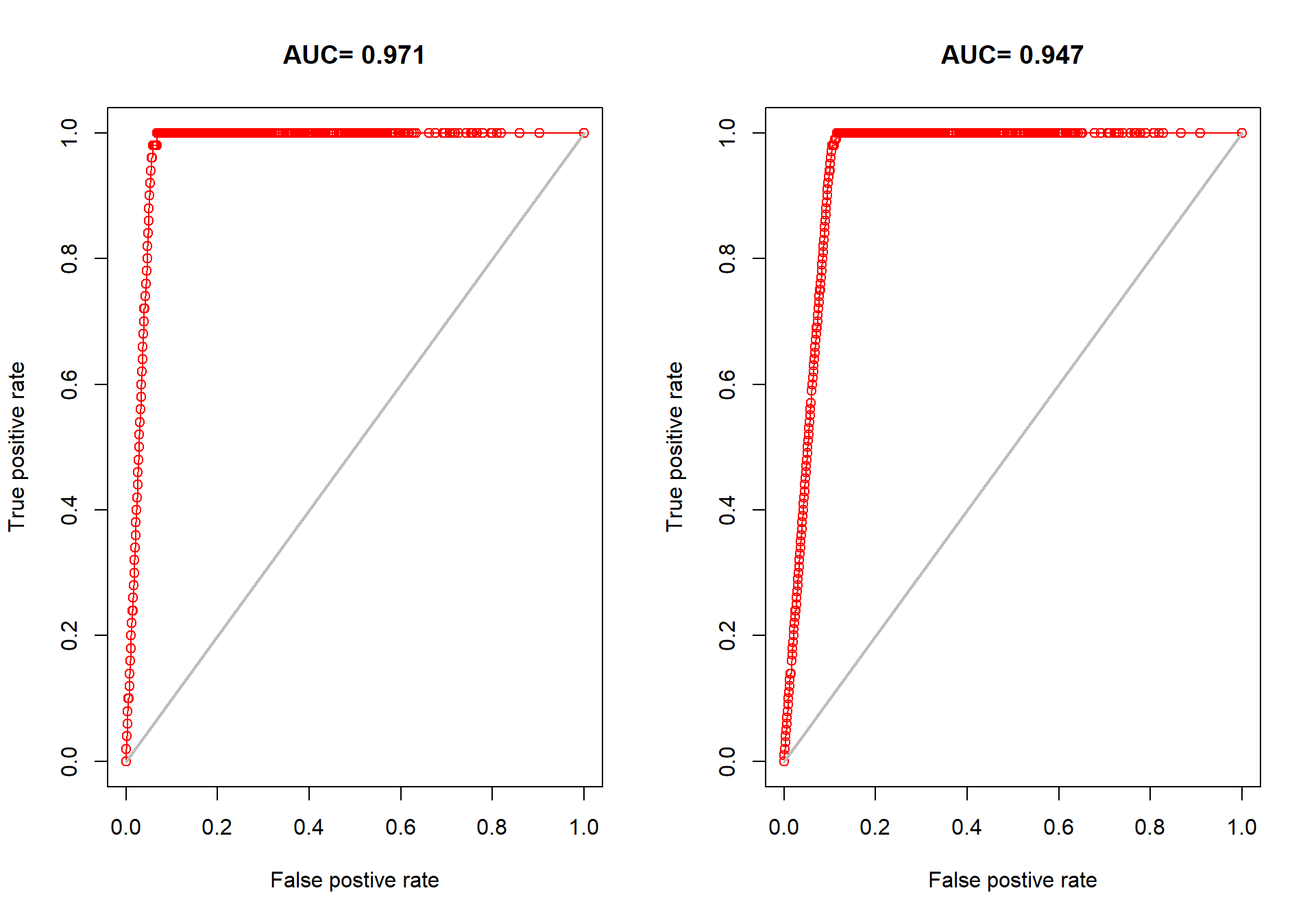

### review-model-6.png

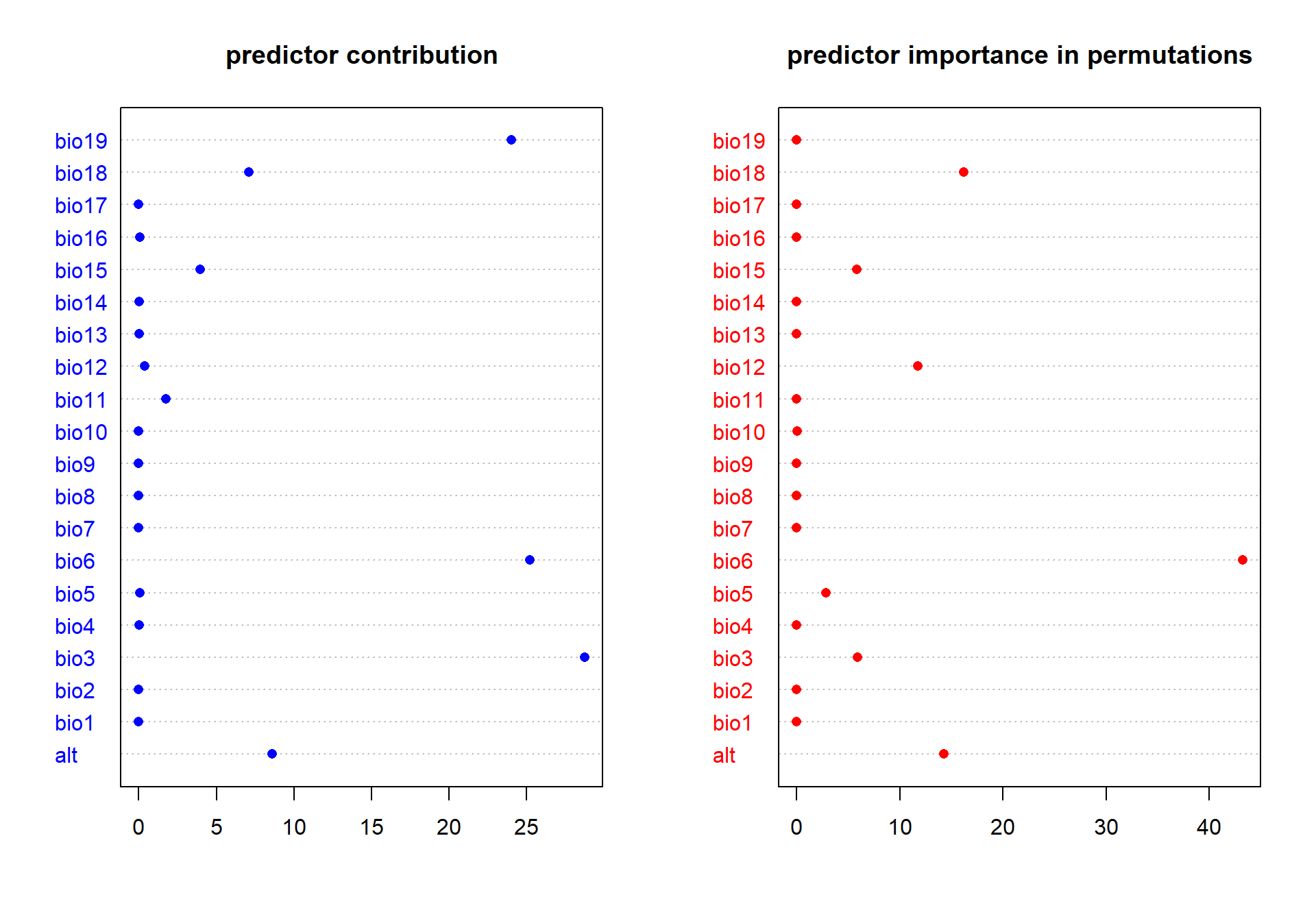

### review-model-7.png

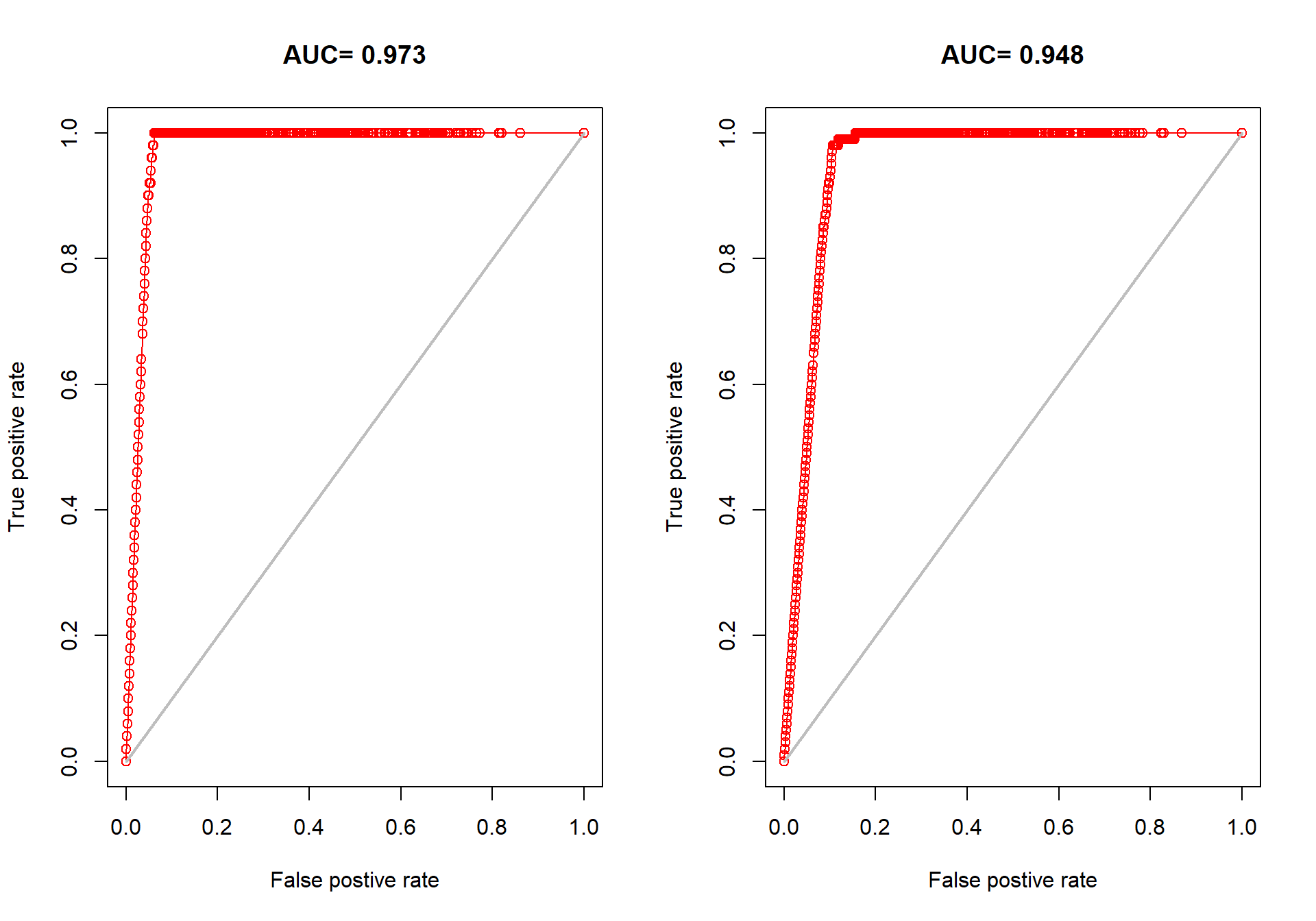

### review-model-8.png

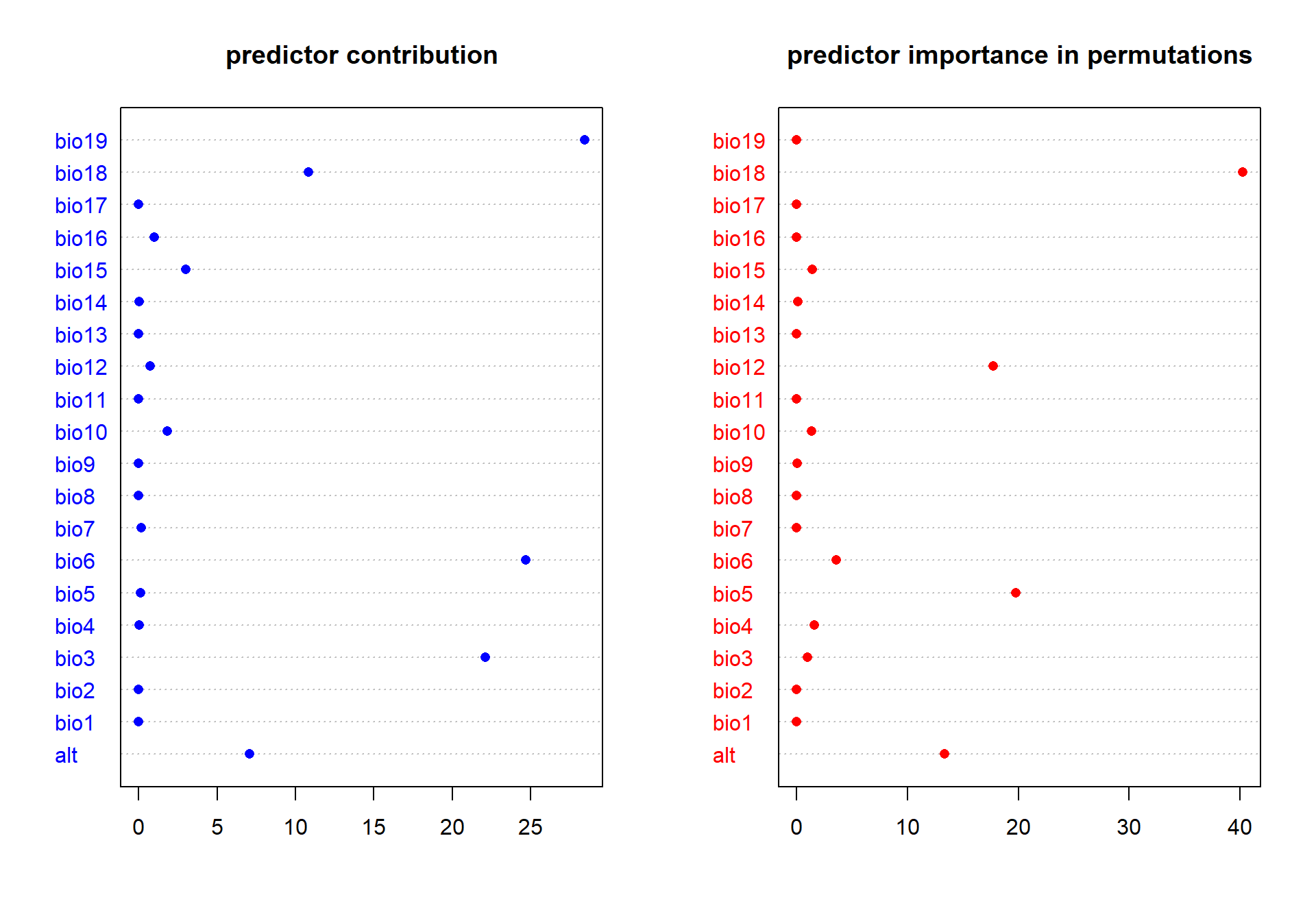

### review-model-9.png

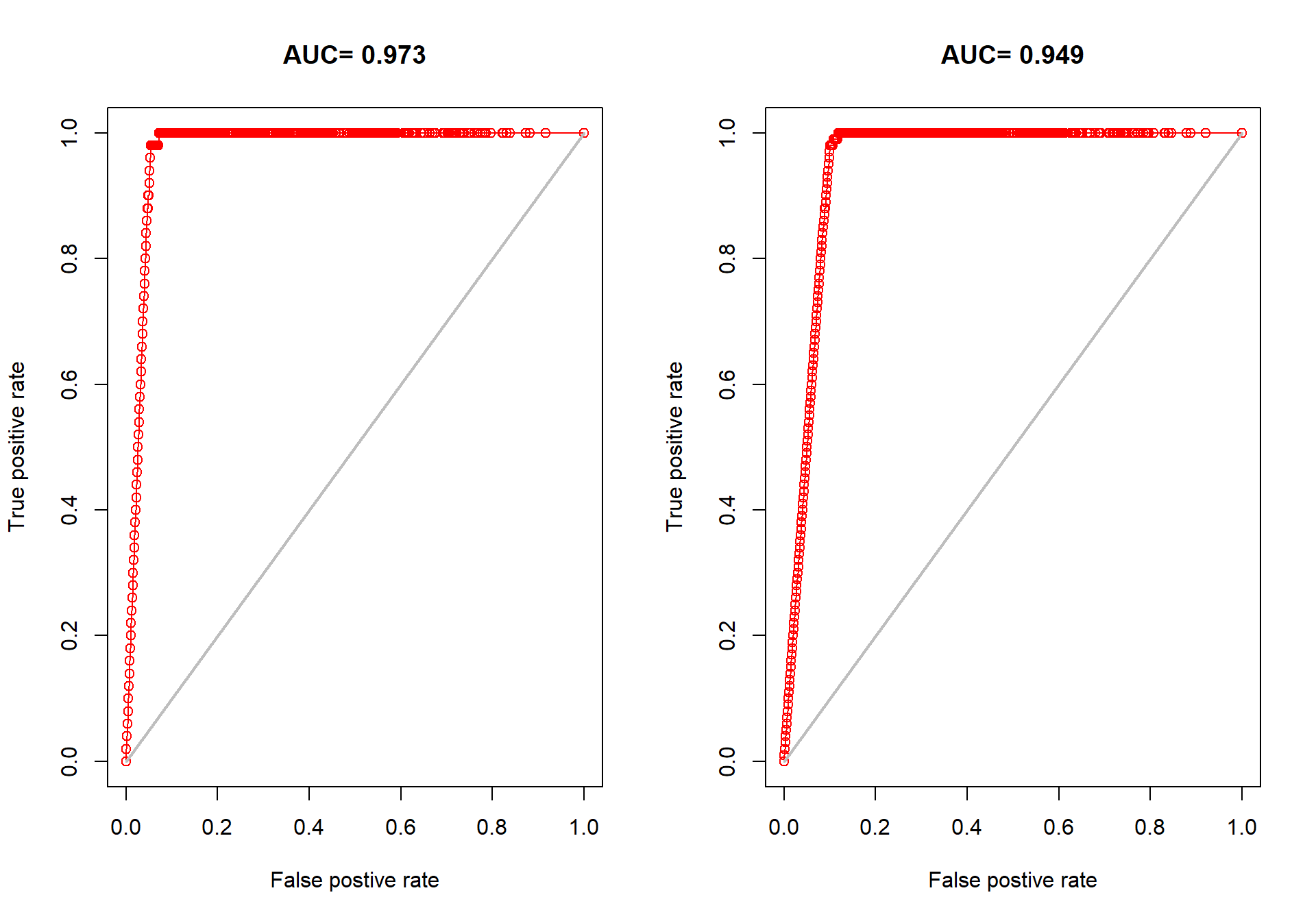

### review-model-10.png

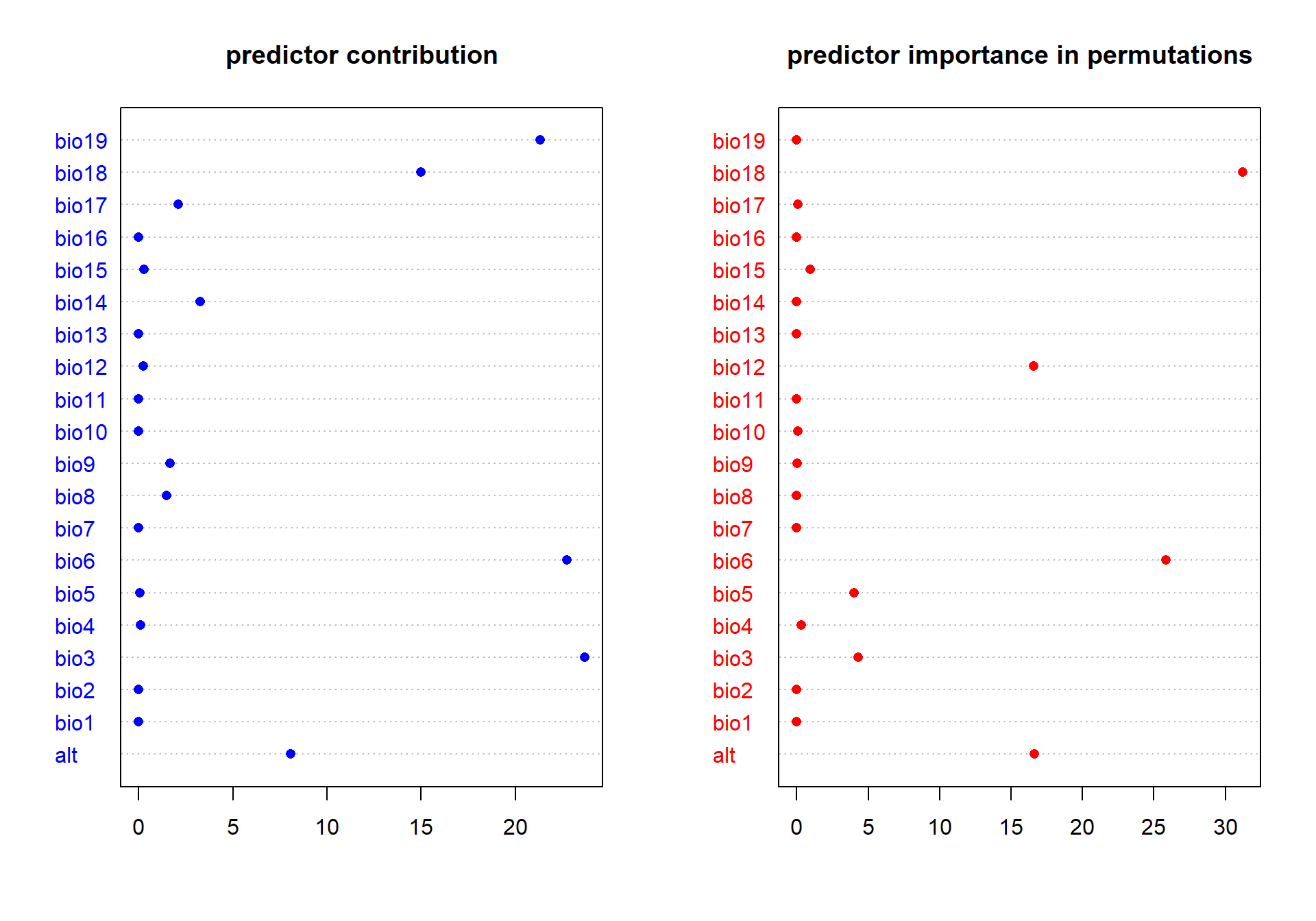

### review-model-11.png

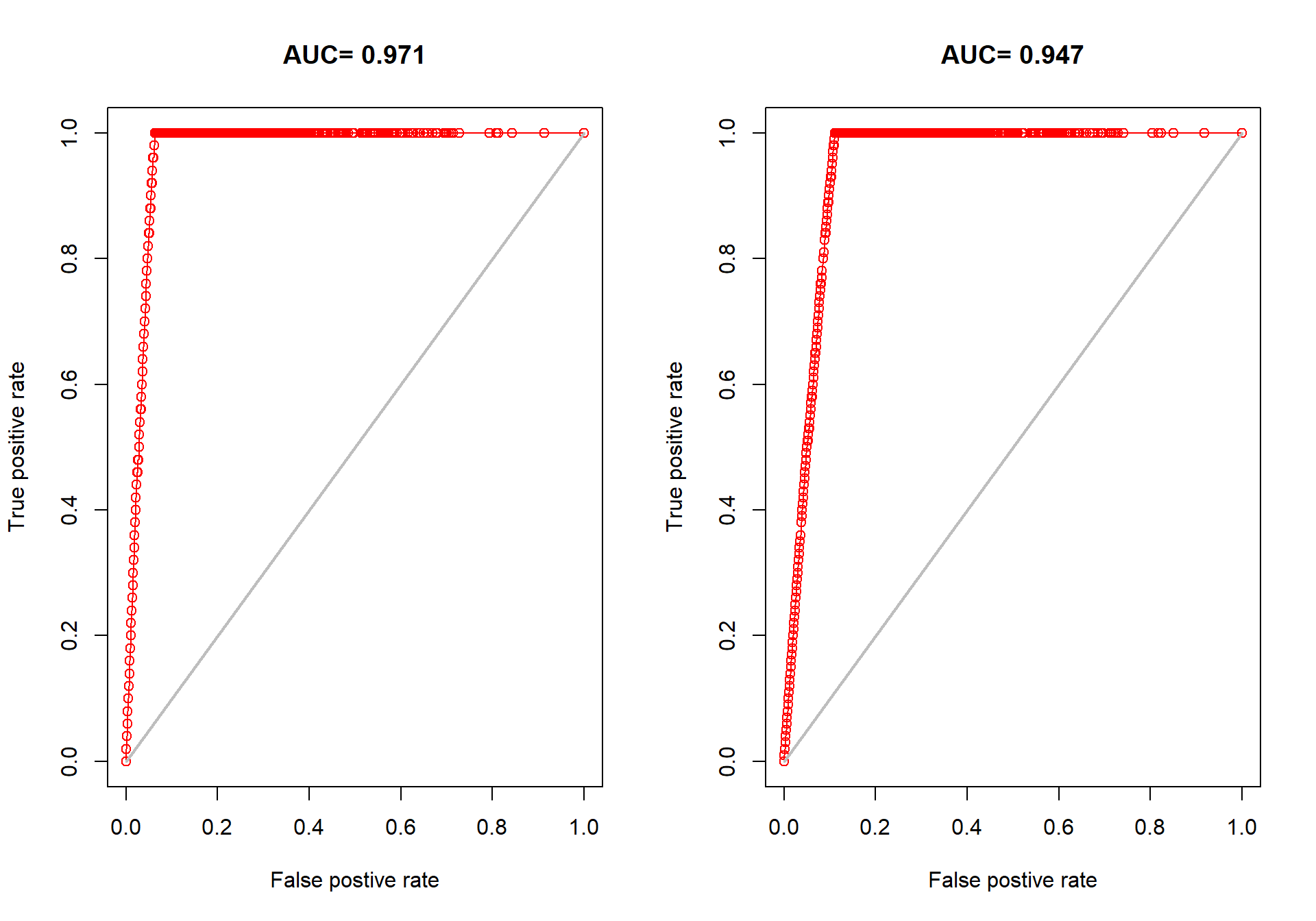

### review-model-12.png

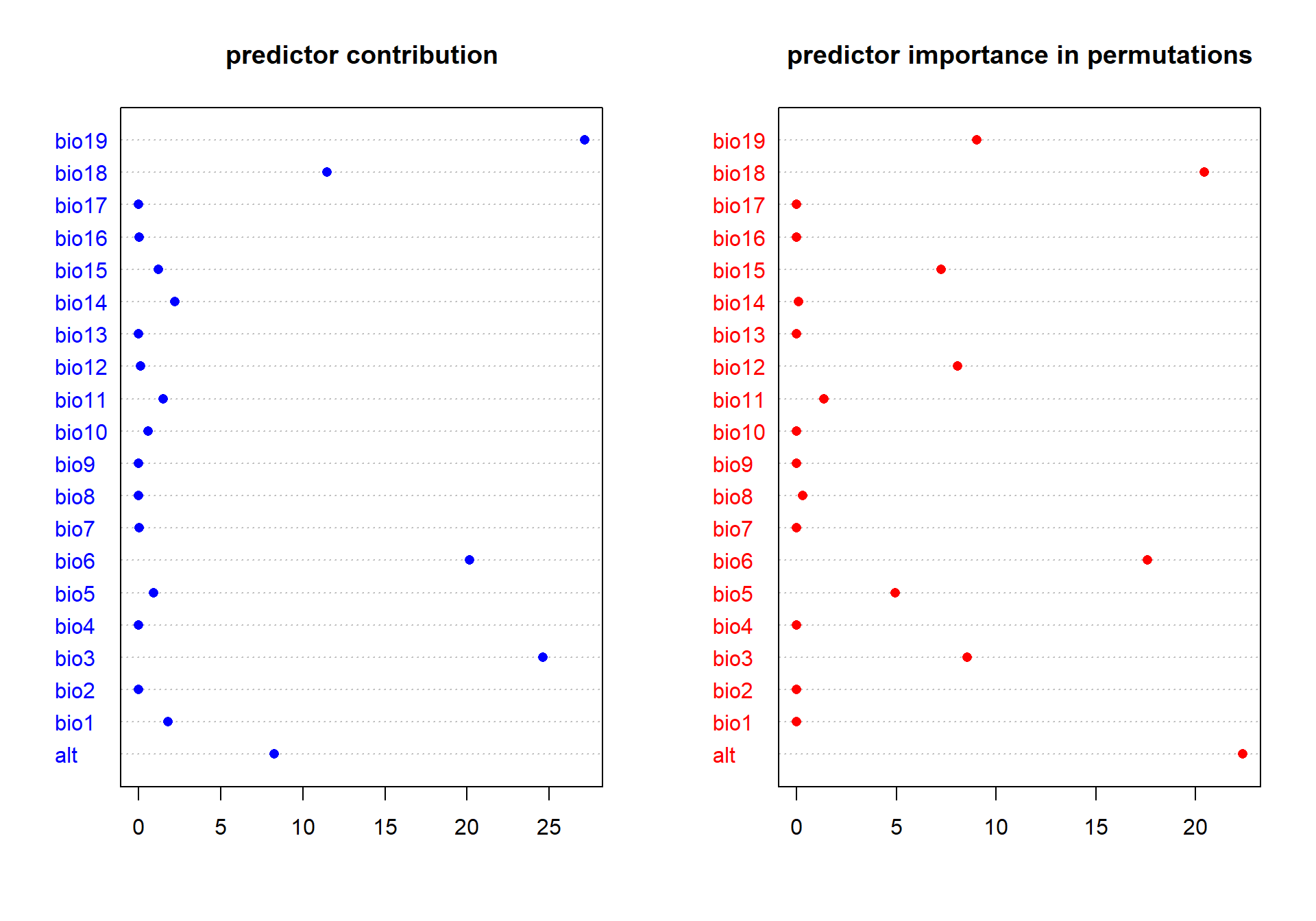

### review-model-13.png

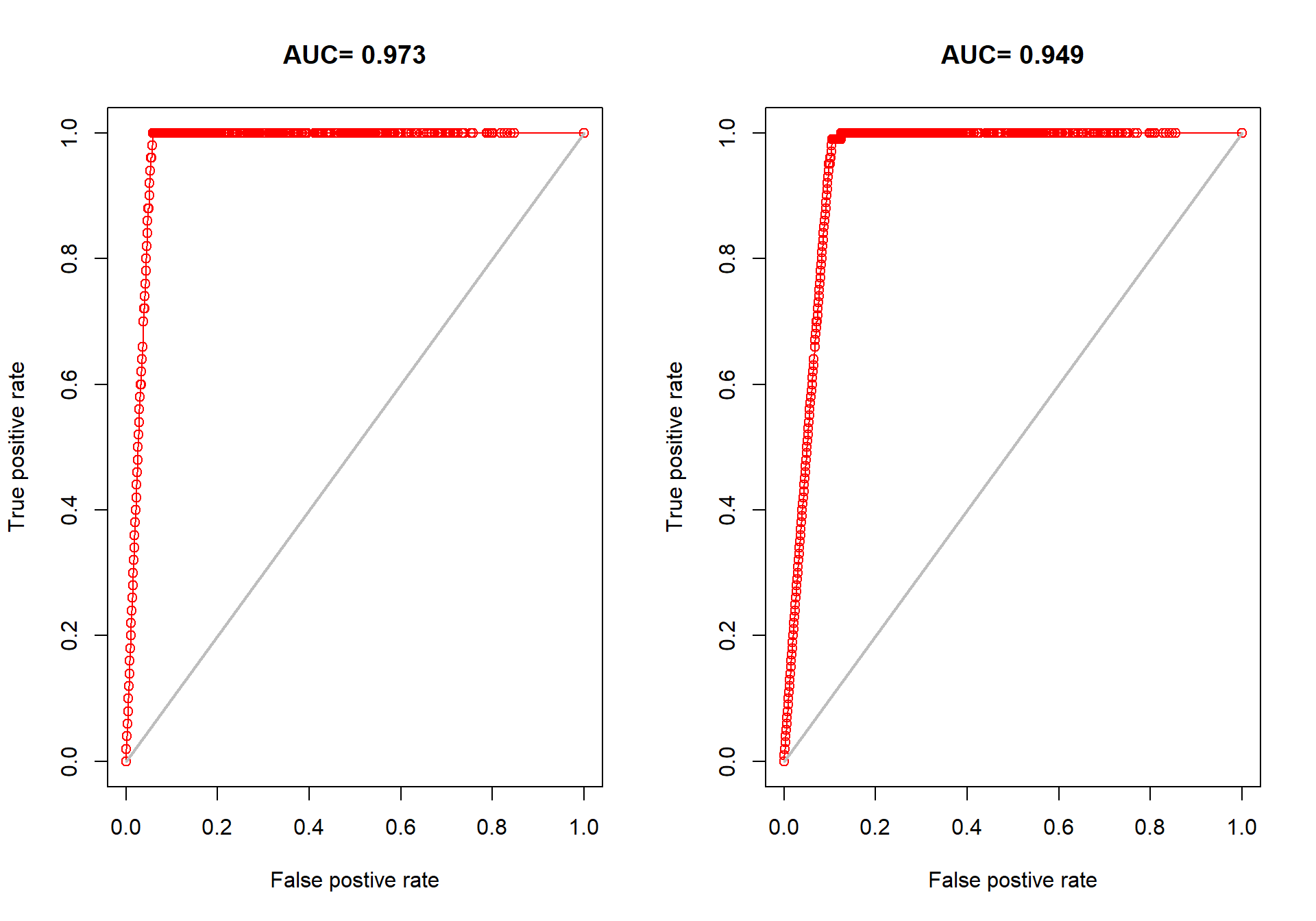

### review-model-14.png

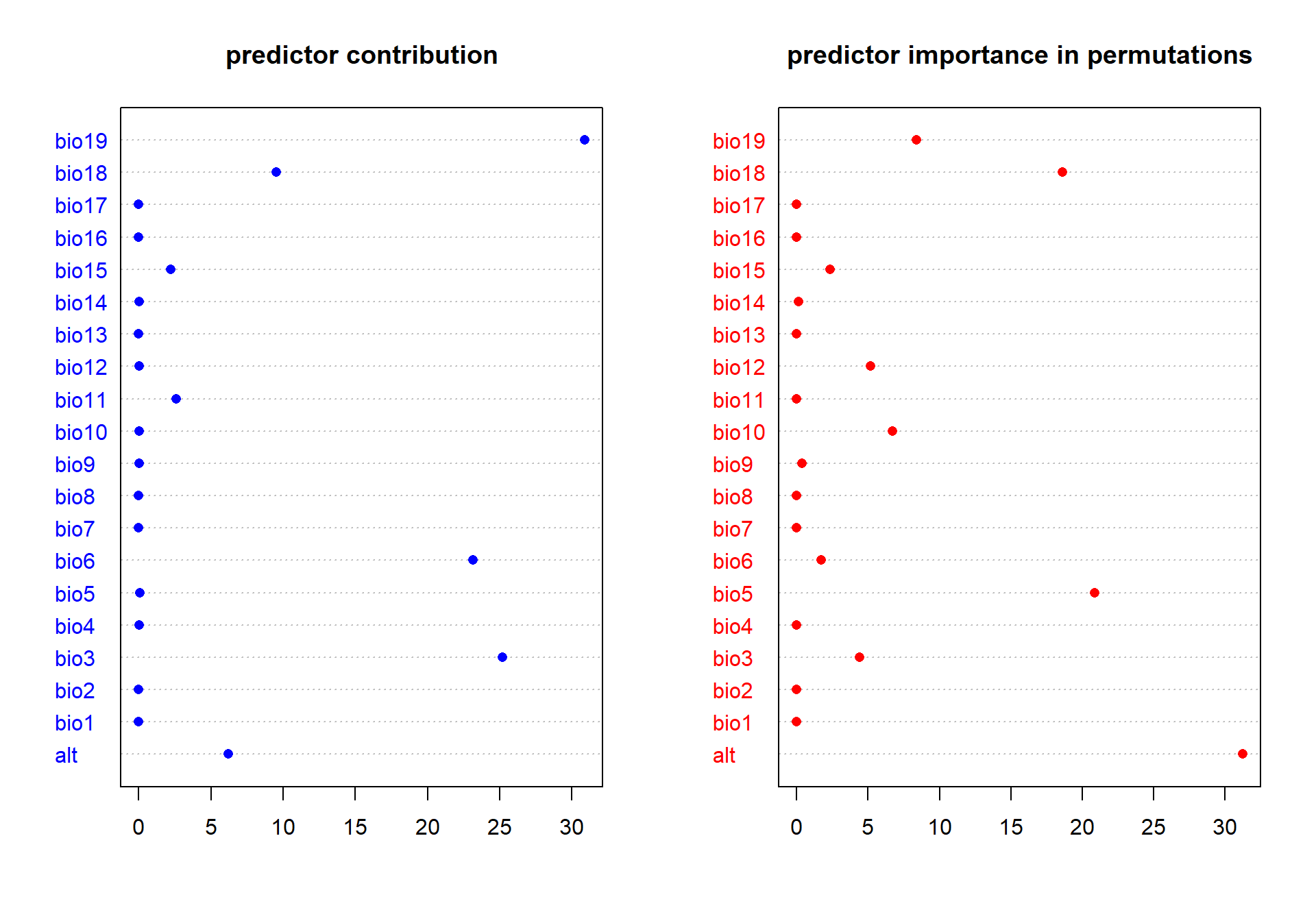

### review-model-15.png

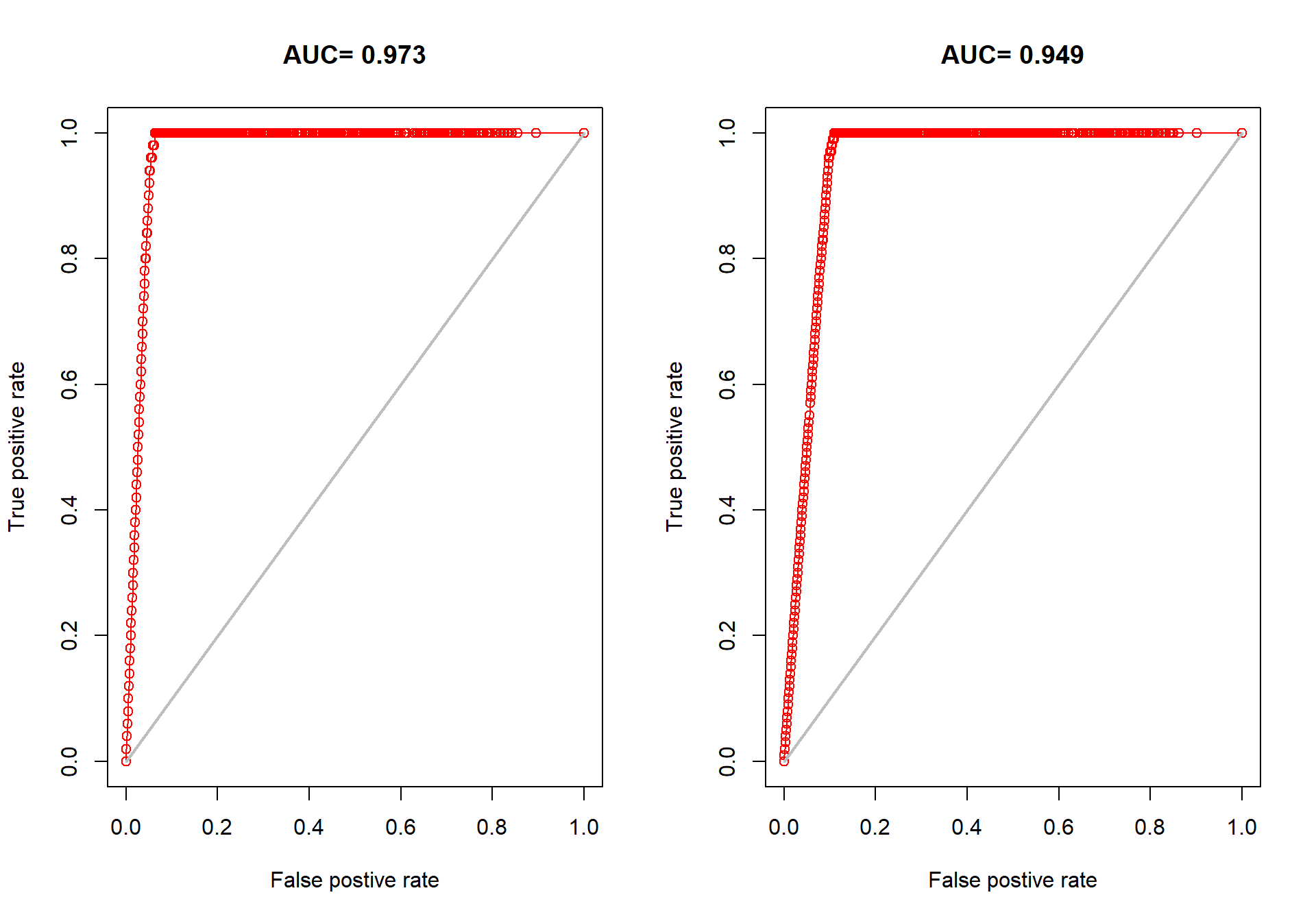

### review-model-16.png

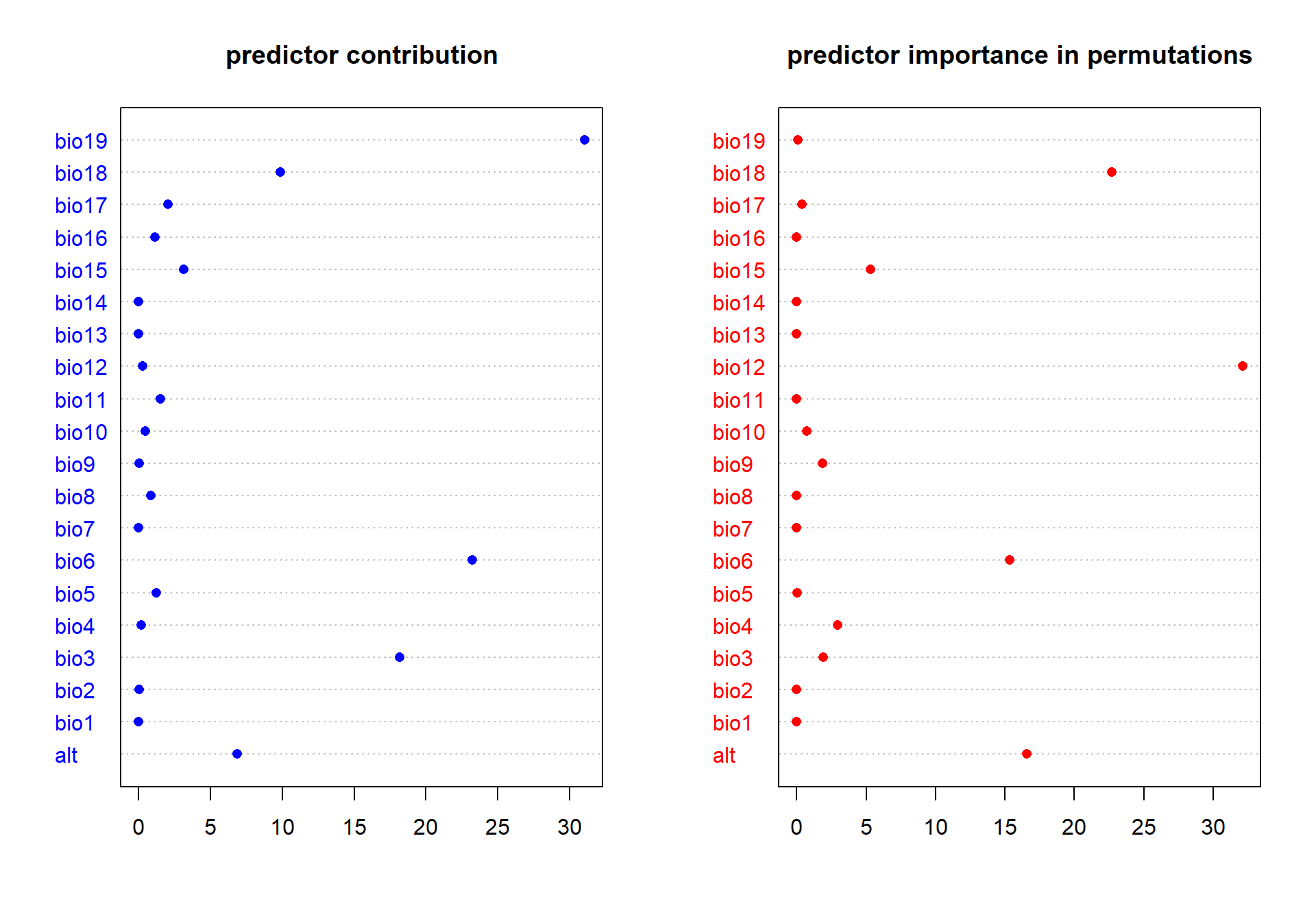

### review-model-17.png

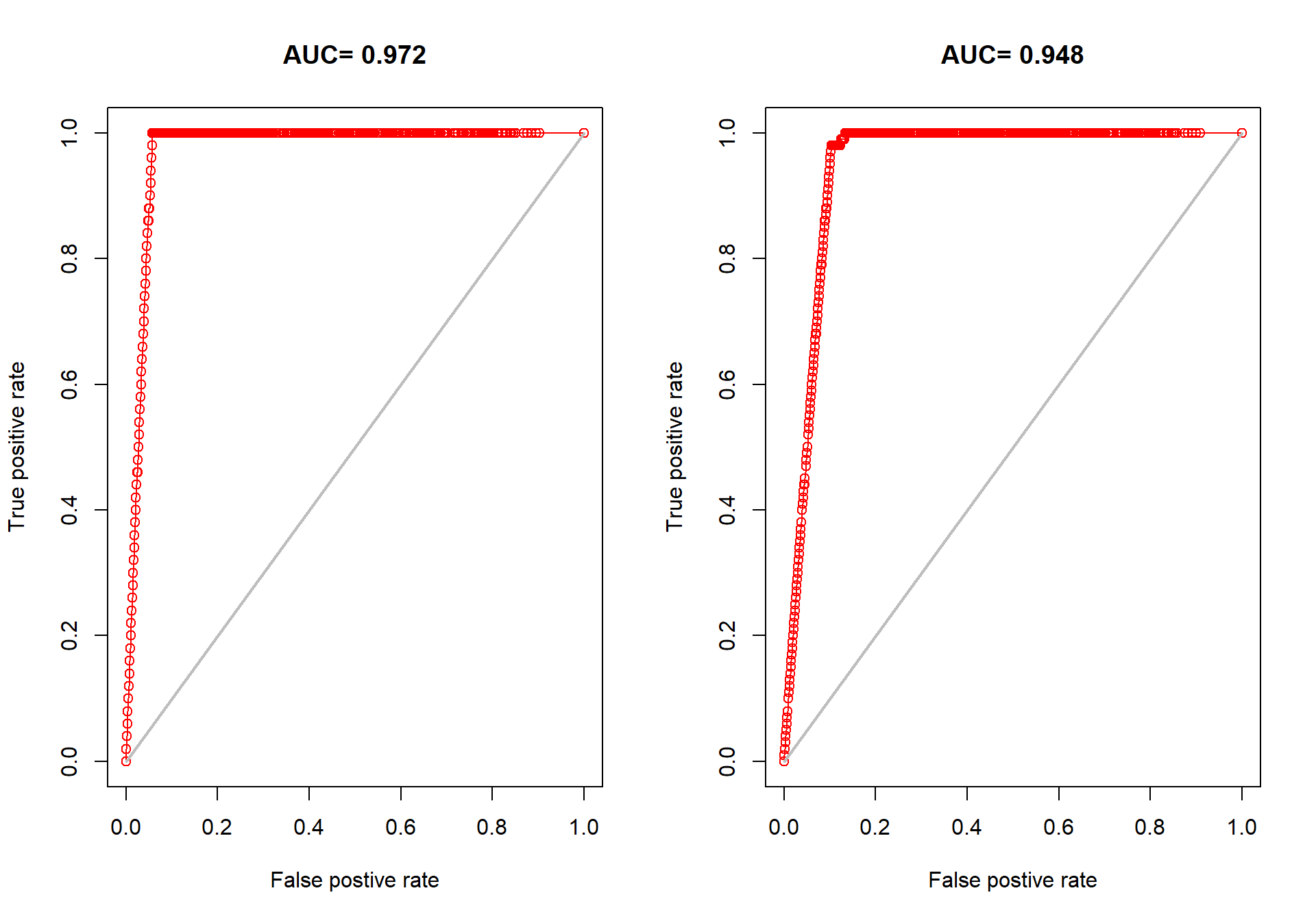

### review-model-18.png

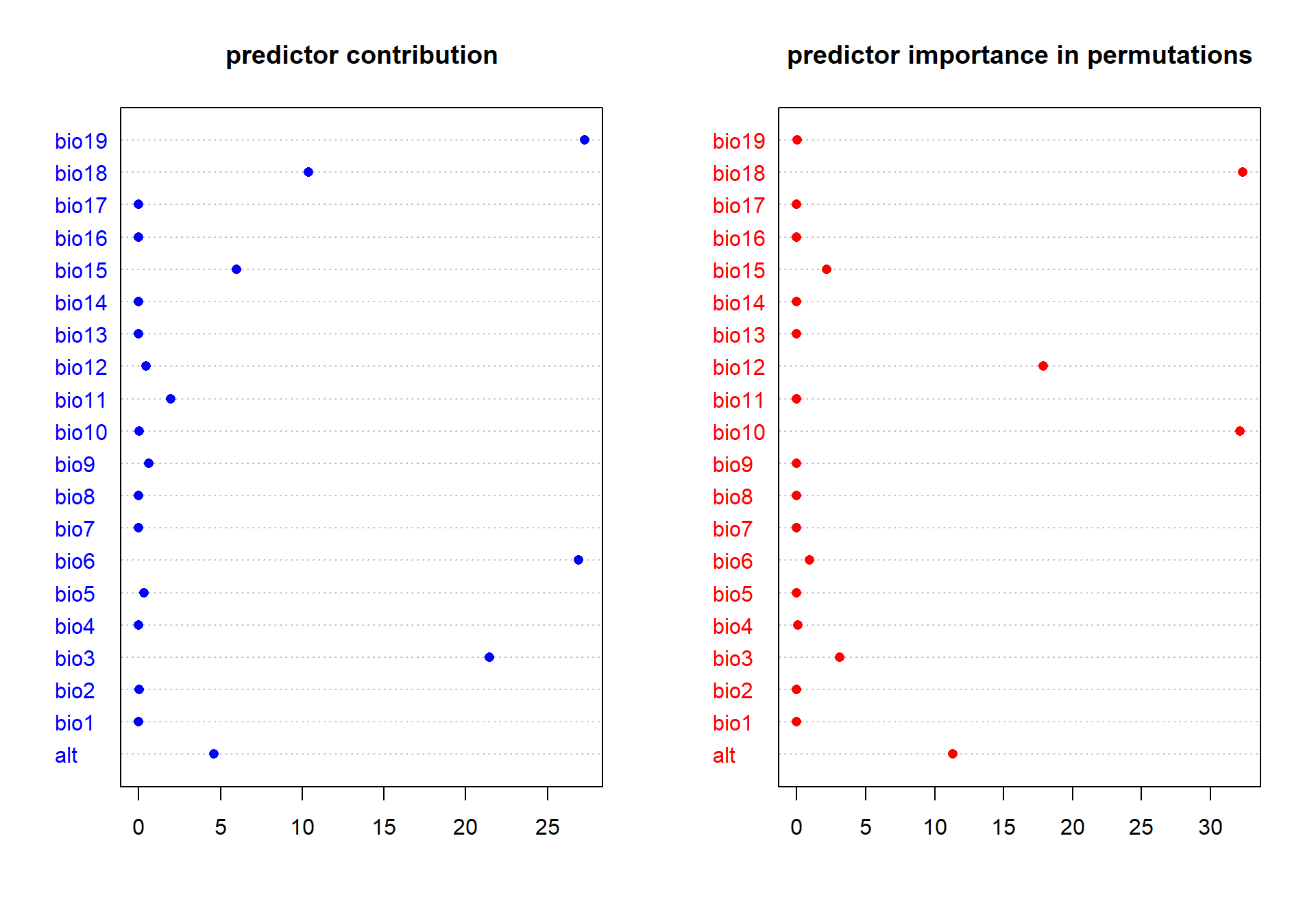

### review-model-19.png

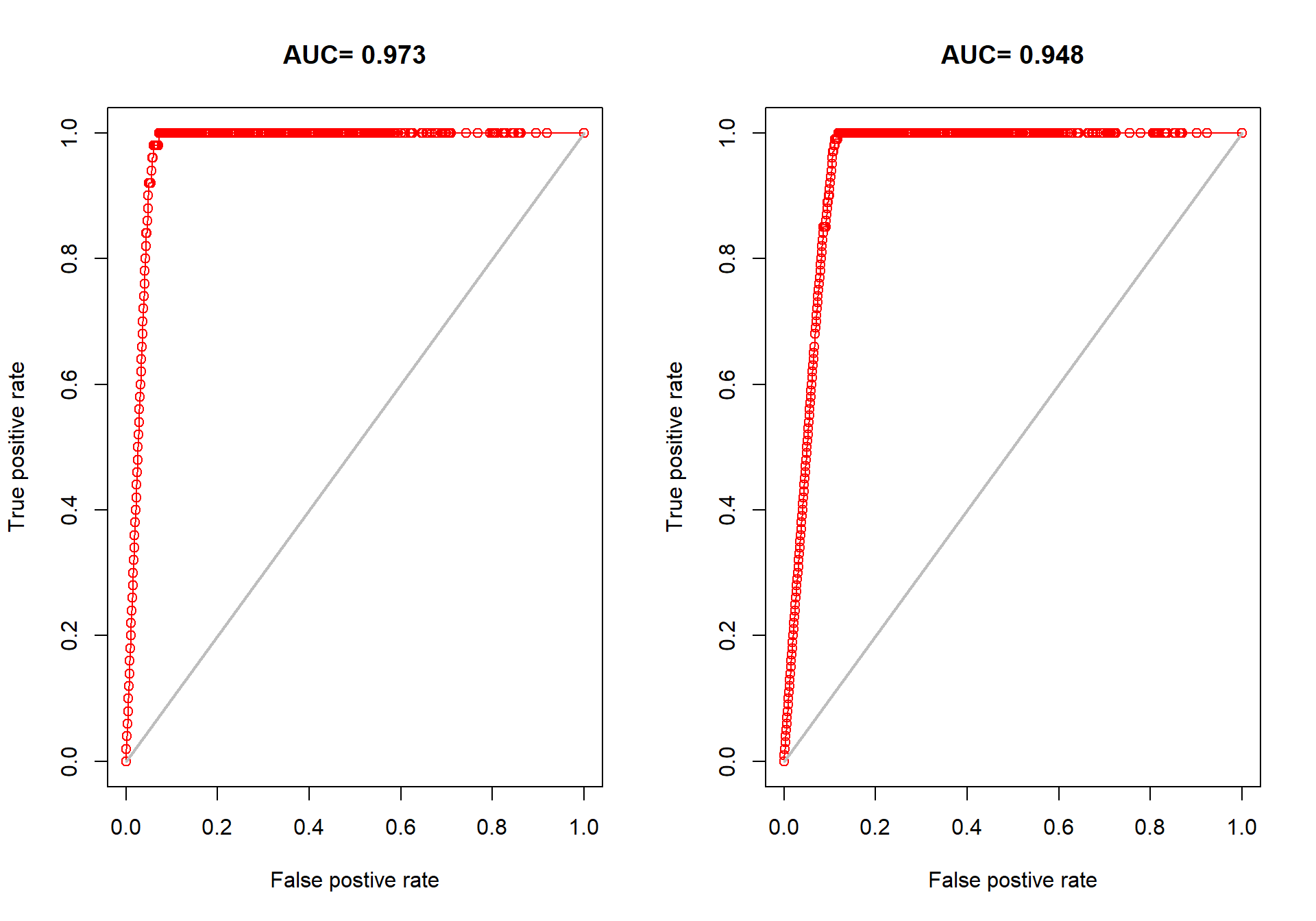

### review-model-20.png

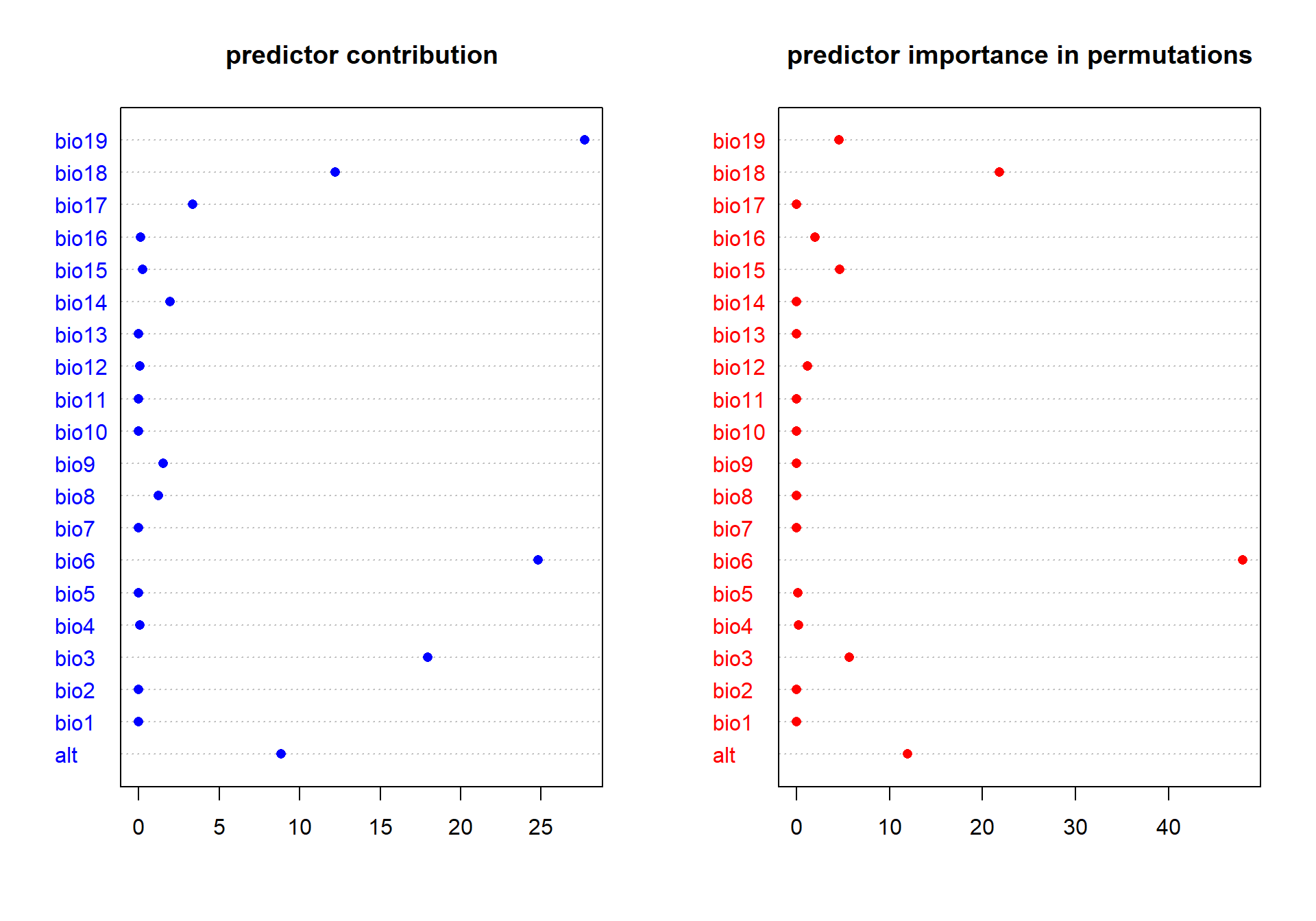

### unnamed-chunk-3-1.png

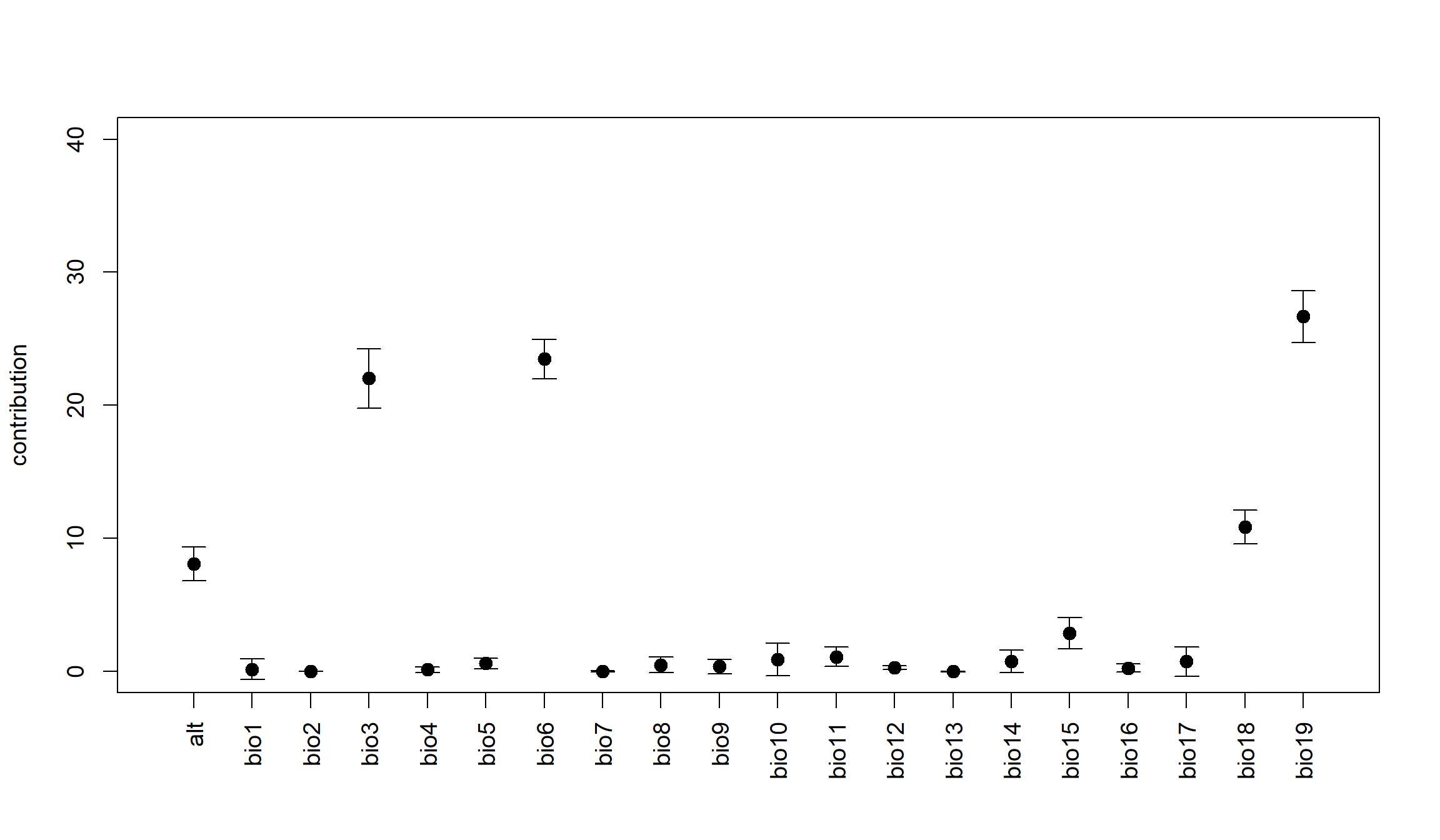

### unnamed-chunk-3-2.png

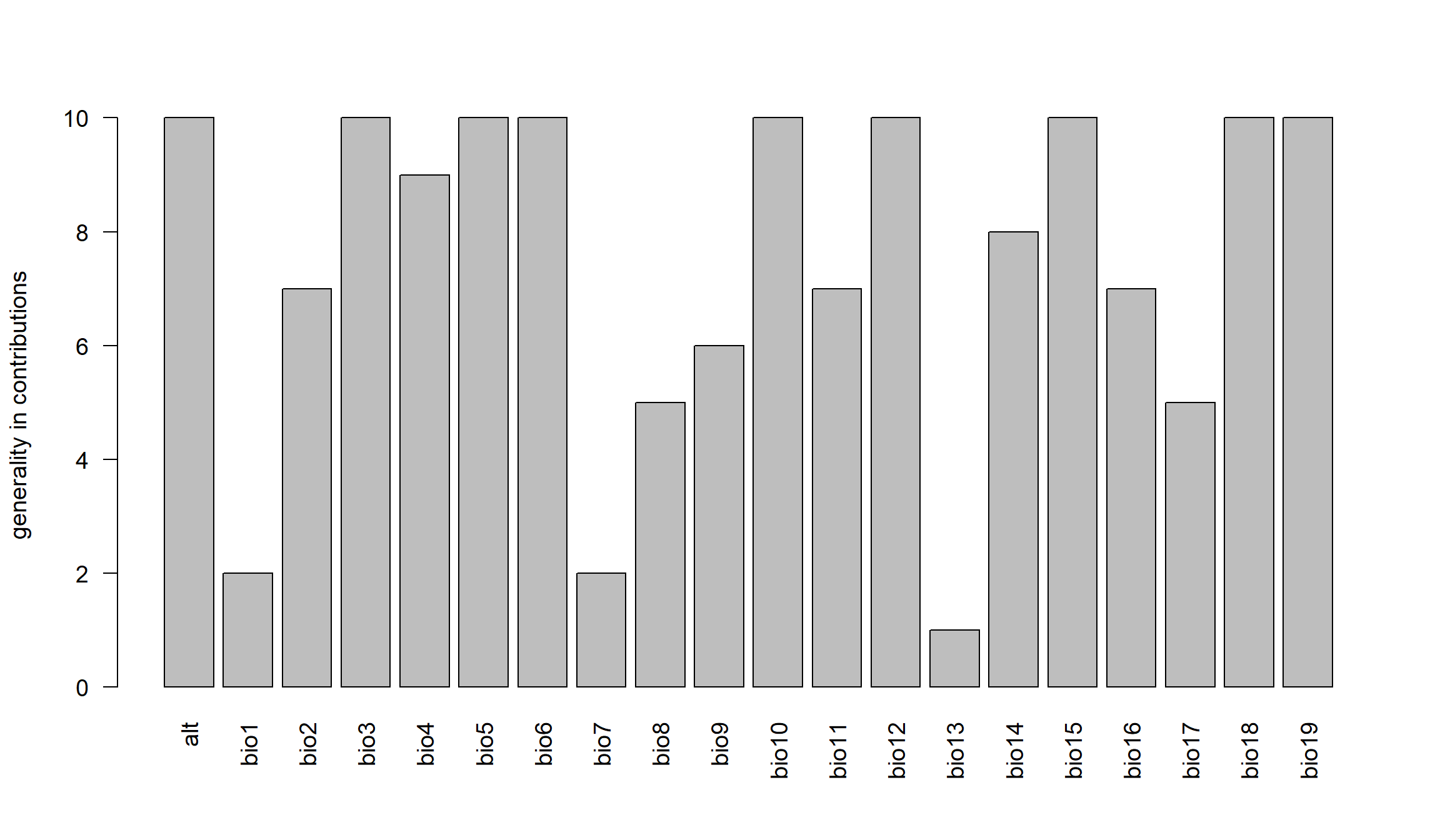

### unnamed-chunk-3-3.png

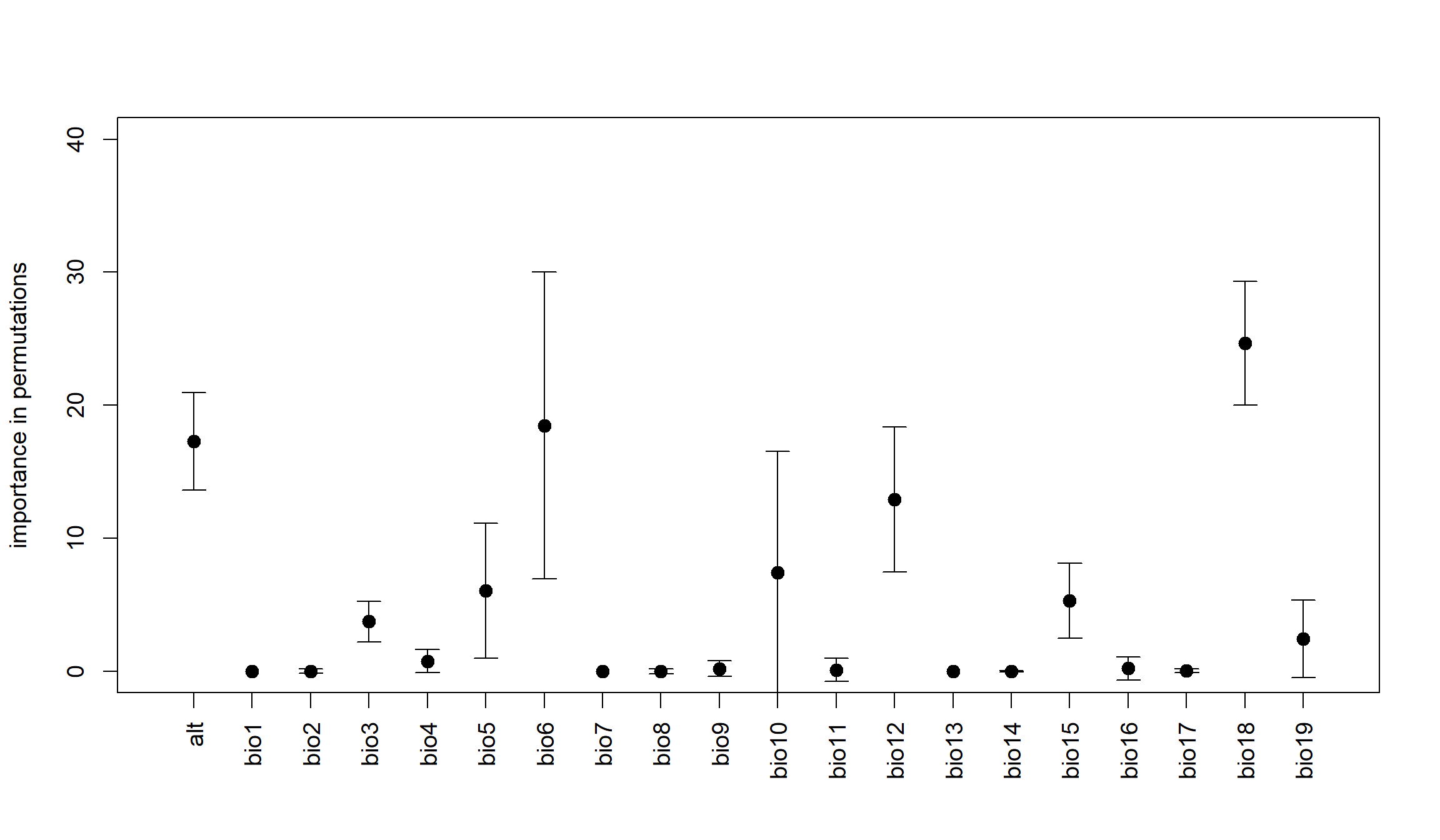

### unnamed-chunk-3-4.png

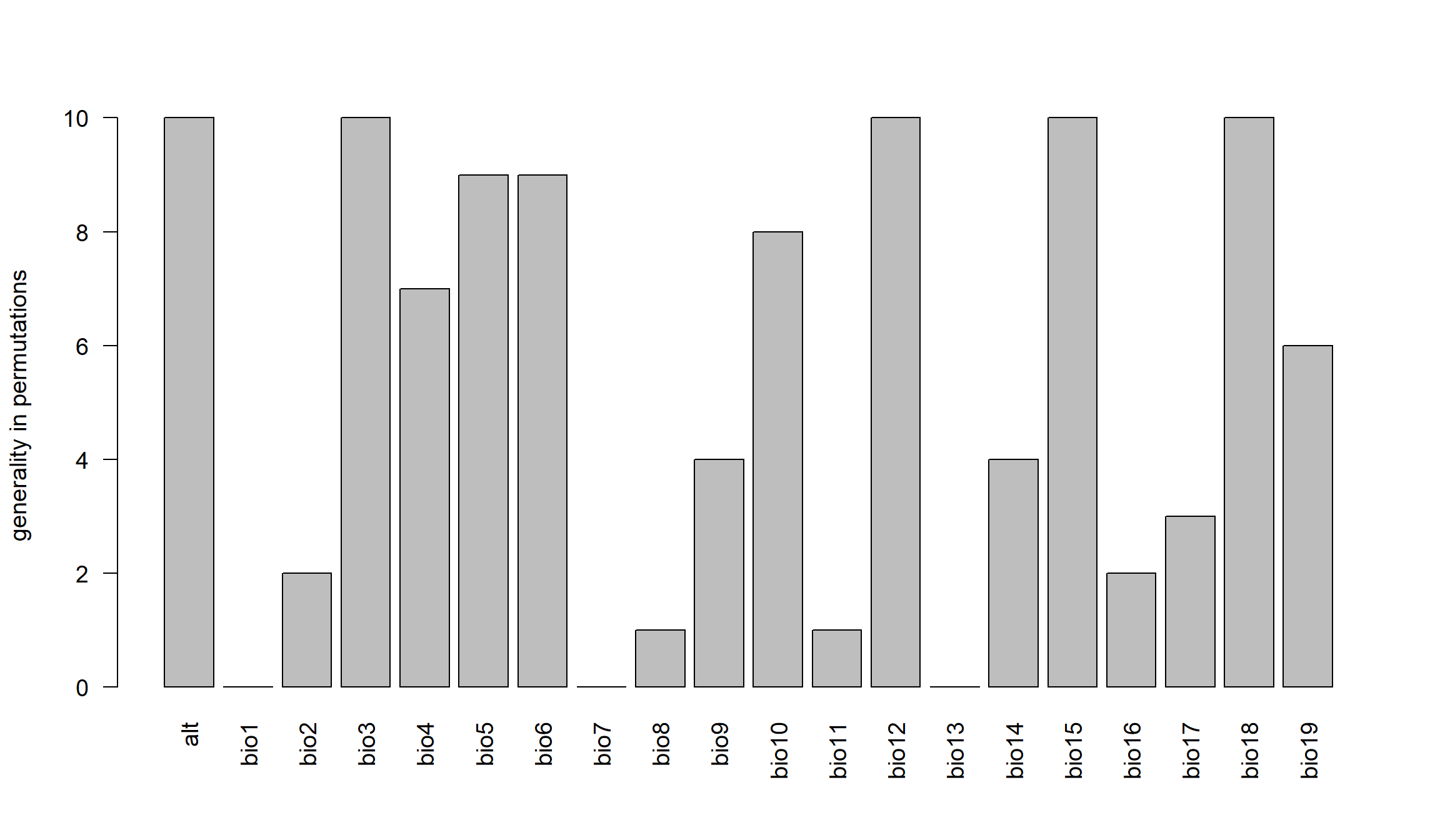

### unnamed-chunk-3-5.png

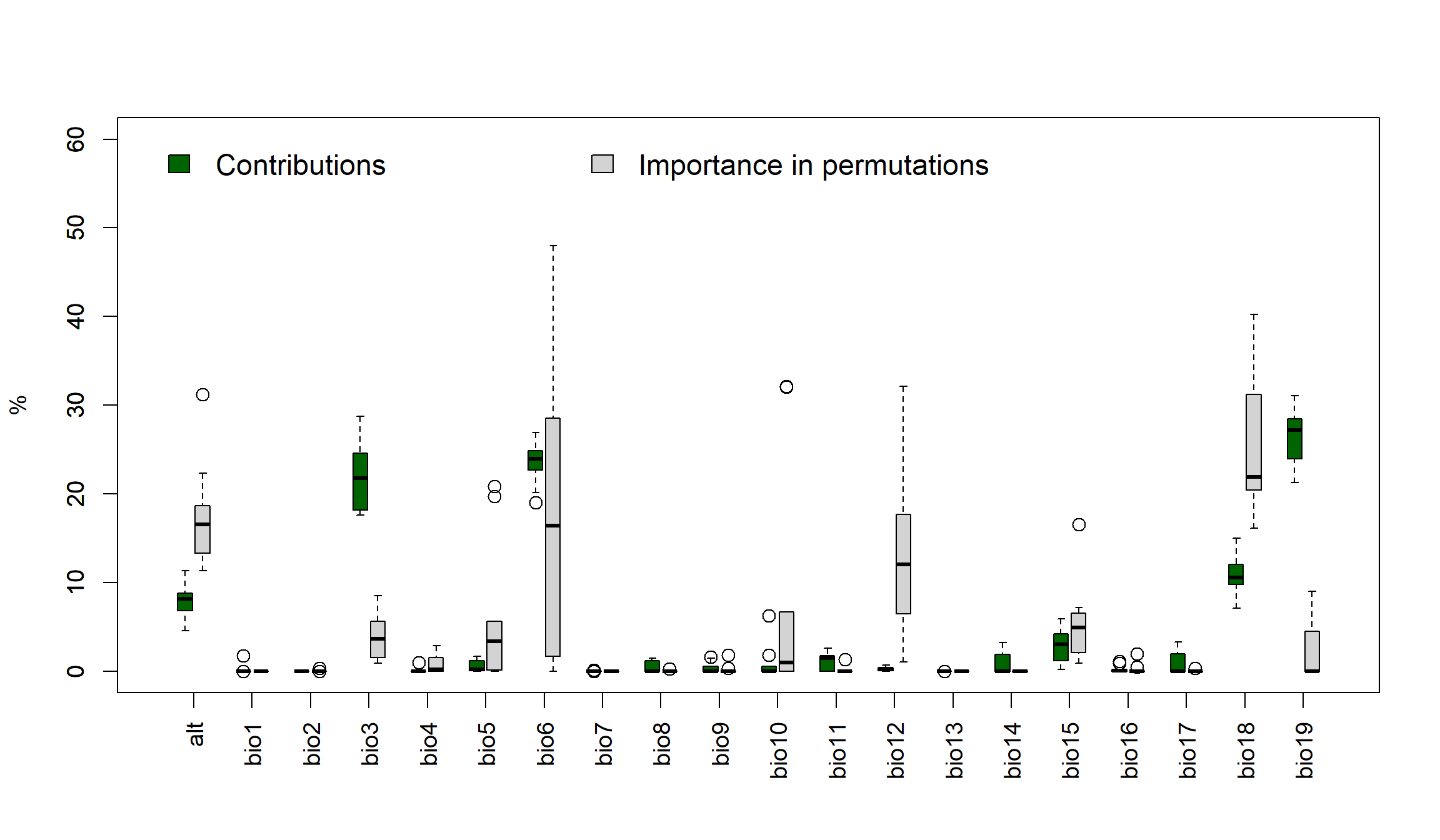

### unnamed-chunk-4-1.png

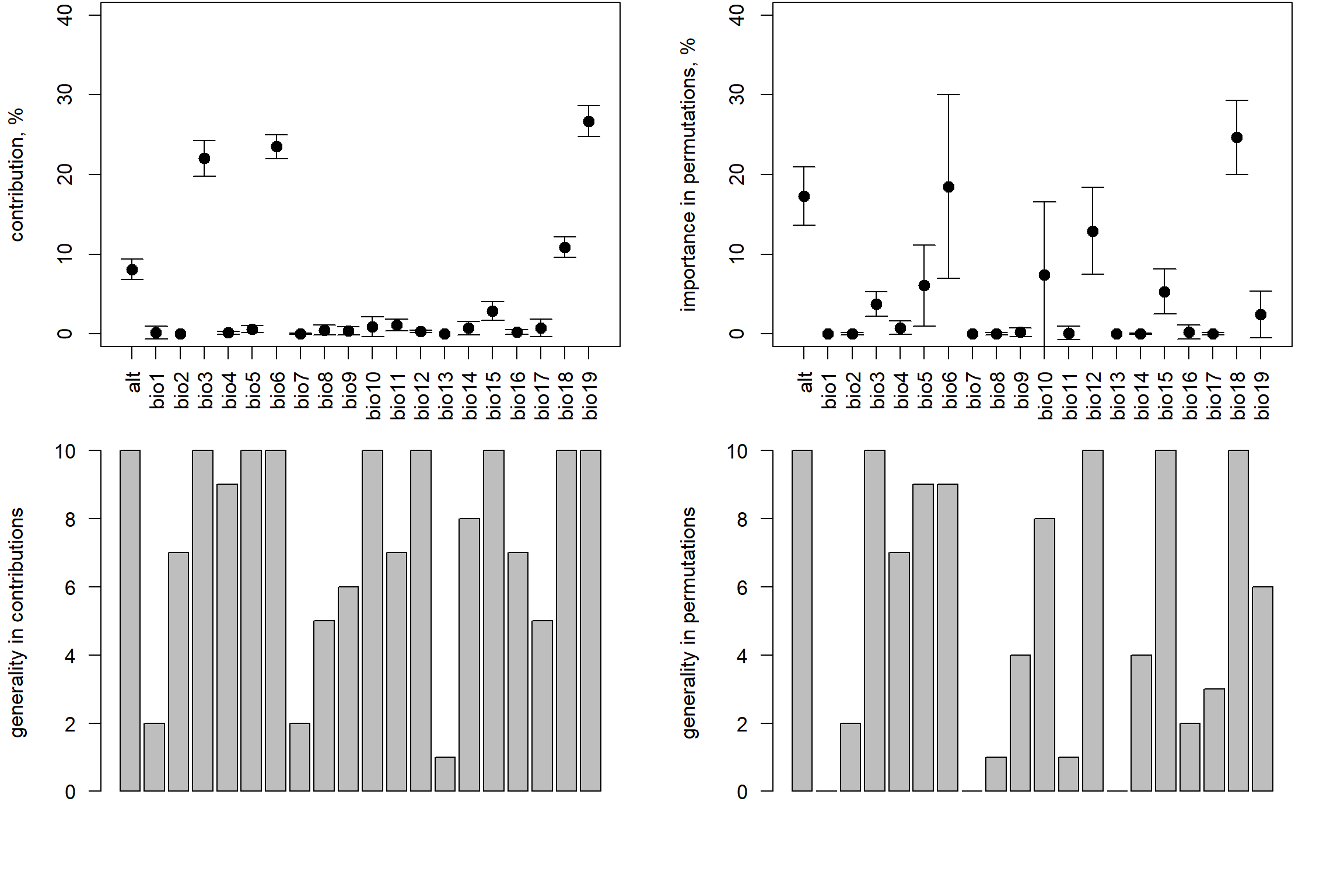

### unnamed-chunk-5-1.png

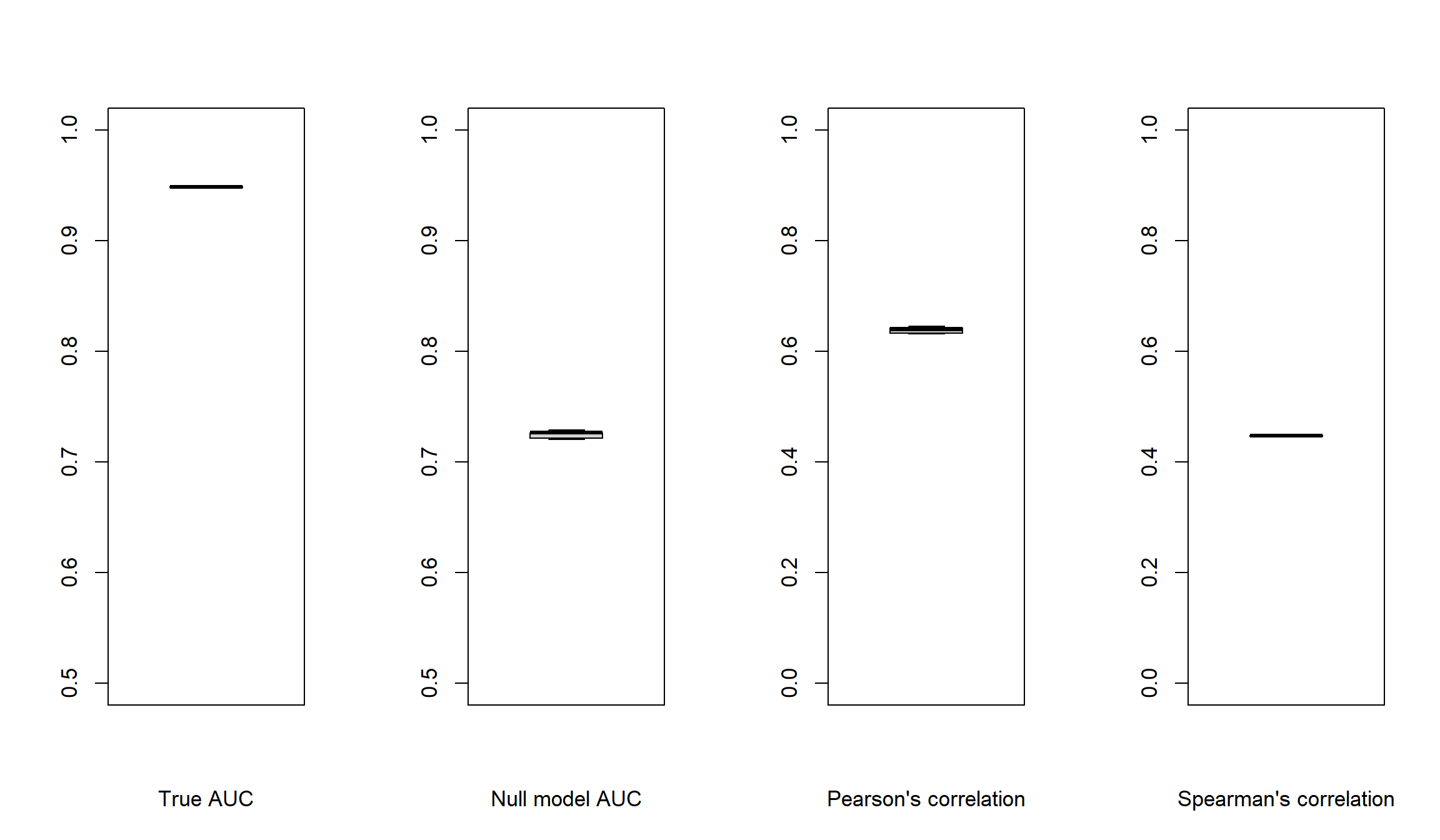

### unnamed-chunk-5-2.png

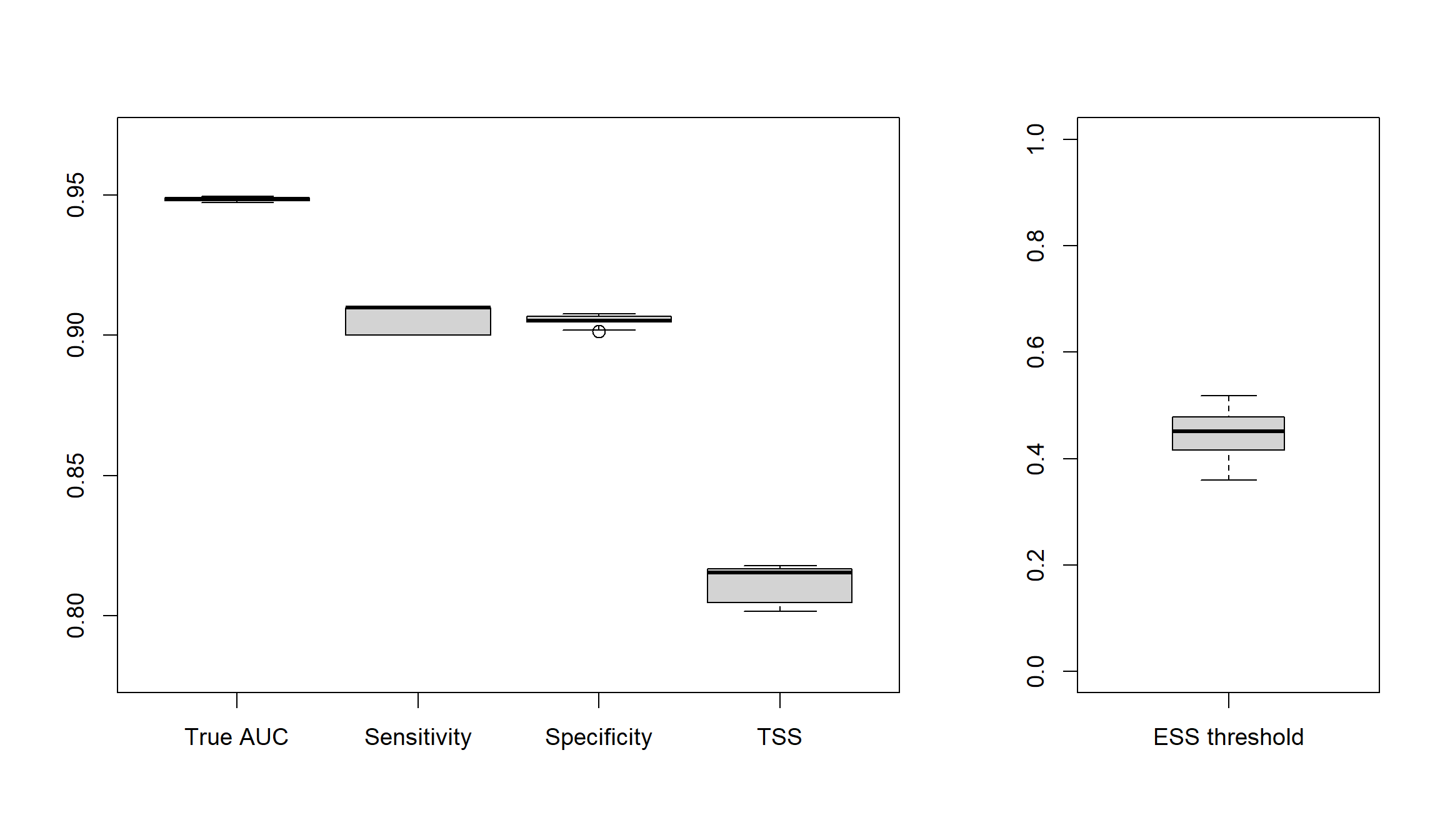

### unnamed-chunk-6-1.png

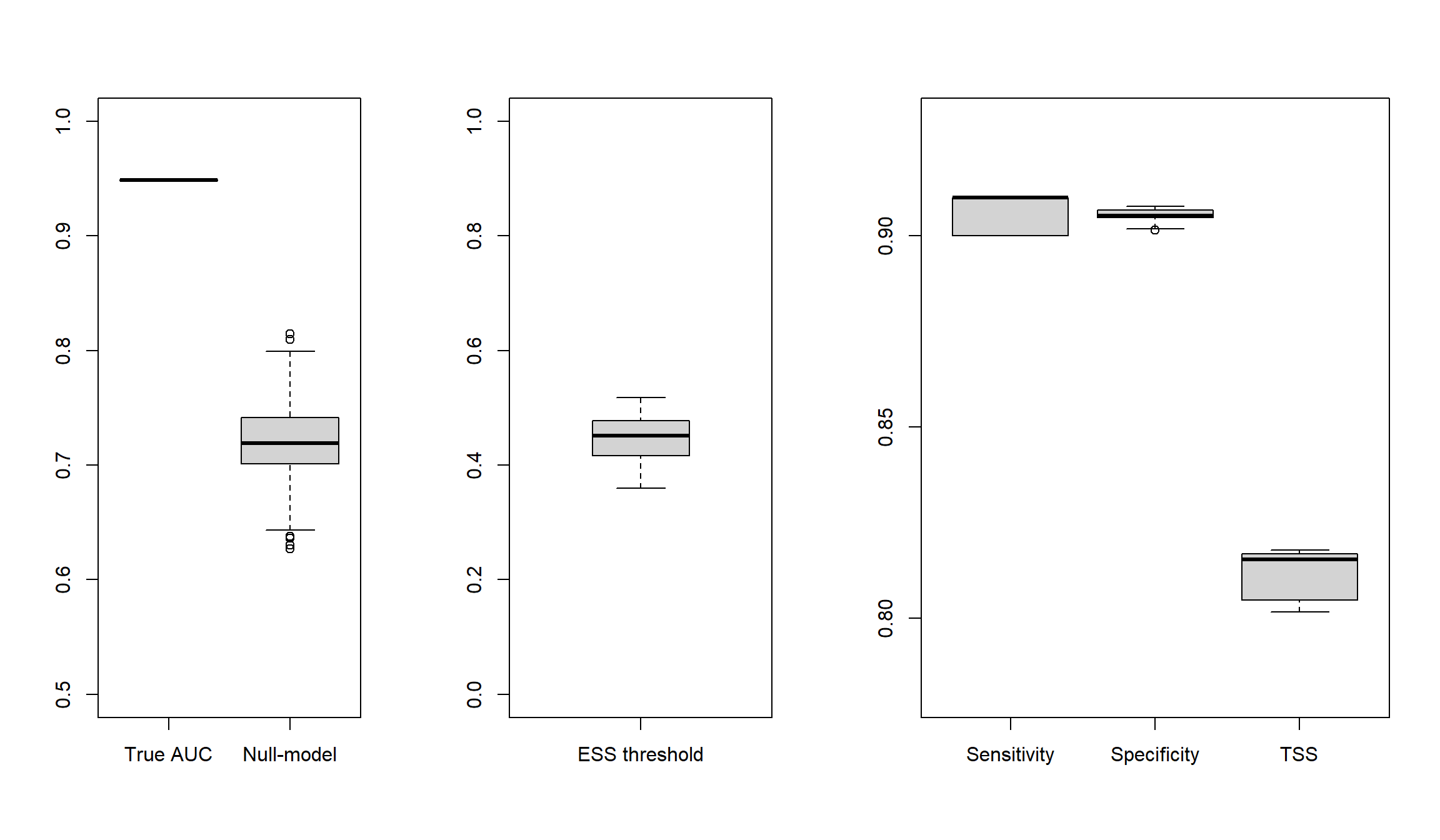

### visualization-1-1.png

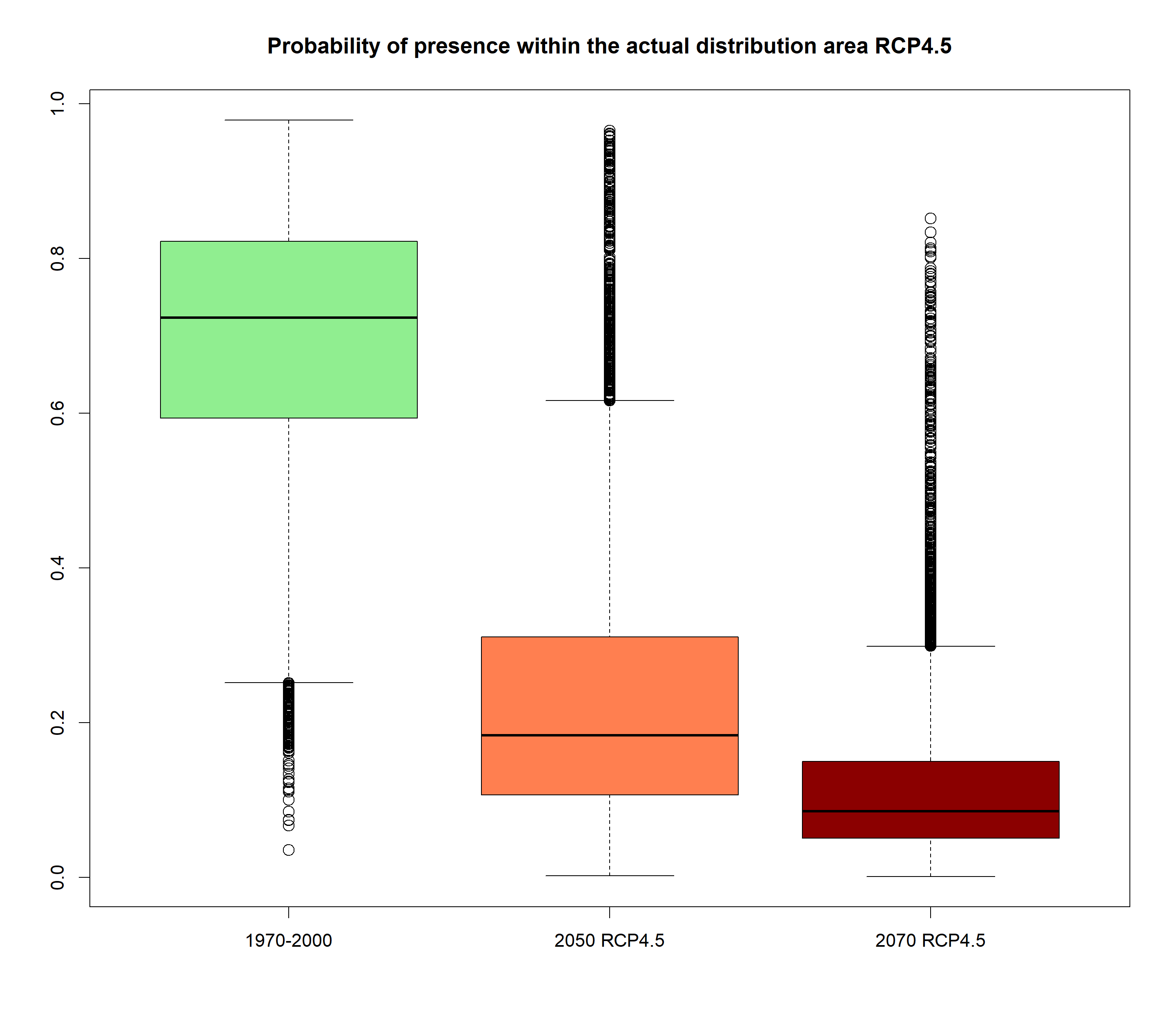

### visualization-1-2.png

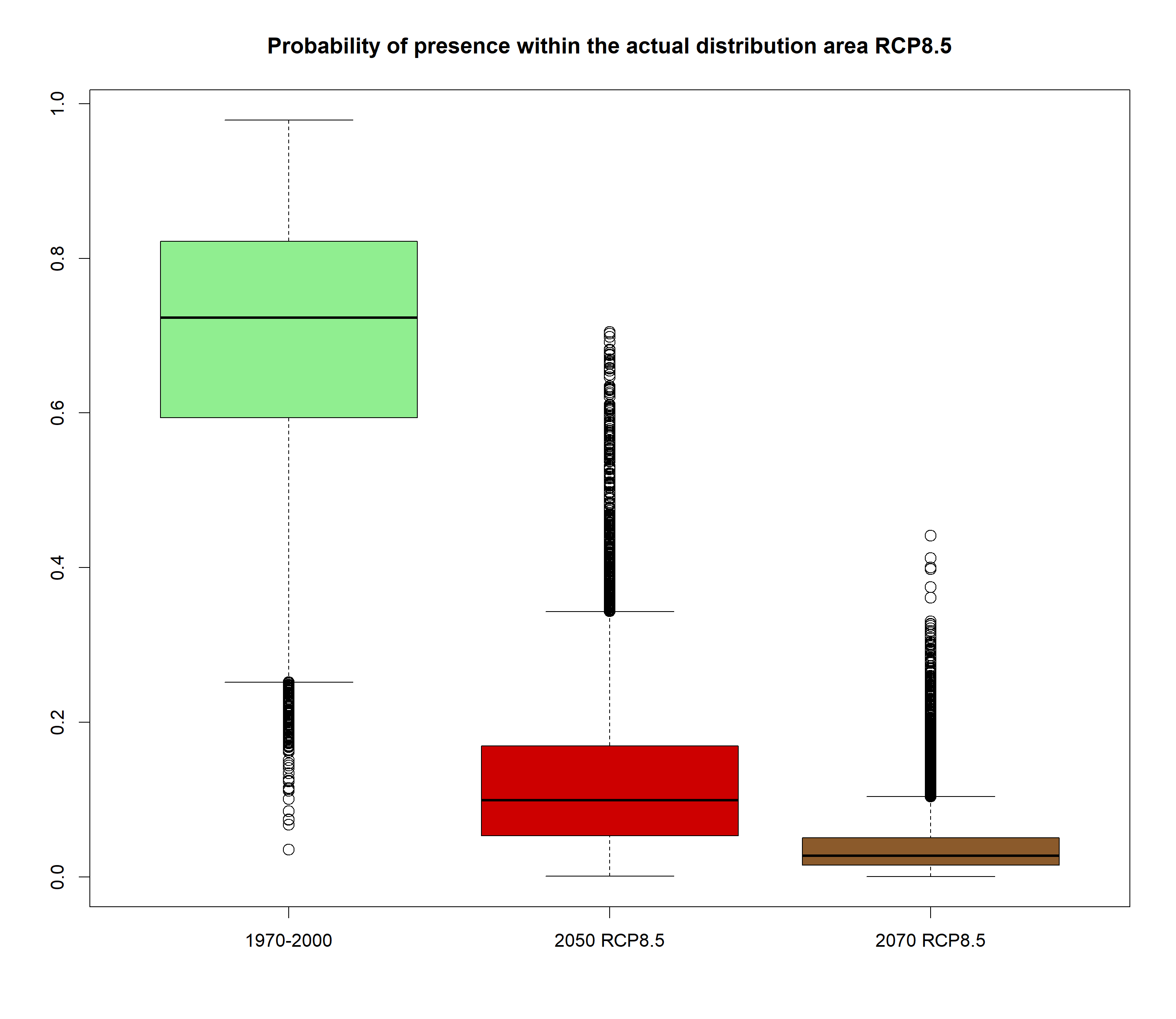

### visualization-1-3.png

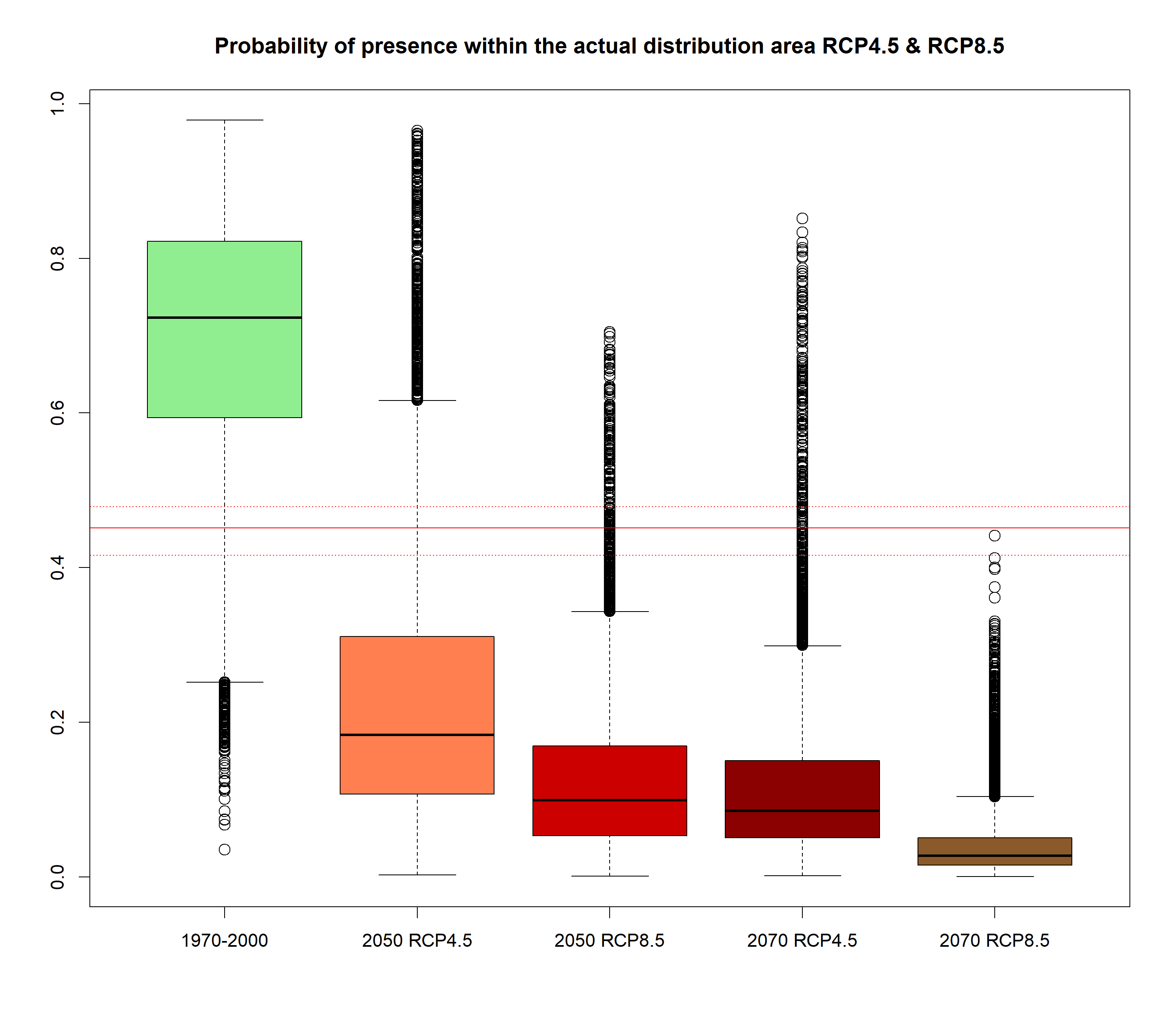

### visualization-5-1.png

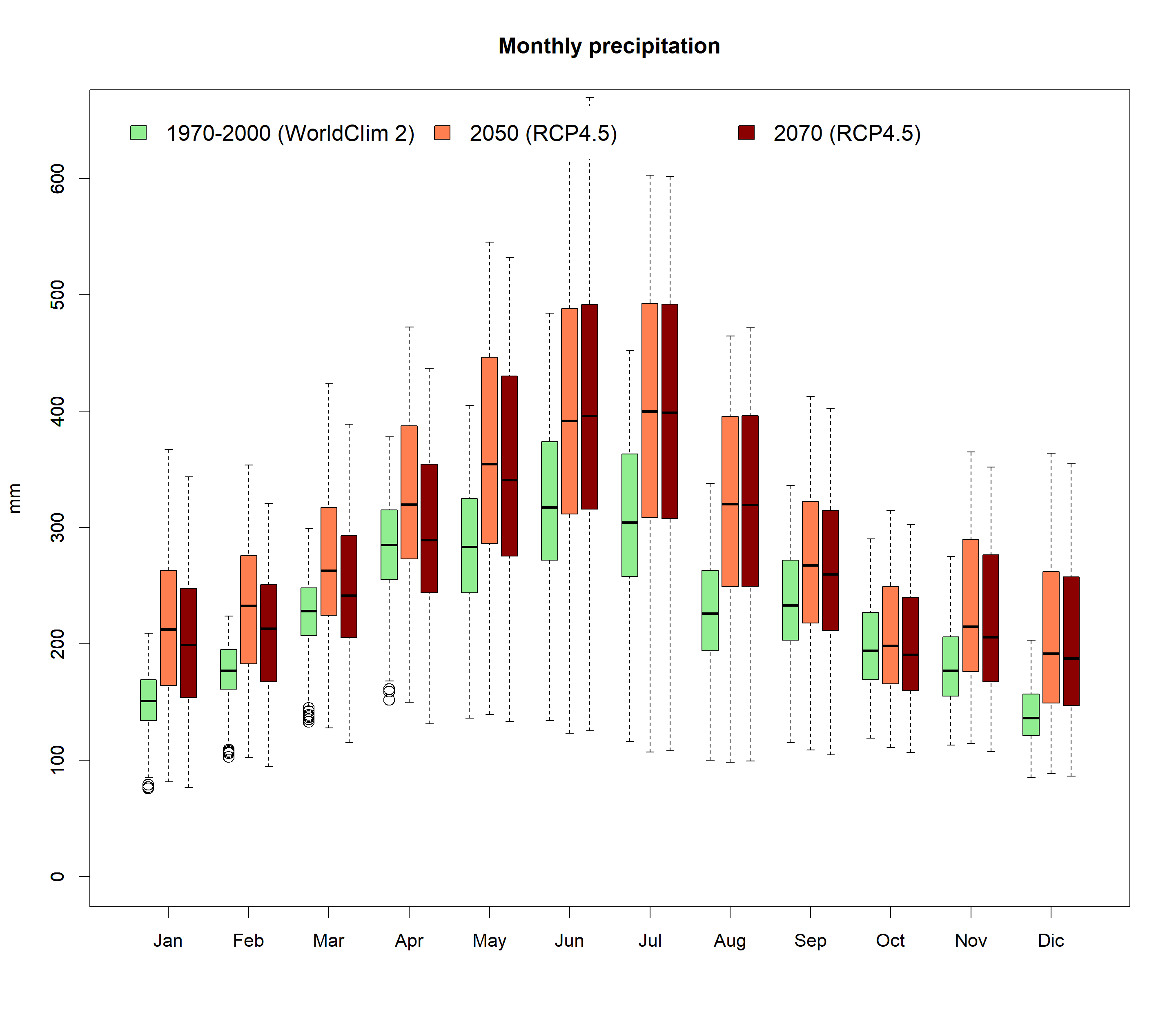

### visualization-5A-1.png

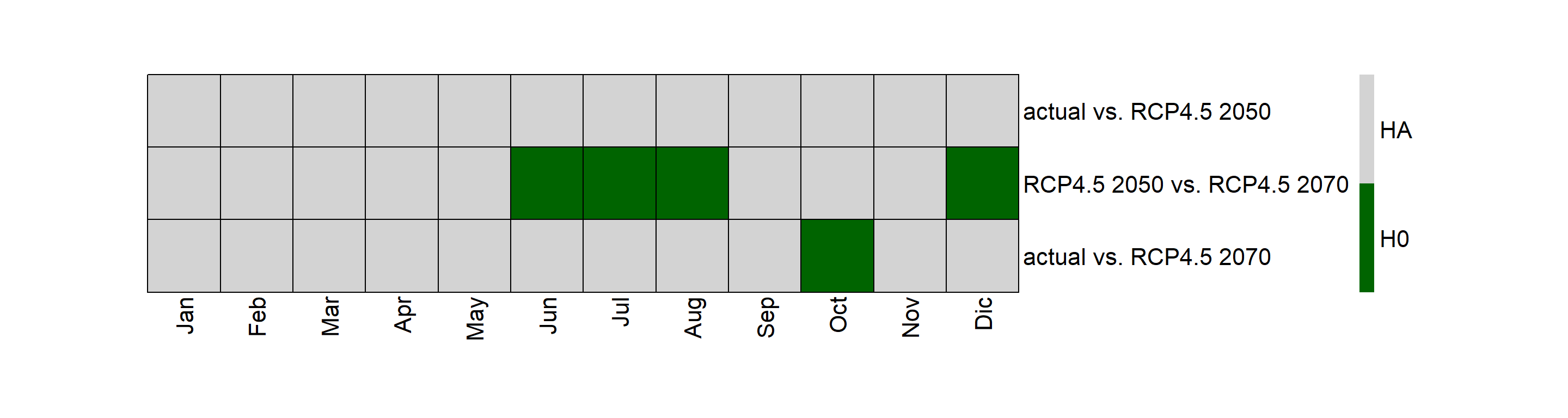

### visualization-5A-2.png

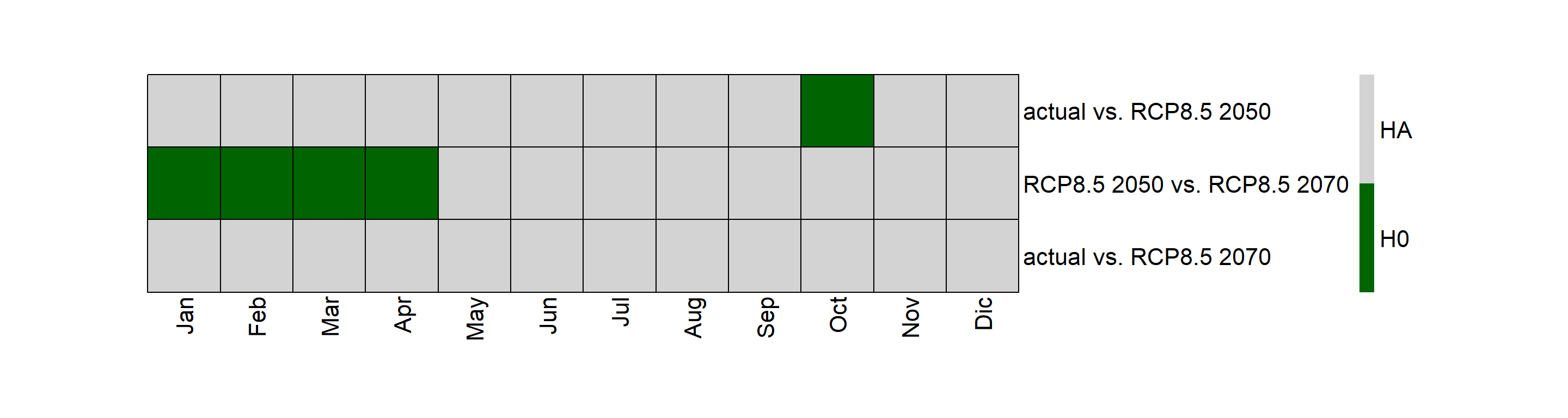

### visualization-5B-1.png

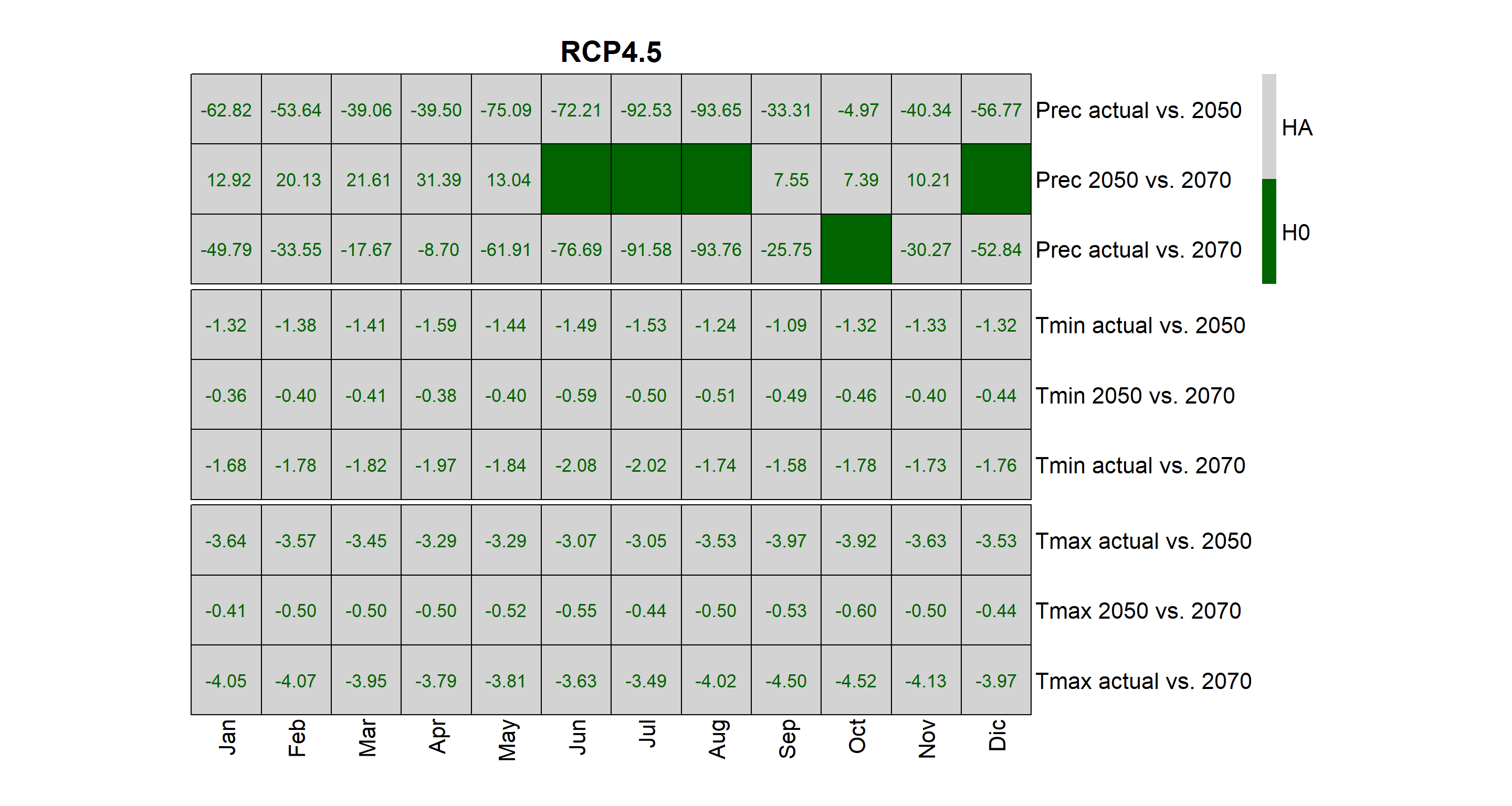

### visualization-5B-2.png

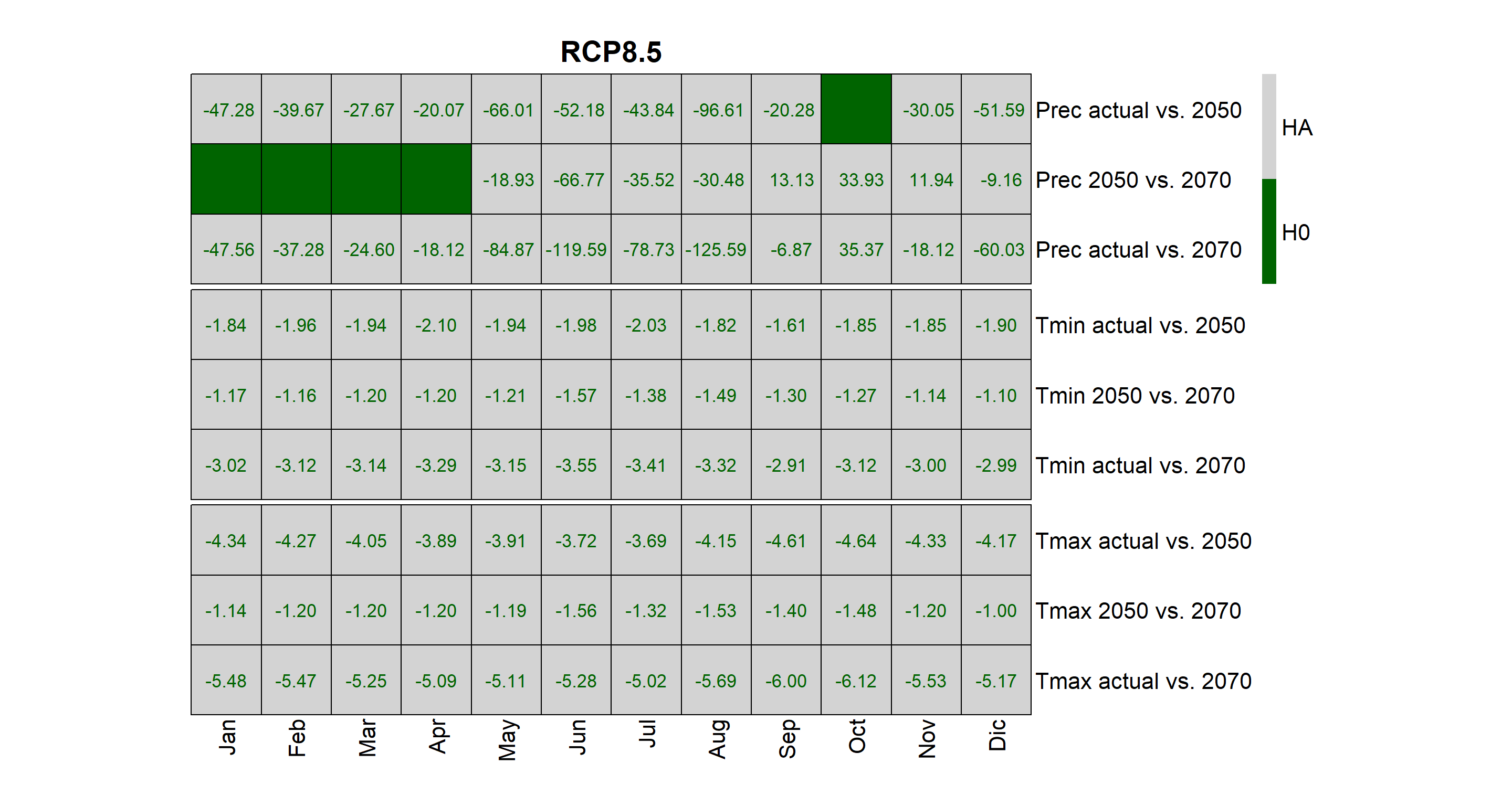

### visualization-5C-1.png

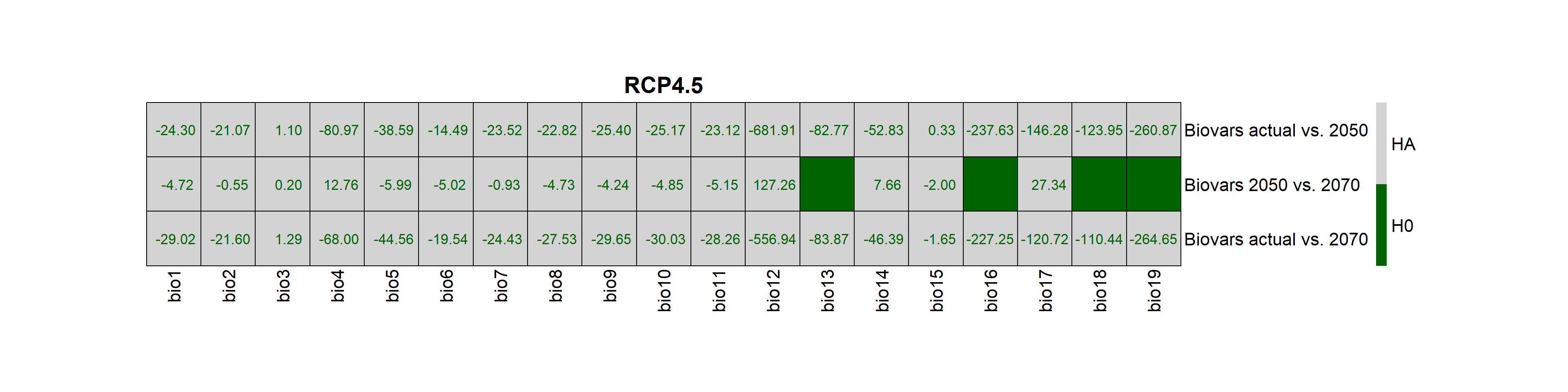

### visualization-5C-2.png

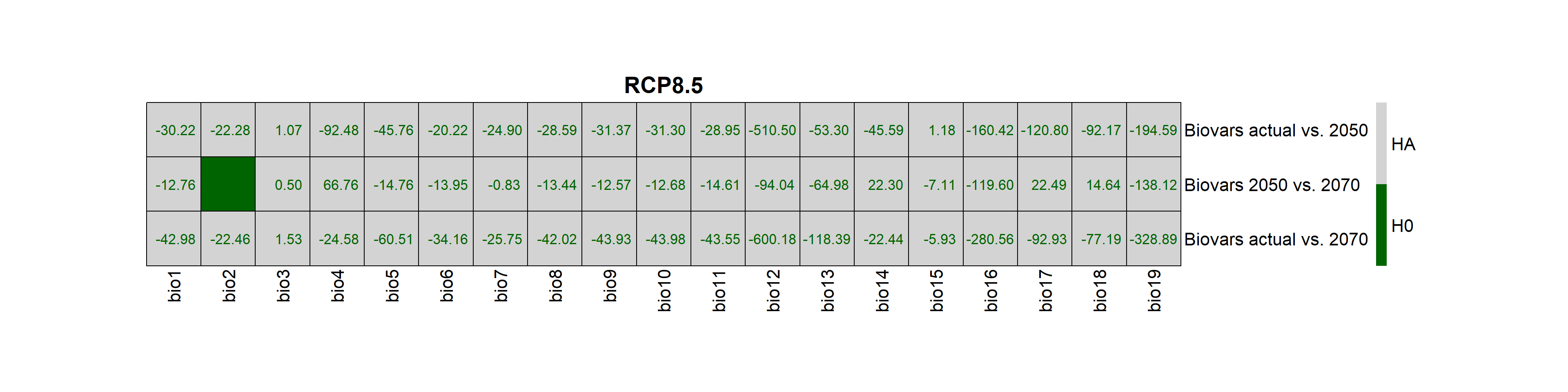

### visualization-6-1.png

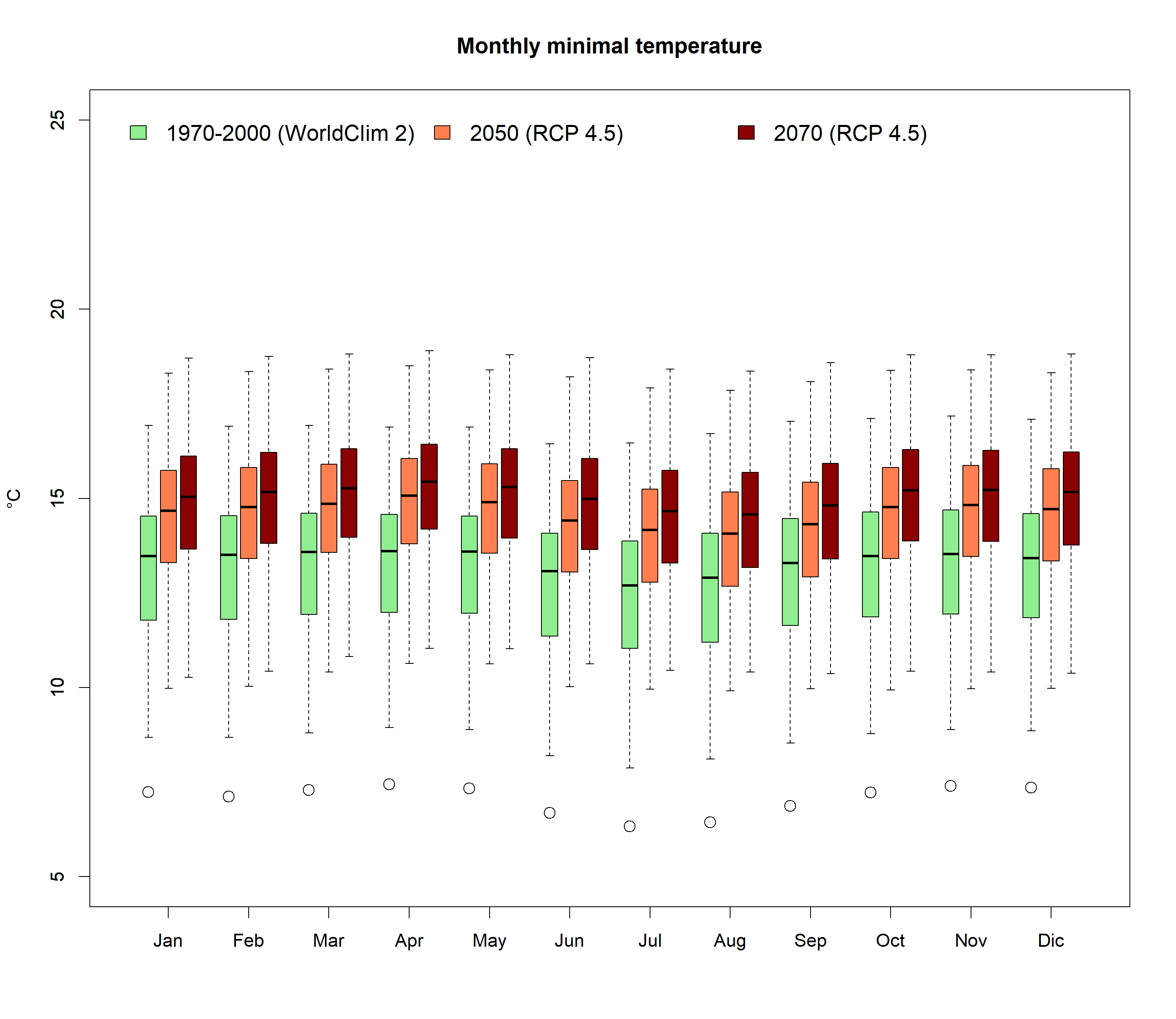

### visualization-7-1.png

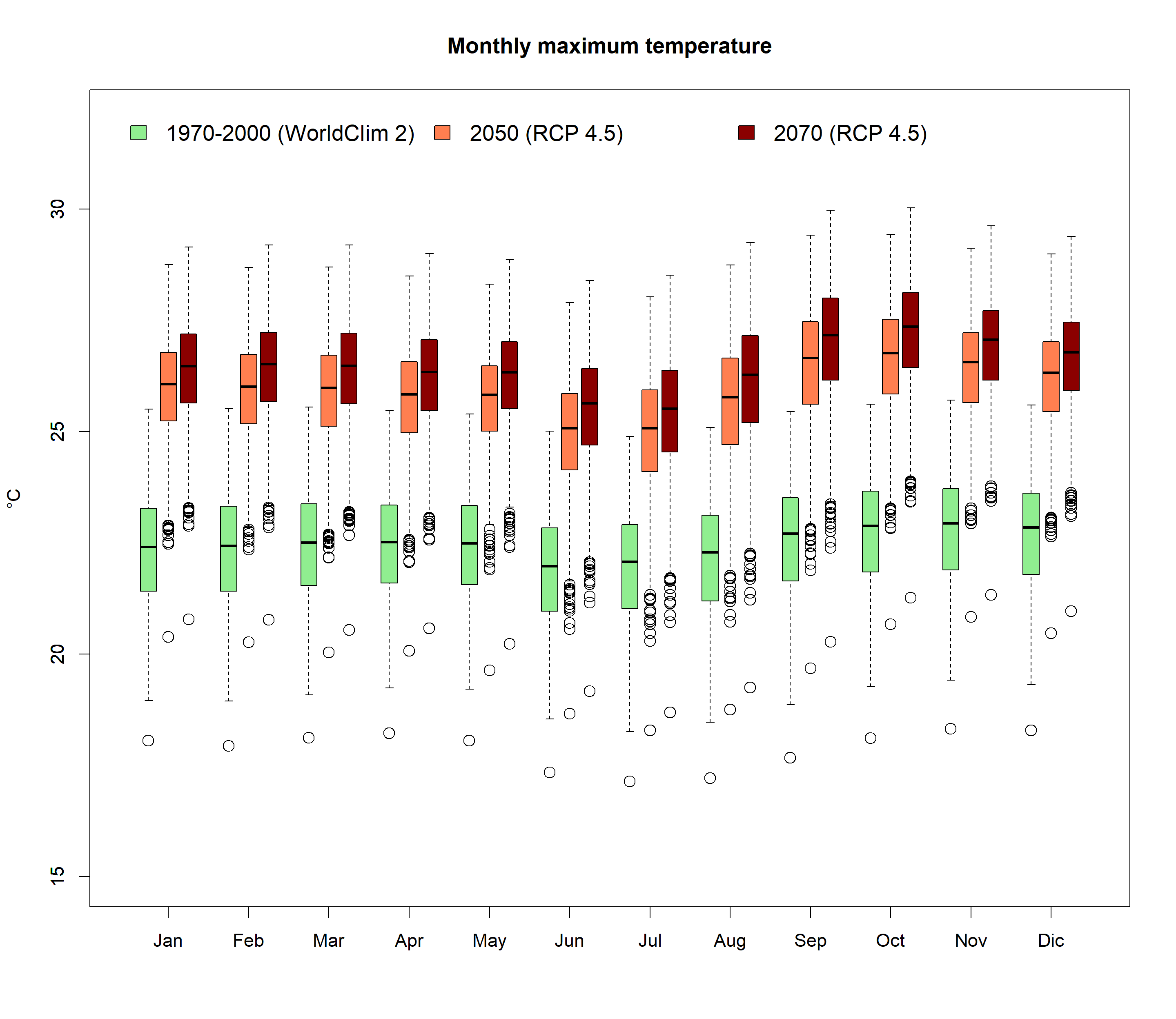

### visualization-8-1.png

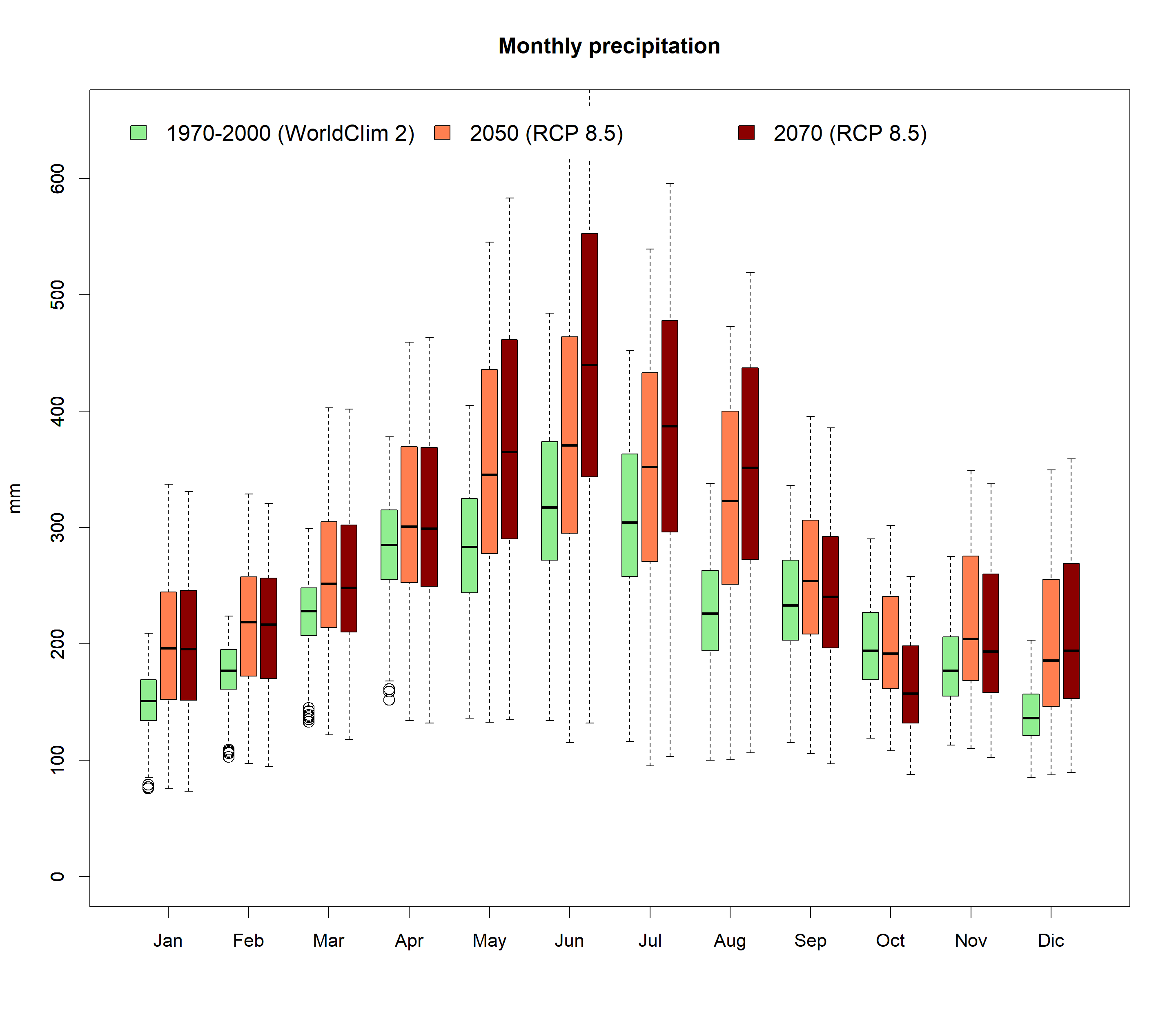

### visualization-9-1.png

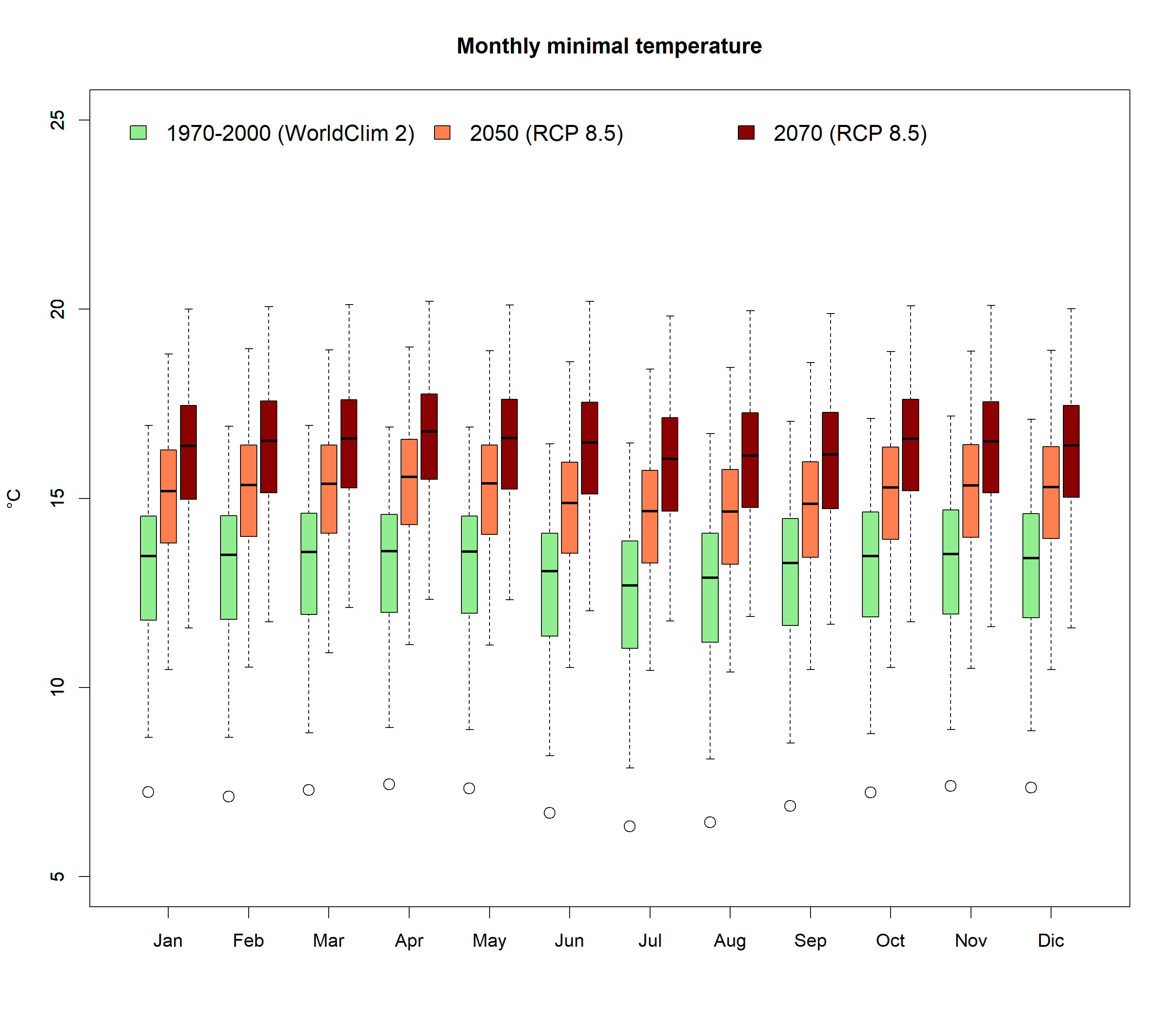

### visualization-10-1.png

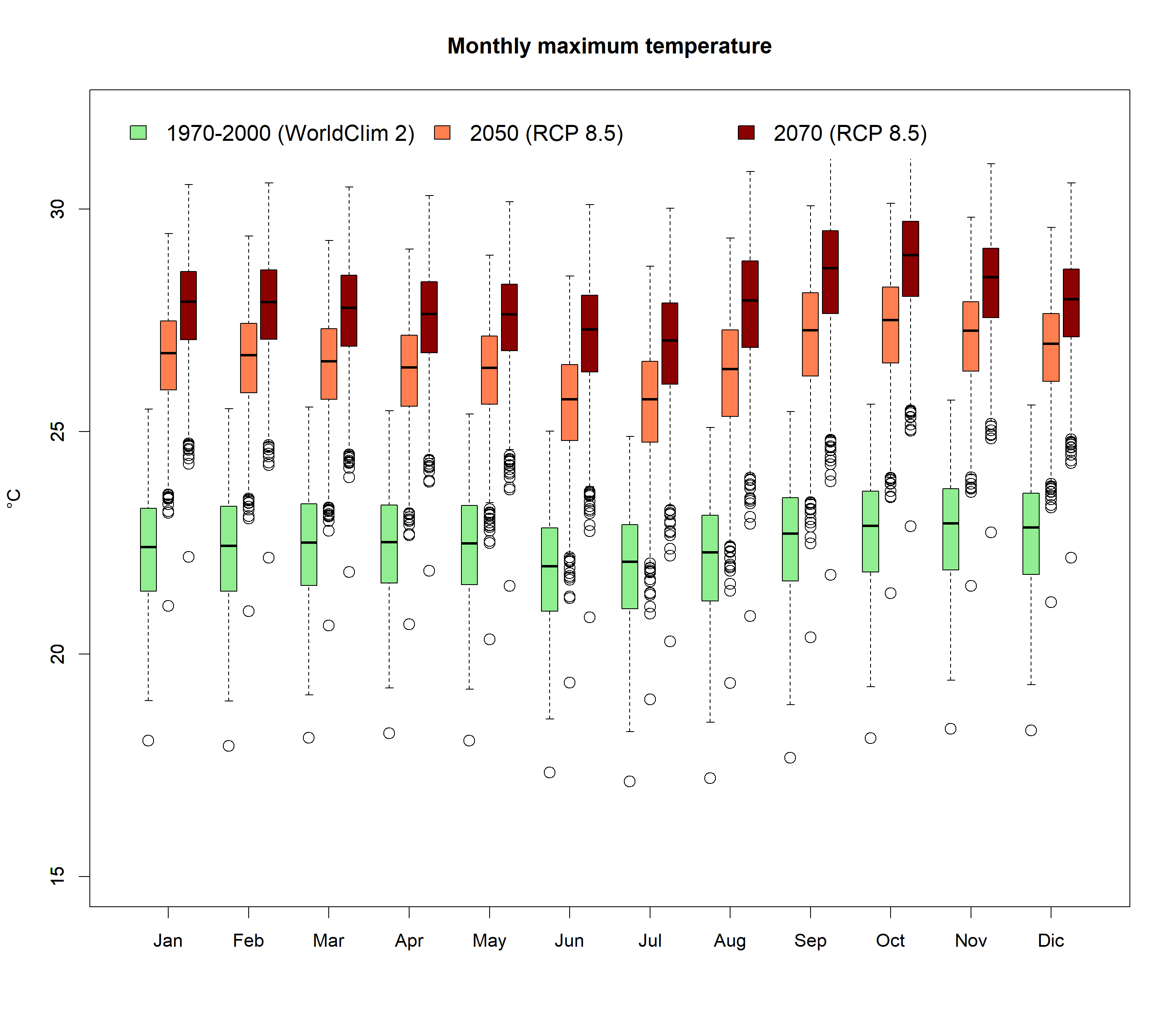

### visualization-11-1.png

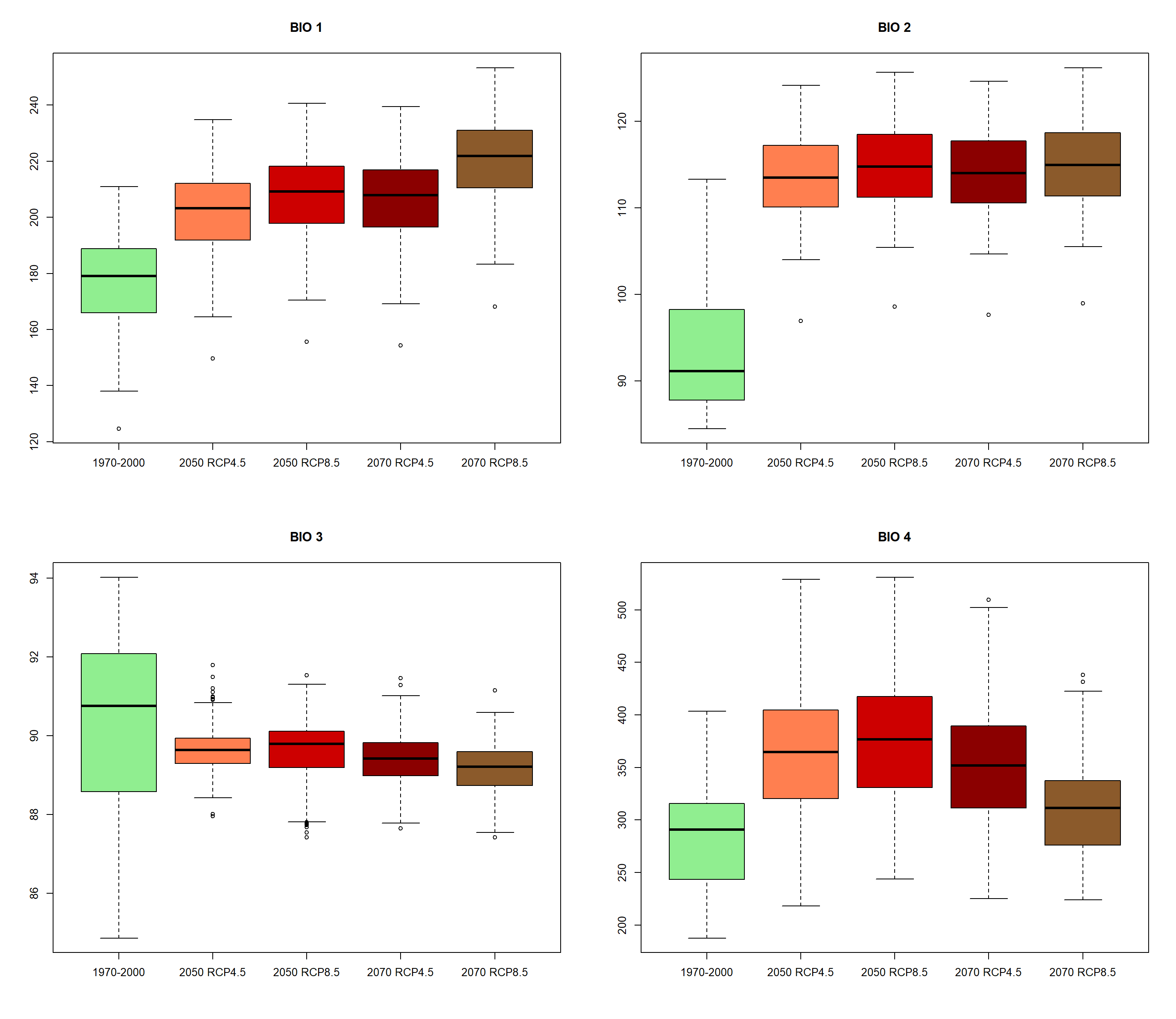

### visualization-11-2.png

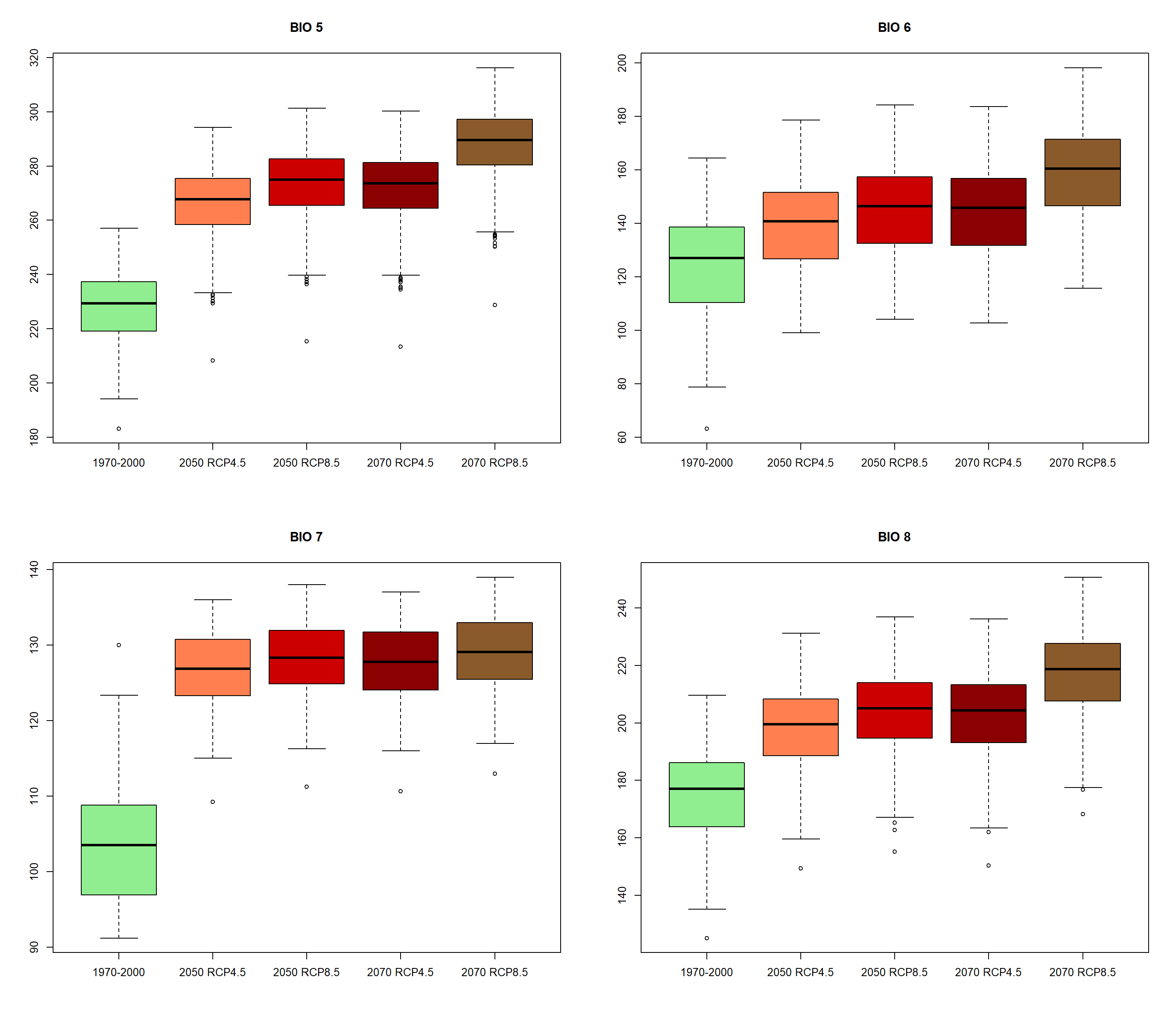

### visualization-11-3.png

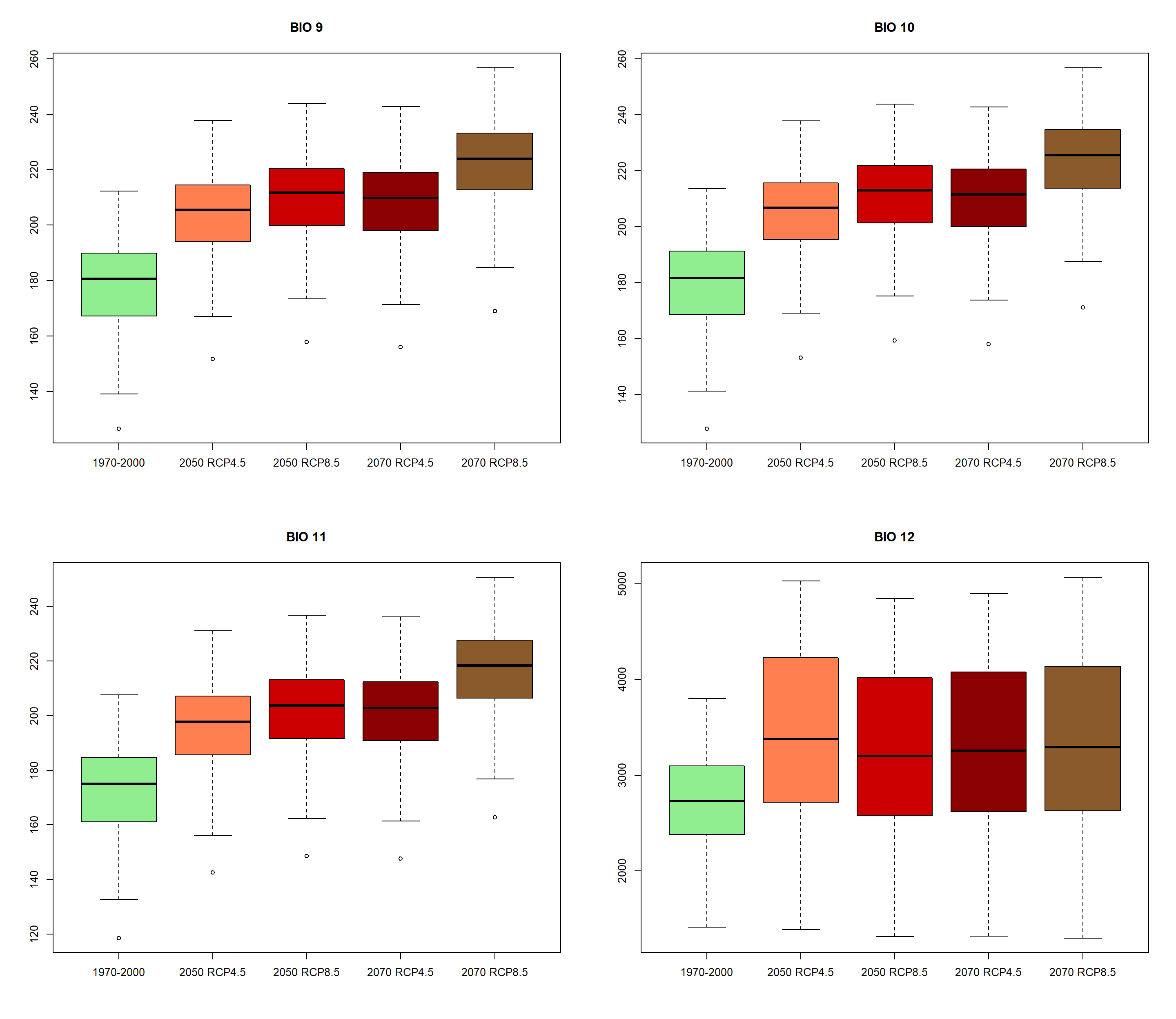

### visualization-11-4.png

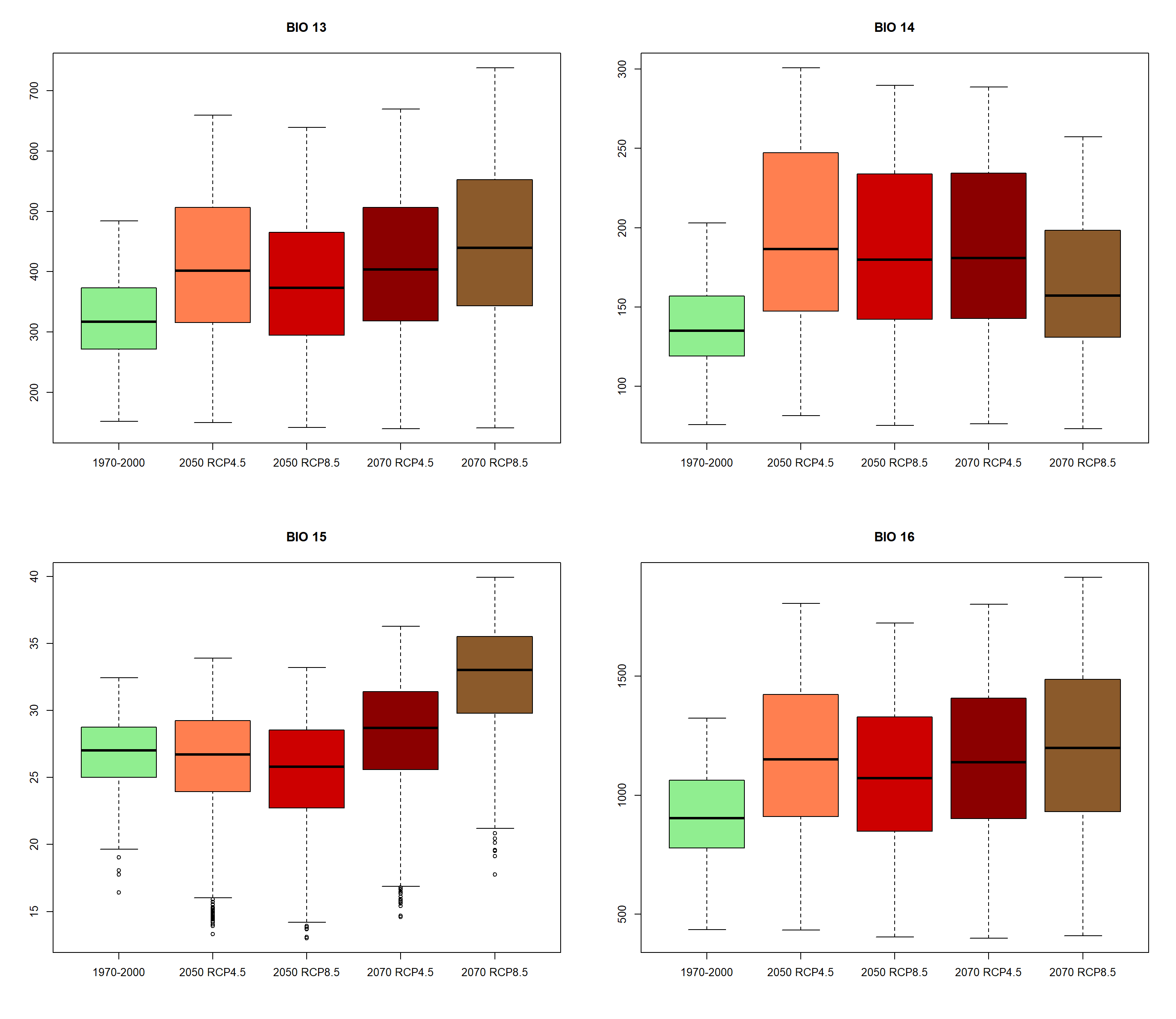

### visualization-11-5.png

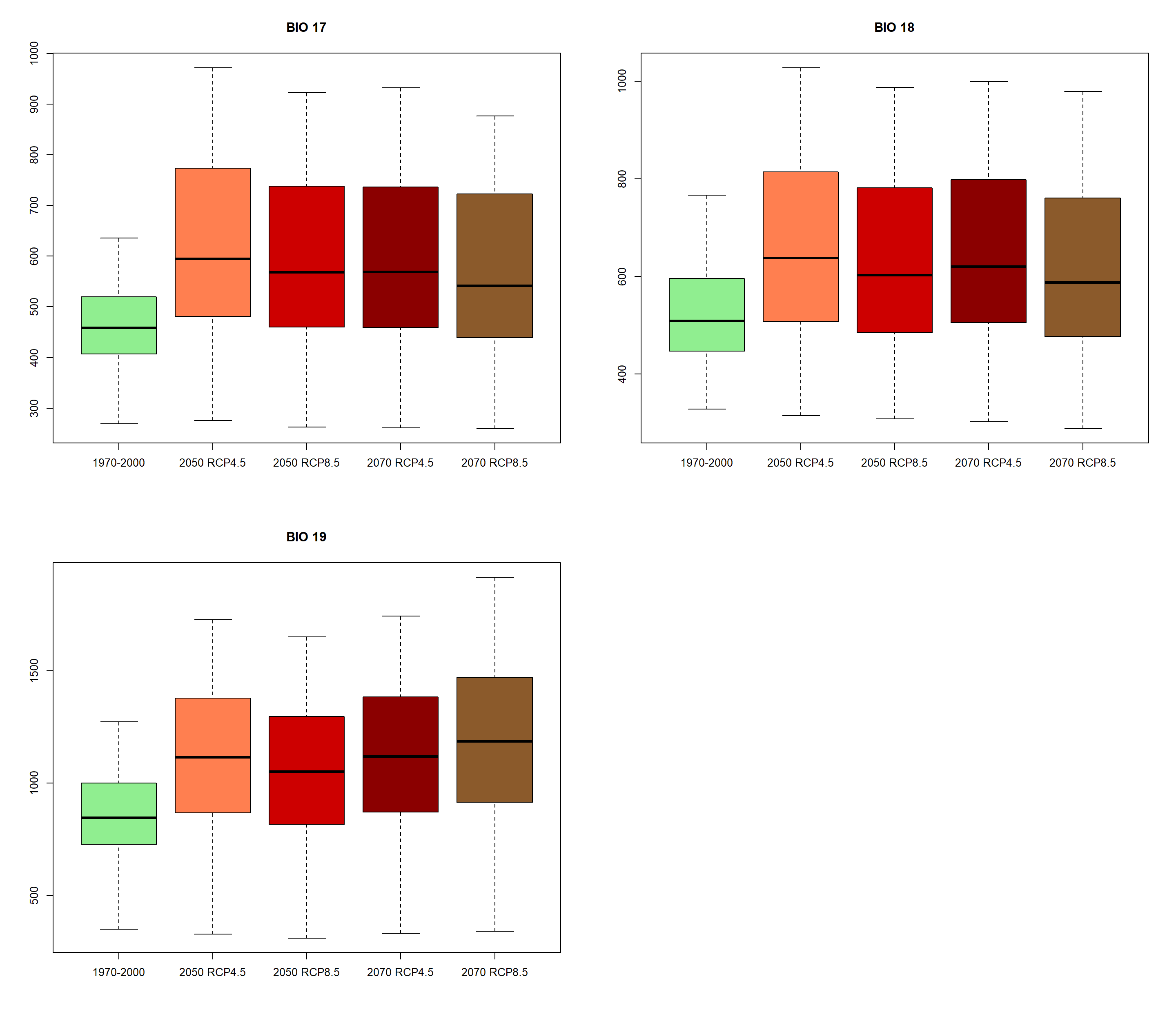
