## Supplementary material for "Vulnerability to climate change for narrowly ranged species: the case of Ecuadorian endemic *Magnolia mercedesiarum*": spatial_autocorrelation_test.html

Auxiliar script for Vazquez et al. 2018


### Script for Shalisko et al. 2018 - Spatial autocorrelation test

###### *Viacheslav Shalisko*

###### *22 of october 2017*

### *Magnolia mercedesiarum* spatial autocorrelation in predictor variables - spatial correlation index

###### Load modules

```
library(maptools)
```

```
## Loading required package: sp
```

```
## Checking rgeos availability: FALSE
##      Note: when rgeos is not available, polygon geometry     computations in maptools depend on gpclib,
##      which has a restricted licence. It is disabled by default;
##      to enable gpclib, type gpclibPermit()
```

```
library(rworldmap)    # worldmap datasets
```

```
## Warning: package 'rworldmap' was built under R version 3.4.2
```

```
## ### Welcome to rworldmap ###
```

```
## For a short introduction type :   vignette('rworldmap')
```

```
library(rworldxtra)   # hires worldmap spatial dataframe
```

```
## Warning: package 'rworldxtra' was built under R version 3.4.2
```

```
library(sp)
library(raster)
library(dismo)
library(rgdal)
```

```
## rgdal: version: 1.2-11, (SVN revision 676)
##  Geospatial Data Abstraction Library extensions to R successfully loaded
##  Loaded GDAL runtime: GDAL 2.2.0, released 2017/04/28
##  Path to GDAL shared files: C:/Users/vshal/Documents/R/win-library/3.4/rgdal/gdal
##  Loaded PROJ.4 runtime: Rel. 4.9.3, 15 August 2016, [PJ_VERSION: 493]
##  Path to PROJ.4 shared files: C:/Users/vshal/Documents/R/win-library/3.4/rgdal/proj
##  Linking to sp version: 1.2-5
```

```
library(rJava)
library(ade4)
```

```
## Warning: package 'ade4' was built under R version 3.4.4
```

```
library(spdep)
```

```
## Warning: package 'spdep' was built under R version 3.4.4
```

```
## Loading required package: Matrix
```

```
## Loading required package: spData
```

```
## Warning: package 'spData' was built under R version 3.4.4
```

```
## To access larger datasets in this package, install the spDataLarge
## package with: `install.packages('spDataLarge',
## repos='https://nowosad.github.io/drat/', type='source'))`
```

```
## 
## Attaching package: 'spdep'
```

```
## The following object is masked from 'package:ade4':
## 
##     mstree
```

```
library(deldir)
```

```
## Warning: package 'deldir' was built under R version 3.4.4
```

```
## deldir 0.1-15
```

#### Data preparation

###### Set adjustable variables

```
run_code <- '29'
basepath <- 'C:/Users/vshal/Downloads/M_mercedesiarum_rasters/run29_climond2'
basename <- 'Magnolia_mercedesiarum_MAXENT_climond2_HR_'

scriptpath <- 'C:/Users/vshal/Downloads/MaxEnt_test_runs/run29_climond2_R'

random_background_number <- 450
presence_points_count <- 50
double_presence_points_count <- 100

#cross_validation_lenght <- 10
cross_validation_lenght <- 10

#layers_to_drop <- c(2,6,7,9,11,12,14,15,17,18,19,21)  # unsignificant layers
layers_to_drop <- c(21)    # the last layer no. 21 is dummy (repetition of the first, included just to drop it when nothing to drop)

# load raster of predicted presence

presence_raster_mask <- stack("C:/Users/vshal/GD/Projects_actual/bot_Magnolia_mercedesiarum/Phytotaxa_2017/Analysis/SIG/Rasters_run_22_23/Magnolia_mercedesiarum_ETSS_raster_small.tif")
```

###### Load model run results

```
iterations <- readRDS(paste(scriptpath, "/iterations_object_",run_code,".rds",sep=""))

#str(iterations)
```

##### Spatial autocorrelation index

```
iterations_spatial_test <- list()
spatil_autocorr_index <- c()

for (j in 1:cross_validation_lenght) {
  my_obj <- iterations[[j]]
  
    coords <- my_obj$entr
    predictors <- my_obj$model@presence

    container <- list()
    container$xy <- coords
    container$predictors <- predictors
    #container$predictors <- container$predictors[,c(1,4,6,7,11,13,16,19,20)]
     

    provi <- deldir::deldir(container$xy[,1],container$xy[,2])
    provi.neig <- neig(edges = as.matrix(provi$delsgs[, 5:6]))
    #print(provi.neig)

    magnolia.listw <- nb2listw(neig2nb(provi.neig))
    #magnolia.listw
        
    magnolia.pca <- dudi.pca(container$predictors, scannf = FALSE)
    #summary(magnolia.pca)
    container$magnolia.pca <- magnolia.pca
    
    magnolia.rtest <- multispati.rtest(magnolia.pca, magnolia.listw)
    spatil_autocorr_index <- c(spatil_autocorr_index, magnolia.rtest$obs)
    
    print(magnolia.rtest)
    #str(magnolia.rtest)
    #summary(magnolia.rtest)
    
    magnolia.pca.ms <- multispati(magnolia.pca, magnolia.listw, scannf=FALSE)
    container$magnolia.pca.ms <- magnolia.pca.ms
    #summary(magnolia.pca.ms)
    #plot(magnolia.pca.ms)
    
    iterations_spatial_test[[j]] <- container 
  
  
}
```

```
## Monte-Carlo test
## Call: multispati.rtest(dudi = magnolia.pca, listw = magnolia.listw)
## 
## Observation: 0.4993246 
## 
## Based on 99 replicates
## Simulated p-value: 0.01 
## Alternative hypothesis: greater 
## 
##      Std.Obs  Expectation     Variance 
##  9.567880939 -0.011323002  0.002848466 
## Monte-Carlo test
## Call: multispati.rtest(dudi = magnolia.pca, listw = magnolia.listw)
## 
## Observation: 0.5322096 
## 
## Based on 99 replicates
## Simulated p-value: 0.01 
## Alternative hypothesis: greater 
## 
##      Std.Obs  Expectation     Variance 
##  9.302391250 -0.019497120  0.003517445 
## Monte-Carlo test
## Call: multispati.rtest(dudi = magnolia.pca, listw = magnolia.listw)
## 
## Observation: 0.6342862 
## 
## Based on 99 replicates
## Simulated p-value: 0.01 
## Alternative hypothesis: greater 
## 
##      Std.Obs  Expectation     Variance 
## 11.311402394 -0.008309009  0.003227320 
## Monte-Carlo test
## Call: multispati.rtest(dudi = magnolia.pca, listw = magnolia.listw)
## 
## Observation: 0.578446 
## 
## Based on 99 replicates
## Simulated p-value: 0.01 
## Alternative hypothesis: greater 
## 
##     Std.Obs Expectation    Variance 
## 10.78409641 -0.02109552  0.00309080 
## Monte-Carlo test
## Call: multispati.rtest(dudi = magnolia.pca, listw = magnolia.listw)
## 
## Observation: 0.5731522 
## 
## Based on 99 replicates
## Simulated p-value: 0.01 
## Alternative hypothesis: greater 
## 
##       Std.Obs   Expectation      Variance 
##  8.7883539473 -0.0006980344  0.0042636543 
## Monte-Carlo test
## Call: multispati.rtest(dudi = magnolia.pca, listw = magnolia.listw)
## 
## Observation: 0.6493364 
## 
## Based on 99 replicates
## Simulated p-value: 0.01 
## Alternative hypothesis: greater 
## 
##      Std.Obs  Expectation     Variance 
## 10.254332593 -0.015734576  0.004206503 
## Monte-Carlo test
## Call: multispati.rtest(dudi = magnolia.pca, listw = magnolia.listw)
## 
## Observation: 0.5737399 
## 
## Based on 99 replicates
## Simulated p-value: 0.01 
## Alternative hypothesis: greater 
## 
##      Std.Obs  Expectation     Variance 
## 10.195899526 -0.016297456  0.003348945 
## Monte-Carlo test
## Call: multispati.rtest(dudi = magnolia.pca, listw = magnolia.listw)
## 
## Observation: 0.6019741 
## 
## Based on 99 replicates
## Simulated p-value: 0.01 
## Alternative hypothesis: greater 
## 
##      Std.Obs  Expectation     Variance 
## 10.417258586 -0.030337472  0.003684303 
## Monte-Carlo test
## Call: multispati.rtest(dudi = magnolia.pca, listw = magnolia.listw)
## 
## Observation: 0.6405521 
## 
## Based on 99 replicates
## Simulated p-value: 0.01 
## Alternative hypothesis: greater 
## 
##      Std.Obs  Expectation     Variance 
## 10.492348767 -0.015757424  0.003912658 
## Monte-Carlo test
## Call: multispati.rtest(dudi = magnolia.pca, listw = magnolia.listw)
## 
## Observation: 0.5309663 
## 
## Based on 99 replicates
## Simulated p-value: 0.01 
## Alternative hypothesis: greater 
## 
##      Std.Obs  Expectation     Variance 
##  9.594472095 -0.020036223  0.003298109
```

```
for (j in 1:cross_validation_lenght) {
  cat("\n\n Iteration ",j,"\n")
  cat("Spatial autocorrelation index R = ",spatil_autocorr_index[j],"\n")
  my_obj <- iterations_spatial_test[[j]]
  plot(my_obj$magnolia.pca.ms)
}
```

```
## 
## 
##  Iteration  1 
## Spatial autocorrelation index R =  0.4993246
```

```
## 
## 
##  Iteration  2 
## Spatial autocorrelation index R =  0.5322096
```

```
## 
## 
##  Iteration  3 
## Spatial autocorrelation index R =  0.6342862
```

```
## 
## 
##  Iteration  4 
## Spatial autocorrelation index R =  0.578446
```

```
## 
## 
##  Iteration  5 
## Spatial autocorrelation index R =  0.5731522
```

```
## 
## 
##  Iteration  6 
## Spatial autocorrelation index R =  0.6493364
```

```
## 
## 
##  Iteration  7 
## Spatial autocorrelation index R =  0.5737399
```

```
## 
## 
##  Iteration  8 
## Spatial autocorrelation index R =  0.6019741
```

```
## 
## 
##  Iteration  9 
## Spatial autocorrelation index R =  0.6405521
```

```
## 
## 
##  Iteration  10 
## Spatial autocorrelation index R =  0.5309663
```

```
spatil_autocorr_index
```

```
##  [1] 0.4993246 0.5322096 0.6342862 0.5784460 0.5731522 0.6493364 0.5737399
##  [8] 0.6019741 0.6405521 0.5309663
```
