## Supplementary material for "Vulnerability to climate change for narrowly ranged species: the case of Ecuadorian endemic *Magnolia mercedesiarum*": spatial_autocorrelation_test.Rmd

````
---
title: "Auxiliar script for Vazquez et al. 2018"
author: "Viacheslav Shalisko"
date: "March 1, 2018"
output: html_document
---

##---------------------------------------------------------------------------

This program is free software; you can redistribute it and/or modify
it under the terms of the GNU General Public License as published by
the Free Software Foundation; either version 2 of the License, or
(at your option) any later version.  

This program is distributed in the hope that it will be useful,
but WITHOUT ANY WARRANTY; without even the implied warranty of
MERCHANTABILITY or FITNESS FOR A PARTICULAR PURPOSE. See the
GNU General Public License for more details.  

##---------------------------------------------------------------------------

*Magnolia mercedesiarum* climate change: spatial autocorrelation
========================================================


#### Load modules

```{r load-modulos}
library(maptools)
library(rworldmap)    # worldmap datasets
library(rworldxtra)   # hires worldmap spatial dataframe

library(sp)
library(raster)
library(dismo)
library(rgdal)
library(rJava)
library(ade4)
library(spdep)
library(deldir)

```

```{r}
knitr::opts_chunk$set(echo = TRUE)
knitr::opts_chunk$set(error = TRUE)

evaluate.mdl <- function (pnt,bkg,rast,mdl) {
  # pnt - presence points, bkg - background points, rast - raster dataset, mdl - model
  result <- list()
  values_bkg <- extract(rast, bkg)
  values_pnt <- extract(rast, pnt)
  result$v_bkg <- values_bkg
  result$v_pnt <- values_pnt  
  
  values_pnt_bkg <- rbind(values_bkg, values_pnt)

  result$e <- evaluate(values_pnt, values_pnt_bkg, mdl)
  #print(result$e)

  par(mfrow=c(2, 2))
  plot(result$e, 'TPR')
  plot(result$e, 'ROC')
  density(result$e)
  boxplot(result$e, col=c('lightblue','coral'), notch=TRUE)
  
  result$t <- threshold(result$e)
  print(result$t)
  
  # use equal specificity & sensitivity threshold
  result$et <- evaluate(values_pnt, values_pnt_bkg, mdl, tr = result$t$equal_sens_spec)
  print(result$et)

  return(result)
}

maxent.select.contribution <- function (m,n) {
  # parametros: m - objeto del model maxent, n - vector de nombres de variables raster
  resultados <- m@results
  tabla_resultados <- data.frame(t(rep(NA,3)))
  names(tabla_resultados) <- c('variable','contribution','permutation.importance')
  for (i in 1:length(n)) {
    variable1_name <- paste(n[i],'.contribution',sep='')
    variable1_value <- resultados[variable1_name,1]
    variable2_name <- paste(n[i],'.permutation.importance',sep='')
    variable2_value <- resultados[variable2_name,1]
    tabla_resultados <- rbind(tabla_resultados,c(n[i],variable1_value,variable2_value))
  }
  tabla_resultados <- tabla_resultados[-1,]
  return(tabla_resultados)
}

maxent.select.AUC <- function (m) {
  resultados <- m@results
  return(resultados['Training.AUC',1])
}

```

## Data preparation
#### Set adjustable variables

```{r set-variables}
run_code <- '29'
basepath <- 'C:/Users/vshal/Downloads/M_mercedesiarum_rasters/run29_climond2'
basename <- 'Magnolia_mercedesiarum_MAXENT_climond2_HR_'

scriptpath <- 'C:/Users/vshal/Downloads/MaxEnt_test_runs/run29_climond2_R'

random_background_number <- 450
presence_points_count <- 50
double_presence_points_count <- 100

#cross_validation_lenght <- 10
cross_validation_lenght <- 10

#layers_to_drop <- c(2,6,7,9,11,12,14,15,17,18,19,21)  # unsignificant layers
layers_to_drop <- c(21)    # the last layer no. 21 is dummy (repetition of the first, included just to drop it when nothing to drop)

# load raster of predicted presence

presence_raster_mask <- stack("C:/Users/vshal/GD/Projects_actual/bot_Magnolia_mercedesiarum/Phytotaxa_2017/Analysis/SIG/Rasters_run_22_23/Magnolia_mercedesiarum_ETSS_raster_small.tif")
```

#### Load raster predictor variables

```{r load-rasters}

names <- c("alt","bio1","bio2","bio3","bio4",
              "bio5","bio6","bio7","bio8","bio9",
              "bio10","bio11","bio12","bio13","bio14",
              "bio15","bio16","bio17","bio18","bio19",
              "dummy")

capas_raster<-stack(c("C:/Users/vshal/Downloads/GMTED2010/GMTED2010_075_Andes.tif",
                      "C:/Users/vshal/Downloads/CliMond/biovars_climond2_Andes_HR.tif",
                      "C:/Users/vshal/Downloads/GMTED2010/GMTED2010_075_Andes.tif"))

names(capas_raster) <- names

capas_raster

#summary(capas_raster)

bio10 <- subset(capas_raster,10)
```

#### Drop unsignificant raster layers (based on preliminary test run)

```{r drop-layers}
# drop unsignificant or dummy layers
capas_raster_drop <- dropLayer(capas_raster,layers_to_drop)
names_after_drop <- names(capas_raster_drop)
```

#### Load model run results

```{r cross-validation-load}
iterations <- readRDS(paste(scriptpath, "/iterations_object_",run_code,".rds",sep=""))

#str(iterations)
```

### Spatial autocorrelation index

```{r fig.width = 12, fig.height = 10, message=FALSE, warning=FALSE}
iterations_spatial_test <- list()
spatil_autocorr_index <- c()

for (j in 1:cross_validation_lenght) {
  my_obj <- iterations[[j]]
  
    coords <- my_obj$entr
    predictors <- my_obj$model@presence

    container <- list()
    container$xy <- coords
    container$predictors <- predictors
    #container$predictors <- container$predictors[,c(1,4,6,7,11,13,16,19,20)]
     

    provi <- deldir::deldir(container$xy[,1],container$xy[,2])
    provi.neig <- neig(edges = as.matrix(provi$delsgs[, 5:6]))
    #print(provi.neig)

    magnolia.listw <- nb2listw(neig2nb(provi.neig))
    #magnolia.listw
        
    magnolia.pca <- dudi.pca(container$predictors, scannf = FALSE)
    #summary(magnolia.pca)
    container$magnolia.pca <- magnolia.pca
    
    magnolia.rtest <- multispati.rtest(magnolia.pca, magnolia.listw)
    spatil_autocorr_index <- c(spatil_autocorr_index, magnolia.rtest$obs)
    
    print(magnolia.rtest)
    #str(magnolia.rtest)
    #summary(magnolia.rtest)
    
    magnolia.pca.ms <- multispati(magnolia.pca, magnolia.listw, scannf=FALSE)
    container$magnolia.pca.ms <- magnolia.pca.ms
    #summary(magnolia.pca.ms)
    #plot(magnolia.pca.ms)
    
    iterations_spatial_test[[j]] <- container 
  
  
}

for (j in 1:cross_validation_lenght) {
  cat("\n\n Iteration ",j,"\n")
  cat("Spatial autocorrelation index R = ",spatil_autocorr_index[j],"\n")
  my_obj <- iterations_spatial_test[[j]]
  plot(my_obj$magnolia.pca.ms)
}

spatil_autocorr_index
```

## Example

```{r}


#    data(mafragh)
#    #mafragh
#    maf.listw <- nb2listw(neig2nb(mafragh$neig))
#    #maf.listw
    
#    maf.pca <- dudi.pca(mafragh$flo, scannf = FALSE)
#    multispati.rtest(maf.pca, maf.listw)
#    maf.pca.ms <- multispati(maf.pca, maf.listw, scannf=FALSE)
#    summary(maf.pca.ms)
#    plot(maf.pca.ms)

```
````
