## Supplementary material for "Vulnerability to climate change for narrowly ranged species: the case of Ecuadorian endemic *Magnolia mercedesiarum*": Downscaling_tmin_01_climond.Rmd

````
---
title: "Downscaling T"
author: "Viacheslav Shalisko"
date: "27 of November 2017"
output: html_document
---

```{r setup, include=FALSE}
knitr::opts_chunk$set(echo = TRUE)
knitr::opts_chunk$set(error = TRUE)

library(maptools)
library(rworldmap)    # worldmap datasets
library(rworldxtra)   # hires worldmap spatial dataframe

library(sp)
library(raster)
```

```{r}
  sampleClasses <- function(r = raster, n = 8) {
   # function gets a raster object and a sample size. It samples n cells from each class
   # in the raster and return a vector with the cells indices
   vals <- unique(getValues(r)) # Get all classes
   vals <- vals[!is.na(vals)]   # Exclude NAs
   for (val in vals) {
    cellVal <- which(t(as.matrix(r)) == val) # get All cells for each class
    #print(cellVal)
    if(!exists("samples")) {
      samples <- sample(cellVal, n) # sample class locations
    } else {
      samples <- c(samples, sample(cellVal, n))
    }
   }
   return(samples)
  }
```

# Variables

```{r set-variables}
# sample size within each altitude class
sample_size <- 50

basepath <- 'C:/Users/vshal/Downscaling'
basename <- 'tmin_01_a100m_current'

# original lowresolution temperature raster
temp_name_LR <- "C:/Users/vshal/Downscaling/tmin_01_a1km.tif"
# high resolution DEM
mde_name_HR <- "C:/Users/vshal/Downscaling/DEM_HR_GEO.tif"
# low resolution DEM
mde_name_LR <- "C:/Users/vshal/Downscaling/DEM_LR_GEO.tif"
```

## Load raster data

```{r load-raster}
# load LR raster
temp_LR <- stack(temp_name_LR)
# load LR mde
mde_LR <- stack(mde_name_LR)
# load HR mde
mde_HR <- stack(mde_name_HR)
```

## Check raster data

```{r verify-raster, fig.width = 10, fig.height = 4}
temp_LR
mde_LR
mde_HR

par(mfrow = c(1,3))

plot(temp_LR, legend = FALSE, main = "Temp")
plot(mde_LR, legend = FALSE, main = "MDE LR")
plot(mde_HR, legend = FALSE, main = "MDE HR")

```

# Produce sampling points

```{r sampling-design, fig.width = 10, fig.height = 8}

# relassify raster to altitude classes and sample N points from each altitude class
# set reclass table
reclass_table <- c(-20, 500, 1,  500, 1000, 2,  1000, 1500, 3, 1500, 2000, 4, 2000, 2500, 5, 2500, 3000, 6, 3000, 5000, 7)
reclass_table <- matrix(reclass_table, ncol=3, byrow=TRUE)

# reclassify
mde_LR_reclass <- reclassify(mde_LR, reclass_table, include.lowest=FALSE, right=TRUE)

# transform raster to categorical variable
mde_LR_reclass <- ratify(mde_LR_reclass)

# produce index of pixels to sample
sampling_points_index <- sampleClasses(r = mde_LR_reclass, n = sample_size)

# get XY coordinates of pixels to sample
sampling_points <- xyFromCell(mde_LR_reclass, sampling_points_index, spatial=TRUE)

# visualiza classified raster and points
plot(mde_LR_reclass)
points(sampling_points)
```

# Sampling

```{r sampling, fig.width = 10, fig.height = 4}
# sample source raster values with pixel index
sampling_points_mde <- mde_LR[sampling_points_index]
attr(sampling_points_mde, "dimnames") <- list(NULL, "mde")
#str(sampling_points_mde)

sampling_points$mde <- sampling_points_mde

# sample raster with same pixel parameters as source with pixel index
sampling_points_temp <- temp_LR[sampling_points_index]
attr(sampling_points_temp,"dimnames") <- list(NULL, "temp")
#str(sampling_points_temp)

sampling_points$temp <- sampling_points_temp

sampling_points
#str(sampling_points)

par(mfrow = c(1,2))  
hist(sampling_points$mde, col = "lightgray")
hist(sampling_points$temp, col = "lightgray")

```

# Generate model fr relation between predictor (elevation) and dependent variable (temperature)

```{r produce-model, fig.width = 10, fig.height = 8}

model_mde <- sampling_points$mde
model_temp <- sampling_points$temp

regression <- glm(model_temp ~ model_mde)

regression
plot(regression)

sequence_mde <- data.frame(model_mde = seq(from = 0, to = 4500, by = 100))

regression_predict <- predict(regression, sequence_mde, se.fit = TRUE)

plot(sampling_points$mde, sampling_points$temp, xlim = c(0, 4500), ylim = c(-5,40))

polygon(c(sequence_mde$model_mde,rev(sequence_mde$model_mde)),
        c(regression_predict$fit - 1.96*regression_predict$se.fit, 
        rev(regression_predict$fit + 1.96*regression_predict$se.fit)),
        col = "lightgray", border = NA, density = 20)
lines(sequence_mde$model_mde , regression_predict$fit + 1.96*regression_predict$se.fit, lwd = 2 , col = "lightgray")
lines(sequence_mde$model_mde , regression_predict$fit - 1.96*regression_predict$se.fit, lwd = 2 , col = "lightgray")
lines(sequence_mde$model_mde , regression_predict$fit, lwd = 3 , col = "red")
```

## Estimate temperature with model at high resolution

```{r temp-predict, fig.width = 10, fig.height = 8}
names(mde_HR) <- c("model_mde")

predicted_surface <- predict(mde_HR, regression, se.fit = TRUE)

plot(predicted_surface)
```

# Produce low resolution pixel index

```{r produce-pixel-index, fig.width = 10, fig.height = 8}
# produce pixel index with same dimensions as source LR raster
lr_ext <- extent(mde_LR)
lr_crs <- crs(mde_LR)
index_raster <- raster(nrows = dim(mde_LR)[1], ncols = dim(mde_LR)[2], ext = lr_ext)
crs(index_raster) <- lr_crs
index_raster[] <- seq(from = 1, to = dim(mde_LR)[1] * dim(mde_LR)[2], by = 1)

# visualize index
index_raster
plot(index_raster)
```

## Change resolution from low to high

```{r resample-pixel-index}
resampled_index_raster <- resample(index_raster,predicted_surface,method='ngb')

# compare extent and pixel count for index and prediction rasters
resampled_index_raster
predicted_surface
```

# Calculate mean for each low resolution pixel

```{r zonal-means}
zonal_stats <- zonal(predicted_surface, resampled_index_raster, 'mean')
dim(zonal_stats)
```

## Transfom xonal sampling to low resolution raster

```{r produce-zonal-raster, fig.width = 10, fig.height = 8}
# produce zonal stats raster with same dimensions as source LR raster
zonal_stats_raster <- raster(nrows = dim(mde_LR)[1], ncols = dim(mde_LR)[2], ext = lr_ext)
crs(zonal_stats_raster) <- lr_crs
zonal_stats_raster[] <- zonal_stats[,2]

# visualize zonal stats
zonal_stats_raster
plot(zonal_stats_raster)
```

# Calculate difference between two low resolution resolution rasters

```{r map-algebra-lr}
mean_differences_raster <- temp_LR - zonal_stats_raster
```

## Resample to high resolution

```{r resample-zonal-raster}
mean_differences_raster_HR1 <- resample(mean_differences_raster,predicted_surface,method='ngb')

# compare extent and pixel count for differences and prediction rasters
mean_differences_raster_HR1
predicted_surface
```

# Apply Gauss filter

```{r focal-statistics}
gf <- focalWeight(mean_differences_raster_HR1, 0.0083333333, "Gauss")
mean_differences_raster_HR <- focal(mean_differences_raster_HR1, w=gf)   # Gauss filter
#mean_differences_raster_HR <- focal(mean_differences_raster_HR1, w=matrix(1/49,nrow=7,ncol=7))   # 7x7 square mask equal weight
```

## View raster after filter

```{r view-differences-raster, fig.width = 10, fig.height = 8}
# visualize differences raster
mean_differences_raster_HR
plot(mean_differences_raster_HR)
```

# Merge adjuste raster with model prediction

```{r map-algebra-hr, fig.width = 10, fig.height = 8}
corrected_raster <- predicted_surface + mean_differences_raster_HR

# visualize corrected raster
corrected_raster
plot(corrected_raster)
```

# Save as GeoTiff

```{r save-downscaled-raster}
# write GeoTIFF with float 32 and LZW compression
  writeRaster(corrected_raster, filename=paste(basepath, '/' ,basename ,'.tif',sep=''), 
           format="GTiff", overwrite=TRUE, datatype = 'FLT4S', options=c("COMPRESS=LZW"))
```
````
