## Supplementary material for "Vulnerability to climate change for narrowly ranged species: the case of Ecuadorian endemic *Magnolia mercedesiarum*": M_mercedesiarum_climate_change_with_U.html

Auxiliar script for Shalisko et al. 2018


### Script for Shalisko et al. 2018 - Comparison of current and future conditions

###### *Viacheslav Shalisko*

###### *15 of november 2017*

### *Magnolia mercedesiarum* climate change: comparison of actual and future conditions

##### Current climate conditions WorldClim ver. 2 (1970-2000)

##### Predictive climatic model HadGEM2 - ES, scenarios RCP45 and RCP85 in 2050 and 2070

```
knitr::opts_chunk$set(echo = TRUE)
knitr::opts_chunk$set(error = TRUE)

library(maptools)
```

```
## Loading required package: sp
```

```
## Checking rgeos availability: FALSE
##      Note: when rgeos is not available, polygon geometry     computations in maptools depend on gpclib,
##      which has a restricted licence. It is disabled by default;
##      to enable gpclib, type gpclibPermit()
```

```
library(rworldmap)    # worldmap datasets
```

```
## Warning: package 'rworldmap' was built under R version 3.4.2
```

```
## ### Welcome to rworldmap ###
```

```
## For a short introduction type :   vignette('rworldmap')
```

```
library(rworldxtra)   # hires worldmap spatial dataframe
```

```
## Warning: package 'rworldxtra' was built under R version 3.4.2
```

```
library(sp)
library(raster)
library(dismo)
library(rgdal)
```

```
## rgdal: version: 1.2-11, (SVN revision 676)
##  Geospatial Data Abstraction Library extensions to R successfully loaded
##  Loaded GDAL runtime: GDAL 2.2.0, released 2017/04/28
##  Path to GDAL shared files: C:/Users/vshal/Documents/R/win-library/3.4/rgdal/gdal
##  Loaded PROJ.4 runtime: Rel. 4.9.3, 15 August 2016, [PJ_VERSION: 493]
##  Path to PROJ.4 shared files: C:/Users/vshal/Documents/R/win-library/3.4/rgdal/proj
##  Linking to sp version: 1.2-5
```

```
compareMWT = function(x, y, test_title, alternative='two.sided') {
  wilcoxTest_result <- wilcox.test(x,y,alternative=alternative,conf.int=TRUE)
  cat('Mann-Whitney test results for',test_title,"\n",sep=' ')
  if (wilcoxTest_result$p.value < 0.05) {
      cat('The H0 is rejected as p < 0.05, the HA accepted (medians unequal)',"\n")
  } else {
      cat('The HA is rejected as p >= 0.05, the H0 accepted (medians equal)',"\n")
  }  
  print(wilcoxTest_result)
}
```

```
presence_points_count <- 1000
month_labels <- c('Jan','Feb','Mar','Apr','May','Jun','Jul','Aug','Sep','Oct','Nov','Dic')
biovars_names <- c("bio1","bio2","bio3","bio4",
              "bio5","bio6","bio7","bio8","bio9",
              "bio10","bio11","bio12","bio13","bio14",
              "bio15","bio16","bio17","bio18","bio19")
month_list <- c('Jan','Feb','Mar','Apr','May','Jun','Jul','Aug','Sep','Oct','Nov','Dic')
```

```
# load raster of predicted presence
presence_raster_mask <- stack("C:/Users/vshal/OneDrive/Documentos_para_GD/Temp_Magnolia/Magnolia_mercedesiarum_ETSS_raster_small.tif")

mde <- stack("C:/Users/vshal/Downloads/GMTED2010/GMTED2010_075_Andes.tif")

# load actual conditions
my_path0 <- 'C:\\Users\\vshal\\Downloads\\CliMond\\'

prec00 <- stack(paste(my_path0,"wc2.0_HR_prec_stack.tif",sep=""))
tmin00 <- stack(paste(my_path0,"wc2.0_HR_tmin_stack.tif",sep=""))
tmax00 <- stack(paste(my_path0,"wc2.0_HR_tmax_stack.tif",sep=""))

biovars00 <- stack(paste(my_path0,"biovars_climond2_Andes_HR.tif",sep=""))
names(biovars00) <- biovars_names

# load future conditions
my_path1 <- 'C:\\Users\\vshal\\Downloads\\future_conditions\\'

prec50 <- stack(paste(my_path1,"he45pr50_stack.tif",sep=""))
tmin50 <- stack(paste(my_path1,"he45tn50_stack.tif",sep=""))
tmax50 <- stack(paste(my_path1,"he45tx50_stack.tif",sep=""))

biovars50 <- stack(paste(my_path1,"biovars_he45_50_Andes_HR.tif",sep=""))
names(biovars50) <- biovars_names

prec50A <- stack(paste(my_path1,"he85pr50_stack.tif",sep=""))
tmin50A <- stack(paste(my_path1,"he85tn50_stack.tif",sep=""))
tmax50A <- stack(paste(my_path1,"he85tx50_stack.tif",sep=""))

biovars50A <- stack(paste(my_path1,"biovars_he85_50_Andes_HR.tif",sep=""))
names(biovars50A) <- biovars_names

prec70 <- stack(paste(my_path1,"he45pr70_stack.tif",sep=""))
tmin70 <- stack(paste(my_path1,"he45tn70_stack.tif",sep=""))
tmax70 <- stack(paste(my_path1,"he45tx70_stack.tif",sep=""))

biovars70 <- stack(paste(my_path1,"biovars_he45_70_Andes_HR.tif",sep=""))
names(biovars70) <- biovars_names

prec70A <- stack(paste(my_path1,"he85pr70_stack.tif",sep=""))
tmin70A <- stack(paste(my_path1,"he85tn70_stack.tif",sep=""))
tmax70A <- stack(paste(my_path1,"he85tx70_stack.tif",sep=""))

biovars70A <- stack(paste(my_path1,"biovars_he85_70_Andes_HR.tif",sep=""))
names(biovars70A) <- biovars_names

# predictions
my_path2 <- 'C:/Users/vshal/Downloads/M_mercedesiarum_rasters/run29_climond2/'
prediction00_1 <- stack(paste(my_path2,"Magnolia_mercedesiarum_MAXENT_climond2_HR_run_29_1.tif",sep=""))
prediction00_2 <- stack(paste(my_path2,"Magnolia_mercedesiarum_MAXENT_climond2_HR_run_29_2.tif",sep=""))
prediction00_3 <- stack(paste(my_path2,"Magnolia_mercedesiarum_MAXENT_climond2_HR_run_29_3.tif",sep=""))
prediction00_4 <- stack(paste(my_path2,"Magnolia_mercedesiarum_MAXENT_climond2_HR_run_29_4.tif",sep=""))
prediction00_5 <- stack(paste(my_path2,"Magnolia_mercedesiarum_MAXENT_climond2_HR_run_29_5.tif",sep=""))
prediction00_6 <- stack(paste(my_path2,"Magnolia_mercedesiarum_MAXENT_climond2_HR_run_29_6.tif",sep=""))
prediction00_7 <- stack(paste(my_path2,"Magnolia_mercedesiarum_MAXENT_climond2_HR_run_29_7.tif",sep=""))
prediction00_8 <- stack(paste(my_path2,"Magnolia_mercedesiarum_MAXENT_climond2_HR_run_29_8.tif",sep=""))
prediction00_9 <- stack(paste(my_path2,"Magnolia_mercedesiarum_MAXENT_climond2_HR_run_29_9.tif",sep=""))
prediction00_10 <- stack(paste(my_path2,"Magnolia_mercedesiarum_MAXENT_climond2_HR_run_29_10.tif",sep=""))

prediction50_1 <- stack(paste(my_path2,"Magnolia_mercedesiarum_MAXENT_he45_50_HR_run_29_1.tif",sep=""))
prediction50_2 <- stack(paste(my_path2,"Magnolia_mercedesiarum_MAXENT_he45_50_HR_run_29_2.tif",sep=""))
prediction50_3 <- stack(paste(my_path2,"Magnolia_mercedesiarum_MAXENT_he45_50_HR_run_29_3.tif",sep=""))
prediction50_4 <- stack(paste(my_path2,"Magnolia_mercedesiarum_MAXENT_he45_50_HR_run_29_4.tif",sep=""))
prediction50_5 <- stack(paste(my_path2,"Magnolia_mercedesiarum_MAXENT_he45_50_HR_run_29_5.tif",sep=""))
prediction50_6 <- stack(paste(my_path2,"Magnolia_mercedesiarum_MAXENT_he45_50_HR_run_29_6.tif",sep=""))
prediction50_7 <- stack(paste(my_path2,"Magnolia_mercedesiarum_MAXENT_he45_50_HR_run_29_7.tif",sep=""))
prediction50_8 <- stack(paste(my_path2,"Magnolia_mercedesiarum_MAXENT_he45_50_HR_run_29_8.tif",sep=""))
prediction50_9 <- stack(paste(my_path2,"Magnolia_mercedesiarum_MAXENT_he45_50_HR_run_29_9.tif",sep=""))
prediction50_10 <- stack(paste(my_path2,"Magnolia_mercedesiarum_MAXENT_he45_50_HR_run_29_10.tif",sep=""))

prediction50A_1 <- stack(paste(my_path2,"Magnolia_mercedesiarum_MAXENT_he85_50_HR_run_29_1.tif",sep=""))
prediction50A_2 <- stack(paste(my_path2,"Magnolia_mercedesiarum_MAXENT_he85_50_HR_run_29_2.tif",sep=""))
prediction50A_3 <- stack(paste(my_path2,"Magnolia_mercedesiarum_MAXENT_he85_50_HR_run_29_3.tif",sep=""))
prediction50A_4 <- stack(paste(my_path2,"Magnolia_mercedesiarum_MAXENT_he85_50_HR_run_29_4.tif",sep=""))
prediction50A_5 <- stack(paste(my_path2,"Magnolia_mercedesiarum_MAXENT_he85_50_HR_run_29_5.tif",sep=""))
prediction50A_6 <- stack(paste(my_path2,"Magnolia_mercedesiarum_MAXENT_he85_50_HR_run_29_6.tif",sep=""))
prediction50A_7 <- stack(paste(my_path2,"Magnolia_mercedesiarum_MAXENT_he85_50_HR_run_29_7.tif",sep=""))
prediction50A_8 <- stack(paste(my_path2,"Magnolia_mercedesiarum_MAXENT_he85_50_HR_run_29_8.tif",sep=""))
prediction50A_9 <- stack(paste(my_path2,"Magnolia_mercedesiarum_MAXENT_he85_50_HR_run_29_9.tif",sep=""))
prediction50A_10 <- stack(paste(my_path2,"Magnolia_mercedesiarum_MAXENT_he85_50_HR_run_29_10.tif",sep=""))

prediction70_1 <- stack(paste(my_path2,"Magnolia_mercedesiarum_MAXENT_he45_70_HR_run_29_1.tif",sep=""))
prediction70_2 <- stack(paste(my_path2,"Magnolia_mercedesiarum_MAXENT_he45_70_HR_run_29_2.tif",sep=""))
prediction70_3 <- stack(paste(my_path2,"Magnolia_mercedesiarum_MAXENT_he45_70_HR_run_29_3.tif",sep=""))
prediction70_4 <- stack(paste(my_path2,"Magnolia_mercedesiarum_MAXENT_he45_70_HR_run_29_4.tif",sep=""))
prediction70_5 <- stack(paste(my_path2,"Magnolia_mercedesiarum_MAXENT_he45_70_HR_run_29_5.tif",sep=""))
prediction70_6 <- stack(paste(my_path2,"Magnolia_mercedesiarum_MAXENT_he45_70_HR_run_29_6.tif",sep=""))
prediction70_7 <- stack(paste(my_path2,"Magnolia_mercedesiarum_MAXENT_he45_70_HR_run_29_7.tif",sep=""))
prediction70_8 <- stack(paste(my_path2,"Magnolia_mercedesiarum_MAXENT_he45_70_HR_run_29_8.tif",sep=""))
prediction70_9 <- stack(paste(my_path2,"Magnolia_mercedesiarum_MAXENT_he45_70_HR_run_29_9.tif",sep=""))
prediction70_10 <- stack(paste(my_path2,"Magnolia_mercedesiarum_MAXENT_he45_70_HR_run_29_10.tif",sep=""))

prediction70A_1 <- stack(paste(my_path2,"Magnolia_mercedesiarum_MAXENT_he85_70_HR_run_29_1.tif",sep=""))
prediction70A_2 <- stack(paste(my_path2,"Magnolia_mercedesiarum_MAXENT_he85_70_HR_run_29_2.tif",sep=""))
prediction70A_3 <- stack(paste(my_path2,"Magnolia_mercedesiarum_MAXENT_he85_70_HR_run_29_3.tif",sep=""))
prediction70A_4 <- stack(paste(my_path2,"Magnolia_mercedesiarum_MAXENT_he85_70_HR_run_29_4.tif",sep=""))
prediction70A_5 <- stack(paste(my_path2,"Magnolia_mercedesiarum_MAXENT_he85_70_HR_run_29_5.tif",sep=""))
prediction70A_6 <- stack(paste(my_path2,"Magnolia_mercedesiarum_MAXENT_he85_70_HR_run_29_6.tif",sep=""))
prediction70A_7 <- stack(paste(my_path2,"Magnolia_mercedesiarum_MAXENT_he85_70_HR_run_29_7.tif",sep=""))
prediction70A_8 <- stack(paste(my_path2,"Magnolia_mercedesiarum_MAXENT_he85_70_HR_run_29_8.tif",sep=""))
prediction70A_9 <- stack(paste(my_path2,"Magnolia_mercedesiarum_MAXENT_he85_70_HR_run_29_9.tif",sep=""))
prediction70A_10 <- stack(paste(my_path2,"Magnolia_mercedesiarum_MAXENT_he85_70_HR_run_29_10.tif",sep=""))
```

```
set.seed(1)
random_presence_points <- randomPoints(presence_raster_mask, 
                                        n = presence_points_count, 
                                        tryf = 100)
```

###### Render presencce data

```
par(mfrow=c(1, 2))

mask_reclass_table <- matrix(c(-Inf, 1, 1, 1, Inf, NA), ncol=3, byrow=TRUE)
b_mask <- reclassify(subset(mde,1),mask_reclass_table)

plot(mde, xlim=c(-82,-70), ylim=c(-10,10), col = c(terrain.colors(256)))
plot(b_mask, add=TRUE, col=c('white'), legend = FALSE)
plot(presence_raster_mask, add=TRUE, alpha=0.9, col= gray(0.2), legend = FALSE)
world_high <- getMap(resolution = "high")
plot(world_high, add=TRUE, axes = TRUE)
#points(random_presence_points,col='black',cex=0.7)


plot(mde, xlim=c(-78.5,-76.5), ylim=c(-1.75,1.75), axes=TRUE, col = terrain.colors(256), legend = FALSE)
plot(presence_raster_mask, add=TRUE, alpha=0.4, col= gray(0.2), legend = FALSE)
world_high <- getMap(resolution = "high")
plot(world_high, add=TRUE)
points(random_presence_points,col='black',cex=0.7)
```

```
Magn_mde <- extract(mde, random_presence_points)

Magn_biovars00 <- extract(biovars00, random_presence_points)
Magn_biovars50 <- extract(biovars50, random_presence_points)
Magn_biovars70 <- extract(biovars70, random_presence_points)
Magn_biovars50A <- extract(biovars50A, random_presence_points)
Magn_biovars70A <- extract(biovars70A, random_presence_points)

Magn_prec00 <- extract(prec00, random_presence_points)
Magn_prec50 <- extract(prec50, random_presence_points)
Magn_prec70 <- extract(prec70, random_presence_points)
Magn_prec50A <- extract(prec50A, random_presence_points)
Magn_prec70A <- extract(prec70A, random_presence_points)

Magn_tmin00 <- extract(tmin00, random_presence_points) / 10
Magn_tmin50 <- extract(tmin50, random_presence_points) / 10
Magn_tmin70 <- extract(tmin70, random_presence_points) / 10
Magn_tmin50A <- extract(tmin50A, random_presence_points) / 10
Magn_tmin70A <- extract(tmin70A, random_presence_points) / 10

Magn_tmax00 <- extract(tmax00, random_presence_points) / 10
Magn_tmax50 <- extract(tmax50, random_presence_points) / 10
Magn_tmax70 <- extract(tmax70, random_presence_points) / 10
Magn_tmax50A <- extract(tmax50A, random_presence_points) / 10
Magn_tmax70A <- extract(tmax70A, random_presence_points) / 10
```

```
par(cex = 1.4)

Magn_pred00_1 <- extract(prediction00_1, random_presence_points)
Magn_pred00_2 <- extract(prediction00_2, random_presence_points)
Magn_pred00_3 <- extract(prediction00_3, random_presence_points)
Magn_pred00_4 <- extract(prediction00_4, random_presence_points)
Magn_pred00_5 <- extract(prediction00_5, random_presence_points)
Magn_pred00_6 <- extract(prediction00_6, random_presence_points)
Magn_pred00_7 <- extract(prediction00_7, random_presence_points)
Magn_pred00_8 <- extract(prediction00_8, random_presence_points)
Magn_pred00_9 <- extract(prediction00_9, random_presence_points)
Magn_pred00_10 <- extract(prediction00_10, random_presence_points)

Magn_pred50_1 <- extract(prediction50_1, random_presence_points)
Magn_pred50_2 <- extract(prediction50_2, random_presence_points)
Magn_pred50_3 <- extract(prediction50_3, random_presence_points)
Magn_pred50_4 <- extract(prediction50_4, random_presence_points)
Magn_pred50_5 <- extract(prediction50_5, random_presence_points)
Magn_pred50_6 <- extract(prediction50_6, random_presence_points)
Magn_pred50_7 <- extract(prediction50_7, random_presence_points)
Magn_pred50_8 <- extract(prediction50_8, random_presence_points)
Magn_pred50_9 <- extract(prediction50_9, random_presence_points)
Magn_pred50_10 <- extract(prediction50_10, random_presence_points)

Magn_pred50A_1 <- extract(prediction50A_1, random_presence_points)
Magn_pred50A_2 <- extract(prediction50A_2, random_presence_points)
Magn_pred50A_3 <- extract(prediction50A_3, random_presence_points)
Magn_pred50A_4 <- extract(prediction50A_4, random_presence_points)
Magn_pred50A_5 <- extract(prediction50A_5, random_presence_points)
Magn_pred50A_6 <- extract(prediction50A_6, random_presence_points)
Magn_pred50A_7 <- extract(prediction50A_7, random_presence_points)
Magn_pred50A_8 <- extract(prediction50A_8, random_presence_points)
Magn_pred50A_9 <- extract(prediction50A_9, random_presence_points)
Magn_pred50A_10 <- extract(prediction50A_10, random_presence_points)

Magn_pred70_1 <- extract(prediction70_1, random_presence_points)
Magn_pred70_2 <- extract(prediction70_2, random_presence_points)
Magn_pred70_3 <- extract(prediction70_3, random_presence_points)
Magn_pred70_4 <- extract(prediction70_4, random_presence_points)
Magn_pred70_5 <- extract(prediction70_5, random_presence_points)
Magn_pred70_6 <- extract(prediction70_6, random_presence_points)
Magn_pred70_7 <- extract(prediction70_7, random_presence_points)
Magn_pred70_8 <- extract(prediction70_8, random_presence_points)
Magn_pred70_9 <- extract(prediction70_9, random_presence_points)
Magn_pred70_10 <- extract(prediction70_10, random_presence_points)

Magn_pred70A_1 <- extract(prediction70A_1, random_presence_points)
Magn_pred70A_2 <- extract(prediction70A_2, random_presence_points)
Magn_pred70A_3 <- extract(prediction70A_3, random_presence_points)
Magn_pred70A_4 <- extract(prediction70A_4, random_presence_points)
Magn_pred70A_5 <- extract(prediction70A_5, random_presence_points)
Magn_pred70A_6 <- extract(prediction70A_6, random_presence_points)
Magn_pred70A_7 <- extract(prediction70A_7, random_presence_points)
Magn_pred70A_8 <- extract(prediction70A_8, random_presence_points)
Magn_pred70A_9 <- extract(prediction70A_9, random_presence_points)
Magn_pred70A_10 <- extract(prediction70A_10, random_presence_points)

Magn_pred00 <- c(Magn_pred00_1,Magn_pred00_2,Magn_pred00_3,Magn_pred00_4,Magn_pred00_5,
                 Magn_pred00_6,Magn_pred00_7,Magn_pred00_8,Magn_pred00_9,Magn_pred00_10)
Magn_pred50 <- c(Magn_pred50_1,Magn_pred50_2,Magn_pred50_3,Magn_pred50_4,Magn_pred50_5,
                 Magn_pred50_6,Magn_pred50_7,Magn_pred50_8,Magn_pred50_9,Magn_pred50_10)
Magn_pred50A <- c(Magn_pred50A_1,Magn_pred50A_2,Magn_pred50A_3,Magn_pred50A_4,Magn_pred50A_5,
                 Magn_pred50A_6,Magn_pred50A_7,Magn_pred50A_8,Magn_pred50A_9,Magn_pred50A_10)
Magn_pred70 <- c(Magn_pred70_1,Magn_pred70_2,Magn_pred70_3,Magn_pred70_4,Magn_pred70_5,
                 Magn_pred70_6,Magn_pred70_7,Magn_pred70_8,Magn_pred70_9,Magn_pred70_10)
Magn_pred70A <- c(Magn_pred70A_1,Magn_pred70A_2,Magn_pred70A_3,Magn_pred70A_4,Magn_pred70A_5,
                 Magn_pred70A_6,Magn_pred70A_7,Magn_pred70A_8,Magn_pred70A_9,Magn_pred70A_10)
```

#### Elevation

```
summary(Magn_mde)
```

```
##  GMTED2010_075_Andes
##  Min.   :1038       
##  1st Qu.:1525       
##  Median :1732       
##  Mean   :1754       
##  3rd Qu.:1998       
##  Max.   :2691
```

```
quantile(Magn_mde, probs = c(0.05, 0.95))
```

```
##      5%     95% 
## 1208.90 2303.15
```

```
boxplot(Magn_mde, boxfill = 'lightgray')
```

```
par(mfrow=c(1, 2))

qqnorm(Magn_mde, col="black", main='Normality QQ plot')
qqline(Magn_mde, col="red")

plot(density(Magn_mde), type="n", main = "Distribution along the altitudinal range")
polygon(density(Magn_mde), col="lightgray")
```

#### Effect on habitat suitabilit as presence probability (PP), Scenarios RCP 4.5 & RCP 8.5

```
# etss vector from other script (temporal solution)
etss_vector <- c(0.4777498,0.4717498,0.5180254,0.4159884,0.4304767,0.4204661,0.4722541,0.4856447,0.4036238,0.3599727)


summary(Magn_pred00)
```

```
##    Min. 1st Qu.  Median    Mean 3rd Qu.    Max. 
## 0.03546 0.59384 0.72327 0.68950 0.82188 0.97878
```

```
quantile(Magn_pred00, probs = c(0.05,0.1,0.9,0.95))
```

```
##        5%       10%       90%       95% 
## 0.3305005 0.4293763 0.8835472 0.9092352
```

```
summary(Magn_pred50)
```

```
##     Min.  1st Qu.   Median     Mean  3rd Qu.     Max. 
## 0.002335 0.106989 0.183765 0.232799 0.310708 0.965332
```

```
quantile(Magn_pred50, probs = c(0.05,0.1,0.9,0.95))
```

```
##         5%        10%        90%        95% 
## 0.03888427 0.05876544 0.48180654 0.58160847
```

```
summary(Magn_pred50A)
```

```
##     Min.  1st Qu.   Median     Mean  3rd Qu.     Max. 
## 0.001163 0.053221 0.098909 0.126259 0.169245 0.704904
```

```
quantile(Magn_pred50A, probs = c(0.05,0.1,0.9,0.95))
```

```
##         5%        10%        90%        95% 
## 0.01858120 0.02886854 0.25961016 0.32559102
```

```
summary(Magn_pred70)
```

```
##    Min. 1st Qu.  Median    Mean 3rd Qu.    Max. 
## 0.00121 0.05082 0.08560 0.11963 0.15021 0.85150
```

```
quantile(Magn_pred70, probs = c(0.05,0.1,0.9,0.95))
```

```
##         5%        10%        90%        95% 
## 0.01764701 0.02679474 0.25038290 0.32668721
```

```
summary(Magn_pred70A)
```

```
##      Min.   1st Qu.    Median      Mean   3rd Qu.      Max. 
## 0.0003005 0.0150034 0.0271530 0.0422623 0.0505465 0.4412384
```

```
quantile(Magn_pred70A, probs = c(0.05,0.1,0.9,0.95))
```

```
##          5%         10%         90%         95% 
## 0.005458507 0.008093949 0.093708216 0.134155174
```

```
par(cex = 1.4)

boxplot(Magn_pred00, Magn_pred50, Magn_pred70, 
        boxfill = c('lightgreen','coral','red4'),
        names = c('1970-2000','2050 RCP4.5','2070 RCP4.5'),
        main = 'Probability of presence within the actual distribution area RCP4.5')
```

```
boxplot(Magn_pred00, Magn_pred50A, Magn_pred70A, 
        boxfill = c('lightgreen','red3','tan4'),
        names = c('1970-2000','2050 RCP8.5','2070 RCP8.5'),
        main = 'Probability of presence within the actual distribution area RCP8.5')
```

```
boxplot(Magn_pred00, Magn_pred50, Magn_pred50A, Magn_pred70, Magn_pred70A, 
        boxfill = c('lightgreen','coral','red3','red4','tan4'),
        names = c('1970-2000','2050 RCP4.5','2050 RCP8.5','2070 RCP4.5','2070 RCP8.5'),
        main = 'Probability of presence within the actual distribution area RCP4.5 & RCP8.5')

etss_vector_median <- median(etss_vector)
etss_vector_conf <- wilcox.test(etss_vector, conf.int=TRUE)$conf.int
abline(h=etss_vector_median, col="red")
abline(h=etss_vector_conf, lty="dotted", col="red")
```

```
compareMWT(Magn_pred00,Magn_pred50,'actual PP vs. RCP4.5 2050 PP')
```

```
## Mann-Whitney test results for actual PP vs. RCP4.5 2050 PP 
## The H0 is rejected as p < 0.05, the HA accepted (medians unequal) 
## 
##  Wilcoxon rank sum test with continuity correction
## 
## data:  x and y
## W = 94859000, p-value < 2.2e-16
## alternative hypothesis: true location shift is not equal to 0
## 95 percent confidence interval:
##  0.4921298 0.5020087
## sample estimates:
## difference in location 
##              0.4970886
```

```
compareMWT(Magn_pred50,Magn_pred70,'RCP4.5 2050 PP vs. RCP4.5 2070 PP')
```

```
## Mann-Whitney test results for RCP4.5 2050 PP vs. RCP4.5 2070 PP 
## The H0 is rejected as p < 0.05, the HA accepted (medians unequal) 
## 
##  Wilcoxon rank sum test with continuity correction
## 
## data:  x and y
## W = 73423000, p-value < 2.2e-16
## alternative hypothesis: true location shift is not equal to 0
## 95 percent confidence interval:
##  0.08353288 0.08967508
## sample estimates:
## difference in location 
##             0.08659209
```

```
compareMWT(Magn_pred00,Magn_pred70,'actual PP vs. RCP45 2070 PP')
```

```
## Mann-Whitney test results for actual PP vs. RCP45 2070 PP 
## The H0 is rejected as p < 0.05, the HA accepted (medians unequal) 
## 
##  Wilcoxon rank sum test with continuity correction
## 
## data:  x and y
## W = 98853000, p-value < 2.2e-16
## alternative hypothesis: true location shift is not equal to 0
## 95 percent confidence interval:
##  0.6032760 0.6110524
## sample estimates:
## difference in location 
##              0.6071808
```

```
compareMWT(Magn_pred00,Magn_pred50A,'actual PP vs. RCP85 2050 PP')
```

```
## Mann-Whitney test results for actual PP vs. RCP85 2050 PP 
## The H0 is rejected as p < 0.05, the HA accepted (medians unequal) 
## 
##  Wilcoxon rank sum test with continuity correction
## 
## data:  x and y
## W = 99073000, p-value < 2.2e-16
## alternative hypothesis: true location shift is not equal to 0
## 95 percent confidence interval:
##  0.5932638 0.6011408
## sample estimates:
## difference in location 
##              0.5972246
```

```
compareMWT(Magn_pred50A,Magn_pred70A,'RCP8.5 2050 PP vs. RCP8.5 2070 PP')
```

```
## Mann-Whitney test results for RCP8.5 2050 PP vs. RCP8.5 2070 PP 
## The H0 is rejected as p < 0.05, the HA accepted (medians unequal) 
## 
##  Wilcoxon rank sum test with continuity correction
## 
## data:  x and y
## W = 82561000, p-value < 2.2e-16
## alternative hypothesis: true location shift is not equal to 0
## 95 percent confidence interval:
##  0.06084872 0.06425564
## sample estimates:
## difference in location 
##             0.06253067
```

```
compareMWT(Magn_pred00,Magn_pred70A,'actual PP vs. RCP8.5 2070 PP')
```

```
## Mann-Whitney test results for actual PP vs. RCP8.5 2070 PP 
## The H0 is rejected as p < 0.05, the HA accepted (medians unequal) 
## 
##  Wilcoxon rank sum test with continuity correction
## 
## data:  x and y
## W = 99943000, p-value < 2.2e-16
## alternative hypothesis: true location shift is not equal to 0
## 95 percent confidence interval:
##  0.6768992 0.6838107
## sample estimates:
## difference in location 
##              0.6803603
```

```
#compareMWT(Magn_pred50A,Magn_pred70,'RCP85 2050 PP vs. RCP45 2070 PP')
```

#### Climatic variables, Model HadGEM2 - ES, Scenario RCP 4.5

###### Precipitation

```
par(cex = 1.4)

summary(Magn_prec00)
```

```
##  wc2.0_HR_prec_stack.1 wc2.0_HR_prec_stack.2 wc2.0_HR_prec_stack.3
##  Min.   : 76.0         Min.   :103.0         Min.   :133.0        
##  1st Qu.:134.0         1st Qu.:161.0         1st Qu.:207.0        
##  Median :151.0         Median :177.0         Median :228.0        
##  Mean   :150.4         Mean   :176.9         Mean   :227.1        
##  3rd Qu.:169.0         3rd Qu.:195.0         3rd Qu.:248.0        
##  Max.   :209.0         Max.   :224.0         Max.   :299.0        
##  wc2.0_HR_prec_stack.4 wc2.0_HR_prec_stack.5 wc2.0_HR_prec_stack.6
##  Min.   :152.0         Min.   :136.0         Min.   :134.0        
##  1st Qu.:255.0         1st Qu.:244.0         1st Qu.:272.0        
##  Median :285.0         Median :283.0         Median :317.0        
##  Mean   :284.4         Mean   :284.5         Mean   :321.9        
##  3rd Qu.:315.0         3rd Qu.:325.0         3rd Qu.:373.2        
##  Max.   :378.0         Max.   :405.0         Max.   :484.0        
##  wc2.0_HR_prec_stack.7 wc2.0_HR_prec_stack.8 wc2.0_HR_prec_stack.9
##  Min.   :116.0         Min.   :100           Min.   :115.0        
##  1st Qu.:258.0         1st Qu.:194           1st Qu.:203.0        
##  Median :304.0         Median :226           Median :233.0        
##  Mean   :307.8         Mean   :227           Mean   :235.8        
##  3rd Qu.:363.0         3rd Qu.:263           3rd Qu.:272.0        
##  Max.   :452.0         Max.   :338           Max.   :336.0        
##  wc2.0_HR_prec_stack.10 wc2.0_HR_prec_stack.11 wc2.0_HR_prec_stack.12
##  Min.   :119            Min.   :113.0          Min.   : 85.0         
##  1st Qu.:169            1st Qu.:155.0          1st Qu.:121.0         
##  Median :194            Median :177.0          Median :136.0         
##  Mean   :199            Mean   :182.2          Mean   :139.2         
##  3rd Qu.:227            3rd Qu.:206.0          3rd Qu.:157.0         
##  Max.   :290            Max.   :275.0          Max.   :203.0
```

```
summary(Magn_prec50)
```

```
##  he45pr50_stack.1 he45pr50_stack.2 he45pr50_stack.3 he45pr50_stack.4
##  Min.   : 81.53   Min.   :102.3    Min.   :127.9    Min.   :150.0   
##  1st Qu.:164.18   1st Qu.:183.0    1st Qu.:224.5    1st Qu.:272.9   
##  Median :212.44   Median :232.8    Median :262.6    Median :319.5   
##  Mean   :214.67   Mean   :229.9    Mean   :267.6    Mean   :328.2   
##  3rd Qu.:262.85   3rd Qu.:275.7    3rd Qu.:317.2    3rd Qu.:387.4   
##  Max.   :366.85   Max.   :353.5    Max.   :423.5    Max.   :472.2   
##  he45pr50_stack.5 he45pr50_stack.6 he45pr50_stack.7 he45pr50_stack.8
##  Min.   :139.4    Min.   :123.1    Min.   :107.1    Min.   : 98.24  
##  1st Qu.:286.6    1st Qu.:311.5    1st Qu.:308.3    1st Qu.:249.24  
##  Median :354.3    Median :391.6    Median :399.6    Median :320.03  
##  Mean   :361.8    Mean   :397.6    Mean   :398.5    Mean   :317.72  
##  3rd Qu.:446.3    3rd Qu.:487.7    3rd Qu.:492.6    3rd Qu.:395.45  
##  Max.   :545.0    Max.   :659.2    Max.   :602.7    Max.   :464.51  
##  he45pr50_stack.9 he45pr50_stack.10 he45pr50_stack.11 he45pr50_stack.12
##  Min.   :108.7    Min.   :110.9     Min.   :114.3     Min.   : 88.4    
##  1st Qu.:218.0    1st Qu.:165.6     1st Qu.:176.1     1st Qu.:149.3    
##  Median :267.2    Median :198.1     Median :214.8     Median :191.6    
##  Mean   :269.9    Mean   :206.4     Mean   :229.4     Mean   :204.4    
##  3rd Qu.:322.4    3rd Qu.:249.2     3rd Qu.:289.8     3rd Qu.:261.9    
##  Max.   :412.4    Max.   :314.8     Max.   :364.9     Max.   :363.8
```

```
summary(Magn_prec70)
```

```
##  he45pr70_stack.1 he45pr70_stack.2 he45pr70_stack.3 he45pr70_stack.4
##  Min.   : 76.48   Min.   : 94.43   Min.   :115.0    Min.   :131.2   
##  1st Qu.:154.26   1st Qu.:167.56   1st Qu.:205.2    1st Qu.:244.0   
##  Median :198.96   Median :212.83   Median :241.2    Median :289.2   
##  Mean   :201.55   Mean   :209.77   Mean   :246.0    Mean   :296.7   
##  3rd Qu.:247.50   3rd Qu.:250.92   3rd Qu.:292.9    3rd Qu.:354.4   
##  Max.   :343.56   Max.   :320.49   Max.   :388.6    Max.   :436.6   
##  he45pr70_stack.5 he45pr70_stack.6 he45pr70_stack.7 he45pr70_stack.8
##  Min.   :133.5    Min.   :125.2    Min.   :108.1    Min.   : 99.18  
##  1st Qu.:275.7    1st Qu.:316.0    1st Qu.:307.7    1st Qu.:249.50  
##  Median :340.6    Median :395.9    Median :398.5    Median :319.29  
##  Mean   :348.8    Mean   :402.6    Mean   :397.3    Mean   :317.95  
##  3rd Qu.:430.1    3rd Qu.:491.4    3rd Qu.:491.8    3rd Qu.:396.11  
##  Max.   :532.0    Max.   :669.2    Max.   :601.7    Max.   :471.50  
##  he45pr70_stack.9 he45pr70_stack.10 he45pr70_stack.11 he45pr70_stack.12
##  Min.   :104.7    Min.   :106.9     Min.   :107.3     Min.   : 86.4    
##  1st Qu.:211.5    1st Qu.:159.6     1st Qu.:167.3     1st Qu.:147.0    
##  Median :259.7    Median :190.5     Median :205.6     Median :187.5    
##  Mean   :262.4    Mean   :198.6     Mean   :218.8     Mean   :200.4    
##  3rd Qu.:314.5    3rd Qu.:239.9     3rd Qu.:276.5     3rd Qu.:257.5    
##  Max.   :402.5    Max.   :302.4     Max.   :351.9     Max.   :354.7
```

```
boxplot(Magn_prec00, ylim = c(0,650), xlim = c(0.5, 12.5), names = month_labels,
        boxfill=rgb(1, 1, 1, alpha=1), border=rgb(1, 1, 1, alpha=1),
        main = 'Monthly precipitation', ylab = 'mm')
boxplot(Magn_prec00, at = 1:12 - 0.25, boxwex=0.2, xaxt = "n", add = TRUE, boxfill = 'lightgreen')
boxplot(Magn_prec50, at = 1:12, boxwex=0.2, xaxt = "n", add = TRUE, boxfill = 'coral')
boxplot(Magn_prec70, at = 1:12 + 0.25, boxwex=0.2, xaxt = "n", add = TRUE, boxfill = 'red4')

legend("topleft", c("1970-2000 (WorldClim 2)","2050 (RCP4.5)","2070 (RCP4.5)"), 
       inset=.02, box.lty=0, fill=c('lightgreen','coral','red4'), horiz=TRUE, cex=1.2)
```

```
par(cex = 1.4)

summary(Magn_tmin00)
```

```
##  wc2.0_HR_tmin_stack.1 wc2.0_HR_tmin_stack.2 wc2.0_HR_tmin_stack.3
##  Min.   : 7.24         Min.   : 7.115        Min.   : 7.295       
##  1st Qu.:11.78         1st Qu.:11.800        1st Qu.:11.936       
##  Median :13.47         Median :13.503        Median :13.586       
##  Mean   :13.20         Mean   :13.213        Mean   :13.294       
##  3rd Qu.:14.53         3rd Qu.:14.546        3rd Qu.:14.605       
##  Max.   :16.93         Max.   :16.904        Max.   :16.925       
##  wc2.0_HR_tmin_stack.4 wc2.0_HR_tmin_stack.5 wc2.0_HR_tmin_stack.6
##  Min.   : 7.444        Min.   : 7.333        Min.   : 6.688       
##  1st Qu.:11.989        1st Qu.:11.965        1st Qu.:11.361       
##  Median :13.604        Median :13.588        Median :13.071       
##  Mean   :13.312        Mean   :13.283        Mean   :12.752       
##  3rd Qu.:14.579        3rd Qu.:14.531        3rd Qu.:14.072       
##  Max.   :16.887        Max.   :16.883        Max.   :16.443       
##  wc2.0_HR_tmin_stack.7 wc2.0_HR_tmin_stack.8 wc2.0_HR_tmin_stack.9
##  Min.   : 6.333        Min.   : 6.436        Min.   : 6.869       
##  1st Qu.:11.038        1st Qu.:11.199        1st Qu.:11.642       
##  Median :12.702        Median :12.898        Median :13.295       
##  Mean   :12.514        Mean   :12.717        Mean   :13.120       
##  3rd Qu.:13.869        3rd Qu.:14.081        3rd Qu.:14.459       
##  Max.   :16.467        Max.   :16.709        Max.   :17.037       
##  wc2.0_HR_tmin_stack.10 wc2.0_HR_tmin_stack.11 wc2.0_HR_tmin_stack.12
##  Min.   : 7.223         Min.   : 7.40          Min.   : 7.354        
##  1st Qu.:11.873         1st Qu.:11.94          1st Qu.:11.845        
##  Median :13.473         Median :13.53          Median :13.423        
##  Mean   :13.304         Mean   :13.37          Mean   :13.273        
##  3rd Qu.:14.634         3rd Qu.:14.69          3rd Qu.:14.592        
##  Max.   :17.113         Max.   :17.17          Max.   :17.089
```

```
summary(Magn_tmin50)
```

```
##  he45tn50_stack.1 he45tn50_stack.2 he45tn50_stack.3 he45tn50_stack.4
##  Min.   : 9.973   Min.   :10.04    Min.   :10.41    Min.   :10.63   
##  1st Qu.:13.304   1st Qu.:13.41    1st Qu.:13.57    1st Qu.:13.80   
##  Median :14.677   Median :14.77    Median :14.86    Median :15.07   
##  Mean   :14.572   Mean   :14.66    Mean   :14.78    Mean   :14.98   
##  3rd Qu.:15.737   3rd Qu.:15.81    3rd Qu.:15.90    3rd Qu.:16.06   
##  Max.   :18.308   Max.   :18.35    Max.   :18.42    Max.   :18.50   
##  he45tn50_stack.5 he45tn50_stack.6 he45tn50_stack.7 he45tn50_stack.8
##  Min.   :10.62    Min.   :10.02    Min.   : 9.956   Min.   : 9.913  
##  1st Qu.:13.55    1st Qu.:13.05    1st Qu.:12.786   1st Qu.:12.675  
##  Median :14.89    Median :14.41    Median :14.166   Median :14.072  
##  Mean   :14.79    Mean   :14.32    Mean   :14.081   Mean   :13.991  
##  3rd Qu.:15.91    3rd Qu.:15.47    3rd Qu.:15.245   3rd Qu.:15.162  
##  Max.   :18.40    Max.   :18.21    Max.   :17.918   Max.   :17.859  
##  he45tn50_stack.9 he45tn50_stack.10 he45tn50_stack.11 he45tn50_stack.12
##  Min.   : 9.97    Min.   : 9.932    Min.   : 9.972    Min.   : 9.975   
##  1st Qu.:12.93    1st Qu.:13.406    1st Qu.:13.468    1st Qu.:13.349   
##  Median :14.32    Median :14.771    Median :14.821    Median :14.717   
##  Mean   :14.24    Mean   :14.657    Mean   :14.715    Mean   :14.613   
##  3rd Qu.:15.42    3rd Qu.:15.811    3rd Qu.:15.870    3rd Qu.:15.781   
##  Max.   :18.09    Max.   :18.380    Max.   :18.395    Max.   :18.317
```

```
summary(Magn_tmin70)
```

```
##  he45tn70_stack.1 he45tn70_stack.2 he45tn70_stack.3 he45tn70_stack.4
##  Min.   :10.27    Min.   :10.44    Min.   :10.81    Min.   :11.03   
##  1st Qu.:13.65    1st Qu.:13.81    1st Qu.:13.98    1st Qu.:14.18   
##  Median :15.04    Median :15.17    Median :15.26    Median :15.44   
##  Mean   :14.93    Mean   :15.06    Mean   :15.19    Mean   :15.36   
##  3rd Qu.:16.12    3rd Qu.:16.21    3rd Qu.:16.31    3rd Qu.:16.43   
##  Max.   :18.71    Max.   :18.75    Max.   :18.82    Max.   :18.90   
##  he45tn70_stack.5 he45tn70_stack.6 he45tn70_stack.7 he45tn70_stack.8
##  Min.   :11.02    Min.   :10.62    Min.   :10.46    Min.   :10.41   
##  1st Qu.:13.95    1st Qu.:13.65    1st Qu.:13.29    1st Qu.:13.18   
##  Median :15.29    Median :14.98    Median :14.66    Median :14.57   
##  Mean   :15.19    Mean   :14.91    Mean   :14.58    Mean   :14.50   
##  3rd Qu.:16.31    3rd Qu.:16.05    3rd Qu.:15.74    3rd Qu.:15.68   
##  Max.   :18.80    Max.   :18.71    Max.   :18.42    Max.   :18.37   
##  he45tn70_stack.9 he45tn70_stack.10 he45tn70_stack.11 he45tn70_stack.12
##  Min.   :10.37    Min.   :10.43     Min.   :10.41     Min.   :10.37    
##  1st Qu.:13.41    1st Qu.:13.87     1st Qu.:13.87     1st Qu.:13.77    
##  Median :14.82    Median :15.22     Median :15.22     Median :15.17    
##  Mean   :14.73    Mean   :15.11     Mean   :15.12     Mean   :15.05    
##  3rd Qu.:15.92    3rd Qu.:16.30     3rd Qu.:16.27     3rd Qu.:16.23    
##  Max.   :18.59    Max.   :18.79     Max.   :18.80     Max.   :18.82
```

```
boxplot(Magn_tmin00, ylim = c(5,25), xlim = c(0.5, 12.5), names = month_labels,
        boxfill=rgb(1, 1, 1, alpha=1), border=rgb(1, 1, 1, alpha=1),
        main = 'Monthly minimal temperature', ylab = '°C')
boxplot(Magn_tmin00, at = 1:12 - 0.25, boxwex=0.2, xaxt = "n", add = TRUE, boxfill = 'lightgreen')
boxplot(Magn_tmin50, at = 1:12, boxwex=0.2, xaxt = "n", add = TRUE, boxfill = 'coral')
boxplot(Magn_tmin70, at = 1:12 + 0.25, boxwex=0.2, xaxt = "n", add = TRUE, boxfill = 'red4')

legend("topleft", c("1970-2000 (WorldClim 2)","2050 (RCP 4.5)","2070 (RCP 4.5)"), 
       inset=.02, box.lty=0, fill=c('lightgreen','coral','red4'), horiz=TRUE, cex=1.2)
```

```
par(cex = 1.4)

summary(Magn_tmax00)
```

```
##  wc2.0_HR_tmax_stack.1 wc2.0_HR_tmax_stack.2 wc2.0_HR_tmax_stack.3
##  Min.   :18.06         Min.   :17.94         Min.   :18.12        
##  1st Qu.:21.41         1st Qu.:21.41         1st Qu.:21.55        
##  Median :22.41         Median :22.43         Median :22.50        
##  Mean   :22.38         Mean   :22.40         Mean   :22.48        
##  3rd Qu.:23.27         3rd Qu.:23.32         3rd Qu.:23.38        
##  Max.   :25.50         Max.   :25.51         Max.   :25.56        
##  wc2.0_HR_tmax_stack.4 wc2.0_HR_tmax_stack.5 wc2.0_HR_tmax_stack.6
##  Min.   :18.22         Min.   :18.06         Min.   :17.34        
##  1st Qu.:21.60         1st Qu.:21.56         1st Qu.:20.96        
##  Median :22.51         Median :22.49         Median :21.98        
##  Mean   :22.50         Mean   :22.47         Mean   :21.94        
##  3rd Qu.:23.35         3rd Qu.:23.34         3rd Qu.:22.84        
##  Max.   :25.47         Max.   :25.40         Max.   :25.01        
##  wc2.0_HR_tmax_stack.7 wc2.0_HR_tmax_stack.8 wc2.0_HR_tmax_stack.9
##  Min.   :17.14         Min.   :17.21         Min.   :17.67        
##  1st Qu.:21.02         1st Qu.:21.19         1st Qu.:21.64        
##  Median :22.08         Median :22.29         Median :22.71        
##  Mean   :21.94         Mean   :22.15         Mean   :22.55        
##  3rd Qu.:22.91         3rd Qu.:23.12         3rd Qu.:23.51        
##  Max.   :24.89         Max.   :25.09         Max.   :25.45        
##  wc2.0_HR_tmax_stack.10 wc2.0_HR_tmax_stack.11 wc2.0_HR_tmax_stack.12
##  Min.   :18.11          Min.   :18.32          Min.   :18.28         
##  1st Qu.:21.84          1st Qu.:21.90          1st Qu.:21.79         
##  Median :22.89          Median :22.93          Median :22.84         
##  Mean   :22.74          Mean   :22.80          Mean   :22.71         
##  3rd Qu.:23.66          3rd Qu.:23.71          3rd Qu.:23.62         
##  Max.   :25.62          Max.   :25.70          Max.   :25.59
```

```
summary(Magn_tmax50)
```

```
##  he45tx50_stack.1 he45tx50_stack.2 he45tx50_stack.3 he45tx50_stack.4
##  Min.   :20.38    Min.   :20.27    Min.   :20.04    Min.   :20.08   
##  1st Qu.:25.24    1st Qu.:25.18    1st Qu.:25.12    1st Qu.:24.98   
##  Median :26.06    Median :26.01    Median :25.98    Median :25.84   
##  Mean   :26.00    Mean   :25.94    Mean   :25.90    Mean   :25.75   
##  3rd Qu.:26.78    3rd Qu.:26.73    3rd Qu.:26.72    3rd Qu.:26.56   
##  Max.   :28.75    Max.   :28.69    Max.   :28.70    Max.   :28.50   
##  he45tx50_stack.5 he45tx50_stack.6 he45tx50_stack.7 he45tx50_stack.8
##  Min.   :19.63    Min.   :18.66    Min.   :18.29    Min.   :18.75   
##  1st Qu.:25.01    1st Qu.:24.14    1st Qu.:24.10    1st Qu.:24.71   
##  Median :25.83    Median :25.08    Median :25.08    Median :25.78   
##  Mean   :25.72    Mean   :24.97    Mean   :24.98    Mean   :25.65   
##  3rd Qu.:26.48    3rd Qu.:25.85    3rd Qu.:25.94    3rd Qu.:26.66   
##  Max.   :28.32    Max.   :27.90    Max.   :28.03    Max.   :28.75   
##  he45tx50_stack.9 he45tx50_stack.10 he45tx50_stack.11 he45tx50_stack.12
##  Min.   :19.68    Min.   :20.67     Min.   :20.83     Min.   :20.46    
##  1st Qu.:25.62    1st Qu.:25.84     1st Qu.:25.66     1st Qu.:25.46    
##  Median :26.65    Median :26.76     Median :26.56     Median :26.32    
##  Mean   :26.50    Mean   :26.66     Mean   :26.43     Mean   :26.24    
##  3rd Qu.:27.47    3rd Qu.:27.52     3rd Qu.:27.22     3rd Qu.:27.02    
##  Max.   :29.41    Max.   :29.43     Max.   :29.12     Max.   :28.99
```

```
summary(Magn_tmax70)
```

```
##  he45tx70_stack.1 he45tx70_stack.2 he45tx70_stack.3 he45tx70_stack.4
##  Min.   :20.78    Min.   :20.77    Min.   :20.54    Min.   :20.58   
##  1st Qu.:25.64    1st Qu.:25.68    1st Qu.:25.62    1st Qu.:25.48   
##  Median :26.46    Median :26.51    Median :26.48    Median :26.34   
##  Mean   :26.41    Mean   :26.44    Mean   :26.40    Mean   :26.25   
##  3rd Qu.:27.19    3rd Qu.:27.23    3rd Qu.:27.22    3rd Qu.:27.06   
##  Max.   :29.15    Max.   :29.19    Max.   :29.20    Max.   :29.00   
##  he45tx70_stack.5 he45tx70_stack.6 he45tx70_stack.7 he45tx70_stack.8
##  Min.   :20.23    Min.   :19.16    Min.   :18.69    Min.   :19.25   
##  1st Qu.:25.52    1st Qu.:24.71    1st Qu.:24.55    1st Qu.:25.21   
##  Median :26.34    Median :25.63    Median :25.51    Median :26.27   
##  Mean   :26.24    Mean   :25.52    Mean   :25.43    Mean   :26.14   
##  3rd Qu.:27.02    3rd Qu.:26.41    3rd Qu.:26.37    3rd Qu.:27.16   
##  Max.   :28.86    Max.   :28.40    Max.   :28.52    Max.   :29.25   
##  he45tx70_stack.9 he45tx70_stack.10 he45tx70_stack.11 he45tx70_stack.12
##  Min.   :20.28    Min.   :21.27     Min.   :21.33     Min.   :20.96    
##  1st Qu.:26.15    1st Qu.:26.44     1st Qu.:26.16     1st Qu.:25.93    
##  Median :27.17    Median :27.36     Median :27.06     Median :26.78    
##  Mean   :27.03    Mean   :27.26     Mean   :26.93     Mean   :26.68    
##  3rd Qu.:28.00    3rd Qu.:28.12     3rd Qu.:27.72     3rd Qu.:27.46    
##  Max.   :29.97    Max.   :30.03     Max.   :29.62     Max.   :29.39
```

```
boxplot(Magn_tmin00, ylim = c(15,32), xlim = c(0.5, 12.5), names = month_labels,
        boxfill=rgb(1, 1, 1, alpha=1), border=rgb(1, 1, 1, alpha=1),
        main = 'Monthly maximum temperature', ylab = '°C')
boxplot(Magn_tmax00, at = 1:12 - 0.25, boxwex=0.2, xaxt = "n", add = TRUE, boxfill = 'lightgreen')
boxplot(Magn_tmax50, at = 1:12, boxwex=0.2, xaxt = "n", add = TRUE, boxfill = 'coral')
boxplot(Magn_tmax70, at = 1:12 + 0.25, boxwex=0.2, xaxt = "n", add = TRUE, boxfill = 'red4')

legend("topleft", c("1970-2000 (WorldClim 2)","2050 (RCP 4.5)","2070 (RCP 4.5)"), 
       inset=.02, box.lty=0, fill=c('lightgreen','coral','red4'), horiz=TRUE, cex=1.2)
```

#### Climatic variables, Model HadGEM2 - ES, Scenario RCP 8.5

```
par(cex = 1.4)

summary(Magn_prec00)
```

```
##  wc2.0_HR_prec_stack.1 wc2.0_HR_prec_stack.2 wc2.0_HR_prec_stack.3
##  Min.   : 76.0         Min.   :103.0         Min.   :133.0        
##  1st Qu.:134.0         1st Qu.:161.0         1st Qu.:207.0        
##  Median :151.0         Median :177.0         Median :228.0        
##  Mean   :150.4         Mean   :176.9         Mean   :227.1        
##  3rd Qu.:169.0         3rd Qu.:195.0         3rd Qu.:248.0        
##  Max.   :209.0         Max.   :224.0         Max.   :299.0        
##  wc2.0_HR_prec_stack.4 wc2.0_HR_prec_stack.5 wc2.0_HR_prec_stack.6
##  Min.   :152.0         Min.   :136.0         Min.   :134.0        
##  1st Qu.:255.0         1st Qu.:244.0         1st Qu.:272.0        
##  Median :285.0         Median :283.0         Median :317.0        
##  Mean   :284.4         Mean   :284.5         Mean   :321.9        
##  3rd Qu.:315.0         3rd Qu.:325.0         3rd Qu.:373.2        
##  Max.   :378.0         Max.   :405.0         Max.   :484.0        
##  wc2.0_HR_prec_stack.7 wc2.0_HR_prec_stack.8 wc2.0_HR_prec_stack.9
##  Min.   :116.0         Min.   :100           Min.   :115.0        
##  1st Qu.:258.0         1st Qu.:194           1st Qu.:203.0        
##  Median :304.0         Median :226           Median :233.0        
##  Mean   :307.8         Mean   :227           Mean   :235.8        
##  3rd Qu.:363.0         3rd Qu.:263           3rd Qu.:272.0        
##  Max.   :452.0         Max.   :338           Max.   :336.0        
##  wc2.0_HR_prec_stack.10 wc2.0_HR_prec_stack.11 wc2.0_HR_prec_stack.12
##  Min.   :119            Min.   :113.0          Min.   : 85.0         
##  1st Qu.:169            1st Qu.:155.0          1st Qu.:121.0         
##  Median :194            Median :177.0          Median :136.0         
##  Mean   :199            Mean   :182.2          Mean   :139.2         
##  3rd Qu.:227            3rd Qu.:206.0          3rd Qu.:157.0         
##  Max.   :290            Max.   :275.0          Max.   :203.0
```

```
summary(Magn_prec50A)
```

```
##  he85pr50_stack.1 he85pr50_stack.2 he85pr50_stack.3 he85pr50_stack.4
##  Min.   : 75.41   Min.   : 97.38   Min.   :121.9    Min.   :134.1   
##  1st Qu.:152.39   1st Qu.:172.45   1st Qu.:214.2    1st Qu.:252.8   
##  Median :196.28   Median :218.45   Median :251.5    Median :300.5   
##  Mean   :198.97   Mean   :215.76   Mean   :256.2    Mean   :308.6   
##  3rd Qu.:244.45   3rd Qu.:257.55   3rd Qu.:304.8    3rd Qu.:369.4   
##  Max.   :337.05   Max.   :328.71   Max.   :402.8    Max.   :459.2   
##  he85pr50_stack.5 he85pr50_stack.6 he85pr50_stack.7 he85pr50_stack.8
##  Min.   :132.5    Min.   :115.1    Min.   : 95.16   Min.   :100.2   
##  1st Qu.:277.4    1st Qu.:295.1    1st Qu.:271.01   1st Qu.:251.2   
##  Median :345.3    Median :370.5    Median :351.72   Median :322.7   
##  Mean   :353.2    Mean   :377.9    Mean   :350.51   Mean   :320.8   
##  3rd Qu.:435.7    3rd Qu.:463.5    3rd Qu.:432.76   3rd Qu.:400.0   
##  Max.   :545.0    Max.   :639.3    Max.   :539.19   Max.   :472.5   
##  he85pr50_stack.9 he85pr50_stack.10 he85pr50_stack.11 he85pr50_stack.12
##  Min.   :105.8    Min.   :108.3     Min.   :110.4     Min.   : 87.46   
##  1st Qu.:208.5    1st Qu.:161.3     1st Qu.:168.5     1st Qu.:146.52   
##  Median :254.0    Median :191.6     Median :204.3     Median :185.61   
##  Mean   :257.2    Mean   :199.6     Mean   :218.4     Mean   :198.90   
##  3rd Qu.:306.3    3rd Qu.:240.6     3rd Qu.:275.4     3rd Qu.:255.24   
##  Max.   :395.5    Max.   :301.7     Max.   :348.9     Max.   :349.58
```

```
summary(Magn_prec70A)
```

```
##  he85pr70_stack.1 he85pr70_stack.2 he85pr70_stack.3 he85pr70_stack.4
##  Min.   : 73.42   Min.   : 94.33   Min.   :117.9    Min.   :132.0   
##  1st Qu.:151.77   1st Qu.:170.06   1st Qu.:210.3    1st Qu.:249.4   
##  Median :195.44   Median :216.33   Median :248.0    Median :298.9   
##  Mean   :199.37   Mean   :213.48   Mean   :253.2    Mean   :306.6   
##  3rd Qu.:245.88   3rd Qu.:256.32   3rd Qu.:302.2    3rd Qu.:368.8   
##  Max.   :330.91   Max.   :320.49   Max.   :401.8    Max.   :463.2   
##  he85pr70_stack.5 he85pr70_stack.6 he85pr70_stack.7 he85pr70_stack.8
##  Min.   :134.6    Min.   :132.0    Min.   :103.1    Min.   :106.2   
##  1st Qu.:290.3    1st Qu.:343.4    1st Qu.:296.1    1st Qu.:272.7   
##  Median :364.9    Median :439.6    Median :387.0    Median :351.3   
##  Mean   :372.6    Mean   :445.4    Mean   :385.4    Mean   :349.8   
##  3rd Qu.:461.1    3rd Qu.:552.2    3rd Qu.:477.5    3rd Qu.:437.0   
##  Max.   :583.0    Max.   :738.0    Max.   :595.7    Max.   :519.0   
##  he85pr70_stack.9 he85pr70_stack.10 he85pr70_stack.11 he85pr70_stack.12
##  Min.   : 96.72   Min.   : 87.8     Min.   :102.3     Min.   : 89.49   
##  1st Qu.:196.51   1st Qu.:132.1     1st Qu.:158.3     1st Qu.:153.02   
##  Median :240.29   Median :157.4     Median :193.4     Median :194.10   
##  Mean   :244.15   Mean   :164.2     Mean   :206.0     Mean   :208.52   
##  3rd Qu.:292.22   3rd Qu.:198.3     3rd Qu.:260.1     3rd Qu.:268.95   
##  Max.   :385.54   Max.   :257.8     Max.   :337.5     Max.   :358.77
```

```
boxplot(Magn_prec00, ylim = c(0,650), xlim = c(0.5, 12.5), names = month_labels,
        boxfill=rgb(1, 1, 1, alpha=1), border=rgb(1, 1, 1, alpha=1),
        main = 'Monthly precipitation', ylab = 'mm')
boxplot(Magn_prec00, at = 1:12 - 0.25, boxwex=0.2, xaxt = "n", add = TRUE, boxfill = 'lightgreen')
boxplot(Magn_prec50A, at = 1:12, boxwex=0.2, xaxt = "n", add = TRUE, boxfill = 'coral')
boxplot(Magn_prec70A, at = 1:12 + 0.25, boxwex=0.2, xaxt = "n", add = TRUE, boxfill = 'red4')

legend("topleft", c("1970-2000 (WorldClim 2)","2050 (RCP 8.5)","2070 (RCP 8.5)"), 
       inset=.02, box.lty=0, fill=c('lightgreen','coral','red4'), horiz=TRUE, cex=1.2)
```

```
par(cex = 1.4)

summary(Magn_tmin00)
```

```
##  wc2.0_HR_tmin_stack.1 wc2.0_HR_tmin_stack.2 wc2.0_HR_tmin_stack.3
##  Min.   : 7.24         Min.   : 7.115        Min.   : 7.295       
##  1st Qu.:11.78         1st Qu.:11.800        1st Qu.:11.936       
##  Median :13.47         Median :13.503        Median :13.586       
##  Mean   :13.20         Mean   :13.213        Mean   :13.294       
##  3rd Qu.:14.53         3rd Qu.:14.546        3rd Qu.:14.605       
##  Max.   :16.93         Max.   :16.904        Max.   :16.925       
##  wc2.0_HR_tmin_stack.4 wc2.0_HR_tmin_stack.5 wc2.0_HR_tmin_stack.6
##  Min.   : 7.444        Min.   : 7.333        Min.   : 6.688       
##  1st Qu.:11.989        1st Qu.:11.965        1st Qu.:11.361       
##  Median :13.604        Median :13.588        Median :13.071       
##  Mean   :13.312        Mean   :13.283        Mean   :12.752       
##  3rd Qu.:14.579        3rd Qu.:14.531        3rd Qu.:14.072       
##  Max.   :16.887        Max.   :16.883        Max.   :16.443       
##  wc2.0_HR_tmin_stack.7 wc2.0_HR_tmin_stack.8 wc2.0_HR_tmin_stack.9
##  Min.   : 6.333        Min.   : 6.436        Min.   : 6.869       
##  1st Qu.:11.038        1st Qu.:11.199        1st Qu.:11.642       
##  Median :12.702        Median :12.898        Median :13.295       
##  Mean   :12.514        Mean   :12.717        Mean   :13.120       
##  3rd Qu.:13.869        3rd Qu.:14.081        3rd Qu.:14.459       
##  Max.   :16.467        Max.   :16.709        Max.   :17.037       
##  wc2.0_HR_tmin_stack.10 wc2.0_HR_tmin_stack.11 wc2.0_HR_tmin_stack.12
##  Min.   : 7.223         Min.   : 7.40          Min.   : 7.354        
##  1st Qu.:11.873         1st Qu.:11.94          1st Qu.:11.845        
##  Median :13.473         Median :13.53          Median :13.423        
##  Mean   :13.304         Mean   :13.37          Mean   :13.273        
##  3rd Qu.:14.634         3rd Qu.:14.69          3rd Qu.:14.592        
##  Max.   :17.113         Max.   :17.17          Max.   :17.089
```

```
summary(Magn_tmin50A)
```

```
##  he85tn50_stack.1 he85tn50_stack.2 he85tn50_stack.3 he85tn50_stack.4
##  Min.   :10.47    Min.   :10.54    Min.   :10.91    Min.   :11.13   
##  1st Qu.:13.82    1st Qu.:13.99    1st Qu.:14.08    1st Qu.:14.30   
##  Median :15.19    Median :15.35    Median :15.39    Median :15.57   
##  Mean   :15.10    Mean   :15.24    Mean   :15.30    Mean   :15.48   
##  3rd Qu.:16.27    3rd Qu.:16.41    3rd Qu.:16.41    3rd Qu.:16.56   
##  Max.   :18.82    Max.   :18.95    Max.   :18.92    Max.   :19.00   
##  he85tn50_stack.5 he85tn50_stack.6 he85tn50_stack.7 he85tn50_stack.8
##  Min.   :11.12    Min.   :10.52    Min.   :10.46    Min.   :10.41   
##  1st Qu.:14.05    1st Qu.:13.55    1st Qu.:13.29    1st Qu.:13.26   
##  Median :15.39    Median :14.88    Median :14.67    Median :14.65   
##  Mean   :15.29    Mean   :14.81    Mean   :14.58    Mean   :14.58   
##  3rd Qu.:16.41    3rd Qu.:15.95    3rd Qu.:15.75    3rd Qu.:15.76   
##  Max.   :18.90    Max.   :18.61    Max.   :18.42    Max.   :18.46   
##  he85tn50_stack.9 he85tn50_stack.10 he85tn50_stack.11 he85tn50_stack.12
##  Min.   :10.47    Min.   :10.53     Min.   :10.51     Min.   :10.47    
##  1st Qu.:13.44    1st Qu.:13.92     1st Qu.:13.98     1st Qu.:13.94    
##  Median :14.86    Median :15.28     Median :15.34     Median :15.29    
##  Mean   :14.76    Mean   :15.18     Mean   :15.24     Mean   :15.19    
##  3rd Qu.:15.97    3rd Qu.:16.36     3rd Qu.:16.43     3rd Qu.:16.36    
##  Max.   :18.59    Max.   :18.88     Max.   :18.90     Max.   :18.92
```

```
summary(Magn_tmin70A)
```

```
##  he85tn70_stack.1 he85tn70_stack.2 he85tn70_stack.3 he85tn70_stack.4
##  Min.   :11.57    Min.   :11.74    Min.   :12.11    Min.   :12.33   
##  1st Qu.:14.98    1st Qu.:15.15    1st Qu.:15.28    1st Qu.:15.50   
##  Median :16.38    Median :16.52    Median :16.59    Median :16.77   
##  Mean   :16.27    Mean   :16.40    Mean   :16.50    Mean   :16.68   
##  3rd Qu.:17.46    3rd Qu.:17.57    3rd Qu.:17.61    3rd Qu.:17.76   
##  Max.   :20.01    Max.   :20.06    Max.   :20.12    Max.   :20.20   
##  he85tn70_stack.5 he85tn70_stack.6 he85tn70_stack.7 he85tn70_stack.8
##  Min.   :12.32    Min.   :12.02    Min.   :11.76    Min.   :11.88   
##  1st Qu.:15.25    1st Qu.:15.11    1st Qu.:14.66    1st Qu.:14.76   
##  Median :16.60    Median :16.47    Median :16.04    Median :16.13   
##  Mean   :16.50    Mean   :16.38    Mean   :15.96    Mean   :16.07   
##  3rd Qu.:17.62    3rd Qu.:17.54    3rd Qu.:17.13    3rd Qu.:17.26   
##  Max.   :20.11    Max.   :20.21    Max.   :19.82    Max.   :19.96   
##  he85tn70_stack.9 he85tn70_stack.10 he85tn70_stack.11 he85tn70_stack.12
##  Min.   :11.67    Min.   :11.73     Min.   :11.61     Min.   :11.57    
##  1st Qu.:14.73    1st Qu.:15.20     1st Qu.:15.15     1st Qu.:15.04    
##  Median :16.16    Median :16.57     Median :16.51     Median :16.40    
##  Mean   :16.07    Mean   :16.45     Mean   :16.39     Mean   :16.28    
##  3rd Qu.:17.27    3rd Qu.:17.61     3rd Qu.:17.56     3rd Qu.:17.45    
##  Max.   :19.89    Max.   :20.09     Max.   :20.10     Max.   :20.02
```

```
boxplot(Magn_tmin00, ylim = c(5,25), xlim = c(0.5, 12.5), names = month_labels,
        boxfill=rgb(1, 1, 1, alpha=1), border=rgb(1, 1, 1, alpha=1),
        main = 'Monthly minimal temperature', ylab = '°C')
boxplot(Magn_tmin00, at = 1:12 - 0.25, boxwex=0.2, xaxt = "n", add = TRUE, boxfill = 'lightgreen')
boxplot(Magn_tmin50A, at = 1:12, boxwex=0.2, xaxt = "n", add = TRUE, boxfill = 'coral')
boxplot(Magn_tmin70A, at = 1:12 + 0.25, boxwex=0.2, xaxt = "n", add = TRUE, boxfill = 'red4')

legend("topleft", c("1970-2000 (WorldClim 2)","2050 (RCP 8.5)","2070 (RCP 8.5)"), 
       inset=.02, box.lty=0, fill=c('lightgreen','coral','red4'), horiz=TRUE, cex=1.2)
```

```
par(cex = 1.4)

summary(Magn_tmax00)
```

```
##  wc2.0_HR_tmax_stack.1 wc2.0_HR_tmax_stack.2 wc2.0_HR_tmax_stack.3
##  Min.   :18.06         Min.   :17.94         Min.   :18.12        
##  1st Qu.:21.41         1st Qu.:21.41         1st Qu.:21.55        
##  Median :22.41         Median :22.43         Median :22.50        
##  Mean   :22.38         Mean   :22.40         Mean   :22.48        
##  3rd Qu.:23.27         3rd Qu.:23.32         3rd Qu.:23.38        
##  Max.   :25.50         Max.   :25.51         Max.   :25.56        
##  wc2.0_HR_tmax_stack.4 wc2.0_HR_tmax_stack.5 wc2.0_HR_tmax_stack.6
##  Min.   :18.22         Min.   :18.06         Min.   :17.34        
##  1st Qu.:21.60         1st Qu.:21.56         1st Qu.:20.96        
##  Median :22.51         Median :22.49         Median :21.98        
##  Mean   :22.50         Mean   :22.47         Mean   :21.94        
##  3rd Qu.:23.35         3rd Qu.:23.34         3rd Qu.:22.84        
##  Max.   :25.47         Max.   :25.40         Max.   :25.01        
##  wc2.0_HR_tmax_stack.7 wc2.0_HR_tmax_stack.8 wc2.0_HR_tmax_stack.9
##  Min.   :17.14         Min.   :17.21         Min.   :17.67        
##  1st Qu.:21.02         1st Qu.:21.19         1st Qu.:21.64        
##  Median :22.08         Median :22.29         Median :22.71        
##  Mean   :21.94         Mean   :22.15         Mean   :22.55        
##  3rd Qu.:22.91         3rd Qu.:23.12         3rd Qu.:23.51        
##  Max.   :24.89         Max.   :25.09         Max.   :25.45        
##  wc2.0_HR_tmax_stack.10 wc2.0_HR_tmax_stack.11 wc2.0_HR_tmax_stack.12
##  Min.   :18.11          Min.   :18.32          Min.   :18.28         
##  1st Qu.:21.84          1st Qu.:21.90          1st Qu.:21.79         
##  Median :22.89          Median :22.93          Median :22.84         
##  Mean   :22.74          Mean   :22.80          Mean   :22.71         
##  3rd Qu.:23.66          3rd Qu.:23.71          3rd Qu.:23.62         
##  Max.   :25.62          Max.   :25.70          Max.   :25.59
```

```
summary(Magn_tmax50A)
```

```
##  he85tx50_stack.1 he85tx50_stack.2 he85tx50_stack.3 he85tx50_stack.4
##  Min.   :21.08    Min.   :20.97    Min.   :20.64    Min.   :20.68   
##  1st Qu.:25.94    1st Qu.:25.88    1st Qu.:25.72    1st Qu.:25.58   
##  Median :26.76    Median :26.71    Median :26.58    Median :26.44   
##  Mean   :26.70    Mean   :26.64    Mean   :26.50    Mean   :26.35   
##  3rd Qu.:27.48    3rd Qu.:27.43    3rd Qu.:27.32    3rd Qu.:27.17   
##  Max.   :29.45    Max.   :29.39    Max.   :29.30    Max.   :29.10   
##  he85tx50_stack.5 he85tx50_stack.6 he85tx50_stack.7 he85tx50_stack.8
##  Min.   :20.33    Min.   :19.36    Min.   :18.99    Min.   :19.35   
##  1st Qu.:25.62    1st Qu.:24.80    1st Qu.:24.76    1st Qu.:25.34   
##  Median :26.44    Median :25.72    Median :25.72    Median :26.41   
##  Mean   :26.34    Mean   :25.62    Mean   :25.63    Mean   :26.28   
##  3rd Qu.:27.14    3rd Qu.:26.50    3rd Qu.:26.57    3rd Qu.:27.29   
##  Max.   :28.96    Max.   :28.50    Max.   :28.72    Max.   :29.35   
##  he85tx50_stack.9 he85tx50_stack.10 he85tx50_stack.11 he85tx50_stack.12
##  Min.   :20.38    Min.   :21.37     Min.   :21.53     Min.   :21.16    
##  1st Qu.:26.25    1st Qu.:26.54     1st Qu.:26.36     1st Qu.:26.13    
##  Median :27.27    Median :27.50     Median :27.26     Median :26.98    
##  Mean   :27.13    Mean   :27.37     Mean   :27.13     Mean   :26.88    
##  3rd Qu.:28.12    3rd Qu.:28.25     3rd Qu.:27.92     3rd Qu.:27.66    
##  Max.   :30.07    Max.   :30.13     Max.   :29.82     Max.   :29.59
```

```
summary(Magn_tmax70A)
```

```
##  he85tx70_stack.1 he85tx70_stack.2 he85tx70_stack.3 he85tx70_stack.4
##  Min.   :22.18    Min.   :22.17    Min.   :21.84    Min.   :21.88   
##  1st Qu.:27.07    1st Qu.:27.08    1st Qu.:26.92    1st Qu.:26.78   
##  Median :27.92    Median :27.91    Median :27.78    Median :27.64   
##  Mean   :27.84    Mean   :27.84    Mean   :27.70    Mean   :27.55   
##  3rd Qu.:28.60    3rd Qu.:28.63    3rd Qu.:28.52    3rd Qu.:28.36   
##  Max.   :30.55    Max.   :30.59    Max.   :30.50    Max.   :30.30   
##  he85tx70_stack.5 he85tx70_stack.6 he85tx70_stack.7 he85tx70_stack.8
##  Min.   :21.53    Min.   :20.82    Min.   :20.29    Min.   :20.85   
##  1st Qu.:26.82    1st Qu.:26.34    1st Qu.:26.07    1st Qu.:26.90   
##  Median :27.64    Median :27.29    Median :27.05    Median :27.95   
##  Mean   :27.53    Mean   :27.17    Mean   :26.95    Mean   :27.81   
##  3rd Qu.:28.31    3rd Qu.:28.06    3rd Qu.:27.89    3rd Qu.:28.84   
##  Max.   :30.16    Max.   :30.10    Max.   :30.02    Max.   :30.85   
##  he85tx70_stack.9 he85tx70_stack.10 he85tx70_stack.11 he85tx70_stack.12
##  Min.   :21.78    Min.   :22.87     Min.   :22.73     Min.   :22.16    
##  1st Qu.:27.65    1st Qu.:28.04     1st Qu.:27.56     1st Qu.:27.13    
##  Median :28.67    Median :28.96     Median :28.46     Median :27.98    
##  Mean   :28.53    Mean   :28.86     Mean   :28.33     Mean   :27.88    
##  3rd Qu.:29.52    3rd Qu.:29.72     3rd Qu.:29.12     3rd Qu.:28.66    
##  Max.   :31.47    Max.   :31.63     Max.   :31.02     Max.   :30.59
```

```
boxplot(Magn_tmin00, ylim = c(15,32), xlim = c(0.5, 12.5), names = month_labels,
        boxfill=rgb(1, 1, 1, alpha=1), border=rgb(1, 1, 1, alpha=1),
        main = 'Monthly maximum temperature', ylab = '°C')
boxplot(Magn_tmax00, at = 1:12 - 0.25, boxwex=0.2, xaxt = "n", add = TRUE, boxfill = 'lightgreen')
boxplot(Magn_tmax50A, at = 1:12, boxwex=0.2, xaxt = "n", add = TRUE, boxfill = 'coral')
boxplot(Magn_tmax70A, at = 1:12 + 0.25, boxwex=0.2, xaxt = "n", add = TRUE, boxfill = 'red4')

legend("topleft", c("1970-2000 (WorldClim 2)","2050 (RCP 8.5)","2070 (RCP 8.5)"), 
       inset=.02, box.lty=0, fill=c('lightgreen','coral','red4'), horiz=TRUE, cex=1.2)
```

#### Climatic variables, Model HadGEM2 - ES, Scenarios RCP 4.5 & RCP 8.5

```
par(cex = 1.4, mfrow=c(2, 2))

summary(Magn_biovars00)
```

```
##       bio1            bio2             bio3            bio4      
##  Min.   :124.7   Min.   : 84.50   Min.   :84.86   Min.   :187.6  
##  1st Qu.:165.9   1st Qu.: 87.78   1st Qu.:88.57   1st Qu.:243.3  
##  Median :179.0   Median : 91.12   Median :90.75   Median :290.8  
##  Mean   :177.7   Mean   : 93.10   Mean   :90.28   Mean   :281.5  
##  3rd Qu.:188.8   3rd Qu.: 98.27   3rd Qu.:92.07   3rd Qu.:315.5  
##  Max.   :211.0   Max.   :113.30   Max.   :94.02   Max.   :403.4  
##       bio5            bio6             bio7             bio8      
##  Min.   :183.2   Min.   : 63.33   Min.   : 91.20   Min.   :125.1  
##  1st Qu.:219.1   1st Qu.:110.37   1st Qu.: 96.93   1st Qu.:163.8  
##  Median :229.3   Median :126.95   Median :103.52   Median :177.0  
##  Mean   :228.2   Mean   :124.98   Mean   :103.17   Mean   :175.5  
##  3rd Qu.:237.3   3rd Qu.:138.67   3rd Qu.:108.81   3rd Qu.:186.1  
##  Max.   :257.0   Max.   :164.43   Max.   :130.00   Max.   :209.6  
##       bio9           bio10           bio11           bio12     
##  Min.   :126.7   Min.   :127.8   Min.   :118.6   Min.   :1409  
##  1st Qu.:167.2   1st Qu.:168.7   1st Qu.:161.1   1st Qu.:2380  
##  Median :180.5   Median :181.6   Median :175.0   Median :2728  
##  Mean   :179.0   Mean   :180.3   Mean   :173.4   Mean   :2736  
##  3rd Qu.:189.9   3rd Qu.:191.2   3rd Qu.:184.7   3rd Qu.:3097  
##  Max.   :212.3   Max.   :213.6   Max.   :207.5   Max.   :3801  
##      bio13           bio14           bio15           bio16       
##  Min.   :152.0   Min.   : 76.0   Min.   :16.40   Min.   : 436.0  
##  1st Qu.:272.0   1st Qu.:119.0   1st Qu.:25.01   1st Qu.: 779.0  
##  Median :317.0   Median :135.0   Median :27.00   Median : 904.0  
##  Mean   :323.0   Mean   :138.3   Mean   :26.81   Mean   : 918.3  
##  3rd Qu.:373.2   3rd Qu.:157.0   3rd Qu.:28.76   3rd Qu.:1062.5  
##  Max.   :484.0   Max.   :203.0   Max.   :32.43   Max.   :1324.0  
##      bio17           bio18           bio19       
##  Min.   :269.0   Min.   :328.0   Min.   : 350.0  
##  1st Qu.:407.0   1st Qu.:447.0   1st Qu.: 728.8  
##  Median :458.5   Median :509.0   Median : 845.0  
##  Mean   :463.1   Mean   :523.9   Mean   : 856.7  
##  3rd Qu.:519.2   3rd Qu.:596.0   3rd Qu.:1000.0  
##  Max.   :636.0   Max.   :767.0   Max.   :1273.0
```

```
summary(Magn_biovars50)
```

```
##       bio1            bio2             bio3            bio4      
##  Min.   :149.7   Min.   : 96.94   Min.   :87.96   Min.   :217.9  
##  1st Qu.:191.9   1st Qu.:110.09   1st Qu.:89.29   1st Qu.:320.3  
##  Median :203.2   Median :113.52   Median :89.63   Median :364.7  
##  Mean   :202.1   Mean   :113.60   Mean   :89.62   Mean   :361.7  
##  3rd Qu.:212.1   3rd Qu.:117.23   3rd Qu.:89.93   3rd Qu.:404.5  
##  Max.   :234.8   Max.   :124.19   Max.   :91.79   Max.   :529.1  
##       bio5            bio6             bio7            bio8      
##  Min.   :208.3   Min.   : 99.13   Min.   :109.2   Min.   :149.4  
##  1st Qu.:258.5   1st Qu.:126.75   1st Qu.:123.3   1st Qu.:188.6  
##  Median :267.7   Median :140.72   Median :126.8   Median :199.5  
##  Mean   :266.7   Mean   :139.91   Mean   :126.8   Mean   :198.5  
##  3rd Qu.:275.4   3rd Qu.:151.62   3rd Qu.:130.7   3rd Qu.:208.2  
##  Max.   :294.3   Max.   :178.59   Max.   :136.0   Max.   :231.2  
##       bio9           bio10           bio11           bio12     
##  Min.   :151.8   Min.   :153.1   Min.   :142.7   Min.   :1383  
##  1st Qu.:194.2   1st Qu.:195.2   1st Qu.:185.7   1st Qu.:2717  
##  Median :205.4   Median :206.7   Median :197.8   Median :3379  
##  Mean   :204.6   Mean   :205.6   Mean   :196.7   Mean   :3426  
##  3rd Qu.:214.5   3rd Qu.:215.6   3rd Qu.:207.2   3rd Qu.:4227  
##  Max.   :237.8   Max.   :237.8   Max.   :231.1   Max.   :5029  
##      bio13           bio14            bio15           bio16       
##  Min.   :150.0   Min.   : 81.53   Min.   :13.31   Min.   : 434.3  
##  1st Qu.:315.6   1st Qu.:147.50   1st Qu.:23.93   1st Qu.: 910.5  
##  Median :401.3   Median :186.50   Median :26.70   Median :1150.2  
##  Mean   :407.8   Mean   :195.30   Mean   :26.04   Mean   :1160.6  
##  3rd Qu.:506.7   3rd Qu.:247.24   3rd Qu.:29.25   3rd Qu.:1423.9  
##  Max.   :659.2   Max.   :300.94   Max.   :33.91   Max.   :1806.9  
##      bio17           bio18            bio19       
##  Min.   :275.6   Min.   : 314.7   Min.   : 328.5  
##  1st Qu.:480.8   1st Qu.: 507.3   1st Qu.: 867.1  
##  Median :594.4   Median : 637.5   Median :1115.4  
##  Mean   :623.4   Mean   : 660.2   Mean   :1114.4  
##  3rd Qu.:773.5   3rd Qu.: 814.2   3rd Qu.:1378.7  
##  Max.   :972.0   Max.   :1028.3   Max.   :1726.5
```

```
summary(Magn_biovars50A)
```

```
##       bio1            bio2             bio3            bio4      
##  Min.   :155.6   Min.   : 98.57   Min.   :87.41   Min.   :243.8  
##  1st Qu.:197.8   1st Qu.:111.22   1st Qu.:89.19   1st Qu.:330.7  
##  Median :209.2   Median :114.76   Median :89.79   Median :376.6  
##  Mean   :208.1   Mean   :114.84   Mean   :89.60   Mean   :372.4  
##  3rd Qu.:218.2   3rd Qu.:118.49   3rd Qu.:90.11   3rd Qu.:417.4  
##  Max.   :240.6   Max.   :125.67   Max.   :91.53   Max.   :531.2  
##       bio5            bio6            bio7            bio8      
##  Min.   :215.3   Min.   :104.1   Min.   :111.2   Min.   :155.3  
##  1st Qu.:265.5   1st Qu.:132.6   1st Qu.:124.8   1st Qu.:194.6  
##  Median :275.0   Median :146.4   Median :128.3   Median :205.1  
##  Mean   :273.8   Mean   :145.6   Mean   :128.2   Mean   :204.2  
##  3rd Qu.:282.7   3rd Qu.:157.5   3rd Qu.:131.9   3rd Qu.:213.9  
##  Max.   :301.3   Max.   :184.2   Max.   :138.0   Max.   :236.8  
##       bio9           bio10           bio11           bio12     
##  Min.   :157.8   Min.   :159.3   Min.   :148.5   Min.   :1310  
##  1st Qu.:199.9   1st Qu.:201.3   1st Qu.:191.5   1st Qu.:2580  
##  Median :211.6   Median :212.9   Median :203.6   Median :3201  
##  Mean   :210.6   Mean   :211.8   Mean   :202.5   Mean   :3256  
##  3rd Qu.:220.4   3rd Qu.:221.8   3rd Qu.:213.1   3rd Qu.:4018  
##  Max.   :243.8   Max.   :243.8   Max.   :236.8   Max.   :4849  
##      bio13           bio14            bio15           bio16       
##  Min.   :141.8   Min.   : 75.41   Min.   :13.02   Min.   : 404.2  
##  1st Qu.:295.1   1st Qu.:142.32   1st Qu.:22.73   1st Qu.: 848.3  
##  Median :373.0   Median :179.77   Median :25.79   Median :1071.1  
##  Mean   :380.0   Mean   :186.67   Mean   :25.21   Mean   :1084.7  
##  3rd Qu.:464.9   3rd Qu.:233.83   3rd Qu.:28.53   3rd Qu.:1328.8  
##  Max.   :639.3   Max.   :289.81   Max.   :33.18   Max.   :1723.4  
##      bio17           bio18           bio19       
##  Min.   :262.8   Min.   :307.8   Min.   : 310.5  
##  1st Qu.:460.4   1st Qu.:484.9   1st Qu.: 816.3  
##  Median :568.3   Median :602.5   Median :1050.6  
##  Mean   :595.2   Mean   :629.9   Mean   :1049.7  
##  3rd Qu.:737.8   3rd Qu.:781.6   3rd Qu.:1296.6  
##  Max.   :922.3   Max.   :987.9   Max.   :1651.0
```

```
summary(Magn_biovars70)
```

```
##       bio1            bio2             bio3            bio4      
##  Min.   :154.4   Min.   : 97.66   Min.   :87.65   Min.   :225.2  
##  1st Qu.:196.6   1st Qu.:110.56   1st Qu.:88.99   1st Qu.:311.3  
##  Median :207.9   Median :114.03   Median :89.42   Median :351.8  
##  Mean   :206.9   Mean   :114.15   Mean   :89.42   Mean   :349.4  
##  3rd Qu.:216.9   3rd Qu.:117.76   3rd Qu.:89.82   3rd Qu.:389.7  
##  Max.   :239.5   Max.   :124.65   Max.   :91.45   Max.   :509.5  
##       bio5            bio6            bio7            bio8      
##  Min.   :213.3   Min.   :102.7   Min.   :110.6   Min.   :150.3  
##  1st Qu.:264.5   1st Qu.:131.8   1st Qu.:124.0   1st Qu.:193.1  
##  Median :273.6   Median :145.7   Median :127.8   Median :204.3  
##  Mean   :272.6   Mean   :145.0   Mean   :127.7   Mean   :203.2  
##  3rd Qu.:281.3   3rd Qu.:156.7   3rd Qu.:131.7   3rd Qu.:213.3  
##  Max.   :300.3   Max.   :183.7   Max.   :137.0   Max.   :236.2  
##       bio9           bio10           bio11           bio12     
##  Min.   :156.0   Min.   :158.0   Min.   :147.7   Min.   :1315  
##  1st Qu.:197.9   1st Qu.:199.9   1st Qu.:190.8   1st Qu.:2616  
##  Median :209.8   Median :211.6   Median :202.8   Median :3255  
##  Mean   :208.8   Mean   :210.5   Mean   :201.8   Mean   :3301  
##  3rd Qu.:219.0   3rd Qu.:220.5   3rd Qu.:212.4   3rd Qu.:4077  
##  Max.   :242.8   Max.   :242.8   Max.   :236.1   Max.   :4896  
##      bio13           bio14            bio15           bio16       
##  Min.   :140.2   Min.   : 76.48   Min.   :14.58   Min.   : 400.0  
##  1st Qu.:318.1   1st Qu.:142.80   1st Qu.:25.59   1st Qu.: 901.9  
##  Median :403.5   Median :181.01   Median :28.69   Median :1139.2  
##  Mean   :408.7   Mean   :187.26   Mean   :27.95   Mean   :1149.1  
##  3rd Qu.:506.2   3rd Qu.:234.32   3rd Qu.:31.40   3rd Qu.:1408.7  
##  Max.   :669.2   Max.   :288.83   Max.   :36.28   Max.   :1802.9  
##      bio17           bio18           bio19       
##  Min.   :260.7   Min.   :301.6   Min.   : 332.5  
##  1st Qu.:459.2   1st Qu.:505.9   1st Qu.: 871.8  
##  Median :569.1   Median :620.1   Median :1118.7  
##  Mean   :595.3   Mean   :647.9   Mean   :1118.8  
##  3rd Qu.:736.3   3rd Qu.:798.2   3rd Qu.:1382.8  
##  Max.   :932.2   Max.   :999.7   Max.   :1742.5
```

```
summary(Magn_biovars70A)
```

```
##       bio1            bio2             bio3            bio4      
##  Min.   :168.1   Min.   : 98.99   Min.   :87.41   Min.   :223.9  
##  1st Qu.:210.5   1st Qu.:111.37   1st Qu.:88.73   1st Qu.:275.9  
##  Median :221.9   Median :114.96   Median :89.21   Median :311.3  
##  Mean   :220.8   Mean   :115.03   Mean   :89.17   Mean   :307.5  
##  3rd Qu.:231.0   3rd Qu.:118.67   3rd Qu.:89.59   3rd Qu.:337.4  
##  Max.   :253.4   Max.   :126.22   Max.   :91.14   Max.   :438.1  
##       bio5            bio6            bio7            bio8      
##  Min.   :228.7   Min.   :115.7   Min.   :113.0   Min.   :168.2  
##  1st Qu.:280.4   1st Qu.:146.6   1st Qu.:125.5   1st Qu.:207.5  
##  Median :289.6   Median :160.4   Median :129.1   Median :218.7  
##  Mean   :288.6   Mean   :159.6   Mean   :129.0   Mean   :217.6  
##  3rd Qu.:297.2   3rd Qu.:171.3   3rd Qu.:132.9   3rd Qu.:227.6  
##  Max.   :316.3   Max.   :198.2   Max.   :139.0   Max.   :250.7  
##       bio9           bio10           bio11           bio12     
##  Min.   :169.0   Min.   :171.1   Min.   :162.7   Min.   :1296  
##  1st Qu.:212.7   1st Qu.:213.7   1st Qu.:206.4   1st Qu.:2627  
##  Median :223.8   Median :225.5   Median :218.3   Median :3293  
##  Mean   :223.1   Mean   :224.4   Mean   :217.0   Mean   :3349  
##  3rd Qu.:233.2   3rd Qu.:234.6   3rd Qu.:227.5   3rd Qu.:4139  
##  Max.   :256.8   Max.   :256.8   Max.   :250.7   Max.   :5071  
##      bio13           bio14            bio15           bio16       
##  Min.   :141.5   Min.   : 73.42   Min.   :17.76   Min.   : 409.0  
##  1st Qu.:343.4   1st Qu.:130.90   1st Qu.:29.80   1st Qu.: 931.7  
##  Median :439.6   Median :157.28   Median :33.02   Median :1198.9  
##  Mean   :445.4   Mean   :163.23   Mean   :32.32   Mean   :1204.4  
##  3rd Qu.:552.2   3rd Qu.:198.35   3rd Qu.:35.52   3rd Qu.:1486.8  
##  Max.   :738.0   Max.   :257.21   Max.   :39.94   Max.   :1916.7  
##      bio17           bio18           bio19       
##  Min.   :259.8   Min.   :288.1   Min.   : 341.4  
##  1st Qu.:439.1   1st Qu.:477.6   1st Qu.: 915.0  
##  Median :541.2   Median :587.4   Median :1185.3  
##  Mean   :572.0   Mean   :613.0   Mean   :1188.0  
##  3rd Qu.:722.2   3rd Qu.:760.4   3rd Qu.:1470.2  
##  Max.   :876.2   Max.   :978.9   Max.   :1916.7
```

```
boxplot(Magn_biovars00[,'bio1'], Magn_biovars50[,'bio1'], Magn_biovars50A[,'bio1'], Magn_biovars70[,'bio1'], Magn_biovars70A[,'bio1'], 
        boxfill = c('lightgreen','coral','red3','red4','tan4'),
        names = c('1970-2000','2050 RCP4.5','2050 RCP8.5','2070 RCP4.5','2070 RCP8.5'),
        main = 'BIO 1')
boxplot(Magn_biovars00[,'bio2'], Magn_biovars50[,'bio2'], Magn_biovars50A[,'bio2'], Magn_biovars70[,'bio2'], Magn_biovars70A[,'bio2'], 
        boxfill = c('lightgreen','coral','red3','red4','tan4'),
        names = c('1970-2000','2050 RCP4.5','2050 RCP8.5','2070 RCP4.5','2070 RCP8.5'),
        main = 'BIO 2')
boxplot(Magn_biovars00[,'bio3'], Magn_biovars50[,'bio3'], Magn_biovars50A[,'bio3'], Magn_biovars70[,'bio3'], Magn_biovars70A[,'bio3'], 
        boxfill = c('lightgreen','coral','red3','red4','tan4'),
        names = c('1970-2000','2050 RCP4.5','2050 RCP8.5','2070 RCP4.5','2070 RCP8.5'),
        main = 'BIO 3')
boxplot(Magn_biovars00[,'bio4'], Magn_biovars50[,'bio4'], Magn_biovars50A[,'bio4'], Magn_biovars70[,'bio4'], Magn_biovars70A[,'bio4'], 
        boxfill = c('lightgreen','coral','red3','red4','tan4'),
        names = c('1970-2000','2050 RCP4.5','2050 RCP8.5','2070 RCP4.5','2070 RCP8.5'),
        main = 'BIO 4')
```

```
boxplot(Magn_biovars00[,'bio5'], Magn_biovars50[,'bio5'], Magn_biovars50A[,'bio5'], Magn_biovars70[,'bio5'], Magn_biovars70A[,'bio5'], 
        boxfill = c('lightgreen','coral','red3','red4','tan4'),
        names = c('1970-2000','2050 RCP4.5','2050 RCP8.5','2070 RCP4.5','2070 RCP8.5'),
        main = 'BIO 5')
boxplot(Magn_biovars00[,'bio6'], Magn_biovars50[,'bio6'], Magn_biovars50A[,'bio6'], Magn_biovars70[,'bio6'], Magn_biovars70A[,'bio6'], 
        boxfill = c('lightgreen','coral','red3','red4','tan4'),
        names = c('1970-2000','2050 RCP4.5','2050 RCP8.5','2070 RCP4.5','2070 RCP8.5'),
        main = 'BIO 6')
boxplot(Magn_biovars00[,'bio7'], Magn_biovars50[,'bio7'], Magn_biovars50A[,'bio7'], Magn_biovars70[,'bio7'], Magn_biovars70A[,'bio7'], 
        boxfill = c('lightgreen','coral','red3','red4','tan4'),
        names = c('1970-2000','2050 RCP4.5','2050 RCP8.5','2070 RCP4.5','2070 RCP8.5'),
        main = 'BIO 7')
boxplot(Magn_biovars00[,'bio8'], Magn_biovars50[,'bio8'], Magn_biovars50A[,'bio8'], Magn_biovars70[,'bio8'], Magn_biovars70A[,'bio8'], 
        boxfill = c('lightgreen','coral','red3','red4','tan4'),
        names = c('1970-2000','2050 RCP4.5','2050 RCP8.5','2070 RCP4.5','2070 RCP8.5'),
        main = 'BIO 8')
```

```
boxplot(Magn_biovars00[,'bio9'], Magn_biovars50[,'bio9'], Magn_biovars50A[,'bio9'], Magn_biovars70[,'bio9'], Magn_biovars70A[,'bio9'], 
        boxfill = c('lightgreen','coral','red3','red4','tan4'),
        names = c('1970-2000','2050 RCP4.5','2050 RCP8.5','2070 RCP4.5','2070 RCP8.5'),
        main = 'BIO 9')
boxplot(Magn_biovars00[,'bio10'], Magn_biovars50[,'bio10'], Magn_biovars50A[,'bio10'], Magn_biovars70[,'bio10'], Magn_biovars70A[,'bio10'], 
        boxfill = c('lightgreen','coral','red3','red4','tan4'),
        names = c('1970-2000','2050 RCP4.5','2050 RCP8.5','2070 RCP4.5','2070 RCP8.5'),
        main = 'BIO 10')
boxplot(Magn_biovars00[,'bio11'], Magn_biovars50[,'bio11'], Magn_biovars50A[,'bio11'], Magn_biovars70[,'bio11'], Magn_biovars70A[,'bio11'], 
        boxfill = c('lightgreen','coral','red3','red4','tan4'),
        names = c('1970-2000','2050 RCP4.5','2050 RCP8.5','2070 RCP4.5','2070 RCP8.5'),
        main = 'BIO 11')
boxplot(Magn_biovars00[,'bio12'], Magn_biovars50[,'bio12'], Magn_biovars50A[,'bio12'], Magn_biovars70[,'bio12'], Magn_biovars70A[,'bio12'], 
        boxfill = c('lightgreen','coral','red3','red4','tan4'),
        names = c('1970-2000','2050 RCP4.5','2050 RCP8.5','2070 RCP4.5','2070 RCP8.5'),
        main = 'BIO 12')
```

```
boxplot(Magn_biovars00[,'bio13'], Magn_biovars50[,'bio13'], Magn_biovars50A[,'bio13'], Magn_biovars70[,'bio13'], Magn_biovars70A[,'bio13'], 
        boxfill = c('lightgreen','coral','red3','red4','tan4'),
        names = c('1970-2000','2050 RCP4.5','2050 RCP8.5','2070 RCP4.5','2070 RCP8.5'),
        main = 'BIO 13')
boxplot(Magn_biovars00[,'bio14'], Magn_biovars50[,'bio14'], Magn_biovars50A[,'bio14'], Magn_biovars70[,'bio14'], Magn_biovars70A[,'bio14'], 
        boxfill = c('lightgreen','coral','red3','red4','tan4'),
        names = c('1970-2000','2050 RCP4.5','2050 RCP8.5','2070 RCP4.5','2070 RCP8.5'),
        main = 'BIO 14')
boxplot(Magn_biovars00[,'bio15'], Magn_biovars50[,'bio15'], Magn_biovars50A[,'bio15'], Magn_biovars70[,'bio15'], Magn_biovars70A[,'bio15'], 
        boxfill = c('lightgreen','coral','red3','red4','tan4'),
        names = c('1970-2000','2050 RCP4.5','2050 RCP8.5','2070 RCP4.5','2070 RCP8.5'),
        main = 'BIO 15')
boxplot(Magn_biovars00[,'bio16'], Magn_biovars50[,'bio16'], Magn_biovars50A[,'bio16'], Magn_biovars70[,'bio16'], Magn_biovars70A[,'bio16'], 
        boxfill = c('lightgreen','coral','red3','red4','tan4'),
        names = c('1970-2000','2050 RCP4.5','2050 RCP8.5','2070 RCP4.5','2070 RCP8.5'),
        main = 'BIO 16')
```

```
boxplot(Magn_biovars00[,'bio17'], Magn_biovars50[,'bio17'], Magn_biovars50A[,'bio17'], Magn_biovars70[,'bio17'], Magn_biovars70A[,'bio17'], 
        boxfill = c('lightgreen','coral','red3','red4','tan4'),
        names = c('1970-2000','2050 RCP4.5','2050 RCP8.5','2070 RCP4.5','2070 RCP8.5'),
        main = 'BIO 17')
boxplot(Magn_biovars00[,'bio18'], Magn_biovars50[,'bio18'], Magn_biovars50A[,'bio18'], Magn_biovars70[,'bio18'], Magn_biovars70A[,'bio18'], 
        boxfill = c('lightgreen','coral','red3','red4','tan4'),
        names = c('1970-2000','2050 RCP4.5','2050 RCP8.5','2070 RCP4.5','2070 RCP8.5'),
        main = 'BIO 18')
boxplot(Magn_biovars00[,'bio19'], Magn_biovars50[,'bio19'], Magn_biovars50A[,'bio19'], Magn_biovars70[,'bio19'], Magn_biovars70A[,'bio19'], 
        boxfill = c('lightgreen','coral','red3','red4','tan4'),
        names = c('1970-2000','2050 RCP4.5','2050 RCP8.5','2070 RCP4.5','2070 RCP8.5'),
        main = 'BIO 19')
```

###### Mann-Whitney U tests

```
# true-false H0 vs. HA matrices
MWU_prec_RCP45_matrix <- matrix(nrow = 12, ncol = 3)
MWU_prec_RCP85_matrix <- matrix(nrow = 12, ncol = 3)
MWU_tmin_RCP45_matrix <- matrix(nrow = 12, ncol = 3)
MWU_tmin_RCP85_matrix <- matrix(nrow = 12, ncol = 3)
MWU_tmax_RCP45_matrix <- matrix(nrow = 12, ncol = 3)
MWU_tmax_RCP85_matrix <- matrix(nrow = 12, ncol = 3)
MWU_biovars_RCP45_matrix <- matrix(nrow = 19, ncol = 3)
MWU_biovars_RCP85_matrix <- matrix(nrow = 19, ncol = 3)

# location shift estimation matrices
MWU_prec_RCP45_matrix_s <- matrix(nrow = 12, ncol = 3)
MWU_prec_RCP85_matrix_s <- matrix(nrow = 12, ncol = 3)
MWU_tmin_RCP45_matrix_s <- matrix(nrow = 12, ncol = 3)
MWU_tmin_RCP85_matrix_s <- matrix(nrow = 12, ncol = 3)
MWU_tmax_RCP45_matrix_s <- matrix(nrow = 12, ncol = 3)
MWU_tmax_RCP85_matrix_s <- matrix(nrow = 12, ncol = 3)
MWU_biovars_RCP45_matrix_s <- matrix(nrow = 19, ncol = 3)
MWU_biovars_RCP85_matrix_s <- matrix(nrow = 19, ncol = 3)


# Mann-Whitney test for prec, tmin, tmax
for (i in 1:12) {
  my_test1 <- compareMWT(Magn_prec00[,i],Magn_prec50[,i],paste(month_list[i],'precipitation: actual vs. RCP4.5 2050'))
  my_test2 <- compareMWT(Magn_prec50[,i],Magn_prec70[,i],paste(month_list[i],'precipitation: RCP4.5 2050 vs. RCP4.5 2070'))
  my_test3 <- compareMWT(Magn_prec00[,i],Magn_prec70[,i],paste(month_list[i],'precipitation: actual vs. RCP4.5 2070'))

  my_test4 <- compareMWT(Magn_prec00[,i],Magn_prec50A[,i],paste(month_list[i],'precipitation: actual vs. RCP8.5 2050'))
  my_test5 <- compareMWT(Magn_prec50A[,i],Magn_prec70A[,i],paste(month_list[i],'precipitation: RCP8.5 2050 vs. RCP8.5 2070'))
  my_test6 <- compareMWT(Magn_prec00[,i],Magn_prec70A[,i],paste(month_list[i],'precipitation: actual vs. RCP8.5 2070'))
 
  my_test7 <- compareMWT(Magn_tmin00[,i],Magn_tmin50[,i],paste(month_list[i],'tmin: actual vs. RCP4.5 2050'))
  my_test8 <- compareMWT(Magn_tmin50[,i],Magn_tmin70[,i],paste(month_list[i],'tmin: RCP4.5 2050 vs. RCP4.5 2070'))
  my_test9 <- compareMWT(Magn_tmin00[,i],Magn_tmin70[,i],paste(month_list[i],'tmin: actual vs. RCP4.5 2070'))

  my_test10 <- compareMWT(Magn_tmin00[,i],Magn_tmin50A[,i],paste(month_list[i],'tmin: actual vs. RCP8.5 2050'))
  my_test11 <- compareMWT(Magn_tmin50A[,i],Magn_tmin70A[,i],paste(month_list[i],'tmin: RCP8.5 2050 vs. RCP8.5 2070'))
  my_test12 <- compareMWT(Magn_tmin00[,i],Magn_tmin70A[,i],paste(month_list[i],'tmin: actual vs. RCP8.5 2070'))   
  
  my_test13 <- compareMWT(Magn_tmax00[,i],Magn_tmax50[,i],paste(month_list[i],'tmax: actual vs. RCP4.5 2050'))
  my_test14 <- compareMWT(Magn_tmax50[,i],Magn_tmax70[,i],paste(month_list[i],'tmax: RCP4.5 2050 vs. RCP4.5 2070'))
  my_test15 <- compareMWT(Magn_tmax00[,i],Magn_tmax70[,i],paste(month_list[i],'tmax: actual vs. RCP4.5 2070'))

  my_test16 <- compareMWT(Magn_tmax00[,i],Magn_tmax50A[,i],paste(month_list[i],'tmax: actual vs. RCP8.5 2050'))
  my_test17 <- compareMWT(Magn_tmax50A[,i],Magn_tmax70A[,i],paste(month_list[i],'tmax: RCP8.5 2050 vs. RCP8.5 2070'))
  my_test18 <- compareMWT(Magn_tmax00[,i],Magn_tmax70A[,i],paste(month_list[i],'tmax: actual vs. RCP8.5 2070'))   
  
  if (my_test1$p.value < 0.05) {
    MWU_prec_RCP45_matrix[i,1] <- 1
    MWU_prec_RCP45_matrix_s[i,1] <- my_test1$estimate
  } else {
    MWU_prec_RCP45_matrix[i,1] <- 0
  }
  if (my_test2$p.value < 0.05) {
    MWU_prec_RCP45_matrix[i,2] <- 1
    MWU_prec_RCP45_matrix_s[i,2] <- my_test2$estimate    
  } else {
    MWU_prec_RCP45_matrix[i,2] <- 0
  }
  if (my_test3$p.value < 0.05) {
    MWU_prec_RCP45_matrix[i,3] <- 1
    MWU_prec_RCP45_matrix_s[i,3] <- my_test3$estimate    
  } else {
    MWU_prec_RCP45_matrix[i,3] <- 0
  }
  if (my_test4$p.value < 0.05) {
    MWU_prec_RCP85_matrix[i,1] <- 1
    MWU_prec_RCP85_matrix_s[i,1] <- my_test4$estimate    
  } else {
    MWU_prec_RCP85_matrix[i,1] <- 0
  }
  if (my_test5$p.value < 0.05) {
    MWU_prec_RCP85_matrix[i,2] <- 1
    MWU_prec_RCP85_matrix_s[i,2] <- my_test5$estimate      
  } else {
    MWU_prec_RCP85_matrix[i,2] <- 0
  }
  if (my_test6$p.value < 0.05) {
    MWU_prec_RCP85_matrix[i,3] <- 1
    MWU_prec_RCP85_matrix_s[i,3] <- my_test6$estimate          
  } else {
    MWU_prec_RCP85_matrix[i,3] <- 0
  }
  
  if (my_test7$p.value < 0.05) {
    MWU_tmin_RCP45_matrix[i,1] <- 1
    MWU_tmin_RCP45_matrix_s[i,1] <- my_test7$estimate          
  } else {
    MWU_tmin_RCP45_matrix[i,1] <- 0
  }
  if (my_test8$p.value < 0.05) {
    MWU_tmin_RCP45_matrix[i,2] <- 1
    MWU_tmin_RCP45_matrix_s[i,2] <- my_test8$estimate       
  } else {
    MWU_tmin_RCP45_matrix[i,2] <- 0
  }
  if (my_test9$p.value < 0.05) {
    MWU_tmin_RCP45_matrix[i,3] <- 1
    MWU_tmin_RCP45_matrix_s[i,3] <- my_test9$estimate       
  } else {
    MWU_tmin_RCP45_matrix[i,3] <- 0
  }
  if (my_test10$p.value < 0.05) {
    MWU_tmin_RCP85_matrix[i,1] <- 1
    MWU_tmin_RCP85_matrix_s[i,1] <- my_test10$estimate         
  } else {
    MWU_tmin_RCP85_matrix[i,1] <- 0
  }
  if (my_test11$p.value < 0.05) {
    MWU_tmin_RCP85_matrix[i,2] <- 1
    MWU_tmin_RCP85_matrix_s[i,2] <- my_test11$estimate         
  } else {
    MWU_tmin_RCP85_matrix[i,2] <- 0
  }
  if (my_test12$p.value < 0.05) {
    MWU_tmin_RCP85_matrix[i,3] <- 1
    MWU_tmin_RCP85_matrix_s[i,3] <- my_test12$estimate         
  } else {
    MWU_tmin_RCP85_matrix[i,3] <- 0
  }

  if (my_test13$p.value < 0.05) {
    MWU_tmax_RCP45_matrix[i,1] <- 1
    MWU_tmax_RCP45_matrix_s[i,1] <- my_test13$estimate       
  } else {
    MWU_tmax_RCP45_matrix[i,1] <- 0
  }
  if (my_test14$p.value < 0.05) {
    MWU_tmax_RCP45_matrix[i,2] <- 1
    MWU_tmax_RCP45_matrix_s[i,2] <- my_test14$estimate    
  } else {
    MWU_tmax_RCP45_matrix[i,2] <- 0
  }
  if (my_test15$p.value < 0.05) {
    MWU_tmax_RCP45_matrix[i,3] <- 1
    MWU_tmax_RCP45_matrix_s[i,3] <- my_test15$estimate    
  } else {
    MWU_tmax_RCP45_matrix[i,3] <- 0
  }
  if (my_test16$p.value < 0.05) {
    MWU_tmax_RCP85_matrix[i,1] <- 1
    MWU_tmax_RCP85_matrix_s[i,1] <- my_test16$estimate        
  } else {
    MWU_tmax_RCP85_matrix[i,1] <- 0
  }
  if (my_test17$p.value < 0.05) {
    MWU_tmax_RCP85_matrix[i,2] <- 1
    MWU_tmax_RCP85_matrix_s[i,2] <- my_test17$estimate       
  } else {
    MWU_tmax_RCP85_matrix[i,2] <- 0
  }
  if (my_test18$p.value < 0.05) {
    MWU_tmax_RCP85_matrix[i,3] <- 1
    MWU_tmax_RCP85_matrix_s[i,3] <- my_test18$estimate       
  } else {
    MWU_tmax_RCP85_matrix[i,3] <- 0
  }
  #print(str(my_test1))
}
```

```
## Mann-Whitney test results for Jan precipitation: actual vs. RCP4.5 2050 
## The H0 is rejected as p < 0.05, the HA accepted (medians unequal) 
## 
##  Wilcoxon rank sum test with continuity correction
## 
## data:  x and y
## W = 189380, p-value < 2.2e-16
## alternative hypothesis: true location shift is not equal to 0
## 95 percent confidence interval:
##  -68.14878 -57.55221
## sample estimates:
## difference in location 
##              -62.82329 
## 
## Mann-Whitney test results for Jan precipitation: RCP4.5 2050 vs. RCP4.5 2070 
## The H0 is rejected as p < 0.05, the HA accepted (medians unequal) 
## 
##  Wilcoxon rank sum test with continuity correction
## 
## data:  x and y
## W = 560120, p-value = 3.227e-06
## alternative hypothesis: true location shift is not equal to 0
## 95 percent confidence interval:
##   7.591309 18.261733
## sample estimates:
## difference in location 
##               12.91624 
## 
## Mann-Whitney test results for Jan precipitation: actual vs. RCP4.5 2070 
## The H0 is rejected as p < 0.05, the HA accepted (medians unequal) 
## 
##  Wilcoxon rank sum test with continuity correction
## 
## data:  x and y
## W = 238410, p-value < 2.2e-16
## alternative hypothesis: true location shift is not equal to 0
## 95 percent confidence interval:
##  -54.76497 -44.84815
## sample estimates:
## difference in location 
##              -49.78519 
## 
## Mann-Whitney test results for Jan precipitation: actual vs. RCP8.5 2050 
## The H0 is rejected as p < 0.05, the HA accepted (medians unequal) 
## 
##  Wilcoxon rank sum test with continuity correction
## 
## data:  x and y
## W = 248660, p-value < 2.2e-16
## alternative hypothesis: true location shift is not equal to 0
## 95 percent confidence interval:
##  -52.20628 -42.41095
## sample estimates:
## difference in location 
##              -47.27904 
## 
## Mann-Whitney test results for Jan precipitation: RCP8.5 2050 vs. RCP8.5 2070 
## The HA is rejected as p >= 0.05, the H0 accepted (medians equal) 
## 
##  Wilcoxon rank sum test with continuity correction
## 
## data:  x and y
## W = 497340, p-value = 0.8366
## alternative hypothesis: true location shift is not equal to 0
## 95 percent confidence interval:
##  -5.697823  4.609907
## sample estimates:
## difference in location 
##             -0.5405613 
## 
## Mann-Whitney test results for Jan precipitation: actual vs. RCP8.5 2070 
## The H0 is rejected as p < 0.05, the HA accepted (medians unequal) 
## 
##  Wilcoxon rank sum test with continuity correction
## 
## data:  x and y
## W = 250770, p-value < 2.2e-16
## alternative hypothesis: true location shift is not equal to 0
## 95 percent confidence interval:
##  -52.58994 -42.55239
## sample estimates:
## difference in location 
##              -47.56407 
## 
## Mann-Whitney test results for Jan tmin: actual vs. RCP4.5 2050 
## The H0 is rejected as p < 0.05, the HA accepted (medians unequal) 
## 
##  Wilcoxon rank sum test with continuity correction
## 
## data:  x and y
## W = 302350, p-value < 2.2e-16
## alternative hypothesis: true location shift is not equal to 0
## 95 percent confidence interval:
##  -1.478626 -1.155824
## sample estimates:
## difference in location 
##              -1.317149 
## 
## Mann-Whitney test results for Jan tmin: RCP4.5 2050 vs. RCP4.5 2070 
## The H0 is rejected as p < 0.05, the HA accepted (medians unequal) 
## 
##  Wilcoxon rank sum test with continuity correction
## 
## data:  x and y
## W = 440640, p-value = 4.296e-06
## alternative hypothesis: true location shift is not equal to 0
## 95 percent confidence interval:
##  -0.5128313 -0.2097363
## sample estimates:
## difference in location 
##             -0.3607165 
## 
## Mann-Whitney test results for Jan tmin: actual vs. RCP4.5 2070 
## The H0 is rejected as p < 0.05, the HA accepted (medians unequal) 
## 
##  Wilcoxon rank sum test with continuity correction
## 
## data:  x and y
## W = 256530, p-value < 2.2e-16
## alternative hypothesis: true location shift is not equal to 0
## 95 percent confidence interval:
##  -1.841747 -1.517226
## sample estimates:
## difference in location 
##              -1.679153 
## 
## Mann-Whitney test results for Jan tmin: actual vs. RCP8.5 2050 
## The H0 is rejected as p < 0.05, the HA accepted (medians unequal) 
## 
##  Wilcoxon rank sum test with continuity correction
## 
## data:  x and y
## W = 236050, p-value < 2.2e-16
## alternative hypothesis: true location shift is not equal to 0
## 95 percent confidence interval:
##  -2.005779 -1.681098
## sample estimates:
## difference in location 
##              -1.843089 
## 
## Mann-Whitney test results for Jan tmin: RCP8.5 2050 vs. RCP8.5 2070 
## The H0 is rejected as p < 0.05, the HA accepted (medians unequal) 
## 
##  Wilcoxon rank sum test with continuity correction
## 
## data:  x and y
## W = 316570, p-value < 2.2e-16
## alternative hypothesis: true location shift is not equal to 0
## 95 percent confidence interval:
##  -1.326985 -1.021958
## sample estimates:
## difference in location 
##              -1.174802 
## 
## Mann-Whitney test results for Jan tmin: actual vs. RCP8.5 2070 
## The H0 is rejected as p < 0.05, the HA accepted (medians unequal) 
## 
##  Wilcoxon rank sum test with continuity correction
## 
## data:  x and y
## W = 114730, p-value < 2.2e-16
## alternative hypothesis: true location shift is not equal to 0
## 95 percent confidence interval:
##  -3.180730 -2.855781
## sample estimates:
## difference in location 
##              -3.018058 
## 
## Mann-Whitney test results for Jan tmax: actual vs. RCP4.5 2050 
## The H0 is rejected as p < 0.05, the HA accepted (medians unequal) 
## 
##  Wilcoxon rank sum test with continuity correction
## 
## data:  x and y
## W = 23783, p-value < 2.2e-16
## alternative hypothesis: true location shift is not equal to 0
## 95 percent confidence interval:
##  -3.757739 -3.528780
## sample estimates:
## difference in location 
##              -3.643493 
## 
## Mann-Whitney test results for Jan tmax: RCP4.5 2050 vs. RCP4.5 2070 
## The H0 is rejected as p < 0.05, the HA accepted (medians unequal) 
## 
##  Wilcoxon rank sum test with continuity correction
## 
## data:  x and y
## W = 400350, p-value = 1.191e-14
## alternative hypothesis: true location shift is not equal to 0
## 95 percent confidence interval:
##  -0.5084337 -0.3070042
## sample estimates:
## difference in location 
##             -0.4061376 
## 
## Mann-Whitney test results for Jan tmax: actual vs. RCP4.5 2070 
## The H0 is rejected as p < 0.05, the HA accepted (medians unequal) 
## 
##  Wilcoxon rank sum test with continuity correction
## 
## data:  x and y
## W = 13731, p-value < 2.2e-16
## alternative hypothesis: true location shift is not equal to 0
## 95 percent confidence interval:
##  -4.165872 -3.935889
## sample estimates:
## difference in location 
##              -4.051103 
## 
## Mann-Whitney test results for Jan tmax: actual vs. RCP8.5 2050 
## The H0 is rejected as p < 0.05, the HA accepted (medians unequal) 
## 
##  Wilcoxon rank sum test with continuity correction
## 
## data:  x and y
## W = 8833, p-value < 2.2e-16
## alternative hypothesis: true location shift is not equal to 0
## 95 percent confidence interval:
##  -4.457739 -4.228780
## sample estimates:
## difference in location 
##              -4.343493 
## 
## Mann-Whitney test results for Jan tmax: RCP8.5 2050 vs. RCP8.5 2070 
## The H0 is rejected as p < 0.05, the HA accepted (medians unequal) 
## 
##  Wilcoxon rank sum test with continuity correction
## 
## data:  x and y
## W = 244030, p-value < 2.2e-16
## alternative hypothesis: true location shift is not equal to 0
## 95 percent confidence interval:
##  -1.241125 -1.039759
## sample estimates:
## difference in location 
##              -1.140169 
## 
## Mann-Whitney test results for Jan tmax: actual vs. RCP8.5 2070 
## The H0 is rejected as p < 0.05, the HA accepted (medians unequal) 
## 
##  Wilcoxon rank sum test with continuity correction
## 
## data:  x and y
## W = 1560, p-value < 2.2e-16
## alternative hypothesis: true location shift is not equal to 0
## 95 percent confidence interval:
##  -5.596633 -5.368047
## sample estimates:
## difference in location 
##              -5.482635 
## 
## Mann-Whitney test results for Feb precipitation: actual vs. RCP4.5 2050 
## The H0 is rejected as p < 0.05, the HA accepted (medians unequal) 
## 
##  Wilcoxon rank sum test with continuity correction
## 
## data:  x and y
## W = 221180, p-value < 2.2e-16
## alternative hypothesis: true location shift is not equal to 0
## 95 percent confidence interval:
##  -58.48613 -48.78493
## sample estimates:
## difference in location 
##              -53.63743 
## 
## Mann-Whitney test results for Feb precipitation: RCP4.5 2050 vs. RCP4.5 2070 
## The H0 is rejected as p < 0.05, the HA accepted (medians unequal) 
## 
##  Wilcoxon rank sum test with continuity correction
## 
## data:  x and y
## W = 602140, p-value = 2.583e-15
## alternative hypothesis: true location shift is not equal to 0
## 95 percent confidence interval:
##  15.20754 25.00944
## sample estimates:
## difference in location 
##               20.12772 
## 
## Mann-Whitney test results for Feb precipitation: actual vs. RCP4.5 2070 
## The H0 is rejected as p < 0.05, the HA accepted (medians unequal) 
## 
##  Wilcoxon rank sum test with continuity correction
## 
## data:  x and y
## W = 305960, p-value < 2.2e-16
## alternative hypothesis: true location shift is not equal to 0
## 95 percent confidence interval:
##  -37.96808 -29.22503
## sample estimates:
## difference in location 
##              -33.55265 
## 
## Mann-Whitney test results for Feb precipitation: actual vs. RCP8.5 2050 
## The H0 is rejected as p < 0.05, the HA accepted (medians unequal) 
## 
##  Wilcoxon rank sum test with continuity correction
## 
## data:  x and y
## W = 277020, p-value < 2.2e-16
## alternative hypothesis: true location shift is not equal to 0
## 95 percent confidence interval:
##  -44.17432 -35.19337
## sample estimates:
## difference in location 
##              -39.66561 
## 
## Mann-Whitney test results for Feb precipitation: RCP8.5 2050 vs. RCP8.5 2070 
## The HA is rejected as p >= 0.05, the H0 accepted (medians equal) 
## 
##  Wilcoxon rank sum test with continuity correction
## 
## data:  x and y
## W = 511730, p-value = 0.3638
## alternative hypothesis: true location shift is not equal to 0
## 95 percent confidence interval:
##  -2.503462  6.790612
## sample estimates:
## difference in location 
##               2.116774 
## 
## Mann-Whitney test results for Feb precipitation: actual vs. RCP8.5 2070 
## The H0 is rejected as p < 0.05, the HA accepted (medians unequal) 
## 
##  Wilcoxon rank sum test with continuity correction
## 
## data:  x and y
## W = 291860, p-value < 2.2e-16
## alternative hypothesis: true location shift is not equal to 0
## 95 percent confidence interval:
##  -41.85784 -32.74775
## sample estimates:
## difference in location 
##              -37.28469 
## 
## Mann-Whitney test results for Feb tmin: actual vs. RCP4.5 2050 
## The H0 is rejected as p < 0.05, the HA accepted (medians unequal) 
## 
##  Wilcoxon rank sum test with continuity correction
## 
## data:  x and y
## W = 292740, p-value < 2.2e-16
## alternative hypothesis: true location shift is not equal to 0
## 95 percent confidence interval:
##  -1.543120 -1.221364
## sample estimates:
## difference in location 
##              -1.381484 
## 
## Mann-Whitney test results for Feb tmin: RCP4.5 2050 vs. RCP4.5 2070 
## The H0 is rejected as p < 0.05, the HA accepted (medians unequal) 
## 
##  Wilcoxon rank sum test with continuity correction
## 
## data:  x and y
## W = 433040, p-value = 2.159e-07
## alternative hypothesis: true location shift is not equal to 0
## 95 percent confidence interval:
##  -0.5483602 -0.2516362
## sample estimates:
## difference in location 
##             -0.3999992 
## 
## Mann-Whitney test results for Feb tmin: actual vs. RCP4.5 2070 
## The H0 is rejected as p < 0.05, the HA accepted (medians unequal) 
## 
##  Wilcoxon rank sum test with continuity correction
## 
## data:  x and y
## W = 241150, p-value < 2.2e-16
## alternative hypothesis: true location shift is not equal to 0
## 95 percent confidence interval:
##  -1.943119 -1.621369
## sample estimates:
## difference in location 
##              -1.781489 
## 
## Mann-Whitney test results for Feb tmin: actual vs. RCP8.5 2050 
## The H0 is rejected as p < 0.05, the HA accepted (medians unequal) 
## 
##  Wilcoxon rank sum test with continuity correction
## 
## data:  x and y
## W = 220200, p-value < 2.2e-16
## alternative hypothesis: true location shift is not equal to 0
## 95 percent confidence interval:
##  -2.123968 -1.800519
## sample estimates:
## difference in location 
##              -1.961522 
## 
## Mann-Whitney test results for Feb tmin: RCP8.5 2050 vs. RCP8.5 2070 
## The H0 is rejected as p < 0.05, the HA accepted (medians unequal) 
## 
##  Wilcoxon rank sum test with continuity correction
## 
## data:  x and y
## W = 315580, p-value < 2.2e-16
## alternative hypothesis: true location shift is not equal to 0
## 95 percent confidence interval:
##  -1.312457 -1.011996
## sample estimates:
## difference in location 
##              -1.162564 
## 
## Mann-Whitney test results for Feb tmin: actual vs. RCP8.5 2070 
## The H0 is rejected as p < 0.05, the HA accepted (medians unequal) 
## 
##  Wilcoxon rank sum test with continuity correction
## 
## data:  x and y
## W = 104320, p-value < 2.2e-16
## alternative hypothesis: true location shift is not equal to 0
## 95 percent confidence interval:
##  -3.286400 -2.962311
## sample estimates:
## difference in location 
##              -3.124255 
## 
## Mann-Whitney test results for Feb tmax: actual vs. RCP4.5 2050 
## The H0 is rejected as p < 0.05, the HA accepted (medians unequal) 
## 
##  Wilcoxon rank sum test with continuity correction
## 
## data:  x and y
## W = 27779, p-value < 2.2e-16
## alternative hypothesis: true location shift is not equal to 0
## 95 percent confidence interval:
##  -3.680606 -3.450480
## sample estimates:
## difference in location 
##              -3.565342 
## 
## Mann-Whitney test results for Feb tmax: RCP4.5 2050 vs. RCP4.5 2070 
## The H0 is rejected as p < 0.05, the HA accepted (medians unequal) 
## 
##  Wilcoxon rank sum test with continuity correction
## 
## data:  x and y
## W = 380740, p-value < 2.2e-16
## alternative hypothesis: true location shift is not equal to 0
## 95 percent confidence interval:
##  -0.6027017 -0.3972983
## sample estimates:
## difference in location 
##                   -0.5 
## 
## Mann-Whitney test results for Feb tmax: actual vs. RCP4.5 2070 
## The H0 is rejected as p < 0.05, the HA accepted (medians unequal) 
## 
##  Wilcoxon rank sum test with continuity correction
## 
## data:  x and y
## W = 14352, p-value < 2.2e-16
## alternative hypothesis: true location shift is not equal to 0
## 95 percent confidence interval:
##  -4.180606 -3.950480
## sample estimates:
## difference in location 
##              -4.065342 
## 
## Mann-Whitney test results for Feb tmax: actual vs. RCP8.5 2050 
## The H0 is rejected as p < 0.05, the HA accepted (medians unequal) 
## 
##  Wilcoxon rank sum test with continuity correction
## 
## data:  x and y
## W = 10801, p-value < 2.2e-16
## alternative hypothesis: true location shift is not equal to 0
## 95 percent confidence interval:
##  -4.380606 -4.150480
## sample estimates:
## difference in location 
##              -4.265342 
## 
## Mann-Whitney test results for Feb tmax: RCP8.5 2050 vs. RCP8.5 2070 
## The H0 is rejected as p < 0.05, the HA accepted (medians unequal) 
## 
##  Wilcoxon rank sum test with continuity correction
## 
## data:  x and y
## W = 236890, p-value < 2.2e-16
## alternative hypothesis: true location shift is not equal to 0
## 95 percent confidence interval:
##  -1.302553 -1.097175
## sample estimates:
## difference in location 
##                   -1.2 
## 
## Mann-Whitney test results for Feb tmax: actual vs. RCP8.5 2070 
## The H0 is rejected as p < 0.05, the HA accepted (medians unequal) 
## 
##  Wilcoxon rank sum test with continuity correction
## 
## data:  x and y
## W = 1734, p-value < 2.2e-16
## alternative hypothesis: true location shift is not equal to 0
## 95 percent confidence interval:
##  -5.580541 -5.350375
## sample estimates:
## difference in location 
##              -5.465197 
## 
## Mann-Whitney test results for Mar precipitation: actual vs. RCP4.5 2050 
## The H0 is rejected as p < 0.05, the HA accepted (medians unequal) 
## 
##  Wilcoxon rank sum test with continuity correction
## 
## data:  x and y
## W = 281640, p-value < 2.2e-16
## alternative hypothesis: true location shift is not equal to 0
## 95 percent confidence interval:
##  -43.74855 -34.41127
## sample estimates:
## difference in location 
##              -39.06025 
## 
## Mann-Whitney test results for Mar precipitation: RCP4.5 2050 vs. RCP4.5 2070 
## The H0 is rejected as p < 0.05, the HA accepted (medians unequal) 
## 
##  Wilcoxon rank sum test with continuity correction
## 
## data:  x and y
## W = 612520, p-value < 2.2e-16
## alternative hypothesis: true location shift is not equal to 0
## 95 percent confidence interval:
##  17.02773 26.20353
## sample estimates:
## difference in location 
##               21.61491 
## 
## Mann-Whitney test results for Mar precipitation: actual vs. RCP4.5 2070 
## The H0 is rejected as p < 0.05, the HA accepted (medians unequal) 
## 
##  Wilcoxon rank sum test with continuity correction
## 
## data:  x and y
## W = 396810, p-value = 1.335e-15
## alternative hypothesis: true location shift is not equal to 0
## 95 percent confidence interval:
##  -22.05887 -13.31625
## sample estimates:
## difference in location 
##              -17.67441 
## 
## Mann-Whitney test results for Mar precipitation: actual vs. RCP8.5 2050 
## The H0 is rejected as p < 0.05, the HA accepted (medians unequal) 
## 
##  Wilcoxon rank sum test with continuity correction
## 
## data:  x and y
## W = 340150, p-value < 2.2e-16
## alternative hypothesis: true location shift is not equal to 0
## 95 percent confidence interval:
##  -32.20111 -23.21214
## sample estimates:
## difference in location 
##               -27.6655 
## 
## Mann-Whitney test results for Mar precipitation: RCP8.5 2050 vs. RCP8.5 2070 
## The HA is rejected as p >= 0.05, the H0 accepted (medians equal) 
## 
##  Wilcoxon rank sum test with continuity correction
## 
## data:  x and y
## W = 517190, p-value = 0.1832
## alternative hypothesis: true location shift is not equal to 0
## 95 percent confidence interval:
##  -1.488437  7.729692
## sample estimates:
## difference in location 
##               3.085777 
## 
## Mann-Whitney test results for Mar precipitation: actual vs. RCP8.5 2070 
## The H0 is rejected as p < 0.05, the HA accepted (medians unequal) 
## 
##  Wilcoxon rank sum test with continuity correction
## 
## data:  x and y
## W = 361180, p-value < 2.2e-16
## alternative hypothesis: true location shift is not equal to 0
## 95 percent confidence interval:
##  -29.22238 -20.00940
## sample estimates:
## difference in location 
##              -24.60054 
## 
## Mann-Whitney test results for Mar tmin: actual vs. RCP4.5 2050 
## The H0 is rejected as p < 0.05, the HA accepted (medians unequal) 
## 
##  Wilcoxon rank sum test with continuity correction
## 
## data:  x and y
## W = 283790, p-value < 2.2e-16
## alternative hypothesis: true location shift is not equal to 0
## 95 percent confidence interval:
##  -1.569244 -1.255354
## sample estimates:
## difference in location 
##              -1.412785 
## 
## Mann-Whitney test results for Mar tmin: RCP4.5 2050 vs. RCP4.5 2070 
## The H0 is rejected as p < 0.05, the HA accepted (medians unequal) 
## 
##  Wilcoxon rank sum test with continuity correction
## 
## data:  x and y
## W = 429150, p-value = 4.094e-08
## alternative hypothesis: true location shift is not equal to 0
## 95 percent confidence interval:
##  -0.5568461 -0.2659892
## sample estimates:
## difference in location 
##              -0.409002 
## 
## Mann-Whitney test results for Mar tmin: actual vs. RCP4.5 2070 
## The H0 is rejected as p < 0.05, the HA accepted (medians unequal) 
## 
##  Wilcoxon rank sum test with continuity correction
## 
## data:  x and y
## W = 229690, p-value < 2.2e-16
## alternative hypothesis: true location shift is not equal to 0
## 95 percent confidence interval:
##  -1.980150 -1.666411
## sample estimates:
## difference in location 
##              -1.823656 
## 
## Mann-Whitney test results for Mar tmin: actual vs. RCP8.5 2050 
## The H0 is rejected as p < 0.05, the HA accepted (medians unequal) 
## 
##  Wilcoxon rank sum test with continuity correction
## 
## data:  x and y
## W = 215260, p-value < 2.2e-16
## alternative hypothesis: true location shift is not equal to 0
## 95 percent confidence interval:
##  -2.095766 -1.781871
## sample estimates:
## difference in location 
##              -1.938802 
## 
## Mann-Whitney test results for Mar tmin: RCP8.5 2050 vs. RCP8.5 2070 
## The H0 is rejected as p < 0.05, the HA accepted (medians unequal) 
## 
##  Wilcoxon rank sum test with continuity correction
## 
## data:  x and y
## W = 303430, p-value < 2.2e-16
## alternative hypothesis: true location shift is not equal to 0
## 95 percent confidence interval:
##  -1.340861 -1.052067
## sample estimates:
## difference in location 
##              -1.198925 
## 
## Mann-Whitney test results for Mar tmin: actual vs. RCP8.5 2070 
## The H0 is rejected as p < 0.05, the HA accepted (medians unequal) 
## 
##  Wilcoxon rank sum test with continuity correction
## 
## data:  x and y
## W = 95001, p-value < 2.2e-16
## alternative hypothesis: true location shift is not equal to 0
## 95 percent confidence interval:
##  -3.292527 -2.979080
## sample estimates:
## difference in location 
##              -3.135638 
## 
## Mann-Whitney test results for Mar tmax: actual vs. RCP4.5 2050 
## The H0 is rejected as p < 0.05, the HA accepted (medians unequal) 
## 
##  Wilcoxon rank sum test with continuity correction
## 
## data:  x and y
## W = 32250, p-value < 2.2e-16
## alternative hypothesis: true location shift is not equal to 0
## 95 percent confidence interval:
##  -3.568139 -3.339505
## sample estimates:
## difference in location 
##              -3.454325 
## 
## Mann-Whitney test results for Mar tmax: RCP4.5 2050 vs. RCP4.5 2070 
## The H0 is rejected as p < 0.05, the HA accepted (medians unequal) 
## 
##  Wilcoxon rank sum test with continuity correction
## 
## data:  x and y
## W = 382930, p-value < 2.2e-16
## alternative hypothesis: true location shift is not equal to 0
## 95 percent confidence interval:
##  -0.6044347 -0.3955653
## sample estimates:
## difference in location 
##                   -0.5 
## 
## Mann-Whitney test results for Mar tmax: actual vs. RCP4.5 2070 
## The H0 is rejected as p < 0.05, the HA accepted (medians unequal) 
## 
##  Wilcoxon rank sum test with continuity correction
## 
## data:  x and y
## W = 17190, p-value < 2.2e-16
## alternative hypothesis: true location shift is not equal to 0
## 95 percent confidence interval:
##  -4.068139 -3.839505
## sample estimates:
## difference in location 
##              -3.954325 
## 
## Mann-Whitney test results for Mar tmax: actual vs. RCP8.5 2050 
## The H0 is rejected as p < 0.05, the HA accepted (medians unequal) 
## 
##  Wilcoxon rank sum test with continuity correction
## 
## data:  x and y
## W = 15084, p-value < 2.2e-16
## alternative hypothesis: true location shift is not equal to 0
## 95 percent confidence interval:
##  -4.168139 -3.939505
## sample estimates:
## difference in location 
##              -4.054325 
## 
## Mann-Whitney test results for Mar tmax: RCP8.5 2050 vs. RCP8.5 2070 
## The H0 is rejected as p < 0.05, the HA accepted (medians unequal) 
## 
##  Wilcoxon rank sum test with continuity correction
## 
## data:  x and y
## W = 241220, p-value < 2.2e-16
## alternative hypothesis: true location shift is not equal to 0
## 95 percent confidence interval:
##  -1.304435 -1.095565
## sample estimates:
## difference in location 
##                   -1.2 
## 
## Mann-Whitney test results for Mar tmax: actual vs. RCP8.5 2070 
## The H0 is rejected as p < 0.05, the HA accepted (medians unequal) 
## 
##  Wilcoxon rank sum test with continuity correction
## 
## data:  x and y
## W = 2633, p-value < 2.2e-16
## alternative hypothesis: true location shift is not equal to 0
## 95 percent confidence interval:
##  -5.368139 -5.139546
## sample estimates:
## difference in location 
##              -5.254325 
## 
## Mann-Whitney test results for Apr precipitation: actual vs. RCP4.5 2050 
## The H0 is rejected as p < 0.05, the HA accepted (medians unequal) 
## 
##  Wilcoxon rank sum test with continuity correction
## 
## data:  x and y
## W = 317400, p-value < 2.2e-16
## alternative hypothesis: true location shift is not equal to 0
## 95 percent confidence interval:
##  -45.21552 -33.76145
## sample estimates:
## difference in location 
##              -39.50453 
## 
## Mann-Whitney test results for Apr precipitation: RCP4.5 2050 vs. RCP4.5 2070 
## The H0 is rejected as p < 0.05, the HA accepted (medians unequal) 
## 
##  Wilcoxon rank sum test with continuity correction
## 
## data:  x and y
## W = 625620, p-value < 2.2e-16
## alternative hypothesis: true location shift is not equal to 0
## 95 percent confidence interval:
##  25.45193 37.40365
## sample estimates:
## difference in location 
##               31.39042 
## 
## Mann-Whitney test results for Apr precipitation: actual vs. RCP4.5 2070 
## The H0 is rejected as p < 0.05, the HA accepted (medians unequal) 
## 
##  Wilcoxon rank sum test with continuity correction
## 
## data:  x and y
## W = 459970, p-value = 0.001934
## alternative hypothesis: true location shift is not equal to 0
## 95 percent confidence interval:
##  -14.381634  -3.124609
## sample estimates:
## difference in location 
##              -8.702567 
## 
## Mann-Whitney test results for Apr precipitation: actual vs. RCP8.5 2050 
## The H0 is rejected as p < 0.05, the HA accepted (medians unequal) 
## 
##  Wilcoxon rank sum test with continuity correction
## 
## data:  x and y
## W = 410070, p-value = 3.303e-12
## alternative hypothesis: true location shift is not equal to 0
## 95 percent confidence interval:
##  -26.03097 -14.20310
## sample estimates:
## difference in location 
##              -20.07402 
## 
## Mann-Whitney test results for Apr precipitation: RCP8.5 2050 vs. RCP8.5 2070 
## The HA is rejected as p >= 0.05, the H0 accepted (medians equal) 
## 
##  Wilcoxon rank sum test with continuity correction
## 
## data:  x and y
## W = 509110, p-value = 0.4803
## alternative hypothesis: true location shift is not equal to 0
## 95 percent confidence interval:
##  -3.846026  8.035148
## sample estimates:
## difference in location 
##               2.072403 
## 
## Mann-Whitney test results for Apr precipitation: actual vs. RCP8.5 2070 
## The H0 is rejected as p < 0.05, the HA accepted (medians unequal) 
## 
##  Wilcoxon rank sum test with continuity correction
## 
## data:  x and y
## W = 420060, p-value = 5.998e-10
## alternative hypothesis: true location shift is not equal to 0
## 95 percent confidence interval:
##  -24.19197 -12.18753
## sample estimates:
## difference in location 
##               -18.1185 
## 
## Mann-Whitney test results for Apr tmin: actual vs. RCP4.5 2050 
## The H0 is rejected as p < 0.05, the HA accepted (medians unequal) 
## 
##  Wilcoxon rank sum test with continuity correction
## 
## data:  x and y
## W = 252770, p-value < 2.2e-16
## alternative hypothesis: true location shift is not equal to 0
## 95 percent confidence interval:
##  -1.743202 -1.438721
## sample estimates:
## difference in location 
##              -1.591587 
## 
## Mann-Whitney test results for Apr tmin: RCP4.5 2050 vs. RCP4.5 2070 
## The H0 is rejected as p < 0.05, the HA accepted (medians unequal) 
## 
##  Wilcoxon rank sum test with continuity correction
## 
## data:  x and y
## W = 432540, p-value = 1.749e-07
## alternative hypothesis: true location shift is not equal to 0
## 95 percent confidence interval:
##  -0.5182076 -0.2376898
## sample estimates:
## difference in location 
##             -0.3788826 
## 
## Mann-Whitney test results for Apr tmin: actual vs. RCP4.5 2070 
## The H0 is rejected as p < 0.05, the HA accepted (medians unequal) 
## 
##  Wilcoxon rank sum test with continuity correction
## 
## data:  x and y
## W = 203940, p-value < 2.2e-16
## alternative hypothesis: true location shift is not equal to 0
## 95 percent confidence interval:
##  -2.121990 -1.817591
## sample estimates:
## difference in location 
##              -1.969394 
## 
## Mann-Whitney test results for Apr tmin: actual vs. RCP8.5 2050 
## The H0 is rejected as p < 0.05, the HA accepted (medians unequal) 
## 
##  Wilcoxon rank sum test with continuity correction
## 
## data:  x and y
## W = 188910, p-value < 2.2e-16
## alternative hypothesis: true location shift is not equal to 0
## 95 percent confidence interval:
##  -2.247044 -1.941933
## sample estimates:
## difference in location 
##              -2.095092 
## 
## Mann-Whitney test results for Apr tmin: RCP8.5 2050 vs. RCP8.5 2070 
## The H0 is rejected as p < 0.05, the HA accepted (medians unequal) 
## 
##  Wilcoxon rank sum test with continuity correction
## 
## data:  x and y
## W = 297300, p-value < 2.2e-16
## alternative hypothesis: true location shift is not equal to 0
## 95 percent confidence interval:
##  -1.338818 -1.059820
## sample estimates:
## difference in location 
##                   -1.2 
## 
## Mann-Whitney test results for Apr tmin: actual vs. RCP8.5 2070 
## The H0 is rejected as p < 0.05, the HA accepted (medians unequal) 
## 
##  Wilcoxon rank sum test with continuity correction
## 
## data:  x and y
## W = 76074, p-value < 2.2e-16
## alternative hypothesis: true location shift is not equal to 0
## 95 percent confidence interval:
##  -3.446490 -3.141269
## sample estimates:
## difference in location 
##               -3.29448 
## 
## Mann-Whitney test results for Apr tmax: actual vs. RCP4.5 2050 
## The H0 is rejected as p < 0.05, the HA accepted (medians unequal) 
## 
##  Wilcoxon rank sum test with continuity correction
## 
## data:  x and y
## W = 36197, p-value < 2.2e-16
## alternative hypothesis: true location shift is not equal to 0
## 95 percent confidence interval:
##  -3.399704 -3.176260
## sample estimates:
## difference in location 
##               -3.28836 
## 
## Mann-Whitney test results for Apr tmax: RCP4.5 2050 vs. RCP4.5 2070 
## The H0 is rejected as p < 0.05, the HA accepted (medians unequal) 
## 
##  Wilcoxon rank sum test with continuity correction
## 
## data:  x and y
## W = 382770, p-value < 2.2e-16
## alternative hypothesis: true location shift is not equal to 0
## 95 percent confidence interval:
##  -0.6046438 -0.3953562
## sample estimates:
## difference in location 
##                   -0.5 
## 
## Mann-Whitney test results for Apr tmax: actual vs. RCP4.5 2070 
## The H0 is rejected as p < 0.05, the HA accepted (medians unequal) 
## 
##  Wilcoxon rank sum test with continuity correction
## 
## data:  x and y
## W = 19386, p-value < 2.2e-16
## alternative hypothesis: true location shift is not equal to 0
## 95 percent confidence interval:
##  -3.899704 -3.676260
## sample estimates:
## difference in location 
##               -3.78836 
## 
## Mann-Whitney test results for Apr tmax: actual vs. RCP8.5 2050 
## The H0 is rejected as p < 0.05, the HA accepted (medians unequal) 
## 
##  Wilcoxon rank sum test with continuity correction
## 
## data:  x and y
## W = 16918, p-value < 2.2e-16
## alternative hypothesis: true location shift is not equal to 0
## 95 percent confidence interval:
##  -4.000910 -3.777144
## sample estimates:
## difference in location 
##              -3.889281 
## 
## Mann-Whitney test results for Apr tmax: RCP8.5 2050 vs. RCP8.5 2070 
## The H0 is rejected as p < 0.05, the HA accepted (medians unequal) 
## 
##  Wilcoxon rank sum test with continuity correction
## 
## data:  x and y
## W = 241000, p-value < 2.2e-16
## alternative hypothesis: true location shift is not equal to 0
## 95 percent confidence interval:
##  -1.303680 -1.094219
## sample estimates:
## difference in location 
##                   -1.2 
## 
## Mann-Whitney test results for Apr tmax: actual vs. RCP8.5 2070 
## The H0 is rejected as p < 0.05, the HA accepted (medians unequal) 
## 
##  Wilcoxon rank sum test with continuity correction
## 
## data:  x and y
## W = 2841, p-value < 2.2e-16
## alternative hypothesis: true location shift is not equal to 0
## 95 percent confidence interval:
##  -5.199704 -4.976260
## sample estimates:
## difference in location 
##               -5.08836 
## 
## Mann-Whitney test results for May precipitation: actual vs. RCP4.5 2050 
## The H0 is rejected as p < 0.05, the HA accepted (medians unequal) 
## 
##  Wilcoxon rank sum test with continuity correction
## 
## data:  x and y
## W = 252280, p-value < 2.2e-16
## alternative hypothesis: true location shift is not equal to 0
## 95 percent confidence interval:
##  -83.06798 -67.24347
## sample estimates:
## difference in location 
##              -75.08513 
## 
## Mann-Whitney test results for May precipitation: RCP4.5 2050 vs. RCP4.5 2070 
## The H0 is rejected as p < 0.05, the HA accepted (medians unequal) 
## 
##  Wilcoxon rank sum test with continuity correction
## 
## data:  x and y
## W = 542650, p-value = 0.0009579
## alternative hypothesis: true location shift is not equal to 0
## 95 percent confidence interval:
##   5.413146 20.710962
## sample estimates:
## difference in location 
##               13.04254 
## 
## Mann-Whitney test results for May precipitation: actual vs. RCP4.5 2070 
## The H0 is rejected as p < 0.05, the HA accepted (medians unequal) 
## 
##  Wilcoxon rank sum test with continuity correction
## 
## data:  x and y
## W = 287740, p-value < 2.2e-16
## alternative hypothesis: true location shift is not equal to 0
## 95 percent confidence interval:
##  -69.64010 -54.26802
## sample estimates:
## difference in location 
##              -61.90883 
## 
## Mann-Whitney test results for May precipitation: actual vs. RCP8.5 2050 
## The H0 is rejected as p < 0.05, the HA accepted (medians unequal) 
## 
##  Wilcoxon rank sum test with continuity correction
## 
## data:  x and y
## W = 279390, p-value < 2.2e-16
## alternative hypothesis: true location shift is not equal to 0
## 95 percent confidence interval:
##  -73.95381 -58.16446
## sample estimates:
## difference in location 
##              -66.01313 
## 
## Mann-Whitney test results for May precipitation: RCP8.5 2050 vs. RCP8.5 2070 
## The H0 is rejected as p < 0.05, the HA accepted (medians unequal) 
## 
##  Wilcoxon rank sum test with continuity correction
## 
## data:  x and y
## W = 443020, p-value = 1.022e-05
## alternative hypothesis: true location shift is not equal to 0
## 95 percent confidence interval:
##  -27.27757 -10.55704
## sample estimates:
## difference in location 
##              -18.92853 
## 
## Mann-Whitney test results for May precipitation: actual vs. RCP8.5 2070 
## The H0 is rejected as p < 0.05, the HA accepted (medians unequal) 
## 
##  Wilcoxon rank sum test with continuity correction
## 
## data:  x and y
## W = 239450, p-value < 2.2e-16
## alternative hypothesis: true location shift is not equal to 0
## 95 percent confidence interval:
##  -93.57849 -76.29179
## sample estimates:
## difference in location 
##              -84.87028 
## 
## Mann-Whitney test results for May tmin: actual vs. RCP4.5 2050 
## The H0 is rejected as p < 0.05, the HA accepted (medians unequal) 
## 
##  Wilcoxon rank sum test with continuity correction
## 
## data:  x and y
## W = 276670, p-value < 2.2e-16
## alternative hypothesis: true location shift is not equal to 0
## 95 percent confidence interval:
##  -1.591019 -1.282879
## sample estimates:
## difference in location 
##              -1.437616 
## 
## Mann-Whitney test results for May tmin: RCP4.5 2050 vs. RCP4.5 2070 
## The H0 is rejected as p < 0.05, the HA accepted (medians unequal) 
## 
##  Wilcoxon rank sum test with continuity correction
## 
## data:  x and y
## W = 430640, p-value = 7.81e-08
## alternative hypothesis: true location shift is not equal to 0
## 95 percent confidence interval:
##  -0.5444225 -0.2566420
## sample estimates:
## difference in location 
##                   -0.4 
## 
## Mann-Whitney test results for May tmin: actual vs. RCP4.5 2070 
## The H0 is rejected as p < 0.05, the HA accepted (medians unequal) 
## 
##  Wilcoxon rank sum test with continuity correction
## 
## data:  x and y
## W = 223850, p-value < 2.2e-16
## alternative hypothesis: true location shift is not equal to 0
## 95 percent confidence interval:
##  -1.991759 -1.683438
## sample estimates:
## difference in location 
##              -1.838116 
## 
## Mann-Whitney test results for May tmin: actual vs. RCP8.5 2050 
## The H0 is rejected as p < 0.05, the HA accepted (medians unequal) 
## 
##  Wilcoxon rank sum test with continuity correction
## 
## data:  x and y
## W = 211170, p-value < 2.2e-16
## alternative hypothesis: true location shift is not equal to 0
## 95 percent confidence interval:
##  -2.092519 -1.784196
## sample estimates:
## difference in location 
##              -1.938574 
## 
## Mann-Whitney test results for May tmin: RCP8.5 2050 vs. RCP8.5 2070 
## The H0 is rejected as p < 0.05, the HA accepted (medians unequal) 
## 
##  Wilcoxon rank sum test with continuity correction
## 
## data:  x and y
## W = 301720, p-value < 2.2e-16
## alternative hypothesis: true location shift is not equal to 0
## 95 percent confidence interval:
##  -1.354558 -1.066023
## sample estimates:
## difference in location 
##              -1.207763 
## 
## Mann-Whitney test results for May tmin: actual vs. RCP8.5 2070 
## The H0 is rejected as p < 0.05, the HA accepted (medians unequal) 
## 
##  Wilcoxon rank sum test with continuity correction
## 
## data:  x and y
## W = 90125, p-value < 2.2e-16
## alternative hypothesis: true location shift is not equal to 0
## 95 percent confidence interval:
##  -3.305201 -2.995177
## sample estimates:
## difference in location 
##              -3.149611 
## 
## Mann-Whitney test results for May tmax: actual vs. RCP4.5 2050 
## The H0 is rejected as p < 0.05, the HA accepted (medians unequal) 
## 
##  Wilcoxon rank sum test with continuity correction
## 
## data:  x and y
## W = 34247, p-value < 2.2e-16
## alternative hypothesis: true location shift is not equal to 0
## 95 percent confidence interval:
##  -3.399689 -3.181138
## sample estimates:
## difference in location 
##              -3.291144 
## 
## Mann-Whitney test results for May tmax: RCP4.5 2050 vs. RCP4.5 2070 
## The H0 is rejected as p < 0.05, the HA accepted (medians unequal) 
## 
##  Wilcoxon rank sum test with continuity correction
## 
## data:  x and y
## W = 371970, p-value < 2.2e-16
## alternative hypothesis: true location shift is not equal to 0
## 95 percent confidence interval:
##  -0.6171100 -0.4201196
## sample estimates:
## difference in location 
##             -0.5174696 
## 
## Mann-Whitney test results for May tmax: actual vs. RCP4.5 2070 
## The H0 is rejected as p < 0.05, the HA accepted (medians unequal) 
## 
##  Wilcoxon rank sum test with continuity correction
## 
## data:  x and y
## W = 18122, p-value < 2.2e-16
## alternative hypothesis: true location shift is not equal to 0
## 95 percent confidence interval:
##  -3.918784 -3.700042
## sample estimates:
## difference in location 
##              -3.809452 
## 
## Mann-Whitney test results for May tmax: actual vs. RCP8.5 2050 
## The H0 is rejected as p < 0.05, the HA accepted (medians unequal) 
## 
##  Wilcoxon rank sum test with continuity correction
## 
## data:  x and y
## W = 15842, p-value < 2.2e-16
## alternative hypothesis: true location shift is not equal to 0
## 95 percent confidence interval:
##  -4.024303 -3.804997
## sample estimates:
## difference in location 
##              -3.914807 
## 
## Mann-Whitney test results for May tmax: RCP8.5 2050 vs. RCP8.5 2070 
## The H0 is rejected as p < 0.05, the HA accepted (medians unequal) 
## 
##  Wilcoxon rank sum test with continuity correction
## 
## data:  x and y
## W = 231430, p-value < 2.2e-16
## alternative hypothesis: true location shift is not equal to 0
## 95 percent confidence interval:
##  -1.291378 -1.093303
## sample estimates:
## difference in location 
##              -1.193845 
## 
## Mann-Whitney test results for May tmax: actual vs. RCP8.5 2070 
## The H0 is rejected as p < 0.05, the HA accepted (medians unequal) 
## 
##  Wilcoxon rank sum test with continuity correction
## 
## data:  x and y
## W = 2844, p-value < 2.2e-16
## alternative hypothesis: true location shift is not equal to 0
## 95 percent confidence interval:
##  -5.216066 -4.998140
## sample estimates:
## difference in location 
##              -5.107272 
## 
## Mann-Whitney test results for Jun precipitation: actual vs. RCP4.5 2050 
## The H0 is rejected as p < 0.05, the HA accepted (medians unequal) 
## 
##  Wilcoxon rank sum test with continuity correction
## 
## data:  x and y
## W = 294340, p-value < 2.2e-16
## alternative hypothesis: true location shift is not equal to 0
## 95 percent confidence interval:
##  -81.23977 -63.11891
## sample estimates:
## difference in location 
##              -72.20869 
## 
## Mann-Whitney test results for Jun precipitation: RCP4.5 2050 vs. RCP4.5 2070 
## The HA is rejected as p >= 0.05, the H0 accepted (medians equal) 
## 
##  Wilcoxon rank sum test with continuity correction
## 
## data:  x and y
## W = 487100, p-value = 0.3178
## alternative hypothesis: true location shift is not equal to 0
## 95 percent confidence interval:
##  -14.761922   4.902003
## sample estimates:
## difference in location 
##              -4.941371 
## 
## Mann-Whitney test results for Jun precipitation: actual vs. RCP4.5 2070 
## The H0 is rejected as p < 0.05, the HA accepted (medians unequal) 
## 
##  Wilcoxon rank sum test with continuity correction
## 
## data:  x and y
## W = 282460, p-value < 2.2e-16
## alternative hypothesis: true location shift is not equal to 0
## 95 percent confidence interval:
##  -85.88447 -67.61795
## sample estimates:
## difference in location 
##              -76.69199 
## 
## Mann-Whitney test results for Jun precipitation: actual vs. RCP8.5 2050 
## The H0 is rejected as p < 0.05, the HA accepted (medians unequal) 
## 
##  Wilcoxon rank sum test with continuity correction
## 
## data:  x and y
## W = 344800, p-value < 2.2e-16
## alternative hypothesis: true location shift is not equal to 0
## 95 percent confidence interval:
##  -60.92428 -43.47564
## sample estimates:
## difference in location 
##              -52.18239 
## 
## Mann-Whitney test results for Jun precipitation: RCP8.5 2050 vs. RCP8.5 2070 
## The H0 is rejected as p < 0.05, the HA accepted (medians unequal) 
## 
##  Wilcoxon rank sum test with continuity correction
## 
## data:  x and y
## W = 347140, p-value < 2.2e-16
## alternative hypothesis: true location shift is not equal to 0
## 95 percent confidence interval:
##  -77.48668 -56.02856
## sample estimates:
## difference in location 
##              -66.76635 
## 
## Mann-Whitney test results for Jun precipitation: actual vs. RCP8.5 2070 
## The H0 is rejected as p < 0.05, the HA accepted (medians unequal) 
## 
##  Wilcoxon rank sum test with continuity correction
## 
## data:  x and y
## W = 209520, p-value < 2.2e-16
## alternative hypothesis: true location shift is not equal to 0
## 95 percent confidence interval:
##  -130.2544 -109.0997
## sample estimates:
## difference in location 
##              -119.5863 
## 
## Mann-Whitney test results for Jun tmin: actual vs. RCP4.5 2050 
## The H0 is rejected as p < 0.05, the HA accepted (medians unequal) 
## 
##  Wilcoxon rank sum test with continuity correction
## 
## data:  x and y
## W = 277760, p-value < 2.2e-16
## alternative hypothesis: true location shift is not equal to 0
## 95 percent confidence interval:
##  -1.652584 -1.331420
## sample estimates:
## difference in location 
##              -1.491937 
## 
## Mann-Whitney test results for Jun tmin: RCP4.5 2050 vs. RCP4.5 2070 
## The H0 is rejected as p < 0.05, the HA accepted (medians unequal) 
## 
##  Wilcoxon rank sum test with continuity correction
## 
## data:  x and y
## W = 402980, p-value = 5.766e-14
## alternative hypothesis: true location shift is not equal to 0
## 95 percent confidence interval:
##  -0.7396457 -0.4394855
## sample estimates:
## difference in location 
##             -0.5921295 
## 
## Mann-Whitney test results for Jun tmin: actual vs. RCP4.5 2070 
## The H0 is rejected as p < 0.05, the HA accepted (medians unequal) 
## 
##  Wilcoxon rank sum test with continuity correction
## 
## data:  x and y
## W = 203230, p-value < 2.2e-16
## alternative hypothesis: true location shift is not equal to 0
## 95 percent confidence interval:
##  -2.240650 -1.920601
## sample estimates:
## difference in location 
##              -2.080645 
## 
## Mann-Whitney test results for Jun tmin: actual vs. RCP8.5 2050 
## The H0 is rejected as p < 0.05, the HA accepted (medians unequal) 
## 
##  Wilcoxon rank sum test with continuity correction
## 
## data:  x and y
## W = 215380, p-value < 2.2e-16
## alternative hypothesis: true location shift is not equal to 0
## 95 percent confidence interval:
##  -2.141287 -1.821315
## sample estimates:
## difference in location 
##              -1.981308 
## 
## Mann-Whitney test results for Jun tmin: RCP8.5 2050 vs. RCP8.5 2070 
## The H0 is rejected as p < 0.05, the HA accepted (medians unequal) 
## 
##  Wilcoxon rank sum test with continuity correction
## 
## data:  x and y
## W = 259290, p-value < 2.2e-16
## alternative hypothesis: true location shift is not equal to 0
## 95 percent confidence interval:
##  -1.718289 -1.417254
## sample estimates:
## difference in location 
##              -1.568385 
## 
## Mann-Whitney test results for Jun tmin: actual vs. RCP8.5 2070 
## The H0 is rejected as p < 0.05, the HA accepted (medians unequal) 
## 
##  Wilcoxon rank sum test with continuity correction
## 
## data:  x and y
## W = 71119, p-value < 2.2e-16
## alternative hypothesis: true location shift is not equal to 0
## 95 percent confidence interval:
##  -3.711493 -3.390343
## sample estimates:
## difference in location 
##              -3.550015 
## 
## Mann-Whitney test results for Jun tmax: actual vs. RCP4.5 2050 
## The H0 is rejected as p < 0.05, the HA accepted (medians unequal) 
## 
##  Wilcoxon rank sum test with continuity correction
## 
## data:  x and y
## W = 58992, p-value < 2.2e-16
## alternative hypothesis: true location shift is not equal to 0
## 95 percent confidence interval:
##  -3.193794 -2.953119
## sample estimates:
## difference in location 
##              -3.073387 
## 
## Mann-Whitney test results for Jun tmax: RCP4.5 2050 vs. RCP4.5 2070 
## The H0 is rejected as p < 0.05, the HA accepted (medians unequal) 
## 
##  Wilcoxon rank sum test with continuity correction
## 
## data:  x and y
## W = 380320, p-value < 2.2e-16
## alternative hypothesis: true location shift is not equal to 0
## 95 percent confidence interval:
##  -0.6654539 -0.4400799
## sample estimates:
## difference in location 
##             -0.5531984 
## 
## Mann-Whitney test results for Jun tmax: actual vs. RCP4.5 2070 
## The H0 is rejected as p < 0.05, the HA accepted (medians unequal) 
## 
##  Wilcoxon rank sum test with continuity correction
## 
## data:  x and y
## W = 33111, p-value < 2.2e-16
## alternative hypothesis: true location shift is not equal to 0
## 95 percent confidence interval:
##  -3.745314 -3.507767
## sample estimates:
## difference in location 
##              -3.626544 
## 
## Mann-Whitney test results for Jun tmax: actual vs. RCP8.5 2050 
## The H0 is rejected as p < 0.05, the HA accepted (medians unequal) 
## 
##  Wilcoxon rank sum test with continuity correction
## 
## data:  x and y
## W = 29572, p-value < 2.2e-16
## alternative hypothesis: true location shift is not equal to 0
## 95 percent confidence interval:
##  -3.842376 -3.604338
## sample estimates:
## difference in location 
##              -3.723806 
## 
## Mann-Whitney test results for Jun tmax: RCP8.5 2050 vs. RCP8.5 2070 
## The H0 is rejected as p < 0.05, the HA accepted (medians unequal) 
## 
##  Wilcoxon rank sum test with continuity correction
## 
## data:  x and y
## W = 199720, p-value < 2.2e-16
## alternative hypothesis: true location shift is not equal to 0
## 95 percent confidence interval:
##  -1.668970 -1.444638
## sample estimates:
## difference in location 
##              -1.555992 
## 
## Mann-Whitney test results for Jun tmax: actual vs. RCP8.5 2070 
## The H0 is rejected as p < 0.05, the HA accepted (medians unequal) 
## 
##  Wilcoxon rank sum test with continuity correction
## 
## data:  x and y
## W = 4306, p-value < 2.2e-16
## alternative hypothesis: true location shift is not equal to 0
## 95 percent confidence interval:
##  -5.398402 -5.158663
## sample estimates:
## difference in location 
##              -5.278783 
## 
## Mann-Whitney test results for Jul precipitation: actual vs. RCP4.5 2050 
## The H0 is rejected as p < 0.05, the HA accepted (medians unequal) 
## 
##  Wilcoxon rank sum test with continuity correction
## 
## data:  x and y
## W = 255680, p-value < 2.2e-16
## alternative hypothesis: true location shift is not equal to 0
## 95 percent confidence interval:
##  -101.99035  -83.10093
## sample estimates:
## difference in location 
##               -92.5334 
## 
## Mann-Whitney test results for Jul precipitation: RCP4.5 2050 vs. RCP4.5 2070 
## The HA is rejected as p >= 0.05, the H0 accepted (medians equal) 
## 
##  Wilcoxon rank sum test with continuity correction
## 
## data:  x and y
## W = 503620, p-value = 0.7793
## alternative hypothesis: true location shift is not equal to 0
## 95 percent confidence interval:
##  -8.266498 10.757231
## sample estimates:
## difference in location 
##               1.320574 
## 
## Mann-Whitney test results for Jul precipitation: actual vs. RCP4.5 2070 
## The H0 is rejected as p < 0.05, the HA accepted (medians unequal) 
## 
##  Wilcoxon rank sum test with continuity correction
## 
## data:  x and y
## W = 255330, p-value < 2.2e-16
## alternative hypothesis: true location shift is not equal to 0
## 95 percent confidence interval:
##  -100.92336  -82.30876
## sample estimates:
## difference in location 
##              -91.57532 
## 
## Mann-Whitney test results for Jul precipitation: actual vs. RCP8.5 2050 
## The H0 is rejected as p < 0.05, the HA accepted (medians unequal) 
## 
##  Wilcoxon rank sum test with continuity correction
## 
## data:  x and y
## W = 365380, p-value < 2.2e-16
## alternative hypothesis: true location shift is not equal to 0
## 95 percent confidence interval:
##  -52.03305 -35.63505
## sample estimates:
## difference in location 
##              -43.83594 
## 
## Mann-Whitney test results for Jul precipitation: RCP8.5 2050 vs. RCP8.5 2070 
## The H0 is rejected as p < 0.05, the HA accepted (medians unequal) 
## 
##  Wilcoxon rank sum test with continuity correction
## 
## data:  x and y
## W = 405160, p-value = 2.072e-13
## alternative hypothesis: true location shift is not equal to 0
## 95 percent confidence interval:
##  -44.75884 -26.34628
## sample estimates:
## difference in location 
##              -35.52443 
## 
## Mann-Whitney test results for Jul precipitation: actual vs. RCP8.5 2070 
## The H0 is rejected as p < 0.05, the HA accepted (medians unequal) 
## 
##  Wilcoxon rank sum test with continuity correction
## 
## data:  x and y
## W = 284610, p-value < 2.2e-16
## alternative hypothesis: true location shift is not equal to 0
## 95 percent confidence interval:
##  -87.92123 -69.46707
## sample estimates:
## difference in location 
##               -78.7283 
## 
## Mann-Whitney test results for Jul tmin: actual vs. RCP4.5 2050 
## The H0 is rejected as p < 0.05, the HA accepted (medians unequal) 
## 
##  Wilcoxon rank sum test with continuity correction
## 
## data:  x and y
## W = 281180, p-value < 2.2e-16
## alternative hypothesis: true location shift is not equal to 0
## 95 percent confidence interval:
##  -1.694382 -1.362100
## sample estimates:
## difference in location 
##               -1.52825 
## 
## Mann-Whitney test results for Jul tmin: RCP4.5 2050 vs. RCP4.5 2070 
## The H0 is rejected as p < 0.05, the HA accepted (medians unequal) 
## 
##  Wilcoxon rank sum test with continuity correction
## 
## data:  x and y
## W = 418760, p-value = 3.153e-10
## alternative hypothesis: true location shift is not equal to 0
## 95 percent confidence interval:
##  -0.6468536 -0.3456331
## sample estimates:
## difference in location 
##             -0.4988544 
## 
## Mann-Whitney test results for Jul tmin: actual vs. RCP4.5 2070 
## The H0 is rejected as p < 0.05, the HA accepted (medians unequal) 
## 
##  Wilcoxon rank sum test with continuity correction
## 
## data:  x and y
## W = 220860, p-value < 2.2e-16
## alternative hypothesis: true location shift is not equal to 0
## 95 percent confidence interval:
##  -2.190066 -1.858627
## sample estimates:
## difference in location 
##              -2.023982 
## 
## Mann-Whitney test results for Jul tmin: actual vs. RCP8.5 2050 
## The H0 is rejected as p < 0.05, the HA accepted (medians unequal) 
## 
##  Wilcoxon rank sum test with continuity correction
## 
## data:  x and y
## W = 220600, p-value < 2.2e-16
## alternative hypothesis: true location shift is not equal to 0
## 95 percent confidence interval:
##  -2.194287 -1.862029
## sample estimates:
## difference in location 
##              -2.028149 
## 
## Mann-Whitney test results for Jul tmin: RCP8.5 2050 vs. RCP8.5 2070 
## The H0 is rejected as p < 0.05, the HA accepted (medians unequal) 
## 
##  Wilcoxon rank sum test with continuity correction
## 
## data:  x and y
## W = 287790, p-value < 2.2e-16
## alternative hypothesis: true location shift is not equal to 0
## 95 percent confidence interval:
##  -1.528188 -1.224984
## sample estimates:
## difference in location 
##              -1.378459 
## 
## Mann-Whitney test results for Jul tmin: actual vs. RCP8.5 2070 
## The H0 is rejected as p < 0.05, the HA accepted (medians unequal) 
## 
##  Wilcoxon rank sum test with continuity correction
## 
## data:  x and y
## W = 90801, p-value < 2.2e-16
## alternative hypothesis: true location shift is not equal to 0
## 95 percent confidence interval:
##  -3.572294 -3.239679
## sample estimates:
## difference in location 
##              -3.405919 
## 
## Mann-Whitney test results for Jul tmax: actual vs. RCP4.5 2050 
## The H0 is rejected as p < 0.05, the HA accepted (medians unequal) 
## 
##  Wilcoxon rank sum test with continuity correction
## 
## data:  x and y
## W = 65610, p-value < 2.2e-16
## alternative hypothesis: true location shift is not equal to 0
## 95 percent confidence interval:
##  -3.172465 -2.923519
## sample estimates:
## difference in location 
##               -3.04905 
## 
## Mann-Whitney test results for Jul tmax: RCP4.5 2050 vs. RCP4.5 2070 
## The H0 is rejected as p < 0.05, the HA accepted (medians unequal) 
## 
##  Wilcoxon rank sum test with continuity correction
## 
## data:  x and y
## W = 410120, p-value = 3.388e-12
## alternative hypothesis: true location shift is not equal to 0
## 95 percent confidence interval:
##  -0.5626666 -0.3210696
## sample estimates:
## difference in location 
##             -0.4418916 
## 
## Mann-Whitney test results for Jul tmax: actual vs. RCP4.5 2070 
## The H0 is rejected as p < 0.05, the HA accepted (medians unequal) 
## 
##  Wilcoxon rank sum test with continuity correction
## 
## data:  x and y
## W = 43227, p-value < 2.2e-16
## alternative hypothesis: true location shift is not equal to 0
## 95 percent confidence interval:
##  -3.615567 -3.365478
## sample estimates:
## difference in location 
##               -3.49107 
## 
## Mann-Whitney test results for Jul tmax: actual vs. RCP8.5 2050 
## The H0 is rejected as p < 0.05, the HA accepted (medians unequal) 
## 
##  Wilcoxon rank sum test with continuity correction
## 
## data:  x and y
## W = 35106, p-value < 2.2e-16
## alternative hypothesis: true location shift is not equal to 0
## 95 percent confidence interval:
##  -3.817813 -3.568102
## sample estimates:
## difference in location 
##               -3.69366 
## 
## Mann-Whitney test results for Jul tmax: RCP8.5 2050 vs. RCP8.5 2070 
## The H0 is rejected as p < 0.05, the HA accepted (medians unequal) 
## 
##  Wilcoxon rank sum test with continuity correction
## 
## data:  x and y
## W = 252480, p-value < 2.2e-16
## alternative hypothesis: true location shift is not equal to 0
## 95 percent confidence interval:
##  -1.444914 -1.204619
## sample estimates:
## difference in location 
##               -1.32335 
## 
## Mann-Whitney test results for Jul tmax: actual vs. RCP8.5 2070 
## The H0 is rejected as p < 0.05, the HA accepted (medians unequal) 
## 
##  Wilcoxon rank sum test with continuity correction
## 
## data:  x and y
## W = 7505, p-value < 2.2e-16
## alternative hypothesis: true location shift is not equal to 0
## 95 percent confidence interval:
##  -5.141433 -4.892096
## sample estimates:
## difference in location 
##              -5.017369 
## 
## Mann-Whitney test results for Aug precipitation: actual vs. RCP4.5 2050 
## The H0 is rejected as p < 0.05, the HA accepted (medians unequal) 
## 
##  Wilcoxon rank sum test with continuity correction
## 
## data:  x and y
## W = 187740, p-value < 2.2e-16
## alternative hypothesis: true location shift is not equal to 0
## 95 percent confidence interval:
##  -100.99272  -86.25798
## sample estimates:
## difference in location 
##              -93.64889 
## 
## Mann-Whitney test results for Aug precipitation: RCP4.5 2050 vs. RCP4.5 2070 
## The HA is rejected as p >= 0.05, the H0 accepted (medians equal) 
## 
##  Wilcoxon rank sum test with continuity correction
## 
## data:  x and y
## W = 499720, p-value = 0.9826
## alternative hypothesis: true location shift is not equal to 0
## 95 percent confidence interval:
##  -7.029890  6.914812
## sample estimates:
## difference in location 
##            -0.06753131 
## 
## Mann-Whitney test results for Aug precipitation: actual vs. RCP4.5 2070 
## The H0 is rejected as p < 0.05, the HA accepted (medians unequal) 
## 
##  Wilcoxon rank sum test with continuity correction
## 
## data:  x and y
## W = 185110, p-value < 2.2e-16
## alternative hypothesis: true location shift is not equal to 0
## 95 percent confidence interval:
##  -101.06069  -86.43041
## sample estimates:
## difference in location 
##              -93.75963 
## 
## Mann-Whitney test results for Aug precipitation: actual vs. RCP8.5 2050 
## The H0 is rejected as p < 0.05, the HA accepted (medians unequal) 
## 
##  Wilcoxon rank sum test with continuity correction
## 
## data:  x and y
## W = 180380, p-value < 2.2e-16
## alternative hypothesis: true location shift is not equal to 0
## 95 percent confidence interval:
##  -104.0710  -89.2036
## sample estimates:
## difference in location 
##              -96.60792 
## 
## Mann-Whitney test results for Aug precipitation: RCP8.5 2050 vs. RCP8.5 2070 
## The H0 is rejected as p < 0.05, the HA accepted (medians unequal) 
## 
##  Wilcoxon rank sum test with continuity correction
## 
## data:  x and y
## W = 405660, p-value = 2.756e-13
## alternative hypothesis: true location shift is not equal to 0
## 95 percent confidence interval:
##  -38.16282 -22.60276
## sample estimates:
## difference in location 
##              -30.48305 
## 
## Mann-Whitney test results for Aug precipitation: actual vs. RCP8.5 2070 
## The H0 is rejected as p < 0.05, the HA accepted (medians unequal) 
## 
##  Wilcoxon rank sum test with continuity correction
## 
## data:  x and y
## W = 133460, p-value < 2.2e-16
## alternative hypothesis: true location shift is not equal to 0
## 95 percent confidence interval:
##  -133.8545 -117.3416
## sample estimates:
## difference in location 
##              -125.5864 
## 
## Mann-Whitney test results for Aug tmin: actual vs. RCP4.5 2050 
## The H0 is rejected as p < 0.05, the HA accepted (medians unequal) 
## 
##  Wilcoxon rank sum test with continuity correction
## 
## data:  x and y
## W = 320850, p-value < 2.2e-16
## alternative hypothesis: true location shift is not equal to 0
## 95 percent confidence interval:
##  -1.404278 -1.068941
## sample estimates:
## difference in location 
##               -1.23627 
## 
## Mann-Whitney test results for Aug tmin: RCP4.5 2050 vs. RCP4.5 2070 
## The H0 is rejected as p < 0.05, the HA accepted (medians unequal) 
## 
##  Wilcoxon rank sum test with continuity correction
## 
## data:  x and y
## W = 418190, p-value = 2.364e-10
## alternative hypothesis: true location shift is not equal to 0
## 95 percent confidence interval:
##  -0.6614465 -0.3539778
## sample estimates:
## difference in location 
##             -0.5053493 
## 
## Mann-Whitney test results for Aug tmin: actual vs. RCP4.5 2070 
## The H0 is rejected as p < 0.05, the HA accepted (medians unequal) 
## 
##  Wilcoxon rank sum test with continuity correction
## 
## data:  x and y
## W = 256890, p-value < 2.2e-16
## alternative hypothesis: true location shift is not equal to 0
## 95 percent confidence interval:
##  -1.912898 -1.576867
## sample estimates:
## difference in location 
##              -1.744606 
## 
## Mann-Whitney test results for Aug tmin: actual vs. RCP8.5 2050 
## The H0 is rejected as p < 0.05, the HA accepted (medians unequal) 
## 
##  Wilcoxon rank sum test with continuity correction
## 
## data:  x and y
## W = 247130, p-value < 2.2e-16
## alternative hypothesis: true location shift is not equal to 0
## 95 percent confidence interval:
##  -1.992547 -1.656070
## sample estimates:
## difference in location 
##              -1.824142 
## 
## Mann-Whitney test results for Aug tmin: RCP8.5 2050 vs. RCP8.5 2070 
## The H0 is rejected as p < 0.05, the HA accepted (medians unequal) 
## 
##  Wilcoxon rank sum test with continuity correction
## 
## data:  x and y
## W = 274970, p-value < 2.2e-16
## alternative hypothesis: true location shift is not equal to 0
## 95 percent confidence interval:
##  -1.646828 -1.337526
## sample estimates:
## difference in location 
##              -1.494273 
## 
## Mann-Whitney test results for Aug tmin: actual vs. RCP8.5 2070 
## The H0 is rejected as p < 0.05, the HA accepted (medians unequal) 
## 
##  Wilcoxon rank sum test with continuity correction
## 
## data:  x and y
## W = 99832, p-value < 2.2e-16
## alternative hypothesis: true location shift is not equal to 0
## 95 percent confidence interval:
##  -3.487157 -3.149285
## sample estimates:
## difference in location 
##              -3.318241 
## 
## Mann-Whitney test results for Aug tmax: actual vs. RCP4.5 2050 
## The H0 is rejected as p < 0.05, the HA accepted (medians unequal) 
## 
##  Wilcoxon rank sum test with continuity correction
## 
## data:  x and y
## W = 47667, p-value < 2.2e-16
## alternative hypothesis: true location shift is not equal to 0
## 95 percent confidence interval:
##  -3.655320 -3.399388
## sample estimates:
## difference in location 
##              -3.527472 
## 
## Mann-Whitney test results for Aug tmax: RCP4.5 2050 vs. RCP4.5 2070 
## The H0 is rejected as p < 0.05, the HA accepted (medians unequal) 
## 
##  Wilcoxon rank sum test with continuity correction
## 
## data:  x and y
## W = 403680, p-value = 8.712e-14
## alternative hypothesis: true location shift is not equal to 0
## 95 percent confidence interval:
##  -0.6218945 -0.3679298
## sample estimates:
## difference in location 
##             -0.4973079 
## 
## Mann-Whitney test results for Aug tmax: actual vs. RCP4.5 2070 
## The H0 is rejected as p < 0.05, the HA accepted (medians unequal) 
## 
##  Wilcoxon rank sum test with continuity correction
## 
## data:  x and y
## W = 29094, p-value < 2.2e-16
## alternative hypothesis: true location shift is not equal to 0
## 95 percent confidence interval:
##  -4.150742 -3.894683
## sample estimates:
## difference in location 
##               -4.02277 
## 
## Mann-Whitney test results for Aug tmax: actual vs. RCP8.5 2050 
## The H0 is rejected as p < 0.05, the HA accepted (medians unequal) 
## 
##  Wilcoxon rank sum test with continuity correction
## 
## data:  x and y
## W = 24733, p-value < 2.2e-16
## alternative hypothesis: true location shift is not equal to 0
## 95 percent confidence interval:
##  -4.282562 -4.027897
## sample estimates:
## difference in location 
##              -4.154773 
## 
## Mann-Whitney test results for Aug tmax: RCP8.5 2050 vs. RCP8.5 2070 
## The H0 is rejected as p < 0.05, the HA accepted (medians unequal) 
## 
##  Wilcoxon rank sum test with continuity correction
## 
## data:  x and y
## W = 228190, p-value < 2.2e-16
## alternative hypothesis: true location shift is not equal to 0
## 95 percent confidence interval:
##  -1.661552 -1.408758
## sample estimates:
## difference in location 
##              -1.534363 
## 
## Mann-Whitney test results for Aug tmax: actual vs. RCP8.5 2070 
## The H0 is rejected as p < 0.05, the HA accepted (medians unequal) 
## 
##  Wilcoxon rank sum test with continuity correction
## 
## data:  x and y
## W = 3852, p-value < 2.2e-16
## alternative hypothesis: true location shift is not equal to 0
## 95 percent confidence interval:
##  -5.818053 -5.563234
## sample estimates:
## difference in location 
##              -5.690325 
## 
## Mann-Whitney test results for Sep precipitation: actual vs. RCP4.5 2050 
## The H0 is rejected as p < 0.05, the HA accepted (medians unequal) 
## 
##  Wilcoxon rank sum test with continuity correction
## 
## data:  x and y
## W = 341850, p-value < 2.2e-16
## alternative hypothesis: true location shift is not equal to 0
## 95 percent confidence interval:
##  -38.84341 -27.94960
## sample estimates:
## difference in location 
##              -33.31498 
## 
## Mann-Whitney test results for Sep precipitation: RCP4.5 2050 vs. RCP4.5 2070 
## The H0 is rejected as p < 0.05, the HA accepted (medians unequal) 
## 
##  Wilcoxon rank sum test with continuity correction
## 
## data:  x and y
## W = 534260, p-value = 0.007973
## alternative hypothesis: true location shift is not equal to 0
## 95 percent confidence interval:
##   1.995953 13.106127
## sample estimates:
## difference in location 
##               7.549098 
## 
## Mann-Whitney test results for Sep precipitation: actual vs. RCP4.5 2070 
## The H0 is rejected as p < 0.05, the HA accepted (medians unequal) 
## 
##  Wilcoxon rank sum test with continuity correction
## 
## data:  x and y
## W = 376010, p-value < 2.2e-16
## alternative hypothesis: true location shift is not equal to 0
## 95 percent confidence interval:
##  -31.14511 -20.39534
## sample estimates:
## difference in location 
##               -25.7484 
## 
## Mann-Whitney test results for Sep precipitation: actual vs. RCP8.5 2050 
## The H0 is rejected as p < 0.05, the HA accepted (medians unequal) 
## 
##  Wilcoxon rank sum test with continuity correction
## 
## data:  x and y
## W = 397980, p-value = 2.782e-15
## alternative hypothesis: true location shift is not equal to 0
## 95 percent confidence interval:
##  -25.47412 -15.16997
## sample estimates:
## difference in location 
##               -20.2786 
## 
## Mann-Whitney test results for Sep precipitation: RCP8.5 2050 vs. RCP8.5 2070 
## The H0 is rejected as p < 0.05, the HA accepted (medians unequal) 
## 
##  Wilcoxon rank sum test with continuity correction
## 
## data:  x and y
## W = 562210, p-value = 1.452e-06
## alternative hypothesis: true location shift is not equal to 0
## 95 percent confidence interval:
##   7.908177 18.412716
## sample estimates:
## difference in location 
##               13.12537 
## 
## Mann-Whitney test results for Sep precipitation: actual vs. RCP8.5 2070 
## The H0 is rejected as p < 0.05, the HA accepted (medians unequal) 
## 
##  Wilcoxon rank sum test with continuity correction
## 
## data:  x and y
## W = 465300, p-value = 0.007211
## alternative hypothesis: true location shift is not equal to 0
## 95 percent confidence interval:
##  -12.006788  -1.828628
## sample estimates:
## difference in location 
##              -6.869505 
## 
## Mann-Whitney test results for Sep tmin: actual vs. RCP4.5 2050 
## The H0 is rejected as p < 0.05, the HA accepted (medians unequal) 
## 
##  Wilcoxon rank sum test with continuity correction
## 
## data:  x and y
## W = 339770, p-value < 2.2e-16
## alternative hypothesis: true location shift is not equal to 0
## 95 percent confidence interval:
##  -1.2542931 -0.9218163
## sample estimates:
## difference in location 
##              -1.088573 
## 
## Mann-Whitney test results for Sep tmin: RCP4.5 2050 vs. RCP4.5 2070 
## The H0 is rejected as p < 0.05, the HA accepted (medians unequal) 
## 
##  Wilcoxon rank sum test with continuity correction
## 
## data:  x and y
## W = 421270, p-value = 1.081e-09
## alternative hypothesis: true location shift is not equal to 0
## 95 percent confidence interval:
##  -0.6448913 -0.3365555
## sample estimates:
## difference in location 
##             -0.4932037 
## 
## Mann-Whitney test results for Sep tmin: actual vs. RCP4.5 2070 
## The H0 is rejected as p < 0.05, the HA accepted (medians unequal) 
## 
##  Wilcoxon rank sum test with continuity correction
## 
## data:  x and y
## W = 275590, p-value < 2.2e-16
## alternative hypothesis: true location shift is not equal to 0
## 95 percent confidence interval:
##  -1.744990 -1.412274
## sample estimates:
## difference in location 
##              -1.579581 
## 
## Mann-Whitney test results for Sep tmin: actual vs. RCP8.5 2050 
## The H0 is rejected as p < 0.05, the HA accepted (medians unequal) 
## 
##  Wilcoxon rank sum test with continuity correction
## 
## data:  x and y
## W = 272480, p-value < 2.2e-16
## alternative hypothesis: true location shift is not equal to 0
## 95 percent confidence interval:
##  -1.776421 -1.441856
## sample estimates:
## difference in location 
##              -1.608963 
## 
## Mann-Whitney test results for Sep tmin: RCP8.5 2050 vs. RCP8.5 2070 
## The H0 is rejected as p < 0.05, the HA accepted (medians unequal) 
## 
##  Wilcoxon rank sum test with continuity correction
## 
## data:  x and y
## W = 301630, p-value < 2.2e-16
## alternative hypothesis: true location shift is not equal to 0
## 95 percent confidence interval:
##  -1.459713 -1.148766
## sample estimates:
## difference in location 
##              -1.301899 
## 
## Mann-Whitney test results for Sep tmin: actual vs. RCP8.5 2070 
## The H0 is rejected as p < 0.05, the HA accepted (medians unequal) 
## 
##  Wilcoxon rank sum test with continuity correction
## 
## data:  x and y
## W = 131460, p-value < 2.2e-16
## alternative hypothesis: true location shift is not equal to 0
## 95 percent confidence interval:
##  -3.080339 -2.745433
## sample estimates:
## difference in location 
##              -2.912977 
## 
## Mann-Whitney test results for Sep tmax: actual vs. RCP4.5 2050 
## The H0 is rejected as p < 0.05, the HA accepted (medians unequal) 
## 
##  Wilcoxon rank sum test with continuity correction
## 
## data:  x and y
## W = 26110, p-value < 2.2e-16
## alternative hypothesis: true location shift is not equal to 0
## 95 percent confidence interval:
##  -4.092325 -3.847035
## sample estimates:
## difference in location 
##              -3.969225 
## 
## Mann-Whitney test results for Sep tmax: RCP4.5 2050 vs. RCP4.5 2070 
## The H0 is rejected as p < 0.05, the HA accepted (medians unequal) 
## 
##  Wilcoxon rank sum test with continuity correction
## 
## data:  x and y
## W = 390940, p-value < 2.2e-16
## alternative hypothesis: true location shift is not equal to 0
## 95 percent confidence interval:
##  -0.6507400 -0.4109078
## sample estimates:
## difference in location 
##             -0.5303098 
## 
## Mann-Whitney test results for Sep tmax: actual vs. RCP4.5 2070 
## The H0 is rejected as p < 0.05, the HA accepted (medians unequal) 
## 
##  Wilcoxon rank sum test with continuity correction
## 
## data:  x and y
## W = 14228, p-value < 2.2e-16
## alternative hypothesis: true location shift is not equal to 0
## 95 percent confidence interval:
##  -4.623181 -4.377867
## sample estimates:
## difference in location 
##              -4.500324 
## 
## Mann-Whitney test results for Sep tmax: actual vs. RCP8.5 2050 
## The H0 is rejected as p < 0.05, the HA accepted (medians unequal) 
## 
##  Wilcoxon rank sum test with continuity correction
## 
## data:  x and y
## W = 12593, p-value < 2.2e-16
## alternative hypothesis: true location shift is not equal to 0
## 95 percent confidence interval:
##  -4.729349 -4.483571
## sample estimates:
## difference in location 
##              -4.606161 
## 
## Mann-Whitney test results for Sep tmax: RCP8.5 2050 vs. RCP8.5 2070 
## The H0 is rejected as p < 0.05, the HA accepted (medians unequal) 
## 
##  Wilcoxon rank sum test with continuity correction
## 
## data:  x and y
## W = 237990, p-value < 2.2e-16
## alternative hypothesis: true location shift is not equal to 0
## 95 percent confidence interval:
##  -1.519294 -1.277676
## sample estimates:
## difference in location 
##              -1.399999 
## 
## Mann-Whitney test results for Sep tmax: actual vs. RCP8.5 2070 
## The H0 is rejected as p < 0.05, the HA accepted (medians unequal) 
## 
##  Wilcoxon rank sum test with continuity correction
## 
## data:  x and y
## W = 2035, p-value < 2.2e-16
## alternative hypothesis: true location shift is not equal to 0
## 95 percent confidence interval:
##  -6.127921 -5.882076
## sample estimates:
## difference in location 
##              -6.004746 
## 
## Mann-Whitney test results for Oct precipitation: actual vs. RCP4.5 2050 
## The H0 is rejected as p < 0.05, the HA accepted (medians unequal) 
## 
##  Wilcoxon rank sum test with continuity correction
## 
## data:  x and y
## W = 467980, p-value = 0.01316
## alternative hypothesis: true location shift is not equal to 0
## 95 percent confidence interval:
##  -9.035886 -1.034567
## sample estimates:
## difference in location 
##              -4.970523 
## 
## Mann-Whitney test results for Oct precipitation: RCP4.5 2050 vs. RCP4.5 2070 
## The H0 is rejected as p < 0.05, the HA accepted (medians unequal) 
## 
##  Wilcoxon rank sum test with continuity correction
## 
## data:  x and y
## W = 547580, p-value = 0.0002294
## alternative hypothesis: true location shift is not equal to 0
## 95 percent confidence interval:
##   3.545023 11.307380
## sample estimates:
## difference in location 
##               7.387732 
## 
## Mann-Whitney test results for Oct precipitation: actual vs. RCP4.5 2070 
## The HA is rejected as p >= 0.05, the H0 accepted (medians equal) 
## 
##  Wilcoxon rank sum test with continuity correction
## 
## data:  x and y
## W = 515930, p-value = 0.2173
## alternative hypothesis: true location shift is not equal to 0
## 95 percent confidence interval:
##  -1.465824  6.221040
## sample estimates:
## difference in location 
##               2.415847 
## 
## Mann-Whitney test results for Oct precipitation: actual vs. RCP8.5 2050 
## The HA is rejected as p >= 0.05, the H0 accepted (medians equal) 
## 
##  Wilcoxon rank sum test with continuity correction
## 
## data:  x and y
## W = 508790, p-value = 0.496
## alternative hypothesis: true location shift is not equal to 0
## 95 percent confidence interval:
##  -2.521305  5.120459
## sample estimates:
## difference in location 
##               1.347207 
## 
## Mann-Whitney test results for Oct precipitation: RCP8.5 2050 vs. RCP8.5 2070 
## The H0 is rejected as p < 0.05, the HA accepted (medians unequal) 
## 
##  Wilcoxon rank sum test with continuity correction
## 
## data:  x and y
## W = 715760, p-value < 2.2e-16
## alternative hypothesis: true location shift is not equal to 0
## 95 percent confidence interval:
##  30.36456 37.62580
## sample estimates:
## difference in location 
##               33.93222 
## 
## Mann-Whitney test results for Oct precipitation: actual vs. RCP8.5 2070 
## The H0 is rejected as p < 0.05, the HA accepted (medians unequal) 
## 
##  Wilcoxon rank sum test with continuity correction
## 
## data:  x and y
## W = 736730, p-value < 2.2e-16
## alternative hypothesis: true location shift is not equal to 0
## 95 percent confidence interval:
##  32.00315 38.72705
## sample estimates:
## difference in location 
##               35.36735 
## 
## Mann-Whitney test results for Oct tmin: actual vs. RCP4.5 2050 
## The H0 is rejected as p < 0.05, the HA accepted (medians unequal) 
## 
##  Wilcoxon rank sum test with continuity correction
## 
## data:  x and y
## W = 302730, p-value < 2.2e-16
## alternative hypothesis: true location shift is not equal to 0
## 95 percent confidence interval:
##  -1.485849 -1.162243
## sample estimates:
## difference in location 
##              -1.323821 
## 
## Mann-Whitney test results for Oct tmin: RCP4.5 2050 vs. RCP4.5 2070 
## The H0 is rejected as p < 0.05, the HA accepted (medians unequal) 
## 
##  Wilcoxon rank sum test with continuity correction
## 
## data:  x and y
## W = 424500, p-value = 5.019e-09
## alternative hypothesis: true location shift is not equal to 0
## 95 percent confidence interval:
##  -0.6065884 -0.3064507
## sample estimates:
## difference in location 
##             -0.4561758 
## 
## Mann-Whitney test results for Oct tmin: actual vs. RCP4.5 2070 
## The H0 is rejected as p < 0.05, the HA accepted (medians unequal) 
## 
##  Wilcoxon rank sum test with continuity correction
## 
## data:  x and y
## W = 244450, p-value < 2.2e-16
## alternative hypothesis: true location shift is not equal to 0
## 95 percent confidence interval:
##  -1.943273 -1.618454
## sample estimates:
## difference in location 
##              -1.779936 
## 
## Mann-Whitney test results for Oct tmin: actual vs. RCP8.5 2050 
## The H0 is rejected as p < 0.05, the HA accepted (medians unequal) 
## 
##  Wilcoxon rank sum test with continuity correction
## 
## data:  x and y
## W = 235310, p-value < 2.2e-16
## alternative hypothesis: true location shift is not equal to 0
## 95 percent confidence interval:
##  -2.014165 -1.689962
## sample estimates:
## difference in location 
##              -1.851583 
## 
## Mann-Whitney test results for Oct tmin: RCP8.5 2050 vs. RCP8.5 2070 
## The H0 is rejected as p < 0.05, the HA accepted (medians unequal) 
## 
##  Wilcoxon rank sum test with continuity correction
## 
## data:  x and y
## W = 300050, p-value < 2.2e-16
## alternative hypothesis: true location shift is not equal to 0
## 95 percent confidence interval:
##  -1.415721 -1.116485
## sample estimates:
## difference in location 
##              -1.267475 
## 
## Mann-Whitney test results for Oct tmin: actual vs. RCP8.5 2070 
## The H0 is rejected as p < 0.05, the HA accepted (medians unequal) 
## 
##  Wilcoxon rank sum test with continuity correction
## 
## data:  x and y
## W = 107090, p-value < 2.2e-16
## alternative hypothesis: true location shift is not equal to 0
## 95 percent confidence interval:
##  -3.279949 -2.956303
## sample estimates:
## difference in location 
##              -3.117362 
## 
## Mann-Whitney test results for Oct tmax: actual vs. RCP4.5 2050 
## The H0 is rejected as p < 0.05, the HA accepted (medians unequal) 
## 
##  Wilcoxon rank sum test with continuity correction
## 
## data:  x and y
## W = 17181, p-value < 2.2e-16
## alternative hypothesis: true location shift is not equal to 0
## 95 percent confidence interval:
##  -4.036842 -3.805393
## sample estimates:
## difference in location 
##              -3.921671 
## 
## Mann-Whitney test results for Oct tmax: RCP4.5 2050 vs. RCP4.5 2070 
## The H0 is rejected as p < 0.05, the HA accepted (medians unequal) 
## 
##  Wilcoxon rank sum test with continuity correction
## 
## data:  x and y
## W = 365370, p-value < 2.2e-16
## alternative hypothesis: true location shift is not equal to 0
## 95 percent confidence interval:
##  -0.7083903 -0.4916097
## sample estimates:
## difference in location 
##                   -0.6 
## 
## Mann-Whitney test results for Oct tmax: actual vs. RCP4.5 2070 
## The H0 is rejected as p < 0.05, the HA accepted (medians unequal) 
## 
##  Wilcoxon rank sum test with continuity correction
## 
## data:  x and y
## W = 7357, p-value < 2.2e-16
## alternative hypothesis: true location shift is not equal to 0
## 95 percent confidence interval:
##  -4.636842 -4.405393
## sample estimates:
## difference in location 
##              -4.521692 
## 
## Mann-Whitney test results for Oct tmax: actual vs. RCP8.5 2050 
## The H0 is rejected as p < 0.05, the HA accepted (medians unequal) 
## 
##  Wilcoxon rank sum test with continuity correction
## 
## data:  x and y
## W = 6344, p-value < 2.2e-16
## alternative hypothesis: true location shift is not equal to 0
## 95 percent confidence interval:
##  -4.756071 -4.524742
## sample estimates:
## difference in location 
##              -4.641206 
## 
## Mann-Whitney test results for Oct tmax: RCP8.5 2050 vs. RCP8.5 2070 
## The H0 is rejected as p < 0.05, the HA accepted (medians unequal) 
## 
##  Wilcoxon rank sum test with continuity correction
## 
## data:  x and y
## W = 201550, p-value < 2.2e-16
## alternative hypothesis: true location shift is not equal to 0
## 95 percent confidence interval:
##  -1.588729 -1.371366
## sample estimates:
## difference in location 
##              -1.481894 
## 
## Mann-Whitney test results for Oct tmax: actual vs. RCP8.5 2070 
## The H0 is rejected as p < 0.05, the HA accepted (medians unequal) 
## 
##  Wilcoxon rank sum test with continuity correction
## 
## data:  x and y
## W = 665, p-value < 2.2e-16
## alternative hypothesis: true location shift is not equal to 0
## 95 percent confidence interval:
##  -6.236842 -6.005393
## sample estimates:
## difference in location 
##              -6.121671 
## 
## Mann-Whitney test results for Nov precipitation: actual vs. RCP4.5 2050 
## The H0 is rejected as p < 0.05, the HA accepted (medians unequal) 
## 
##  Wilcoxon rank sum test with continuity correction
## 
## data:  x and y
## W = 278960, p-value < 2.2e-16
## alternative hypothesis: true location shift is not equal to 0
## 95 percent confidence interval:
##  -45.60254 -35.26236
## sample estimates:
## difference in location 
##              -40.34177 
## 
## Mann-Whitney test results for Nov precipitation: RCP4.5 2050 vs. RCP4.5 2070 
## The H0 is rejected as p < 0.05, the HA accepted (medians unequal) 
## 
##  Wilcoxon rank sum test with continuity correction
## 
## data:  x and y
## W = 552480, p-value = 4.823e-05
## alternative hypothesis: true location shift is not equal to 0
## 95 percent confidence interval:
##   5.352663 15.093957
## sample estimates:
## difference in location 
##               10.21016 
## 
## Mann-Whitney test results for Nov precipitation: actual vs. RCP4.5 2070 
## The H0 is rejected as p < 0.05, the HA accepted (medians unequal) 
## 
##  Wilcoxon rank sum test with continuity correction
## 
## data:  x and y
## W = 329250, p-value < 2.2e-16
## alternative hypothesis: true location shift is not equal to 0
## 95 percent confidence interval:
##  -35.31966 -25.34429
## sample estimates:
## difference in location 
##              -30.27031 
## 
## Mann-Whitney test results for Nov precipitation: actual vs. RCP8.5 2050 
## The H0 is rejected as p < 0.05, the HA accepted (medians unequal) 
## 
##  Wilcoxon rank sum test with continuity correction
## 
## data:  x and y
## W = 326700, p-value < 2.2e-16
## alternative hypothesis: true location shift is not equal to 0
## 95 percent confidence interval:
##  -34.95788 -25.31053
## sample estimates:
## difference in location 
##              -30.05469 
## 
## Mann-Whitney test results for Nov precipitation: RCP8.5 2050 vs. RCP8.5 2070 
## The H0 is rejected as p < 0.05, the HA accepted (medians unequal) 
## 
##  Wilcoxon rank sum test with continuity correction
## 
## data:  x and y
## W = 565130, p-value = 4.575e-07
## alternative hypothesis: true location shift is not equal to 0
## 95 percent confidence interval:
##   7.373793 16.477984
## sample estimates:
## difference in location 
##               11.93575 
## 
## Mann-Whitney test results for Nov precipitation: actual vs. RCP8.5 2070 
## The H0 is rejected as p < 0.05, the HA accepted (medians unequal) 
## 
##  Wilcoxon rank sum test with continuity correction
## 
## data:  x and y
## W = 392680, p-value < 2.2e-16
## alternative hypothesis: true location shift is not equal to 0
## 95 percent confidence interval:
##  -22.81529 -13.66287
## sample estimates:
## difference in location 
##              -18.11706 
## 
## Mann-Whitney test results for Nov tmin: actual vs. RCP4.5 2050 
## The H0 is rejected as p < 0.05, the HA accepted (medians unequal) 
## 
##  Wilcoxon rank sum test with continuity correction
## 
## data:  x and y
## W = 301840, p-value < 2.2e-16
## alternative hypothesis: true location shift is not equal to 0
## 95 percent confidence interval:
##  -1.486648 -1.164407
## sample estimates:
## difference in location 
##              -1.325242 
## 
## Mann-Whitney test results for Nov tmin: RCP4.5 2050 vs. RCP4.5 2070 
## The H0 is rejected as p < 0.05, the HA accepted (medians unequal) 
## 
##  Wilcoxon rank sum test with continuity correction
## 
## data:  x and y
## W = 432940, p-value = 2.064e-07
## alternative hypothesis: true location shift is not equal to 0
## 95 percent confidence interval:
##  -0.5493071 -0.2521532
## sample estimates:
## difference in location 
##             -0.4000175 
## 
## Mann-Whitney test results for Nov tmin: actual vs. RCP4.5 2070 
## The H0 is rejected as p < 0.05, the HA accepted (medians unequal) 
## 
##  Wilcoxon rank sum test with continuity correction
## 
## data:  x and y
## W = 250140, p-value < 2.2e-16
## alternative hypothesis: true location shift is not equal to 0
## 95 percent confidence interval:
##  -1.887312 -1.565103
## sample estimates:
## difference in location 
##              -1.725946 
## 
## Mann-Whitney test results for Nov tmin: actual vs. RCP8.5 2050 
## The H0 is rejected as p < 0.05, the HA accepted (medians unequal) 
## 
##  Wilcoxon rank sum test with continuity correction
## 
## data:  x and y
## W = 235410, p-value < 2.2e-16
## alternative hypothesis: true location shift is not equal to 0
## 95 percent confidence interval:
##  -2.014045 -1.691126
## sample estimates:
## difference in location 
##              -1.852465 
## 
## Mann-Whitney test results for Nov tmin: RCP8.5 2050 vs. RCP8.5 2070 
## The H0 is rejected as p < 0.05, the HA accepted (medians unequal) 
## 
##  Wilcoxon rank sum test with continuity correction
## 
## data:  x and y
## W = 317940, p-value < 2.2e-16
## alternative hypothesis: true location shift is not equal to 0
## 95 percent confidence interval:
##  -1.2940259 -0.9940073
## sample estimates:
## difference in location 
##                -1.1445 
## 
## Mann-Whitney test results for Nov tmin: actual vs. RCP8.5 2070 
## The H0 is rejected as p < 0.05, the HA accepted (medians unequal) 
## 
##  Wilcoxon rank sum test with continuity correction
## 
## data:  x and y
## W = 116330, p-value < 2.2e-16
## alternative hypothesis: true location shift is not equal to 0
## 95 percent confidence interval:
##  -3.157733 -2.835767
## sample estimates:
## difference in location 
##              -2.996623 
## 
## Mann-Whitney test results for Nov tmax: actual vs. RCP4.5 2050 
## The H0 is rejected as p < 0.05, the HA accepted (medians unequal) 
## 
##  Wilcoxon rank sum test with continuity correction
## 
## data:  x and y
## W = 20465, p-value < 2.2e-16
## alternative hypothesis: true location shift is not equal to 0
## 95 percent confidence interval:
##  -3.742256 -3.519402
## sample estimates:
## difference in location 
##              -3.631001 
## 
## Mann-Whitney test results for Nov tmax: RCP4.5 2050 vs. RCP4.5 2070 
## The H0 is rejected as p < 0.05, the HA accepted (medians unequal) 
## 
##  Wilcoxon rank sum test with continuity correction
## 
## data:  x and y
## W = 379730, p-value < 2.2e-16
## alternative hypothesis: true location shift is not equal to 0
## 95 percent confidence interval:
##  -0.6022343 -0.3977657
## sample estimates:
## difference in location 
##                   -0.5 
## 
## Mann-Whitney test results for Nov tmax: actual vs. RCP4.5 2070 
## The H0 is rejected as p < 0.05, the HA accepted (medians unequal) 
## 
##  Wilcoxon rank sum test with continuity correction
## 
## data:  x and y
## W = 9826, p-value < 2.2e-16
## alternative hypothesis: true location shift is not equal to 0
## 95 percent confidence interval:
##  -4.242256 -4.019398
## sample estimates:
## difference in location 
##              -4.131001 
## 
## Mann-Whitney test results for Nov tmax: actual vs. RCP8.5 2050 
## The H0 is rejected as p < 0.05, the HA accepted (medians unequal) 
## 
##  Wilcoxon rank sum test with continuity correction
## 
## data:  x and y
## W = 7244, p-value < 2.2e-16
## alternative hypothesis: true location shift is not equal to 0
## 95 percent confidence interval:
##  -4.442736 -4.219730
## sample estimates:
## difference in location 
##              -4.331266 
## 
## Mann-Whitney test results for Nov tmax: RCP8.5 2050 vs. RCP8.5 2070 
## The H0 is rejected as p < 0.05, the HA accepted (medians unequal) 
## 
##  Wilcoxon rank sum test with continuity correction
## 
## data:  x and y
## W = 234740, p-value < 2.2e-16
## alternative hypothesis: true location shift is not equal to 0
## 95 percent confidence interval:
##  -1.301901 -1.097307
## sample estimates:
## difference in location 
##                   -1.2 
## 
## Mann-Whitney test results for Nov tmax: actual vs. RCP8.5 2070 
## The H0 is rejected as p < 0.05, the HA accepted (medians unequal) 
## 
##  Wilcoxon rank sum test with continuity correction
## 
## data:  x and y
## W = 1077, p-value < 2.2e-16
## alternative hypothesis: true location shift is not equal to 0
## 95 percent confidence interval:
##  -5.642252 -5.419398
## sample estimates:
## difference in location 
##              -5.531001 
## 
## Mann-Whitney test results for Dic precipitation: actual vs. RCP4.5 2050 
## The H0 is rejected as p < 0.05, the HA accepted (medians unequal) 
## 
##  Wilcoxon rank sum test with continuity correction
## 
## data:  x and y
## W = 179090, p-value < 2.2e-16
## alternative hypothesis: true location shift is not equal to 0
## 95 percent confidence interval:
##  -62.32155 -51.47465
## sample estimates:
## difference in location 
##              -56.77018 
## 
## Mann-Whitney test results for Dic precipitation: RCP4.5 2050 vs. RCP4.5 2070 
## The HA is rejected as p >= 0.05, the H0 accepted (medians equal) 
## 
##  Wilcoxon rank sum test with continuity correction
## 
## data:  x and y
## W = 519390, p-value = 0.1331
## alternative hypothesis: true location shift is not equal to 0
## 95 percent confidence interval:
##  -1.166857  8.616983
## sample estimates:
## difference in location 
##               3.745275 
## 
## Mann-Whitney test results for Dic precipitation: actual vs. RCP4.5 2070 
## The H0 is rejected as p < 0.05, the HA accepted (medians unequal) 
## 
##  Wilcoxon rank sum test with continuity correction
## 
## data:  x and y
## W = 194680, p-value < 2.2e-16
## alternative hypothesis: true location shift is not equal to 0
## 95 percent confidence interval:
##  -58.29537 -47.71118
## sample estimates:
## difference in location 
##              -52.83946 
## 
## Mann-Whitney test results for Dic precipitation: actual vs. RCP8.5 2050 
## The H0 is rejected as p < 0.05, the HA accepted (medians unequal) 
## 
##  Wilcoxon rank sum test with continuity correction
## 
## data:  x and y
## W = 197710, p-value < 2.2e-16
## alternative hypothesis: true location shift is not equal to 0
## 95 percent confidence interval:
##  -56.92656 -46.54621
## sample estimates:
## difference in location 
##              -51.59168 
## 
## Mann-Whitney test results for Dic precipitation: RCP8.5 2050 vs. RCP8.5 2070 
## The H0 is rejected as p < 0.05, the HA accepted (medians unequal) 
## 
##  Wilcoxon rank sum test with continuity correction
## 
## data:  x and y
## W = 453530, p-value = 0.0003202
## alternative hypothesis: true location shift is not equal to 0
## 95 percent confidence interval:
##  -14.17565  -4.20220
## sample estimates:
## difference in location 
##              -9.158194 
## 
## Mann-Whitney test results for Dic precipitation: actual vs. RCP8.5 2070 
## The H0 is rejected as p < 0.05, the HA accepted (medians unequal) 
## 
##  Wilcoxon rank sum test with continuity correction
## 
## data:  x and y
## W = 166040, p-value < 2.2e-16
## alternative hypothesis: true location shift is not equal to 0
## 95 percent confidence interval:
##  -65.77142 -54.62470
## sample estimates:
## difference in location 
##              -60.02914 
## 
## Mann-Whitney test results for Dic tmin: actual vs. RCP4.5 2050 
## The H0 is rejected as p < 0.05, the HA accepted (medians unequal) 
## 
##  Wilcoxon rank sum test with continuity correction
## 
## data:  x and y
## W = 303350, p-value < 2.2e-16
## alternative hypothesis: true location shift is not equal to 0
## 95 percent confidence interval:
##  -1.483315 -1.159977
## sample estimates:
## difference in location 
##              -1.321395 
## 
## Mann-Whitney test results for Dic tmin: RCP4.5 2050 vs. RCP4.5 2070 
## The H0 is rejected as p < 0.05, the HA accepted (medians unequal) 
## 
##  Wilcoxon rank sum test with continuity correction
## 
## data:  x and y
## W = 427340, p-value = 1.839e-08
## alternative hypothesis: true location shift is not equal to 0
## 95 percent confidence interval:
##  -0.5906993 -0.2894341
## sample estimates:
## difference in location 
##             -0.4393557 
## 
## Mann-Whitney test results for Dic tmin: actual vs. RCP4.5 2070 
## The H0 is rejected as p < 0.05, the HA accepted (medians unequal) 
## 
##  Wilcoxon rank sum test with continuity correction
## 
## data:  x and y
## W = 248490, p-value < 2.2e-16
## alternative hypothesis: true location shift is not equal to 0
## 95 percent confidence interval:
##  -1.924361 -1.598857
## sample estimates:
## difference in location 
##              -1.761673 
## 
## Mann-Whitney test results for Dic tmin: actual vs. RCP8.5 2050 
## The H0 is rejected as p < 0.05, the HA accepted (medians unequal) 
## 
##  Wilcoxon rank sum test with continuity correction
## 
## data:  x and y
## W = 231430, p-value < 2.2e-16
## alternative hypothesis: true location shift is not equal to 0
## 95 percent confidence interval:
##  -2.057370 -1.733214
## sample estimates:
## difference in location 
##              -1.895376 
## 
## Mann-Whitney test results for Dic tmin: RCP8.5 2050 vs. RCP8.5 2070 
## The H0 is rejected as p < 0.05, the HA accepted (medians unequal) 
## 
##  Wilcoxon rank sum test with continuity correction
## 
## data:  x and y
## W = 325690, p-value < 2.2e-16
## alternative hypothesis: true location shift is not equal to 0
## 95 percent confidence interval:
##  -1.2436274 -0.9437993
## sample estimates:
## difference in location 
##               -1.09579 
## 
## Mann-Whitney test results for Dic tmin: actual vs. RCP8.5 2070 
## The H0 is rejected as p < 0.05, the HA accepted (medians unequal) 
## 
##  Wilcoxon rank sum test with continuity correction
## 
## data:  x and y
## W = 119120, p-value < 2.2e-16
## alternative hypothesis: true location shift is not equal to 0
## 95 percent confidence interval:
##  -3.150751 -2.826199
## sample estimates:
## difference in location 
##               -2.98841 
## 
## Mann-Whitney test results for Dic tmax: actual vs. RCP4.5 2050 
## The H0 is rejected as p < 0.05, the HA accepted (medians unequal) 
## 
##  Wilcoxon rank sum test with continuity correction
## 
## data:  x and y
## W = 24891, p-value < 2.2e-16
## alternative hypothesis: true location shift is not equal to 0
## 95 percent confidence interval:
##  -3.638023 -3.412683
## sample estimates:
## difference in location 
##              -3.525228 
## 
## Mann-Whitney test results for Dic tmax: RCP4.5 2050 vs. RCP4.5 2070 
## The H0 is rejected as p < 0.05, the HA accepted (medians unequal) 
## 
##  Wilcoxon rank sum test with continuity correction
## 
## data:  x and y
## W = 393000, p-value < 2.2e-16
## alternative hypothesis: true location shift is not equal to 0
## 95 percent confidence interval:
##  -0.5424593 -0.3398208
## sample estimates:
## difference in location 
##              -0.440747 
## 
## Mann-Whitney test results for Dic tmax: actual vs. RCP4.5 2070 
## The H0 is rejected as p < 0.05, the HA accepted (medians unequal) 
## 
##  Wilcoxon rank sum test with continuity correction
## 
## data:  x and y
## W = 12918, p-value < 2.2e-16
## alternative hypothesis: true location shift is not equal to 0
## 95 percent confidence interval:
##  -4.078160 -3.854775
## sample estimates:
## difference in location 
##              -3.965745 
## 
## Mann-Whitney test results for Dic tmax: actual vs. RCP8.5 2050 
## The H0 is rejected as p < 0.05, the HA accepted (medians unequal) 
## 
##  Wilcoxon rank sum test with continuity correction
## 
## data:  x and y
## W = 9598, p-value < 2.2e-16
## alternative hypothesis: true location shift is not equal to 0
## 95 percent confidence interval:
##  -4.278232 -4.054829
## sample estimates:
## difference in location 
##              -4.165893 
## 
## Mann-Whitney test results for Dic tmax: RCP8.5 2050 vs. RCP8.5 2070 
## The H0 is rejected as p < 0.05, the HA accepted (medians unequal) 
## 
##  Wilcoxon rank sum test with continuity correction
## 
## data:  x and y
## W = 270280, p-value < 2.2e-16
## alternative hypothesis: true location shift is not equal to 0
## 95 percent confidence interval:
##  -1.1004554 -0.8995446
## sample estimates:
## difference in location 
##                     -1 
## 
## Mann-Whitney test results for Dic tmax: actual vs. RCP8.5 2070 
## The H0 is rejected as p < 0.05, the HA accepted (medians unequal) 
## 
##  Wilcoxon rank sum test with continuity correction
## 
## data:  x and y
## W = 2060, p-value < 2.2e-16
## alternative hypothesis: true location shift is not equal to 0
## 95 percent confidence interval:
##  -5.278228 -5.054829
## sample estimates:
## difference in location 
##              -5.165893
```

```
# Mann-Whitney test for biovars
for (i in 1:19) {
  my_test19 <- compareMWT(Magn_biovars00[,i],Magn_biovars50[,i],paste(month_list[i],'biovars: actual vs. RCP4.5 2050'))
  my_test20 <- compareMWT(Magn_biovars50[,i],Magn_biovars70[,i],paste(month_list[i],'biovars: RCP4.5 2050 vs. RCP4.5 2070'))
  my_test21 <- compareMWT(Magn_biovars00[,i],Magn_biovars70[,i],paste(month_list[i],'biovars: actual vs. RCP4.5 2070'))
  
  my_test22 <- compareMWT(Magn_biovars00[,i],Magn_biovars50A[,i],paste(month_list[i],'biovars: actual vs. RCP8.5 2050'))
  my_test23 <- compareMWT(Magn_biovars50A[,i],Magn_biovars70A[,i],paste(month_list[i],'biovars: RCP8.5 2050 vs. RCP8.5 2070'))
  my_test24 <- compareMWT(Magn_biovars00[,i],Magn_biovars70A[,i],paste(month_list[i],'biovars: actual vs. RCP8.5 2070'))    
  
  if (my_test19$p.value < 0.05) {
    MWU_biovars_RCP45_matrix[i,1] <- 1
    MWU_biovars_RCP45_matrix_s[i,1] <- my_test19$estimate
  } else {
    MWU_biovars_RCP45_matrix[i,1] <- 0
  }
  if (my_test20$p.value < 0.05) {
    MWU_biovars_RCP45_matrix[i,2] <- 1
    MWU_biovars_RCP45_matrix_s[i,2] <- my_test20$estimate
  } else {
    MWU_biovars_RCP45_matrix[i,2] <- 0
  }  
  if (my_test21$p.value < 0.05) {
    MWU_biovars_RCP45_matrix[i,3] <- 1
    MWU_biovars_RCP45_matrix_s[i,3] <- my_test21$estimate
  } else {
    MWU_biovars_RCP45_matrix[i,3] <- 0
  } 
  if (my_test22$p.value < 0.05) {
    MWU_biovars_RCP85_matrix[i,1] <- 1
    MWU_biovars_RCP85_matrix_s[i,1] <- my_test22$estimate
  } else {
    MWU_biovars_RCP85_matrix[i,1] <- 0
  } 
  if (my_test23$p.value < 0.05) {
    MWU_biovars_RCP85_matrix[i,2] <- 1
    MWU_biovars_RCP85_matrix_s[i,2] <- my_test23$estimate
  } else {
    MWU_biovars_RCP85_matrix[i,2] <- 0
  } 
  if (my_test24$p.value < 0.05) {
    MWU_biovars_RCP85_matrix[i,3] <- 1
    MWU_biovars_RCP85_matrix_s[i,3] <- my_test24$estimate
  } else {
    MWU_biovars_RCP85_matrix[i,3] <- 0
  }   
     
}
```

```
## Mann-Whitney test results for Jan biovars: actual vs. RCP4.5 2050 
## The H0 is rejected as p < 0.05, the HA accepted (medians unequal) 
## 
##  Wilcoxon rank sum test with continuity correction
## 
## data:  x and y
## W = 131800, p-value < 2.2e-16
## alternative hypothesis: true location shift is not equal to 0
## 95 percent confidence interval:
##  -25.68508 -22.91888
## sample estimates:
## difference in location 
##              -24.29676 
## 
## Mann-Whitney test results for Jan biovars: RCP4.5 2050 vs. RCP4.5 2070 
## The H0 is rejected as p < 0.05, the HA accepted (medians unequal) 
## 
##  Wilcoxon rank sum test with continuity correction
## 
## data:  x and y
## W = 409460, p-value = 2.366e-12
## alternative hypothesis: true location shift is not equal to 0
## 95 percent confidence interval:
##  -6.000874 -3.432399
## sample estimates:
## difference in location 
##              -4.719334 
## 
## Mann-Whitney test results for Jan biovars: actual vs. RCP4.5 2070 
## The H0 is rejected as p < 0.05, the HA accepted (medians unequal) 
## 
##  Wilcoxon rank sum test with continuity correction
## 
## data:  x and y
## W = 89347, p-value < 2.2e-16
## alternative hypothesis: true location shift is not equal to 0
## 95 percent confidence interval:
##  -30.40368 -27.63401
## sample estimates:
## difference in location 
##              -29.01579 
## 
## Mann-Whitney test results for Jan biovars: actual vs. RCP8.5 2050 
## The H0 is rejected as p < 0.05, the HA accepted (medians unequal) 
## 
##  Wilcoxon rank sum test with continuity correction
## 
## data:  x and y
## W = 80338, p-value < 2.2e-16
## alternative hypothesis: true location shift is not equal to 0
## 95 percent confidence interval:
##  -31.60940 -28.83988
## sample estimates:
## difference in location 
##              -30.21841 
## 
## Mann-Whitney test results for Jan biovars: RCP8.5 2050 vs. RCP8.5 2070 
## The H0 is rejected as p < 0.05, the HA accepted (medians unequal) 
## 
##  Wilcoxon rank sum test with continuity correction
## 
## data:  x and y
## W = 270860, p-value < 2.2e-16
## alternative hypothesis: true location shift is not equal to 0
## 95 percent confidence interval:
##  -14.03868 -11.47124
## sample estimates:
## difference in location 
##              -12.75955 
## 
## Mann-Whitney test results for Jan biovars: actual vs. RCP8.5 2070 
## The H0 is rejected as p < 0.05, the HA accepted (medians unequal) 
## 
##  Wilcoxon rank sum test with continuity correction
## 
## data:  x and y
## W = 21338, p-value < 2.2e-16
## alternative hypothesis: true location shift is not equal to 0
## 95 percent confidence interval:
##  -44.36527 -41.59398
## sample estimates:
## difference in location 
##              -42.97522 
## 
## Mann-Whitney test results for Feb biovars: actual vs. RCP4.5 2050 
## The H0 is rejected as p < 0.05, the HA accepted (medians unequal) 
## 
##  Wilcoxon rank sum test with continuity correction
## 
## data:  x and y
## W = 4155, p-value < 2.2e-16
## alternative hypothesis: true location shift is not equal to 0
## 95 percent confidence interval:
##  -21.57599 -20.55421
## sample estimates:
## difference in location 
##              -21.06508 
## 
## Mann-Whitney test results for Feb biovars: RCP4.5 2050 vs. RCP4.5 2070 
## The H0 is rejected as p < 0.05, the HA accepted (medians unequal) 
## 
##  Wilcoxon rank sum test with continuity correction
## 
## data:  x and y
## W = 466720, p-value = 0.00995
## alternative hypothesis: true location shift is not equal to 0
## 95 percent confidence interval:
##  -0.9590616 -0.1340881
## sample estimates:
## difference in location 
##             -0.5450447 
## 
## Mann-Whitney test results for Feb biovars: actual vs. RCP4.5 2070 
## The H0 is rejected as p < 0.05, the HA accepted (medians unequal) 
## 
##  Wilcoxon rank sum test with continuity correction
## 
## data:  x and y
## W = 3462, p-value < 2.2e-16
## alternative hypothesis: true location shift is not equal to 0
## 95 percent confidence interval:
##  -22.11052 -21.08578
## sample estimates:
## difference in location 
##              -21.59572 
## 
## Mann-Whitney test results for Feb biovars: actual vs. RCP8.5 2050 
## The H0 is rejected as p < 0.05, the HA accepted (medians unequal) 
## 
##  Wilcoxon rank sum test with continuity correction
## 
## data:  x and y
## W = 2603, p-value < 2.2e-16
## alternative hypothesis: true location shift is not equal to 0
## 95 percent confidence interval:
##  -22.79167 -21.76206
## sample estimates:
## difference in location 
##              -22.27606 
## 
## Mann-Whitney test results for Feb biovars: RCP8.5 2050 vs. RCP8.5 2070 
## The HA is rejected as p >= 0.05, the H0 accepted (medians equal) 
## 
##  Wilcoxon rank sum test with continuity correction
## 
## data:  x and y
## W = 488590, p-value = 0.3768
## alternative hypothesis: true location shift is not equal to 0
## 95 percent confidence interval:
##  -0.6109646  0.2385904
## sample estimates:
## difference in location 
##             -0.1872062 
## 
## Mann-Whitney test results for Feb biovars: actual vs. RCP8.5 2070 
## The H0 is rejected as p < 0.05, the HA accepted (medians unequal) 
## 
##  Wilcoxon rank sum test with continuity correction
## 
## data:  x and y
## W = 2485, p-value < 2.2e-16
## alternative hypothesis: true location shift is not equal to 0
## 95 percent confidence interval:
##  -22.97421 -21.94012
## sample estimates:
## difference in location 
##               -22.4571 
## 
## Mann-Whitney test results for Mar biovars: actual vs. RCP4.5 2050 
## The H0 is rejected as p < 0.05, the HA accepted (medians unequal) 
## 
##  Wilcoxon rank sum test with continuity correction
## 
## data:  x and y
## W = 644740, p-value < 2.2e-16
## alternative hypothesis: true location shift is not equal to 0
## 95 percent confidence interval:
##  0.9369608 1.2598933
## sample estimates:
## difference in location 
##               1.101375 
## 
## Mann-Whitney test results for Mar biovars: RCP4.5 2050 vs. RCP4.5 2070 
## The H0 is rejected as p < 0.05, the HA accepted (medians unequal) 
## 
##  Wilcoxon rank sum test with continuity correction
## 
## data:  x and y
## W = 608080, p-value < 2.2e-16
## alternative hypothesis: true location shift is not equal to 0
## 95 percent confidence interval:
##  0.1523906 0.2420225
## sample estimates:
## difference in location 
##              0.1973121 
## 
## Mann-Whitney test results for Mar biovars: actual vs. RCP4.5 2070 
## The H0 is rejected as p < 0.05, the HA accepted (medians unequal) 
## 
##  Wilcoxon rank sum test with continuity correction
## 
## data:  x and y
## W = 665850, p-value < 2.2e-16
## alternative hypothesis: true location shift is not equal to 0
## 95 percent confidence interval:
##  1.127760 1.445735
## sample estimates:
## difference in location 
##               1.291993 
## 
## Mann-Whitney test results for Mar biovars: actual vs. RCP8.5 2050 
## The H0 is rejected as p < 0.05, the HA accepted (medians unequal) 
## 
##  Wilcoxon rank sum test with continuity correction
## 
## data:  x and y
## W = 643690, p-value < 2.2e-16
## alternative hypothesis: true location shift is not equal to 0
## 95 percent confidence interval:
##  0.9100512 1.2322341
## sample estimates:
## difference in location 
##               1.074351 
## 
## Mann-Whitney test results for Mar biovars: RCP8.5 2050 vs. RCP8.5 2070 
## The H0 is rejected as p < 0.05, the HA accepted (medians unequal) 
## 
##  Wilcoxon rank sum test with continuity correction
## 
## data:  x and y
## W = 700530, p-value < 2.2e-16
## alternative hypothesis: true location shift is not equal to 0
## 95 percent confidence interval:
##  0.4459601 0.5587755
## sample estimates:
## difference in location 
##              0.5027078 
## 
## Mann-Whitney test results for Mar biovars: actual vs. RCP8.5 2070 
## The H0 is rejected as p < 0.05, the HA accepted (medians unequal) 
## 
##  Wilcoxon rank sum test with continuity correction
## 
## data:  x and y
## W = 691120, p-value < 2.2e-16
## alternative hypothesis: true location shift is not equal to 0
## 95 percent confidence interval:
##  1.365018 1.689065
## sample estimates:
## difference in location 
##               1.530811 
## 
## Mann-Whitney test results for Apr biovars: actual vs. RCP4.5 2050 
## The H0 is rejected as p < 0.05, the HA accepted (medians unequal) 
## 
##  Wilcoxon rank sum test with continuity correction
## 
## data:  x and y
## W = 142880, p-value < 2.2e-16
## alternative hypothesis: true location shift is not equal to 0
## 95 percent confidence interval:
##  -85.60713 -76.32925
## sample estimates:
## difference in location 
##              -80.96703 
## 
## Mann-Whitney test results for Apr biovars: RCP4.5 2050 vs. RCP4.5 2070 
## The H0 is rejected as p < 0.05, the HA accepted (medians unequal) 
## 
##  Wilcoxon rank sum test with continuity correction
## 
## data:  x and y
## W = 562980, p-value = 1.075e-06
## alternative hypothesis: true location shift is not equal to 0
## 95 percent confidence interval:
##   7.681207 17.896084
## sample estimates:
## difference in location 
##               12.75689 
## 
## Mann-Whitney test results for Apr biovars: actual vs. RCP4.5 2070 
## The H0 is rejected as p < 0.05, the HA accepted (medians unequal) 
## 
##  Wilcoxon rank sum test with continuity correction
## 
## data:  x and y
## W = 172130, p-value < 2.2e-16
## alternative hypothesis: true location shift is not equal to 0
## 95 percent confidence interval:
##  -72.51936 -63.50198
## sample estimates:
## difference in location 
##              -67.99959 
## 
## Mann-Whitney test results for Apr biovars: actual vs. RCP8.5 2050 
## The H0 is rejected as p < 0.05, the HA accepted (medians unequal) 
## 
##  Wilcoxon rank sum test with continuity correction
## 
## data:  x and y
## W = 115350, p-value < 2.2e-16
## alternative hypothesis: true location shift is not equal to 0
## 95 percent confidence interval:
##  -97.23555 -87.61630
## sample estimates:
## difference in location 
##              -92.48163 
## 
## Mann-Whitney test results for Apr biovars: RCP8.5 2050 vs. RCP8.5 2070 
## The H0 is rejected as p < 0.05, the HA accepted (medians unequal) 
## 
##  Wilcoxon rank sum test with continuity correction
## 
## data:  x and y
## W = 805800, p-value < 2.2e-16
## alternative hypothesis: true location shift is not equal to 0
## 95 percent confidence interval:
##  61.85529 71.57215
## sample estimates:
## difference in location 
##               66.75842 
## 
## Mann-Whitney test results for Apr biovars: actual vs. RCP8.5 2070 
## The H0 is rejected as p < 0.05, the HA accepted (medians unequal) 
## 
##  Wilcoxon rank sum test with continuity correction
## 
## data:  x and y
## W = 338460, p-value < 2.2e-16
## alternative hypothesis: true location shift is not equal to 0
## 95 percent confidence interval:
##  -28.28194 -20.92811
## sample estimates:
## difference in location 
##              -24.58276 
## 
## Mann-Whitney test results for May biovars: actual vs. RCP4.5 2050 
## The H0 is rejected as p < 0.05, the HA accepted (medians unequal) 
## 
##  Wilcoxon rank sum test with continuity correction
## 
## data:  x and y
## W = 18491, p-value < 2.2e-16
## alternative hypothesis: true location shift is not equal to 0
## 95 percent confidence interval:
##  -39.73614 -37.43403
## sample estimates:
## difference in location 
##              -38.58569 
## 
## Mann-Whitney test results for May biovars: RCP4.5 2050 vs. RCP4.5 2070 
## The H0 is rejected as p < 0.05, the HA accepted (medians unequal) 
## 
##  Wilcoxon rank sum test with continuity correction
## 
## data:  x and y
## W = 366660, p-value < 2.2e-16
## alternative hypothesis: true location shift is not equal to 0
## 95 percent confidence interval:
##  -7.064900 -4.890134
## sample estimates:
## difference in location 
##              -5.993927 
## 
## Mann-Whitney test results for May biovars: actual vs. RCP4.5 2070 
## The H0 is rejected as p < 0.05, the HA accepted (medians unequal) 
## 
##  Wilcoxon rank sum test with continuity correction
## 
## data:  x and y
## W = 8123, p-value < 2.2e-16
## alternative hypothesis: true location shift is not equal to 0
## 95 percent confidence interval:
##  -45.71323 -43.40931
## sample estimates:
## difference in location 
##               -44.5638 
## 
## Mann-Whitney test results for May biovars: actual vs. RCP8.5 2050 
## The H0 is rejected as p < 0.05, the HA accepted (medians unequal) 
## 
##  Wilcoxon rank sum test with continuity correction
## 
## data:  x and y
## W = 6768, p-value < 2.2e-16
## alternative hypothesis: true location shift is not equal to 0
## 95 percent confidence interval:
##  -46.90541 -44.60782
## sample estimates:
## difference in location 
##              -45.76211 
## 
## Mann-Whitney test results for May biovars: RCP8.5 2050 vs. RCP8.5 2070 
## The H0 is rejected as p < 0.05, the HA accepted (medians unequal) 
## 
##  Wilcoxon rank sum test with continuity correction
## 
## data:  x and y
## W = 202830, p-value < 2.2e-16
## alternative hypothesis: true location shift is not equal to 0
## 95 percent confidence interval:
##  -15.83536 -13.65612
## sample estimates:
## difference in location 
##              -14.75508 
## 
## Mann-Whitney test results for May biovars: actual vs. RCP8.5 2070 
## The H0 is rejected as p < 0.05, the HA accepted (medians unequal) 
## 
##  Wilcoxon rank sum test with continuity correction
## 
## data:  x and y
## W = 741, p-value < 2.2e-16
## alternative hypothesis: true location shift is not equal to 0
## 95 percent confidence interval:
##  -61.65597 -59.35761
## sample estimates:
## difference in location 
##              -60.51341 
## 
## Mann-Whitney test results for Jun biovars: actual vs. RCP4.5 2050 
## The H0 is rejected as p < 0.05, the HA accepted (medians unequal) 
## 
##  Wilcoxon rank sum test with continuity correction
## 
## data:  x and y
## W = 290940, p-value < 2.2e-16
## alternative hypothesis: true location shift is not equal to 0
## 95 percent confidence interval:
##  -16.15061 -12.84025
## sample estimates:
## difference in location 
##              -14.49005 
## 
## Mann-Whitney test results for Jun biovars: RCP4.5 2050 vs. RCP4.5 2070 
## The H0 is rejected as p < 0.05, the HA accepted (medians unequal) 
## 
##  Wilcoxon rank sum test with continuity correction
## 
## data:  x and y
## W = 418540, p-value = 2.826e-10
## alternative hypothesis: true location shift is not equal to 0
## 95 percent confidence interval:
##  -6.578099 -3.512983
## sample estimates:
## difference in location 
##              -5.020359 
## 
## Mann-Whitney test results for Jun biovars: actual vs. RCP4.5 2070 
## The H0 is rejected as p < 0.05, the HA accepted (medians unequal) 
## 
##  Wilcoxon rank sum test with continuity correction
## 
## data:  x and y
## W = 229080, p-value < 2.2e-16
## alternative hypothesis: true location shift is not equal to 0
## 95 percent confidence interval:
##  -21.20386 -17.88574
## sample estimates:
## difference in location 
##              -19.54115 
## 
## Mann-Whitney test results for Jun biovars: actual vs. RCP8.5 2050 
## The H0 is rejected as p < 0.05, the HA accepted (medians unequal) 
## 
##  Wilcoxon rank sum test with continuity correction
## 
## data:  x and y
## W = 220500, p-value < 2.2e-16
## alternative hypothesis: true location shift is not equal to 0
## 95 percent confidence interval:
##  -21.88377 -18.57682
## sample estimates:
## difference in location 
##              -20.22126 
## 
## Mann-Whitney test results for Jun biovars: RCP8.5 2050 vs. RCP8.5 2070 
## The H0 is rejected as p < 0.05, the HA accepted (medians unequal) 
## 
##  Wilcoxon rank sum test with continuity correction
## 
## data:  x and y
## W = 286260, p-value < 2.2e-16
## alternative hypothesis: true location shift is not equal to 0
## 95 percent confidence interval:
##  -15.45795 -12.41092
## sample estimates:
## difference in location 
##              -13.95073 
## 
## Mann-Whitney test results for Jun biovars: actual vs. RCP8.5 2070 
## The H0 is rejected as p < 0.05, the HA accepted (medians unequal) 
## 
##  Wilcoxon rank sum test with continuity correction
## 
## data:  x and y
## W = 88750, p-value < 2.2e-16
## alternative hypothesis: true location shift is not equal to 0
## 95 percent confidence interval:
##  -35.81732 -32.51095
## sample estimates:
## difference in location 
##               -34.1609 
## 
## Mann-Whitney test results for Jul biovars: actual vs. RCP4.5 2050 
## The H0 is rejected as p < 0.05, the HA accepted (medians unequal) 
## 
##  Wilcoxon rank sum test with continuity correction
## 
## data:  x and y
## W = 2133, p-value < 2.2e-16
## alternative hypothesis: true location shift is not equal to 0
## 95 percent confidence interval:
##  -24.10228 -22.93551
## sample estimates:
## difference in location 
##              -23.51537 
## 
## Mann-Whitney test results for Jul biovars: RCP4.5 2050 vs. RCP4.5 2070 
## The H0 is rejected as p < 0.05, the HA accepted (medians unequal) 
## 
##  Wilcoxon rank sum test with continuity correction
## 
## data:  x and y
## W = 447520, p-value = 4.815e-05
## alternative hypothesis: true location shift is not equal to 0
## 95 percent confidence interval:
##  -1.3642667 -0.4820413
## sample estimates:
## difference in location 
##               -0.93111 
## 
## Mann-Whitney test results for Jul biovars: actual vs. RCP4.5 2070 
## The H0 is rejected as p < 0.05, the HA accepted (medians unequal) 
## 
##  Wilcoxon rank sum test with continuity correction
## 
## data:  x and y
## W = 1647, p-value < 2.2e-16
## alternative hypothesis: true location shift is not equal to 0
## 95 percent confidence interval:
##  -25.00736 -23.84201
## sample estimates:
## difference in location 
##              -24.42511 
## 
## Mann-Whitney test results for Jul biovars: actual vs. RCP8.5 2050 
## The H0 is rejected as p < 0.05, the HA accepted (medians unequal) 
## 
##  Wilcoxon rank sum test with continuity correction
## 
## data:  x and y
## W = 1351, p-value < 2.2e-16
## alternative hypothesis: true location shift is not equal to 0
## 95 percent confidence interval:
##  -25.49418 -24.31673
## sample estimates:
## difference in location 
##              -24.90351 
## 
## Mann-Whitney test results for Jul biovars: RCP8.5 2050 vs. RCP8.5 2070 
## The H0 is rejected as p < 0.05, the HA accepted (medians unequal) 
## 
##  Wilcoxon rank sum test with continuity correction
## 
## data:  x and y
## W = 453460, p-value = 0.0003135
## alternative hypothesis: true location shift is not equal to 0
## 95 percent confidence interval:
##  -1.2695545 -0.3802507
## sample estimates:
## difference in location 
##             -0.8290996 
## 
## Mann-Whitney test results for Jul biovars: actual vs. RCP8.5 2070 
## The H0 is rejected as p < 0.05, the HA accepted (medians unequal) 
## 
##  Wilcoxon rank sum test with continuity correction
## 
## data:  x and y
## W = 1077, p-value < 2.2e-16
## alternative hypothesis: true location shift is not equal to 0
## 95 percent confidence interval:
##  -26.34974 -25.16363
## sample estimates:
## difference in location 
##              -25.75422 
## 
## Mann-Whitney test results for Aug biovars: actual vs. RCP4.5 2050 
## The H0 is rejected as p < 0.05, the HA accepted (medians unequal) 
## 
##  Wilcoxon rank sum test with continuity correction
## 
## data:  x and y
## W = 143530, p-value < 2.2e-16
## alternative hypothesis: true location shift is not equal to 0
## 95 percent confidence interval:
##  -24.19629 -21.46166
## sample estimates:
## difference in location 
##              -22.82045 
## 
## Mann-Whitney test results for Aug biovars: RCP4.5 2050 vs. RCP4.5 2070 
## The H0 is rejected as p < 0.05, the HA accepted (medians unequal) 
## 
##  Wilcoxon rank sum test with continuity correction
## 
## data:  x and y
## W = 408810, p-value = 1.642e-12
## alternative hypothesis: true location shift is not equal to 0
## 95 percent confidence interval:
##  -5.986316 -3.424124
## sample estimates:
## difference in location 
##              -4.726727 
## 
## Mann-Whitney test results for Aug biovars: actual vs. RCP4.5 2070 
## The H0 is rejected as p < 0.05, the HA accepted (medians unequal) 
## 
##  Wilcoxon rank sum test with continuity correction
## 
## data:  x and y
## W = 99910, p-value < 2.2e-16
## alternative hypothesis: true location shift is not equal to 0
## 95 percent confidence interval:
##  -28.91399 -26.17174
## sample estimates:
## difference in location 
##              -27.53189 
## 
## Mann-Whitney test results for Aug biovars: actual vs. RCP8.5 2050 
## The H0 is rejected as p < 0.05, the HA accepted (medians unequal) 
## 
##  Wilcoxon rank sum test with continuity correction
## 
## data:  x and y
## W = 89228, p-value < 2.2e-16
## alternative hypothesis: true location shift is not equal to 0
## 95 percent confidence interval:
##  -29.95815 -27.23128
## sample estimates:
## difference in location 
##              -28.59004 
## 
## Mann-Whitney test results for Aug biovars: RCP8.5 2050 vs. RCP8.5 2070 
## The H0 is rejected as p < 0.05, the HA accepted (medians unequal) 
## 
##  Wilcoxon rank sum test with continuity correction
## 
## data:  x and y
## W = 258010, p-value < 2.2e-16
## alternative hypothesis: true location shift is not equal to 0
## 95 percent confidence interval:
##  -14.70062 -12.15194
## sample estimates:
## difference in location 
##              -13.43782 
## 
## Mann-Whitney test results for Aug biovars: actual vs. RCP8.5 2070 
## The H0 is rejected as p < 0.05, the HA accepted (medians unequal) 
## 
##  Wilcoxon rank sum test with continuity correction
## 
## data:  x and y
## W = 23865, p-value < 2.2e-16
## alternative hypothesis: true location shift is not equal to 0
## 95 percent confidence interval:
##  -43.40783 -40.65048
## sample estimates:
## difference in location 
##              -42.02242 
## 
## Mann-Whitney test results for Sep biovars: actual vs. RCP4.5 2050 
## The H0 is rejected as p < 0.05, the HA accepted (medians unequal) 
## 
##  Wilcoxon rank sum test with continuity correction
## 
## data:  x and y
## W = 121080, p-value < 2.2e-16
## alternative hypothesis: true location shift is not equal to 0
## 95 percent confidence interval:
##  -26.76717 -24.03173
## sample estimates:
## difference in location 
##              -25.40392 
## 
## Mann-Whitney test results for Sep biovars: RCP4.5 2050 vs. RCP4.5 2070 
## The H0 is rejected as p < 0.05, the HA accepted (medians unequal) 
## 
##  Wilcoxon rank sum test with continuity correction
## 
## data:  x and y
## W = 419850, p-value = 5.405e-10
## alternative hypothesis: true location shift is not equal to 0
## 95 percent confidence interval:
##  -5.548039 -2.924998
## sample estimates:
## difference in location 
##              -4.239073 
## 
## Mann-Whitney test results for Sep biovars: actual vs. RCP4.5 2070 
## The H0 is rejected as p < 0.05, the HA accepted (medians unequal) 
## 
##  Wilcoxon rank sum test with continuity correction
## 
## data:  x and y
## W = 86667, p-value < 2.2e-16
## alternative hypothesis: true location shift is not equal to 0
## 95 percent confidence interval:
##  -31.02303 -28.26915
## sample estimates:
## difference in location 
##              -29.64955 
## 
## Mann-Whitney test results for Sep biovars: actual vs. RCP8.5 2050 
## The H0 is rejected as p < 0.05, the HA accepted (medians unequal) 
## 
##  Wilcoxon rank sum test with continuity correction
## 
## data:  x and y
## W = 72728, p-value < 2.2e-16
## alternative hypothesis: true location shift is not equal to 0
## 95 percent confidence interval:
##  -32.73461 -29.98442
## sample estimates:
## difference in location 
##              -31.36568 
## 
## Mann-Whitney test results for Sep biovars: RCP8.5 2050 vs. RCP8.5 2070 
## The H0 is rejected as p < 0.05, the HA accepted (medians unequal) 
## 
##  Wilcoxon rank sum test with continuity correction
## 
## data:  x and y
## W = 277030, p-value < 2.2e-16
## alternative hypothesis: true location shift is not equal to 0
## 95 percent confidence interval:
##  -13.88823 -11.25556
## sample estimates:
## difference in location 
##              -12.56842 
## 
## Mann-Whitney test results for Sep biovars: actual vs. RCP8.5 2070 
## The H0 is rejected as p < 0.05, the HA accepted (medians unequal) 
## 
##  Wilcoxon rank sum test with continuity correction
## 
## data:  x and y
## W = 19587, p-value < 2.2e-16
## alternative hypothesis: true location shift is not equal to 0
## 95 percent confidence interval:
##  -45.30167 -42.55626
## sample estimates:
## difference in location 
##              -43.92882 
## 
## Mann-Whitney test results for Oct biovars: actual vs. RCP4.5 2050 
## The H0 is rejected as p < 0.05, the HA accepted (medians unequal) 
## 
##  Wilcoxon rank sum test with continuity correction
## 
## data:  x and y
## W = 121530, p-value < 2.2e-16
## alternative hypothesis: true location shift is not equal to 0
## 95 percent confidence interval:
##  -26.55296 -23.79950
## sample estimates:
## difference in location 
##              -25.17219 
## 
## Mann-Whitney test results for Oct biovars: RCP4.5 2050 vs. RCP4.5 2070 
## The H0 is rejected as p < 0.05, the HA accepted (medians unequal) 
## 
##  Wilcoxon rank sum test with continuity correction
## 
## data:  x and y
## W = 406800, p-value = 5.293e-13
## alternative hypothesis: true location shift is not equal to 0
## 95 percent confidence interval:
##  -6.148804 -3.570572
## sample estimates:
## difference in location 
##              -4.853944 
## 
## Mann-Whitney test results for Oct biovars: actual vs. RCP4.5 2070 
## The H0 is rejected as p < 0.05, the HA accepted (medians unequal) 
## 
##  Wilcoxon rank sum test with continuity correction
## 
## data:  x and y
## W = 80965, p-value < 2.2e-16
## alternative hypothesis: true location shift is not equal to 0
## 95 percent confidence interval:
##  -31.41729 -28.65274
## sample estimates:
## difference in location 
##              -30.03171 
## 
## Mann-Whitney test results for Oct biovars: actual vs. RCP8.5 2050 
## The H0 is rejected as p < 0.05, the HA accepted (medians unequal) 
## 
##  Wilcoxon rank sum test with continuity correction
## 
## data:  x and y
## W = 71558, p-value < 2.2e-16
## alternative hypothesis: true location shift is not equal to 0
## 95 percent confidence interval:
##  -32.68636 -29.93132
## sample estimates:
## difference in location 
##              -31.30104 
## 
## Mann-Whitney test results for Oct biovars: RCP8.5 2050 vs. RCP8.5 2070 
## The H0 is rejected as p < 0.05, the HA accepted (medians unequal) 
## 
##  Wilcoxon rank sum test with continuity correction
## 
## data:  x and y
## W = 272990, p-value < 2.2e-16
## alternative hypothesis: true location shift is not equal to 0
## 95 percent confidence interval:
##  -13.98506 -11.38488
## sample estimates:
## difference in location 
##              -12.68165 
## 
## Mann-Whitney test results for Oct biovars: actual vs. RCP8.5 2070 
## The H0 is rejected as p < 0.05, the HA accepted (medians unequal) 
## 
##  Wilcoxon rank sum test with continuity correction
## 
## data:  x and y
## W = 19023, p-value < 2.2e-16
## alternative hypothesis: true location shift is not equal to 0
## 95 percent confidence interval:
##  -45.37873 -42.60586
## sample estimates:
## difference in location 
##              -43.98333 
## 
## Mann-Whitney test results for Nov biovars: actual vs. RCP4.5 2050 
## The H0 is rejected as p < 0.05, the HA accepted (medians unequal) 
## 
##  Wilcoxon rank sum test with continuity correction
## 
## data:  x and y
## W = 154390, p-value < 2.2e-16
## alternative hypothesis: true location shift is not equal to 0
## 95 percent confidence interval:
##  -24.55361 -21.67731
## sample estimates:
## difference in location 
##              -23.12207 
## 
## Mann-Whitney test results for Nov biovars: RCP4.5 2050 vs. RCP4.5 2070 
## The H0 is rejected as p < 0.05, the HA accepted (medians unequal) 
## 
##  Wilcoxon rank sum test with continuity correction
## 
## data:  x and y
## W = 406580, p-value = 4.667e-13
## alternative hypothesis: true location shift is not equal to 0
## 95 percent confidence interval:
##  -6.497265 -3.780453
## sample estimates:
## difference in location 
##              -5.148205 
## 
## Mann-Whitney test results for Nov biovars: actual vs. RCP4.5 2070 
## The H0 is rejected as p < 0.05, the HA accepted (medians unequal) 
## 
##  Wilcoxon rank sum test with continuity correction
## 
## data:  x and y
## W = 104520, p-value < 2.2e-16
## alternative hypothesis: true location shift is not equal to 0
## 95 percent confidence interval:
##  -29.69249 -26.81557
## sample estimates:
## difference in location 
##              -28.25764 
## 
## Mann-Whitney test results for Nov biovars: actual vs. RCP8.5 2050 
## The H0 is rejected as p < 0.05, the HA accepted (medians unequal) 
## 
##  Wilcoxon rank sum test with continuity correction
## 
## data:  x and y
## W = 98524, p-value < 2.2e-16
## alternative hypothesis: true location shift is not equal to 0
## 95 percent confidence interval:
##  -30.38403 -27.51192
## sample estimates:
## difference in location 
##               -28.9541 
## 
## Mann-Whitney test results for Nov biovars: RCP8.5 2050 vs. RCP8.5 2070 
## The H0 is rejected as p < 0.05, the HA accepted (medians unequal) 
## 
##  Wilcoxon rank sum test with continuity correction
## 
## data:  x and y
## W = 252550, p-value < 2.2e-16
## alternative hypothesis: true location shift is not equal to 0
## 95 percent confidence interval:
##  -15.95834 -13.25531
## sample estimates:
## difference in location 
##              -14.60613 
## 
## Mann-Whitney test results for Nov biovars: actual vs. RCP8.5 2070 
## The H0 is rejected as p < 0.05, the HA accepted (medians unequal) 
## 
##  Wilcoxon rank sum test with continuity correction
## 
## data:  x and y
## W = 24083, p-value < 2.2e-16
## alternative hypothesis: true location shift is not equal to 0
## 95 percent confidence interval:
##  -44.97962 -42.12257
## sample estimates:
## difference in location 
##              -43.55361 
## 
## Mann-Whitney test results for Dic biovars: actual vs. RCP4.5 2050 
## The H0 is rejected as p < 0.05, the HA accepted (medians unequal) 
## 
##  Wilcoxon rank sum test with continuity correction
## 
## data:  x and y
## W = 261420, p-value < 2.2e-16
## alternative hypothesis: true location shift is not equal to 0
## 95 percent confidence interval:
##  -756.5976 -607.4252
## sample estimates:
## difference in location 
##              -681.9129 
## 
## Mann-Whitney test results for Dic biovars: RCP4.5 2050 vs. RCP4.5 2070 
## The H0 is rejected as p < 0.05, the HA accepted (medians unequal) 
## 
##  Wilcoxon rank sum test with continuity correction
## 
## data:  x and y
## W = 545670, p-value = 0.0004051
## alternative hypothesis: true location shift is not equal to 0
## 95 percent confidence interval:
##   58.44318 195.93729
## sample estimates:
## difference in location 
##               127.2624 
## 
## Mann-Whitney test results for Dic biovars: actual vs. RCP4.5 2070 
## The H0 is rejected as p < 0.05, the HA accepted (medians unequal) 
## 
##  Wilcoxon rank sum test with continuity correction
## 
## data:  x and y
## W = 298670, p-value < 2.2e-16
## alternative hypothesis: true location shift is not equal to 0
## 95 percent confidence interval:
##  -629.1275 -484.6494
## sample estimates:
## difference in location 
##              -556.9354 
## 
## Mann-Whitney test results for Dic biovars: actual vs. RCP8.5 2050 
## The H0 is rejected as p < 0.05, the HA accepted (medians unequal) 
## 
##  Wilcoxon rank sum test with continuity correction
## 
## data:  x and y
## W = 312060, p-value < 2.2e-16
## alternative hypothesis: true location shift is not equal to 0
## 95 percent confidence interval:
##  -581.4917 -439.4580
## sample estimates:
## difference in location 
##              -510.4962 
## 
## Mann-Whitney test results for Dic biovars: RCP8.5 2050 vs. RCP8.5 2070 
## The H0 is rejected as p < 0.05, the HA accepted (medians unequal) 
## 
##  Wilcoxon rank sum test with continuity correction
## 
## data:  x and y
## W = 466140, p-value = 0.008739
## alternative hypothesis: true location shift is not equal to 0
## 95 percent confidence interval:
##  -163.94933  -24.09229
## sample estimates:
## difference in location 
##              -94.04197 
## 
## Mann-Whitney test results for Dic biovars: actual vs. RCP8.5 2070 
## The H0 is rejected as p < 0.05, the HA accepted (medians unequal) 
## 
##  Wilcoxon rank sum test with continuity correction
## 
## data:  x and y
## W = 290010, p-value < 2.2e-16
## alternative hypothesis: true location shift is not equal to 0
## 95 percent confidence interval:
##  -675.6953 -524.9788
## sample estimates:
## difference in location 
##              -600.1776 
## 
## Mann-Whitney test results for NA biovars: actual vs. RCP4.5 2050 
## The H0 is rejected as p < 0.05, the HA accepted (medians unequal) 
## 
##  Wilcoxon rank sum test with continuity correction
## 
## data:  x and y
## W = 275460, p-value < 2.2e-16
## alternative hypothesis: true location shift is not equal to 0
## 95 percent confidence interval:
##  -92.54357 -73.33440
## sample estimates:
## difference in location 
##              -82.77215 
## 
## Mann-Whitney test results for NA biovars: RCP4.5 2050 vs. RCP4.5 2070 
## The HA is rejected as p >= 0.05, the H0 accepted (medians equal) 
## 
##  Wilcoxon rank sum test with continuity correction
## 
## data:  x and y
## W = 496610, p-value = 0.7932
## alternative hypothesis: true location shift is not equal to 0
## 95 percent confidence interval:
##  -11.229256   8.456531
## sample estimates:
## difference in location 
##               -1.26536 
## 
## Mann-Whitney test results for NA biovars: actual vs. RCP4.5 2070 
## The H0 is rejected as p < 0.05, the HA accepted (medians unequal) 
## 
##  Wilcoxon rank sum test with continuity correction
## 
## data:  x and y
## W = 272810, p-value < 2.2e-16
## alternative hypothesis: true location shift is not equal to 0
## 95 percent confidence interval:
##  -93.45634 -74.33431
## sample estimates:
## difference in location 
##              -83.87102 
## 
## Mann-Whitney test results for NA biovars: actual vs. RCP8.5 2050 
## The H0 is rejected as p < 0.05, the HA accepted (medians unequal) 
## 
##  Wilcoxon rank sum test with continuity correction
## 
## data:  x and y
## W = 340860, p-value < 2.2e-16
## alternative hypothesis: true location shift is not equal to 0
## 95 percent confidence interval:
##  -62.06203 -44.64321
## sample estimates:
## difference in location 
##              -53.30454 
## 
## Mann-Whitney test results for NA biovars: RCP8.5 2050 vs. RCP8.5 2070 
## The H0 is rejected as p < 0.05, the HA accepted (medians unequal) 
## 
##  Wilcoxon rank sum test with continuity correction
## 
## data:  x and y
## W = 351070, p-value < 2.2e-16
## alternative hypothesis: true location shift is not equal to 0
## 95 percent confidence interval:
##  -75.65746 -54.29405
## sample estimates:
## difference in location 
##              -64.98185 
## 
## Mann-Whitney test results for NA biovars: actual vs. RCP8.5 2070 
## The H0 is rejected as p < 0.05, the HA accepted (medians unequal) 
## 
##  Wilcoxon rank sum test with continuity correction
## 
## data:  x and y
## W = 210410, p-value < 2.2e-16
## alternative hypothesis: true location shift is not equal to 0
## 95 percent confidence interval:
##  -129.0001 -107.9992
## sample estimates:
## difference in location 
##              -118.3921 
## 
## Mann-Whitney test results for NA biovars: actual vs. RCP4.5 2050 
## The H0 is rejected as p < 0.05, the HA accepted (medians unequal) 
## 
##  Wilcoxon rank sum test with continuity correction
## 
## data:  x and y
## W = 194270, p-value < 2.2e-16
## alternative hypothesis: true location shift is not equal to 0
## 95 percent confidence interval:
##  -57.66047 -48.15162
## sample estimates:
## difference in location 
##              -52.83223 
## 
## Mann-Whitney test results for NA biovars: RCP4.5 2050 vs. RCP4.5 2070 
## The H0 is rejected as p < 0.05, the HA accepted (medians unequal) 
## 
##  Wilcoxon rank sum test with continuity correction
## 
## data:  x and y
## W = 541550, p-value = 0.001294
## alternative hypothesis: true location shift is not equal to 0
## 95 percent confidence interval:
##   3.034419 12.309174
## sample estimates:
## difference in location 
##               7.663441 
## 
## Mann-Whitney test results for NA biovars: actual vs. RCP4.5 2070 
## The H0 is rejected as p < 0.05, the HA accepted (medians unequal) 
## 
##  Wilcoxon rank sum test with continuity correction
## 
## data:  x and y
## W = 221330, p-value < 2.2e-16
## alternative hypothesis: true location shift is not equal to 0
## 95 percent confidence interval:
##  -50.85781 -41.96467
## sample estimates:
## difference in location 
##               -46.3919 
## 
## Mann-Whitney test results for NA biovars: actual vs. RCP8.5 2050 
## The H0 is rejected as p < 0.05, the HA accepted (medians unequal) 
## 
##  Wilcoxon rank sum test with continuity correction
## 
## data:  x and y
## W = 226690, p-value < 2.2e-16
## alternative hypothesis: true location shift is not equal to 0
## 95 percent confidence interval:
##  -50.05381 -41.15616
## sample estimates:
## difference in location 
##              -45.59133 
## 
## Mann-Whitney test results for NA biovars: RCP8.5 2050 vs. RCP8.5 2070 
## The H0 is rejected as p < 0.05, the HA accepted (medians unequal) 
## 
##  Wilcoxon rank sum test with continuity correction
## 
## data:  x and y
## W = 631280, p-value < 2.2e-16
## alternative hypothesis: true location shift is not equal to 0
## 95 percent confidence interval:
##  17.92515 26.70316
## sample estimates:
## difference in location 
##               22.30046 
## 
## Mann-Whitney test results for NA biovars: actual vs. RCP8.5 2070 
## The H0 is rejected as p < 0.05, the HA accepted (medians unequal) 
## 
##  Wilcoxon rank sum test with continuity correction
## 
## data:  x and y
## W = 319160, p-value < 2.2e-16
## alternative hypothesis: true location shift is not equal to 0
## 95 percent confidence interval:
##  -25.71765 -19.28678
## sample estimates:
## difference in location 
##              -22.44217 
## 
## Mann-Whitney test results for NA biovars: actual vs. RCP4.5 2050 
## The H0 is rejected as p < 0.05, the HA accepted (medians unequal) 
## 
##  Wilcoxon rank sum test with continuity correction
## 
## data:  x and y
## W = 529190, p-value = 0.02379
## alternative hypothesis: true location shift is not equal to 0
## 95 percent confidence interval:
##  0.0431220 0.6226461
## sample estimates:
## difference in location 
##              0.3294316 
## 
## Mann-Whitney test results for NA biovars: RCP4.5 2050 vs. RCP4.5 2070 
## The H0 is rejected as p < 0.05, the HA accepted (medians unequal) 
## 
##  Wilcoxon rank sum test with continuity correction
## 
## data:  x and y
## W = 358400, p-value < 2.2e-16
## alternative hypothesis: true location shift is not equal to 0
## 95 percent confidence interval:
##  -2.328546 -1.664417
## sample estimates:
## difference in location 
##              -2.000695 
## 
## Mann-Whitney test results for NA biovars: actual vs. RCP4.5 2070 
## The H0 is rejected as p < 0.05, the HA accepted (medians unequal) 
## 
##  Wilcoxon rank sum test with continuity correction
## 
## data:  x and y
## W = 372880, p-value < 2.2e-16
## alternative hypothesis: true location shift is not equal to 0
## 95 percent confidence interval:
##  -1.956442 -1.337928
## sample estimates:
## difference in location 
##              -1.650917 
## 
## Mann-Whitney test results for NA biovars: actual vs. RCP8.5 2050 
## The H0 is rejected as p < 0.05, the HA accepted (medians unequal) 
## 
##  Wilcoxon rank sum test with continuity correction
## 
## data:  x and y
## W = 600560, p-value = 6.842e-15
## alternative hypothesis: true location shift is not equal to 0
## 95 percent confidence interval:
##  0.8844389 1.4811355
## sample estimates:
## difference in location 
##               1.180858 
## 
## Mann-Whitney test results for NA biovars: RCP8.5 2050 vs. RCP8.5 2070 
## The H0 is rejected as p < 0.05, the HA accepted (medians unequal) 
## 
##  Wilcoxon rank sum test with continuity correction
## 
## data:  x and y
## W = 113560, p-value < 2.2e-16
## alternative hypothesis: true location shift is not equal to 0
## 95 percent confidence interval:
##  -7.462828 -6.756958
## sample estimates:
## difference in location 
##              -7.111779 
## 
## Mann-Whitney test results for NA biovars: actual vs. RCP8.5 2070 
## The H0 is rejected as p < 0.05, the HA accepted (medians unequal) 
## 
##  Wilcoxon rank sum test with continuity correction
## 
## data:  x and y
## W = 140800, p-value < 2.2e-16
## alternative hypothesis: true location shift is not equal to 0
## 95 percent confidence interval:
##  -6.240010 -5.614965
## sample estimates:
## difference in location 
##              -5.931531 
## 
## Mann-Whitney test results for NA biovars: actual vs. RCP4.5 2050 
## The H0 is rejected as p < 0.05, the HA accepted (medians unequal) 
## 
##  Wilcoxon rank sum test with continuity correction
## 
## data:  x and y
## W = 266130, p-value < 2.2e-16
## alternative hypothesis: true location shift is not equal to 0
## 95 percent confidence interval:
##  -263.9710 -211.6128
## sample estimates:
## difference in location 
##              -237.6319 
## 
## Mann-Whitney test results for NA biovars: RCP4.5 2050 vs. RCP4.5 2070 
## The HA is rejected as p >= 0.05, the H0 accepted (medians equal) 
## 
##  Wilcoxon rank sum test with continuity correction
## 
## data:  x and y
## W = 510620, p-value = 0.4107
## alternative hypothesis: true location shift is not equal to 0
## 95 percent confidence interval:
##  -15.93983  37.57158
## sample estimates:
## difference in location 
##               10.78233 
## 
## Mann-Whitney test results for NA biovars: actual vs. RCP4.5 2070 
## The H0 is rejected as p < 0.05, the HA accepted (medians unequal) 
## 
##  Wilcoxon rank sum test with continuity correction
## 
## data:  x and y
## W = 274730, p-value < 2.2e-16
## alternative hypothesis: true location shift is not equal to 0
## 95 percent confidence interval:
##  -253.2739 -201.3122
## sample estimates:
## difference in location 
##              -227.2514 
## 
## Mann-Whitney test results for NA biovars: actual vs. RCP8.5 2050 
## The H0 is rejected as p < 0.05, the HA accepted (medians unequal) 
## 
##  Wilcoxon rank sum test with continuity correction
## 
## data:  x and y
## W = 329350, p-value < 2.2e-16
## alternative hypothesis: true location shift is not equal to 0
## 95 percent confidence interval:
##  -185.1366 -136.1145
## sample estimates:
## difference in location 
##              -160.4242 
## 
## Mann-Whitney test results for NA biovars: RCP8.5 2050 vs. RCP8.5 2070 
## The H0 is rejected as p < 0.05, the HA accepted (medians unequal) 
## 
##  Wilcoxon rank sum test with continuity correction
## 
## data:  x and y
## W = 394850, p-value = 3.867e-16
## alternative hypothesis: true location shift is not equal to 0
## 95 percent confidence interval:
##  -147.86311  -91.58593
## sample estimates:
## difference in location 
##              -119.5965 
## 
## Mann-Whitney test results for NA biovars: actual vs. RCP8.5 2070 
## The H0 is rejected as p < 0.05, the HA accepted (medians unequal) 
## 
##  Wilcoxon rank sum test with continuity correction
## 
## data:  x and y
## W = 243250, p-value < 2.2e-16
## alternative hypothesis: true location shift is not equal to 0
## 95 percent confidence interval:
##  -309.0779 -252.4757
## sample estimates:
## difference in location 
##              -280.5629 
## 
## Mann-Whitney test results for NA biovars: actual vs. RCP4.5 2050 
## The H0 is rejected as p < 0.05, the HA accepted (medians unequal) 
## 
##  Wilcoxon rank sum test with continuity correction
## 
## data:  x and y
## W = 217220, p-value < 2.2e-16
## alternative hypothesis: true location shift is not equal to 0
## 95 percent confidence interval:
##  -161.2960 -131.8932
## sample estimates:
## difference in location 
##              -146.2824 
## 
## Mann-Whitney test results for NA biovars: RCP4.5 2050 vs. RCP4.5 2070 
## The H0 is rejected as p < 0.05, the HA accepted (medians unequal) 
## 
##  Wilcoxon rank sum test with continuity correction
## 
## data:  x and y
## W = 548710, p-value = 0.000162
## alternative hypothesis: true location shift is not equal to 0
## 95 percent confidence interval:
##  13.37478 41.42604
## sample estimates:
## difference in location 
##                27.3378 
## 
## Mann-Whitney test results for NA biovars: actual vs. RCP4.5 2070 
## The H0 is rejected as p < 0.05, the HA accepted (medians unequal) 
## 
##  Wilcoxon rank sum test with continuity correction
## 
## data:  x and y
## W = 260170, p-value < 2.2e-16
## alternative hypothesis: true location shift is not equal to 0
## 95 percent confidence interval:
##  -134.9756 -106.9439
## sample estimates:
## difference in location 
##               -120.723 
## 
## Mann-Whitney test results for NA biovars: actual vs. RCP8.5 2050 
## The H0 is rejected as p < 0.05, the HA accepted (medians unequal) 
## 
##  Wilcoxon rank sum test with continuity correction
## 
## data:  x and y
## W = 258190, p-value < 2.2e-16
## alternative hypothesis: true location shift is not equal to 0
## 95 percent confidence interval:
##  -135.0177 -107.1219
## sample estimates:
## difference in location 
##              -120.8046 
## 
## Mann-Whitney test results for NA biovars: RCP8.5 2050 vs. RCP8.5 2070 
## The H0 is rejected as p < 0.05, the HA accepted (medians unequal) 
## 
##  Wilcoxon rank sum test with continuity correction
## 
## data:  x and y
## W = 542840, p-value = 0.000907
## alternative hypothesis: true location shift is not equal to 0
## 95 percent confidence interval:
##   9.228801 35.928096
## sample estimates:
## difference in location 
##               22.49199 
## 
## Mann-Whitney test results for NA biovars: actual vs. RCP8.5 2070 
## The H0 is rejected as p < 0.05, the HA accepted (medians unequal) 
## 
##  Wilcoxon rank sum test with continuity correction
## 
## data:  x and y
## W = 302050, p-value < 2.2e-16
## alternative hypothesis: true location shift is not equal to 0
## 95 percent confidence interval:
##  -106.47756  -79.81596
## sample estimates:
## difference in location 
##              -92.92699 
## 
## Mann-Whitney test results for NA biovars: actual vs. RCP4.5 2050 
## The H0 is rejected as p < 0.05, the HA accepted (medians unequal) 
## 
##  Wilcoxon rank sum test with continuity correction
## 
## data:  x and y
## W = 275580, p-value < 2.2e-16
## alternative hypothesis: true location shift is not equal to 0
## 95 percent confidence interval:
##  -138.8839 -109.4071
## sample estimates:
## difference in location 
##              -123.9463 
## 
## Mann-Whitney test results for NA biovars: RCP4.5 2050 vs. RCP4.5 2070 
## The HA is rejected as p >= 0.05, the H0 accepted (medians equal) 
## 
##  Wilcoxon rank sum test with continuity correction
## 
## data:  x and y
## W = 519200, p-value = 0.1371
## alternative hypothesis: true location shift is not equal to 0
## 95 percent confidence interval:
##  -3.677377 26.447192
## sample estimates:
## difference in location 
##               11.44144 
## 
## Mann-Whitney test results for NA biovars: actual vs. RCP4.5 2070 
## The H0 is rejected as p < 0.05, the HA accepted (medians unequal) 
## 
##  Wilcoxon rank sum test with continuity correction
## 
## data:  x and y
## W = 289800, p-value < 2.2e-16
## alternative hypothesis: true location shift is not equal to 0
## 95 percent confidence interval:
##  -124.62379  -96.80859
## sample estimates:
## difference in location 
##              -110.4433 
## 
## Mann-Whitney test results for NA biovars: actual vs. RCP8.5 2050 
## The H0 is rejected as p < 0.05, the HA accepted (medians unequal) 
## 
##  Wilcoxon rank sum test with continuity correction
## 
## data:  x and y
## W = 320600, p-value < 2.2e-16
## alternative hypothesis: true location shift is not equal to 0
## 95 percent confidence interval:
##  -106.17160  -78.20463
## sample estimates:
## difference in location 
##              -92.17437 
## 
## Mann-Whitney test results for NA biovars: RCP8.5 2050 vs. RCP8.5 2070 
## The H0 is rejected as p < 0.05, the HA accepted (medians unequal) 
## 
##  Wilcoxon rank sum test with continuity correction
## 
## data:  x and y
## W = 526890, p-value = 0.03734
## alternative hypothesis: true location shift is not equal to 0
## 95 percent confidence interval:
##   0.8575242 28.7831643
## sample estimates:
## difference in location 
##               14.64223 
## 
## Mann-Whitney test results for NA biovars: actual vs. RCP8.5 2070 
## The H0 is rejected as p < 0.05, the HA accepted (medians unequal) 
## 
##  Wilcoxon rank sum test with continuity correction
## 
## data:  x and y
## W = 342440, p-value < 2.2e-16
## alternative hypothesis: true location shift is not equal to 0
## 95 percent confidence interval:
##  -90.64549 -64.11195
## sample estimates:
## difference in location 
##              -77.18968 
## 
## Mann-Whitney test results for NA biovars: actual vs. RCP4.5 2050 
## The H0 is rejected as p < 0.05, the HA accepted (medians unequal) 
## 
##  Wilcoxon rank sum test with continuity correction
## 
## data:  x and y
## W = 247070, p-value < 2.2e-16
## alternative hypothesis: true location shift is not equal to 0
## 95 percent confidence interval:
##  -286.8819 -235.1138
## sample estimates:
## difference in location 
##              -260.8661 
## 
## Mann-Whitney test results for NA biovars: RCP4.5 2050 vs. RCP4.5 2070 
## The HA is rejected as p >= 0.05, the H0 accepted (medians equal) 
## 
##  Wilcoxon rank sum test with continuity correction
## 
## data:  x and y
## W = 495610, p-value = 0.7339
## alternative hypothesis: true location shift is not equal to 0
## 95 percent confidence interval:
##  -30.24458  21.60073
## sample estimates:
## difference in location 
##              -4.323329 
## 
## Mann-Whitney test results for NA biovars: actual vs. RCP4.5 2070 
## The H0 is rejected as p < 0.05, the HA accepted (medians unequal) 
## 
##  Wilcoxon rank sum test with continuity correction
## 
## data:  x and y
## W = 242110, p-value < 2.2e-16
## alternative hypothesis: true location shift is not equal to 0
## 95 percent confidence interval:
##  -290.6256 -238.9875
## sample estimates:
## difference in location 
##              -264.6519 
## 
## Mann-Whitney test results for NA biovars: actual vs. RCP8.5 2050 
## The H0 is rejected as p < 0.05, the HA accepted (medians unequal) 
## 
##  Wilcoxon rank sum test with continuity correction
## 
## data:  x and y
## W = 296740, p-value < 2.2e-16
## alternative hypothesis: true location shift is not equal to 0
## 95 percent confidence interval:
##  -219.0201 -170.3920
## sample estimates:
## difference in location 
##              -194.5876 
## 
## Mann-Whitney test results for NA biovars: RCP8.5 2050 vs. RCP8.5 2070 
## The H0 is rejected as p < 0.05, the HA accepted (medians unequal) 
## 
##  Wilcoxon rank sum test with continuity correction
## 
## data:  x and y
## W = 380530, p-value < 2.2e-16
## alternative hypothesis: true location shift is not equal to 0
## 95 percent confidence interval:
##  -166.6901 -109.8155
## sample estimates:
## difference in location 
##              -138.1153 
## 
## Mann-Whitney test results for NA biovars: actual vs. RCP8.5 2070 
## The H0 is rejected as p < 0.05, the HA accepted (medians unequal) 
## 
##  Wilcoxon rank sum test with continuity correction
## 
## data:  x and y
## W = 207280, p-value < 2.2e-16
## alternative hypothesis: true location shift is not equal to 0
## 95 percent confidence interval:
##  -357.1485 -300.8912
## sample estimates:
## difference in location 
##              -328.8912
```

###### Mann-Whitney U test results visualization

- H0 - null hypothesis (medians equal)
- HA - alternative hypothesis (medians significantly different)

```
library(pheatmap)
```

```
## Warning: package 'pheatmap' was built under R version 3.4.3
```

```
draw_colnames_90 <- function(coln, gaps, ...) {
    coord = pheatmap:::find_coordinates(length(coln), gaps)
    x = coord$coord - 0.5 * coord$size
    res = grid:::textGrob(coln, x = x, y = grid:::unit(1, "npc") - grid:::unit(3, "bigpts"), 
                          vjust = 0.5, hjust = 1, rot = 90, gp = grid:::gpar(...))
    return(res)
}  

# override draw_colnames function from pheatmap with custom draw_colnames_90 function
# works for current session
assignInNamespace(x="draw_colnames", value="draw_colnames_90", ns=asNamespace("pheatmap"))  


pheatmap(t(MWU_prec_RCP45_matrix), color = c('darkgreen','lightgray'), border_color = 'black', 
         cellwidth = 50, cellheight = 50,
         symm = FALSE, cluster_rows = FALSE, cluster_cols = FALSE,
         legend_breaks = c(0,0.25,0.5,0.75,1), legend_labels = c('','H0','','HA',''),
         labels_col = month_list,
         labels_row = c('actual vs. RCP4.5 2050','RCP4.5 2050 vs. RCP4.5 2070','actual vs. RCP4.5 2070'),
         fontsize = 16)
```

```
pheatmap(t(MWU_prec_RCP85_matrix), color = c('darkgreen','lightgray'), border_color = 'black', 
         cellwidth = 50, cellheight = 50,
         symm = FALSE, cluster_rows = FALSE, cluster_cols = FALSE,
         legend_breaks = c(0,0.25,0.5,0.75,1), legend_labels = c('','H0','','HA',''),
         labels_col = month_list,
         labels_row = c('actual vs. RCP8.5 2050','RCP8.5 2050 vs. RCP8.5 2070','actual vs. RCP8.5 2070'),
         fontsize = 16)
```

###### Mann-Whitney U test results for climatic variables

- H0 - null hypothesis (medians equal)
- HA - alternative hypothesis (medians significantly different)

```
MWU_prec_RCP45_matrix_s <- format(MWU_prec_RCP45_matrix_s, digits=2, nsmall=2)
MWU_tmin_RCP45_matrix_s <- format(MWU_tmin_RCP45_matrix_s, digits=2, nsmall=2)
MWU_tmax_RCP45_matrix_s <- format(MWU_tmax_RCP45_matrix_s, digits=2, nsmall=2)

pheatmap(rbind(t(MWU_prec_RCP45_matrix),
               t(MWU_tmin_RCP45_matrix),
               t(MWU_tmax_RCP45_matrix)), 
         display_numbers = rbind(
               t(MWU_prec_RCP45_matrix_s),
               t(MWU_tmin_RCP45_matrix_s),
               t(MWU_tmax_RCP45_matrix_s)),
         fontsize_number = 13, number_color = 'darkgreen',
         color = c('darkgreen','lightgray'), border_color = 'black', 
         cellwidth = 50, cellheight = 50,
         symm = FALSE, cluster_rows = FALSE, cluster_cols = FALSE,
         gaps_row = c(3,6),
         legend_breaks = c(0,0.25,0.5,0.75,1), legend_labels = c('','H0','','HA',''),
         labels_col = month_list,
         labels_row = c('Prec actual vs. 2050','Prec 2050 vs. 2070','Prec actual vs. 2070',
                        'Tmin actual vs. 2050','Tmin 2050 vs. 2070','Tmin actual vs. 2070',
                        'Tmax actual vs. 2050','Tmax 2050 vs. 2070','Tmax actual vs. 2070'),
         main = 'RCP4.5', fontsize = 16)
```

```
MWU_prec_RCP85_matrix_s <- format(MWU_prec_RCP85_matrix_s, digits=2, nsmall=2)
MWU_tmin_RCP85_matrix_s <- format(MWU_tmin_RCP85_matrix_s, digits=2, nsmall=2)
MWU_tmax_RCP85_matrix_s <- format(MWU_tmax_RCP85_matrix_s, digits=2, nsmall=2)

pheatmap(rbind(t(MWU_prec_RCP85_matrix),
               t(MWU_tmin_RCP85_matrix),
               t(MWU_tmax_RCP85_matrix)), 
         display_numbers = rbind(
               t(MWU_prec_RCP85_matrix_s),
               t(MWU_tmin_RCP85_matrix_s),
               t(MWU_tmax_RCP85_matrix_s)),
         fontsize_number = 13, number_color = 'darkgreen',
         color = c('darkgreen','lightgray'), border_color = 'black', 
         cellwidth = 50, cellheight = 50,
         symm = FALSE, cluster_rows = FALSE, cluster_cols = FALSE,
         gaps_row = c(3,6),
         legend_breaks = c(0,0.25,0.5,0.75,1), legend_labels = c('','H0','','HA',''),
         labels_col = month_list,
         labels_row = c('Prec actual vs. 2050','Prec 2050 vs. 2070','Prec actual vs. 2070',
                        'Tmin actual vs. 2050','Tmin 2050 vs. 2070','Tmin actual vs. 2070',
                        'Tmax actual vs. 2050','Tmax 2050 vs. 2070','Tmax actual vs. 2070'),
         main = 'RCP8.5', fontsize = 16)
```

###### Mann-Whitney U test results for bioclimatic predictors

- H0 - null hypothesis (medians equal)
- HA - alternative hypothesis (medians significantly different)

```
MWU_biovars_RCP45_matrix_s <- format(MWU_biovars_RCP45_matrix_s, digits=2, nsmall=2)
MWU_biovars_RCP85_matrix_s <- format(MWU_biovars_RCP85_matrix_s, digits=2, nsmall=2)

pheatmap(t(MWU_biovars_RCP45_matrix), 
         display_numbers = t(MWU_biovars_RCP45_matrix_s),
         fontsize_number = 13, number_color = 'darkgreen',
         color = c('darkgreen','lightgray'), border_color = 'black', 
         cellwidth = 50, cellheight = 50,
         symm = FALSE, cluster_rows = FALSE, cluster_cols = FALSE,
         legend_breaks = c(0,0.25,0.5,0.75,1), legend_labels = c('','H0','','HA',''),
         labels_col = biovars_names,
         labels_row = c('Biovars actual vs. 2050','Biovars 2050 vs. 2070','Biovars actual vs. 2070'),
         main = 'RCP4.5', fontsize = 16)
```

```
pheatmap(t(MWU_biovars_RCP85_matrix),
         display_numbers = t(MWU_biovars_RCP85_matrix_s),
         fontsize_number = 13, number_color = 'darkgreen',
         color = c('darkgreen','lightgray'), border_color = 'black', 
         cellwidth = 50, cellheight = 50,
         symm = FALSE, cluster_rows = FALSE, cluster_cols = FALSE,
         legend_breaks = c(0,0.25,0.5,0.75,1), legend_labels = c('','H0','','HA',''),
         labels_col = biovars_names,
         labels_row = c('Biovars actual vs. 2050','Biovars 2050 vs. 2070','Biovars actual vs. 2070'),
         main = 'RCP8.5', fontsize = 16)
```
