## Supplementary material for "Vulnerability to climate change for narrowly ranged species: the case of Ecuadorian endemic *Magnolia mercedesiarum*": M_mercedesiarum_climate_change_with_U.Rmd

````
---
title: "Auxiliar script for Shalisko et al. 2018"
author: "Viacheslav Shalisko"
date: "November 15, 2017"
output: html_document
---

*Script for Shalisko et al. 2018 - Comparison of current and future conditions
========================================================

### Current climate conditions WorldClim ver. 2 (1970-2000)
### Predictive climatic model HadGEM2 - ES, scenarios RCP45 and RCP85 in 2050 and 2070 


```{r load-libs}
knitr::opts_chunk$set(echo = TRUE)
knitr::opts_chunk$set(error = TRUE)

library(maptools)
library(rworldmap)    # worldmap datasets
library(rworldxtra)   # hires worldmap spatial dataframe

library(sp)
library(raster)
library(dismo)
library(rgdal)
```

```{r load-subs}
compareMWT = function(x, y, test_title, alternative='two.sided') {
  wilcoxTest_result <- wilcox.test(x,y,alternative=alternative,conf.int=TRUE)
  cat('Mann-Whitney test results for',test_title,"\n",sep=' ')
  if (wilcoxTest_result$p.value < 0.05) {
      cat('The H0 is rejected as p < 0.05, the HA accepted (medians unequal)',"\n")
  } else {
      cat('The HA is rejected as p >= 0.05, the H0 accepted (medians equal)',"\n")
  }  
  print(wilcoxTest_result)
}
```


```{r set-variables}
presence_points_count <- 1000
month_labels <- c('Jan','Feb','Mar','Apr','May','Jun','Jul','Aug','Sep','Oct','Nov','Dic')
biovars_names <- c("bio1","bio2","bio3","bio4",
              "bio5","bio6","bio7","bio8","bio9",
              "bio10","bio11","bio12","bio13","bio14",
              "bio15","bio16","bio17","bio18","bio19")
month_list <- c('Jan','Feb','Mar','Apr','May','Jun','Jul','Aug','Sep','Oct','Nov','Dic')
```


```{r load-rasters}
# load raster of predicted presence
presence_raster_mask <- stack("C:/Users/vshal/GD/Projects_actual/bot_Magnolia_mercedesiarum/Phytotaxa_2017/Analysis/SIG/Rasters_run_22_23/Magnolia_mercedesiarum_ETSS_raster_small.tif")

mde <- stack("C:/Users/vshal/Downloads/GMTED2010/GMTED2010_075_Andes.tif")

# load actual conditions
my_path0 <- 'C:\\Users\\vshal\\Downloads\\CliMond\\'

prec00 <- stack(paste(my_path0,"wc2.0_HR_prec_stack.tif",sep=""))
tmin00 <- stack(paste(my_path0,"wc2.0_HR_tmin_stack.tif",sep=""))
tmax00 <- stack(paste(my_path0,"wc2.0_HR_tmax_stack.tif",sep=""))

biovars00 <- stack(paste(my_path0,"biovars_climond2_Andes_HR.tif",sep=""))
names(biovars00) <- biovars_names

# load future conditions
my_path1 <- 'C:\\Users\\vshal\\Downloads\\future_conditions\\'

prec50 <- stack(paste(my_path1,"he45pr50_stack.tif",sep=""))
tmin50 <- stack(paste(my_path1,"he45tn50_stack.tif",sep=""))
tmax50 <- stack(paste(my_path1,"he45tx50_stack.tif",sep=""))

biovars50 <- stack(paste(my_path1,"biovars_he45_50_Andes_HR.tif",sep=""))
names(biovars50) <- biovars_names

prec50A <- stack(paste(my_path1,"he85pr50_stack.tif",sep=""))
tmin50A <- stack(paste(my_path1,"he85tn50_stack.tif",sep=""))
tmax50A <- stack(paste(my_path1,"he85tx50_stack.tif",sep=""))

biovars50A <- stack(paste(my_path1,"biovars_he85_50_Andes_HR.tif",sep=""))
names(biovars50A) <- biovars_names

prec70 <- stack(paste(my_path1,"he45pr70_stack.tif",sep=""))
tmin70 <- stack(paste(my_path1,"he45tn70_stack.tif",sep=""))
tmax70 <- stack(paste(my_path1,"he45tx70_stack.tif",sep=""))

biovars70 <- stack(paste(my_path1,"biovars_he45_70_Andes_HR.tif",sep=""))
names(biovars70) <- biovars_names

prec70A <- stack(paste(my_path1,"he85pr70_stack.tif",sep=""))
tmin70A <- stack(paste(my_path1,"he85tn70_stack.tif",sep=""))
tmax70A <- stack(paste(my_path1,"he85tx70_stack.tif",sep=""))

biovars70A <- stack(paste(my_path1,"biovars_he85_70_Andes_HR.tif",sep=""))
names(biovars70A) <- biovars_names

# predictions
my_path2 <- 'C:/Users/vshal/Downloads/M_mercedesiarum_rasters/run29_climond2/'
prediction00_1 <- stack(paste(my_path2,"Magnolia_mercedesiarum_MAXENT_climond2_HR_run_29_1.tif",sep=""))
prediction00_2 <- stack(paste(my_path2,"Magnolia_mercedesiarum_MAXENT_climond2_HR_run_29_2.tif",sep=""))
prediction00_3 <- stack(paste(my_path2,"Magnolia_mercedesiarum_MAXENT_climond2_HR_run_29_3.tif",sep=""))
prediction00_4 <- stack(paste(my_path2,"Magnolia_mercedesiarum_MAXENT_climond2_HR_run_29_4.tif",sep=""))
prediction00_5 <- stack(paste(my_path2,"Magnolia_mercedesiarum_MAXENT_climond2_HR_run_29_5.tif",sep=""))
prediction00_6 <- stack(paste(my_path2,"Magnolia_mercedesiarum_MAXENT_climond2_HR_run_29_6.tif",sep=""))
prediction00_7 <- stack(paste(my_path2,"Magnolia_mercedesiarum_MAXENT_climond2_HR_run_29_7.tif",sep=""))
prediction00_8 <- stack(paste(my_path2,"Magnolia_mercedesiarum_MAXENT_climond2_HR_run_29_8.tif",sep=""))
prediction00_9 <- stack(paste(my_path2,"Magnolia_mercedesiarum_MAXENT_climond2_HR_run_29_9.tif",sep=""))
prediction00_10 <- stack(paste(my_path2,"Magnolia_mercedesiarum_MAXENT_climond2_HR_run_29_10.tif",sep=""))

prediction50_1 <- stack(paste(my_path2,"Magnolia_mercedesiarum_MAXENT_he45_50_HR_run_29_1.tif",sep=""))
prediction50_2 <- stack(paste(my_path2,"Magnolia_mercedesiarum_MAXENT_he45_50_HR_run_29_2.tif",sep=""))
prediction50_3 <- stack(paste(my_path2,"Magnolia_mercedesiarum_MAXENT_he45_50_HR_run_29_3.tif",sep=""))
prediction50_4 <- stack(paste(my_path2,"Magnolia_mercedesiarum_MAXENT_he45_50_HR_run_29_4.tif",sep=""))
prediction50_5 <- stack(paste(my_path2,"Magnolia_mercedesiarum_MAXENT_he45_50_HR_run_29_5.tif",sep=""))
prediction50_6 <- stack(paste(my_path2,"Magnolia_mercedesiarum_MAXENT_he45_50_HR_run_29_6.tif",sep=""))
prediction50_7 <- stack(paste(my_path2,"Magnolia_mercedesiarum_MAXENT_he45_50_HR_run_29_7.tif",sep=""))
prediction50_8 <- stack(paste(my_path2,"Magnolia_mercedesiarum_MAXENT_he45_50_HR_run_29_8.tif",sep=""))
prediction50_9 <- stack(paste(my_path2,"Magnolia_mercedesiarum_MAXENT_he45_50_HR_run_29_9.tif",sep=""))
prediction50_10 <- stack(paste(my_path2,"Magnolia_mercedesiarum_MAXENT_he45_50_HR_run_29_10.tif",sep=""))

prediction50A_1 <- stack(paste(my_path2,"Magnolia_mercedesiarum_MAXENT_he85_50_HR_run_29_1.tif",sep=""))
prediction50A_2 <- stack(paste(my_path2,"Magnolia_mercedesiarum_MAXENT_he85_50_HR_run_29_2.tif",sep=""))
prediction50A_3 <- stack(paste(my_path2,"Magnolia_mercedesiarum_MAXENT_he85_50_HR_run_29_3.tif",sep=""))
prediction50A_4 <- stack(paste(my_path2,"Magnolia_mercedesiarum_MAXENT_he85_50_HR_run_29_4.tif",sep=""))
prediction50A_5 <- stack(paste(my_path2,"Magnolia_mercedesiarum_MAXENT_he85_50_HR_run_29_5.tif",sep=""))
prediction50A_6 <- stack(paste(my_path2,"Magnolia_mercedesiarum_MAXENT_he85_50_HR_run_29_6.tif",sep=""))
prediction50A_7 <- stack(paste(my_path2,"Magnolia_mercedesiarum_MAXENT_he85_50_HR_run_29_7.tif",sep=""))
prediction50A_8 <- stack(paste(my_path2,"Magnolia_mercedesiarum_MAXENT_he85_50_HR_run_29_8.tif",sep=""))
prediction50A_9 <- stack(paste(my_path2,"Magnolia_mercedesiarum_MAXENT_he85_50_HR_run_29_9.tif",sep=""))
prediction50A_10 <- stack(paste(my_path2,"Magnolia_mercedesiarum_MAXENT_he85_50_HR_run_29_10.tif",sep=""))

prediction70_1 <- stack(paste(my_path2,"Magnolia_mercedesiarum_MAXENT_he45_70_HR_run_29_1.tif",sep=""))
prediction70_2 <- stack(paste(my_path2,"Magnolia_mercedesiarum_MAXENT_he45_70_HR_run_29_2.tif",sep=""))
prediction70_3 <- stack(paste(my_path2,"Magnolia_mercedesiarum_MAXENT_he45_70_HR_run_29_3.tif",sep=""))
prediction70_4 <- stack(paste(my_path2,"Magnolia_mercedesiarum_MAXENT_he45_70_HR_run_29_4.tif",sep=""))
prediction70_5 <- stack(paste(my_path2,"Magnolia_mercedesiarum_MAXENT_he45_70_HR_run_29_5.tif",sep=""))
prediction70_6 <- stack(paste(my_path2,"Magnolia_mercedesiarum_MAXENT_he45_70_HR_run_29_6.tif",sep=""))
prediction70_7 <- stack(paste(my_path2,"Magnolia_mercedesiarum_MAXENT_he45_70_HR_run_29_7.tif",sep=""))
prediction70_8 <- stack(paste(my_path2,"Magnolia_mercedesiarum_MAXENT_he45_70_HR_run_29_8.tif",sep=""))
prediction70_9 <- stack(paste(my_path2,"Magnolia_mercedesiarum_MAXENT_he45_70_HR_run_29_9.tif",sep=""))
prediction70_10 <- stack(paste(my_path2,"Magnolia_mercedesiarum_MAXENT_he45_70_HR_run_29_10.tif",sep=""))

prediction70A_1 <- stack(paste(my_path2,"Magnolia_mercedesiarum_MAXENT_he85_70_HR_run_29_1.tif",sep=""))
prediction70A_2 <- stack(paste(my_path2,"Magnolia_mercedesiarum_MAXENT_he85_70_HR_run_29_2.tif",sep=""))
prediction70A_3 <- stack(paste(my_path2,"Magnolia_mercedesiarum_MAXENT_he85_70_HR_run_29_3.tif",sep=""))
prediction70A_4 <- stack(paste(my_path2,"Magnolia_mercedesiarum_MAXENT_he85_70_HR_run_29_4.tif",sep=""))
prediction70A_5 <- stack(paste(my_path2,"Magnolia_mercedesiarum_MAXENT_he85_70_HR_run_29_5.tif",sep=""))
prediction70A_6 <- stack(paste(my_path2,"Magnolia_mercedesiarum_MAXENT_he85_70_HR_run_29_6.tif",sep=""))
prediction70A_7 <- stack(paste(my_path2,"Magnolia_mercedesiarum_MAXENT_he85_70_HR_run_29_7.tif",sep=""))
prediction70A_8 <- stack(paste(my_path2,"Magnolia_mercedesiarum_MAXENT_he85_70_HR_run_29_8.tif",sep=""))
prediction70A_9 <- stack(paste(my_path2,"Magnolia_mercedesiarum_MAXENT_he85_70_HR_run_29_9.tif",sep=""))
prediction70A_10 <- stack(paste(my_path2,"Magnolia_mercedesiarum_MAXENT_he85_70_HR_run_29_10.tif",sep=""))
```

```{r random-points-generation}
set.seed(1)
random_presence_points <- randomPoints(presence_raster_mask, 
                                        n = presence_points_count, 
                                        tryf = 100)
```

#### Render presencce data

```{r points-visualization, fig.height=13, fig.width=16}
par(mfrow=c(1, 2))

mask_reclass_table <- matrix(c(-Inf, 1, 1, 1, Inf, NA), ncol=3, byrow=TRUE)
b_mask <- reclassify(subset(mde,1),mask_reclass_table)

plot(mde, xlim=c(-82,-70), ylim=c(-10,10), col = c(terrain.colors(256)))
plot(b_mask, add=TRUE, col=c('white'), legend = FALSE)
plot(presence_raster_mask, add=TRUE, alpha=0.9, col= gray(0.2), legend = FALSE)
world_high <- getMap(resolution = "high")
plot(world_high, add=TRUE, axes = TRUE)
#points(random_presence_points,col='black',cex=0.7)


plot(mde, xlim=c(-78.5,-76.5), ylim=c(-1.75,1.75), axes=TRUE, col = terrain.colors(256), legend = FALSE)
plot(presence_raster_mask, add=TRUE, alpha=0.4, col= gray(0.2), legend = FALSE)
world_high <- getMap(resolution = "high")
plot(world_high, add=TRUE)
points(random_presence_points,col='black',cex=0.7)
```

```{r sampling, fig.height=13, fig.width=15, cache=TRUE}

Magn_mde <- extract(mde, random_presence_points)

Magn_biovars00 <- extract(biovars00, random_presence_points)
Magn_biovars50 <- extract(biovars50, random_presence_points)
Magn_biovars70 <- extract(biovars70, random_presence_points)
Magn_biovars50A <- extract(biovars50A, random_presence_points)
Magn_biovars70A <- extract(biovars70A, random_presence_points)

Magn_prec00 <- extract(prec00, random_presence_points)
Magn_prec50 <- extract(prec50, random_presence_points)
Magn_prec70 <- extract(prec70, random_presence_points)
Magn_prec50A <- extract(prec50A, random_presence_points)
Magn_prec70A <- extract(prec70A, random_presence_points)

Magn_tmin00 <- extract(tmin00, random_presence_points) / 10
Magn_tmin50 <- extract(tmin50, random_presence_points) / 10
Magn_tmin70 <- extract(tmin70, random_presence_points) / 10
Magn_tmin50A <- extract(tmin50A, random_presence_points) / 10
Magn_tmin70A <- extract(tmin70A, random_presence_points) / 10

Magn_tmax00 <- extract(tmax00, random_presence_points) / 10
Magn_tmax50 <- extract(tmax50, random_presence_points) / 10
Magn_tmax70 <- extract(tmax70, random_presence_points) / 10
Magn_tmax50A <- extract(tmax50A, random_presence_points) / 10
Magn_tmax70A <- extract(tmax70A, random_presence_points) / 10

```

```{r sampling0, fig.height=13, fig.width=15, cache=TRUE}
par(cex = 1.4)

Magn_pred00_1 <- extract(prediction00_1, random_presence_points)
Magn_pred00_2 <- extract(prediction00_2, random_presence_points)
Magn_pred00_3 <- extract(prediction00_3, random_presence_points)
Magn_pred00_4 <- extract(prediction00_4, random_presence_points)
Magn_pred00_5 <- extract(prediction00_5, random_presence_points)
Magn_pred00_6 <- extract(prediction00_6, random_presence_points)
Magn_pred00_7 <- extract(prediction00_7, random_presence_points)
Magn_pred00_8 <- extract(prediction00_8, random_presence_points)
Magn_pred00_9 <- extract(prediction00_9, random_presence_points)
Magn_pred00_10 <- extract(prediction00_10, random_presence_points)

Magn_pred50_1 <- extract(prediction50_1, random_presence_points)
Magn_pred50_2 <- extract(prediction50_2, random_presence_points)
Magn_pred50_3 <- extract(prediction50_3, random_presence_points)
Magn_pred50_4 <- extract(prediction50_4, random_presence_points)
Magn_pred50_5 <- extract(prediction50_5, random_presence_points)
Magn_pred50_6 <- extract(prediction50_6, random_presence_points)
Magn_pred50_7 <- extract(prediction50_7, random_presence_points)
Magn_pred50_8 <- extract(prediction50_8, random_presence_points)
Magn_pred50_9 <- extract(prediction50_9, random_presence_points)
Magn_pred50_10 <- extract(prediction50_10, random_presence_points)

Magn_pred50A_1 <- extract(prediction50A_1, random_presence_points)
Magn_pred50A_2 <- extract(prediction50A_2, random_presence_points)
Magn_pred50A_3 <- extract(prediction50A_3, random_presence_points)
Magn_pred50A_4 <- extract(prediction50A_4, random_presence_points)
Magn_pred50A_5 <- extract(prediction50A_5, random_presence_points)
Magn_pred50A_6 <- extract(prediction50A_6, random_presence_points)
Magn_pred50A_7 <- extract(prediction50A_7, random_presence_points)
Magn_pred50A_8 <- extract(prediction50A_8, random_presence_points)
Magn_pred50A_9 <- extract(prediction50A_9, random_presence_points)
Magn_pred50A_10 <- extract(prediction50A_10, random_presence_points)

Magn_pred70_1 <- extract(prediction70_1, random_presence_points)
Magn_pred70_2 <- extract(prediction70_2, random_presence_points)
Magn_pred70_3 <- extract(prediction70_3, random_presence_points)
Magn_pred70_4 <- extract(prediction70_4, random_presence_points)
Magn_pred70_5 <- extract(prediction70_5, random_presence_points)
Magn_pred70_6 <- extract(prediction70_6, random_presence_points)
Magn_pred70_7 <- extract(prediction70_7, random_presence_points)
Magn_pred70_8 <- extract(prediction70_8, random_presence_points)
Magn_pred70_9 <- extract(prediction70_9, random_presence_points)
Magn_pred70_10 <- extract(prediction70_10, random_presence_points)

Magn_pred70A_1 <- extract(prediction70A_1, random_presence_points)
Magn_pred70A_2 <- extract(prediction70A_2, random_presence_points)
Magn_pred70A_3 <- extract(prediction70A_3, random_presence_points)
Magn_pred70A_4 <- extract(prediction70A_4, random_presence_points)
Magn_pred70A_5 <- extract(prediction70A_5, random_presence_points)
Magn_pred70A_6 <- extract(prediction70A_6, random_presence_points)
Magn_pred70A_7 <- extract(prediction70A_7, random_presence_points)
Magn_pred70A_8 <- extract(prediction70A_8, random_presence_points)
Magn_pred70A_9 <- extract(prediction70A_9, random_presence_points)
Magn_pred70A_10 <- extract(prediction70A_10, random_presence_points)

Magn_pred00 <- c(Magn_pred00_1,Magn_pred00_2,Magn_pred00_3,Magn_pred00_4,Magn_pred00_5,
                 Magn_pred00_6,Magn_pred00_7,Magn_pred00_8,Magn_pred00_9,Magn_pred00_10)
Magn_pred50 <- c(Magn_pred50_1,Magn_pred50_2,Magn_pred50_3,Magn_pred50_4,Magn_pred50_5,
                 Magn_pred50_6,Magn_pred50_7,Magn_pred50_8,Magn_pred50_9,Magn_pred50_10)
Magn_pred50A <- c(Magn_pred50A_1,Magn_pred50A_2,Magn_pred50A_3,Magn_pred50A_4,Magn_pred50A_5,
                 Magn_pred50A_6,Magn_pred50A_7,Magn_pred50A_8,Magn_pred50A_9,Magn_pred50A_10)
Magn_pred70 <- c(Magn_pred70_1,Magn_pred70_2,Magn_pred70_3,Magn_pred70_4,Magn_pred70_5,
                 Magn_pred70_6,Magn_pred70_7,Magn_pred70_8,Magn_pred70_9,Magn_pred70_10)
Magn_pred70A <- c(Magn_pred70A_1,Magn_pred70A_2,Magn_pred70A_3,Magn_pred70A_4,Magn_pred70A_5,
                 Magn_pred70A_6,Magn_pred70A_7,Magn_pred70A_8,Magn_pred70A_9,Magn_pred70A_10)

```

## Elevation

```{r elevation, fig.height=13, fig.width=15}
summary(Magn_mde)
quantile(Magn_mde, probs = c(0.05, 0.95))
boxplot(Magn_mde, boxfill = 'lightgray')
```

```{r elevation2, fig.height=13, fig.width=15}
par(mfrow=c(1, 2))

qqnorm(Magn_mde, col="black", main='Normality QQ plot')
qqline(Magn_mde, col="red")

plot(density(Magn_mde), type="n", main = "Distribution along the altitudinal range")
polygon(density(Magn_mde), col="lightgray")
```


## Effect on habitat suitabilit as presence probability (PP), Scenarios RCP 4.5 & RCP 8.5

```{r visualization-1, fig.height=13, fig.width=15}

# etss vector from other script (temporal solution)
etss_vector <- c(0.4777498,0.4717498,0.5180254,0.4159884,0.4304767,0.4204661,0.4722541,0.4856447,0.4036238,0.3599727)


summary(Magn_pred00)
quantile(Magn_pred00, probs = c(0.05,0.1,0.9,0.95))

summary(Magn_pred50)
quantile(Magn_pred50, probs = c(0.05,0.1,0.9,0.95))

summary(Magn_pred50A)
quantile(Magn_pred50A, probs = c(0.05,0.1,0.9,0.95))

summary(Magn_pred70)
quantile(Magn_pred70, probs = c(0.05,0.1,0.9,0.95))

summary(Magn_pred70A)
quantile(Magn_pred70A, probs = c(0.05,0.1,0.9,0.95))

par(cex = 1.4)

boxplot(Magn_pred00, Magn_pred50, Magn_pred70, 
        boxfill = c('lightgreen','coral','red4'),
        names = c('1970-2000','2050 RCP4.5','2070 RCP4.5'),
        main = 'Probability of presence within the actual distribution area RCP4.5')

boxplot(Magn_pred00, Magn_pred50A, Magn_pred70A, 
        boxfill = c('lightgreen','red3','tan4'),
        names = c('1970-2000','2050 RCP8.5','2070 RCP8.5'),
        main = 'Probability of presence within the actual distribution area RCP8.5')

boxplot(Magn_pred00, Magn_pred50, Magn_pred50A, Magn_pred70, Magn_pred70A, 
        boxfill = c('lightgreen','coral','red3','red4','tan4'),
        names = c('1970-2000','2050 RCP4.5','2050 RCP8.5','2070 RCP4.5','2070 RCP8.5'),
        main = 'Probability of presence within the actual distribution area RCP4.5 & RCP8.5')

etss_vector_median <- median(etss_vector)
etss_vector_conf <- wilcox.test(etss_vector, conf.int=TRUE)$conf.int
abline(h=etss_vector_median, col="red")
abline(h=etss_vector_conf, lty="dotted", col="red")


compareMWT(Magn_pred00,Magn_pred50,'actual PP vs. RCP4.5 2050 PP')
compareMWT(Magn_pred50,Magn_pred70,'RCP4.5 2050 PP vs. RCP4.5 2070 PP')
compareMWT(Magn_pred00,Magn_pred70,'actual PP vs. RCP45 2070 PP')

compareMWT(Magn_pred00,Magn_pred50A,'actual PP vs. RCP85 2050 PP')
compareMWT(Magn_pred50A,Magn_pred70A,'RCP8.5 2050 PP vs. RCP8.5 2070 PP')
compareMWT(Magn_pred00,Magn_pred70A,'actual PP vs. RCP8.5 2070 PP')

#compareMWT(Magn_pred50A,Magn_pred70,'RCP85 2050 PP vs. RCP45 2070 PP')

```


## Climatic variables, Model HadGEM2 - ES, Scenario RCP 4.5
#### Precipitation

```{r visualization-5, fig.height=13, fig.width=15}
par(cex = 1.4)

summary(Magn_prec00)
summary(Magn_prec50)
summary(Magn_prec70)

boxplot(Magn_prec00, ylim = c(0,650), xlim = c(0.5, 12.5), names = month_labels,
        boxfill=rgb(1, 1, 1, alpha=1), border=rgb(1, 1, 1, alpha=1),
        main = 'Monthly precipitation', ylab = 'mm')
boxplot(Magn_prec00, at = 1:12 - 0.25, boxwex=0.2, xaxt = "n", add = TRUE, boxfill = 'lightgreen')
boxplot(Magn_prec50, at = 1:12, boxwex=0.2, xaxt = "n", add = TRUE, boxfill = 'coral')
boxplot(Magn_prec70, at = 1:12 + 0.25, boxwex=0.2, xaxt = "n", add = TRUE, boxfill = 'red4')

legend("topleft", c("1970-2000 (WorldClim 2)","2050 (RCP4.5)","2070 (RCP4.5)"), 
       inset=.02, box.lty=0, fill=c('lightgreen','coral','red4'), horiz=TRUE, cex=1.2)

```


```{r visualization-6, fig.height=13, fig.width=15}
par(cex = 1.4)

summary(Magn_tmin00)
summary(Magn_tmin50)
summary(Magn_tmin70)

boxplot(Magn_tmin00, ylim = c(5,25), xlim = c(0.5, 12.5), names = month_labels,
        boxfill=rgb(1, 1, 1, alpha=1), border=rgb(1, 1, 1, alpha=1),
        main = 'Monthly minimal temperature', ylab = '°C')
boxplot(Magn_tmin00, at = 1:12 - 0.25, boxwex=0.2, xaxt = "n", add = TRUE, boxfill = 'lightgreen')
boxplot(Magn_tmin50, at = 1:12, boxwex=0.2, xaxt = "n", add = TRUE, boxfill = 'coral')
boxplot(Magn_tmin70, at = 1:12 + 0.25, boxwex=0.2, xaxt = "n", add = TRUE, boxfill = 'red4')

legend("topleft", c("1970-2000 (WorldClim 2)","2050 (RCP 4.5)","2070 (RCP 4.5)"), 
       inset=.02, box.lty=0, fill=c('lightgreen','coral','red4'), horiz=TRUE, cex=1.2)
```

```{r visualization-7, fig.height=13, fig.width=15}
par(cex = 1.4)

summary(Magn_tmax00)
summary(Magn_tmax50)
summary(Magn_tmax70)

boxplot(Magn_tmin00, ylim = c(15,32), xlim = c(0.5, 12.5), names = month_labels,
        boxfill=rgb(1, 1, 1, alpha=1), border=rgb(1, 1, 1, alpha=1),
        main = 'Monthly maximum temperature', ylab = '°C')
boxplot(Magn_tmax00, at = 1:12 - 0.25, boxwex=0.2, xaxt = "n", add = TRUE, boxfill = 'lightgreen')
boxplot(Magn_tmax50, at = 1:12, boxwex=0.2, xaxt = "n", add = TRUE, boxfill = 'coral')
boxplot(Magn_tmax70, at = 1:12 + 0.25, boxwex=0.2, xaxt = "n", add = TRUE, boxfill = 'red4')

legend("topleft", c("1970-2000 (WorldClim 2)","2050 (RCP 4.5)","2070 (RCP 4.5)"), 
       inset=.02, box.lty=0, fill=c('lightgreen','coral','red4'), horiz=TRUE, cex=1.2)
```

## Climatic variables, Model HadGEM2 - ES, Scenario RCP 8.5

```{r visualization-8, fig.height=13, fig.width=15}
par(cex = 1.4)

summary(Magn_prec00)
summary(Magn_prec50A)
summary(Magn_prec70A)

boxplot(Magn_prec00, ylim = c(0,650), xlim = c(0.5, 12.5), names = month_labels,
        boxfill=rgb(1, 1, 1, alpha=1), border=rgb(1, 1, 1, alpha=1),
        main = 'Monthly precipitation', ylab = 'mm')
boxplot(Magn_prec00, at = 1:12 - 0.25, boxwex=0.2, xaxt = "n", add = TRUE, boxfill = 'lightgreen')
boxplot(Magn_prec50A, at = 1:12, boxwex=0.2, xaxt = "n", add = TRUE, boxfill = 'coral')
boxplot(Magn_prec70A, at = 1:12 + 0.25, boxwex=0.2, xaxt = "n", add = TRUE, boxfill = 'red4')

legend("topleft", c("1970-2000 (WorldClim 2)","2050 (RCP 8.5)","2070 (RCP 8.5)"), 
       inset=.02, box.lty=0, fill=c('lightgreen','coral','red4'), horiz=TRUE, cex=1.2)
```

```{r visualization-9, fig.height=13, fig.width=15}
par(cex = 1.4)

summary(Magn_tmin00)
summary(Magn_tmin50A)
summary(Magn_tmin70A)

boxplot(Magn_tmin00, ylim = c(5,25), xlim = c(0.5, 12.5), names = month_labels,
        boxfill=rgb(1, 1, 1, alpha=1), border=rgb(1, 1, 1, alpha=1),
        main = 'Monthly minimal temperature', ylab = '°C')
boxplot(Magn_tmin00, at = 1:12 - 0.25, boxwex=0.2, xaxt = "n", add = TRUE, boxfill = 'lightgreen')
boxplot(Magn_tmin50A, at = 1:12, boxwex=0.2, xaxt = "n", add = TRUE, boxfill = 'coral')
boxplot(Magn_tmin70A, at = 1:12 + 0.25, boxwex=0.2, xaxt = "n", add = TRUE, boxfill = 'red4')

legend("topleft", c("1970-2000 (WorldClim 2)","2050 (RCP 8.5)","2070 (RCP 8.5)"), 
       inset=.02, box.lty=0, fill=c('lightgreen','coral','red4'), horiz=TRUE, cex=1.2)
```

```{r visualization-10, fig.height=13, fig.width=15}
par(cex = 1.4)

summary(Magn_tmax00)
summary(Magn_tmax50A)
summary(Magn_tmax70A)

boxplot(Magn_tmin00, ylim = c(15,32), xlim = c(0.5, 12.5), names = month_labels,
        boxfill=rgb(1, 1, 1, alpha=1), border=rgb(1, 1, 1, alpha=1),
        main = 'Monthly maximum temperature', ylab = '°C')
boxplot(Magn_tmax00, at = 1:12 - 0.25, boxwex=0.2, xaxt = "n", add = TRUE, boxfill = 'lightgreen')
boxplot(Magn_tmax50A, at = 1:12, boxwex=0.2, xaxt = "n", add = TRUE, boxfill = 'coral')
boxplot(Magn_tmax70A, at = 1:12 + 0.25, boxwex=0.2, xaxt = "n", add = TRUE, boxfill = 'red4')

legend("topleft", c("1970-2000 (WorldClim 2)","2050 (RCP 8.5)","2070 (RCP 8.5)"), 
       inset=.02, box.lty=0, fill=c('lightgreen','coral','red4'), horiz=TRUE, cex=1.2)
```

## Climatic variables, Model HadGEM2 - ES, Scenarios RCP 4.5 & RCP 8.5

```{r visualization-11, fig.height=13, fig.width=15}
par(cex = 1.4, mfrow=c(2, 2))

summary(Magn_biovars00)
summary(Magn_biovars50)
summary(Magn_biovars50A)
summary(Magn_biovars70)
summary(Magn_biovars70A)

boxplot(Magn_biovars00[,'bio1'], Magn_biovars50[,'bio1'], Magn_biovars50A[,'bio1'], Magn_biovars70[,'bio1'], Magn_biovars70A[,'bio1'], 
        boxfill = c('lightgreen','coral','red3','red4','tan4'),
        names = c('1970-2000','2050 RCP4.5','2050 RCP8.5','2070 RCP4.5','2070 RCP8.5'),
        main = 'BIO 1')
boxplot(Magn_biovars00[,'bio2'], Magn_biovars50[,'bio2'], Magn_biovars50A[,'bio2'], Magn_biovars70[,'bio2'], Magn_biovars70A[,'bio2'], 
        boxfill = c('lightgreen','coral','red3','red4','tan4'),
        names = c('1970-2000','2050 RCP4.5','2050 RCP8.5','2070 RCP4.5','2070 RCP8.5'),
        main = 'BIO 2')
boxplot(Magn_biovars00[,'bio3'], Magn_biovars50[,'bio3'], Magn_biovars50A[,'bio3'], Magn_biovars70[,'bio3'], Magn_biovars70A[,'bio3'], 
        boxfill = c('lightgreen','coral','red3','red4','tan4'),
        names = c('1970-2000','2050 RCP4.5','2050 RCP8.5','2070 RCP4.5','2070 RCP8.5'),
        main = 'BIO 3')
boxplot(Magn_biovars00[,'bio4'], Magn_biovars50[,'bio4'], Magn_biovars50A[,'bio4'], Magn_biovars70[,'bio4'], Magn_biovars70A[,'bio4'], 
        boxfill = c('lightgreen','coral','red3','red4','tan4'),
        names = c('1970-2000','2050 RCP4.5','2050 RCP8.5','2070 RCP4.5','2070 RCP8.5'),
        main = 'BIO 4')

boxplot(Magn_biovars00[,'bio5'], Magn_biovars50[,'bio5'], Magn_biovars50A[,'bio5'], Magn_biovars70[,'bio5'], Magn_biovars70A[,'bio5'], 
        boxfill = c('lightgreen','coral','red3','red4','tan4'),
        names = c('1970-2000','2050 RCP4.5','2050 RCP8.5','2070 RCP4.5','2070 RCP8.5'),
        main = 'BIO 5')
boxplot(Magn_biovars00[,'bio6'], Magn_biovars50[,'bio6'], Magn_biovars50A[,'bio6'], Magn_biovars70[,'bio6'], Magn_biovars70A[,'bio6'], 
        boxfill = c('lightgreen','coral','red3','red4','tan4'),
        names = c('1970-2000','2050 RCP4.5','2050 RCP8.5','2070 RCP4.5','2070 RCP8.5'),
        main = 'BIO 6')
boxplot(Magn_biovars00[,'bio7'], Magn_biovars50[,'bio7'], Magn_biovars50A[,'bio7'], Magn_biovars70[,'bio7'], Magn_biovars70A[,'bio7'], 
        boxfill = c('lightgreen','coral','red3','red4','tan4'),
        names = c('1970-2000','2050 RCP4.5','2050 RCP8.5','2070 RCP4.5','2070 RCP8.5'),
        main = 'BIO 7')
boxplot(Magn_biovars00[,'bio8'], Magn_biovars50[,'bio8'], Magn_biovars50A[,'bio8'], Magn_biovars70[,'bio8'], Magn_biovars70A[,'bio8'], 
        boxfill = c('lightgreen','coral','red3','red4','tan4'),
        names = c('1970-2000','2050 RCP4.5','2050 RCP8.5','2070 RCP4.5','2070 RCP8.5'),
        main = 'BIO 8')

boxplot(Magn_biovars00[,'bio9'], Magn_biovars50[,'bio9'], Magn_biovars50A[,'bio9'], Magn_biovars70[,'bio9'], Magn_biovars70A[,'bio9'], 
        boxfill = c('lightgreen','coral','red3','red4','tan4'),
        names = c('1970-2000','2050 RCP4.5','2050 RCP8.5','2070 RCP4.5','2070 RCP8.5'),
        main = 'BIO 9')
boxplot(Magn_biovars00[,'bio10'], Magn_biovars50[,'bio10'], Magn_biovars50A[,'bio10'], Magn_biovars70[,'bio10'], Magn_biovars70A[,'bio10'], 
        boxfill = c('lightgreen','coral','red3','red4','tan4'),
        names = c('1970-2000','2050 RCP4.5','2050 RCP8.5','2070 RCP4.5','2070 RCP8.5'),
        main = 'BIO 10')
boxplot(Magn_biovars00[,'bio11'], Magn_biovars50[,'bio11'], Magn_biovars50A[,'bio11'], Magn_biovars70[,'bio11'], Magn_biovars70A[,'bio11'], 
        boxfill = c('lightgreen','coral','red3','red4','tan4'),
        names = c('1970-2000','2050 RCP4.5','2050 RCP8.5','2070 RCP4.5','2070 RCP8.5'),
        main = 'BIO 11')
boxplot(Magn_biovars00[,'bio12'], Magn_biovars50[,'bio12'], Magn_biovars50A[,'bio12'], Magn_biovars70[,'bio12'], Magn_biovars70A[,'bio12'], 
        boxfill = c('lightgreen','coral','red3','red4','tan4'),
        names = c('1970-2000','2050 RCP4.5','2050 RCP8.5','2070 RCP4.5','2070 RCP8.5'),
        main = 'BIO 12')

boxplot(Magn_biovars00[,'bio13'], Magn_biovars50[,'bio13'], Magn_biovars50A[,'bio13'], Magn_biovars70[,'bio13'], Magn_biovars70A[,'bio13'], 
        boxfill = c('lightgreen','coral','red3','red4','tan4'),
        names = c('1970-2000','2050 RCP4.5','2050 RCP8.5','2070 RCP4.5','2070 RCP8.5'),
        main = 'BIO 13')
boxplot(Magn_biovars00[,'bio14'], Magn_biovars50[,'bio14'], Magn_biovars50A[,'bio14'], Magn_biovars70[,'bio14'], Magn_biovars70A[,'bio14'], 
        boxfill = c('lightgreen','coral','red3','red4','tan4'),
        names = c('1970-2000','2050 RCP4.5','2050 RCP8.5','2070 RCP4.5','2070 RCP8.5'),
        main = 'BIO 14')
boxplot(Magn_biovars00[,'bio15'], Magn_biovars50[,'bio15'], Magn_biovars50A[,'bio15'], Magn_biovars70[,'bio15'], Magn_biovars70A[,'bio15'], 
        boxfill = c('lightgreen','coral','red3','red4','tan4'),
        names = c('1970-2000','2050 RCP4.5','2050 RCP8.5','2070 RCP4.5','2070 RCP8.5'),
        main = 'BIO 15')
boxplot(Magn_biovars00[,'bio16'], Magn_biovars50[,'bio16'], Magn_biovars50A[,'bio16'], Magn_biovars70[,'bio16'], Magn_biovars70A[,'bio16'], 
        boxfill = c('lightgreen','coral','red3','red4','tan4'),
        names = c('1970-2000','2050 RCP4.5','2050 RCP8.5','2070 RCP4.5','2070 RCP8.5'),
        main = 'BIO 16')

boxplot(Magn_biovars00[,'bio17'], Magn_biovars50[,'bio17'], Magn_biovars50A[,'bio17'], Magn_biovars70[,'bio17'], Magn_biovars70A[,'bio17'], 
        boxfill = c('lightgreen','coral','red3','red4','tan4'),
        names = c('1970-2000','2050 RCP4.5','2050 RCP8.5','2070 RCP4.5','2070 RCP8.5'),
        main = 'BIO 17')
boxplot(Magn_biovars00[,'bio18'], Magn_biovars50[,'bio18'], Magn_biovars50A[,'bio18'], Magn_biovars70[,'bio18'], Magn_biovars70A[,'bio18'], 
        boxfill = c('lightgreen','coral','red3','red4','tan4'),
        names = c('1970-2000','2050 RCP4.5','2050 RCP8.5','2070 RCP4.5','2070 RCP8.5'),
        main = 'BIO 18')
boxplot(Magn_biovars00[,'bio19'], Magn_biovars50[,'bio19'], Magn_biovars50A[,'bio19'], Magn_biovars70[,'bio19'], Magn_biovars70A[,'bio19'], 
        boxfill = c('lightgreen','coral','red3','red4','tan4'),
        names = c('1970-2000','2050 RCP4.5','2050 RCP8.5','2070 RCP4.5','2070 RCP8.5'),
        main = 'BIO 19')
```

#### Mann-Whitney U tests 

```{r cache=TRUE}
# true-false H0 vs. HA matrices
MWU_prec_RCP45_matrix <- matrix(nrow = 12, ncol = 3)
MWU_prec_RCP85_matrix <- matrix(nrow = 12, ncol = 3)
MWU_tmin_RCP45_matrix <- matrix(nrow = 12, ncol = 3)
MWU_tmin_RCP85_matrix <- matrix(nrow = 12, ncol = 3)
MWU_tmax_RCP45_matrix <- matrix(nrow = 12, ncol = 3)
MWU_tmax_RCP85_matrix <- matrix(nrow = 12, ncol = 3)
MWU_biovars_RCP45_matrix <- matrix(nrow = 19, ncol = 3)
MWU_biovars_RCP85_matrix <- matrix(nrow = 19, ncol = 3)

# location shift estimation matrices
MWU_prec_RCP45_matrix_s <- matrix(nrow = 12, ncol = 3)
MWU_prec_RCP85_matrix_s <- matrix(nrow = 12, ncol = 3)
MWU_tmin_RCP45_matrix_s <- matrix(nrow = 12, ncol = 3)
MWU_tmin_RCP85_matrix_s <- matrix(nrow = 12, ncol = 3)
MWU_tmax_RCP45_matrix_s <- matrix(nrow = 12, ncol = 3)
MWU_tmax_RCP85_matrix_s <- matrix(nrow = 12, ncol = 3)
MWU_biovars_RCP45_matrix_s <- matrix(nrow = 19, ncol = 3)
MWU_biovars_RCP85_matrix_s <- matrix(nrow = 19, ncol = 3)


# Mann-Whitney test for prec, tmin, tmax
for (i in 1:12) {
  my_test1 <- compareMWT(Magn_prec00[,i],Magn_prec50[,i],paste(month_list[i],'precipitation: actual vs. RCP4.5 2050'))
  my_test2 <- compareMWT(Magn_prec50[,i],Magn_prec70[,i],paste(month_list[i],'precipitation: RCP4.5 2050 vs. RCP4.5 2070'))
  my_test3 <- compareMWT(Magn_prec00[,i],Magn_prec70[,i],paste(month_list[i],'precipitation: actual vs. RCP4.5 2070'))

  my_test4 <- compareMWT(Magn_prec00[,i],Magn_prec50A[,i],paste(month_list[i],'precipitation: actual vs. RCP8.5 2050'))
  my_test5 <- compareMWT(Magn_prec50A[,i],Magn_prec70A[,i],paste(month_list[i],'precipitation: RCP8.5 2050 vs. RCP8.5 2070'))
  my_test6 <- compareMWT(Magn_prec00[,i],Magn_prec70A[,i],paste(month_list[i],'precipitation: actual vs. RCP8.5 2070'))
 
  my_test7 <- compareMWT(Magn_tmin00[,i],Magn_tmin50[,i],paste(month_list[i],'tmin: actual vs. RCP4.5 2050'))
  my_test8 <- compareMWT(Magn_tmin50[,i],Magn_tmin70[,i],paste(month_list[i],'tmin: RCP4.5 2050 vs. RCP4.5 2070'))
  my_test9 <- compareMWT(Magn_tmin00[,i],Magn_tmin70[,i],paste(month_list[i],'tmin: actual vs. RCP4.5 2070'))

  my_test10 <- compareMWT(Magn_tmin00[,i],Magn_tmin50A[,i],paste(month_list[i],'tmin: actual vs. RCP8.5 2050'))
  my_test11 <- compareMWT(Magn_tmin50A[,i],Magn_tmin70A[,i],paste(month_list[i],'tmin: RCP8.5 2050 vs. RCP8.5 2070'))
  my_test12 <- compareMWT(Magn_tmin00[,i],Magn_tmin70A[,i],paste(month_list[i],'tmin: actual vs. RCP8.5 2070'))   
  
  my_test13 <- compareMWT(Magn_tmax00[,i],Magn_tmax50[,i],paste(month_list[i],'tmax: actual vs. RCP4.5 2050'))
  my_test14 <- compareMWT(Magn_tmax50[,i],Magn_tmax70[,i],paste(month_list[i],'tmax: RCP4.5 2050 vs. RCP4.5 2070'))
  my_test15 <- compareMWT(Magn_tmax00[,i],Magn_tmax70[,i],paste(month_list[i],'tmax: actual vs. RCP4.5 2070'))

  my_test16 <- compareMWT(Magn_tmax00[,i],Magn_tmax50A[,i],paste(month_list[i],'tmax: actual vs. RCP8.5 2050'))
  my_test17 <- compareMWT(Magn_tmax50A[,i],Magn_tmax70A[,i],paste(month_list[i],'tmax: RCP8.5 2050 vs. RCP8.5 2070'))
  my_test18 <- compareMWT(Magn_tmax00[,i],Magn_tmax70A[,i],paste(month_list[i],'tmax: actual vs. RCP8.5 2070'))   
  
  if (my_test1$p.value < 0.05) {
    MWU_prec_RCP45_matrix[i,1] <- 1
    MWU_prec_RCP45_matrix_s[i,1] <- my_test1$estimate
  } else {
    MWU_prec_RCP45_matrix[i,1] <- 0
  }
  if (my_test2$p.value < 0.05) {
    MWU_prec_RCP45_matrix[i,2] <- 1
    MWU_prec_RCP45_matrix_s[i,2] <- my_test2$estimate    
  } else {
    MWU_prec_RCP45_matrix[i,2] <- 0
  }
  if (my_test3$p.value < 0.05) {
    MWU_prec_RCP45_matrix[i,3] <- 1
    MWU_prec_RCP45_matrix_s[i,3] <- my_test3$estimate    
  } else {
    MWU_prec_RCP45_matrix[i,3] <- 0
  }
  if (my_test4$p.value < 0.05) {
    MWU_prec_RCP85_matrix[i,1] <- 1
    MWU_prec_RCP85_matrix_s[i,1] <- my_test4$estimate    
  } else {
    MWU_prec_RCP85_matrix[i,1] <- 0
  }
  if (my_test5$p.value < 0.05) {
    MWU_prec_RCP85_matrix[i,2] <- 1
    MWU_prec_RCP85_matrix_s[i,2] <- my_test5$estimate      
  } else {
    MWU_prec_RCP85_matrix[i,2] <- 0
  }
  if (my_test6$p.value < 0.05) {
    MWU_prec_RCP85_matrix[i,3] <- 1
    MWU_prec_RCP85_matrix_s[i,3] <- my_test6$estimate          
  } else {
    MWU_prec_RCP85_matrix[i,3] <- 0
  }
  
  if (my_test7$p.value < 0.05) {
    MWU_tmin_RCP45_matrix[i,1] <- 1
    MWU_tmin_RCP45_matrix_s[i,1] <- my_test7$estimate          
  } else {
    MWU_tmin_RCP45_matrix[i,1] <- 0
  }
  if (my_test8$p.value < 0.05) {
    MWU_tmin_RCP45_matrix[i,2] <- 1
    MWU_tmin_RCP45_matrix_s[i,2] <- my_test8$estimate       
  } else {
    MWU_tmin_RCP45_matrix[i,2] <- 0
  }
  if (my_test9$p.value < 0.05) {
    MWU_tmin_RCP45_matrix[i,3] <- 1
    MWU_tmin_RCP45_matrix_s[i,3] <- my_test9$estimate       
  } else {
    MWU_tmin_RCP45_matrix[i,3] <- 0
  }
  if (my_test10$p.value < 0.05) {
    MWU_tmin_RCP85_matrix[i,1] <- 1
    MWU_tmin_RCP85_matrix_s[i,1] <- my_test10$estimate         
  } else {
    MWU_tmin_RCP85_matrix[i,1] <- 0
  }
  if (my_test11$p.value < 0.05) {
    MWU_tmin_RCP85_matrix[i,2] <- 1
    MWU_tmin_RCP85_matrix_s[i,2] <- my_test11$estimate         
  } else {
    MWU_tmin_RCP85_matrix[i,2] <- 0
  }
  if (my_test12$p.value < 0.05) {
    MWU_tmin_RCP85_matrix[i,3] <- 1
    MWU_tmin_RCP85_matrix_s[i,3] <- my_test12$estimate         
  } else {
    MWU_tmin_RCP85_matrix[i,3] <- 0
  }

  if (my_test13$p.value < 0.05) {
    MWU_tmax_RCP45_matrix[i,1] <- 1
    MWU_tmax_RCP45_matrix_s[i,1] <- my_test13$estimate       
  } else {
    MWU_tmax_RCP45_matrix[i,1] <- 0
  }
  if (my_test14$p.value < 0.05) {
    MWU_tmax_RCP45_matrix[i,2] <- 1
    MWU_tmax_RCP45_matrix_s[i,2] <- my_test14$estimate    
  } else {
    MWU_tmax_RCP45_matrix[i,2] <- 0
  }
  if (my_test15$p.value < 0.05) {
    MWU_tmax_RCP45_matrix[i,3] <- 1
    MWU_tmax_RCP45_matrix_s[i,3] <- my_test15$estimate    
  } else {
    MWU_tmax_RCP45_matrix[i,3] <- 0
  }
  if (my_test16$p.value < 0.05) {
    MWU_tmax_RCP85_matrix[i,1] <- 1
    MWU_tmax_RCP85_matrix_s[i,1] <- my_test16$estimate        
  } else {
    MWU_tmax_RCP85_matrix[i,1] <- 0
  }
  if (my_test17$p.value < 0.05) {
    MWU_tmax_RCP85_matrix[i,2] <- 1
    MWU_tmax_RCP85_matrix_s[i,2] <- my_test17$estimate       
  } else {
    MWU_tmax_RCP85_matrix[i,2] <- 0
  }
  if (my_test18$p.value < 0.05) {
    MWU_tmax_RCP85_matrix[i,3] <- 1
    MWU_tmax_RCP85_matrix_s[i,3] <- my_test18$estimate       
  } else {
    MWU_tmax_RCP85_matrix[i,3] <- 0
  }
  #print(str(my_test1))
}

# Mann-Whitney test for biovars
for (i in 1:19) {
  my_test19 <- compareMWT(Magn_biovars00[,i],Magn_biovars50[,i],paste(month_list[i],'biovars: actual vs. RCP4.5 2050'))
  my_test20 <- compareMWT(Magn_biovars50[,i],Magn_biovars70[,i],paste(month_list[i],'biovars: RCP4.5 2050 vs. RCP4.5 2070'))
  my_test21 <- compareMWT(Magn_biovars00[,i],Magn_biovars70[,i],paste(month_list[i],'biovars: actual vs. RCP4.5 2070'))
  
  my_test22 <- compareMWT(Magn_biovars00[,i],Magn_biovars50A[,i],paste(month_list[i],'biovars: actual vs. RCP8.5 2050'))
  my_test23 <- compareMWT(Magn_biovars50A[,i],Magn_biovars70A[,i],paste(month_list[i],'biovars: RCP8.5 2050 vs. RCP8.5 2070'))
  my_test24 <- compareMWT(Magn_biovars00[,i],Magn_biovars70A[,i],paste(month_list[i],'biovars: actual vs. RCP8.5 2070'))    
  
  if (my_test19$p.value < 0.05) {
    MWU_biovars_RCP45_matrix[i,1] <- 1
    MWU_biovars_RCP45_matrix_s[i,1] <- my_test19$estimate
  } else {
    MWU_biovars_RCP45_matrix[i,1] <- 0
  }
  if (my_test20$p.value < 0.05) {
    MWU_biovars_RCP45_matrix[i,2] <- 1
    MWU_biovars_RCP45_matrix_s[i,2] <- my_test20$estimate
  } else {
    MWU_biovars_RCP45_matrix[i,2] <- 0
  }  
  if (my_test21$p.value < 0.05) {
    MWU_biovars_RCP45_matrix[i,3] <- 1
    MWU_biovars_RCP45_matrix_s[i,3] <- my_test21$estimate
  } else {
    MWU_biovars_RCP45_matrix[i,3] <- 0
  } 
  if (my_test22$p.value < 0.05) {
    MWU_biovars_RCP85_matrix[i,1] <- 1
    MWU_biovars_RCP85_matrix_s[i,1] <- my_test22$estimate
  } else {
    MWU_biovars_RCP85_matrix[i,1] <- 0
  } 
  if (my_test23$p.value < 0.05) {
    MWU_biovars_RCP85_matrix[i,2] <- 1
    MWU_biovars_RCP85_matrix_s[i,2] <- my_test23$estimate
  } else {
    MWU_biovars_RCP85_matrix[i,2] <- 0
  } 
  if (my_test24$p.value < 0.05) {
    MWU_biovars_RCP85_matrix[i,3] <- 1
    MWU_biovars_RCP85_matrix_s[i,3] <- my_test24$estimate
  } else {
    MWU_biovars_RCP85_matrix[i,3] <- 0
  }   
     
}
```


#### Mann-Whitney U test results visualization

* H0 - null hypothesis (medians equal)
* HA - alternative hypothesis (medians significantly different)

```{r visualization-5A, fig.height=4, fig.width=15}

library(pheatmap)

draw_colnames_90 <- function(coln, gaps, ...) {
    coord = pheatmap:::find_coordinates(length(coln), gaps)
    x = coord$coord - 0.5 * coord$size
    res = grid:::textGrob(coln, x = x, y = grid:::unit(1, "npc") - grid:::unit(3, "bigpts"), 
                          vjust = 0.5, hjust = 1, rot = 90, gp = grid:::gpar(...))
    return(res)
}  

# override draw_colnames function from pheatmap with custom draw_colnames_90 function
# works for current session
assignInNamespace(x="draw_colnames", value="draw_colnames_90", ns=asNamespace("pheatmap"))  


pheatmap(t(MWU_prec_RCP45_matrix), color = c('darkgreen','lightgray'), border_color = 'black', 
         cellwidth = 50, cellheight = 50,
         symm = FALSE, cluster_rows = FALSE, cluster_cols = FALSE,
         legend_breaks = c(0,0.25,0.5,0.75,1), legend_labels = c('','H0','','HA',''),
         labels_col = month_list,
         labels_row = c('actual vs. RCP4.5 2050','RCP4.5 2050 vs. RCP4.5 2070','actual vs. RCP4.5 2070'),
         fontsize = 16)

pheatmap(t(MWU_prec_RCP85_matrix), color = c('darkgreen','lightgray'), border_color = 'black', 
         cellwidth = 50, cellheight = 50,
         symm = FALSE, cluster_rows = FALSE, cluster_cols = FALSE,
         legend_breaks = c(0,0.25,0.5,0.75,1), legend_labels = c('','H0','','HA',''),
         labels_col = month_list,
         labels_row = c('actual vs. RCP8.5 2050','RCP8.5 2050 vs. RCP8.5 2070','actual vs. RCP8.5 2070'),
         fontsize = 16)

```


#### Mann-Whitney U test results for climatic variables

* H0 - null hypothesis (medians equal)
* HA - alternative hypothesis (medians significantly different)

```{r visualization-5B, fig.height=8, fig.width=15}

MWU_prec_RCP45_matrix_s <- format(MWU_prec_RCP45_matrix_s, digits=2, nsmall=2)
MWU_tmin_RCP45_matrix_s <- format(MWU_tmin_RCP45_matrix_s, digits=2, nsmall=2)
MWU_tmax_RCP45_matrix_s <- format(MWU_tmax_RCP45_matrix_s, digits=2, nsmall=2)

pheatmap(rbind(t(MWU_prec_RCP45_matrix),
               t(MWU_tmin_RCP45_matrix),
               t(MWU_tmax_RCP45_matrix)), 
         display_numbers = rbind(
               t(MWU_prec_RCP45_matrix_s),
               t(MWU_tmin_RCP45_matrix_s),
               t(MWU_tmax_RCP45_matrix_s)),
         fontsize_number = 13, number_color = 'darkgreen',
         color = c('darkgreen','lightgray'), border_color = 'black', 
         cellwidth = 50, cellheight = 50,
         symm = FALSE, cluster_rows = FALSE, cluster_cols = FALSE,
         gaps_row = c(3,6),
         legend_breaks = c(0,0.25,0.5,0.75,1), legend_labels = c('','H0','','HA',''),
         labels_col = month_list,
         labels_row = c('Prec actual vs. 2050','Prec 2050 vs. 2070','Prec actual vs. 2070',
                        'Tmin actual vs. 2050','Tmin 2050 vs. 2070','Tmin actual vs. 2070',
                        'Tmax actual vs. 2050','Tmax 2050 vs. 2070','Tmax actual vs. 2070'),
         main = 'RCP4.5', fontsize = 16)

MWU_prec_RCP85_matrix_s <- format(MWU_prec_RCP85_matrix_s, digits=2, nsmall=2)
MWU_tmin_RCP85_matrix_s <- format(MWU_tmin_RCP85_matrix_s, digits=2, nsmall=2)
MWU_tmax_RCP85_matrix_s <- format(MWU_tmax_RCP85_matrix_s, digits=2, nsmall=2)

pheatmap(rbind(t(MWU_prec_RCP85_matrix),
               t(MWU_tmin_RCP85_matrix),
               t(MWU_tmax_RCP85_matrix)), 
         display_numbers = rbind(
               t(MWU_prec_RCP85_matrix_s),
               t(MWU_tmin_RCP85_matrix_s),
               t(MWU_tmax_RCP85_matrix_s)),
         fontsize_number = 13, number_color = 'darkgreen',
         color = c('darkgreen','lightgray'), border_color = 'black', 
         cellwidth = 50, cellheight = 50,
         symm = FALSE, cluster_rows = FALSE, cluster_cols = FALSE,
         gaps_row = c(3,6),
         legend_breaks = c(0,0.25,0.5,0.75,1), legend_labels = c('','H0','','HA',''),
         labels_col = month_list,
         labels_row = c('Prec actual vs. 2050','Prec 2050 vs. 2070','Prec actual vs. 2070',
                        'Tmin actual vs. 2050','Tmin 2050 vs. 2070','Tmin actual vs. 2070',
                        'Tmax actual vs. 2050','Tmax 2050 vs. 2070','Tmax actual vs. 2070'),
         main = 'RCP8.5', fontsize = 16)


```

#### Mann-Whitney U test results for bioclimatic predictors

* H0 - null hypothesis (medians equal)
* HA - alternative hypothesis (medians significantly different)

```{r visualization-5C, fig.height=5, fig.width=20}
MWU_biovars_RCP45_matrix_s <- format(MWU_biovars_RCP45_matrix_s, digits=2, nsmall=2)
MWU_biovars_RCP85_matrix_s <- format(MWU_biovars_RCP85_matrix_s, digits=2, nsmall=2)

pheatmap(t(MWU_biovars_RCP45_matrix), 
         display_numbers = t(MWU_biovars_RCP45_matrix_s),
         fontsize_number = 13, number_color = 'darkgreen',
         color = c('darkgreen','lightgray'), border_color = 'black', 
         cellwidth = 50, cellheight = 50,
         symm = FALSE, cluster_rows = FALSE, cluster_cols = FALSE,
         legend_breaks = c(0,0.25,0.5,0.75,1), legend_labels = c('','H0','','HA',''),
         labels_col = biovars_names,
         labels_row = c('Biovars actual vs. 2050','Biovars 2050 vs. 2070','Biovars actual vs. 2070'),
         main = 'RCP4.5', fontsize = 16)

pheatmap(t(MWU_biovars_RCP85_matrix),
         display_numbers = t(MWU_biovars_RCP85_matrix_s),
         fontsize_number = 13, number_color = 'darkgreen',
         color = c('darkgreen','lightgray'), border_color = 'black', 
         cellwidth = 50, cellheight = 50,
         symm = FALSE, cluster_rows = FALSE, cluster_cols = FALSE,
         legend_breaks = c(0,0.25,0.5,0.75,1), legend_labels = c('','H0','','HA',''),
         labels_col = biovars_names,
         labels_row = c('Biovars actual vs. 2050','Biovars 2050 vs. 2070','Biovars actual vs. 2070'),
         main = 'RCP8.5', fontsize = 16)
```
````
