## Supplementary material for "Vulnerability to climate change for narrowly ranged species: the case of Ecuadorian endemic *Magnolia mercedesiarum*": evaluation_current.html

Script for Shalisko et al. 2018 - Cross-validated SDM script 3


### Script for Shalisko et al. 2018 - Cross-validated SDM script 3

###### *Viacheslav Shalisko*

###### *15 of november 2017*

### *Magnolia mercedesiarum* SDM on current conditions

##### Model evaluation

###### Load modules

```
library(maptools)
```

```
## Loading required package: sp
```

```
## Checking rgeos availability: FALSE
##      Note: when rgeos is not available, polygon geometry     computations in maptools depend on gpclib,
##      which has a restricted licence. It is disabled by default;
##      to enable gpclib, type gpclibPermit()
```

```
library(rworldmap)    # worldmap datasets
```

```
## Warning: package 'rworldmap' was built under R version 3.4.2
```

```
## ### Welcome to rworldmap ###
```

```
## For a short introduction type :   vignette('rworldmap')
```

```
library(rworldxtra)   # hires worldmap spatial dataframe
```

```
## Warning: package 'rworldxtra' was built under R version 3.4.2
```

```
library(sp)
library(raster)
library(dismo)
library(rgdal)
```

```
## rgdal: version: 1.2-11, (SVN revision 676)
##  Geospatial Data Abstraction Library extensions to R successfully loaded
##  Loaded GDAL runtime: GDAL 2.2.0, released 2017/04/28
##  Path to GDAL shared files: C:/Users/vshal/Documents/R/win-library/3.4/rgdal/gdal
##  Loaded PROJ.4 runtime: Rel. 4.9.3, 15 August 2016, [PJ_VERSION: 493]
##  Path to PROJ.4 shared files: C:/Users/vshal/Documents/R/win-library/3.4/rgdal/proj
##  Linking to sp version: 1.2-5
```

```
library(rJava)
```

```
knitr::opts_chunk$set(echo = TRUE)
knitr::opts_chunk$set(error = TRUE)

evaluate.mdl <- function (pnt,bkg,rast,mdl) {
  # pnt - presence points, bkg - background points, rast - raster dataset, mdl - model
  result <- list()
  values_bkg <- extract(rast, bkg)
  values_pnt <- extract(rast, pnt)
  result$v_bkg <- values_bkg
  result$v_pnt <- values_pnt  
  
  values_pnt_bkg <- rbind(values_bkg, values_pnt)

  result$e <- evaluate(values_pnt, values_pnt_bkg, mdl)
  #print(result$e)

  par(mfrow=c(2, 2))
  plot(result$e, 'TPR')
  plot(result$e, 'ROC')
  density(result$e)
  boxplot(result$e, col=c('lightblue','coral'), notch=TRUE)
  
  result$t <- threshold(result$e)
  print(result$t)
  
  # use equal specificity & sensitivity threshold
  result$et <- evaluate(values_pnt, values_pnt_bkg, mdl, tr = result$t$equal_sens_spec)
  print(result$et)

  return(result)
}

maxent.select.contribution <- function (m,n) {
  # parametros: m - objeto del model maxent, n - vector de nombres de variables raster
  resultados <- m@results
  tabla_resultados <- data.frame(t(rep(NA,3)))
  names(tabla_resultados) <- c('variable','contribution','permutation.importance')
  for (i in 1:length(n)) {
    variable1_name <- paste(n[i],'.contribution',sep='')
    variable1_value <- resultados[variable1_name,1]
    variable2_name <- paste(n[i],'.permutation.importance',sep='')
    variable2_value <- resultados[variable2_name,1]
    tabla_resultados <- rbind(tabla_resultados,c(n[i],variable1_value,variable2_value))
  }
  tabla_resultados <- tabla_resultados[-1,]
  return(tabla_resultados)
}

maxent.select.AUC <- function (m) {
  resultados <- m@results
  return(resultados['Training.AUC',1])
}
```

#### Data preparation

###### Set adjustable variables

```
run_code <- '29dup'
basepath <- 'C:/Users/vshal/Downloads/MaxEnt_test_runs/run29_duplicate'
basename <- 'Magnolia_mercedesiarum_MAXENT_climond2_HR_'

scriptpath <- 'C:/Users/vshal/Downloads/MaxEnt_test_runs/run29_duplicate'

random_background_number <- 450
presence_points_count <- 50
double_presence_points_count <- 100

cross_validation_lenght <- 10

#layers_to_drop <- c(2,6,7,9,11,12,14,15,17,18,19,21)
layers_to_drop <- c(21)    # the last layer no. 21 is dummy (repetition of the first, included just to drop it when nothing to drop)
```

###### Load presence mask

```
# load raster of predicted presence
presence_raster_mask <- stack("C:/Users/vshal/GD/Projects_actual/bot_Magnolia_mercedesiarum/Phytotaxa_2017/Analysis/SIG/Rasters_run_22_23/Magnolia_mercedesiarum_ETSS_raster_small.tif")
```

###### Load raster predictor variables

```
names <- c("alt","bio1","bio2","bio3","bio4",
              "bio5","bio6","bio7","bio8","bio9",
              "bio10","bio11","bio12","bio13","bio14",
              "bio15","bio16","bio17","bio18","bio19",
              "dummy")

capas_raster<-stack(c("C:/Users/vshal/Downloads/GMTED2010/GMTED2010_075_Andes.tif",
                      "C:/Users/vshal/Downloads/SDM_highres_Andes/biovars_climond2_Andes_HR.tif",
                      "C:/Users/vshal/Downloads/GMTED2010/GMTED2010_075_Andes.tif"))

names(capas_raster) <- names

capas_raster
```

```
## class       : RasterStack 
## dimensions  : 9600, 5760, 55296000, 21  (nrow, ncol, ncell, nlayers)
## resolution  : 0.002083333, 0.002083333  (x, y)
## extent      : -82.00014, -70.00014, -10.00014, 9.999861  (xmin, xmax, ymin, ymax)
## coord. ref. : +proj=longlat +datum=WGS84 +no_defs +ellps=WGS84 +towgs84=0,0,0 
## names       :         alt,        bio1,        bio2,        bio3,        bio4,        bio5,        bio6,        bio7,        bio8,        bio9,       bio10,       bio11,       bio12,       bio13,       bio14, ... 
## min values  :  -50.000000,  -67.965177,   51.788566,   46.986875,   77.333818,   11.052460, -164.658356,   62.161942,  -56.866343,  -81.849262,  -56.866343,  -81.849262,    1.000000,    1.000000,   -4.000000, ... 
## max values  :  6544.00000,   290.01051,   166.67886,    97.66176,  2813.85039,   353.26230,   240.15173,   195.08090,   294.50851,   297.26141,   297.26141,   282.18530,  8007.00000,   965.00000,   537.00000, ...
```

```
#summary(capas_raster)

bio10 <- subset(capas_raster,10)
```

###### Drop unsignificant raster layers (based on preliminary test run)

```
# drop unsignificant or dummy layers
capas_raster_drop <- dropLayer(capas_raster,layers_to_drop)
names_after_drop <- names(capas_raster_drop)
```

###### Load model run results

```
iterations <- readRDS(paste(scriptpath, "/iterations_object_",run_code,".rds",sep=""))

#str(iterations)
```

#### Model evaluation & store in reusable object

```
iterations_evaluation <- list()
  
  
for (j in 1:cross_validation_lenght) {
  my_obj <- iterations[[j]]

  set.seed(j)
  random_presence_points <- randomPoints(presence_raster_mask, 
                                        n = double_presence_points_count, 
                                        tryf = 10)
  # fold in 2 subsets
  fold <- kfold(random_presence_points, k = 2)
  # set one subset for training
  puntos_entrenamiento_coordenadas <- random_presence_points[ fold == 1, ]
  # set another subset for model evaluation
  puntos_control_coordenadas <- random_presence_points[ fold == 2, ]

  test_points <- rbind(my_obj$entr,puntos_control_coordenadas)  
  evalutaion_new <- evaluate.mdl(test_points,my_obj$bkg,capas_raster_drop,my_obj$model)

  container <- list()
  container$e_new <- evalutaion_new
  iterations_evaluation[[j]] <- container
  
}
```

```
## Warning in bxp(structure(list(stats = structure(c(0,
## 6.49829644316924e-06, : some notches went outside hinges ('box'): maybe set
## notch=FALSE
```

```
##                kappa spec_sens no_omission prevalence equal_sens_spec
## thresholds 0.3194446  0.280841    0.280841 0.08962266       0.4777498
##            sensitivity
## thresholds   0.4994363
## class          : ModelEvaluation 
## n presences    : 100 
## n absences     : 997 
## AUC            : 0.9495988 
## cor            : 0.6456129 
## max TPR+TNR at : 0.4777498
```

```
##                kappa spec_sens no_omission prevalence equal_sens_spec
## thresholds 0.3329513 0.2857366   0.2857366  0.1028378       0.4717498
##            sensitivity
## thresholds   0.4895669
## class          : ModelEvaluation 
## n presences    : 100 
## n absences     : 997 
## AUC            : 0.9486058 
## cor            : 0.6423122 
## max TPR+TNR at : 0.4717498
```

```
##                kappa spec_sens no_omission prevalence equal_sens_spec
## thresholds 0.3641435 0.2226662   0.2226662  0.0921749       0.5180254
##            sensitivity
## thresholds   0.5193262
## class          : ModelEvaluation 
## n presences    : 100 
## n absences     : 999 
## AUC            : 0.9474775 
## cor            : 0.6398904 
## max TPR+TNR at : 0.5180254
```

```
##               kappa spec_sens no_omission prevalence equal_sens_spec
## thresholds 0.223376  0.223376  0.03817012 0.09344866       0.4159884
##            sensitivity
## thresholds   0.4159884
## class          : ModelEvaluation 
## n presences    : 100 
## n absences     : 997 
## AUC            : 0.9482949 
## cor            : 0.6321078 
## max TPR+TNR at : 0.4159884
```

```
##                kappa spec_sens no_omission prevalence equal_sens_spec
## thresholds 0.3126245  0.235911   0.1569168 0.09217318       0.4304767
##            sensitivity
## thresholds   0.4704748
## class          : ModelEvaluation 
## n presences    : 100 
## n absences     : 995 
## AUC            : 0.9492563 
## cor            : 0.6419186 
## max TPR+TNR at : 0.4304767
```

```
## Warning in bxp(structure(list(stats = structure(c(4.81836792687318e-14, :
## some notches went outside hinges ('box'): maybe set notch=FALSE
```

```
##                kappa spec_sens no_omission prevalence equal_sens_spec
## thresholds 0.2325936 0.2325936   0.2325936 0.09042663       0.4204661
##            sensitivity
## thresholds   0.4204661
## class          : ModelEvaluation 
## n presences    : 100 
## n absences     : 998 
## AUC            : 0.9472846 
## cor            : 0.6318832 
## max TPR+TNR at : 0.4204661
```

```
##                kappa spec_sens no_omission prevalence equal_sens_spec
## thresholds 0.3094583 0.3094583  0.09491595 0.09041204       0.4722541
##            sensitivity
## thresholds   0.4806095
## class          : ModelEvaluation 
## n presences    : 100 
## n absences     : 997 
## AUC            : 0.948656 
## cor            : 0.6393436 
## max TPR+TNR at : 0.4722541
```

```
##                kappa spec_sens no_omission prevalence equal_sens_spec
## thresholds 0.4503702  0.227728    0.227728  0.0913518       0.4856447
##            sensitivity
## thresholds   0.4906887
## class          : ModelEvaluation 
## n presences    : 100 
## n absences     : 997 
## AUC            : 0.9492879 
## cor            : 0.6447679 
## max TPR+TNR at : 0.4856447
```

```
##                kappa spec_sens no_omission prevalence equal_sens_spec
## thresholds 0.2912626 0.2912626  0.08720537 0.09170825       0.4036238
##            sensitivity
## thresholds   0.4036238
## class          : ModelEvaluation 
## n presences    : 100 
## n absences     : 999 
## AUC            : 0.9481181 
## cor            : 0.6344718 
## max TPR+TNR at : 0.4036238
```

```
##                kappa spec_sens no_omission prevalence equal_sens_spec
## thresholds 0.3007302 0.1861077   0.1861077 0.09052775       0.3599727
##            sensitivity
## thresholds   0.3599727
## class          : ModelEvaluation 
## n presences    : 100 
## n absences     : 995 
## AUC            : 0.9483317 
## cor            : 0.6327752 
## max TPR+TNR at : 0.3599727
```

```
saveRDS(iterations_evaluation, paste(scriptpath, "/iterations_evaluation_object_",run_code,".rds",sep=""))
```

###### Optional: Check & compare first objects

```
#str(iterations[[1]])
#str(iterations_evaluation[[1]])
```

#### Results of model evaluation

###### Visualization 1

```
contrib_df1 <- data.frame(as.list(c(1:20)))
names(contrib_df1) <- names[1:20]
contrib_df1 <- contrib_df1[FALSE,]
contrib_df2 <- contrib_df1[FALSE,]
auc_vector <- c()
ess_vector <- c()
sen_vector <- c()
sel_vector <- c()
tss_vector <- c()
cor_vector <- c()
auc_null_model_vector <- c()

for (j in 1:cross_validation_lenght) {

  my_obj <- iterations[[j]]
  my_obj2 <- iterations_evaluation[[j]]

  par(mfrow=c(1, 2))  
  
  plot(my_obj$e_train$e, 'ROC')
  plot(my_obj2$e_new$e, 'ROC')
  
  contribuciones <- maxent.select.contribution(my_obj$model,names_after_drop)

  dotchart(as.numeric(contribuciones[,2]),
        col='blue', pch=16,
        labels=contribuciones[,1],
        main='predictor contribution')
  dotchart(as.numeric(contribuciones[,3]),
        col='red', pch=16,
        labels=contribuciones[,1],
        main='predictor importance in permutations')
  
  contrib_temp <- as.data.frame(as.list(contribuciones[,2]), col.names = contribuciones[,1])
  contrib_df1 <- rbind(contrib_df1, contrib_temp)
  permut_temp <- as.data.frame(as.list(contribuciones[,3]), col.names = contribuciones[,1])
  contrib_df2 <- rbind(contrib_df2, permut_temp)
  
  auc_vector <- c(auc_vector, my_obj2$e_new$e@auc)
  ess_vector <- c(ess_vector, my_obj2$e_new$t$equal_sens_spec) 
  cor_vector <- c(cor_vector, my_obj2$e_new$e@cor)
  auc_null_model_vector <- c(auc_null_model_vector, my_obj$auc_null_model_mean)
  
  sensitivity <- my_obj2$e_new$et@TPR / (my_obj2$e_new$et@TPR + my_obj2$e_new$et@FNR)
  specificity <- my_obj2$e_new$et@TNR / (my_obj2$e_new$et@TNR + my_obj2$e_new$et@FPR)
  tss <-  sensitivity + specificity - 1
  
  sen_vector <- c(sen_vector, sensitivity)
  sel_vector <- c(sel_vector, specificity)
  tss_vector <- c(tss_vector, tss)

}
```

###### Visualization 2

```
par(cex = 1.2)

contrib_df1[1:20] <- lapply(contrib_df1[1:20], function(x) as.numeric(as.character(x)))
contrib_df2[1:20] <- lapply(contrib_df2[1:20], function(x) as.numeric(as.character(x)))

summary(contrib_df1)
```

```
##       alt              bio1             bio2               bio3      
##  Min.   : 4.614   Min.   :0.0000   Min.   :0.000000   Min.   :17.62  
##  1st Qu.: 6.906   1st Qu.:0.0000   1st Qu.:0.000525   1st Qu.:18.82  
##  Median : 8.172   Median :0.0000   Median :0.003950   Median :21.79  
##  Mean   : 8.096   Mean   :0.1797   Mean   :0.006490   Mean   :22.03  
##  3rd Qu.: 8.771   3rd Qu.:0.0000   3rd Qu.:0.010725   3rd Qu.:24.38  
##  Max.   :11.377   Max.   :1.7912   Max.   :0.020500   Max.   :28.74  
##       bio4              bio5             bio6            bio7        
##  Min.   :0.00000   Min.   :0.0034   Min.   :19.08   Min.   :0.00000  
##  1st Qu.:0.02077   1st Qu.:0.0842   1st Qu.:22.83   1st Qu.:0.00000  
##  Median :0.04225   Median :0.2291   Median :23.95   Median :0.00000  
##  Mean   :0.14861   Mean   :0.6107   Mean   :23.48   Mean   :0.01857  
##  3rd Qu.:0.09248   3rd Qu.:1.1575   3rd Qu.:24.85   3rd Qu.:0.00000  
##  Max.   :1.02260   Max.   :1.6913   Max.   :26.90   Max.   :0.14170  
##       bio8              bio9             bio10              bio11        
##  Min.   :0.00000   Min.   :0.00000   Min.   :0.000200   Min.   :0.00000  
##  1st Qu.:0.00000   1st Qu.:0.00000   1st Qu.:0.003575   1st Qu.:0.05532  
##  Median :0.00195   Median :0.00595   Median :0.016800   Median :1.49805  
##  Mean   :0.50476   Mean   :0.38483   Mean   :0.927720   Mean   :1.11565  
##  3rd Qu.:1.12252   3rd Qu.:0.46992   3rd Qu.:0.563425   3rd Qu.:1.73237  
##  Max.   :1.50770   Max.   :1.67280   Max.   :6.322700   Max.   :2.60580  
##      bio12            bio13             bio14             bio15       
##  Min.   :0.0440   Min.   :0.00000   Min.   :0.00000   Min.   :0.2335  
##  1st Qu.:0.1176   1st Qu.:0.00000   1st Qu.:0.00455   1st Qu.:1.4535  
##  Median :0.2646   Median :0.00000   Median :0.02360   Median :3.0644  
##  Mean   :0.2990   Mean   :0.00093   Mean   :0.75332   Mean   :2.8859  
##  3rd Qu.:0.4446   3rd Qu.:0.00000   3rd Qu.:1.47480   3rd Qu.:4.2030  
##  Max.   :0.7091   Max.   :0.00930   Max.   :3.25980   Max.   :5.9715  
##      bio16            bio17            bio18            bio19      
##  Min.   :0.0000   Min.   :0.0000   Min.   : 7.110   Min.   :21.32  
##  1st Qu.:0.0015   1st Qu.:0.0000   1st Qu.: 9.936   1st Qu.:24.33  
##  Median :0.0897   Median :0.0015   Median :10.608   Median :27.22  
##  Mean   :0.2639   Mean   :0.7490   Mean   :10.867   Mean   :26.68  
##  3rd Qu.:0.2039   3rd Qu.:1.5251   3rd Qu.:11.951   3rd Qu.:28.26  
##  Max.   :1.1255   Max.   :3.3550   Max.   :15.002   Max.   :31.06
```

```
length(contrib_df1)
```

```
## [1] 20
```

```
summary(contrib_df2)
```

```
##       alt             bio1        bio2              bio3       
##  Min.   :11.33   Min.   :0   Min.   :0.00000   Min.   :0.9668  
##  1st Qu.:13.57   1st Qu.:0   1st Qu.:0.00000   1st Qu.:1.6757  
##  Median :16.60   Median :0   Median :0.00000   Median :3.7159  
##  Mean   :17.30   Mean   :0   Mean   :0.03974   Mean   :3.7696  
##  3rd Qu.:18.18   3rd Qu.:0   3rd Qu.:0.00000   3rd Qu.:5.3362  
##  Max.   :31.25   Max.   :0   Max.   :0.38880   Max.   :8.5654  
##       bio4             bio5              bio6             bio7  
##  Min.   :0.0000   Min.   : 0.0000   Min.   : 0.000   Min.   :0  
##  1st Qu.:0.0143   1st Qu.: 0.7657   1st Qu.: 2.165   1st Qu.:0  
##  Median :0.2180   Median : 3.4265   Median :16.459   Median :0  
##  Mean   :0.7942   Mean   : 6.0877   Mean   :18.475   Mean   :0  
##  3rd Qu.:1.2798   3rd Qu.: 5.4751   3rd Qu.:27.892   3rd Qu.:0  
##  Max.   :2.9463   Max.   :20.9053   Max.   :47.986   Max.   :0  
##       bio8              bio9             bio10              bio11       
##  Min.   :0.00000   Min.   :0.00000   Min.   : 0.00000   Min.   :0.0000  
##  1st Qu.:0.00000   1st Qu.:0.00000   1st Qu.: 0.02852   1st Qu.:0.0000  
##  Median :0.00000   Median :0.00000   Median : 1.04145   Median :0.0000  
##  Mean   :0.02866   Mean   :0.22643   Mean   : 7.45167   Mean   :0.1366  
##  3rd Qu.:0.00000   3rd Qu.:0.02485   3rd Qu.: 5.39900   3rd Qu.:0.0000  
##  Max.   :0.28660   Max.   :1.84600   Max.   :32.13670   Max.   :1.3657  
##      bio12            bio13       bio14             bio15       
##  Min.   : 1.120   Min.   :0   Min.   :0.00000   Min.   : 0.962  
##  1st Qu.: 6.911   1st Qu.:0   1st Qu.:0.00000   1st Qu.: 2.200  
##  Median :12.092   Median :0   Median :0.00000   Median : 4.984  
##  Mean   :12.936   Mean   :0   Mean   :0.03311   Mean   : 5.305  
##  3rd Qu.:17.446   3rd Qu.:0   3rd Qu.:0.05805   3rd Qu.: 6.403  
##  Max.   :32.115   Max.   :0   Max.   :0.13730   Max.   :16.583  
##      bio16            bio17             bio18           bio19        
##  Min.   :0.0000   Min.   :0.00000   Min.   :16.18   Min.   :0.00000  
##  1st Qu.:0.0000   1st Qu.:0.00000   1st Qu.:20.59   1st Qu.:0.00000  
##  Median :0.0000   Median :0.00000   Median :21.94   Median :0.05975  
##  Mean   :0.2485   Mean   :0.05093   Mean   :24.67   Mean   :2.44927  
##  3rd Qu.:0.0000   3rd Qu.:0.04448   3rd Qu.:29.09   3rd Qu.:4.02485  
##  Max.   :1.9715   Max.   :0.37890   Max.   :40.24   Max.   :9.03190
```

```
mean.contrib_df1 <- apply(contrib_df1, 2, mean)
sd.contrib_df1 <- apply(contrib_df1, 2, sd)
na.contrib_df1 <- contrib_df1
na.contrib_df1[na.contrib_df1 == 0] <- NA
c.contrib_df1 <- apply(na.contrib_df1, 2, function(x){sum(!is.na(x))})
se.contrib_df1 <- sd.contrib_df1 / sqrt(c.contrib_df1)

c.contrib_df2 <- apply(na.contrib_df1, 1, function(x){sum(!is.na(x))})
c.contrib_df2
```

```
##  [1] 14 17 15 15 14 15 14 16 13 16
```

```
plot(c(1:20), mean.contrib_df1, col = "black", pch = 19, cex = 1.2,
     ylim = c(0,40), xlim = c(0.5, 20.5), axes = FALSE,
     ylab = 'contribution', xlab = '')
axis(2)
axis(1, at = c(1:20), labels = colnames(contrib_df1), las = 2)
box(lty = "solid")
arrows(c(1:20), mean.contrib_df1 - 1.96 * se.contrib_df1, 
       c(1:20), mean.contrib_df1 + 1.96 * se.contrib_df1,
       code = 3, col = "black", angle = 90, length = .1)
```

```
barplot(c.contrib_df1, ylab = 'generality in contributions', las = 2)
```

```
mean.contrib_df2 <- apply(contrib_df2, 2, mean)
sd.contrib_df2 <- apply(contrib_df2, 2, sd)
na.contrib_df2 <- contrib_df2
na.contrib_df2[na.contrib_df2 == 0] <- NA
c.contrib_df2 <- apply(na.contrib_df2, 2, function(x){sum(!is.na(x))})
se.contrib_df2 <- sd.contrib_df2 / sqrt(c.contrib_df2)

plot(c(1:20), mean.contrib_df2, col = "black", pch = 19, cex = 1.2,
     ylim = c(0,40), xlim = c(0.5, 20.5), axes = FALSE,
     ylab = 'importance in permutations', xlab = '')
axis(2)
axis(1, at = c(1:20), labels = colnames(contrib_df2), las = 2)
box(lty = "solid")
arrows(c(1:20), mean.contrib_df2 - 1.96 * se.contrib_df2, 
       c(1:20), mean.contrib_df2 + 1.96 * se.contrib_df2,
       code = 3, col = "black", angle = 90, length = .1)
```

```
barplot(c.contrib_df2, ylab = 'generality in permutations', las = 2)
```

```
boxplot(contrib_df2, ylim = c(0,60), xlim = c(0.5, 20.5), 
        names = colnames(contrib_df1), las = 3,
        boxfill=rgb(1, 1, 1, alpha=1), border=rgb(1, 1, 1, alpha=1),
        ylab = '%')
boxplot(contrib_df1, at = 1:20 - 0.15, boxwex=0.25, xaxt = "n", add = TRUE, boxfill = 'darkgreen')
boxplot(contrib_df2, at = 1:20 + 0.15, boxwex=0.25, xaxt = "n", add = TRUE, boxfill = 'lightgray')

legend("topleft", c("Contributions","Importance in permutations"), 
       inset=.02, box.lty=0, fill=c('darkgreen','lightgray'), horiz=TRUE, cex=1.2)
```

###### Visualization 3

```
par(mfrow=c(2, 2), mar = c(4, 4, 0, 2) + 0.1, cex = 1.1)

plot(c(1:20), mean.contrib_df1, col = "black", pch = 19, cex = 1.2,
     ylim = c(0,40), xlim = c(0.5, 20.5), axes = FALSE,
     ylab = 'contribution, %', xlab = '')
axis(2)
axis(1, at = c(1:20), labels = colnames(contrib_df1), las = 2)
box(lty = "solid")
arrows(c(1:20), mean.contrib_df1 - 1.96 * se.contrib_df1, 
       c(1:20), mean.contrib_df1 + 1.96 * se.contrib_df1,
       code = 3, col = "black", angle = 90, length = .1)
```

```
## Warning in arrows(c(1:20), mean.contrib_df1 - 1.96 * se.contrib_df1,
## c(1:20), : zero-length arrow is of indeterminate angle and so skipped

## Warning in arrows(c(1:20), mean.contrib_df1 - 1.96 * se.contrib_df1,
## c(1:20), : zero-length arrow is of indeterminate angle and so skipped
```

```
mean.contrib_df2 <- apply(contrib_df2, 2, mean)
sd.contrib_df2 <- apply(contrib_df2, 2, sd)
na.contrib_df2 <- contrib_df2
na.contrib_df2[na.contrib_df2 == 0] <- NA
c.contrib_df2 <- apply(na.contrib_df2, 2, function(x){sum(!is.na(x))})
se.contrib_df2 <- sd.contrib_df2 / sqrt(c.contrib_df2)

plot(c(1:20), mean.contrib_df2, col = "black", pch = 19, cex = 1.2,
     ylim = c(0,40), xlim = c(0.5, 20.5), axes = FALSE,
     ylab = 'importance in permutations, %', xlab = '')
axis(2)
axis(1, at = c(1:20), labels = colnames(contrib_df2), las = 2)
box(lty = "solid")
arrows(c(1:20), mean.contrib_df2 - 1.96 * se.contrib_df2, 
       c(1:20), mean.contrib_df2 + 1.96 * se.contrib_df2,
       code = 3, col = "black", angle = 90, length = .1)

barplot(c.contrib_df1, ylab = 'generality in contributions', 
        las = 2, names.arg = rep("",20))

barplot(c.contrib_df2, ylab = 'generality in permutations', 
        las = 2, names.arg = rep("",20))
```

###### Visualization 4

```
pearson_vector <- c()
spearman_vector <- c()

for (j in 1:cross_validation_lenght) {
   # realculate correlation coefficients
   # probability vales given in evaluate object
   p <- iterations_evaluation[[j]]$e_new$e@presence
   a <- iterations_evaluation[[j]]$e_new$e@absence

  # Pearson correlation (cor)
    crp <- cor.test(c(p,a), c(rep(1, length(p)), rep(0, length(a))))
    pearson_vector <- c(pearson_vector, crp$estimate)
    print(crp$estimate)
  print(crp$p.value)
  
  # Spearman correlation (rho)
  crs <- cor.test(c(p,a), c(rep(1, length(p)), rep(0, length(a))), method = "spearman", exact = FALSE)
    spearman_vector <- c(spearman_vector, crs$estimate)
    print(crs$estimate)
  print(crs$p.value)
}
```

```
##       cor 
## 0.6456129 
## [1] 2.23948e-130
##       rho 
## 0.4482879 
## [1] 2.424818e-55
##       cor 
## 0.6423122 
## [1] 1.200512e-128
##       rho 
## 0.4472978 
## [1] 4.460156e-55
##       cor 
## 0.6398904 
## [1] 1.274777e-127
##       rho 
## 0.4458073 
## [1] 8.902953e-55
##       cor 
## 0.6321078 
## [1] 1.962241e-123
##       rho 
## 0.4469878 
## [1] 5.395685e-55
##       cor 
## 0.6419186 
## [1] 3.274753e-128
##       rho 
## 0.4483142 
## [1] 2.988648e-55
##       cor 
## 0.6318832 
## [1] 1.969904e-123
##       rho 
## 0.4457976 
## [1] 1.000935e-54
##       cor 
## 0.6393436 
## [1] 4.134401e-127
##       rho 
## 0.4473478 
## [1] 4.325176e-55
##       cor 
## 0.6447679 
## [1] 6.23595e-130
##       rho 
## 0.4479778 
## [1] 2.935289e-55
##       cor 
## 0.6344718 
## [1] 7.556915e-125
##       rho 
## 0.4464455 
## [1] 6.019408e-55
##       cor 
## 0.6327752 
## [1] 1.514592e-123
##       rho 
## 0.4473915 
## [1] 5.268987e-55
```

```
summary(auc_vector)
```

```
##    Min. 1st Qu.  Median    Mean 3rd Qu.    Max. 
##  0.9473  0.9482  0.9485  0.9485  0.9491  0.9496
```

```
summary(auc_null_model_vector)
```

```
##    Min. 1st Qu.  Median    Mean 3rd Qu.    Max. 
##  0.7202  0.7225  0.7262  0.7249  0.7266  0.7286
```

```
summary(sen_vector)
```

```
##    Min. 1st Qu.  Median    Mean 3rd Qu.    Max. 
##   0.900   0.900   0.910   0.906   0.910   0.910
```

```
summary(sel_vector)
```

```
##    Min. 1st Qu.  Median    Mean 3rd Qu.    Max. 
##  0.9015  0.9048  0.9053  0.9051  0.9067  0.9077
```

```
summary(tss_vector)
```

```
##    Min. 1st Qu.  Median    Mean 3rd Qu.    Max. 
##  0.8015  0.8048  0.8153  0.8111  0.8167  0.8177
```

```
summary(ess_vector)
```

```
##    Min. 1st Qu.  Median    Mean 3rd Qu.    Max. 
##  0.3600  0.4171  0.4511  0.4456  0.4764  0.5180
```

```
summary(cor_vector)
```

```
##    Min. 1st Qu.  Median    Mean 3rd Qu.    Max. 
##  0.6319  0.6332  0.6396  0.6385  0.6422  0.6456
```

```
summary(pearson_vector)
```

```
##    Min. 1st Qu.  Median    Mean 3rd Qu.    Max. 
##  0.6319  0.6332  0.6396  0.6385  0.6422  0.6456
```

```
summary(spearman_vector)
```

```
##    Min. 1st Qu.  Median    Mean 3rd Qu.    Max. 
##  0.4458  0.4466  0.4473  0.4472  0.4478  0.4483
```

```
par(mfrow=c(1, 4), cex = 1.1)

boxplot(auc_vector, boxfill = 'lightgray', ylim = c(0.5,1), xlab="True AUC")
boxplot(auc_null_model_vector, boxfill = 'lightgray', ylim = c(0.5,1), xlab="Null model AUC")
boxplot(pearson_vector, boxfill = 'lightgray', ylim = c(0,1), xlab="Pearson's correlation")
boxplot(spearman_vector, boxfill = 'lightgray', ylim = c(0,1), xlab="Spearman's correlation")
```

```
#boxplot(sel_vector, boxfill = 'lightgray', ylim = c(0.75,1), xlab="Specificity")

layout(matrix(c(1,1,2),1,3,byrow = TRUE))
par(cex = 1.2)

boxplot(cbind(auc_vector,sen_vector,sel_vector,tss_vector),
        boxfill = 'lightgray', ylim = c(0.78,0.97),
        names=c("True AUC","Sensitivity","Specificity","TSS"))

boxplot(cbind(ess_vector), boxfill = 'lightgray', ylim = c(0,1), 
        names=c("ESS threshold"),show.names=TRUE)
```

###### Rank model AUC comparing to 99 null-models

```
null_model_auc_vector <- c()
null_model_rank_vector <- c()

for (j in 1:cross_validation_lenght) {

  my_obj2 <- iterations_evaluation[[j]]
  valores_puntos_aleatorios_fondo_y_entrenamiento <- rbind(my_obj2$e_new$v_pnt, my_obj2$e_new$v_bkg)

  rep <- 99     # number or null models to produce

  null_models <- list()
  for (r in 1:rep) {
    # following Robert J. Hijmans nullRandom() function from 'dismo'
        # sample presence records and set the rest of records as absence
        index <- sample(nrow(valores_puntos_aleatorios_fondo_y_entrenamiento), presence_points_count)
        pres <- valores_puntos_aleatorios_fondo_y_entrenamiento[index, ]
        absc <- valores_puntos_aleatorios_fondo_y_entrenamiento[-index, ]
        d <- data.frame(rbind(pres, absc))
        v <- c(rep(1, nrow(pres)), rep(0, nrow(absc)))
        # make null-model with presence and absence sets
        m <- maxent(d, v)
        # evaluate the null-model
        null_models[[r]] <- evaluate(pres, absc, m)
  }

  #null_models
  auc_list_vector <- sapply(null_models, function(x) x@auc)

  null_model_auc_vector <- c(null_model_auc_vector, auc_list_vector)  
  
  #auc_list_vector
  #mean(auc_list_vector)

  auc_list_real_vector <- c(my_obj2$e_new$e@auc, auc_list_vector)
  auc_list_real_vector_ranks <- rank(auc_list_real_vector, ties.method= "max")

  #print('Rank of real AUC vs null-models')
  #auc_list_real_vector_ranks[1]
  null_model_rank_vector <- c(null_model_rank_vector, auc_list_real_vector_ranks[1])   
}
```

###### Visualization 5

```
#null_model_rank_vector
summary(null_model_auc_vector)
```

```
##    Min. 1st Qu.  Median    Mean 3rd Qu.    Max. 
##  0.6268  0.7009  0.7192  0.7208  0.7413  0.8147
```

```
layout(matrix(c(1,1,2,2,3,3,3),1,7,byrow = TRUE))
par(cex = 1.0)

boxplot(cbind(auc_vector,null_model_auc_vector),
        boxfill = 'lightgray', ylim = c(0.5,1),
        names=c("True AUC","Null-model"))

boxplot(cbind(ess_vector), boxfill = 'lightgray', ylim = c(0,1), 
        names=c("ESS threshold"),show.names=TRUE)

boxplot(cbind(sen_vector,sel_vector,tss_vector),
        boxfill = 'lightgray', ylim = c(0.78,0.93),
        names=c("Sensitivity","Specificity","TSS"))
```
