## Supplementary material for "Vulnerability to climate change for narrowly ranged species: the case of Ecuadorian endemic *Magnolia mercedesiarum*": integrator.html

Integrator (run 29 duplicate)


### Script for Shalisko et al. 2018 - Cross-validated SDM script 1

###### *Viacheslav Shalisko*

###### *22 of october 2017*

### *Magnolia mercedesiarum* SDM on current conditions

##### Model production

##### Note: run 29 duplicate

##### Parameters: MaxEnt model, high resolution predictors (7.5“) CliMond2 current conditions, 50 simulated presence points (from presence mask), 900 random background ponts (half within 500 km distance)

```
knitr::opts_chunk$set(echo = TRUE)
knitr::opts_chunk$set(error = TRUE)

library(maptools)
```

```
## Loading required package: sp
```

```
## Checking rgeos availability: FALSE
##      Note: when rgeos is not available, polygon geometry     computations in maptools depend on gpclib,
##      which has a restricted licence. It is disabled by default;
##      to enable gpclib, type gpclibPermit()
```

```
library(rworldmap)    # worldmap datasets
```

```
## Warning: package 'rworldmap' was built under R version 3.4.2
```

```
## ### Welcome to rworldmap ###
```

```
## For a short introduction type :   vignette('rworldmap')
```

```
library(rworldxtra)   # hires worldmap spatial dataframe
```

```
## Warning: package 'rworldxtra' was built under R version 3.4.2
```

```
library(sp)
library(raster)
library(dismo)
library(rgdal)
```

```
## rgdal: version: 1.2-11, (SVN revision 676)
##  Geospatial Data Abstraction Library extensions to R successfully loaded
##  Loaded GDAL runtime: GDAL 2.2.0, released 2017/04/28
##  Path to GDAL shared files: C:/Users/vshal/Documents/R/win-library/3.4/rgdal/gdal
##  Loaded PROJ.4 runtime: Rel. 4.9.3, 15 August 2016, [PJ_VERSION: 493]
##  Path to PROJ.4 shared files: C:/Users/vshal/Documents/R/win-library/3.4/rgdal/proj
##  Linking to sp version: 1.2-5
```

```
library(rJava)
```

```
maxent.select.contribution <- function (m,n) {
  # parametros: m - objeto del model maxent, n - vector de nombres de variables raster
  resultados <- m@results
  tabla_resultados <- data.frame(t(rep(NA,3)))
  names(tabla_resultados) <- c('variable','contribution','permutation.importance')
  for (i in 1:length(n)) {
    variable1_name <- paste(n[i],'.contribution',sep='')
    variable1_value <- resultados[variable1_name,1]
    variable2_name <- paste(n[i],'.permutation.importance',sep='')
    variable2_value <- resultados[variable2_name,1]
    tabla_resultados <- rbind(tabla_resultados,c(n[i],variable1_value,variable2_value))
  }
  tabla_resultados <- tabla_resultados[-1,]
  return(tabla_resultados)
}


evaluate.mdl <- function (pnt,bkg,rast,mdl) {
  # pnt - presence points, bkg - background points, rast - raster dataset, mdl - model
  result <- list()
  values_bkg <- extract(rast, bkg)
  values_pnt <- extract(rast, pnt)
  values_pnt_bkg <- rbind(values_bkg, values_pnt)

  result$e <- evaluate(values_pnt, values_pnt_bkg, mdl)
  print(result$e)

  par(mfrow=c(2, 2))
  plot(result$e, 'TPR')
  plot(result$e, 'ROC')
  density(result$e)
  boxplot(result$e, col=c('lightblue','coral'), notch=TRUE)
  
  result$t <- threshold(result$e)
  print(result$t)
  
  result$v_bkg <- values_bkg
  result$v_pnt <- values_pnt
  
  return(result)
}
```

#### Data preparation

###### Set adjustable variables

```
run_code <- '29dup'
basepath <- 'C:/Users/vshal/Downloads/MaxEnt_test_runs/run29_duplicate'
basename <- 'Magnolia_mercedesiarum_MAXENT_climond2_HR_'

scriptpath <- 'C:/Users/vshal/Downloads/MaxEnt_test_runs/run29_duplicate'

random_background_number <- 450
presence_points_count <- 50
double_presence_points_count <- 100

cross_validation_lenght <- 10

#layers_to_drop <- c(2,6,7,9,11,12,14,15,17,18,19,21)
layers_to_drop <- c(21)    # the last layer no. 21 is dummy (repetition of the first, included just to drop it when nothing to drop)
```

###### Load raster predictor variables

```
names <- c("alt","bio1","bio2","bio3","bio4",
              "bio5","bio6","bio7","bio8","bio9",
              "bio10","bio11","bio12","bio13","bio14",
              "bio15","bio16","bio17","bio18","bio19",
              "dummy")

capas_raster<-stack(c("C:/Users/vshal/Downloads/GMTED2010/GMTED2010_075_Andes.tif",
                      "C:/Users/vshal/Downloads/SDM_highres_Andes/biovars_climond2_Andes_HR.tif",
                      "C:/Users/vshal/Downloads/GMTED2010/GMTED2010_075_Andes.tif"))

names(capas_raster) <- names

capas_raster
```

```
## class       : RasterStack 
## dimensions  : 9600, 5760, 55296000, 21  (nrow, ncol, ncell, nlayers)
## resolution  : 0.002083333, 0.002083333  (x, y)
## extent      : -82.00014, -70.00014, -10.00014, 9.999861  (xmin, xmax, ymin, ymax)
## coord. ref. : +proj=longlat +datum=WGS84 +no_defs +ellps=WGS84 +towgs84=0,0,0 
## names       :         alt,        bio1,        bio2,        bio3,        bio4,        bio5,        bio6,        bio7,        bio8,        bio9,       bio10,       bio11,       bio12,       bio13,       bio14, ... 
## min values  :  -50.000000,  -67.965177,   51.788566,   46.986875,   77.333818,   11.052460, -164.658356,   62.161942,  -56.866343,  -81.849262,  -56.866343,  -81.849262,    1.000000,    1.000000,   -4.000000, ... 
## max values  :  6544.00000,   290.01051,   166.67886,    97.66176,  2813.85039,   353.26230,   240.15173,   195.08090,   294.50851,   297.26141,   297.26141,   282.18530,  8007.00000,   965.00000,   537.00000, ...
```

```
#summary(capas_raster)

bio10 <- subset(capas_raster,10)
```

###### Load TGS raster

```
# load raster of TGS mask in full extent
tgs_raster_full <- stack("C:/Users/vshal/GD/Projects_actual/bot_Magnolia_mercedesiarum/Phytotaxa_2017/Analysis/SIG/Rasters_complementary/NWSouthamerica.tif")
# load raster of TGS mask buffered with 500 km
tgs_raster_500 <- stack("C:/Users/vshal/GD/Projects_actual/bot_Magnolia_mercedesiarum/Phytotaxa_2017/Analysis/SIG/Rasters_complementary/Magnolia_mercedesiarum_logistic_raster_clip.tif")
# load raster of TGS mask buffered with 750 km
#tgs_raster_750 <- stack("C:/Users/vshal/GD/Projects_actual/bot_Magnolia_mercedesiarum/Phytotaxa_2017/Analysis/SIG/Rasters_TGS/GBif_magnoliopsida_KD_Andes_5km_2only_MM_B750.tif")

tgs_raster_full
```

```
## class       : RasterStack 
## dimensions  : 9600, 5760, 55296000, 1  (nrow, ncol, ncell, nlayers)
## resolution  : 0.002083333, 0.002083333  (x, y)
## extent      : -82.00014, -70.00014, -10.00014, 9.999861  (xmin, xmax, ymin, ymax)
## coord. ref. : +proj=longlat +datum=WGS84 +no_defs +ellps=WGS84 +towgs84=0,0,0 
## names       : NWSouthamerica 
## min values  :              1 
## max values  :              1
```

```
#tgs_raster_750
tgs_raster_500
```

```
## class       : RasterStack 
## dimensions  : 4762, 4073, 19395626, 1  (nrow, ncol, ncell, nlayers)
## resolution  : 0.002083333, 0.002083333  (x, y)
## extent      : -81.51681, -73.03139, -5.302222, 4.618611  (xmin, xmax, ymin, ymax)
## coord. ref. : +proj=longlat +datum=WGS84 +no_defs +ellps=WGS84 +towgs84=0,0,0 
## names       : Magnolia_mercedesiarum_logistic_raster_clip 
## min values  :                                           1 
## max values  :                                           1
```

```
par(mfrow=c(1, 2))
world_high <- getMap(resolution = "high")

plot(tgs_raster_500, main="TGS 500 km", legend=FALSE)
#plot(tgs_raster_500, legend.only=TRUE)
plot(world_high, add=TRUE)

#plot(tgs_raster_750, main="TGS 750 km", legend=FALSE)
#plot(world_high, add=TRUE)

plot(tgs_raster_full, main="TGS full extent", legend=FALSE)
plot(world_high, add=TRUE)
```

###### Load presence data: 1) main dataset, 2) additional fixed control dataset

```
# load main presence dataset
puntos_entrenamiento_preliminar<- read.csv("C:/Users/vshal/GD/Projects_actual/bot_Magnolia_mercedesiarum/Phytotaxa_2017/Analysis/Magnolia_mercedesiarum_extra.csv")
dim(puntos_entrenamiento_preliminar)
```

```
## [1] 68  3
```

```
head(puntos_entrenamiento_preliminar)
```

```
##                ï..Specie       Lat      Lon
## 1 Magnolia_mercedesiarum -0.620767 -77.6281
## 2 Magnolia_mercedesiarum -0.619102 -77.6282
## 3 Magnolia_mercedesiarum -0.622135 -77.6280
## 4 Magnolia_mercedesiarum -0.620767 -77.6298
## 5 Magnolia_mercedesiarum -0.620767 -77.6263
## 6 Magnolia_mercedesiarum -0.619161 -77.6264
```

```
tail(puntos_entrenamiento_preliminar)
```

```
##                 ï..Specie       Lat      Lon
## 63 Magnolia_mercedesiarum -0.185026 -77.6415
## 64 Magnolia_mercedesiarum -0.186682 -77.6436
## 65 Magnolia_mercedesiarum -0.182913 -77.6453
## 66 Magnolia_mercedesiarum -0.178859 -77.6437
## 67 Magnolia_mercedesiarum -0.186697 -77.6399
## 68                        -0.635844 -77.8384
```

```
#attach(puntos_entrenamiento)
puntos_entrenamiento_coordenadas_preliminar<-data.frame(cbind(
                              puntos_entrenamiento_preliminar[,'Lon'],
                              puntos_entrenamiento_preliminar[,'Lat']))


# load additional fixed control points
puntos_control<- read.csv("C:/Users/vshal/GD/Projects_actual/bot_Magnolia_mercedesiarum/Phytotaxa_2017/Analysis/Magnolia_mercedesiarum.csv")
dim(puntos_control)
```

```
## [1] 4 7
```

```
head(puntos_control)
```

```
##                   Specie         Lat       Lon               Lat_Lon_text
## 1 Magnolia_mercedesiarum  0.01194444 -77.53611           00°03'N 077°35'W
## 2 Magnolia_mercedesiarum -0.63584354 -77.83844           00°38'S 077°50'W
## 3 Magnolia_mercedesiarum -0.62083333 -77.62778           00°36'S 077°35'W
## 4 Magnolia_mercedesiarum -0.18277778 -77.64333 0°10'58.00"S 77°38'36.00"O
##   Num_Colecta                      Colector               Lat_lon_mios
## 1        9896 Cerón Martínez Carlos Eduardo   00°00'43''N 077°32'10''W
## 2        3135               Homeier Juergen   00°37'05''S 077°50'14''W
## 3        1869               Homeier Juergen   00°37'15''S 077°37'40''W
## 4         s.n                       Vazquez 0°10'58.00"S 77°38'36.00"O
```

```
tail(puntos_control)
```

```
##                   Specie         Lat       Lon               Lat_Lon_text
## 1 Magnolia_mercedesiarum  0.01194444 -77.53611           00°03'N 077°35'W
## 2 Magnolia_mercedesiarum -0.63584354 -77.83844           00°38'S 077°50'W
## 3 Magnolia_mercedesiarum -0.62083333 -77.62778           00°36'S 077°35'W
## 4 Magnolia_mercedesiarum -0.18277778 -77.64333 0°10'58.00"S 77°38'36.00"O
##   Num_Colecta                      Colector               Lat_lon_mios
## 1        9896 Cerón Martínez Carlos Eduardo   00°00'43''N 077°32'10''W
## 2        3135               Homeier Juergen   00°37'05''S 077°50'14''W
## 3        1869               Homeier Juergen   00°37'15''S 077°37'40''W
## 4         s.n                       Vazquez 0°10'58.00"S 77°38'36.00"O
```

```
#attach(puntos_control)
puntos_control_extra_coordenadas<-data.frame(cbind(
                            puntos_control[,'Lon'],
                            puntos_control[,'Lat']))

# independent fixd control dataset
puntos_control_extra_coordenadas <- rbind(puntos_control_extra_coordenadas,puntos_entrenamiento_coordenadas_preliminar)

# load raster of predicted presence
presence_raster_mask <- stack("C:/Users/vshal/GD/Projects_actual/bot_Magnolia_mercedesiarum/Phytotaxa_2017/Analysis/SIG/Rasters_run_22_23/Magnolia_mercedesiarum_ETSS_raster_small.tif")
```

#### Cross-validated SDM (render iterations)

```
iterations <- list()

for (i in 1:cross_validation_lenght){
  random_seed_folding <- i
  random_seed_background <- i
  
  rmarkdown::render(paste(scriptpath,'/renderer.Rmd',sep=""), 
                   output_file =  paste("step_", run_code, '_', i, ".html", sep=''), 
                   output_dir = scriptpath)
}
```

```
## 
## 
## processing file: renderer.Rmd
```

```
## 
  |                                                                       
  |                                                                 |   0%
  |                                                                       
  |..                                                               |   3%
##    inline R code fragments
## 
## 
  |                                                                       
  |....                                                             |   7%
## label: set-knitr
## 
  |                                                                       
  |.......                                                          |  10%
##   ordinary text without R code
## 
## 
  |                                                                       
  |.........                                                        |  14%
## label: fold-presence-data
## 
  |                                                                       
  |...........                                                      |  17%
##   ordinary text without R code
## 
## 
  |                                                                       
  |.............                                                    |  21%
## label: background-points
## 
  |                                                                       
  |................                                                 |  24%
##   ordinary text without R code
## 
## 
  |                                                                       
  |..................                                               |  28%
## label: data-visualization (with options) 
## List of 2
##  $ fig.height: num 16
##  $ fig.width : num 10
```

```
## 
  |                                                                       
  |....................                                             |  31%
##   ordinary text without R code
## 
## 
  |                                                                       
  |......................                                           |  34%
## label: drop-layers
## 
  |                                                                       
  |.........................                                        |  38%
##   ordinary text without R code
## 
## 
  |                                                                       
  |...........................                                      |  41%
## label: ajustar-modelo (with options) 
## List of 2
##  $ fig.height: num 10
##  $ fig.width : num 10
## 
## 
  |                                                                       
  |.............................                                    |  45%
##   ordinary text without R code
## 
## 
  |                                                                       
  |...............................                                  |  48%
## label: variable-importance (with options) 
## List of 2
##  $ fig.height: num 7
##  $ fig.width : num 10
```

```
## 
  |                                                                       
  |..................................                               |  52%
##   ordinary text without R code
## 
## 
  |                                                                       
  |....................................                             |  55%
## label: curvas-respuesta (with options) 
## List of 2
##  $ fig.height: num 10
##  $ fig.width : num 10
```

```
## 
  |                                                                       
  |......................................                           |  59%
##   ordinary text without R code
## 
## 
  |                                                                       
  |........................................                         |  62%
## label: revisar-datos-maxent-en-html
## 
  |                                                                       
  |...........................................                      |  66%
##   ordinary text without R code
## 
## 
  |                                                                       
  |.............................................                    |  69%
## label: evaluate-model-training-data (with options) 
## List of 2
##  $ fig.width : num 10
##  $ fig.height: num 10
```

```
## 
  |                                                                       
  |...............................................                  |  72%
##   ordinary text without R code
## 
## 
  |                                                                       
  |.................................................                |  76%
## label: evaluate-model-control-data (with options) 
## List of 2
##  $ fig.width : num 10
##  $ fig.height: num 10
```

```
## 
  |                                                                       
  |....................................................             |  79%
##   ordinary text without R code
## 
## 
  |                                                                       
  |......................................................           |  83%
## label: evaluate-model-control-data-2 (with options) 
## List of 2
##  $ fig.width : num 10
##  $ fig.height: num 10
```

```
## 
  |                                                                       
  |........................................................         |  86%
##   ordinary text without R code
## 
## 
  |                                                                       
  |..........................................................       |  90%
## label: null-model-auc-training
## 
  |                                                                       
  |.............................................................    |  93%
##   ordinary text without R code
## 
## 
  |                                                                       
  |...............................................................  |  97%
## label: store-result
## 
  |                                                                       
  |.................................................................| 100%
##   ordinary text without R code
```

```
## output file: renderer.knit.md
```

```
## "C:/Program Files/RStudio/bin/pandoc/pandoc" +RTS -K512m -RTS renderer.utf8.md --to html --from markdown+autolink_bare_uris+ascii_identifiers+tex_math_single_backslash --output pandoc6e07c652757.html --smart --email-obfuscation none --self-contained --standalone --section-divs --template "C:\Users\vshal\Documents\R\win-library\3.4\rmarkdown\rmd\h\default.html" --no-highlight --variable highlightjs=1 --variable "theme:bootstrap" --include-in-header "C:\Users\vshal\AppData\Local\Temp\RtmpWiYTWp\rmarkdown-str6e028fd2327.html" --mathjax --variable "mathjax-url:https://mathjax.rstudio.com/latest/MathJax.js?config=TeX-AMS-MML_HTMLorMML"
```

```
## 
## Output created: step_29dup_1.html
## 
## 
## processing file: renderer.Rmd
```

```
## 
  |                                                                       
  |                                                                 |   0%
  |                                                                       
  |..                                                               |   3%
##    inline R code fragments
## 
## 
  |                                                                       
  |....                                                             |   7%
## label: set-knitr
## 
  |                                                                       
  |.......                                                          |  10%
##   ordinary text without R code
## 
## 
  |                                                                       
  |.........                                                        |  14%
## label: fold-presence-data
## 
  |                                                                       
  |...........                                                      |  17%
##   ordinary text without R code
## 
## 
  |                                                                       
  |.............                                                    |  21%
## label: background-points
## 
  |                                                                       
  |................                                                 |  24%
##   ordinary text without R code
## 
## 
  |                                                                       
  |..................                                               |  28%
## label: data-visualization (with options) 
## List of 2
##  $ fig.height: num 16
##  $ fig.width : num 10
```

```
## 
  |                                                                       
  |....................                                             |  31%
##   ordinary text without R code
## 
## 
  |                                                                       
  |......................                                           |  34%
## label: drop-layers
## 
  |                                                                       
  |.........................                                        |  38%
##   ordinary text without R code
## 
## 
  |                                                                       
  |...........................                                      |  41%
## label: ajustar-modelo (with options) 
## List of 2
##  $ fig.height: num 10
##  $ fig.width : num 10
## 
## 
  |                                                                       
  |.............................                                    |  45%
##   ordinary text without R code
## 
## 
  |                                                                       
  |...............................                                  |  48%
## label: variable-importance (with options) 
## List of 2
##  $ fig.height: num 7
##  $ fig.width : num 10
```

```
## 
  |                                                                       
  |..................................                               |  52%
##   ordinary text without R code
## 
## 
  |                                                                       
  |....................................                             |  55%
## label: curvas-respuesta (with options) 
## List of 2
##  $ fig.height: num 10
##  $ fig.width : num 10
```

```
## 
  |                                                                       
  |......................................                           |  59%
##   ordinary text without R code
## 
## 
  |                                                                       
  |........................................                         |  62%
## label: revisar-datos-maxent-en-html
## 
  |                                                                       
  |...........................................                      |  66%
##   ordinary text without R code
## 
## 
  |                                                                       
  |.............................................                    |  69%
## label: evaluate-model-training-data (with options) 
## List of 2
##  $ fig.width : num 10
##  $ fig.height: num 10
```

```
## 
  |                                                                       
  |...............................................                  |  72%
##   ordinary text without R code
## 
## 
  |                                                                       
  |.................................................                |  76%
## label: evaluate-model-control-data (with options) 
## List of 2
##  $ fig.width : num 10
##  $ fig.height: num 10
```

```
## 
  |                                                                       
  |....................................................             |  79%
##   ordinary text without R code
## 
## 
  |                                                                       
  |......................................................           |  83%
## label: evaluate-model-control-data-2 (with options) 
## List of 2
##  $ fig.width : num 10
##  $ fig.height: num 10
```

```
## 
  |                                                                       
  |........................................................         |  86%
##   ordinary text without R code
## 
## 
  |                                                                       
  |..........................................................       |  90%
## label: null-model-auc-training
## 
  |                                                                       
  |.............................................................    |  93%
##   ordinary text without R code
## 
## 
  |                                                                       
  |...............................................................  |  97%
## label: store-result
## 
  |                                                                       
  |.................................................................| 100%
##   ordinary text without R code
```

```
## output file: renderer.knit.md
```

```
## "C:/Program Files/RStudio/bin/pandoc/pandoc" +RTS -K512m -RTS renderer.utf8.md --to html --from markdown+autolink_bare_uris+ascii_identifiers+tex_math_single_backslash --output pandoc6e019224ba.html --smart --email-obfuscation none --self-contained --standalone --section-divs --template "C:\Users\vshal\Documents\R\win-library\3.4\rmarkdown\rmd\h\default.html" --no-highlight --variable highlightjs=1 --variable "theme:bootstrap" --include-in-header "C:\Users\vshal\AppData\Local\Temp\RtmpWiYTWp\rmarkdown-str6e042612ba1.html" --mathjax --variable "mathjax-url:https://mathjax.rstudio.com/latest/MathJax.js?config=TeX-AMS-MML_HTMLorMML"
```

```
## 
## Output created: step_29dup_2.html
## 
## 
## processing file: renderer.Rmd
```

```
## 
  |                                                                       
  |                                                                 |   0%
  |                                                                       
  |..                                                               |   3%
##    inline R code fragments
## 
## 
  |                                                                       
  |....                                                             |   7%
## label: set-knitr
## 
  |                                                                       
  |.......                                                          |  10%
##   ordinary text without R code
## 
## 
  |                                                                       
  |.........                                                        |  14%
## label: fold-presence-data
## 
  |                                                                       
  |...........                                                      |  17%
##   ordinary text without R code
## 
## 
  |                                                                       
  |.............                                                    |  21%
## label: background-points
## 
  |                                                                       
  |................                                                 |  24%
##   ordinary text without R code
## 
## 
  |                                                                       
  |..................                                               |  28%
## label: data-visualization (with options) 
## List of 2
##  $ fig.height: num 16
##  $ fig.width : num 10
```

```
## 
  |                                                                       
  |....................                                             |  31%
##   ordinary text without R code
## 
## 
  |                                                                       
  |......................                                           |  34%
## label: drop-layers
## 
  |                                                                       
  |.........................                                        |  38%
##   ordinary text without R code
## 
## 
  |                                                                       
  |...........................                                      |  41%
## label: ajustar-modelo (with options) 
## List of 2
##  $ fig.height: num 10
##  $ fig.width : num 10
## 
## 
  |                                                                       
  |.............................                                    |  45%
##   ordinary text without R code
## 
## 
  |                                                                       
  |...............................                                  |  48%
## label: variable-importance (with options) 
## List of 2
##  $ fig.height: num 7
##  $ fig.width : num 10
```

```
## 
  |                                                                       
  |..................................                               |  52%
##   ordinary text without R code
## 
## 
  |                                                                       
  |....................................                             |  55%
## label: curvas-respuesta (with options) 
## List of 2
##  $ fig.height: num 10
##  $ fig.width : num 10
```

```
## 
  |                                                                       
  |......................................                           |  59%
##   ordinary text without R code
## 
## 
  |                                                                       
  |........................................                         |  62%
## label: revisar-datos-maxent-en-html
## 
  |                                                                       
  |...........................................                      |  66%
##   ordinary text without R code
## 
## 
  |                                                                       
  |.............................................                    |  69%
## label: evaluate-model-training-data (with options) 
## List of 2
##  $ fig.width : num 10
##  $ fig.height: num 10
```

```
## 
  |                                                                       
  |...............................................                  |  72%
##   ordinary text without R code
## 
## 
  |                                                                       
  |.................................................                |  76%
## label: evaluate-model-control-data (with options) 
## List of 2
##  $ fig.width : num 10
##  $ fig.height: num 10
```

```
## 
  |                                                                       
  |....................................................             |  79%
##   ordinary text without R code
## 
## 
  |                                                                       
  |......................................................           |  83%
## label: evaluate-model-control-data-2 (with options) 
## List of 2
##  $ fig.width : num 10
##  $ fig.height: num 10
```

```
## 
  |                                                                       
  |........................................................         |  86%
##   ordinary text without R code
## 
## 
  |                                                                       
  |..........................................................       |  90%
## label: null-model-auc-training
## 
  |                                                                       
  |.............................................................    |  93%
##   ordinary text without R code
## 
## 
  |                                                                       
  |...............................................................  |  97%
## label: store-result
## 
  |                                                                       
  |.................................................................| 100%
##   ordinary text without R code
```

```
## output file: renderer.knit.md
```

```
## "C:/Program Files/RStudio/bin/pandoc/pandoc" +RTS -K512m -RTS renderer.utf8.md --to html --from markdown+autolink_bare_uris+ascii_identifiers+tex_math_single_backslash --output pandoc6e0734a4903.html --smart --email-obfuscation none --self-contained --standalone --section-divs --template "C:\Users\vshal\Documents\R\win-library\3.4\rmarkdown\rmd\h\default.html" --no-highlight --variable highlightjs=1 --variable "theme:bootstrap" --include-in-header "C:\Users\vshal\AppData\Local\Temp\RtmpWiYTWp\rmarkdown-str6e07ce3b7b.html" --mathjax --variable "mathjax-url:https://mathjax.rstudio.com/latest/MathJax.js?config=TeX-AMS-MML_HTMLorMML"
```

```
## 
## Output created: step_29dup_3.html
## 
## 
## processing file: renderer.Rmd
```

```
## 
  |                                                                       
  |                                                                 |   0%
  |                                                                       
  |..                                                               |   3%
##    inline R code fragments
## 
## 
  |                                                                       
  |....                                                             |   7%
## label: set-knitr
## 
  |                                                                       
  |.......                                                          |  10%
##   ordinary text without R code
## 
## 
  |                                                                       
  |.........                                                        |  14%
## label: fold-presence-data
## 
  |                                                                       
  |...........                                                      |  17%
##   ordinary text without R code
## 
## 
  |                                                                       
  |.............                                                    |  21%
## label: background-points
## 
  |                                                                       
  |................                                                 |  24%
##   ordinary text without R code
## 
## 
  |                                                                       
  |..................                                               |  28%
## label: data-visualization (with options) 
## List of 2
##  $ fig.height: num 16
##  $ fig.width : num 10
```

```
## 
  |                                                                       
  |....................                                             |  31%
##   ordinary text without R code
## 
## 
  |                                                                       
  |......................                                           |  34%
## label: drop-layers
## 
  |                                                                       
  |.........................                                        |  38%
##   ordinary text without R code
## 
## 
  |                                                                       
  |...........................                                      |  41%
## label: ajustar-modelo (with options) 
## List of 2
##  $ fig.height: num 10
##  $ fig.width : num 10
## 
## 
  |                                                                       
  |.............................                                    |  45%
##   ordinary text without R code
## 
## 
  |                                                                       
  |...............................                                  |  48%
## label: variable-importance (with options) 
## List of 2
##  $ fig.height: num 7
##  $ fig.width : num 10
```

```
## 
  |                                                                       
  |..................................                               |  52%
##   ordinary text without R code
## 
## 
  |                                                                       
  |....................................                             |  55%
## label: curvas-respuesta (with options) 
## List of 2
##  $ fig.height: num 10
##  $ fig.width : num 10
```

```
## 
  |                                                                       
  |......................................                           |  59%
##   ordinary text without R code
## 
## 
  |                                                                       
  |........................................                         |  62%
## label: revisar-datos-maxent-en-html
## 
  |                                                                       
  |...........................................                      |  66%
##   ordinary text without R code
## 
## 
  |                                                                       
  |.............................................                    |  69%
## label: evaluate-model-training-data (with options) 
## List of 2
##  $ fig.width : num 10
##  $ fig.height: num 10
```

```
## 
  |                                                                       
  |...............................................                  |  72%
##   ordinary text without R code
## 
## 
  |                                                                       
  |.................................................                |  76%
## label: evaluate-model-control-data (with options) 
## List of 2
##  $ fig.width : num 10
##  $ fig.height: num 10
```

```
## 
  |                                                                       
  |....................................................             |  79%
##   ordinary text without R code
## 
## 
  |                                                                       
  |......................................................           |  83%
## label: evaluate-model-control-data-2 (with options) 
## List of 2
##  $ fig.width : num 10
##  $ fig.height: num 10
```

```
## 
  |                                                                       
  |........................................................         |  86%
##   ordinary text without R code
## 
## 
  |                                                                       
  |..........................................................       |  90%
## label: null-model-auc-training
## 
  |                                                                       
  |.............................................................    |  93%
##   ordinary text without R code
## 
## 
  |                                                                       
  |...............................................................  |  97%
## label: store-result
## 
  |                                                                       
  |.................................................................| 100%
##   ordinary text without R code
```

```
## output file: renderer.knit.md
```

```
## "C:/Program Files/RStudio/bin/pandoc/pandoc" +RTS -K512m -RTS renderer.utf8.md --to html --from markdown+autolink_bare_uris+ascii_identifiers+tex_math_single_backslash --output pandoc6e0e74353c.html --smart --email-obfuscation none --self-contained --standalone --section-divs --template "C:\Users\vshal\Documents\R\win-library\3.4\rmarkdown\rmd\h\default.html" --no-highlight --variable highlightjs=1 --variable "theme:bootstrap" --include-in-header "C:\Users\vshal\AppData\Local\Temp\RtmpWiYTWp\rmarkdown-str6e0582a4d7e.html" --mathjax --variable "mathjax-url:https://mathjax.rstudio.com/latest/MathJax.js?config=TeX-AMS-MML_HTMLorMML"
```

```
## 
## Output created: step_29dup_4.html
## 
## 
## processing file: renderer.Rmd
```

```
## 
  |                                                                       
  |                                                                 |   0%
  |                                                                       
  |..                                                               |   3%
##    inline R code fragments
## 
## 
  |                                                                       
  |....                                                             |   7%
## label: set-knitr
## 
  |                                                                       
  |.......                                                          |  10%
##   ordinary text without R code
## 
## 
  |                                                                       
  |.........                                                        |  14%
## label: fold-presence-data
## 
  |                                                                       
  |...........                                                      |  17%
##   ordinary text without R code
## 
## 
  |                                                                       
  |.............                                                    |  21%
## label: background-points
## 
  |                                                                       
  |................                                                 |  24%
##   ordinary text without R code
## 
## 
  |                                                                       
  |..................                                               |  28%
## label: data-visualization (with options) 
## List of 2
##  $ fig.height: num 16
##  $ fig.width : num 10
```

```
## 
  |                                                                       
  |....................                                             |  31%
##   ordinary text without R code
## 
## 
  |                                                                       
  |......................                                           |  34%
## label: drop-layers
## 
  |                                                                       
  |.........................                                        |  38%
##   ordinary text without R code
## 
## 
  |                                                                       
  |...........................                                      |  41%
## label: ajustar-modelo (with options) 
## List of 2
##  $ fig.height: num 10
##  $ fig.width : num 10
## 
## 
  |                                                                       
  |.............................                                    |  45%
##   ordinary text without R code
## 
## 
  |                                                                       
  |...............................                                  |  48%
## label: variable-importance (with options) 
## List of 2
##  $ fig.height: num 7
##  $ fig.width : num 10
```

```
## 
  |                                                                       
  |..................................                               |  52%
##   ordinary text without R code
## 
## 
  |                                                                       
  |....................................                             |  55%
## label: curvas-respuesta (with options) 
## List of 2
##  $ fig.height: num 10
##  $ fig.width : num 10
```

```
## 
  |                                                                       
  |......................................                           |  59%
##   ordinary text without R code
## 
## 
  |                                                                       
  |........................................                         |  62%
## label: revisar-datos-maxent-en-html
## 
  |                                                                       
  |...........................................                      |  66%
##   ordinary text without R code
## 
## 
  |                                                                       
  |.............................................                    |  69%
## label: evaluate-model-training-data (with options) 
## List of 2
##  $ fig.width : num 10
##  $ fig.height: num 10
```

```
## 
  |                                                                       
  |...............................................                  |  72%
##   ordinary text without R code
## 
## 
  |                                                                       
  |.................................................                |  76%
## label: evaluate-model-control-data (with options) 
## List of 2
##  $ fig.width : num 10
##  $ fig.height: num 10
```

```
## 
  |                                                                       
  |....................................................             |  79%
##   ordinary text without R code
## 
## 
  |                                                                       
  |......................................................           |  83%
## label: evaluate-model-control-data-2 (with options) 
## List of 2
##  $ fig.width : num 10
##  $ fig.height: num 10
```

```
## 
  |                                                                       
  |........................................................         |  86%
##   ordinary text without R code
## 
## 
  |                                                                       
  |..........................................................       |  90%
## label: null-model-auc-training
## 
  |                                                                       
  |.............................................................    |  93%
##   ordinary text without R code
## 
## 
  |                                                                       
  |...............................................................  |  97%
## label: store-result
## 
  |                                                                       
  |.................................................................| 100%
##   ordinary text without R code
```

```
## output file: renderer.knit.md
```

```
## "C:/Program Files/RStudio/bin/pandoc/pandoc" +RTS -K512m -RTS renderer.utf8.md --to html --from markdown+autolink_bare_uris+ascii_identifiers+tex_math_single_backslash --output pandoc6e033f87e6c.html --smart --email-obfuscation none --self-contained --standalone --section-divs --template "C:\Users\vshal\Documents\R\win-library\3.4\rmarkdown\rmd\h\default.html" --no-highlight --variable highlightjs=1 --variable "theme:bootstrap" --include-in-header "C:\Users\vshal\AppData\Local\Temp\RtmpWiYTWp\rmarkdown-str6e057df5072.html" --mathjax --variable "mathjax-url:https://mathjax.rstudio.com/latest/MathJax.js?config=TeX-AMS-MML_HTMLorMML"
```

```
## 
## Output created: step_29dup_5.html
## 
## 
## processing file: renderer.Rmd
```

```
## 
  |                                                                       
  |                                                                 |   0%
  |                                                                       
  |..                                                               |   3%
##    inline R code fragments
## 
## 
  |                                                                       
  |....                                                             |   7%
## label: set-knitr
## 
  |                                                                       
  |.......                                                          |  10%
##   ordinary text without R code
## 
## 
  |                                                                       
  |.........                                                        |  14%
## label: fold-presence-data
## 
  |                                                                       
  |...........                                                      |  17%
##   ordinary text without R code
## 
## 
  |                                                                       
  |.............                                                    |  21%
## label: background-points
## 
  |                                                                       
  |................                                                 |  24%
##   ordinary text without R code
## 
## 
  |                                                                       
  |..................                                               |  28%
## label: data-visualization (with options) 
## List of 2
##  $ fig.height: num 16
##  $ fig.width : num 10
```

```
## 
  |                                                                       
  |....................                                             |  31%
##   ordinary text without R code
## 
## 
  |                                                                       
  |......................                                           |  34%
## label: drop-layers
## 
  |                                                                       
  |.........................                                        |  38%
##   ordinary text without R code
## 
## 
  |                                                                       
  |...........................                                      |  41%
## label: ajustar-modelo (with options) 
## List of 2
##  $ fig.height: num 10
##  $ fig.width : num 10
## 
## 
  |                                                                       
  |.............................                                    |  45%
##   ordinary text without R code
## 
## 
  |                                                                       
  |...............................                                  |  48%
## label: variable-importance (with options) 
## List of 2
##  $ fig.height: num 7
##  $ fig.width : num 10
```

```
## 
  |                                                                       
  |..................................                               |  52%
##   ordinary text without R code
## 
## 
  |                                                                       
  |....................................                             |  55%
## label: curvas-respuesta (with options) 
## List of 2
##  $ fig.height: num 10
##  $ fig.width : num 10
```

```
## 
  |                                                                       
  |......................................                           |  59%
##   ordinary text without R code
## 
## 
  |                                                                       
  |........................................                         |  62%
## label: revisar-datos-maxent-en-html
## 
  |                                                                       
  |...........................................                      |  66%
##   ordinary text without R code
## 
## 
  |                                                                       
  |.............................................                    |  69%
## label: evaluate-model-training-data (with options) 
## List of 2
##  $ fig.width : num 10
##  $ fig.height: num 10
```

```
## 
  |                                                                       
  |...............................................                  |  72%
##   ordinary text without R code
## 
## 
  |                                                                       
  |.................................................                |  76%
## label: evaluate-model-control-data (with options) 
## List of 2
##  $ fig.width : num 10
##  $ fig.height: num 10
```

```
## 
  |                                                                       
  |....................................................             |  79%
##   ordinary text without R code
## 
## 
  |                                                                       
  |......................................................           |  83%
## label: evaluate-model-control-data-2 (with options) 
## List of 2
##  $ fig.width : num 10
##  $ fig.height: num 10
```

```
## 
  |                                                                       
  |........................................................         |  86%
##   ordinary text without R code
## 
## 
  |                                                                       
  |..........................................................       |  90%
## label: null-model-auc-training
## 
  |                                                                       
  |.............................................................    |  93%
##   ordinary text without R code
## 
## 
  |                                                                       
  |...............................................................  |  97%
## label: store-result
## 
  |                                                                       
  |.................................................................| 100%
##   ordinary text without R code
```

```
## output file: renderer.knit.md
```

```
## "C:/Program Files/RStudio/bin/pandoc/pandoc" +RTS -K512m -RTS renderer.utf8.md --to html --from markdown+autolink_bare_uris+ascii_identifiers+tex_math_single_backslash --output pandoc6e068be2d9c.html --smart --email-obfuscation none --self-contained --standalone --section-divs --template "C:\Users\vshal\Documents\R\win-library\3.4\rmarkdown\rmd\h\default.html" --no-highlight --variable highlightjs=1 --variable "theme:bootstrap" --include-in-header "C:\Users\vshal\AppData\Local\Temp\RtmpWiYTWp\rmarkdown-str6e03a9471f.html" --mathjax --variable "mathjax-url:https://mathjax.rstudio.com/latest/MathJax.js?config=TeX-AMS-MML_HTMLorMML"
```

```
## 
## Output created: step_29dup_6.html
## 
## 
## processing file: renderer.Rmd
```

```
## 
  |                                                                       
  |                                                                 |   0%
  |                                                                       
  |..                                                               |   3%
##    inline R code fragments
## 
## 
  |                                                                       
  |....                                                             |   7%
## label: set-knitr
## 
  |                                                                       
  |.......                                                          |  10%
##   ordinary text without R code
## 
## 
  |                                                                       
  |.........                                                        |  14%
## label: fold-presence-data
## 
  |                                                                       
  |...........                                                      |  17%
##   ordinary text without R code
## 
## 
  |                                                                       
  |.............                                                    |  21%
## label: background-points
## 
  |                                                                       
  |................                                                 |  24%
##   ordinary text without R code
## 
## 
  |                                                                       
  |..................                                               |  28%
## label: data-visualization (with options) 
## List of 2
##  $ fig.height: num 16
##  $ fig.width : num 10
```

```
## 
  |                                                                       
  |....................                                             |  31%
##   ordinary text without R code
## 
## 
  |                                                                       
  |......................                                           |  34%
## label: drop-layers
## 
  |                                                                       
  |.........................                                        |  38%
##   ordinary text without R code
## 
## 
  |                                                                       
  |...........................                                      |  41%
## label: ajustar-modelo (with options) 
## List of 2
##  $ fig.height: num 10
##  $ fig.width : num 10
## 
## 
  |                                                                       
  |.............................                                    |  45%
##   ordinary text without R code
## 
## 
  |                                                                       
  |...............................                                  |  48%
## label: variable-importance (with options) 
## List of 2
##  $ fig.height: num 7
##  $ fig.width : num 10
```

```
## 
  |                                                                       
  |..................................                               |  52%
##   ordinary text without R code
## 
## 
  |                                                                       
  |....................................                             |  55%
## label: curvas-respuesta (with options) 
## List of 2
##  $ fig.height: num 10
##  $ fig.width : num 10
```

```
## 
  |                                                                       
  |......................................                           |  59%
##   ordinary text without R code
## 
## 
  |                                                                       
  |........................................                         |  62%
## label: revisar-datos-maxent-en-html
## 
  |                                                                       
  |...........................................                      |  66%
##   ordinary text without R code
## 
## 
  |                                                                       
  |.............................................                    |  69%
## label: evaluate-model-training-data (with options) 
## List of 2
##  $ fig.width : num 10
##  $ fig.height: num 10
```

```
## 
  |                                                                       
  |...............................................                  |  72%
##   ordinary text without R code
## 
## 
  |                                                                       
  |.................................................                |  76%
## label: evaluate-model-control-data (with options) 
## List of 2
##  $ fig.width : num 10
##  $ fig.height: num 10
```

```
## 
  |                                                                       
  |....................................................             |  79%
##   ordinary text without R code
## 
## 
  |                                                                       
  |......................................................           |  83%
## label: evaluate-model-control-data-2 (with options) 
## List of 2
##  $ fig.width : num 10
##  $ fig.height: num 10
```

```
## 
  |                                                                       
  |........................................................         |  86%
##   ordinary text without R code
## 
## 
  |                                                                       
  |..........................................................       |  90%
## label: null-model-auc-training
## 
  |                                                                       
  |.............................................................    |  93%
##   ordinary text without R code
## 
## 
  |                                                                       
  |...............................................................  |  97%
## label: store-result
## 
  |                                                                       
  |.................................................................| 100%
##   ordinary text without R code
```

```
## output file: renderer.knit.md
```

```
## "C:/Program Files/RStudio/bin/pandoc/pandoc" +RTS -K512m -RTS renderer.utf8.md --to html --from markdown+autolink_bare_uris+ascii_identifiers+tex_math_single_backslash --output pandoc6e055ad3fd3.html --smart --email-obfuscation none --self-contained --standalone --section-divs --template "C:\Users\vshal\Documents\R\win-library\3.4\rmarkdown\rmd\h\default.html" --no-highlight --variable highlightjs=1 --variable "theme:bootstrap" --include-in-header "C:\Users\vshal\AppData\Local\Temp\RtmpWiYTWp\rmarkdown-str6e06731384c.html" --mathjax --variable "mathjax-url:https://mathjax.rstudio.com/latest/MathJax.js?config=TeX-AMS-MML_HTMLorMML"
```

```
## 
## Output created: step_29dup_7.html
## 
## 
## processing file: renderer.Rmd
```

```
## 
  |                                                                       
  |                                                                 |   0%
  |                                                                       
  |..                                                               |   3%
##    inline R code fragments
## 
## 
  |                                                                       
  |....                                                             |   7%
## label: set-knitr
## 
  |                                                                       
  |.......                                                          |  10%
##   ordinary text without R code
## 
## 
  |                                                                       
  |.........                                                        |  14%
## label: fold-presence-data
## 
  |                                                                       
  |...........                                                      |  17%
##   ordinary text without R code
## 
## 
  |                                                                       
  |.............                                                    |  21%
## label: background-points
## 
  |                                                                       
  |................                                                 |  24%
##   ordinary text without R code
## 
## 
  |                                                                       
  |..................                                               |  28%
## label: data-visualization (with options) 
## List of 2
##  $ fig.height: num 16
##  $ fig.width : num 10
```

```
## 
  |                                                                       
  |....................                                             |  31%
##   ordinary text without R code
## 
## 
  |                                                                       
  |......................                                           |  34%
## label: drop-layers
## 
  |                                                                       
  |.........................                                        |  38%
##   ordinary text without R code
## 
## 
  |                                                                       
  |...........................                                      |  41%
## label: ajustar-modelo (with options) 
## List of 2
##  $ fig.height: num 10
##  $ fig.width : num 10
## 
## 
  |                                                                       
  |.............................                                    |  45%
##   ordinary text without R code
## 
## 
  |                                                                       
  |...............................                                  |  48%
## label: variable-importance (with options) 
## List of 2
##  $ fig.height: num 7
##  $ fig.width : num 10
```

```
## 
  |                                                                       
  |..................................                               |  52%
##   ordinary text without R code
## 
## 
  |                                                                       
  |....................................                             |  55%
## label: curvas-respuesta (with options) 
## List of 2
##  $ fig.height: num 10
##  $ fig.width : num 10
```

```
## 
  |                                                                       
  |......................................                           |  59%
##   ordinary text without R code
## 
## 
  |                                                                       
  |........................................                         |  62%
## label: revisar-datos-maxent-en-html
## 
  |                                                                       
  |...........................................                      |  66%
##   ordinary text without R code
## 
## 
  |                                                                       
  |.............................................                    |  69%
## label: evaluate-model-training-data (with options) 
## List of 2
##  $ fig.width : num 10
##  $ fig.height: num 10
```

```
## 
  |                                                                       
  |...............................................                  |  72%
##   ordinary text without R code
## 
## 
  |                                                                       
  |.................................................                |  76%
## label: evaluate-model-control-data (with options) 
## List of 2
##  $ fig.width : num 10
##  $ fig.height: num 10
```

```
## 
  |                                                                       
  |....................................................             |  79%
##   ordinary text without R code
## 
## 
  |                                                                       
  |......................................................           |  83%
## label: evaluate-model-control-data-2 (with options) 
## List of 2
##  $ fig.width : num 10
##  $ fig.height: num 10
```

```
## 
  |                                                                       
  |........................................................         |  86%
##   ordinary text without R code
## 
## 
  |                                                                       
  |..........................................................       |  90%
## label: null-model-auc-training
## 
  |                                                                       
  |.............................................................    |  93%
##   ordinary text without R code
## 
## 
  |                                                                       
  |...............................................................  |  97%
## label: store-result
## 
  |                                                                       
  |.................................................................| 100%
##   ordinary text without R code
```

```
## output file: renderer.knit.md
```

```
## "C:/Program Files/RStudio/bin/pandoc/pandoc" +RTS -K512m -RTS renderer.utf8.md --to html --from markdown+autolink_bare_uris+ascii_identifiers+tex_math_single_backslash --output pandoc6e047af2619.html --smart --email-obfuscation none --self-contained --standalone --section-divs --template "C:\Users\vshal\Documents\R\win-library\3.4\rmarkdown\rmd\h\default.html" --no-highlight --variable highlightjs=1 --variable "theme:bootstrap" --include-in-header "C:\Users\vshal\AppData\Local\Temp\RtmpWiYTWp\rmarkdown-str6e0121f7ec3.html" --mathjax --variable "mathjax-url:https://mathjax.rstudio.com/latest/MathJax.js?config=TeX-AMS-MML_HTMLorMML"
```

```
## 
## Output created: step_29dup_8.html
## 
## 
## processing file: renderer.Rmd
```

```
## 
  |                                                                       
  |                                                                 |   0%
  |                                                                       
  |..                                                               |   3%
##    inline R code fragments
## 
## 
  |                                                                       
  |....                                                             |   7%
## label: set-knitr
## 
  |                                                                       
  |.......                                                          |  10%
##   ordinary text without R code
## 
## 
  |                                                                       
  |.........                                                        |  14%
## label: fold-presence-data
## 
  |                                                                       
  |...........                                                      |  17%
##   ordinary text without R code
## 
## 
  |                                                                       
  |.............                                                    |  21%
## label: background-points
## 
  |                                                                       
  |................                                                 |  24%
##   ordinary text without R code
## 
## 
  |                                                                       
  |..................                                               |  28%
## label: data-visualization (with options) 
## List of 2
##  $ fig.height: num 16
##  $ fig.width : num 10
```

```
## 
  |                                                                       
  |....................                                             |  31%
##   ordinary text without R code
## 
## 
  |                                                                       
  |......................                                           |  34%
## label: drop-layers
## 
  |                                                                       
  |.........................                                        |  38%
##   ordinary text without R code
## 
## 
  |                                                                       
  |...........................                                      |  41%
## label: ajustar-modelo (with options) 
## List of 2
##  $ fig.height: num 10
##  $ fig.width : num 10
## 
## 
  |                                                                       
  |.............................                                    |  45%
##   ordinary text without R code
## 
## 
  |                                                                       
  |...............................                                  |  48%
## label: variable-importance (with options) 
## List of 2
##  $ fig.height: num 7
##  $ fig.width : num 10
```

```
## 
  |                                                                       
  |..................................                               |  52%
##   ordinary text without R code
## 
## 
  |                                                                       
  |....................................                             |  55%
## label: curvas-respuesta (with options) 
## List of 2
##  $ fig.height: num 10
##  $ fig.width : num 10
```

```
## 
  |                                                                       
  |......................................                           |  59%
##   ordinary text without R code
## 
## 
  |                                                                       
  |........................................                         |  62%
## label: revisar-datos-maxent-en-html
## 
  |                                                                       
  |...........................................                      |  66%
##   ordinary text without R code
## 
## 
  |                                                                       
  |.............................................                    |  69%
## label: evaluate-model-training-data (with options) 
## List of 2
##  $ fig.width : num 10
##  $ fig.height: num 10
```

```
## 
  |                                                                       
  |...............................................                  |  72%
##   ordinary text without R code
## 
## 
  |                                                                       
  |.................................................                |  76%
## label: evaluate-model-control-data (with options) 
## List of 2
##  $ fig.width : num 10
##  $ fig.height: num 10
```

```
## 
  |                                                                       
  |....................................................             |  79%
##   ordinary text without R code
## 
## 
  |                                                                       
  |......................................................           |  83%
## label: evaluate-model-control-data-2 (with options) 
## List of 2
##  $ fig.width : num 10
##  $ fig.height: num 10
```

```
## 
  |                                                                       
  |........................................................         |  86%
##   ordinary text without R code
## 
## 
  |                                                                       
  |..........................................................       |  90%
## label: null-model-auc-training
## 
  |                                                                       
  |.............................................................    |  93%
##   ordinary text without R code
## 
## 
  |                                                                       
  |...............................................................  |  97%
## label: store-result
## 
  |                                                                       
  |.................................................................| 100%
##   ordinary text without R code
```

```
## output file: renderer.knit.md
```

```
## "C:/Program Files/RStudio/bin/pandoc/pandoc" +RTS -K512m -RTS renderer.utf8.md --to html --from markdown+autolink_bare_uris+ascii_identifiers+tex_math_single_backslash --output pandoc6e02fab4576.html --smart --email-obfuscation none --self-contained --standalone --section-divs --template "C:\Users\vshal\Documents\R\win-library\3.4\rmarkdown\rmd\h\default.html" --no-highlight --variable highlightjs=1 --variable "theme:bootstrap" --include-in-header "C:\Users\vshal\AppData\Local\Temp\RtmpWiYTWp\rmarkdown-str6e0181a494a.html" --mathjax --variable "mathjax-url:https://mathjax.rstudio.com/latest/MathJax.js?config=TeX-AMS-MML_HTMLorMML"
```

```
## 
## Output created: step_29dup_9.html
## 
## 
## processing file: renderer.Rmd
```

```
## 
  |                                                                       
  |                                                                 |   0%
  |                                                                       
  |..                                                               |   3%
##    inline R code fragments
## 
## 
  |                                                                       
  |....                                                             |   7%
## label: set-knitr
## 
  |                                                                       
  |.......                                                          |  10%
##   ordinary text without R code
## 
## 
  |                                                                       
  |.........                                                        |  14%
## label: fold-presence-data
## 
  |                                                                       
  |...........                                                      |  17%
##   ordinary text without R code
## 
## 
  |                                                                       
  |.............                                                    |  21%
## label: background-points
## 
  |                                                                       
  |................                                                 |  24%
##   ordinary text without R code
## 
## 
  |                                                                       
  |..................                                               |  28%
## label: data-visualization (with options) 
## List of 2
##  $ fig.height: num 16
##  $ fig.width : num 10
```

```
## 
  |                                                                       
  |....................                                             |  31%
##   ordinary text without R code
## 
## 
  |                                                                       
  |......................                                           |  34%
## label: drop-layers
## 
  |                                                                       
  |.........................                                        |  38%
##   ordinary text without R code
## 
## 
  |                                                                       
  |...........................                                      |  41%
## label: ajustar-modelo (with options) 
## List of 2
##  $ fig.height: num 10
##  $ fig.width : num 10
## 
## 
  |                                                                       
  |.............................                                    |  45%
##   ordinary text without R code
## 
## 
  |                                                                       
  |...............................                                  |  48%
## label: variable-importance (with options) 
## List of 2
##  $ fig.height: num 7
##  $ fig.width : num 10
```

```
## 
  |                                                                       
  |..................................                               |  52%
##   ordinary text without R code
## 
## 
  |                                                                       
  |....................................                             |  55%
## label: curvas-respuesta (with options) 
## List of 2
##  $ fig.height: num 10
##  $ fig.width : num 10
```

```
## 
  |                                                                       
  |......................................                           |  59%
##   ordinary text without R code
## 
## 
  |                                                                       
  |........................................                         |  62%
## label: revisar-datos-maxent-en-html
## 
  |                                                                       
  |...........................................                      |  66%
##   ordinary text without R code
## 
## 
  |                                                                       
  |.............................................                    |  69%
## label: evaluate-model-training-data (with options) 
## List of 2
##  $ fig.width : num 10
##  $ fig.height: num 10
```

```
## 
  |                                                                       
  |...............................................                  |  72%
##   ordinary text without R code
## 
## 
  |                                                                       
  |.................................................                |  76%
## label: evaluate-model-control-data (with options) 
## List of 2
##  $ fig.width : num 10
##  $ fig.height: num 10
```

```
## 
  |                                                                       
  |....................................................             |  79%
##   ordinary text without R code
## 
## 
  |                                                                       
  |......................................................           |  83%
## label: evaluate-model-control-data-2 (with options) 
## List of 2
##  $ fig.width : num 10
##  $ fig.height: num 10
```

```
## 
  |                                                                       
  |........................................................         |  86%
##   ordinary text without R code
## 
## 
  |                                                                       
  |..........................................................       |  90%
## label: null-model-auc-training
## 
  |                                                                       
  |.............................................................    |  93%
##   ordinary text without R code
## 
## 
  |                                                                       
  |...............................................................  |  97%
## label: store-result
## 
  |                                                                       
  |.................................................................| 100%
##   ordinary text without R code
```

```
## output file: renderer.knit.md
```

```
## "C:/Program Files/RStudio/bin/pandoc/pandoc" +RTS -K512m -RTS renderer.utf8.md --to html --from markdown+autolink_bare_uris+ascii_identifiers+tex_math_single_backslash --output pandoc6e0228a76f3.html --smart --email-obfuscation none --self-contained --standalone --section-divs --template "C:\Users\vshal\Documents\R\win-library\3.4\rmarkdown\rmd\h\default.html" --no-highlight --variable highlightjs=1 --variable "theme:bootstrap" --include-in-header "C:\Users\vshal\AppData\Local\Temp\RtmpWiYTWp\rmarkdown-str6e010ca1aaa.html" --mathjax --variable "mathjax-url:https://mathjax.rstudio.com/latest/MathJax.js?config=TeX-AMS-MML_HTMLorMML"
```

```
## 
## Output created: step_29dup_10.html
```

#### Store resulting model object

```
#str(iterations)

saveRDS(iterations, paste(scriptpath, "/iterations_object_",run_code,".rds",sep=""))
```
