## Supplementary material for "Vulnerability to climate change for narrowly ranged species: the case of Ecuadorian endemic *Magnolia mercedesiarum*": predictor_current.html

Script for Shalisko et al. 2018 - Cross-validated SDM script 4


### Script for Shalisko et al. 2018 - Cross-validated SDM script 4

###### *Viacheslav Shalisko*

###### *15 of november 2017*

### *Magnolia mercedesiarum* SDM on current conditions

##### Habitat suitability surfaces on current conditions

```
knitr::opts_chunk$set(echo = TRUE)
knitr::opts_chunk$set(error = TRUE)
```

#### Data preparation

###### Set adjustable variables

```
run_code <- '29dup'
basepath <- 'C:/Users/vshal/Downloads/MaxEnt_test_runs/run29_duplicate'
basename <- 'Magnolia_mercedesiarum_MAXENT_climond2_HR_'

scriptpath <- 'C:/Users/vshal/Downloads/MaxEnt_test_runs/run29_duplicate'

random_background_number <- 450
presence_points_count <- 50
double_presence_points_count <- 100

cross_validation_lenght <- 1

#layers_to_drop <- c(2,6,7,9,11,12,14,15,17,18,19,21)
layers_to_drop <- c(21)    # the last layer no. 21 is dummy (repetition of the first, included just to drop it when nothing to drop)
```

###### Load modules

```
library(maptools)
```

```
## Loading required package: sp
```

```
## Checking rgeos availability: FALSE
##      Note: when rgeos is not available, polygon geometry     computations in maptools depend on gpclib,
##      which has a restricted licence. It is disabled by default;
##      to enable gpclib, type gpclibPermit()
```

```
library(rworldmap)    # worldmap datasets
```

```
## Warning: package 'rworldmap' was built under R version 3.4.2
```

```
## ### Welcome to rworldmap ###
```

```
## For a short introduction type :   vignette('rworldmap')
```

```
library(rworldxtra)   # hires worldmap spatial dataframe
```

```
## Warning: package 'rworldxtra' was built under R version 3.4.2
```

```
library(sp)
library(raster)
library(dismo)
library(rgdal)
```

```
## rgdal: version: 1.2-11, (SVN revision 676)
##  Geospatial Data Abstraction Library extensions to R successfully loaded
##  Loaded GDAL runtime: GDAL 2.2.0, released 2017/04/28
##  Path to GDAL shared files: C:/Users/vshal/Documents/R/win-library/3.4/rgdal/gdal
##  Loaded PROJ.4 runtime: Rel. 4.9.3, 15 August 2016, [PJ_VERSION: 493]
##  Path to PROJ.4 shared files: C:/Users/vshal/Documents/R/win-library/3.4/rgdal/proj
##  Linking to sp version: 1.2-5
```

```
library(rJava)
```

###### Load raster predictor variables

```
names <- c("alt","bio1","bio2","bio3","bio4",
              "bio5","bio6","bio7","bio8","bio9",
              "bio10","bio11","bio12","bio13","bio14",
              "bio15","bio16","bio17","bio18","bio19",
              "dummy")

capas_raster<-stack(c("C:/Users/vshal/Downloads/GMTED2010/GMTED2010_075_Andes.tif",
                      "C:/Users/vshal/Downloads/SDM_highres_Andes/biovars_climond2_Andes_HR.tif",
                      "C:/Users/vshal/Downloads/GMTED2010/GMTED2010_075_Andes.tif"))

names(capas_raster) <- names

capas_raster
```

```
## class       : RasterStack 
## dimensions  : 9600, 5760, 55296000, 21  (nrow, ncol, ncell, nlayers)
## resolution  : 0.002083333, 0.002083333  (x, y)
## extent      : -82.00014, -70.00014, -10.00014, 9.999861  (xmin, xmax, ymin, ymax)
## coord. ref. : +proj=longlat +datum=WGS84 +no_defs +ellps=WGS84 +towgs84=0,0,0 
## names       :         alt,        bio1,        bio2,        bio3,        bio4,        bio5,        bio6,        bio7,        bio8,        bio9,       bio10,       bio11,       bio12,       bio13,       bio14, ... 
## min values  :  -50.000000,  -67.965177,   51.788566,   46.986875,   77.333818,   11.052460, -164.658356,   62.161942,  -56.866343,  -81.849262,  -56.866343,  -81.849262,    1.000000,    1.000000,   -4.000000, ... 
## max values  :  6544.00000,   290.01051,   166.67886,    97.66176,  2813.85039,   353.26230,   240.15173,   195.08090,   294.50851,   297.26141,   297.26141,   282.18530,  8007.00000,   965.00000,   537.00000, ...
```

```
#summary(capas_raster)

bio10 <- subset(capas_raster,10)
```

###### Drop unsignificant layers (based on preliminary test run)

```
capas_raster_drop <- dropLayer(capas_raster,layers_to_drop)
names_after_drop <- names(capas_raster_drop)
```

###### Load model

```
iterations <- readRDS(paste(scriptpath, "/iterations_object_",run_code,".rds",sep=""))

#str(iterations)
```

#### Prediction

###### Predict presence probaility surface

```
for (j in 1:cross_validation_lenght) {
  my_obj <- iterations[[j]]
  resultado_maxent <- predict(my_obj$model, capas_raster_drop, 
                           na.rm=TRUE,
                           progress = 'windows') 
  
  writeRaster(resultado_maxent, filename=paste(basepath, '/' ,basename ,'run_',run_code,'_',j,'.tif',sep=''), 
           format="GTiff", overwrite=TRUE)
  
  plot(resultado_maxent, main = paste('probability raster ',j,sep=''))
  #plot(bio10, xlim=c(-82,-70), ylim=c(-10,10), axes=TRUE)
  world_high <- getMap(resolution = "high")
  #plot(world_high, xlim=c(-82,-70), ylim=c(-10,10), axes=TRUE)
  plot(world_high, add=TRUE)
  
  puntos_entrenamiento_coordenadas<-data.frame(cbind(
                              my_obj$entr$X1,
                              my_obj$entr$X2))  
  
  points(puntos_entrenamiento_coordenadas, col='red')
  #str(my_obj)
  #str(my_obj$auc_null_model_mean)

  #plot(my_obj$e_train$e, 'ROC')
  #print(my_obj$e_train)
  #iterations[j]$e_train$e@auc
  
}
```

```
## Loading required namespace: tcltk
```

```
## Error: $ operator is invalid for atomic vectors
```
