## Supplementary material for "Vulnerability to climate change for narrowly ranged species: the case of Ecuadorian endemic *Magnolia mercedesiarum*": predictor_future_he45_70.html

Predictor (run 29) - two steps


### Script for Shalisko et al. 2018 - Cross-validated SDM script 5 (RCP4.5 2070)

###### *Viacheslav Shalisko*

###### *30 of november 2017*

### *Magnolia mercedesiarum* SDM on future conditions

##### Habitat suitability surfaces on projected future conditions in 2070, scenario RCP4.5, HadGEM2-ES model

```
## class       : RasterStack 
## dimensions  : 9600, 5760, 55296000, 21  (nrow, ncol, ncell, nlayers)
## resolution  : 0.002083333, 0.002083333  (x, y)
## extent      : -82.00014, -70.00014, -10.00014, 9.999861  (xmin, xmax, ymin, ymax)
## coord. ref. : +proj=longlat +datum=WGS84 +no_defs +ellps=WGS84 +towgs84=0,0,0 
## names       :          alt,         bio1,         bio2,         bio3,         bio4,         bio5,         bio6,         bio7,         bio8,         bio9,        bio10,        bio11,        bio12,        bio13,        bio14, ... 
## min values  :  -50.0000000,  -50.2039675,   44.5805779,   46.9416416,  103.6553730,   17.3961945, -153.5604553,   60.1606903,  -40.0025525,  -61.5954545,  -40.0025525,  -61.5954545,    0.8121550,    0.8319550,   -1.0774200, ... 
## max values  :   6544.00000,    321.99548,    159.25984,     94.19313,   2625.62311,    393.09305,    264.29697,    202.19723,    320.04180,    333.54686,    333.54686,    316.99212,  11764.69678,   1398.69067,    843.69104, ...
```

```
#summary(capas_raster)

bio10 <- subset(capas_raster,10)
```

###### Drop unsignificant layers (based on preliminary test run)

```
capas_raster_drop <- dropLayer(capas_raster,layers_to_drop)
names_after_drop <- names(capas_raster_drop)
```

```
iterations <- readRDS(paste(scriptpath, "/iterations_object_",run_code,".rds",sep=""))

#str(iterations)
```

#### Prediction

###### Predict presence probaility surface

```
for (j in 1:cross_validation_lenght) {
  my_obj <- iterations[[j]]
  resultado_maxent <- predict(my_obj$model, capas_raster_drop, 
                           na.rm=TRUE,
                           progress = 'windows') 
  
  writeRaster(resultado_maxent, filename=paste(basepath, '/' ,basename ,'run_',run_code,'_',j,'.tif',sep=''), 
           format="GTiff", overwrite=TRUE)
  
  plot(resultado_maxent, main = paste('probability raster ',j,sep=''))
  #plot(bio10, xlim=c(-82,-70), ylim=c(-10,10), axes=TRUE)
  world_high <- getMap(resolution = "high")
  #plot(world_high, xlim=c(-82,-70), ylim=c(-10,10), axes=TRUE)
  plot(world_high, add=TRUE)
  
  #puntos_entrenamiento_coordenadas<-data.frame(cbind(
  #                            my_obj$entr$X1,
  #                            my_obj$entr$X2))  
  
  #points(puntos_entrenamiento_coordenadas, col='red')
  #str(my_obj)
  #str(my_obj$auc_null_model_mean)

  #plot(my_obj$e_train$e, 'ROC')
  #print(my_obj$e_train)
  #iterations[j]$e_train$e@auc
  
}
```

```
## Loading required namespace: tcltk
```
