## Supplementary material for "Vulnerability to climate change for narrowly ranged species: the case of Ecuadorian endemic *Magnolia mercedesiarum*": renderer.Rmd

````
---
title: "Script for Shalisko et al. 2018 - Cross-validated SDM script 2"
author: "Viacheslav Shalisko"
date: "22 of october 2017"
output: html_document
---

## Replication `r i` of MaxEnt SDM

```{r set-knitr}
knitr::opts_chunk$set(echo = TRUE)
knitr::opts_chunk$set(error = TRUE)
```


#### Produce and K-fold presence and test points

```{r fold-presence-data}
set.seed(random_seed_folding)

random_presence_points <- randomPoints(presence_raster_mask, 
                                        n = double_presence_points_count, 
                                        tryf = 10)

# fold in 2 subsets
fold <- kfold(random_presence_points, k = 2)

# set one subset for training
puntos_entrenamiento_coordenadas <- random_presence_points[ fold == 1, ]
dim(puntos_entrenamiento_coordenadas)

# set another subset for initial model evaluation
puntos_control_coordenadas <- random_presence_points[ fold == 2, ]
dim(puntos_control_coordenadas)
```

#### Produce background points within target mask

```{r background-points}
set.seed(random_seed_background)

# "tryf"" parameter is set to ensure the required number of random points, as the points generated outside
# of the tgs_raster are descarted, examples tryf = 50, tryf = 100
puntos_aleatorios_fondo1 <- randomPoints(tgs_raster_full, 
                                        n = random_background_number, 
                                        p = random_presence_points,
                                        tryf = 10)
puntos_aleatorios_fondo2 <- randomPoints(tgs_raster_500, 
                                        n = random_background_number, 
                                        p = random_presence_points,
                                        tryf = 20)

puntos_aleatorios_fondo <- rbind(puntos_aleatorios_fondo1,puntos_aleatorios_fondo2)
dim(puntos_aleatorios_fondo)
```


#### Render presencce, test and background data

```{r data-visualization, fig.height=16, fig.width=10}

plot(bio10, xlim=c(-82,-70), ylim=c(-10,10), axes=TRUE)
world_high <- getMap(resolution = "high")
plot(world_high, add=TRUE)
# data(wrld_simpl)
# plot(wrld_simpl, add = TRUE)
points(puntos_aleatorios_fondo, cex=0.5, col="black")
points(puntos_entrenamiento_coordenadas,col='red')
points(puntos_control_coordenadas,col='blue',cex=0.7)

#points(puntos_control_extra_coordenadas,col='navy',cex=0.5)
```

#### Drop unsignificant layers (based on preliminary test run)

```{r drop-layers}
capas_raster_drop <- dropLayer(capas_raster,layers_to_drop)
names_after_drop <- names(capas_raster_drop)
```

## Modelling

```{r ajustar-modelo, fig.height=10, fig.width=10}
# create maxent output directory
dir.create(paste(basepath, '/maxent_',run_code,'_',i, sep=''), recursive=TRUE)

jar <- paste(system.file(package="dismo"), "/java/maxent.jar", sep='')

modelo_maxent <- maxent(x=capas_raster_drop, 
                        p=puntos_entrenamiento_coordenadas,
                        a=puntos_aleatorios_fondo,
                        path=paste(basepath, '/maxent_',run_code,'_',i, sep=''),
                        args=c(
                          'removeDuplicates=TRUE', 
                          'jackknife=TRUE',
                          'responsecurves=TRUE',
                          'threads=2',
                          'linear=TRUE',
                          'quadratic=TRUE',
                          'hinge=FALSE',
                          'product=FALSE'
                        ))

summary(modelo_maxent)
##str(modelo_maxent)
#representar resultados en forma de una lista
print(modelo_maxent@results)
#str(modelo_maxent@results)
```

#### Get variable importance values

```{r variable-importance, fig.height=7, fig.width=10}
#plot(modelo_maxent)

contribuciones <- maxent.select.contribution(modelo_maxent,names_after_drop)

#contribuciones
par(mfrow=c(1, 2))
dotchart(as.numeric(contribuciones[,2]),
        col='blue', pch=16,
        labels=contribuciones[,1],
        main='predictor contribution')
dotchart(as.numeric(contribuciones[,3]),
        col='red', pch=16,
        labels=contribuciones[,1],
        main='predictor importance in permutations')
```

#### Draw respnse cuves

```{r curvas-respuesta, fig.height=10, fig.width=10}
response(modelo_maxent)
```

#### Output MaxEnt HTML

```{r revisar-datos-maxent-en-html}
#modelo_maxent
```

## Initial model evaluation
#### Evaluate model with training dataset

```{r evaluate-model-training-data, fig.width=10, fig.height=10}
evalutaion_training <- evaluate.mdl(puntos_entrenamiento_coordenadas,puntos_aleatorios_fondo,capas_raster_drop,modelo_maxent)
str(evalutaion_training)
```

#### Evaluate model with control dataset

```{r evaluate-model-control-data, fig.width=10, fig.height=10}
evalutaion_test <- evaluate.mdl(puntos_control_coordenadas,puntos_aleatorios_fondo,capas_raster_drop,modelo_maxent)
str(evalutaion_test)
```

#### Evaluate model with fixed control dataset

```{r evaluate-model-control-data-2, fig.width=10, fig.height=10}
evalutaion_test2 <- evaluate.mdl(puntos_control_extra_coordenadas,puntos_aleatorios_fondo,capas_raster_drop,modelo_maxent)
str(evalutaion_test2)
```


#### Rank model AUC comparing to 99 null-models (with training dataset)

```{r null-model-auc-training}

valores_puntos_aleatorios_fondo_y_entrenamiento <- rbind(evalutaion_training$v_pnt, evalutaion_training$v_bkg)

rep <- 99     # number or null models to produce

null_models <- list()
for (r in 1:rep) {
    # following Robert J. Hijmans nullRandom() function from 'dismo'
		# sample presence records and set the rest of records as absence
		index <- sample(nrow(valores_puntos_aleatorios_fondo_y_entrenamiento), presence_points_count)
		pres <- valores_puntos_aleatorios_fondo_y_entrenamiento[index, ]
		absc <- valores_puntos_aleatorios_fondo_y_entrenamiento[-index, ]
		d <- data.frame(rbind(pres, absc))
		v <- c(rep(1, nrow(pres)), rep(0, nrow(absc)))
		# make null-model with presence and absence sets
		m <- maxent(d, v)
		# evaluate the null-model
		null_models[[r]] <- evaluate(pres, absc, m)
}

#null_models
auc_list_vector <- sapply(null_models, function(x) x@auc)

#auc_list_vector
mean(auc_list_vector)

auc_list_training_vector <- c(evalutaion_training$e@auc, auc_list_vector)
auc_list_training_vector_ranks <- rank(auc_list_training_vector, ties.method= "max")

print('Rank of trainign AUC vs null-models')
auc_list_training_vector_ranks[1]

auc_list_control_vector <- c(evalutaion_test$e@auc, auc_list_vector)
auc_list_control_vector_ranks <- rank(auc_list_control_vector, ties.method= "max")

print('Rank of control AUC vs null-models')
auc_list_control_vector_ranks[1]

```

#### Store the model, parameters and control values in an object for further use

```{r store-result}

container <- list()
container$bkg <- puntos_aleatorios_fondo
container$entr <- puntos_entrenamiento_coordenadas
container$cntrl <- puntos_control_extra_coordenadas
container$model <- modelo_maxent
container$contrib <- contribuciones
container$e_train <- evalutaion_training
container$e_test <- evalutaion_test
container$e_test2 <- evalutaion_test2
container$auc_rank_train <- auc_list_training_vector_ranks[1]
container$auc_rank_test <- auc_list_control_vector_ranks[1]
container$auc_null_model_mean <- mean(auc_list_vector)
  
iterations[[i]] <- container
```
````
