## Supplementary material for "Vulnerability to climate change for narrowly ranged species: the case of Ecuadorian endemic *Magnolia mercedesiarum*": step_29dup_1.html

Cross-validation SDM renderer


### Script for Shalisko et al. 2018 - Cross-validated SDM script 2

###### *Viacheslav Shalisko*

###### *22 of october 2017*

#### Replication 1 of MaxEnt SDM

```
knitr::opts_chunk$set(echo = TRUE)
knitr::opts_chunk$set(error = TRUE)
```

###### Produce and K-fold presence and test points

```
set.seed(random_seed_folding)

random_presence_points <- randomPoints(presence_raster_mask, 
                                        n = double_presence_points_count, 
                                        tryf = 10)

# fold in 2 subsets
fold <- kfold(random_presence_points, k = 2)

# set one subset for training
puntos_entrenamiento_coordenadas <- random_presence_points[ fold == 1, ]
dim(puntos_entrenamiento_coordenadas)
```

```
## [1] 50  2
```

```
# set another subset for initial model evaluation
puntos_control_coordenadas <- random_presence_points[ fold == 2, ]
dim(puntos_control_coordenadas)
```

```
## [1] 50  2
```

###### Produce background points within target mask

```
set.seed(random_seed_background)

# "tryf"" parameter is set to ensure the required number of random points, as the points generated outside
# of the tgs_raster are descarted, examples tryf = 50, tryf = 100
puntos_aleatorios_fondo1 <- randomPoints(tgs_raster_full, 
                                        n = random_background_number, 
                                        p = random_presence_points,
                                        tryf = 10)
puntos_aleatorios_fondo2 <- randomPoints(tgs_raster_500, 
                                        n = random_background_number, 
                                        p = random_presence_points,
                                        tryf = 20)

puntos_aleatorios_fondo <- rbind(puntos_aleatorios_fondo1,puntos_aleatorios_fondo2)
dim(puntos_aleatorios_fondo)
```

```
## [1] 900   2
```

###### Render presencce, test and background data

```
plot(bio10, xlim=c(-82,-70), ylim=c(-10,10), axes=TRUE)
world_high <- getMap(resolution = "high")
plot(world_high, add=TRUE)
# data(wrld_simpl)
# plot(wrld_simpl, add = TRUE)
points(puntos_aleatorios_fondo, cex=0.5, col="black")
points(puntos_entrenamiento_coordenadas,col='red')
points(puntos_control_coordenadas,col='blue',cex=0.7)
```

```
#points(puntos_control_extra_coordenadas,col='navy',cex=0.5)
```

###### Drop unsignificant layers (based on preliminary test run)

```
capas_raster_drop <- dropLayer(capas_raster,layers_to_drop)
names_after_drop <- names(capas_raster_drop)
```

#### Modelling

```
# create maxent output directory
dir.create(paste(basepath, '/maxent_',run_code,'_',i, sep=''), recursive=TRUE)

jar <- paste(system.file(package="dismo"), "/java/maxent.jar", sep='')

modelo_maxent <- maxent(x=capas_raster_drop, 
                        p=puntos_entrenamiento_coordenadas,
                        a=puntos_aleatorios_fondo,
                        path=paste(basepath, '/maxent_',run_code,'_',i, sep=''),
                        args=c(
                          'removeDuplicates=TRUE', 
                          'jackknife=TRUE',
                          'responsecurves=TRUE',
                          'threads=2',
                          'linear=TRUE',
                          'quadratic=TRUE',
                          'hinge=FALSE',
                          'product=FALSE'
                        ))
```

```
## Warning in .local(x, p, ...): 3 (0.33%) of the presence points have NA
## predictor values
```

```
summary(modelo_maxent)
```

```
## Length  Class   Mode 
##      1 MaxEnt     S4
```

```
##str(modelo_maxent)
#representar resultados en forma de una lista
print(modelo_maxent@results)
```

```
##                                                                                        [,1]
## X.Training.samples                                                                  50.0000
## Regularized.training.gain                                                            2.5315
## Unregularized.training.gain                                                          2.6946
## Iterations                                                                         500.0000
## Training.AUC                                                                         0.9734
## X.Background.points                                                                947.0000
## alt.contribution                                                                    11.3772
## bio1.contribution                                                                    0.0000
## bio10.contribution                                                                   0.0086
## bio11.contribution                                                                   1.6198
## bio12.contribution                                                                   0.1341
## bio13.contribution                                                                   0.0000
## bio14.contribution                                                                   0.0020
## bio15.contribution                                                                   4.5821
## bio16.contribution                                                                   0.2305
## bio17.contribution                                                                   0.0000
## bio18.contribution                                                                  10.2022
## bio19.contribution                                                                  23.6364
## bio2.contribution                                                                    0.0047
## bio3.contribution                                                                   20.7343
## bio4.contribution                                                                    1.0226
## bio5.contribution                                                                    1.5748
## bio6.contribution                                                                   24.8708
## bio7.contribution                                                                    0.0000
## bio8.contribution                                                                    0.0000
## bio9.contribution                                                                    0.0000
## alt.permutation.importance                                                          16.6479
## bio1.permutation.importance                                                          0.0000
## bio10.permutation.importance                                                         1.4387
## bio11.permutation.importance                                                         0.0000
## bio12.permutation.importance                                                         6.5220
## bio13.permutation.importance                                                         0.0000
## bio14.permutation.importance                                                         0.0144
## bio15.permutation.importance                                                        16.5831
## bio16.permutation.importance                                                         0.0000
## bio17.permutation.importance                                                         0.0000
## bio18.permutation.importance                                                        21.0382
## bio19.permutation.importance                                                         2.4026
## bio2.permutation.importance                                                          0.0000
## bio3.permutation.importance                                                          1.5921
## bio4.permutation.importance                                                          2.5704
## bio5.permutation.importance                                                          2.6088
## bio6.permutation.importance                                                         28.5817
## bio7.permutation.importance                                                          0.0000
## bio8.permutation.importance                                                          0.0000
## bio9.permutation.importance                                                          0.0000
## Training.gain.without.alt                                                            2.4871
## Training.gain.without.bio1                                                           2.5315
## Training.gain.without.bio10                                                          2.5311
## Training.gain.without.bio11                                                          2.5321
## Training.gain.without.bio12                                                          2.5280
## Training.gain.without.bio13                                                          2.5320
## Training.gain.without.bio14                                                          2.5317
## Training.gain.without.bio15                                                          2.5095
## Training.gain.without.bio16                                                          2.5316
## Training.gain.without.bio17                                                          2.5320
## Training.gain.without.bio18                                                          2.4681
## Training.gain.without.bio19                                                          2.5277
## Training.gain.without.bio2                                                           2.5314
## Training.gain.without.bio3                                                           2.5201
## Training.gain.without.bio4                                                           2.5215
## Training.gain.without.bio5                                                           2.5321
## Training.gain.without.bio6                                                           2.5317
## Training.gain.without.bio7                                                           2.5313
## Training.gain.without.bio8                                                           2.5321
## Training.gain.without.bio9                                                           2.5318
## Training.gain.with.only.alt                                                          1.5831
## Training.gain.with.only.bio1                                                         1.4267
## Training.gain.with.only.bio10                                                        1.4534
## Training.gain.with.only.bio11                                                        1.3877
## Training.gain.with.only.bio12                                                        0.2076
## Training.gain.with.only.bio13                                                        0.0844
## Training.gain.with.only.bio14                                                        1.0050
## Training.gain.with.only.bio15                                                        0.4275
## Training.gain.with.only.bio16                                                        0.0987
## Training.gain.with.only.bio17                                                        0.8987
## Training.gain.with.only.bio18                                                        0.5869
## Training.gain.with.only.bio19                                                        0.3207
## Training.gain.with.only.bio2                                                         0.0752
## Training.gain.with.only.bio3                                                         0.6860
## Training.gain.with.only.bio4                                                         0.8296
## Training.gain.with.only.bio5                                                         1.4111
## Training.gain.with.only.bio6                                                         1.3983
## Training.gain.with.only.bio7                                                         0.1870
## Training.gain.with.only.bio8                                                         1.4387
## Training.gain.with.only.bio9                                                         1.4238
## Entropy                                                                              4.3222
## Prevalence..average.probability.of.presence.over.background.sites.                   0.0467
## Fixed.cumulative.value.1.cumulative.threshold                                        1.0000
## Fixed.cumulative.value.1.Cloglog.threshold                                           0.0164
## Fixed.cumulative.value.1.area                                                        0.1468
## Fixed.cumulative.value.1.training.omission                                           0.0000
## Fixed.cumulative.value.5.cumulative.threshold                                        5.0000
## Fixed.cumulative.value.5.Cloglog.threshold                                           0.1194
## Fixed.cumulative.value.5.area                                                        0.0845
## Fixed.cumulative.value.5.training.omission                                           0.0000
## Fixed.cumulative.value.10.cumulative.threshold                                      10.0000
## Fixed.cumulative.value.10.Cloglog.threshold                                          0.2416
## Fixed.cumulative.value.10.area                                                       0.0612
## Fixed.cumulative.value.10.training.omission                                          0.0000
## Minimum.training.presence.cumulative.threshold                                      12.7182
## Minimum.training.presence.Cloglog.threshold                                          0.3232
## Minimum.training.presence.area                                                       0.0560
## Minimum.training.presence.training.omission                                          0.0000
## X10.percentile.training.presence.cumulative.threshold                               17.5893
## X10.percentile.training.presence.Cloglog.threshold                                   0.4995
## X10.percentile.training.presence.area                                                0.0475
## X10.percentile.training.presence.training.omission                                   0.1000
## Equal.training.sensitivity.and.specificity.cumulative.threshold                     15.8083
## Equal.training.sensitivity.and.specificity.Cloglog.threshold                         0.4104
## Equal.training.sensitivity.and.specificity.area                                      0.0507
## Equal.training.sensitivity.and.specificity.training.omission                         0.0600
## Maximum.training.sensitivity.plus.specificity.cumulative.threshold                  12.7182
## Maximum.training.sensitivity.plus.specificity.Cloglog.threshold                      0.3232
## Maximum.training.sensitivity.plus.specificity.area                                   0.0560
## Maximum.training.sensitivity.plus.specificity.training.omission                      0.0000
## Balance.training.omission..predicted.area.and.threshold.value.cumulative.threshold   1.6460
## Balance.training.omission..predicted.area.and.threshold.value.Cloglog.threshold      0.0313
## Balance.training.omission..predicted.area.and.threshold.value.area                   0.1257
## Balance.training.omission..predicted.area.and.threshold.value.training.omission      0.0000
## Equate.entropy.of.thresholded.and.original.distributions.cumulative.threshold        5.9839
## Equate.entropy.of.thresholded.and.original.distributions.Cloglog.threshold           0.1353
## Equate.entropy.of.thresholded.and.original.distributions.area                        0.0792
## Equate.entropy.of.thresholded.and.original.distributions.training.omission           0.0000
```

```
#str(modelo_maxent@results)
```

###### Get variable importance values

```
#plot(modelo_maxent)

contribuciones <- maxent.select.contribution(modelo_maxent,names_after_drop)

#contribuciones
par(mfrow=c(1, 2))
dotchart(as.numeric(contribuciones[,2]),
        col='blue', pch=16,
        labels=contribuciones[,1],
        main='predictor contribution')
dotchart(as.numeric(contribuciones[,3]),
        col='red', pch=16,
        labels=contribuciones[,1],
        main='predictor importance in permutations')
```

###### Draw respnse cuves

```
response(modelo_maxent)
```

###### Output MaxEnt HTML

```
#modelo_maxent
```

#### Initial model evaluation

###### Evaluate model with training dataset

```
evalutaion_training <- evaluate.mdl(puntos_entrenamiento_coordenadas,puntos_aleatorios_fondo,capas_raster_drop,modelo_maxent)
```

```
## class          : ModelEvaluation 
## n presences    : 50 
## n absences     : 947 
## AUC            : 0.9733685 
## cor            : 0.6575698 
## max TPR+TNR at : 0.3230663
```

```
##                kappa spec_sens no_omission prevalence equal_sens_spec
## thresholds 0.3230663 0.3230663   0.3230663 0.05053335       0.4103319
##            sensitivity
## thresholds   0.4994363
```

```
str(evalutaion_training)
```

```
## List of 4
##  $ e    :Formal class 'ModelEvaluation' [package "dismo"] with 22 slots
##   .. ..@ presence  : num [1:50] 0.898 0.7 0.867 0.879 0.862 ...
##   .. ..@ absence   : num [1:947] 6.11e-08 1.53e-04 2.49e-06 1.88e-05 2.94e-06 ...
##   .. ..@ np        : int 50
##   .. ..@ na        : int 947
##   .. ..@ auc       : num 0.973
##   .. ..@ pauc      : num(0) 
##   .. ..@ cor       : Named num 0.658
##   .. .. ..- attr(*, "names")= chr "cor"
##   .. ..@ pcor      : num 1.66e-124
##   .. ..@ t         : num [1:815] -1e-04 -1e-04 -1e-04 -1e-04 -1e-04 ...
##   .. ..@ confusion : int [1:815, 1:4] 50 50 50 50 50 50 50 50 50 50 ...
##   .. .. ..- attr(*, "dimnames")=List of 2
##   .. .. .. ..$ : NULL
##   .. .. .. ..$ : chr [1:4] "tp" "fp" "fn" "tn"
##   .. ..@ prevalence: num [1:815] 0.0502 0.0502 0.0502 0.0502 0.0502 ...
##   .. ..@ ODP       : num [1:815] 0.95 0.95 0.95 0.95 0.95 ...
##   .. ..@ CCR       : num [1:815] 0.0502 0.0502 0.0502 0.0502 0.0502 ...
##   .. ..@ TPR       : num [1:815] 1 1 1 1 1 1 1 1 1 1 ...
##   .. ..@ TNR       : num [1:815] 0 0 0 0 0 0 0 0 0 0 ...
##   .. ..@ FPR       : num [1:815] 1 1 1 1 1 1 1 1 1 1 ...
##   .. ..@ FNR       : num [1:815] 0 0 0 0 0 0 0 0 0 0 ...
##   .. ..@ PPP       : num [1:815] 0.0502 0.0502 0.0502 0.0502 0.0502 ...
##   .. ..@ NPP       : num [1:815] NaN NaN NaN NaN NaN NaN NaN NaN NaN NaN ...
##   .. ..@ MCR       : num [1:815] 0.95 0.95 0.95 0.95 0.95 ...
##   .. ..@ OR        : num [1:815] NaN NaN NaN NaN NaN NaN NaN NaN NaN NaN ...
##   .. ..@ kappa     : num [1:815] 0 0 0 0 0 0 0 0 0 0 ...
##  $ t    :'data.frame':   1 obs. of  6 variables:
##   ..$ kappa          : num 0.323
##   ..$ spec_sens      : num 0.323
##   ..$ no_omission    : num 0.323
##   ..$ prevalence     : num 0.0505
##   ..$ equal_sens_spec: num 0.41
##   ..$ sensitivity    : num 0.499
##  $ v_bkg: num [1:900, 1:20] 132 192 124 596 2699 ...
##   ..- attr(*, "dimnames")=List of 2
##   .. ..$ : NULL
##   .. ..$ : chr [1:20] "alt" "bio1" "bio2" "bio3" ...
##  $ v_pnt: num [1:50, 1:20] 1982 2078 1894 1930 1907 ...
##   ..- attr(*, "dimnames")=List of 2
##   .. ..$ : NULL
##   .. ..$ : chr [1:20] "alt" "bio1" "bio2" "bio3" ...
```

###### Evaluate model with control dataset

```
evalutaion_test <- evaluate.mdl(puntos_control_coordenadas,puntos_aleatorios_fondo,capas_raster_drop,modelo_maxent)
```

```
## class          : ModelEvaluation 
## n presences    : 50 
## n absences     : 947 
## AUC            : 0.9733052 
## cor            : 0.6595155 
## max TPR+TNR at : 0.280841
```

```
##                kappa spec_sens no_omission prevalence equal_sens_spec
## thresholds 0.3820169  0.280841    0.280841 0.05053335       0.3882545
##            sensitivity
## thresholds   0.5092594
```

```
str(evalutaion_test)
```

```
## List of 4
##  $ e    :Formal class 'ModelEvaluation' [package "dismo"] with 22 slots
##   .. ..@ presence  : num [1:50] 0.727 0.762 0.627 0.816 0.765 ...
##   .. ..@ absence   : num [1:947] 6.11e-08 1.53e-04 2.49e-06 1.88e-05 2.94e-06 ...
##   .. ..@ np        : int 50
##   .. ..@ na        : int 947
##   .. ..@ auc       : num 0.973
##   .. ..@ pauc      : num(0) 
##   .. ..@ cor       : Named num 0.66
##   .. .. ..- attr(*, "names")= chr "cor"
##   .. ..@ pcor      : num 1.75e-125
##   .. ..@ t         : num [1:815] -1e-04 -1e-04 -1e-04 -1e-04 -1e-04 ...
##   .. ..@ confusion : int [1:815, 1:4] 50 50 50 50 50 50 50 50 50 50 ...
##   .. .. ..- attr(*, "dimnames")=List of 2
##   .. .. .. ..$ : NULL
##   .. .. .. ..$ : chr [1:4] "tp" "fp" "fn" "tn"
##   .. ..@ prevalence: num [1:815] 0.0502 0.0502 0.0502 0.0502 0.0502 ...
##   .. ..@ ODP       : num [1:815] 0.95 0.95 0.95 0.95 0.95 ...
##   .. ..@ CCR       : num [1:815] 0.0502 0.0502 0.0502 0.0502 0.0502 ...
##   .. ..@ TPR       : num [1:815] 1 1 1 1 1 1 1 1 1 1 ...
##   .. ..@ TNR       : num [1:815] 0 0 0 0 0 0 0 0 0 0 ...
##   .. ..@ FPR       : num [1:815] 1 1 1 1 1 1 1 1 1 1 ...
##   .. ..@ FNR       : num [1:815] 0 0 0 0 0 0 0 0 0 0 ...
##   .. ..@ PPP       : num [1:815] 0.0502 0.0502 0.0502 0.0502 0.0502 ...
##   .. ..@ NPP       : num [1:815] NaN NaN NaN NaN NaN NaN NaN NaN NaN NaN ...
##   .. ..@ MCR       : num [1:815] 0.95 0.95 0.95 0.95 0.95 ...
##   .. ..@ OR        : num [1:815] NaN NaN NaN NaN NaN NaN NaN NaN NaN NaN ...
##   .. ..@ kappa     : num [1:815] 0 0 0 0 0 0 0 0 0 0 ...
##  $ t    :'data.frame':   1 obs. of  6 variables:
##   ..$ kappa          : num 0.382
##   ..$ spec_sens      : num 0.281
##   ..$ no_omission    : num 0.281
##   ..$ prevalence     : num 0.0505
##   ..$ equal_sens_spec: num 0.388
##   ..$ sensitivity    : num 0.509
##  $ v_bkg: num [1:900, 1:20] 132 192 124 596 2699 ...
##   ..- attr(*, "dimnames")=List of 2
##   .. ..$ : NULL
##   .. ..$ : chr [1:20] "alt" "bio1" "bio2" "bio3" ...
##  $ v_pnt: num [1:50, 1:20] 1956 1819 2225 1815 2039 ...
##   ..- attr(*, "dimnames")=List of 2
##   .. ..$ : NULL
##   .. ..$ : chr [1:20] "alt" "bio1" "bio2" "bio3" ...
```

###### Evaluate model with fixed control dataset

```
evalutaion_test2 <- evaluate.mdl(puntos_control_extra_coordenadas,puntos_aleatorios_fondo,capas_raster_drop,modelo_maxent)
```

```
## class          : ModelEvaluation 
## n presences    : 72 
## n absences     : 969 
## AUC            : 0.9624183 
## cor            : 0.6441127 
## max TPR+TNR at : 0.4068761
```

```
##                kappa spec_sens no_omission prevalence equal_sens_spec
## thresholds 0.4068761 0.4068761   0.4068761 0.06996256       0.4422971
##            sensitivity
## thresholds   0.4465707
```

```
str(evalutaion_test2)
```

```
## List of 4
##  $ e    :Formal class 'ModelEvaluation' [package "dismo"] with 22 slots
##   .. ..@ presence  : num [1:72] 0.522 0.467 0.6 0.863 0.6 ...
##   .. ..@ absence   : num [1:969] 6.11e-08 1.53e-04 2.49e-06 1.88e-05 2.94e-06 ...
##   .. ..@ np        : int 72
##   .. ..@ na        : int 969
##   .. ..@ auc       : num 0.962
##   .. ..@ pauc      : num(0) 
##   .. ..@ cor       : Named num 0.644
##   .. .. ..- attr(*, "names")= chr "cor"
##   .. ..@ pcor      : num 4.65e-123
##   .. ..@ t         : num [1:823] -1e-04 -1e-04 -1e-04 -1e-04 -1e-04 ...
##   .. ..@ confusion : int [1:823, 1:4] 72 72 72 72 72 72 72 72 72 72 ...
##   .. .. ..- attr(*, "dimnames")=List of 2
##   .. .. .. ..$ : NULL
##   .. .. .. ..$ : chr [1:4] "tp" "fp" "fn" "tn"
##   .. ..@ prevalence: num [1:823] 0.0692 0.0692 0.0692 0.0692 0.0692 ...
##   .. ..@ ODP       : num [1:823] 0.931 0.931 0.931 0.931 0.931 ...
##   .. ..@ CCR       : num [1:823] 0.0692 0.0692 0.0692 0.0692 0.0692 ...
##   .. ..@ TPR       : num [1:823] 1 1 1 1 1 1 1 1 1 1 ...
##   .. ..@ TNR       : num [1:823] 0 0 0 0 0 0 0 0 0 0 ...
##   .. ..@ FPR       : num [1:823] 1 1 1 1 1 1 1 1 1 1 ...
##   .. ..@ FNR       : num [1:823] 0 0 0 0 0 0 0 0 0 0 ...
##   .. ..@ PPP       : num [1:823] 0.0692 0.0692 0.0692 0.0692 0.0692 ...
##   .. ..@ NPP       : num [1:823] NaN NaN NaN NaN NaN NaN NaN NaN NaN NaN ...
##   .. ..@ MCR       : num [1:823] 0.931 0.931 0.931 0.931 0.931 ...
##   .. ..@ OR        : num [1:823] NaN NaN NaN NaN NaN NaN NaN NaN NaN NaN ...
##   .. ..@ kappa     : num [1:823] 0 0 0 0 0 0 0 0 0 0 ...
##  $ t    :'data.frame':   1 obs. of  6 variables:
##   ..$ kappa          : num 0.407
##   ..$ spec_sens      : num 0.407
##   ..$ no_omission    : num 0.407
##   ..$ prevalence     : num 0.07
##   ..$ equal_sens_spec: num 0.442
##   ..$ sensitivity    : num 0.447
##  $ v_bkg: num [1:900, 1:20] 132 192 124 596 2699 ...
##   ..- attr(*, "dimnames")=List of 2
##   .. ..$ : NULL
##   .. ..$ : chr [1:20] "alt" "bio1" "bio2" "bio3" ...
##  $ v_pnt: num [1:72, 1:20] 1252 1946 1793 1871 1793 ...
##   ..- attr(*, "dimnames")=List of 2
##   .. ..$ : NULL
##   .. ..$ : chr [1:20] "alt" "bio1" "bio2" "bio3" ...
```

###### Rank model AUC comparing to 99 null-models (with training dataset)

```
valores_puntos_aleatorios_fondo_y_entrenamiento <- rbind(evalutaion_training$v_pnt, evalutaion_training$v_bkg)

rep <- 99     # number or null models to produce

null_models <- list()
for (r in 1:rep) {
    # following Robert J. Hijmans nullRandom() function from 'dismo'
        # sample presence records and set the rest of records as absence
        index <- sample(nrow(valores_puntos_aleatorios_fondo_y_entrenamiento), presence_points_count)
        pres <- valores_puntos_aleatorios_fondo_y_entrenamiento[index, ]
        absc <- valores_puntos_aleatorios_fondo_y_entrenamiento[-index, ]
        d <- data.frame(rbind(pres, absc))
        v <- c(rep(1, nrow(pres)), rep(0, nrow(absc)))
        # make null-model with presence and absence sets
        m <- maxent(d, v)
        # evaluate the null-model
        null_models[[r]] <- evaluate(pres, absc, m)
}

#null_models
auc_list_vector <- sapply(null_models, function(x) x@auc)

#auc_list_vector
mean(auc_list_vector)
```

```
## [1] 0.726552
```

```
auc_list_training_vector <- c(evalutaion_training$e@auc, auc_list_vector)
auc_list_training_vector_ranks <- rank(auc_list_training_vector, ties.method= "max")

print('Rank of trainign AUC vs null-models')
```

```
## [1] "Rank of trainign AUC vs null-models"
```

```
auc_list_training_vector_ranks[1]
```

```
## [1] 100
```

```
auc_list_control_vector <- c(evalutaion_test$e@auc, auc_list_vector)
auc_list_control_vector_ranks <- rank(auc_list_control_vector, ties.method= "max")

print('Rank of control AUC vs null-models')
```

```
## [1] "Rank of control AUC vs null-models"
```

```
auc_list_control_vector_ranks[1]
```

```
## [1] 100
```

###### Store the model, parameters and control values in an object for further use

```
container <- list()
container$bkg <- puntos_aleatorios_fondo
container$entr <- puntos_entrenamiento_coordenadas
container$cntrl <- puntos_control_extra_coordenadas
container$model <- modelo_maxent
container$contrib <- contribuciones
container$e_train <- evalutaion_training
container$e_test <- evalutaion_test
container$e_test2 <- evalutaion_test2
container$auc_rank_train <- auc_list_training_vector_ranks[1]
container$auc_rank_test <- auc_list_control_vector_ranks[1]
container$auc_null_model_mean <- mean(auc_list_vector)
  
iterations[[i]] <- container
```
