## Supplementary material for "Vulnerability to climate change for narrowly ranged species: the case of Ecuadorian endemic *Magnolia mercedesiarum*": step_29dup_2.html

```
##                                                                                        [,1]
## X.Training.samples                                                                  50.0000
## Regularized.training.gain                                                            2.4698
## Unregularized.training.gain                                                          2.6472
## Iterations                                                                         500.0000
## Training.AUC                                                                         0.9723
## X.Background.points                                                                947.0000
## alt.contribution                                                                    11.0953
## bio1.contribution                                                                    0.0056
## bio10.contribution                                                                   6.3227
## bio11.contribution                                                                   0.2213
## bio12.contribution                                                                   0.5314
## bio13.contribution                                                                   0.0000
## bio14.contribution                                                                   0.0921
## bio15.contribution                                                                   4.2802
## bio16.contribution                                                                   0.1022
## bio17.contribution                                                                   0.0045
## bio18.contribution                                                                  12.1058
## bio19.contribution                                                                  25.3079
## bio2.contribution                                                                    0.0205
## bio3.contribution                                                                   17.6185
## bio4.contribution                                                                    0.0165
## bio5.contribution                                                                    1.6913
## bio6.contribution                                                                   19.0765
## bio7.contribution                                                                    0.0000
## bio8.contribution                                                                    1.5077
## bio9.contribution                                                                    0.0000
## alt.permutation.importance                                                          18.6947
## bio1.permutation.importance                                                          0.0000
## bio10.permutation.importance                                                        32.0622
## bio11.permutation.importance                                                         0.0000
## bio12.permutation.importance                                                        12.4631
## bio13.permutation.importance                                                         0.0000
## bio14.permutation.importance                                                         0.0000
## bio15.permutation.importance                                                         6.5857
## bio16.permutation.importance                                                         0.5139
## bio17.permutation.importance                                                         0.0593
## bio18.permutation.importance                                                        22.0798
## bio19.permutation.importance                                                         0.0000
## bio2.permutation.importance                                                          0.3888
## bio3.permutation.importance                                                          1.2618
## bio4.permutation.importance                                                          0.2372
## bio5.permutation.importance                                                          5.6534
## bio6.permutation.importance                                                          0.0000
## bio7.permutation.importance                                                          0.0000
## bio8.permutation.importance                                                          0.0000
## bio9.permutation.importance                                                          0.0000
## Training.gain.without.alt                                                            2.4113
## Training.gain.without.bio1                                                           2.4699
## Training.gain.without.bio10                                                          2.4677
## Training.gain.without.bio11                                                          2.4699
## Training.gain.without.bio12                                                          2.4678
## Training.gain.without.bio13                                                          2.4691
## Training.gain.without.bio14                                                          2.4693
## Training.gain.without.bio15                                                          2.4578
## Training.gain.without.bio16                                                          2.4693
## Training.gain.without.bio17                                                          2.4700
## Training.gain.without.bio18                                                          2.3611
## Training.gain.without.bio19                                                          2.4696
## Training.gain.without.bio2                                                           2.4666
## Training.gain.without.bio3                                                           2.4536
## Training.gain.without.bio4                                                           2.4689
## Training.gain.without.bio5                                                           2.4698
## Training.gain.without.bio6                                                           2.4701
## Training.gain.without.bio7                                                           2.4695
## Training.gain.without.bio8                                                           2.4695
## Training.gain.without.bio9                                                           2.4693
## Training.gain.with.only.alt                                                          1.4642
## Training.gain.with.only.bio1                                                         1.3165
## Training.gain.with.only.bio10                                                        1.3530
## Training.gain.with.only.bio11                                                        1.2609
## Training.gain.with.only.bio12                                                        0.2095
## Training.gain.with.only.bio13                                                        0.1097
## Training.gain.with.only.bio14                                                        0.9531
## Training.gain.with.only.bio15                                                        0.3697
## Training.gain.with.only.bio16                                                        0.1126
## Training.gain.with.only.bio17                                                        0.8778
## Training.gain.with.only.bio18                                                        0.5753
## Training.gain.with.only.bio19                                                        0.3151
## Training.gain.with.only.bio2                                                         0.0263
## Training.gain.with.only.bio3                                                         0.5678
## Training.gain.with.only.bio4                                                         0.7042
## Training.gain.with.only.bio5                                                         1.3487
## Training.gain.with.only.bio6                                                         1.2392
## Training.gain.with.only.bio7                                                         0.1363
## Training.gain.with.only.bio8                                                         1.3503
## Training.gain.with.only.bio9                                                         1.2919
## Entropy                                                                              4.3836
## Prevalence..average.probability.of.presence.over.background.sites.                   0.0490
## Fixed.cumulative.value.1.cumulative.threshold                                        1.0000
## Fixed.cumulative.value.1.Cloglog.threshold                                           0.0077
## Fixed.cumulative.value.1.area                                                        0.2101
## Fixed.cumulative.value.1.training.omission                                           0.0000
## Fixed.cumulative.value.5.cumulative.threshold                                        5.0000
## Fixed.cumulative.value.5.Cloglog.threshold                                           0.1072
## Fixed.cumulative.value.5.area                                                        0.0898
## Fixed.cumulative.value.5.training.omission                                           0.0000
## Fixed.cumulative.value.10.cumulative.threshold                                      10.0000
## Fixed.cumulative.value.10.Cloglog.threshold                                          0.2604
## Fixed.cumulative.value.10.area                                                       0.0644
## Fixed.cumulative.value.10.training.omission                                          0.0000
## Minimum.training.presence.cumulative.threshold                                      13.4971
## Minimum.training.presence.Cloglog.threshold                                          0.3446
## Minimum.training.presence.area                                                       0.0560
## Minimum.training.presence.training.omission                                          0.0000
## X10.percentile.training.presence.cumulative.threshold                               18.1687
## X10.percentile.training.presence.Cloglog.threshold                                   0.4659
## X10.percentile.training.presence.area                                                0.0486
## X10.percentile.training.presence.training.omission                                   0.1000
## Equal.training.sensitivity.and.specificity.cumulative.threshold                     15.2422
## Equal.training.sensitivity.and.specificity.Cloglog.threshold                         0.3869
## Equal.training.sensitivity.and.specificity.area                                      0.0539
## Equal.training.sensitivity.and.specificity.training.omission                         0.0600
## Maximum.training.sensitivity.plus.specificity.cumulative.threshold                  13.4971
## Maximum.training.sensitivity.plus.specificity.Cloglog.threshold                      0.3446
## Maximum.training.sensitivity.plus.specificity.area                                   0.0560
## Maximum.training.sensitivity.plus.specificity.training.omission                      0.0000
## Balance.training.omission..predicted.area.and.threshold.value.cumulative.threshold   2.7570
## Balance.training.omission..predicted.area.and.threshold.value.Cloglog.threshold      0.0323
## Balance.training.omission..predicted.area.and.threshold.value.area                   0.1225
## Balance.training.omission..predicted.area.and.threshold.value.training.omission      0.0000
## Equate.entropy.of.thresholded.and.original.distributions.cumulative.threshold        5.7970
## Equate.entropy.of.thresholded.and.original.distributions.Cloglog.threshold           0.1202
## Equate.entropy.of.thresholded.and.original.distributions.area                        0.0845
## Equate.entropy.of.thresholded.and.original.distributions.training.omission           0.0000
```

```
#str(modelo_maxent@results)
```

##### Get variable importance values

```
#plot(modelo_maxent)

contribuciones <- maxent.select.contribution(modelo_maxent,names_after_drop)

#contribuciones
par(mfrow=c(1, 2))
dotchart(as.numeric(contribuciones[,2]),
        col='blue', pch=16,
        labels=contribuciones[,1],
        main='predictor contribution')
dotchart(as.numeric(contribuciones[,3]),
        col='red', pch=16,
        labels=contribuciones[,1],
        main='predictor importance in permutations')
```

##### Draw respnse cuves

```
response(modelo_maxent)
```

##### Output MaxEnt HTML

```
#modelo_maxent
```

### Initial model evaluation

##### Evaluate model with training dataset

```
evalutaion_training <- evaluate.mdl(puntos_entrenamiento_coordenadas,puntos_aleatorios_fondo,capas_raster_drop,modelo_maxent)
```

```
## class          : ModelEvaluation 
## n presences    : 50 
## n absences     : 947 
## AUC            : 0.9722703 
## cor            : 0.6513998 
## max TPR+TNR at : 0.344535
```

```
##               kappa spec_sens no_omission prevalence equal_sens_spec
## thresholds 0.344535  0.344535    0.344535 0.04948492       0.3867882
##            sensitivity
## thresholds   0.4549082
```

```
str(evalutaion_training)
```

```
## List of 4
##  $ e    :Formal class 'ModelEvaluation' [package "dismo"] with 22 slots
##   .. ..@ presence  : num [1:50] 0.636 0.793 0.618 0.704 0.685 ...
##   .. ..@ absence   : num [1:947] 4.27e-05 2.11e-03 7.01e-05 8.17e-03 9.25e-04 ...
##   .. ..@ np        : int 50
##   .. ..@ na        : int 947
##   .. ..@ auc       : num 0.972
##   .. ..@ pauc      : num(0) 
##   .. ..@ cor       : Named num 0.651
##   .. .. ..- attr(*, "names")= chr "cor"
##   .. ..@ pcor      : num 1.9e-121
##   .. ..@ t         : num [1:875] -1e-04 -1e-04 -1e-04 -1e-04 -1e-04 ...
##   .. ..@ confusion : int [1:875, 1:4] 50 50 50 50 50 50 50 50 50 50 ...
##   .. .. ..- attr(*, "dimnames")=List of 2
##   .. .. .. ..$ : NULL
##   .. .. .. ..$ : chr [1:4] "tp" "fp" "fn" "tn"
##   .. ..@ prevalence: num [1:875] 0.0502 0.0502 0.0502 0.0502 0.0502 ...
##   .. ..@ ODP       : num [1:875] 0.95 0.95 0.95 0.95 0.95 ...
##   .. ..@ CCR       : num [1:875] 0.0502 0.0502 0.0502 0.0502 0.0502 ...
##   .. ..@ TPR       : num [1:875] 1 1 1 1 1 1 1 1 1 1 ...
##   .. ..@ TNR       : num [1:875] 0 0 0 0 0 0 0 0 0 0 ...
##   .. ..@ FPR       : num [1:875] 1 1 1 1 1 1 1 1 1 1 ...
##   .. ..@ FNR       : num [1:875] 0 0 0 0 0 0 0 0 0 0 ...
##   .. ..@ PPP       : num [1:875] 0.0502 0.0502 0.0502 0.0502 0.0502 ...
##   .. ..@ NPP       : num [1:875] NaN NaN NaN NaN NaN NaN NaN NaN NaN NaN ...
##   .. ..@ MCR       : num [1:875] 0.95 0.95 0.95 0.95 0.95 ...
##   .. ..@ OR        : num [1:875] NaN NaN NaN NaN NaN NaN NaN NaN NaN NaN ...
##   .. ..@ kappa     : num [1:875] 0 0 0 0 0 0 0 0 0 0 ...
##  $ t    :'data.frame':   1 obs. of  6 variables:
##   ..$ kappa          : num 0.345
##   ..$ spec_sens      : num 0.345
##   ..$ no_omission    : num 0.345
##   ..$ prevalence     : num 0.0495
##   ..$ equal_sens_spec: num 0.387
##   ..$ sensitivity    : num 0.455
##  $ v_bkg: num [1:900, 1:20] 11 201 161 1984 288 ...
##   ..- attr(*, "dimnames")=List of 2
##   .. ..$ : NULL
##   .. ..$ : chr [1:20] "alt" "bio1" "bio2" "bio3" ...
##  $ v_pnt: num [1:50, 1:20] 2039 1674 1280 1568 1487 ...
##   ..- attr(*, "dimnames")=List of 2
##   .. ..$ : NULL
##   .. ..$ : chr [1:20] "alt" "bio1" "bio2" "bio3" ...
```

##### Evaluate model with control dataset

```
evalutaion_test <- evaluate.mdl(puntos_control_coordenadas,puntos_aleatorios_fondo,capas_raster_drop,modelo_maxent)
```

```
## class          : ModelEvaluation 
## n presences    : 50 
## n absences     : 947 
## AUC            : 0.9723126 
## cor            : 0.6535875 
## max TPR+TNR at : 0.2857366
```

```
##                kappa spec_sens no_omission prevalence equal_sens_spec
## thresholds 0.3329513 0.2857366   0.2857366 0.04948492       0.4549082
##            sensitivity
## thresholds   0.5265526
```

```
str(evalutaion_test)
```

```
## List of 4
##  $ e    :Formal class 'ModelEvaluation' [package "dismo"] with 22 slots
##   .. ..@ presence  : num [1:50] 0.333 0.837 0.817 0.286 0.773 ...
##   .. ..@ absence   : num [1:947] 4.27e-05 2.11e-03 7.01e-05 8.17e-03 9.25e-04 ...
##   .. ..@ np        : int 50
##   .. ..@ na        : int 947
##   .. ..@ auc       : num 0.972
##   .. ..@ pauc      : num(0) 
##   .. ..@ cor       : Named num 0.654
##   .. .. ..- attr(*, "names")= chr "cor"
##   .. ..@ pcor      : num 1.59e-122
##   .. ..@ t         : num [1:875] -1e-04 -1e-04 -1e-04 -1e-04 -1e-04 ...
##   .. ..@ confusion : int [1:875, 1:4] 50 50 50 50 50 50 50 50 50 50 ...
##   .. .. ..- attr(*, "dimnames")=List of 2
##   .. .. .. ..$ : NULL
##   .. .. .. ..$ : chr [1:4] "tp" "fp" "fn" "tn"
##   .. ..@ prevalence: num [1:875] 0.0502 0.0502 0.0502 0.0502 0.0502 ...
##   .. ..@ ODP       : num [1:875] 0.95 0.95 0.95 0.95 0.95 ...
##   .. ..@ CCR       : num [1:875] 0.0502 0.0502 0.0502 0.0502 0.0502 ...
##   .. ..@ TPR       : num [1:875] 1 1 1 1 1 1 1 1 1 1 ...
##   .. ..@ TNR       : num [1:875] 0 0 0 0 0 0 0 0 0 0 ...
##   .. ..@ FPR       : num [1:875] 1 1 1 1 1 1 1 1 1 1 ...
##   .. ..@ FNR       : num [1:875] 0 0 0 0 0 0 0 0 0 0 ...
##   .. ..@ PPP       : num [1:875] 0.0502 0.0502 0.0502 0.0502 0.0502 ...
##   .. ..@ NPP       : num [1:875] NaN NaN NaN NaN NaN NaN NaN NaN NaN NaN ...
##   .. ..@ MCR       : num [1:875] 0.95 0.95 0.95 0.95 0.95 ...
##   .. ..@ OR        : num [1:875] NaN NaN NaN NaN NaN NaN NaN NaN NaN NaN ...
##   .. ..@ kappa     : num [1:875] 0 0 0 0 0 0 0 0 0 0 ...
##  $ t    :'data.frame':   1 obs. of  6 variables:
##   ..$ kappa          : num 0.333
##   ..$ spec_sens      : num 0.286
##   ..$ no_omission    : num 0.286
##   ..$ prevalence     : num 0.0495
##   ..$ equal_sens_spec: num 0.455
##   ..$ sensitivity    : num 0.527
##  $ v_bkg: num [1:900, 1:20] 11 201 161 1984 288 ...
##   ..- attr(*, "dimnames")=List of 2
##   .. ..$ : NULL
##   .. ..$ : chr [1:20] "alt" "bio1" "bio2" "bio3" ...
##  $ v_pnt: num [1:50, 1:20] 2354 1657 1478 2216 1953 ...
##   ..- attr(*, "dimnames")=List of 2
##   .. ..$ : NULL
##   .. ..$ : chr [1:20] "alt" "bio1" "bio2" "bio3" ...
```

##### Evaluate model with fixed control dataset

```
evalutaion_test2 <- evaluate.mdl(puntos_control_extra_coordenadas,puntos_aleatorios_fondo,capas_raster_drop,modelo_maxent)
```

```
## class          : ModelEvaluation 
## n presences    : 72 
## n absences     : 969 
## AUC            : 0.961673 
## cor            : 0.6502098 
## max TPR+TNR at : 0.3811137
```

```
##               kappa spec_sens no_omission prevalence equal_sens_spec
## thresholds 0.401993 0.3811137   0.3811137  0.0658954       0.4895669
##            sensitivity
## thresholds     0.52465
```

```
str(evalutaion_test2)
```

```
## List of 4
##  $ e    :Formal class 'ModelEvaluation' [package "dismo"] with 22 slots
##   .. ..@ presence  : num [1:72] 0.552 0.55 0.678 0.815 0.678 ...
##   .. ..@ absence   : num [1:969] 4.27e-05 2.11e-03 7.01e-05 8.17e-03 9.25e-04 ...
##   .. ..@ np        : int 72
##   .. ..@ na        : int 969
##   .. ..@ auc       : num 0.962
##   .. ..@ pauc      : num(0) 
##   .. ..@ cor       : Named num 0.65
##   .. .. ..- attr(*, "names")= chr "cor"
##   .. ..@ pcor      : num 3.98e-126
##   .. ..@ t         : num [1:883] -1e-04 -1e-04 -1e-04 -1e-04 -1e-04 ...
##   .. ..@ confusion : int [1:883, 1:4] 72 72 72 72 72 72 72 72 72 72 ...
##   .. .. ..- attr(*, "dimnames")=List of 2
##   .. .. .. ..$ : NULL
##   .. .. .. ..$ : chr [1:4] "tp" "fp" "fn" "tn"
##   .. ..@ prevalence: num [1:883] 0.0692 0.0692 0.0692 0.0692 0.0692 ...
##   .. ..@ ODP       : num [1:883] 0.931 0.931 0.931 0.931 0.931 ...
##   .. ..@ CCR       : num [1:883] 0.0692 0.0692 0.0692 0.0692 0.0692 ...
##   .. ..@ TPR       : num [1:883] 1 1 1 1 1 1 1 1 1 1 ...
##   .. ..@ TNR       : num [1:883] 0 0 0 0 0 0 0 0 0 0 ...
##   .. ..@ FPR       : num [1:883] 1 1 1 1 1 1 1 1 1 1 ...
##   .. ..@ FNR       : num [1:883] 0 0 0 0 0 0 0 0 0 0 ...
##   .. ..@ PPP       : num [1:883] 0.0692 0.0692 0.0692 0.0692 0.0692 ...
##   .. ..@ NPP       : num [1:883] NaN NaN NaN NaN NaN NaN NaN NaN NaN NaN ...
##   .. ..@ MCR       : num [1:883] 0.931 0.931 0.931 0.931 0.931 ...
##   .. ..@ OR        : num [1:883] NaN NaN NaN NaN NaN NaN NaN NaN NaN NaN ...
##   .. ..@ kappa     : num [1:883] 0 0 0 0 0 0 0 0 0 0 ...
##  $ t    :'data.frame':   1 obs. of  6 variables:
##   ..$ kappa          : num 0.402
##   ..$ spec_sens      : num 0.381
##   ..$ no_omission    : num 0.381
##   ..$ prevalence     : num 0.0659
##   ..$ equal_sens_spec: num 0.49
##   ..$ sensitivity    : num 0.525
##  $ v_bkg: num [1:900, 1:20] 11 201 161 1984 288 ...
##   ..- attr(*, "dimnames")=List of 2
##   .. ..$ : NULL
##   .. ..$ : chr [1:20] "alt" "bio1" "bio2" "bio3" ...
##  $ v_pnt: num [1:72, 1:20] 1252 1946 1793 1871 1793 ...
##   ..- attr(*, "dimnames")=List of 2
##   .. ..$ : NULL
##   .. ..$ : chr [1:20] "alt" "bio1" "bio2" "bio3" ...
```

##### Rank model AUC comparing to 99 null-models (with training dataset)

```
valores_puntos_aleatorios_fondo_y_entrenamiento <- rbind(evalutaion_training$v_pnt, evalutaion_training$v_bkg)

rep <- 99     # number or null models to produce

null_models <- list()
for (r in 1:rep) {
    # following Robert J. Hijmans nullRandom() function from 'dismo'
        # sample presence records and set the rest of records as absence
        index <- sample(nrow(valores_puntos_aleatorios_fondo_y_entrenamiento), presence_points_count)
        pres <- valores_puntos_aleatorios_fondo_y_entrenamiento[index, ]
        absc <- valores_puntos_aleatorios_fondo_y_entrenamiento[-index, ]
        d <- data.frame(rbind(pres, absc))
        v <- c(rep(1, nrow(pres)), rep(0, nrow(absc)))
        # make null-model with presence and absence sets
        m <- maxent(d, v)
        # evaluate the null-model
        null_models[[r]] <- evaluate(pres, absc, m)
}

#null_models
auc_list_vector <- sapply(null_models, function(x) x@auc)

#auc_list_vector
mean(auc_list_vector)
```

```
## [1] 0.7266066
```

```
auc_list_training_vector <- c(evalutaion_training$e@auc, auc_list_vector)
auc_list_training_vector_ranks <- rank(auc_list_training_vector, ties.method= "max")

print('Rank of trainign AUC vs null-models')
```

```
## [1] "Rank of trainign AUC vs null-models"
```

```
auc_list_training_vector_ranks[1]
```

```
## [1] 100
```

```
auc_list_control_vector <- c(evalutaion_test$e@auc, auc_list_vector)
auc_list_control_vector_ranks <- rank(auc_list_control_vector, ties.method= "max")

print('Rank of control AUC vs null-models')
```

```
## [1] "Rank of control AUC vs null-models"
```

```
auc_list_control_vector_ranks[1]
```

```
## [1] 100
```

##### Store the model, parameters and control values in an object for further use

```
container <- list()
container$bkg <- puntos_aleatorios_fondo
container$entr <- puntos_entrenamiento_coordenadas
container$cntrl <- puntos_control_extra_coordenadas
container$model <- modelo_maxent
container$contrib <- contribuciones
container$e_train <- evalutaion_training
container$e_test <- evalutaion_test
container$e_test2 <- evalutaion_test2
container$auc_rank_train <- auc_list_training_vector_ranks[1]
container$auc_rank_test <- auc_list_control_vector_ranks[1]
container$auc_null_model_mean <- mean(auc_list_vector)
  
iterations[[i]] <- container
```
