## Supplementary material for "Vulnerability to climate change for narrowly ranged species: the case of Ecuadorian endemic *Magnolia mercedesiarum*": step_29dup_3.html

```
## Warning in .local(x, p, ...): 1 (0.11%) of the presence points have NA
## predictor values
```

```
summary(modelo_maxent)
```

```
## Length  Class   Mode 
##      1 MaxEnt     S4
```

```
##str(modelo_maxent)
#representar resultados en forma de una lista
print(modelo_maxent@results)
```

```
##                                                                                        [,1]
## X.Training.samples                                                                  50.0000
## Regularized.training.gain                                                            2.4947
## Unregularized.training.gain                                                          2.6663
## Iterations                                                                         340.0000
## Training.AUC                                                                         0.9710
## X.Background.points                                                                949.0000
## alt.contribution                                                                     8.5996
## bio1.contribution                                                                    0.0000
## bio10.contribution                                                                   0.0002
## bio11.contribution                                                                   1.7699
## bio12.contribution                                                                   0.3922
## bio13.contribution                                                                   0.0093
## bio14.contribution                                                                   0.0285
## bio15.contribution                                                                   3.9715
## bio16.contribution                                                                   0.0772
## bio17.contribution                                                                   0.0000
## bio18.contribution                                                                   7.1097
## bio19.contribution                                                                  23.9996
## bio2.contribution                                                                    0.0021
## bio3.contribution                                                                   28.7356
## bio4.contribution                                                                    0.0362
## bio5.contribution                                                                    0.0790
## bio6.contribution                                                                   25.1894
## bio7.contribution                                                                    0.0000
## bio8.contribution                                                                    0.0000
## bio9.contribution                                                                    0.0000
## alt.permutation.importance                                                          14.2503
## bio1.permutation.importance                                                          0.0000
## bio10.permutation.importance                                                         0.0185
## bio11.permutation.importance                                                         0.0000
## bio12.permutation.importance                                                        11.7200
## bio13.permutation.importance                                                         0.0000
## bio14.permutation.importance                                                         0.0000
## bio15.permutation.importance                                                         5.8530
## bio16.permutation.importance                                                         0.0000
## bio17.permutation.importance                                                         0.0000
## bio18.permutation.importance                                                        16.1758
## bio19.permutation.importance                                                         0.0000
## bio2.permutation.importance                                                          0.0000
## bio3.permutation.importance                                                          5.9087
## bio4.permutation.importance                                                          0.0000
## bio5.permutation.importance                                                          2.8501
## bio6.permutation.importance                                                         43.2236
## bio7.permutation.importance                                                          0.0000
## bio8.permutation.importance                                                          0.0000
## bio9.permutation.importance                                                          0.0000
## Training.gain.without.alt                                                            2.4427
## Training.gain.without.bio1                                                           2.4948
## Training.gain.without.bio10                                                          2.4945
## Training.gain.without.bio11                                                          2.4948
## Training.gain.without.bio12                                                          2.4779
## Training.gain.without.bio13                                                          2.4946
## Training.gain.without.bio14                                                          2.4948
## Training.gain.without.bio15                                                          2.4837
## Training.gain.without.bio16                                                          2.4945
## Training.gain.without.bio17                                                          2.4947
## Training.gain.without.bio18                                                          2.4509
## Training.gain.without.bio19                                                          2.4936
## Training.gain.without.bio2                                                           2.4946
## Training.gain.without.bio3                                                           2.4657
## Training.gain.without.bio4                                                           2.4940
## Training.gain.without.bio5                                                           2.4951
## Training.gain.without.bio6                                                           2.4914
## Training.gain.without.bio7                                                           2.4938
## Training.gain.without.bio8                                                           2.4942
## Training.gain.without.bio9                                                           2.4936
## Training.gain.with.only.alt                                                          1.5122
## Training.gain.with.only.bio1                                                         1.3996
## Training.gain.with.only.bio10                                                        1.4287
## Training.gain.with.only.bio11                                                        1.3564
## Training.gain.with.only.bio12                                                        0.1816
## Training.gain.with.only.bio13                                                        0.0812
## Training.gain.with.only.bio14                                                        0.9620
## Training.gain.with.only.bio15                                                        0.4166
## Training.gain.with.only.bio16                                                        0.0938
## Training.gain.with.only.bio17                                                        0.9033
## Training.gain.with.only.bio18                                                        0.4803
## Training.gain.with.only.bio19                                                        0.3375
## Training.gain.with.only.bio2                                                         0.0362
## Training.gain.with.only.bio3                                                         0.8542
## Training.gain.with.only.bio4                                                         0.9140
## Training.gain.with.only.bio5                                                         1.4161
## Training.gain.with.only.bio6                                                         1.3519
## Training.gain.with.only.bio7                                                         0.1599
## Training.gain.with.only.bio8                                                         1.4209
## Training.gain.with.only.bio9                                                         1.3767
## Entropy                                                                              4.3617
## Prevalence..average.probability.of.presence.over.background.sites.                   0.0488
## Fixed.cumulative.value.1.cumulative.threshold                                        1.0000
## Fixed.cumulative.value.1.Cloglog.threshold                                           0.0109
## Fixed.cumulative.value.1.area                                                        0.1581
## Fixed.cumulative.value.1.training.omission                                           0.0000
## Fixed.cumulative.value.5.cumulative.threshold                                        5.0000
## Fixed.cumulative.value.5.Cloglog.threshold                                           0.1442
## Fixed.cumulative.value.5.area                                                        0.0780
## Fixed.cumulative.value.5.training.omission                                           0.0000
## Fixed.cumulative.value.10.cumulative.threshold                                      10.0000
## Fixed.cumulative.value.10.Cloglog.threshold                                          0.3023
## Fixed.cumulative.value.10.area                                                       0.0611
## Fixed.cumulative.value.10.training.omission                                          0.0200
## Minimum.training.presence.cumulative.threshold                                       7.2248
## Minimum.training.presence.Cloglog.threshold                                          0.2228
## Minimum.training.presence.area                                                       0.0674
## Minimum.training.presence.training.omission                                          0.0000
## X10.percentile.training.presence.cumulative.threshold                               17.0623
## X10.percentile.training.presence.Cloglog.threshold                                   0.5634
## X10.percentile.training.presence.area                                                0.0506
## X10.percentile.training.presence.training.omission                                   0.1000
## Equal.training.sensitivity.and.specificity.cumulative.threshold                     14.1319
## Equal.training.sensitivity.and.specificity.Cloglog.threshold                         0.4831
## Equal.training.sensitivity.and.specificity.area                                      0.0548
## Equal.training.sensitivity.and.specificity.training.omission                         0.0600
## Maximum.training.sensitivity.plus.specificity.cumulative.threshold                   7.2248
## Maximum.training.sensitivity.plus.specificity.Cloglog.threshold                      0.2228
## Maximum.training.sensitivity.plus.specificity.area                                   0.0674
## Maximum.training.sensitivity.plus.specificity.training.omission                      0.0000
## Balance.training.omission..predicted.area.and.threshold.value.cumulative.threshold   2.2135
## Balance.training.omission..predicted.area.and.threshold.value.Cloglog.threshold      0.0310
## Balance.training.omission..predicted.area.and.threshold.value.area                   0.1096
## Balance.training.omission..predicted.area.and.threshold.value.training.omission      0.0000
## Equate.entropy.of.thresholded.and.original.distributions.cumulative.threshold        4.3213
## Equate.entropy.of.thresholded.and.original.distributions.Cloglog.threshold           0.1113
## Equate.entropy.of.thresholded.and.original.distributions.area                        0.0822
## Equate.entropy.of.thresholded.and.original.distributions.training.omission           0.0000
```

```
#str(modelo_maxent@results)
```

##### Get variable importance values

```
#plot(modelo_maxent)

contribuciones <- maxent.select.contribution(modelo_maxent,names_after_drop)

#contribuciones
par(mfrow=c(1, 2))
dotchart(as.numeric(contribuciones[,2]),
        col='blue', pch=16,
        labels=contribuciones[,1],
        main='predictor contribution')
dotchart(as.numeric(contribuciones[,3]),
        col='red', pch=16,
        labels=contribuciones[,1],
        main='predictor importance in permutations')
```

##### Draw respnse cuves

```
response(modelo_maxent)
```

##### Output MaxEnt HTML

```
#modelo_maxent
```

### Initial model evaluation

##### Evaluate model with training dataset

```
evalutaion_training <- evaluate.mdl(puntos_entrenamiento_coordenadas,puntos_aleatorios_fondo,capas_raster_drop,modelo_maxent)
```

```
## class          : ModelEvaluation 
## n presences    : 50 
## n absences     : 949 
## AUC            : 0.9710432 
## cor            : 0.6484945 
## max TPR+TNR at : 0.2226662
```

```
##                kappa spec_sens no_omission prevalence equal_sens_spec
## thresholds 0.4686839 0.2226662   0.2226662 0.04899953       0.4829901
##            sensitivity
## thresholds   0.5632848
```

```
str(evalutaion_training)
```

```
## List of 4
##  $ e    :Formal class 'ModelEvaluation' [package "dismo"] with 22 slots
##   .. ..@ presence  : num [1:50] 0.763 0.834 0.645 0.601 0.804 ...
##   .. ..@ absence   : num [1:949] 3.86e-04 7.00e-07 9.42e-02 4.56e-05 4.05e-06 ...
##   .. ..@ np        : int 50
##   .. ..@ na        : int 949
##   .. ..@ auc       : num 0.971
##   .. ..@ pauc      : num(0) 
##   .. ..@ cor       : Named num 0.648
##   .. .. ..- attr(*, "names")= chr "cor"
##   .. ..@ pcor      : num 2.85e-120
##   .. ..@ t         : num [1:858] -1e-04 -1e-04 -1e-04 -1e-04 -1e-04 ...
##   .. ..@ confusion : int [1:858, 1:4] 50 50 50 50 50 50 50 50 50 50 ...
##   .. .. ..- attr(*, "dimnames")=List of 2
##   .. .. .. ..$ : NULL
##   .. .. .. ..$ : chr [1:4] "tp" "fp" "fn" "tn"
##   .. ..@ prevalence: num [1:858] 0.0501 0.0501 0.0501 0.0501 0.0501 ...
##   .. ..@ ODP       : num [1:858] 0.95 0.95 0.95 0.95 0.95 ...
##   .. ..@ CCR       : num [1:858] 0.0501 0.0501 0.0501 0.0501 0.0501 ...
##   .. ..@ TPR       : num [1:858] 1 1 1 1 1 1 1 1 1 1 ...
##   .. ..@ TNR       : num [1:858] 0 0 0 0 0 0 0 0 0 0 ...
##   .. ..@ FPR       : num [1:858] 1 1 1 1 1 1 1 1 1 1 ...
##   .. ..@ FNR       : num [1:858] 0 0 0 0 0 0 0 0 0 0 ...
##   .. ..@ PPP       : num [1:858] 0.0501 0.0501 0.0501 0.0501 0.0501 ...
##   .. ..@ NPP       : num [1:858] NaN NaN NaN NaN NaN NaN NaN NaN NaN NaN ...
##   .. ..@ MCR       : num [1:858] 0.95 0.95 0.95 0.95 0.95 ...
##   .. ..@ OR        : num [1:858] NaN NaN NaN NaN NaN NaN NaN NaN NaN NaN ...
##   .. ..@ kappa     : num [1:858] 0 0 0 0 0 0 0 0 0 0 ...
##  $ t    :'data.frame':   1 obs. of  6 variables:
##   ..$ kappa          : num 0.469
##   ..$ spec_sens      : num 0.223
##   ..$ no_omission    : num 0.223
##   ..$ prevalence     : num 0.049
##   ..$ equal_sens_spec: num 0.483
##   ..$ sensitivity    : num 0.563
##  $ v_bkg: num [1:900, 1:20] 261 40 1398 255 276 ...
##   ..- attr(*, "dimnames")=List of 2
##   .. ..$ : NULL
##   .. ..$ : chr [1:20] "alt" "bio1" "bio2" "bio3" ...
##  $ v_pnt: num [1:50, 1:20] 1505 2275 2295 2457 2044 ...
##   ..- attr(*, "dimnames")=List of 2
##   .. ..$ : NULL
##   .. ..$ : chr [1:20] "alt" "bio1" "bio2" "bio3" ...
```

##### Evaluate model with control dataset

```
evalutaion_test <- evaluate.mdl(puntos_control_coordenadas,puntos_aleatorios_fondo,capas_raster_drop,modelo_maxent)
```

```
## class          : ModelEvaluation 
## n presences    : 50 
## n absences     : 949 
## AUC            : 0.9710643 
## cor            : 0.6443349 
## max TPR+TNR at : 0.2722717
```

```
##                kappa spec_sens no_omission prevalence equal_sens_spec
## thresholds 0.3641435 0.2722717   0.2722717 0.04899953       0.4362655
##            sensitivity
## thresholds   0.5026834
```

```
str(evalutaion_test)
```

```
## List of 4
##  $ e    :Formal class 'ModelEvaluation' [package "dismo"] with 22 slots
##   .. ..@ presence  : num [1:50] 0.628 0.766 0.93 0.503 0.783 ...
##   .. ..@ absence   : num [1:949] 3.86e-04 7.00e-07 9.42e-02 4.56e-05 4.05e-06 ...
##   .. ..@ np        : int 50
##   .. ..@ na        : int 949
##   .. ..@ auc       : num 0.971
##   .. ..@ pauc      : num(0) 
##   .. ..@ cor       : Named num 0.644
##   .. .. ..- attr(*, "names")= chr "cor"
##   .. ..@ pcor      : num 2.87e-118
##   .. ..@ t         : num [1:858] -1e-04 -1e-04 -1e-04 -1e-04 -1e-04 ...
##   .. ..@ confusion : int [1:858, 1:4] 50 50 50 50 50 50 50 50 50 50 ...
##   .. .. ..- attr(*, "dimnames")=List of 2
##   .. .. .. ..$ : NULL
##   .. .. .. ..$ : chr [1:4] "tp" "fp" "fn" "tn"
##   .. ..@ prevalence: num [1:858] 0.0501 0.0501 0.0501 0.0501 0.0501 ...
##   .. ..@ ODP       : num [1:858] 0.95 0.95 0.95 0.95 0.95 ...
##   .. ..@ CCR       : num [1:858] 0.0501 0.0501 0.0501 0.0501 0.0501 ...
##   .. ..@ TPR       : num [1:858] 1 1 1 1 1 1 1 1 1 1 ...
##   .. ..@ TNR       : num [1:858] 0 0 0 0 0 0 0 0 0 0 ...
##   .. ..@ FPR       : num [1:858] 1 1 1 1 1 1 1 1 1 1 ...
##   .. ..@ FNR       : num [1:858] 0 0 0 0 0 0 0 0 0 0 ...
##   .. ..@ PPP       : num [1:858] 0.0501 0.0501 0.0501 0.0501 0.0501 ...
##   .. ..@ NPP       : num [1:858] NaN NaN NaN NaN NaN NaN NaN NaN NaN NaN ...
##   .. ..@ MCR       : num [1:858] 0.95 0.95 0.95 0.95 0.95 ...
##   .. ..@ OR        : num [1:858] NaN NaN NaN NaN NaN NaN NaN NaN NaN NaN ...
##   .. ..@ kappa     : num [1:858] 0 0 0 0 0 0 0 0 0 0 ...
##  $ t    :'data.frame':   1 obs. of  6 variables:
##   ..$ kappa          : num 0.364
##   ..$ spec_sens      : num 0.272
##   ..$ no_omission    : num 0.272
##   ..$ prevalence     : num 0.049
##   ..$ equal_sens_spec: num 0.436
##   ..$ sensitivity    : num 0.503
##  $ v_bkg: num [1:900, 1:20] 261 40 1398 255 276 ...
##   ..- attr(*, "dimnames")=List of 2
##   .. ..$ : NULL
##   .. ..$ : chr [1:20] "alt" "bio1" "bio2" "bio3" ...
##  $ v_pnt: num [1:50, 1:20] 1285 1705 1993 2284 1927 ...
##   ..- attr(*, "dimnames")=List of 2
##   .. ..$ : NULL
##   .. ..$ : chr [1:20] "alt" "bio1" "bio2" "bio3" ...
```

##### Evaluate model with fixed control dataset

```
evalutaion_test2 <- evaluate.mdl(puntos_control_extra_coordenadas,puntos_aleatorios_fondo,capas_raster_drop,modelo_maxent)
```

```
## class          : ModelEvaluation 
## n presences    : 72 
## n absences     : 971 
## AUC            : 0.9589341 
## cor            : 0.6333403 
## max TPR+TNR at : 0.3843467
```

```
##                kappa spec_sens no_omission prevalence equal_sens_spec
## thresholds 0.3843467 0.3843467   0.3843467 0.06393831       0.4166263
##            sensitivity
## thresholds   0.4275223
```

```
str(evalutaion_test2)
```

```
## List of 4
##  $ e    :Formal class 'ModelEvaluation' [package "dismo"] with 22 slots
##   .. ..@ presence  : num [1:72] 0.502 0.459 0.598 0.803 0.598 ...
##   .. ..@ absence   : num [1:971] 3.86e-04 7.00e-07 9.42e-02 4.56e-05 4.05e-06 ...
##   .. ..@ np        : int 72
##   .. ..@ na        : int 971
##   .. ..@ auc       : num 0.959
##   .. ..@ pauc      : num(0) 
##   .. ..@ cor       : Named num 0.633
##   .. .. ..- attr(*, "names")= chr "cor"
##   .. ..@ pcor      : num 4.97e-118
##   .. ..@ t         : num [1:866] -1e-04 -1e-04 -1e-04 -1e-04 -1e-04 ...
##   .. ..@ confusion : int [1:866, 1:4] 72 72 72 72 72 72 72 72 72 72 ...
##   .. .. ..- attr(*, "dimnames")=List of 2
##   .. .. .. ..$ : NULL
##   .. .. .. ..$ : chr [1:4] "tp" "fp" "fn" "tn"
##   .. ..@ prevalence: num [1:866] 0.069 0.069 0.069 0.069 0.069 ...
##   .. ..@ ODP       : num [1:866] 0.931 0.931 0.931 0.931 0.931 ...
##   .. ..@ CCR       : num [1:866] 0.069 0.069 0.069 0.069 0.069 ...
##   .. ..@ TPR       : num [1:866] 1 1 1 1 1 1 1 1 1 1 ...
##   .. ..@ TNR       : num [1:866] 0 0 0 0 0 0 0 0 0 0 ...
##   .. ..@ FPR       : num [1:866] 1 1 1 1 1 1 1 1 1 1 ...
##   .. ..@ FNR       : num [1:866] 0 0 0 0 0 0 0 0 0 0 ...
##   .. ..@ PPP       : num [1:866] 0.069 0.069 0.069 0.069 0.069 ...
##   .. ..@ NPP       : num [1:866] NaN NaN NaN NaN NaN NaN NaN NaN NaN NaN ...
##   .. ..@ MCR       : num [1:866] 0.931 0.931 0.931 0.931 0.931 ...
##   .. ..@ OR        : num [1:866] NaN NaN NaN NaN NaN NaN NaN NaN NaN NaN ...
##   .. ..@ kappa     : num [1:866] 0 0 0 0 0 0 0 0 0 0 ...
##  $ t    :'data.frame':   1 obs. of  6 variables:
##   ..$ kappa          : num 0.384
##   ..$ spec_sens      : num 0.384
##   ..$ no_omission    : num 0.384
##   ..$ prevalence     : num 0.0639
##   ..$ equal_sens_spec: num 0.417
##   ..$ sensitivity    : num 0.428
##  $ v_bkg: num [1:900, 1:20] 261 40 1398 255 276 ...
##   ..- attr(*, "dimnames")=List of 2
##   .. ..$ : NULL
##   .. ..$ : chr [1:20] "alt" "bio1" "bio2" "bio3" ...
##  $ v_pnt: num [1:72, 1:20] 1252 1946 1793 1871 1793 ...
##   ..- attr(*, "dimnames")=List of 2
##   .. ..$ : NULL
##   .. ..$ : chr [1:20] "alt" "bio1" "bio2" "bio3" ...
```

##### Rank model AUC comparing to 99 null-models (with training dataset)

```
valores_puntos_aleatorios_fondo_y_entrenamiento <- rbind(evalutaion_training$v_pnt, evalutaion_training$v_bkg)

rep <- 99     # number or null models to produce

null_models <- list()
for (r in 1:rep) {
    # following Robert J. Hijmans nullRandom() function from 'dismo'
        # sample presence records and set the rest of records as absence
        index <- sample(nrow(valores_puntos_aleatorios_fondo_y_entrenamiento), presence_points_count)
        pres <- valores_puntos_aleatorios_fondo_y_entrenamiento[index, ]
        absc <- valores_puntos_aleatorios_fondo_y_entrenamiento[-index, ]
        d <- data.frame(rbind(pres, absc))
        v <- c(rep(1, nrow(pres)), rep(0, nrow(absc)))
        # make null-model with presence and absence sets
        m <- maxent(d, v)
        # evaluate the null-model
        null_models[[r]] <- evaluate(pres, absc, m)
}

#null_models
auc_list_vector <- sapply(null_models, function(x) x@auc)

#auc_list_vector
mean(auc_list_vector)
```

```
## [1] 0.7203298
```

```
auc_list_training_vector <- c(evalutaion_training$e@auc, auc_list_vector)
auc_list_training_vector_ranks <- rank(auc_list_training_vector, ties.method= "max")

print('Rank of trainign AUC vs null-models')
```

```
## [1] "Rank of trainign AUC vs null-models"
```

```
auc_list_training_vector_ranks[1]
```

```
## [1] 100
```

```
auc_list_control_vector <- c(evalutaion_test$e@auc, auc_list_vector)
auc_list_control_vector_ranks <- rank(auc_list_control_vector, ties.method= "max")

print('Rank of control AUC vs null-models')
```

```
## [1] "Rank of control AUC vs null-models"
```

```
auc_list_control_vector_ranks[1]
```

```
## [1] 100
```

##### Store the model, parameters and control values in an object for further use

```
container <- list()
container$bkg <- puntos_aleatorios_fondo
container$entr <- puntos_entrenamiento_coordenadas
container$cntrl <- puntos_control_extra_coordenadas
container$model <- modelo_maxent
container$contrib <- contribuciones
container$e_train <- evalutaion_training
container$e_test <- evalutaion_test
container$e_test2 <- evalutaion_test2
container$auc_rank_train <- auc_list_training_vector_ranks[1]
container$auc_rank_test <- auc_list_control_vector_ranks[1]
container$auc_null_model_mean <- mean(auc_list_vector)
  
iterations[[i]] <- container
```
