## Supplementary material for "Vulnerability to climate change for narrowly ranged species: the case of Ecuadorian endemic *Magnolia mercedesiarum*": step_29dup_4.html

```
##                                                                                        [,1]
## X.Training.samples                                                                  50.0000
## Regularized.training.gain                                                            2.5308
## Unregularized.training.gain                                                          2.6992
## Iterations                                                                         500.0000
## Training.AUC                                                                         0.9730
## X.Background.points                                                                947.0000
## alt.contribution                                                                     7.0481
## bio1.contribution                                                                    0.0000
## bio10.contribution                                                                   1.8354
## bio11.contribution                                                                   0.0000
## bio12.contribution                                                                   0.7091
## bio13.contribution                                                                   0.0000
## bio14.contribution                                                                   0.0122
## bio15.contribution                                                                   3.0152
## bio16.contribution                                                                   0.9729
## bio17.contribution                                                                   0.0030
## bio18.contribution                                                                  10.8123
## bio19.contribution                                                                  28.4491
## bio2.contribution                                                                    0.0000
## bio3.contribution                                                                   22.1322
## bio4.contribution                                                                    0.0483
## bio5.contribution                                                                    0.1342
## bio6.contribution                                                                   24.6853
## bio7.contribution                                                                    0.1417
## bio8.contribution                                                                    0.0000
## bio9.contribution                                                                    0.0010
## alt.permutation.importance                                                          13.3459
## bio1.permutation.importance                                                          0.0000
## bio10.permutation.importance                                                         1.3413
## bio11.permutation.importance                                                         0.0000
## bio12.permutation.importance                                                        17.7328
## bio13.permutation.importance                                                         0.0000
## bio14.permutation.importance                                                         0.0726
## bio15.permutation.importance                                                         1.3719
## bio16.permutation.importance                                                         0.0000
## bio17.permutation.importance                                                         0.0000
## bio18.permutation.importance                                                        40.2385
## bio19.permutation.importance                                                         0.0000
## bio2.permutation.importance                                                          0.0000
## bio3.permutation.importance                                                          0.9668
## bio4.permutation.importance                                                          1.5935
## bio5.permutation.importance                                                         19.7524
## bio6.permutation.importance                                                          3.5729
## bio7.permutation.importance                                                          0.0000
## bio8.permutation.importance                                                          0.0000
## bio9.permutation.importance                                                          0.0115
## Training.gain.without.alt                                                            2.4894
## Training.gain.without.bio1                                                           2.5308
## Training.gain.without.bio10                                                          2.5307
## Training.gain.without.bio11                                                          2.5310
## Training.gain.without.bio12                                                          2.5146
## Training.gain.without.bio13                                                          2.5310
## Training.gain.without.bio14                                                          2.5306
## Training.gain.without.bio15                                                          2.5242
## Training.gain.without.bio16                                                          2.5311
## Training.gain.without.bio17                                                          2.5306
## Training.gain.without.bio18                                                          2.4290
## Training.gain.without.bio19                                                          2.5310
## Training.gain.without.bio2                                                           2.5309
## Training.gain.without.bio3                                                           2.5182
## Training.gain.without.bio4                                                           2.5293
## Training.gain.without.bio5                                                           2.5303
## Training.gain.without.bio6                                                           2.5308
## Training.gain.without.bio7                                                           2.5310
## Training.gain.without.bio8                                                           2.5311
## Training.gain.without.bio9                                                           2.5312
## Training.gain.with.only.alt                                                          1.5956
## Training.gain.with.only.bio1                                                         1.4214
## Training.gain.with.only.bio10                                                        1.4608
## Training.gain.with.only.bio11                                                        1.3618
## Training.gain.with.only.bio12                                                        0.2860
## Training.gain.with.only.bio13                                                        0.1295
## Training.gain.with.only.bio14                                                        1.0796
## Training.gain.with.only.bio15                                                        0.3712
## Training.gain.with.only.bio16                                                        0.1451
## Training.gain.with.only.bio17                                                        0.9920
## Training.gain.with.only.bio18                                                        0.5628
## Training.gain.with.only.bio19                                                        0.3684
## Training.gain.with.only.bio2                                                         0.0649
## Training.gain.with.only.bio3                                                         0.6647
## Training.gain.with.only.bio4                                                         0.8672
## Training.gain.with.only.bio5                                                         1.4083
## Training.gain.with.only.bio6                                                         1.3556
## Training.gain.with.only.bio7                                                         0.2764
## Training.gain.with.only.bio8                                                         1.4426
## Training.gain.with.only.bio9                                                         1.3870
## Entropy                                                                              4.3223
## Prevalence..average.probability.of.presence.over.background.sites.                   0.0467
## Fixed.cumulative.value.1.cumulative.threshold                                        1.0000
## Fixed.cumulative.value.1.Cloglog.threshold                                           0.0110
## Fixed.cumulative.value.1.area                                                        0.1584
## Fixed.cumulative.value.1.training.omission                                           0.0000
## Fixed.cumulative.value.5.cumulative.threshold                                        5.0000
## Fixed.cumulative.value.5.Cloglog.threshold                                           0.1089
## Fixed.cumulative.value.5.area                                                        0.0781
## Fixed.cumulative.value.5.training.omission                                           0.0000
## Fixed.cumulative.value.10.cumulative.threshold                                      10.0000
## Fixed.cumulative.value.10.Cloglog.threshold                                          0.3533
## Fixed.cumulative.value.10.area                                                       0.0591
## Fixed.cumulative.value.10.training.omission                                          0.0200
## Minimum.training.presence.cumulative.threshold                                       9.1404
## Minimum.training.presence.Cloglog.threshold                                          0.2235
## Minimum.training.presence.area                                                       0.0602
## Minimum.training.presence.training.omission                                          0.0000
## X10.percentile.training.presence.cumulative.threshold                               17.9668
## X10.percentile.training.presence.Cloglog.threshold                                   0.5101
## X10.percentile.training.presence.area                                                0.0475
## X10.percentile.training.presence.training.omission                                   0.1000
## Equal.training.sensitivity.and.specificity.cumulative.threshold                     12.8660
## Equal.training.sensitivity.and.specificity.Cloglog.threshold                         0.4224
## Equal.training.sensitivity.and.specificity.area                                      0.0549
## Equal.training.sensitivity.and.specificity.training.omission                         0.0600
## Maximum.training.sensitivity.plus.specificity.cumulative.threshold                   9.1404
## Maximum.training.sensitivity.plus.specificity.Cloglog.threshold                      0.2235
## Maximum.training.sensitivity.plus.specificity.area                                   0.0602
## Maximum.training.sensitivity.plus.specificity.training.omission                      0.0000
## Balance.training.omission..predicted.area.and.threshold.value.cumulative.threshold   1.9973
## Balance.training.omission..predicted.area.and.threshold.value.Cloglog.threshold      0.0309
## Balance.training.omission..predicted.area.and.threshold.value.area                   0.1172
## Balance.training.omission..predicted.area.and.threshold.value.training.omission      0.0000
## Equate.entropy.of.thresholded.and.original.distributions.cumulative.threshold        4.9696
## Equate.entropy.of.thresholded.and.original.distributions.Cloglog.threshold           0.1066
## Equate.entropy.of.thresholded.and.original.distributions.area                        0.0792
## Equate.entropy.of.thresholded.and.original.distributions.training.omission           0.0000
```

```
#str(modelo_maxent@results)
```

##### Get variable importance values

```
#plot(modelo_maxent)

contribuciones <- maxent.select.contribution(modelo_maxent,names_after_drop)

#contribuciones
par(mfrow=c(1, 2))
dotchart(as.numeric(contribuciones[,2]),
        col='blue', pch=16,
        labels=contribuciones[,1],
        main='predictor contribution')
dotchart(as.numeric(contribuciones[,3]),
        col='red', pch=16,
        labels=contribuciones[,1],
        main='predictor importance in permutations')
```

##### Draw respnse cuves

```
response(modelo_maxent)
```

##### Output MaxEnt HTML

```
#modelo_maxent
```

### Initial model evaluation

##### Evaluate model with training dataset

```
evalutaion_training <- evaluate.mdl(puntos_entrenamiento_coordenadas,puntos_aleatorios_fondo,capas_raster_drop,modelo_maxent)
```

```
## class          : ModelEvaluation 
## n presences    : 50 
## n absences     : 947 
## AUC            : 0.9730095 
## cor            : 0.6566945 
## max TPR+TNR at : 0.223376
```

```
##                kappa spec_sens no_omission prevalence equal_sens_spec
## thresholds 0.4881428  0.223376    0.223376 0.04947405        0.422268
##            sensitivity
## thresholds   0.4883774
```

```
str(evalutaion_training)
```

```
## List of 4
##  $ e    :Formal class 'ModelEvaluation' [package "dismo"] with 22 slots
##   .. ..@ presence  : num [1:50] 0.878 0.702 0.679 0.659 0.762 ...
##   .. ..@ absence   : num [1:947] 4.62e-05 2.16e-07 1.07e-08 1.49e-01 1.73e-04 ...
##   .. ..@ np        : int 50
##   .. ..@ na        : int 947
##   .. ..@ auc       : num 0.973
##   .. ..@ pauc      : num(0) 
##   .. ..@ cor       : Named num 0.657
##   .. .. ..- attr(*, "names")= chr "cor"
##   .. ..@ pcor      : num 4.56e-124
##   .. ..@ t         : num [1:848] -1e-04 -1e-04 -1e-04 -1e-04 -1e-04 ...
##   .. ..@ confusion : int [1:848, 1:4] 50 50 50 50 50 50 50 50 50 50 ...
##   .. .. ..- attr(*, "dimnames")=List of 2
##   .. .. .. ..$ : NULL
##   .. .. .. ..$ : chr [1:4] "tp" "fp" "fn" "tn"
##   .. ..@ prevalence: num [1:848] 0.0502 0.0502 0.0502 0.0502 0.0502 ...
##   .. ..@ ODP       : num [1:848] 0.95 0.95 0.95 0.95 0.95 ...
##   .. ..@ CCR       : num [1:848] 0.0502 0.0502 0.0502 0.0502 0.0502 ...
##   .. ..@ TPR       : num [1:848] 1 1 1 1 1 1 1 1 1 1 ...
##   .. ..@ TNR       : num [1:848] 0 0 0 0 0 0 0 0 0 0 ...
##   .. ..@ FPR       : num [1:848] 1 1 1 1 1 1 1 1 1 1 ...
##   .. ..@ FNR       : num [1:848] 0 0 0 0 0 0 0 0 0 0 ...
##   .. ..@ PPP       : num [1:848] 0.0502 0.0502 0.0502 0.0502 0.0502 ...
##   .. ..@ NPP       : num [1:848] NaN NaN NaN NaN NaN NaN NaN NaN NaN NaN ...
##   .. ..@ MCR       : num [1:848] 0.95 0.95 0.95 0.95 0.95 ...
##   .. ..@ OR        : num [1:848] NaN NaN NaN NaN NaN NaN NaN NaN NaN NaN ...
##   .. ..@ kappa     : num [1:848] 0 0 0 0 0 0 0 0 0 0 ...
##  $ t    :'data.frame':   1 obs. of  6 variables:
##   ..$ kappa          : num 0.488
##   ..$ spec_sens      : num 0.223
##   ..$ no_omission    : num 0.223
##   ..$ prevalence     : num 0.0495
##   ..$ equal_sens_spec: num 0.422
##   ..$ sensitivity    : num 0.488
##  $ v_bkg: num [1:900, 1:20] 180 57 5 2608 216 ...
##   ..- attr(*, "dimnames")=List of 2
##   .. ..$ : NULL
##   .. ..$ : chr [1:20] "alt" "bio1" "bio2" "bio3" ...
##  $ v_pnt: num [1:50, 1:20] 1546 1784 1740 2049 1785 ...
##   ..- attr(*, "dimnames")=List of 2
##   .. ..$ : NULL
##   .. ..$ : chr [1:20] "alt" "bio1" "bio2" "bio3" ...
```

##### Evaluate model with control dataset

```
evalutaion_test <- evaluate.mdl(puntos_control_coordenadas,puntos_aleatorios_fondo,capas_raster_drop,modelo_maxent)
```

```
## class          : ModelEvaluation 
## n presences    : 50 
## n absences     : 947 
## AUC            : 0.9709187 
## cor            : 0.6327075 
## max TPR+TNR at : 0.1268105
```

```
##                kappa spec_sens no_omission prevalence equal_sens_spec
## thresholds 0.2254459 0.1268105  0.03817012 0.04947405       0.2368941
##            sensitivity
## thresholds   0.2602624
```

```
str(evalutaion_test)
```

```
## List of 4
##  $ e    :Formal class 'ModelEvaluation' [package "dismo"] with 22 slots
##   .. ..@ presence  : num [1:50] 0.237 0.865 0.762 0.671 0.864 ...
##   .. ..@ absence   : num [1:947] 4.62e-05 2.16e-07 1.07e-08 1.49e-01 1.73e-04 ...
##   .. ..@ np        : int 50
##   .. ..@ na        : int 947
##   .. ..@ auc       : num 0.971
##   .. ..@ pauc      : num(0) 
##   .. ..@ cor       : Named num 0.633
##   .. .. ..- attr(*, "names")= chr "cor"
##   .. ..@ pcor      : num 1.31e-112
##   .. ..@ t         : num [1:848] -1e-04 -1e-04 -1e-04 -1e-04 -1e-04 ...
##   .. ..@ confusion : int [1:848, 1:4] 50 50 50 50 50 50 50 50 50 50 ...
##   .. .. ..- attr(*, "dimnames")=List of 2
##   .. .. .. ..$ : NULL
##   .. .. .. ..$ : chr [1:4] "tp" "fp" "fn" "tn"
##   .. ..@ prevalence: num [1:848] 0.0502 0.0502 0.0502 0.0502 0.0502 ...
##   .. ..@ ODP       : num [1:848] 0.95 0.95 0.95 0.95 0.95 ...
##   .. ..@ CCR       : num [1:848] 0.0502 0.0502 0.0502 0.0502 0.0502 ...
##   .. ..@ TPR       : num [1:848] 1 1 1 1 1 1 1 1 1 1 ...
##   .. ..@ TNR       : num [1:848] 0 0 0 0 0 0 0 0 0 0 ...
##   .. ..@ FPR       : num [1:848] 1 1 1 1 1 1 1 1 1 1 ...
##   .. ..@ FNR       : num [1:848] 0 0 0 0 0 0 0 0 0 0 ...
##   .. ..@ PPP       : num [1:848] 0.0502 0.0502 0.0502 0.0502 0.0502 ...
##   .. ..@ NPP       : num [1:848] NaN NaN NaN NaN NaN NaN NaN NaN NaN NaN ...
##   .. ..@ MCR       : num [1:848] 0.95 0.95 0.95 0.95 0.95 ...
##   .. ..@ OR        : num [1:848] NaN NaN NaN NaN NaN NaN NaN NaN NaN NaN ...
##   .. ..@ kappa     : num [1:848] 0 0 0 0 0 0 0 0 0 0 ...
##  $ t    :'data.frame':   1 obs. of  6 variables:
##   ..$ kappa          : num 0.225
##   ..$ spec_sens      : num 0.127
##   ..$ no_omission    : num 0.0382
##   ..$ prevalence     : num 0.0495
##   ..$ equal_sens_spec: num 0.237
##   ..$ sensitivity    : num 0.26
##  $ v_bkg: num [1:900, 1:20] 180 57 5 2608 216 ...
##   ..- attr(*, "dimnames")=List of 2
##   .. ..$ : NULL
##   .. ..$ : chr [1:20] "alt" "bio1" "bio2" "bio3" ...
##  $ v_pnt: num [1:50, 1:20] 1349 1556 1731 1559 1788 ...
##   ..- attr(*, "dimnames")=List of 2
##   .. ..$ : NULL
##   .. ..$ : chr [1:20] "alt" "bio1" "bio2" "bio3" ...
```

##### Evaluate model with fixed control dataset

```
evalutaion_test2 <- evaluate.mdl(puntos_control_extra_coordenadas,puntos_aleatorios_fondo,capas_raster_drop,modelo_maxent)
```

```
## class          : ModelEvaluation 
## n presences    : 72 
## n absences     : 969 
## AUC            : 0.9613433 
## cor            : 0.6405048 
## max TPR+TNR at : 0.2531108
```

```
##                kappa spec_sens no_omission prevalence equal_sens_spec
## thresholds 0.2531108 0.2531108   0.2531108 0.06871727       0.4122998
##            sensitivity
## thresholds   0.4397148
```

```
str(evalutaion_test2)
```

```
## List of 4
##  $ e    :Formal class 'ModelEvaluation' [package "dismo"] with 22 slots
##   .. ..@ presence  : num [1:72] 0.467 0.47 0.734 0.792 0.734 ...
##   .. ..@ absence   : num [1:969] 4.62e-05 2.16e-07 1.07e-08 1.49e-01 1.73e-04 ...
##   .. ..@ np        : int 72
##   .. ..@ na        : int 969
##   .. ..@ auc       : num 0.961
##   .. ..@ pauc      : num(0) 
##   .. ..@ cor       : Named num 0.641
##   .. .. ..- attr(*, "names")= chr "cor"
##   .. ..@ pcor      : num 2.82e-121
##   .. ..@ t         : num [1:856] -1e-04 -1e-04 -1e-04 -1e-04 -1e-04 ...
##   .. ..@ confusion : int [1:856, 1:4] 72 72 72 72 72 72 72 72 72 72 ...
##   .. .. ..- attr(*, "dimnames")=List of 2
##   .. .. .. ..$ : NULL
##   .. .. .. ..$ : chr [1:4] "tp" "fp" "fn" "tn"
##   .. ..@ prevalence: num [1:856] 0.0692 0.0692 0.0692 0.0692 0.0692 ...
##   .. ..@ ODP       : num [1:856] 0.931 0.931 0.931 0.931 0.931 ...
##   .. ..@ CCR       : num [1:856] 0.0692 0.0692 0.0692 0.0692 0.0692 ...
##   .. ..@ TPR       : num [1:856] 1 1 1 1 1 1 1 1 1 1 ...
##   .. ..@ TNR       : num [1:856] 0 0 0 0 0 0 0 0 0 0 ...
##   .. ..@ FPR       : num [1:856] 1 1 1 1 1 1 1 1 1 1 ...
##   .. ..@ FNR       : num [1:856] 0 0 0 0 0 0 0 0 0 0 ...
##   .. ..@ PPP       : num [1:856] 0.0692 0.0692 0.0692 0.0692 0.0692 ...
##   .. ..@ NPP       : num [1:856] NaN NaN NaN NaN NaN NaN NaN NaN NaN NaN ...
##   .. ..@ MCR       : num [1:856] 0.931 0.931 0.931 0.931 0.931 ...
##   .. ..@ OR        : num [1:856] NaN NaN NaN NaN NaN NaN NaN NaN NaN NaN ...
##   .. ..@ kappa     : num [1:856] 0 0 0 0 0 0 0 0 0 0 ...
##  $ t    :'data.frame':   1 obs. of  6 variables:
##   ..$ kappa          : num 0.253
##   ..$ spec_sens      : num 0.253
##   ..$ no_omission    : num 0.253
##   ..$ prevalence     : num 0.0687
##   ..$ equal_sens_spec: num 0.412
##   ..$ sensitivity    : num 0.44
##  $ v_bkg: num [1:900, 1:20] 180 57 5 2608 216 ...
##   ..- attr(*, "dimnames")=List of 2
##   .. ..$ : NULL
##   .. ..$ : chr [1:20] "alt" "bio1" "bio2" "bio3" ...
##  $ v_pnt: num [1:72, 1:20] 1252 1946 1793 1871 1793 ...
##   ..- attr(*, "dimnames")=List of 2
##   .. ..$ : NULL
##   .. ..$ : chr [1:20] "alt" "bio1" "bio2" "bio3" ...
```

##### Rank model AUC comparing to 99 null-models (with training dataset)

```
valores_puntos_aleatorios_fondo_y_entrenamiento <- rbind(evalutaion_training$v_pnt, evalutaion_training$v_bkg)

rep <- 99     # number or null models to produce

null_models <- list()
for (r in 1:rep) {
    # following Robert J. Hijmans nullRandom() function from 'dismo'
        # sample presence records and set the rest of records as absence
        index <- sample(nrow(valores_puntos_aleatorios_fondo_y_entrenamiento), presence_points_count)
        pres <- valores_puntos_aleatorios_fondo_y_entrenamiento[index, ]
        absc <- valores_puntos_aleatorios_fondo_y_entrenamiento[-index, ]
        d <- data.frame(rbind(pres, absc))
        v <- c(rep(1, nrow(pres)), rep(0, nrow(absc)))
        # make null-model with presence and absence sets
        m <- maxent(d, v)
        # evaluate the null-model
        null_models[[r]] <- evaluate(pres, absc, m)
}

#null_models
auc_list_vector <- sapply(null_models, function(x) x@auc)

#auc_list_vector
mean(auc_list_vector)
```

```
## [1] 0.7263127
```

```
auc_list_training_vector <- c(evalutaion_training$e@auc, auc_list_vector)
auc_list_training_vector_ranks <- rank(auc_list_training_vector, ties.method= "max")

print('Rank of trainign AUC vs null-models')
```

```
## [1] "Rank of trainign AUC vs null-models"
```

```
auc_list_training_vector_ranks[1]
```

```
## [1] 100
```

```
auc_list_control_vector <- c(evalutaion_test$e@auc, auc_list_vector)
auc_list_control_vector_ranks <- rank(auc_list_control_vector, ties.method= "max")

print('Rank of control AUC vs null-models')
```

```
## [1] "Rank of control AUC vs null-models"
```

```
auc_list_control_vector_ranks[1]
```

```
## [1] 100
```

##### Store the model, parameters and control values in an object for further use

```
container <- list()
container$bkg <- puntos_aleatorios_fondo
container$entr <- puntos_entrenamiento_coordenadas
container$cntrl <- puntos_control_extra_coordenadas
container$model <- modelo_maxent
container$contrib <- contribuciones
container$e_train <- evalutaion_training
container$e_test <- evalutaion_test
container$e_test2 <- evalutaion_test2
container$auc_rank_train <- auc_list_training_vector_ranks[1]
container$auc_rank_test <- auc_list_control_vector_ranks[1]
container$auc_null_model_mean <- mean(auc_list_vector)
  
iterations[[i]] <- container
```
