## Supplementary material for "Vulnerability to climate change for narrowly ranged species: the case of Ecuadorian endemic *Magnolia mercedesiarum*": step_29dup_5.html

```
## Warning in .local(x, p, ...): 5 (0.56%) of the presence points have NA
## predictor values
```

```
summary(modelo_maxent)
```

```
## Length  Class   Mode 
##      1 MaxEnt     S4
```

```
##str(modelo_maxent)
#representar resultados en forma de una lista
print(modelo_maxent@results)
```

```
##                                                                                        [,1]
## X.Training.samples                                                                  50.0000
## Regularized.training.gain                                                            2.5271
## Unregularized.training.gain                                                          2.6920
## Iterations                                                                         500.0000
## Training.AUC                                                                         0.9730
## X.Background.points                                                                945.0000
## alt.contribution                                                                     8.0773
## bio1.contribution                                                                    0.0000
## bio10.contribution                                                                   0.0013
## bio11.contribution                                                                   0.0000
## bio12.contribution                                                                   0.2480
## bio13.contribution                                                                   0.0000
## bio14.contribution                                                                   3.2598
## bio15.contribution                                                                   0.2718
## bio16.contribution                                                                   0.0000
## bio17.contribution                                                                   2.0959
## bio18.contribution                                                                  15.0021
## bio19.contribution                                                                  21.3242
## bio2.contribution                                                                    0.0000
## bio3.contribution                                                                   23.6799
## bio4.contribution                                                                    0.1054
## bio5.contribution                                                                    0.0648
## bio6.contribution                                                                   22.7234
## bio7.contribution                                                                    0.0000
## bio8.contribution                                                                    1.4733
## bio9.contribution                                                                    1.6728
## alt.permutation.importance                                                          16.6203
## bio1.permutation.importance                                                          0.0000
## bio10.permutation.importance                                                         0.0586
## bio11.permutation.importance                                                         0.0000
## bio12.permutation.importance                                                        16.5848
## bio13.permutation.importance                                                         0.0000
## bio14.permutation.importance                                                         0.0000
## bio15.permutation.importance                                                         0.9620
## bio16.permutation.importance                                                         0.0000
## bio17.permutation.importance                                                         0.0711
## bio18.permutation.importance                                                        31.2120
## bio19.permutation.importance                                                         0.0000
## bio2.permutation.importance                                                          0.0000
## bio3.permutation.importance                                                          4.2957
## bio4.permutation.importance                                                          0.3388
## bio5.permutation.importance                                                          4.0029
## bio6.permutation.importance                                                         25.8245
## bio7.permutation.importance                                                          0.0000
## bio8.permutation.importance                                                          0.0000
## bio9.permutation.importance                                                          0.0293
## Training.gain.without.alt                                                            2.4789
## Training.gain.without.bio1                                                           2.5271
## Training.gain.without.bio10                                                          2.5270
## Training.gain.without.bio11                                                          2.5270
## Training.gain.without.bio12                                                          2.5215
## Training.gain.without.bio13                                                          2.5271
## Training.gain.without.bio14                                                          2.5272
## Training.gain.without.bio15                                                          2.5226
## Training.gain.without.bio16                                                          2.5271
## Training.gain.without.bio17                                                          2.5271
## Training.gain.without.bio18                                                          2.4400
## Training.gain.without.bio19                                                          2.5270
## Training.gain.without.bio2                                                           2.5272
## Training.gain.without.bio3                                                           2.5004
## Training.gain.without.bio4                                                           2.5268
## Training.gain.without.bio5                                                           2.5274
## Training.gain.without.bio6                                                           2.5265
## Training.gain.without.bio7                                                           2.5271
## Training.gain.without.bio8                                                           2.5268
## Training.gain.without.bio9                                                           2.5270
## Training.gain.with.only.alt                                                          1.5416
## Training.gain.with.only.bio1                                                         1.4093
## Training.gain.with.only.bio10                                                        1.4464
## Training.gain.with.only.bio11                                                        1.3568
## Training.gain.with.only.bio12                                                        0.2282
## Training.gain.with.only.bio13                                                        0.1018
## Training.gain.with.only.bio14                                                        1.0289
## Training.gain.with.only.bio15                                                        0.3364
## Training.gain.with.only.bio16                                                        0.1048
## Training.gain.with.only.bio17                                                        0.9062
## Training.gain.with.only.bio18                                                        0.5831
## Training.gain.with.only.bio19                                                        0.3081
## Training.gain.with.only.bio2                                                         0.0326
## Training.gain.with.only.bio3                                                         0.7200
## Training.gain.with.only.bio4                                                         0.8258
## Training.gain.with.only.bio5                                                         1.4312
## Training.gain.with.only.bio6                                                         1.3614
## Training.gain.with.only.bio7                                                         0.1731
## Training.gain.with.only.bio8                                                         1.4539
## Training.gain.with.only.bio9                                                         1.3775
## Entropy                                                                              4.3239
## Prevalence..average.probability.of.presence.over.background.sites.                   0.0466
## Fixed.cumulative.value.1.cumulative.threshold                                        1.0000
## Fixed.cumulative.value.1.Cloglog.threshold                                           0.0099
## Fixed.cumulative.value.1.area                                                        0.1725
## Fixed.cumulative.value.1.training.omission                                           0.0000
## Fixed.cumulative.value.5.cumulative.threshold                                        5.0000
## Fixed.cumulative.value.5.Cloglog.threshold                                           0.1070
## Fixed.cumulative.value.5.area                                                        0.0825
## Fixed.cumulative.value.5.training.omission                                           0.0000
## Fixed.cumulative.value.10.cumulative.threshold                                      10.0000
## Fixed.cumulative.value.10.Cloglog.threshold                                          0.2442
## Fixed.cumulative.value.10.area                                                       0.0603
## Fixed.cumulative.value.10.training.omission                                          0.0200
## Minimum.training.presence.cumulative.threshold                                       6.9108
## Minimum.training.presence.Cloglog.threshold                                          0.1570
## Minimum.training.presence.area                                                       0.0709
## Minimum.training.presence.training.omission                                          0.0000
## X10.percentile.training.presence.cumulative.threshold                               16.8775
## X10.percentile.training.presence.Cloglog.threshold                                   0.4819
## X10.percentile.training.presence.area                                                0.0487
## X10.percentile.training.presence.training.omission                                   0.1000
## Equal.training.sensitivity.and.specificity.cumulative.threshold                     14.3006
## Equal.training.sensitivity.and.specificity.Cloglog.threshold                         0.4274
## Equal.training.sensitivity.and.specificity.area                                      0.0519
## Equal.training.sensitivity.and.specificity.training.omission                         0.0600
## Maximum.training.sensitivity.plus.specificity.cumulative.threshold                   6.9108
## Maximum.training.sensitivity.plus.specificity.Cloglog.threshold                      0.1570
## Maximum.training.sensitivity.plus.specificity.area                                   0.0709
## Maximum.training.sensitivity.plus.specificity.training.omission                      0.0000
## Balance.training.omission..predicted.area.and.threshold.value.cumulative.threshold   2.2310
## Balance.training.omission..predicted.area.and.threshold.value.Cloglog.threshold      0.0314
## Balance.training.omission..predicted.area.and.threshold.value.area                   0.1196
## Balance.training.omission..predicted.area.and.threshold.value.training.omission      0.0000
## Equate.entropy.of.thresholded.and.original.distributions.cumulative.threshold        5.4896
## Equate.entropy.of.thresholded.and.original.distributions.Cloglog.threshold           0.1182
## Equate.entropy.of.thresholded.and.original.distributions.area                        0.0794
## Equate.entropy.of.thresholded.and.original.distributions.training.omission           0.0000
```

```
#str(modelo_maxent@results)
```

##### Get variable importance values

```
#plot(modelo_maxent)

contribuciones <- maxent.select.contribution(modelo_maxent,names_after_drop)

#contribuciones
par(mfrow=c(1, 2))
dotchart(as.numeric(contribuciones[,2]),
        col='blue', pch=16,
        labels=contribuciones[,1],
        main='predictor contribution')
dotchart(as.numeric(contribuciones[,3]),
        col='red', pch=16,
        labels=contribuciones[,1],
        main='predictor importance in permutations')
```

##### Draw respnse cuves

```
response(modelo_maxent)
```

##### Output MaxEnt HTML

```
#modelo_maxent
```

### Initial model evaluation

##### Evaluate model with training dataset

```
evalutaion_training <- evaluate.mdl(puntos_entrenamiento_coordenadas,puntos_aleatorios_fondo,capas_raster_drop,modelo_maxent)
```

```
## class          : ModelEvaluation 
## n presences    : 50 
## n absences     : 945 
## AUC            : 0.9729735 
## cor            : 0.657763 
## max TPR+TNR at : 0.1569168
```

```
##                kappa spec_sens no_omission prevalence equal_sens_spec
## thresholds 0.3126245 0.1569168   0.1569168 0.05002202       0.4273411
##            sensitivity
## thresholds   0.4787077
```

```
str(evalutaion_training)
```

```
## List of 4
##  $ e    :Formal class 'ModelEvaluation' [package "dismo"] with 22 slots
##   .. ..@ presence  : num [1:50] 0.85 0.733 0.659 0.77 0.874 ...
##   .. ..@ absence   : num [1:945] 2.77e-06 1.00e-03 6.38e-05 3.60e-06 1.19e-05 ...
##   .. ..@ np        : int 50
##   .. ..@ na        : int 945
##   .. ..@ auc       : num 0.973
##   .. ..@ pauc      : num(0) 
##   .. ..@ cor       : Named num 0.658
##   .. .. ..- attr(*, "names")= chr "cor"
##   .. ..@ pcor      : num 2.35e-124
##   .. ..@ t         : num [1:875] -1e-04 -1e-04 -1e-04 -1e-04 -1e-04 ...
##   .. ..@ confusion : int [1:875, 1:4] 50 50 50 50 50 50 50 50 50 50 ...
##   .. .. ..- attr(*, "dimnames")=List of 2
##   .. .. .. ..$ : NULL
##   .. .. .. ..$ : chr [1:4] "tp" "fp" "fn" "tn"
##   .. ..@ prevalence: num [1:875] 0.0503 0.0503 0.0503 0.0503 0.0503 ...
##   .. ..@ ODP       : num [1:875] 0.95 0.95 0.95 0.95 0.95 ...
##   .. ..@ CCR       : num [1:875] 0.0503 0.0503 0.0503 0.0503 0.0503 ...
##   .. ..@ TPR       : num [1:875] 1 1 1 1 1 1 1 1 1 1 ...
##   .. ..@ TNR       : num [1:875] 0 0 0 0 0 0 0 0 0 0 ...
##   .. ..@ FPR       : num [1:875] 1 1 1 1 1 1 1 1 1 1 ...
##   .. ..@ FNR       : num [1:875] 0 0 0 0 0 0 0 0 0 0 ...
##   .. ..@ PPP       : num [1:875] 0.0503 0.0503 0.0503 0.0503 0.0503 ...
##   .. ..@ NPP       : num [1:875] NaN NaN NaN NaN NaN NaN NaN NaN NaN NaN ...
##   .. ..@ MCR       : num [1:875] 0.95 0.95 0.95 0.95 0.95 ...
##   .. ..@ OR        : num [1:875] NaN NaN NaN NaN NaN NaN NaN NaN NaN NaN ...
##   .. ..@ kappa     : num [1:875] 0 0 0 0 0 0 0 0 0 0 ...
##  $ t    :'data.frame':   1 obs. of  6 variables:
##   ..$ kappa          : num 0.313
##   ..$ spec_sens      : num 0.157
##   ..$ no_omission    : num 0.157
##   ..$ prevalence     : num 0.05
##   ..$ equal_sens_spec: num 0.427
##   ..$ sensitivity    : num 0.479
##  $ v_bkg: num [1:900, 1:20] 101 261 141 152 336 ...
##   ..- attr(*, "dimnames")=List of 2
##   .. ..$ : NULL
##   .. ..$ : chr [1:20] "alt" "bio1" "bio2" "bio3" ...
##  $ v_pnt: num [1:50, 1:20] 1710 1523 2146 1507 1574 ...
##   ..- attr(*, "dimnames")=List of 2
##   .. ..$ : NULL
##   .. ..$ : chr [1:20] "alt" "bio1" "bio2" "bio3" ...
```

##### Evaluate model with control dataset

```
evalutaion_test <- evaluate.mdl(puntos_control_coordenadas,puntos_aleatorios_fondo,capas_raster_drop,modelo_maxent)
```

```
## class          : ModelEvaluation 
## n presences    : 50 
## n absences     : 945 
## AUC            : 0.9730794 
## cor            : 0.6533174 
## max TPR+TNR at : 0.235911
```

```
##                kappa spec_sens no_omission prevalence equal_sens_spec
## thresholds 0.3375569  0.235911    0.235911 0.05002202       0.4057216
##            sensitivity
## thresholds   0.4304767
```

```
str(evalutaion_test)
```

```
## List of 4
##  $ e    :Formal class 'ModelEvaluation' [package "dismo"] with 22 slots
##   .. ..@ presence  : num [1:50] 0.738 0.822 0.926 0.475 0.725 ...
##   .. ..@ absence   : num [1:945] 2.77e-06 1.00e-03 6.38e-05 3.60e-06 1.19e-05 ...
##   .. ..@ np        : int 50
##   .. ..@ na        : int 945
##   .. ..@ auc       : num 0.973
##   .. ..@ pauc      : num(0) 
##   .. ..@ cor       : Named num 0.653
##   .. .. ..- attr(*, "names")= chr "cor"
##   .. ..@ pcor      : num 3.78e-122
##   .. ..@ t         : num [1:875] -1e-04 -1e-04 -1e-04 -1e-04 -1e-04 ...
##   .. ..@ confusion : int [1:875, 1:4] 50 50 50 50 50 50 50 50 50 50 ...
##   .. .. ..- attr(*, "dimnames")=List of 2
##   .. .. .. ..$ : NULL
##   .. .. .. ..$ : chr [1:4] "tp" "fp" "fn" "tn"
##   .. ..@ prevalence: num [1:875] 0.0503 0.0503 0.0503 0.0503 0.0503 ...
##   .. ..@ ODP       : num [1:875] 0.95 0.95 0.95 0.95 0.95 ...
##   .. ..@ CCR       : num [1:875] 0.0503 0.0503 0.0503 0.0503 0.0503 ...
##   .. ..@ TPR       : num [1:875] 1 1 1 1 1 1 1 1 1 1 ...
##   .. ..@ TNR       : num [1:875] 0 0 0 0 0 0 0 0 0 0 ...
##   .. ..@ FPR       : num [1:875] 1 1 1 1 1 1 1 1 1 1 ...
##   .. ..@ FNR       : num [1:875] 0 0 0 0 0 0 0 0 0 0 ...
##   .. ..@ PPP       : num [1:875] 0.0503 0.0503 0.0503 0.0503 0.0503 ...
##   .. ..@ NPP       : num [1:875] NaN NaN NaN NaN NaN NaN NaN NaN NaN NaN ...
##   .. ..@ MCR       : num [1:875] 0.95 0.95 0.95 0.95 0.95 ...
##   .. ..@ OR        : num [1:875] NaN NaN NaN NaN NaN NaN NaN NaN NaN NaN ...
##   .. ..@ kappa     : num [1:875] 0 0 0 0 0 0 0 0 0 0 ...
##  $ t    :'data.frame':   1 obs. of  6 variables:
##   ..$ kappa          : num 0.338
##   ..$ spec_sens      : num 0.236
##   ..$ no_omission    : num 0.236
##   ..$ prevalence     : num 0.05
##   ..$ equal_sens_spec: num 0.406
##   ..$ sensitivity    : num 0.43
##  $ v_bkg: num [1:900, 1:20] 101 261 141 152 336 ...
##   ..- attr(*, "dimnames")=List of 2
##   .. ..$ : NULL
##   .. ..$ : chr [1:20] "alt" "bio1" "bio2" "bio3" ...
##  $ v_pnt: num [1:50, 1:20] 2143 1606 1799 2517 1941 ...
##   ..- attr(*, "dimnames")=List of 2
##   .. ..$ : NULL
##   .. ..$ : chr [1:20] "alt" "bio1" "bio2" "bio3" ...
```

##### Evaluate model with fixed control dataset

```
evalutaion_test2 <- evaluate.mdl(puntos_control_extra_coordenadas,puntos_aleatorios_fondo,capas_raster_drop,modelo_maxent)
```

```
## class          : ModelEvaluation 
## n presences    : 72 
## n absences     : 967 
## AUC            : 0.9621395 
## cor            : 0.6465875 
## max TPR+TNR at : 0.3170009
```

```
##                kappa spec_sens no_omission prevalence equal_sens_spec
## thresholds 0.3170009 0.3170009   0.3170009 0.06851768       0.4280867
##            sensitivity
## thresholds   0.4644487
```

```
str(evalutaion_test2)
```

```
## List of 4
##  $ e    :Formal class 'ModelEvaluation' [package "dismo"] with 22 slots
##   .. ..@ presence  : num [1:72] 0.485 0.493 0.626 0.83 0.626 ...
##   .. ..@ absence   : num [1:967] 2.77e-06 1.00e-03 6.38e-05 3.60e-06 1.19e-05 ...
##   .. ..@ np        : int 72
##   .. ..@ na        : int 967
##   .. ..@ auc       : num 0.962
##   .. ..@ pauc      : num(0) 
##   .. ..@ cor       : Named num 0.647
##   .. .. ..- attr(*, "names")= chr "cor"
##   .. ..@ pcor      : num 4.64e-124
##   .. ..@ t         : num [1:883] -1e-04 -1e-04 -1e-04 -1e-04 -1e-04 ...
##   .. ..@ confusion : int [1:883, 1:4] 72 72 72 72 72 72 72 72 72 72 ...
##   .. .. ..- attr(*, "dimnames")=List of 2
##   .. .. .. ..$ : NULL
##   .. .. .. ..$ : chr [1:4] "tp" "fp" "fn" "tn"
##   .. ..@ prevalence: num [1:883] 0.0693 0.0693 0.0693 0.0693 0.0693 ...
##   .. ..@ ODP       : num [1:883] 0.931 0.931 0.931 0.931 0.931 ...
##   .. ..@ CCR       : num [1:883] 0.0693 0.0693 0.0693 0.0693 0.0693 ...
##   .. ..@ TPR       : num [1:883] 1 1 1 1 1 1 1 1 1 1 ...
##   .. ..@ TNR       : num [1:883] 0 0 0 0 0 0 0 0 0 0 ...
##   .. ..@ FPR       : num [1:883] 1 1 1 1 1 1 1 1 1 1 ...
##   .. ..@ FNR       : num [1:883] 0 0 0 0 0 0 0 0 0 0 ...
##   .. ..@ PPP       : num [1:883] 0.0693 0.0693 0.0693 0.0693 0.0693 ...
##   .. ..@ NPP       : num [1:883] NaN NaN NaN NaN NaN NaN NaN NaN NaN NaN ...
##   .. ..@ MCR       : num [1:883] 0.931 0.931 0.931 0.931 0.931 ...
##   .. ..@ OR        : num [1:883] NaN NaN NaN NaN NaN NaN NaN NaN NaN NaN ...
##   .. ..@ kappa     : num [1:883] 0 0 0 0 0 0 0 0 0 0 ...
##  $ t    :'data.frame':   1 obs. of  6 variables:
##   ..$ kappa          : num 0.317
##   ..$ spec_sens      : num 0.317
##   ..$ no_omission    : num 0.317
##   ..$ prevalence     : num 0.0685
##   ..$ equal_sens_spec: num 0.428
##   ..$ sensitivity    : num 0.464
##  $ v_bkg: num [1:900, 1:20] 101 261 141 152 336 ...
##   ..- attr(*, "dimnames")=List of 2
##   .. ..$ : NULL
##   .. ..$ : chr [1:20] "alt" "bio1" "bio2" "bio3" ...
##  $ v_pnt: num [1:72, 1:20] 1252 1946 1793 1871 1793 ...
##   ..- attr(*, "dimnames")=List of 2
##   .. ..$ : NULL
##   .. ..$ : chr [1:20] "alt" "bio1" "bio2" "bio3" ...
```

##### Rank model AUC comparing to 99 null-models (with training dataset)

```
valores_puntos_aleatorios_fondo_y_entrenamiento <- rbind(evalutaion_training$v_pnt, evalutaion_training$v_bkg)

rep <- 99     # number or null models to produce

null_models <- list()
for (r in 1:rep) {
    # following Robert J. Hijmans nullRandom() function from 'dismo'
        # sample presence records and set the rest of records as absence
        index <- sample(nrow(valores_puntos_aleatorios_fondo_y_entrenamiento), presence_points_count)
        pres <- valores_puntos_aleatorios_fondo_y_entrenamiento[index, ]
        absc <- valores_puntos_aleatorios_fondo_y_entrenamiento[-index, ]
        d <- data.frame(rbind(pres, absc))
        v <- c(rep(1, nrow(pres)), rep(0, nrow(absc)))
        # make null-model with presence and absence sets
        m <- maxent(d, v)
        # evaluate the null-model
        null_models[[r]] <- evaluate(pres, absc, m)
}

#null_models
auc_list_vector <- sapply(null_models, function(x) x@auc)

#auc_list_vector
mean(auc_list_vector)
```

```
## [1] 0.7260997
```

```
auc_list_training_vector <- c(evalutaion_training$e@auc, auc_list_vector)
auc_list_training_vector_ranks <- rank(auc_list_training_vector, ties.method= "max")

print('Rank of trainign AUC vs null-models')
```

```
## [1] "Rank of trainign AUC vs null-models"
```

```
auc_list_training_vector_ranks[1]
```

```
## [1] 100
```

```
auc_list_control_vector <- c(evalutaion_test$e@auc, auc_list_vector)
auc_list_control_vector_ranks <- rank(auc_list_control_vector, ties.method= "max")

print('Rank of control AUC vs null-models')
```

```
## [1] "Rank of control AUC vs null-models"
```

```
auc_list_control_vector_ranks[1]
```

```
## [1] 100
```

##### Store the model, parameters and control values in an object for further use

```
container <- list()
container$bkg <- puntos_aleatorios_fondo
container$entr <- puntos_entrenamiento_coordenadas
container$cntrl <- puntos_control_extra_coordenadas
container$model <- modelo_maxent
container$contrib <- contribuciones
container$e_train <- evalutaion_training
container$e_test <- evalutaion_test
container$e_test2 <- evalutaion_test2
container$auc_rank_train <- auc_list_training_vector_ranks[1]
container$auc_rank_test <- auc_list_control_vector_ranks[1]
container$auc_null_model_mean <- mean(auc_list_vector)
  
iterations[[i]] <- container
```
