## Supplementary material for "Vulnerability to climate change for narrowly ranged species: the case of Ecuadorian endemic *Magnolia mercedesiarum*": step_29dup_8.html

```
##                                                                                        [,1]
## X.Training.samples                                                                  50.0000
## Regularized.training.gain                                                            2.5122
## Unregularized.training.gain                                                          2.6819
## Iterations                                                                         500.0000
## Training.AUC                                                                         0.9729
## X.Background.points                                                                947.0000
## alt.contribution                                                                     6.8591
## bio1.contribution                                                                    0.0000
## bio10.contribution                                                                   0.4834
## bio11.contribution                                                                   1.5047
## bio12.contribution                                                                   0.2812
## bio13.contribution                                                                   0.0000
## bio14.contribution                                                                   0.0000
## bio15.contribution                                                                   3.1135
## bio16.contribution                                                                   1.1255
## bio17.contribution                                                                   2.0319
## bio18.contribution                                                                   9.8473
## bio19.contribution                                                                  31.0586
## bio2.contribution                                                                    0.0124
## bio3.contribution                                                                   18.1836
## bio4.contribution                                                                    0.1687
## bio5.contribution                                                                    1.2473
## bio6.contribution                                                                   23.2231
## bio7.contribution                                                                    0.0000
## bio8.contribution                                                                    0.8490
## bio9.contribution                                                                    0.0109
## alt.permutation.importance                                                          16.5894
## bio1.permutation.importance                                                          0.0000
## bio10.permutation.importance                                                         0.7416
## bio11.permutation.importance                                                         0.0000
## bio12.permutation.importance                                                        32.1148
## bio13.permutation.importance                                                         0.0000
## bio14.permutation.importance                                                         0.0000
## bio15.permutation.importance                                                         5.3202
## bio16.permutation.importance                                                         0.0000
## bio17.permutation.importance                                                         0.3789
## bio18.permutation.importance                                                        22.7157
## bio19.permutation.importance                                                         0.0766
## bio2.permutation.importance                                                          0.0000
## bio3.permutation.importance                                                          1.9266
## bio4.permutation.importance                                                          2.9463
## bio5.permutation.importance                                                          0.0121
## bio6.permutation.importance                                                         15.3319
## bio7.permutation.importance                                                          0.0000
## bio8.permutation.importance                                                          0.0000
## bio9.permutation.importance                                                          1.8460
## Training.gain.without.alt                                                            2.4808
## Training.gain.without.bio1                                                           2.5120
## Training.gain.without.bio10                                                          2.5124
## Training.gain.without.bio11                                                          2.5124
## Training.gain.without.bio12                                                          2.5066
## Training.gain.without.bio13                                                          2.5123
## Training.gain.without.bio14                                                          2.5119
## Training.gain.without.bio15                                                          2.4989
## Training.gain.without.bio16                                                          2.5122
## Training.gain.without.bio17                                                          2.5119
## Training.gain.without.bio18                                                          2.4366
## Training.gain.without.bio19                                                          2.5123
## Training.gain.without.bio2                                                           2.5122
## Training.gain.without.bio3                                                           2.4945
## Training.gain.without.bio4                                                           2.5076
## Training.gain.without.bio5                                                           2.5123
## Training.gain.without.bio6                                                           2.5122
## Training.gain.without.bio7                                                           2.5121
## Training.gain.without.bio8                                                           2.5124
## Training.gain.without.bio9                                                           2.5120
## Training.gain.with.only.alt                                                          1.4735
## Training.gain.with.only.bio1                                                         1.3111
## Training.gain.with.only.bio10                                                        1.3416
## Training.gain.with.only.bio11                                                        1.2582
## Training.gain.with.only.bio12                                                        0.2289
## Training.gain.with.only.bio13                                                        0.1019
## Training.gain.with.only.bio14                                                        0.9906
## Training.gain.with.only.bio15                                                        0.4288
## Training.gain.with.only.bio16                                                        0.1159
## Training.gain.with.only.bio17                                                        0.9496
## Training.gain.with.only.bio18                                                        0.5244
## Training.gain.with.only.bio19                                                        0.3676
## Training.gain.with.only.bio2                                                         0.0315
## Training.gain.with.only.bio3                                                         0.5587
## Training.gain.with.only.bio4                                                         0.7081
## Training.gain.with.only.bio5                                                         1.3207
## Training.gain.with.only.bio6                                                         1.2451
## Training.gain.with.only.bio7                                                         0.1369
## Training.gain.with.only.bio8                                                         1.3414
## Training.gain.with.only.bio9                                                         1.2880
## Entropy                                                                              4.3436
## Prevalence..average.probability.of.presence.over.background.sites.                   0.0472
## Fixed.cumulative.value.1.cumulative.threshold                                        1.0000
## Fixed.cumulative.value.1.Cloglog.threshold                                           0.0088
## Fixed.cumulative.value.1.area                                                        0.1943
## Fixed.cumulative.value.1.training.omission                                           0.0000
## Fixed.cumulative.value.5.cumulative.threshold                                        5.0000
## Fixed.cumulative.value.5.Cloglog.threshold                                           0.0837
## Fixed.cumulative.value.5.area                                                        0.0855
## Fixed.cumulative.value.5.training.omission                                           0.0000
## Fixed.cumulative.value.10.cumulative.threshold                                      10.0000
## Fixed.cumulative.value.10.Cloglog.threshold                                          0.2798
## Fixed.cumulative.value.10.area                                                       0.0602
## Fixed.cumulative.value.10.training.omission                                          0.0200
## Minimum.training.presence.cumulative.threshold                                       9.1655
## Minimum.training.presence.Cloglog.threshold                                          0.2278
## Minimum.training.presence.area                                                       0.0623
## Minimum.training.presence.training.omission                                          0.0000
## X10.percentile.training.presence.cumulative.threshold                               16.5830
## X10.percentile.training.presence.Cloglog.threshold                                   0.5020
## X10.percentile.training.presence.area                                                0.0486
## X10.percentile.training.presence.training.omission                                   0.1000
## Equal.training.sensitivity.and.specificity.cumulative.threshold                     13.4589
## Equal.training.sensitivity.and.specificity.Cloglog.threshold                         0.3446
## Equal.training.sensitivity.and.specificity.area                                      0.0539
## Equal.training.sensitivity.and.specificity.training.omission                         0.0600
## Maximum.training.sensitivity.plus.specificity.cumulative.threshold                   9.1655
## Maximum.training.sensitivity.plus.specificity.Cloglog.threshold                      0.2278
## Maximum.training.sensitivity.plus.specificity.area                                   0.0623
## Maximum.training.sensitivity.plus.specificity.training.omission                      0.0000
## Balance.training.omission..predicted.area.and.threshold.value.cumulative.threshold   2.6449
## Balance.training.omission..predicted.area.and.threshold.value.Cloglog.threshold      0.0318
## Balance.training.omission..predicted.area.and.threshold.value.area                   0.1193
## Balance.training.omission..predicted.area.and.threshold.value.training.omission      0.0000
## Equate.entropy.of.thresholded.and.original.distributions.cumulative.threshold        5.6418
## Equate.entropy.of.thresholded.and.original.distributions.Cloglog.threshold           0.0999
## Equate.entropy.of.thresholded.and.original.distributions.area                        0.0803
## Equate.entropy.of.thresholded.and.original.distributions.training.omission           0.0000
```

```
#str(modelo_maxent@results)
```

##### Get variable importance values

```
#plot(modelo_maxent)

contribuciones <- maxent.select.contribution(modelo_maxent,names_after_drop)

#contribuciones
par(mfrow=c(1, 2))
dotchart(as.numeric(contribuciones[,2]),
        col='blue', pch=16,
        labels=contribuciones[,1],
        main='predictor contribution')
dotchart(as.numeric(contribuciones[,3]),
        col='red', pch=16,
        labels=contribuciones[,1],
        main='predictor importance in permutations')
```

##### Draw respnse cuves

```
response(modelo_maxent)
```

##### Output MaxEnt HTML

```
#modelo_maxent
```

### Initial model evaluation

##### Evaluate model with training dataset

```
evalutaion_training <- evaluate.mdl(puntos_entrenamiento_coordenadas,puntos_aleatorios_fondo,capas_raster_drop,modelo_maxent)
```

```
## class          : ModelEvaluation 
## n presences    : 50 
## n absences     : 947 
## AUC            : 0.972925 
## cor            : 0.6552535 
## max TPR+TNR at : 0.227728
```

```
##                kappa spec_sens no_omission prevalence equal_sens_spec
## thresholds 0.3445351  0.227728    0.227728 0.05104714       0.3574521
##            sensitivity
## thresholds   0.5018732
```

```
str(evalutaion_training)
```

```
## List of 4
##  $ e    :Formal class 'ModelEvaluation' [package "dismo"] with 22 slots
##   .. ..@ presence  : num [1:50] 0.228 0.689 0.664 0.587 0.815 ...
##   .. ..@ absence   : num [1:947] 6.34e-04 8.25e-04 1.70e-04 8.28e-05 1.71e-04 ...
##   .. ..@ np        : int 50
##   .. ..@ na        : int 947
##   .. ..@ auc       : num 0.973
##   .. ..@ pauc      : num(0) 
##   .. ..@ cor       : Named num 0.655
##   .. .. ..- attr(*, "names")= chr "cor"
##   .. ..@ pcor      : num 2.38e-123
##   .. ..@ t         : num [1:872] -1.00e-04 -1.00e-04 -1.00e-04 -1.00e-04 -9.99e-05 ...
##   .. ..@ confusion : int [1:872, 1:4] 50 50 50 50 50 50 50 50 50 50 ...
##   .. .. ..- attr(*, "dimnames")=List of 2
##   .. .. .. ..$ : NULL
##   .. .. .. ..$ : chr [1:4] "tp" "fp" "fn" "tn"
##   .. ..@ prevalence: num [1:872] 0.0502 0.0502 0.0502 0.0502 0.0502 ...
##   .. ..@ ODP       : num [1:872] 0.95 0.95 0.95 0.95 0.95 ...
##   .. ..@ CCR       : num [1:872] 0.0502 0.0502 0.0502 0.0502 0.0502 ...
##   .. ..@ TPR       : num [1:872] 1 1 1 1 1 1 1 1 1 1 ...
##   .. ..@ TNR       : num [1:872] 0 0 0 0 0 0 0 0 0 0 ...
##   .. ..@ FPR       : num [1:872] 1 1 1 1 1 1 1 1 1 1 ...
##   .. ..@ FNR       : num [1:872] 0 0 0 0 0 0 0 0 0 0 ...
##   .. ..@ PPP       : num [1:872] 0.0502 0.0502 0.0502 0.0502 0.0502 ...
##   .. ..@ NPP       : num [1:872] NaN NaN NaN NaN NaN NaN NaN NaN NaN NaN ...
##   .. ..@ MCR       : num [1:872] 0.95 0.95 0.95 0.95 0.95 ...
##   .. ..@ OR        : num [1:872] NaN NaN NaN NaN NaN NaN NaN NaN NaN NaN ...
##   .. ..@ kappa     : num [1:872] 0 0 0 0 0 0 0 0 0 0 ...
##  $ t    :'data.frame':   1 obs. of  6 variables:
##   ..$ kappa          : num 0.345
##   ..$ spec_sens      : num 0.228
##   ..$ no_omission    : num 0.228
##   ..$ prevalence     : num 0.051
##   ..$ equal_sens_spec: num 0.357
##   ..$ sensitivity    : num 0.502
##  $ v_bkg: num [1:900, 1:20] 2818 184 269 282 212 ...
##   ..- attr(*, "dimnames")=List of 2
##   .. ..$ : NULL
##   .. ..$ : chr [1:20] "alt" "bio1" "bio2" "bio3" ...
##  $ v_pnt: num [1:50, 1:20] 1385 1814 1536 1324 1929 ...
##   ..- attr(*, "dimnames")=List of 2
##   .. ..$ : NULL
##   .. ..$ : chr [1:20] "alt" "bio1" "bio2" "bio3" ...
```

##### Evaluate model with control dataset

```
evalutaion_test <- evaluate.mdl(puntos_control_coordenadas,puntos_aleatorios_fondo,capas_raster_drop,modelo_maxent)
```

```
## class          : ModelEvaluation 
## n presences    : 50 
## n absences     : 947 
## AUC            : 0.973094 
## cor            : 0.6593611 
## max TPR+TNR at : 0.324187
```

```
##                kappa spec_sens no_omission prevalence equal_sens_spec
## thresholds 0.4503702  0.324187    0.324187 0.05104714       0.4786613
##            sensitivity
## thresholds   0.4906887
```

```
str(evalutaion_test)
```

```
## List of 4
##  $ e    :Formal class 'ModelEvaluation' [package "dismo"] with 22 slots
##   .. ..@ presence  : num [1:50] 0.724 0.781 0.616 0.901 0.679 ...
##   .. ..@ absence   : num [1:947] 6.34e-04 8.25e-04 1.70e-04 8.28e-05 1.71e-04 ...
##   .. ..@ np        : int 50
##   .. ..@ na        : int 947
##   .. ..@ auc       : num 0.973
##   .. ..@ pauc      : num(0) 
##   .. ..@ cor       : Named num 0.659
##   .. .. ..- attr(*, "names")= chr "cor"
##   .. ..@ pcor      : num 2.09e-125
##   .. ..@ t         : num [1:872] -1.00e-04 -1.00e-04 -1.00e-04 -1.00e-04 -9.99e-05 ...
##   .. ..@ confusion : int [1:872, 1:4] 50 50 50 50 50 50 50 50 50 50 ...
##   .. .. ..- attr(*, "dimnames")=List of 2
##   .. .. .. ..$ : NULL
##   .. .. .. ..$ : chr [1:4] "tp" "fp" "fn" "tn"
##   .. ..@ prevalence: num [1:872] 0.0502 0.0502 0.0502 0.0502 0.0502 ...
##   .. ..@ ODP       : num [1:872] 0.95 0.95 0.95 0.95 0.95 ...
##   .. ..@ CCR       : num [1:872] 0.0502 0.0502 0.0502 0.0502 0.0502 ...
##   .. ..@ TPR       : num [1:872] 1 1 1 1 1 1 1 1 1 1 ...
##   .. ..@ TNR       : num [1:872] 0 0 0 0 0 0 0 0 0 0 ...
##   .. ..@ FPR       : num [1:872] 1 1 1 1 1 1 1 1 1 1 ...
##   .. ..@ FNR       : num [1:872] 0 0 0 0 0 0 0 0 0 0 ...
##   .. ..@ PPP       : num [1:872] 0.0502 0.0502 0.0502 0.0502 0.0502 ...
##   .. ..@ NPP       : num [1:872] NaN NaN NaN NaN NaN NaN NaN NaN NaN NaN ...
##   .. ..@ MCR       : num [1:872] 0.95 0.95 0.95 0.95 0.95 ...
##   .. ..@ OR        : num [1:872] NaN NaN NaN NaN NaN NaN NaN NaN NaN NaN ...
##   .. ..@ kappa     : num [1:872] 0 0 0 0 0 0 0 0 0 0 ...
##  $ t    :'data.frame':   1 obs. of  6 variables:
##   ..$ kappa          : num 0.45
##   ..$ spec_sens      : num 0.324
##   ..$ no_omission    : num 0.324
##   ..$ prevalence     : num 0.051
##   ..$ equal_sens_spec: num 0.479
##   ..$ sensitivity    : num 0.491
##  $ v_bkg: num [1:900, 1:20] 2818 184 269 282 212 ...
##   ..- attr(*, "dimnames")=List of 2
##   .. ..$ : NULL
##   .. ..$ : chr [1:20] "alt" "bio1" "bio2" "bio3" ...
##  $ v_pnt: num [1:50, 1:20] 1573 2264 1442 1663 2042 ...
##   ..- attr(*, "dimnames")=List of 2
##   .. ..$ : NULL
##   .. ..$ : chr [1:20] "alt" "bio1" "bio2" "bio3" ...
```

##### Evaluate model with fixed control dataset

```
evalutaion_test2 <- evaluate.mdl(puntos_control_extra_coordenadas,puntos_aleatorios_fondo,capas_raster_drop,modelo_maxent)
```

```
## class          : ModelEvaluation 
## n presences    : 72 
## n absences     : 969 
## AUC            : 0.9618593 
## cor            : 0.6475836 
## max TPR+TNR at : 0.3578348
```

```
##                kappa spec_sens no_omission prevalence equal_sens_spec
## thresholds 0.3578348 0.3578348   0.3578348 0.06893302       0.4323911
##            sensitivity
## thresholds   0.4567688
```

```
str(evalutaion_test2)
```

```
## List of 4
##  $ e    :Formal class 'ModelEvaluation' [package "dismo"] with 22 slots
##   .. ..@ presence  : num [1:72] 0.488 0.494 0.723 0.8 0.723 ...
##   .. ..@ absence   : num [1:969] 6.34e-04 8.25e-04 1.70e-04 8.28e-05 1.71e-04 ...
##   .. ..@ np        : int 72
##   .. ..@ na        : int 969
##   .. ..@ auc       : num 0.962
##   .. ..@ pauc      : num(0) 
##   .. ..@ cor       : Named num 0.648
##   .. .. ..- attr(*, "names")= chr "cor"
##   .. ..@ pcor      : num 8.5e-125
##   .. ..@ t         : num [1:880] -1.00e-04 -1.00e-04 -1.00e-04 -1.00e-04 -9.99e-05 ...
##   .. ..@ confusion : int [1:880, 1:4] 72 72 72 72 72 72 72 72 72 72 ...
##   .. .. ..- attr(*, "dimnames")=List of 2
##   .. .. .. ..$ : NULL
##   .. .. .. ..$ : chr [1:4] "tp" "fp" "fn" "tn"
##   .. ..@ prevalence: num [1:880] 0.0692 0.0692 0.0692 0.0692 0.0692 ...
##   .. ..@ ODP       : num [1:880] 0.931 0.931 0.931 0.931 0.931 ...
##   .. ..@ CCR       : num [1:880] 0.0692 0.0692 0.0692 0.0692 0.0692 ...
##   .. ..@ TPR       : num [1:880] 1 1 1 1 1 1 1 1 1 1 ...
##   .. ..@ TNR       : num [1:880] 0 0 0 0 0 0 0 0 0 0 ...
##   .. ..@ FPR       : num [1:880] 1 1 1 1 1 1 1 1 1 1 ...
##   .. ..@ FNR       : num [1:880] 0 0 0 0 0 0 0 0 0 0 ...
##   .. ..@ PPP       : num [1:880] 0.0692 0.0692 0.0692 0.0692 0.0692 ...
##   .. ..@ NPP       : num [1:880] NaN NaN NaN NaN NaN NaN NaN NaN NaN NaN ...
##   .. ..@ MCR       : num [1:880] 0.931 0.931 0.931 0.931 0.931 ...
##   .. ..@ OR        : num [1:880] NaN NaN NaN NaN NaN NaN NaN NaN NaN NaN ...
##   .. ..@ kappa     : num [1:880] 0 0 0 0 0 0 0 0 0 0 ...
##  $ t    :'data.frame':   1 obs. of  6 variables:
##   ..$ kappa          : num 0.358
##   ..$ spec_sens      : num 0.358
##   ..$ no_omission    : num 0.358
##   ..$ prevalence     : num 0.0689
##   ..$ equal_sens_spec: num 0.432
##   ..$ sensitivity    : num 0.457
##  $ v_bkg: num [1:900, 1:20] 2818 184 269 282 212 ...
##   ..- attr(*, "dimnames")=List of 2
##   .. ..$ : NULL
##   .. ..$ : chr [1:20] "alt" "bio1" "bio2" "bio3" ...
##  $ v_pnt: num [1:72, 1:20] 1252 1946 1793 1871 1793 ...
##   ..- attr(*, "dimnames")=List of 2
##   .. ..$ : NULL
##   .. ..$ : chr [1:20] "alt" "bio1" "bio2" "bio3" ...
```

##### Rank model AUC comparing to 99 null-models (with training dataset)

```
valores_puntos_aleatorios_fondo_y_entrenamiento <- rbind(evalutaion_training$v_pnt, evalutaion_training$v_bkg)

rep <- 99     # number or null models to produce

null_models <- list()
for (r in 1:rep) {
    # following Robert J. Hijmans nullRandom() function from 'dismo'
        # sample presence records and set the rest of records as absence
        index <- sample(nrow(valores_puntos_aleatorios_fondo_y_entrenamiento), presence_points_count)
        pres <- valores_puntos_aleatorios_fondo_y_entrenamiento[index, ]
        absc <- valores_puntos_aleatorios_fondo_y_entrenamiento[-index, ]
        d <- data.frame(rbind(pres, absc))
        v <- c(rep(1, nrow(pres)), rep(0, nrow(absc)))
        # make null-model with presence and absence sets
        m <- maxent(d, v)
        # evaluate the null-model
        null_models[[r]] <- evaluate(pres, absc, m)
}

#null_models
auc_list_vector <- sapply(null_models, function(x) x@auc)

#auc_list_vector
mean(auc_list_vector)
```

```
## [1] 0.7274973
```

```
auc_list_training_vector <- c(evalutaion_training$e@auc, auc_list_vector)
auc_list_training_vector_ranks <- rank(auc_list_training_vector, ties.method= "max")

print('Rank of trainign AUC vs null-models')
```

```
## [1] "Rank of trainign AUC vs null-models"
```

```
auc_list_training_vector_ranks[1]
```

```
## [1] 100
```

```
auc_list_control_vector <- c(evalutaion_test$e@auc, auc_list_vector)
auc_list_control_vector_ranks <- rank(auc_list_control_vector, ties.method= "max")

print('Rank of control AUC vs null-models')
```

```
## [1] "Rank of control AUC vs null-models"
```

```
auc_list_control_vector_ranks[1]
```

```
## [1] 100
```

##### Store the model, parameters and control values in an object for further use

```
container <- list()
container$bkg <- puntos_aleatorios_fondo
container$entr <- puntos_entrenamiento_coordenadas
container$cntrl <- puntos_control_extra_coordenadas
container$model <- modelo_maxent
container$contrib <- contribuciones
container$e_train <- evalutaion_training
container$e_test <- evalutaion_test
container$e_test2 <- evalutaion_test2
container$auc_rank_train <- auc_list_training_vector_ranks[1]
container$auc_rank_test <- auc_list_control_vector_ranks[1]
container$auc_null_model_mean <- mean(auc_list_vector)
  
iterations[[i]] <- container
```
