## Supplementary material for "Vulnerability to climate change for narrowly ranged species: the case of Ecuadorian endemic *Magnolia mercedesiarum*": step_29dup_9.html

```
##                                                                                        [,1]
## X.Training.samples                                                                  50.0000
## Regularized.training.gain                                                            2.5072
## Unregularized.training.gain                                                          2.6599
## Iterations                                                                         500.0000
## Training.AUC                                                                         0.9723
## X.Background.points                                                                949.0000
## alt.contribution                                                                     4.6135
## bio1.contribution                                                                    0.0000
## bio10.contribution                                                                   0.0233
## bio11.contribution                                                                   1.9436
## bio12.contribution                                                                   0.4621
## bio13.contribution                                                                   0.0000
## bio14.contribution                                                                   0.0000
## bio15.contribution                                                                   5.9715
## bio16.contribution                                                                   0.0000
## bio17.contribution                                                                   0.0000
## bio18.contribution                                                                  10.4031
## bio19.contribution                                                                  27.2715
## bio2.contribution                                                                    0.0163
## bio3.contribution                                                                   21.4541
## bio4.contribution                                                                    0.0011
## bio5.contribution                                                                    0.3240
## bio6.contribution                                                                   26.9038
## bio7.contribution                                                                    0.0000
## bio8.contribution                                                                    0.0000
## bio9.contribution                                                                    0.6120
## alt.permutation.importance                                                          11.3338
## bio1.permutation.importance                                                          0.0000
## bio10.permutation.importance                                                        32.1367
## bio11.permutation.importance                                                         0.0000
## bio12.permutation.importance                                                        17.8579
## bio13.permutation.importance                                                         0.0000
## bio14.permutation.importance                                                         0.0000
## bio15.permutation.importance                                                         2.1539
## bio16.permutation.importance                                                         0.0000
## bio17.permutation.importance                                                         0.0000
## bio18.permutation.importance                                                        32.3314
## bio19.permutation.importance                                                         0.0429
## bio2.permutation.importance                                                          0.0000
## bio3.permutation.importance                                                          3.1362
## bio4.permutation.importance                                                          0.0572
## bio5.permutation.importance                                                          0.0000
## bio6.permutation.importance                                                          0.9501
## bio7.permutation.importance                                                          0.0000
## bio8.permutation.importance                                                          0.0000
## bio9.permutation.importance                                                          0.0000
## Training.gain.without.alt                                                            2.4659
## Training.gain.without.bio1                                                           2.5068
## Training.gain.without.bio10                                                          2.5064
## Training.gain.without.bio11                                                          2.5067
## Training.gain.without.bio12                                                          2.4956
## Training.gain.without.bio13                                                          2.5069
## Training.gain.without.bio14                                                          2.5066
## Training.gain.without.bio15                                                          2.4956
## Training.gain.without.bio16                                                          2.5069
## Training.gain.without.bio17                                                          2.5067
## Training.gain.without.bio18                                                          2.3752
## Training.gain.without.bio19                                                          2.5064
## Training.gain.without.bio2                                                           2.5066
## Training.gain.without.bio3                                                           2.4674
## Training.gain.without.bio4                                                           2.5061
## Training.gain.without.bio5                                                           2.5069
## Training.gain.without.bio6                                                           2.5071
## Training.gain.without.bio7                                                           2.5065
## Training.gain.without.bio8                                                           2.5064
## Training.gain.without.bio9                                                           2.5069
## Training.gain.with.only.alt                                                          1.5519
## Training.gain.with.only.bio1                                                         1.4140
## Training.gain.with.only.bio10                                                        1.4419
## Training.gain.with.only.bio11                                                        1.3727
## Training.gain.with.only.bio12                                                        0.2199
## Training.gain.with.only.bio13                                                        0.1220
## Training.gain.with.only.bio14                                                        0.9392
## Training.gain.with.only.bio15                                                        0.4566
## Training.gain.with.only.bio16                                                        0.1270
## Training.gain.with.only.bio17                                                        0.8416
## Training.gain.with.only.bio18                                                        0.6827
## Training.gain.with.only.bio19                                                        0.3812
## Training.gain.with.only.bio2                                                         0.0562
## Training.gain.with.only.bio3                                                         0.6921
## Training.gain.with.only.bio4                                                         0.7007
## Training.gain.with.only.bio5                                                         1.4191
## Training.gain.with.only.bio6                                                         1.3615
## Training.gain.with.only.bio7                                                         0.0752
## Training.gain.with.only.bio8                                                         1.4271
## Training.gain.with.only.bio9                                                         1.3957
## Entropy                                                                              4.3506
## Prevalence..average.probability.of.presence.over.background.sites.                   0.0472
## Fixed.cumulative.value.1.cumulative.threshold                                        1.0000
## Fixed.cumulative.value.1.Cloglog.threshold                                           0.0075
## Fixed.cumulative.value.1.area                                                        0.2086
## Fixed.cumulative.value.1.training.omission                                           0.0000
## Fixed.cumulative.value.5.cumulative.threshold                                        5.0000
## Fixed.cumulative.value.5.Cloglog.threshold                                           0.0793
## Fixed.cumulative.value.5.area                                                        0.0896
## Fixed.cumulative.value.5.training.omission                                           0.0000
## Fixed.cumulative.value.10.cumulative.threshold                                      10.0000
## Fixed.cumulative.value.10.Cloglog.threshold                                          0.1975
## Fixed.cumulative.value.10.area                                                       0.0601
## Fixed.cumulative.value.10.training.omission                                          0.0000
## Minimum.training.presence.cumulative.threshold                                      11.0317
## Minimum.training.presence.Cloglog.threshold                                          0.2914
## Minimum.training.presence.area                                                       0.0580
## Minimum.training.presence.training.omission                                          0.0000
## X10.percentile.training.presence.cumulative.threshold                               14.2928
## X10.percentile.training.presence.Cloglog.threshold                                   0.4270
## X10.percentile.training.presence.area                                                0.0516
## X10.percentile.training.presence.training.omission                                   0.1000
## Equal.training.sensitivity.and.specificity.cumulative.threshold                     12.2427
## Equal.training.sensitivity.and.specificity.Cloglog.threshold                         0.3904
## Equal.training.sensitivity.and.specificity.area                                      0.0558
## Equal.training.sensitivity.and.specificity.training.omission                         0.0600
## Maximum.training.sensitivity.plus.specificity.cumulative.threshold                  11.0317
## Maximum.training.sensitivity.plus.specificity.Cloglog.threshold                      0.2914
## Maximum.training.sensitivity.plus.specificity.area                                   0.0580
## Maximum.training.sensitivity.plus.specificity.training.omission                      0.0000
## Balance.training.omission..predicted.area.and.threshold.value.cumulative.threshold   2.7720
## Balance.training.omission..predicted.area.and.threshold.value.Cloglog.threshold      0.0320
## Balance.training.omission..predicted.area.and.threshold.value.area                   0.1254
## Balance.training.omission..predicted.area.and.threshold.value.training.omission      0.0000
## Equate.entropy.of.thresholded.and.original.distributions.cumulative.threshold        6.0608
## Equate.entropy.of.thresholded.and.original.distributions.Cloglog.threshold           0.1059
## Equate.entropy.of.thresholded.and.original.distributions.area                        0.0811
## Equate.entropy.of.thresholded.and.original.distributions.training.omission           0.0000
```

```
#str(modelo_maxent@results)
```

##### Get variable importance values

```
#plot(modelo_maxent)

contribuciones <- maxent.select.contribution(modelo_maxent,names_after_drop)

#contribuciones
par(mfrow=c(1, 2))
dotchart(as.numeric(contribuciones[,2]),
        col='blue', pch=16,
        labels=contribuciones[,1],
        main='predictor contribution')
dotchart(as.numeric(contribuciones[,3]),
        col='red', pch=16,
        labels=contribuciones[,1],
        main='predictor importance in permutations')
```

##### Draw respnse cuves

```
response(modelo_maxent)
```

##### Output MaxEnt HTML

```
#modelo_maxent
```

### Initial model evaluation

##### Evaluate model with training dataset

```
evalutaion_training <- evaluate.mdl(puntos_entrenamiento_coordenadas,puntos_aleatorios_fondo,capas_raster_drop,modelo_maxent)
```

```
## class          : ModelEvaluation 
## n presences    : 50 
## n absences     : 949 
## AUC            : 0.9722866 
## cor            : 0.6515058 
## max TPR+TNR at : 0.2912626
```

```
##                kappa spec_sens no_omission prevalence equal_sens_spec
## thresholds 0.2912626 0.2912626   0.2912626 0.05022478       0.4025569
##            sensitivity
## thresholds   0.4269137
```

```
str(evalutaion_training)
```

```
## List of 4
##  $ e    :Formal class 'ModelEvaluation' [package "dismo"] with 22 slots
##   .. ..@ presence  : num [1:50] 0.659 0.751 0.624 0.761 0.745 ...
##   .. ..@ absence   : num [1:949] 1.40e-04 3.38e-05 9.98e-06 1.23e-04 5.25e-04 ...
##   .. ..@ np        : int 50
##   .. ..@ na        : int 949
##   .. ..@ auc       : num 0.972
##   .. ..@ pauc      : num(0) 
##   .. ..@ cor       : Named num 0.652
##   .. .. ..- attr(*, "names")= chr "cor"
##   .. ..@ pcor      : num 9.68e-122
##   .. ..@ t         : num [1:871] -1e-04 -1e-04 -1e-04 -1e-04 -1e-04 ...
##   .. ..@ confusion : int [1:871, 1:4] 50 50 50 50 50 50 50 50 50 50 ...
##   .. .. ..- attr(*, "dimnames")=List of 2
##   .. .. .. ..$ : NULL
##   .. .. .. ..$ : chr [1:4] "tp" "fp" "fn" "tn"
##   .. ..@ prevalence: num [1:871] 0.0501 0.0501 0.0501 0.0501 0.0501 ...
##   .. ..@ ODP       : num [1:871] 0.95 0.95 0.95 0.95 0.95 ...
##   .. ..@ CCR       : num [1:871] 0.0501 0.0501 0.0501 0.0501 0.0501 ...
##   .. ..@ TPR       : num [1:871] 1 1 1 1 1 1 1 1 1 1 ...
##   .. ..@ TNR       : num [1:871] 0 0 0 0 0 0 0 0 0 0 ...
##   .. ..@ FPR       : num [1:871] 1 1 1 1 1 1 1 1 1 1 ...
##   .. ..@ FNR       : num [1:871] 0 0 0 0 0 0 0 0 0 0 ...
##   .. ..@ PPP       : num [1:871] 0.0501 0.0501 0.0501 0.0501 0.0501 ...
##   .. ..@ NPP       : num [1:871] NaN NaN NaN NaN NaN NaN NaN NaN NaN NaN ...
##   .. ..@ MCR       : num [1:871] 0.95 0.95 0.95 0.95 0.95 ...
##   .. ..@ OR        : num [1:871] NaN NaN NaN NaN NaN NaN NaN NaN NaN NaN ...
##   .. ..@ kappa     : num [1:871] 0 0 0 0 0 0 0 0 0 0 ...
##  $ t    :'data.frame':   1 obs. of  6 variables:
##   ..$ kappa          : num 0.291
##   ..$ spec_sens      : num 0.291
##   ..$ no_omission    : num 0.291
##   ..$ prevalence     : num 0.0502
##   ..$ equal_sens_spec: num 0.403
##   ..$ sensitivity    : num 0.427
##  $ v_bkg: num [1:900, 1:20] 190 577 123 102 3155 ...
##   ..- attr(*, "dimnames")=List of 2
##   .. ..$ : NULL
##   .. ..$ : chr [1:20] "alt" "bio1" "bio2" "bio3" ...
##  $ v_pnt: num [1:50, 1:20] 2135 1928 1275 1490 1784 ...
##   ..- attr(*, "dimnames")=List of 2
##   .. ..$ : NULL
##   .. ..$ : chr [1:20] "alt" "bio1" "bio2" "bio3" ...
```

##### Evaluate model with control dataset

```
evalutaion_test <- evaluate.mdl(puntos_control_coordenadas,puntos_aleatorios_fondo,capas_raster_drop,modelo_maxent)
```

```
## class          : ModelEvaluation 
## n presences    : 50 
## n absences     : 949 
## AUC            : 0.9711697 
## cor            : 0.6405544 
## max TPR+TNR at : 0.08720537
```

```
##                kappa  spec_sens no_omission prevalence equal_sens_spec
## thresholds 0.3251962 0.08720537  0.08720537 0.05022478       0.3406139
##            sensitivity
## thresholds   0.3946025
```

```
str(evalutaion_test)
```

```
## List of 4
##  $ e    :Formal class 'ModelEvaluation' [package "dismo"] with 22 slots
##   .. ..@ presence  : num [1:50] 0.52 0.732 0.843 0.648 0.934 ...
##   .. ..@ absence   : num [1:949] 1.40e-04 3.38e-05 9.98e-06 1.23e-04 5.25e-04 ...
##   .. ..@ np        : int 50
##   .. ..@ na        : int 949
##   .. ..@ auc       : num 0.971
##   .. ..@ pauc      : num(0) 
##   .. ..@ cor       : Named num 0.641
##   .. .. ..- attr(*, "names")= chr "cor"
##   .. ..@ pcor      : num 1.78e-116
##   .. ..@ t         : num [1:871] -1e-04 -1e-04 -1e-04 -1e-04 -1e-04 ...
##   .. ..@ confusion : int [1:871, 1:4] 50 50 50 50 50 50 50 50 50 50 ...
##   .. .. ..- attr(*, "dimnames")=List of 2
##   .. .. .. ..$ : NULL
##   .. .. .. ..$ : chr [1:4] "tp" "fp" "fn" "tn"
##   .. ..@ prevalence: num [1:871] 0.0501 0.0501 0.0501 0.0501 0.0501 ...
##   .. ..@ ODP       : num [1:871] 0.95 0.95 0.95 0.95 0.95 ...
##   .. ..@ CCR       : num [1:871] 0.0501 0.0501 0.0501 0.0501 0.0501 ...
##   .. ..@ TPR       : num [1:871] 1 1 1 1 1 1 1 1 1 1 ...
##   .. ..@ TNR       : num [1:871] 0 0 0 0 0 0 0 0 0 0 ...
##   .. ..@ FPR       : num [1:871] 1 1 1 1 1 1 1 1 1 1 ...
##   .. ..@ FNR       : num [1:871] 0 0 0 0 0 0 0 0 0 0 ...
##   .. ..@ PPP       : num [1:871] 0.0501 0.0501 0.0501 0.0501 0.0501 ...
##   .. ..@ NPP       : num [1:871] NaN NaN NaN NaN NaN NaN NaN NaN NaN NaN ...
##   .. ..@ MCR       : num [1:871] 0.95 0.95 0.95 0.95 0.95 ...
##   .. ..@ OR        : num [1:871] NaN NaN NaN NaN NaN NaN NaN NaN NaN NaN ...
##   .. ..@ kappa     : num [1:871] 0 0 0 0 0 0 0 0 0 0 ...
##  $ t    :'data.frame':   1 obs. of  6 variables:
##   ..$ kappa          : num 0.325
##   ..$ spec_sens      : num 0.0872
##   ..$ no_omission    : num 0.0872
##   ..$ prevalence     : num 0.0502
##   ..$ equal_sens_spec: num 0.341
##   ..$ sensitivity    : num 0.395
##  $ v_bkg: num [1:900, 1:20] 190 577 123 102 3155 ...
##   ..- attr(*, "dimnames")=List of 2
##   .. ..$ : NULL
##   .. ..$ : chr [1:20] "alt" "bio1" "bio2" "bio3" ...
##  $ v_pnt: num [1:50, 1:20] 1742 1592 1604 2001 2220 ...
##   ..- attr(*, "dimnames")=List of 2
##   .. ..$ : NULL
##   .. ..$ : chr [1:20] "alt" "bio1" "bio2" "bio3" ...
```

##### Evaluate model with fixed control dataset

```
evalutaion_test2 <- evaluate.mdl(puntos_control_extra_coordenadas,puntos_aleatorios_fondo,capas_raster_drop,modelo_maxent)
```

```
## class          : ModelEvaluation 
## n presences    : 72 
## n absences     : 971 
## AUC            : 0.9608508 
## cor            : 0.6429759 
## max TPR+TNR at : 0.3738234
```

```
##                kappa spec_sens no_omission prevalence equal_sens_spec
## thresholds 0.3738234 0.3738234   0.3738234 0.06931017       0.4427548
##            sensitivity
## thresholds   0.4571416
```

```
str(evalutaion_test2)
```

```
## List of 4
##  $ e    :Formal class 'ModelEvaluation' [package "dismo"] with 22 slots
##   .. ..@ presence  : num [1:72] 0.571 0.463 0.602 0.86 0.602 ...
##   .. ..@ absence   : num [1:971] 1.40e-04 3.38e-05 9.98e-06 1.23e-04 5.25e-04 ...
##   .. ..@ np        : int 72
##   .. ..@ na        : int 971
##   .. ..@ auc       : num 0.961
##   .. ..@ pauc      : num(0) 
##   .. ..@ cor       : Named num 0.643
##   .. .. ..- attr(*, "names")= chr "cor"
##   .. ..@ pcor      : num 9.99e-123
##   .. ..@ t         : num [1:879] -1e-04 -1e-04 -1e-04 -1e-04 -1e-04 ...
##   .. ..@ confusion : int [1:879, 1:4] 72 72 72 72 72 72 72 72 72 72 ...
##   .. .. ..- attr(*, "dimnames")=List of 2
##   .. .. .. ..$ : NULL
##   .. .. .. ..$ : chr [1:4] "tp" "fp" "fn" "tn"
##   .. ..@ prevalence: num [1:879] 0.069 0.069 0.069 0.069 0.069 ...
##   .. ..@ ODP       : num [1:879] 0.931 0.931 0.931 0.931 0.931 ...
##   .. ..@ CCR       : num [1:879] 0.069 0.069 0.069 0.069 0.069 ...
##   .. ..@ TPR       : num [1:879] 1 1 1 1 1 1 1 1 1 1 ...
##   .. ..@ TNR       : num [1:879] 0 0 0 0 0 0 0 0 0 0 ...
##   .. ..@ FPR       : num [1:879] 1 1 1 1 1 1 1 1 1 1 ...
##   .. ..@ FNR       : num [1:879] 0 0 0 0 0 0 0 0 0 0 ...
##   .. ..@ PPP       : num [1:879] 0.069 0.069 0.069 0.069 0.069 ...
##   .. ..@ NPP       : num [1:879] NaN NaN NaN NaN NaN NaN NaN NaN NaN NaN ...
##   .. ..@ MCR       : num [1:879] 0.931 0.931 0.931 0.931 0.931 ...
##   .. ..@ OR        : num [1:879] NaN NaN NaN NaN NaN NaN NaN NaN NaN NaN ...
##   .. ..@ kappa     : num [1:879] 0 0 0 0 0 0 0 0 0 0 ...
##  $ t    :'data.frame':   1 obs. of  6 variables:
##   ..$ kappa          : num 0.374
##   ..$ spec_sens      : num 0.374
##   ..$ no_omission    : num 0.374
##   ..$ prevalence     : num 0.0693
##   ..$ equal_sens_spec: num 0.443
##   ..$ sensitivity    : num 0.457
##  $ v_bkg: num [1:900, 1:20] 190 577 123 102 3155 ...
##   ..- attr(*, "dimnames")=List of 2
##   .. ..$ : NULL
##   .. ..$ : chr [1:20] "alt" "bio1" "bio2" "bio3" ...
##  $ v_pnt: num [1:72, 1:20] 1252 1946 1793 1871 1793 ...
##   ..- attr(*, "dimnames")=List of 2
##   .. ..$ : NULL
##   .. ..$ : chr [1:20] "alt" "bio1" "bio2" "bio3" ...
```

##### Rank model AUC comparing to 99 null-models (with training dataset)

```
valores_puntos_aleatorios_fondo_y_entrenamiento <- rbind(evalutaion_training$v_pnt, evalutaion_training$v_bkg)

rep <- 99     # number or null models to produce

null_models <- list()
for (r in 1:rep) {
    # following Robert J. Hijmans nullRandom() function from 'dismo'
        # sample presence records and set the rest of records as absence
        index <- sample(nrow(valores_puntos_aleatorios_fondo_y_entrenamiento), presence_points_count)
        pres <- valores_puntos_aleatorios_fondo_y_entrenamiento[index, ]
        absc <- valores_puntos_aleatorios_fondo_y_entrenamiento[-index, ]
        d <- data.frame(rbind(pres, absc))
        v <- c(rep(1, nrow(pres)), rep(0, nrow(absc)))
        # make null-model with presence and absence sets
        m <- maxent(d, v)
        # evaluate the null-model
        null_models[[r]] <- evaluate(pres, absc, m)
}

#null_models
auc_list_vector <- sapply(null_models, function(x) x@auc)

#auc_list_vector
mean(auc_list_vector)
```

```
## [1] 0.7201531
```

```
auc_list_training_vector <- c(evalutaion_training$e@auc, auc_list_vector)
auc_list_training_vector_ranks <- rank(auc_list_training_vector, ties.method= "max")

print('Rank of trainign AUC vs null-models')
```

```
## [1] "Rank of trainign AUC vs null-models"
```

```
auc_list_training_vector_ranks[1]
```

```
## [1] 100
```

```
auc_list_control_vector <- c(evalutaion_test$e@auc, auc_list_vector)
auc_list_control_vector_ranks <- rank(auc_list_control_vector, ties.method= "max")

print('Rank of control AUC vs null-models')
```

```
## [1] "Rank of control AUC vs null-models"
```

```
auc_list_control_vector_ranks[1]
```

```
## [1] 100
```

##### Store the model, parameters and control values in an object for further use

```
container <- list()
container$bkg <- puntos_aleatorios_fondo
container$entr <- puntos_entrenamiento_coordenadas
container$cntrl <- puntos_control_extra_coordenadas
container$model <- modelo_maxent
container$contrib <- contribuciones
container$e_train <- evalutaion_training
container$e_test <- evalutaion_test
container$e_test2 <- evalutaion_test2
container$auc_rank_train <- auc_list_training_vector_ranks[1]
container$auc_rank_test <- auc_list_control_vector_ranks[1]
container$auc_null_model_mean <- mean(auc_list_vector)
  
iterations[[i]] <- container
```
