## Supplementary material for "Vulnerability to climate change for narrowly ranged species: the case of Ecuadorian endemic *Magnolia mercedesiarum*": step_29dup_10.html

```
##                                                                                        [,1]
## X.Training.samples                                                                  50.0000
## Regularized.training.gain                                                            2.5011
## Unregularized.training.gain                                                          2.6659
## Iterations                                                                         500.0000
## Training.AUC                                                                         0.9727
## X.Background.points                                                                945.0000
## alt.contribution                                                                     8.8286
## bio1.contribution                                                                    0.0000
## bio10.contribution                                                                   0.0019
## bio11.contribution                                                                   0.0000
## bio12.contribution                                                                   0.0755
## bio13.contribution                                                                   0.0000
## bio14.contribution                                                                   1.9357
## bio15.contribution                                                                   0.2335
## bio16.contribution                                                                   0.1243
## bio17.contribution                                                                   3.3550
## bio18.contribution                                                                  12.1991
## bio19.contribution                                                                  27.7010
## bio2.contribution                                                                    0.0057
## bio3.contribution                                                                   17.9543
## bio4.contribution                                                                    0.0537
## bio5.contribution                                                                    0.0034
## bio6.contribution                                                                   24.8066
## bio7.contribution                                                                    0.0000
## bio8.contribution                                                                    1.2137
## bio9.contribution                                                                    1.5079
## alt.permutation.importance                                                          11.8917
## bio1.permutation.importance                                                          0.0000
## bio10.permutation.importance                                                         0.0000
## bio11.permutation.importance                                                         0.0000
## bio12.permutation.importance                                                         1.1198
## bio13.permutation.importance                                                         0.0000
## bio14.permutation.importance                                                         0.0000
## bio15.permutation.importance                                                         4.6477
## bio16.permutation.importance                                                         1.9715
## bio17.permutation.importance                                                         0.0000
## bio18.permutation.importance                                                        21.8076
## bio19.permutation.importance                                                         4.5656
## bio2.permutation.importance                                                          0.0086
## bio3.permutation.importance                                                          5.6508
## bio4.permutation.importance                                                          0.1989
## bio5.permutation.importance                                                          0.1513
## bio6.permutation.importance                                                         47.9863
## bio7.permutation.importance                                                          0.0000
## bio8.permutation.importance                                                          0.0000
## bio9.permutation.importance                                                          0.0000
## Training.gain.without.alt                                                            2.4564
## Training.gain.without.bio1                                                           2.5011
## Training.gain.without.bio10                                                          2.5011
## Training.gain.without.bio11                                                          2.5010
## Training.gain.without.bio12                                                          2.5009
## Training.gain.without.bio13                                                          2.5010
## Training.gain.without.bio14                                                          2.5013
## Training.gain.without.bio15                                                          2.4895
## Training.gain.without.bio16                                                          2.5005
## Training.gain.without.bio17                                                          2.5010
## Training.gain.without.bio18                                                          2.4260
## Training.gain.without.bio19                                                          2.4960
## Training.gain.without.bio2                                                           2.5009
## Training.gain.without.bio3                                                           2.4812
## Training.gain.without.bio4                                                           2.4997
## Training.gain.without.bio5                                                           2.5009
## Training.gain.without.bio6                                                           2.4994
## Training.gain.without.bio7                                                           2.5011
## Training.gain.without.bio8                                                           2.5012
## Training.gain.without.bio9                                                           2.5011
## Training.gain.with.only.alt                                                          1.5265
## Training.gain.with.only.bio1                                                         1.3815
## Training.gain.with.only.bio10                                                        1.4026
## Training.gain.with.only.bio11                                                        1.3513
## Training.gain.with.only.bio12                                                        0.1897
## Training.gain.with.only.bio13                                                        0.0977
## Training.gain.with.only.bio14                                                        0.8774
## Training.gain.with.only.bio15                                                        0.3685
## Training.gain.with.only.bio16                                                        0.0984
## Training.gain.with.only.bio17                                                        0.8046
## Training.gain.with.only.bio18                                                        0.5066
## Training.gain.with.only.bio19                                                        0.3242
## Training.gain.with.only.bio2                                                         0.0354
## Training.gain.with.only.bio3                                                         0.5497
## Training.gain.with.only.bio4                                                         0.7451
## Training.gain.with.only.bio5                                                         1.3946
## Training.gain.with.only.bio6                                                         1.3633
## Training.gain.with.only.bio7                                                         0.1241
## Training.gain.with.only.bio8                                                         1.4106
## Training.gain.with.only.bio9                                                         1.3664
## Entropy                                                                              4.3492
## Prevalence..average.probability.of.presence.over.background.sites.                   0.0482
## Fixed.cumulative.value.1.cumulative.threshold                                        1.0000
## Fixed.cumulative.value.1.Cloglog.threshold                                           0.0116
## Fixed.cumulative.value.1.area                                                        0.1693
## Fixed.cumulative.value.1.training.omission                                           0.0000
## Fixed.cumulative.value.5.cumulative.threshold                                        5.0000
## Fixed.cumulative.value.5.Cloglog.threshold                                           0.1257
## Fixed.cumulative.value.5.area                                                        0.0804
## Fixed.cumulative.value.5.training.omission                                           0.0000
## Fixed.cumulative.value.10.cumulative.threshold                                      10.0000
## Fixed.cumulative.value.10.Cloglog.threshold                                          0.2949
## Fixed.cumulative.value.10.area                                                       0.0624
## Fixed.cumulative.value.10.training.omission                                          0.0200
## Minimum.training.presence.cumulative.threshold                                       6.8481
## Minimum.training.presence.Cloglog.threshold                                          0.1862
## Minimum.training.presence.area                                                       0.0709
## Minimum.training.presence.training.omission                                          0.0000
## X10.percentile.training.presence.cumulative.threshold                               19.5798
## X10.percentile.training.presence.Cloglog.threshold                                   0.5081
## X10.percentile.training.presence.area                                                0.0476
## X10.percentile.training.presence.training.omission                                   0.1000
## Equal.training.sensitivity.and.specificity.cumulative.threshold                     13.7628
## Equal.training.sensitivity.and.specificity.Cloglog.threshold                         0.4341
## Equal.training.sensitivity.and.specificity.area                                      0.0561
## Equal.training.sensitivity.and.specificity.training.omission                         0.0600
## Maximum.training.sensitivity.plus.specificity.cumulative.threshold                   6.8481
## Maximum.training.sensitivity.plus.specificity.Cloglog.threshold                      0.1862
## Maximum.training.sensitivity.plus.specificity.area                                   0.0709
## Maximum.training.sensitivity.plus.specificity.training.omission                      0.0000
## Balance.training.omission..predicted.area.and.threshold.value.cumulative.threshold   2.3577
## Balance.training.omission..predicted.area.and.threshold.value.Cloglog.threshold      0.0316
## Balance.training.omission..predicted.area.and.threshold.value.area                   0.1101
## Balance.training.omission..predicted.area.and.threshold.value.training.omission      0.0000
## Equate.entropy.of.thresholded.and.original.distributions.cumulative.threshold        4.8813
## Equate.entropy.of.thresholded.and.original.distributions.Cloglog.threshold           0.1212
## Equate.entropy.of.thresholded.and.original.distributions.area                        0.0815
## Equate.entropy.of.thresholded.and.original.distributions.training.omission           0.0000
```

```
#str(modelo_maxent@results)
```

##### Get variable importance values

```
#plot(modelo_maxent)

contribuciones <- maxent.select.contribution(modelo_maxent,names_after_drop)

#contribuciones
par(mfrow=c(1, 2))
dotchart(as.numeric(contribuciones[,2]),
        col='blue', pch=16,
        labels=contribuciones[,1],
        main='predictor contribution')
dotchart(as.numeric(contribuciones[,3]),
        col='red', pch=16,
        labels=contribuciones[,1],
        main='predictor importance in permutations')
```

##### Draw respnse cuves

```
response(modelo_maxent)
```

##### Output MaxEnt HTML

```
#modelo_maxent
```

### Initial model evaluation

##### Evaluate model with training dataset

```
evalutaion_training <- evaluate.mdl(puntos_entrenamiento_coordenadas,puntos_aleatorios_fondo,capas_raster_drop,modelo_maxent)
```

```
## class          : ModelEvaluation 
## n presences    : 50 
## n absences     : 945 
## AUC            : 0.9726561 
## cor            : 0.6520958 
## max TPR+TNR at : 0.1861077
```

```
##                kappa spec_sens no_omission prevalence equal_sens_spec
## thresholds 0.5014433 0.1861077   0.1861077 0.04957077       0.4340354
##            sensitivity
## thresholds   0.5080043
```

```
str(evalutaion_training)
```

```
## List of 4
##  $ e    :Formal class 'ModelEvaluation' [package "dismo"] with 22 slots
##   .. ..@ presence  : num [1:50] 0.343 0.698 0.652 0.62 0.642 ...
##   .. ..@ absence   : num [1:945] 4.69e-05 1.11e-04 3.63e-07 9.06e-02 1.21e-05 ...
##   .. ..@ np        : int 50
##   .. ..@ na        : int 945
##   .. ..@ auc       : num 0.973
##   .. ..@ pauc      : num(0) 
##   .. ..@ cor       : Named num 0.652
##   .. .. ..- attr(*, "names")= chr "cor"
##   .. ..@ pcor      : num 1.51e-121
##   .. ..@ t         : num [1:874] -1e-04 -1e-04 -1e-04 -1e-04 -1e-04 ...
##   .. ..@ confusion : int [1:874, 1:4] 50 50 50 50 50 50 50 50 50 50 ...
##   .. .. ..- attr(*, "dimnames")=List of 2
##   .. .. .. ..$ : NULL
##   .. .. .. ..$ : chr [1:4] "tp" "fp" "fn" "tn"
##   .. ..@ prevalence: num [1:874] 0.0503 0.0503 0.0503 0.0503 0.0503 ...
##   .. ..@ ODP       : num [1:874] 0.95 0.95 0.95 0.95 0.95 ...
##   .. ..@ CCR       : num [1:874] 0.0503 0.0503 0.0503 0.0503 0.0503 ...
##   .. ..@ TPR       : num [1:874] 1 1 1 1 1 1 1 1 1 1 ...
##   .. ..@ TNR       : num [1:874] 0 0 0 0 0 0 0 0 0 0 ...
##   .. ..@ FPR       : num [1:874] 1 1 1 1 1 1 1 1 1 1 ...
##   .. ..@ FNR       : num [1:874] 0 0 0 0 0 0 0 0 0 0 ...
##   .. ..@ PPP       : num [1:874] 0.0503 0.0503 0.0503 0.0503 0.0503 ...
##   .. ..@ NPP       : num [1:874] NaN NaN NaN NaN NaN NaN NaN NaN NaN NaN ...
##   .. ..@ MCR       : num [1:874] 0.95 0.95 0.95 0.95 0.95 ...
##   .. ..@ OR        : num [1:874] NaN NaN NaN NaN NaN NaN NaN NaN NaN NaN ...
##   .. ..@ kappa     : num [1:874] 0 0 0 0 0 0 0 0 0 0 ...
##  $ t    :'data.frame':   1 obs. of  6 variables:
##   ..$ kappa          : num 0.501
##   ..$ spec_sens      : num 0.186
##   ..$ no_omission    : num 0.186
##   ..$ prevalence     : num 0.0496
##   ..$ equal_sens_spec: num 0.434
##   ..$ sensitivity    : num 0.508
##  $ v_bkg: num [1:900, 1:20] 250 255 17 2801 161 ...
##   ..- attr(*, "dimnames")=List of 2
##   .. ..$ : NULL
##   .. ..$ : chr [1:20] "alt" "bio1" "bio2" "bio3" ...
##  $ v_pnt: num [1:50, 1:20] 2148 1877 1720 1902 1470 ...
##   ..- attr(*, "dimnames")=List of 2
##   .. ..$ : NULL
##   .. ..$ : chr [1:20] "alt" "bio1" "bio2" "bio3" ...
```

##### Evaluate model with control dataset

```
evalutaion_test <- evaluate.mdl(puntos_control_coordenadas,puntos_aleatorios_fondo,capas_raster_drop,modelo_maxent)
```

```
## class          : ModelEvaluation 
## n presences    : 50 
## n absences     : 945 
## AUC            : 0.9714497 
## cor            : 0.6338032 
## max TPR+TNR at : 0.2604912
```

```
##                kappa spec_sens no_omission prevalence equal_sens_spec
## thresholds 0.3007302 0.2604912   0.2604912 0.04957077       0.3179175
##            sensitivity
## thresholds   0.3250952
```

```
str(evalutaion_test)
```

```
## List of 4
##  $ e    :Formal class 'ModelEvaluation' [package "dismo"] with 22 slots
##   .. ..@ presence  : num [1:50] 0.616 0.752 0.758 0.275 0.533 ...
##   .. ..@ absence   : num [1:945] 4.69e-05 1.11e-04 3.63e-07 9.06e-02 1.21e-05 ...
##   .. ..@ np        : int 50
##   .. ..@ na        : int 945
##   .. ..@ auc       : num 0.971
##   .. ..@ pauc      : num(0) 
##   .. ..@ cor       : Named num 0.634
##   .. .. ..- attr(*, "names")= chr "cor"
##   .. ..@ pcor      : num 6.9e-113
##   .. ..@ t         : num [1:874] -1e-04 -1e-04 -1e-04 -1e-04 -1e-04 ...
##   .. ..@ confusion : int [1:874, 1:4] 50 50 50 50 50 50 50 50 50 50 ...
##   .. .. ..- attr(*, "dimnames")=List of 2
##   .. .. .. ..$ : NULL
##   .. .. .. ..$ : chr [1:4] "tp" "fp" "fn" "tn"
##   .. ..@ prevalence: num [1:874] 0.0503 0.0503 0.0503 0.0503 0.0503 ...
##   .. ..@ ODP       : num [1:874] 0.95 0.95 0.95 0.95 0.95 ...
##   .. ..@ CCR       : num [1:874] 0.0503 0.0503 0.0503 0.0503 0.0503 ...
##   .. ..@ TPR       : num [1:874] 1 1 1 1 1 1 1 1 1 1 ...
##   .. ..@ TNR       : num [1:874] 0 0 0 0 0 0 0 0 0 0 ...
##   .. ..@ FPR       : num [1:874] 1 1 1 1 1 1 1 1 1 1 ...
##   .. ..@ FNR       : num [1:874] 0 0 0 0 0 0 0 0 0 0 ...
##   .. ..@ PPP       : num [1:874] 0.0503 0.0503 0.0503 0.0503 0.0503 ...
##   .. ..@ NPP       : num [1:874] NaN NaN NaN NaN NaN NaN NaN NaN NaN NaN ...
##   .. ..@ MCR       : num [1:874] 0.95 0.95 0.95 0.95 0.95 ...
##   .. ..@ OR        : num [1:874] NaN NaN NaN NaN NaN NaN NaN NaN NaN NaN ...
##   .. ..@ kappa     : num [1:874] 0 0 0 0 0 0 0 0 0 0 ...
##  $ t    :'data.frame':   1 obs. of  6 variables:
##   ..$ kappa          : num 0.301
##   ..$ spec_sens      : num 0.26
##   ..$ no_omission    : num 0.26
##   ..$ prevalence     : num 0.0496
##   ..$ equal_sens_spec: num 0.318
##   ..$ sensitivity    : num 0.325
##  $ v_bkg: num [1:900, 1:20] 250 255 17 2801 161 ...
##   ..- attr(*, "dimnames")=List of 2
##   .. ..$ : NULL
##   .. ..$ : chr [1:20] "alt" "bio1" "bio2" "bio3" ...
##  $ v_pnt: num [1:50, 1:20] 1683 1684 1540 1383 1970 ...
##   ..- attr(*, "dimnames")=List of 2
##   .. ..$ : NULL
##   .. ..$ : chr [1:20] "alt" "bio1" "bio2" "bio3" ...
```

##### Evaluate model with fixed control dataset

```
evalutaion_test2 <- evaluate.mdl(puntos_control_extra_coordenadas,puntos_aleatorios_fondo,capas_raster_drop,modelo_maxent)
```

```
## class          : ModelEvaluation 
## n presences    : 72 
## n absences     : 967 
## AUC            : 0.9601143 
## cor            : 0.6277981 
## max TPR+TNR at : 0.2485233
```

```
##                kappa spec_sens no_omission prevalence equal_sens_spec
## thresholds 0.2485233 0.2485233   0.2485233 0.07005003       0.2941936
##            sensitivity
## thresholds   0.3224455
```

```
str(evalutaion_test2)
```

```
## List of 4
##  $ e    :Formal class 'ModelEvaluation' [package "dismo"] with 22 slots
##   .. ..@ presence  : num [1:72] 0.339 0.533 0.744 0.767 0.744 ...
##   .. ..@ absence   : num [1:967] 4.69e-05 1.11e-04 3.63e-07 9.06e-02 1.21e-05 ...
##   .. ..@ np        : int 72
##   .. ..@ na        : int 967
##   .. ..@ auc       : num 0.96
##   .. ..@ pauc      : num(0) 
##   .. ..@ cor       : Named num 0.628
##   .. .. ..- attr(*, "names")= chr "cor"
##   .. ..@ pcor      : num 5.74e-115
##   .. ..@ t         : num [1:882] -1e-04 -1e-04 -1e-04 -1e-04 -1e-04 ...
##   .. ..@ confusion : int [1:882, 1:4] 72 72 72 72 72 72 72 72 72 72 ...
##   .. .. ..- attr(*, "dimnames")=List of 2
##   .. .. .. ..$ : NULL
##   .. .. .. ..$ : chr [1:4] "tp" "fp" "fn" "tn"
##   .. ..@ prevalence: num [1:882] 0.0693 0.0693 0.0693 0.0693 0.0693 ...
##   .. ..@ ODP       : num [1:882] 0.931 0.931 0.931 0.931 0.931 ...
##   .. ..@ CCR       : num [1:882] 0.0693 0.0693 0.0693 0.0693 0.0693 ...
##   .. ..@ TPR       : num [1:882] 1 1 1 1 1 1 1 1 1 1 ...
##   .. ..@ TNR       : num [1:882] 0 0 0 0 0 0 0 0 0 0 ...
##   .. ..@ FPR       : num [1:882] 1 1 1 1 1 1 1 1 1 1 ...
##   .. ..@ FNR       : num [1:882] 0 0 0 0 0 0 0 0 0 0 ...
##   .. ..@ PPP       : num [1:882] 0.0693 0.0693 0.0693 0.0693 0.0693 ...
##   .. ..@ NPP       : num [1:882] NaN NaN NaN NaN NaN NaN NaN NaN NaN NaN ...
##   .. ..@ MCR       : num [1:882] 0.931 0.931 0.931 0.931 0.931 ...
##   .. ..@ OR        : num [1:882] NaN NaN NaN NaN NaN NaN NaN NaN NaN NaN ...
##   .. ..@ kappa     : num [1:882] 0 0 0 0 0 0 0 0 0 0 ...
##  $ t    :'data.frame':   1 obs. of  6 variables:
##   ..$ kappa          : num 0.249
##   ..$ spec_sens      : num 0.249
##   ..$ no_omission    : num 0.249
##   ..$ prevalence     : num 0.0701
##   ..$ equal_sens_spec: num 0.294
##   ..$ sensitivity    : num 0.322
##  $ v_bkg: num [1:900, 1:20] 250 255 17 2801 161 ...
##   ..- attr(*, "dimnames")=List of 2
##   .. ..$ : NULL
##   .. ..$ : chr [1:20] "alt" "bio1" "bio2" "bio3" ...
##  $ v_pnt: num [1:72, 1:20] 1252 1946 1793 1871 1793 ...
##   ..- attr(*, "dimnames")=List of 2
##   .. ..$ : NULL
##   .. ..$ : chr [1:20] "alt" "bio1" "bio2" "bio3" ...
```

##### Rank model AUC comparing to 99 null-models (with training dataset)

```
valores_puntos_aleatorios_fondo_y_entrenamiento <- rbind(evalutaion_training$v_pnt, evalutaion_training$v_bkg)

rep <- 99     # number or null models to produce

null_models <- list()
for (r in 1:rep) {
    # following Robert J. Hijmans nullRandom() function from 'dismo'
        # sample presence records and set the rest of records as absence
        index <- sample(nrow(valores_puntos_aleatorios_fondo_y_entrenamiento), presence_points_count)
        pres <- valores_puntos_aleatorios_fondo_y_entrenamiento[index, ]
        absc <- valores_puntos_aleatorios_fondo_y_entrenamiento[-index, ]
        d <- data.frame(rbind(pres, absc))
        v <- c(rep(1, nrow(pres)), rep(0, nrow(absc)))
        # make null-model with presence and absence sets
        m <- maxent(d, v)
        # evaluate the null-model
        null_models[[r]] <- evaluate(pres, absc, m)
}

#null_models
auc_list_vector <- sapply(null_models, function(x) x@auc)

#auc_list_vector
mean(auc_list_vector)
```

```
## [1] 0.7250983
```

```
auc_list_training_vector <- c(evalutaion_training$e@auc, auc_list_vector)
auc_list_training_vector_ranks <- rank(auc_list_training_vector, ties.method= "max")

print('Rank of trainign AUC vs null-models')
```

```
## [1] "Rank of trainign AUC vs null-models"
```

```
auc_list_training_vector_ranks[1]
```

```
## [1] 100
```

```
auc_list_control_vector <- c(evalutaion_test$e@auc, auc_list_vector)
auc_list_control_vector_ranks <- rank(auc_list_control_vector, ties.method= "max")

print('Rank of control AUC vs null-models')
```

```
## [1] "Rank of control AUC vs null-models"
```

```
auc_list_control_vector_ranks[1]
```

```
## [1] 100
```

##### Store the model, parameters and control values in an object for further use

```
container <- list()
container$bkg <- puntos_aleatorios_fondo
container$entr <- puntos_entrenamiento_coordenadas
container$cntrl <- puntos_control_extra_coordenadas
container$model <- modelo_maxent
container$contrib <- contribuciones
container$e_train <- evalutaion_training
container$e_test <- evalutaion_test
container$e_test2 <- evalutaion_test2
container$auc_rank_train <- auc_list_training_vector_ranks[1]
container$auc_rank_test <- auc_list_control_vector_ranks[1]
container$auc_null_model_mean <- mean(auc_list_vector)
  
iterations[[i]] <- container
```
